# Resource allocation to cell envelopes and the scaling of bacterial growth rate

**DOI:** 10.1101/2022.01.07.475415

**Authors:** Bogi Trickovic, Michael Lynch

## Abstract

Although various empirical studies have reported a positive correlation between the specific growth rate and cell size across bacteria, it is currently unclear what causes this relationship. We conjecture that such scaling occurs because smaller cells have a larger surface-to-volume ratio and thus have to allocate a greater fraction of the total resources to the production of the cell envelope, leaving fewer resources for other biosynthetic processes. To test this theory, we developed a coarse-grained model of bacterial physiology composed of the proteome that converts nutrients into biomass, with the cell envelope acting as a resource sink. Assuming resources are partitioned to maximize the growth rate, the model yields expected scalings. Namely, the growth rate and ribosomal mass fraction scale negatively, while the mass fraction of envelope-producing enzymes scales positively with surface-to-volume. These relationships are compatible with growth measurements and quantitative proteomics data reported in the literature.

## 2 Introduction

The rate of cell growth varies across bacterial species. The inhabitant of salt marshes, *Vibrio natriegens*, divides in 10 minutes [1], whereas the causal agent of gum disease, *Treponema denticola*, takes 20 hours to divide [2]. Given the centrality of the growth rate in physiology, ecology, and evolution, efforts have been invested in understanding the causes of this variation. Previous meta-analyses have reported that the overall pattern of variation is predictable: bacterial species with larger cell volume tend to grow faster. This observation was made in both heterotrophic [3, 4] and autotrophic bacteria [5, 6]. The causes of this scaling are unclear. Eukaryotic growth rates scale negatively with body size, implying that the theories explaining this pattern – such as those invoking transport-related constraints [7, 8] and limits imposed by self-shading of chlorophyll [9] – are fundamentally inadequate in accounting for the opposite scaling observed in bacteria.

Given that the surface-to-volume ratio increases with decreasing cell size, small cells will have more weight sequestered in membranes and walls than their larger counterparts. Therefore, as the cell size decreases, more resources have to be invested in the production of the cell envelope, implying that fewer resources can be invested in the biosynthetic processes that replicate the other features of the cell. It was initially suggested that this constraint might impose the limit on the smallest size that a cell can attain [10]. This idea has been further used to explain the lower limit on the size of the photoautotrophic organism and why phytoplankton growth rates appear to increase with cell volume [11, 12]. Our goal is to formalize this theory and investigate whether it can explain the growth scaling in heterotrophic bacteria. We start by imagining the fastest-growing cell as a bag of self-replicating ribosomes [13]. However, the growth rate is depressed below this perfect state because the cell has to invest resources in (1) machinery that acquires and converts nutrients to fuel ribosomes, and (2) cell envelope, which is necessary for the cell to maintain the proper shape. As the cell becomes smaller, a larger fraction of the resources are diverted to envelope, and fewer resources are left for ribosomes which, in turn, means that cells grow more slowly.

We formalize the afore-mentioned verbal argument in sections 3.1 and 3.2, and then use this framework to obtain the simple analytical solution for maximal attainable growth rate given a particular cell size (Section 3.3). These predictions are tested using quantitative proteomic data of the model bacterium, *Escherichia coli* (Section 4.1). Lastly, these cross-species scaling expectations are compared against data on bacterial growth rates, cell sizes, and proteome compositions (sections 4.3 and 4.2).

## 3 Materials and methods

### 3.1 Derivation of the steady-state growth rate

In our model, a cell is composed of two metabolite species and three protein species (Fig 1). The external nutrient concentration is assumed to be constant, thus mimicking the nutrient-replete conditions when bacteria are grown in the lab. These nutrients are taken up and converted into building block *b* which corresponds to amino acids. Although we refer to species *l* as lipids, this group includes all molecules used in the construction of the cell envelope, such as peptides and saccharides (including components of membrane lipoproteins and peptidoglycan). We use upper-case symbols to refer to absolute abundances, and lower-case symbols for relative abundances or concentrations. All chemical reactions obey Michaelis-Menten kinetics, and we assume that half-saturation constants (*K*_*M*_) for all reactions are identical. We ultimately focus on the special case when all reactions are operating at saturation (*K*_*M*_ = 0) because this permits a simple analytical solution for the steady-state of various cellular features, and we only vary *K*_*M*_ to compare this limiting behavior to a more general case when cellular enzymes are not saturated. Table 1 outlines the meaning of symbols that are used throughout the main text.

**Table 1.** List of symbols used in the text and their meaning.

| Symbol | Meaning |
| --- | --- |
| $p_b(t)$ | The relative abundance of building block-producing enzyme at time $t$ |
| $p_l(t)$ | The relative abundance of envelope-producing enzyme at time $t$ |
| $r(t)$ | The relative abundance of ribosomes at time $t$ |
| $l(t)$ | The relative abundance of envelope component |
| $b(t)$ | The relative abundance of building blocks |
| $P_b(t)$ | The absolute abundance of building block producer |
| $P_l(t)$ | The absolute abundance of envelope producer |
| $R(t)$ | The absolute abundance of ribosomes |
| $L(t)$ | The absolute abundance of envelope component |
| $B(t)$ | The absolute abundance of building blocks |
| $\phi_b$ | The ribosome fraction allocated to translation of building block producers |
| $\phi_l$ | The ribosome fraction allocated to translation of envelope producers |
| $\phi_r$ | The ribosome fraction allocated to translation of ribosomes |
| $\Phi_B$ | The proteome mass fraction allocated to translation of building block producers |
| $\Phi_L$ | The proteome mass fraction allocated to translation of envelope producers |
| $\Phi_R$ | The proteome mass fraction allocated to translation of ribosomes |
| $m_p$ | Length of building block producers and envelope producers in amino acids |
| $m_r$ | Length of ribosome in amino acids |
| $m_s$ | Length of the unit of envelope (lipids, peptides, and sugars) in amino acids |
| $k_n$ | The turnover number of building block producers |
| $k_l$ | The turnover number of envelope producers |
| $k_t$ | The turnover number of ribosomes |
| $d_p$ | Protein degradation rate |
| $d_l$ | Envelope degradation rate |
| $K_M$ | Michaelis-Menten constant for all chemical reactions |
| $\Pi$ | Surface area-to-cytoplasmic volume ratio |
| $\beta$ | The number of envelope units per unit of surface area |
| $\gamma$ | The number of amino acids in proteome per unit of cytoplasmic volume |
| $\epsilon$ | The resource cost of the unit S/V ($m_s\beta/\gamma$) |
| $V(t)$ | Volume of the cytoplasm |
| $\lambda(t)$ | The specific growth rate at time $t$ |

**Figure 1.**
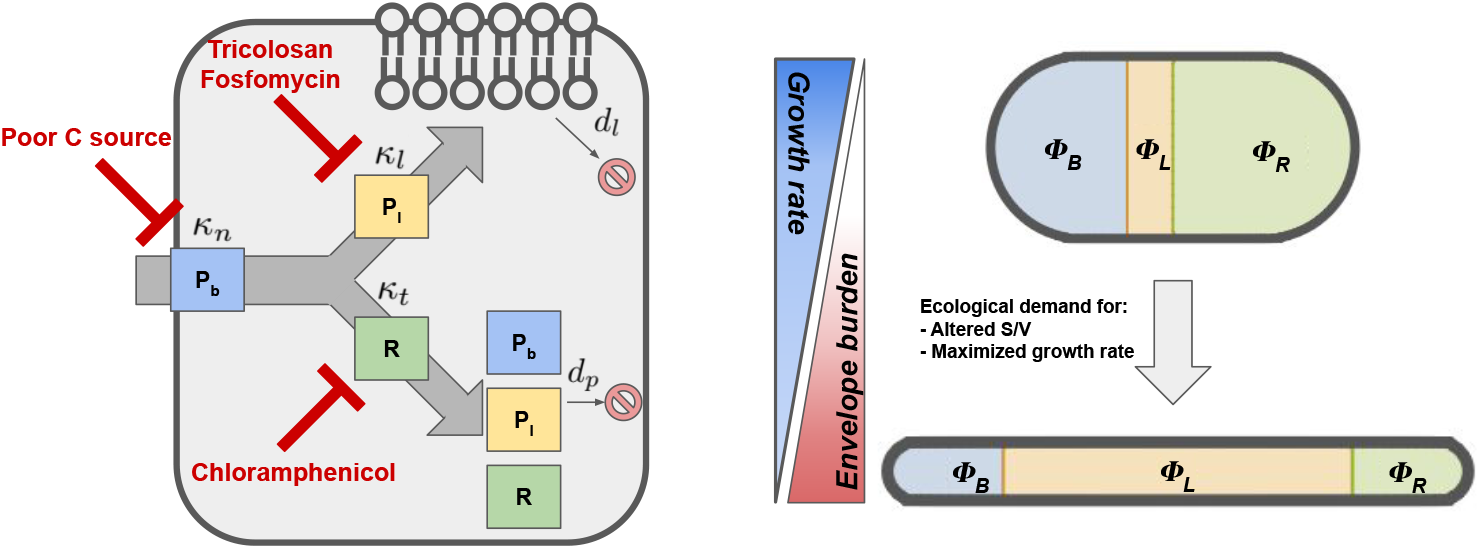
Model of the cell. The key components and processes in the model (left pane). Nutrients are taken up from the external environment via the building block producer (*P*_*b*_). The produced block is then converted into lipids and other cell envelope components via the lipid-producer (*P*_*l*_), and into proteins by the ribosome (*R*). Envelope and protein synthesis production can be inhibited by antibiotics, whereas nutrient uptake can be reduced by growing the culture on a poor carbon source. Envelope and proteins are degraded at rates *d*_*l*_ and *d*_*p*_, respectively. Building blocks are represented as grey arrows flowing through protein machinery. Envelope components are depicted in the membrane. Outline of the hypothesis (right pane).

Our model is similar to those proposed in [14], with two important differences. First, the cell divides once the critical volume is reached, and not the critical abundance of a particular protein controlling cell division. We chose this model because it was the easiest to deal with analytically, and other cell division mechanisms are unlikely to affect the overall scaling patterns, as the envelope imposes a burden regardless of the exact molecular details underlying the cell division. Second, we introduced the degradation of macromolecules, and the production of an additional metabolite, which is the envelope component. Perhaps most importantly, we obtained a simple analytical solution to the maximal growth rate in the limit of saturation, whereas the model in [14] is explored only numerically. This allows one to quickly derive relationships between proteomic mass fractions and the growth rate under various perturbations, and use these to infer model parameters. The model is also similar to the one reported in [15], as these authors also include protein degradation, but differs in that it posits second-order mass action kinetics, whereas we let reactions run according to Michaelis-Menten kinetics. Lastly, we show that their analytical solution for maximal growth rate reduces to our solution, in the limit of no envelope burden and enzyme saturation.

The time-evolution of metabolite concentrations are:

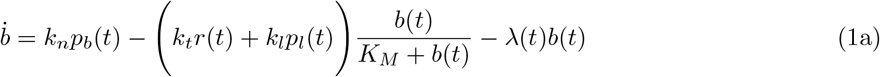

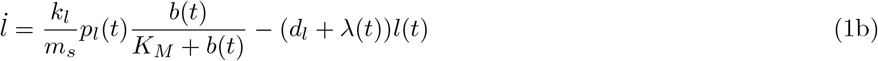

The production of *b* occurs at rate *k*_*n*_, which represents a pseudo-first-order rate constant that depends on the nutrient status of the environment inhabited by the cell. Rate constants *k*_*n*_, *k*_*l*_, and *k*_*t*_ are the turnover numbers for reactions of nutrient processing, envelope synthesis, and translation, while *m*_*s*_ is the size an envelope unit expressed in terms of the number of building blocks. Envelope components are eliminated from the cell by degradation at rate *d*_*l*_, and by dilution due to growth at rate *λ*(*t*). Note that the growth rate is time-dependent because it is a function of state variables (i.e., molecular abundances). We neglect the degradation of the free amino acids and focus only on the turnover of cell envelope components.

Protein species *p*_*b*_, *p*_*l*_, and *r* are produced by ribosomes which represent the autocatalytic part of the cell. All proteins have associated degradation rates *d*_*p*_, with the exception of ribosomes which are reported to be remarkably stable in exponentially growing cells [16]. Sizes of the metabolic protein (*P*_*b*_ and *P*_*l*_) and the ribosome are *m*_*p*_ and *m*_*r*_, respectively. The time-evolution equations are identical in form to equations for metabolites:

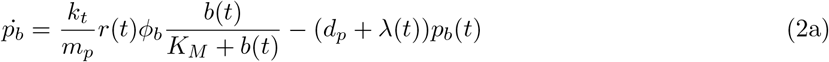

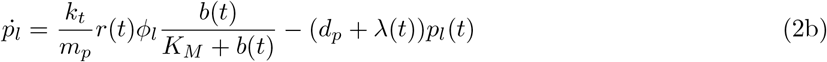

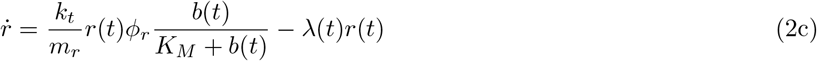

with the only notable difference that – given the finite ribosomal pool – ribosomes have to be partitioned between different protein species, and this is denoted with *ϕ*_*x*_, which represents the fraction of total ribosomal concentration that is allocated to translation of protein species *x*. We will assume that each molecular species contributes to volume proportional to its size in amino acids. For example, ribosomes are about 20x larger than other metabolic proteins, contributing 20x more to the cell volume. Concentrations of each species are defined as the abundance (labeled with the capital letter) divided by cytoplasmic volume *V* :

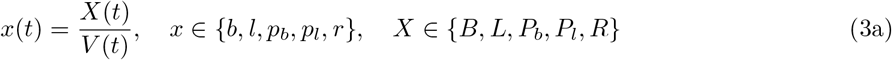

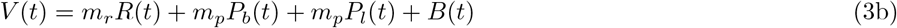

The volume *V* corresponds to the internal volume of the cell where the chemical reactions take place, and including the envelope in the volume of the cell would imply that one can slow down chemical reactions by, say, increasing the thickness of the cell wall. As this is a non-sensical conclusion, we assume that *L* does not contribute to cytoplasmic volume. The cell is assumed to grow exponentially, such that 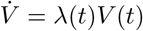. Given that both volume and concentrations are time-dependent, the left hand side of equations (1a–2c) will take the form:

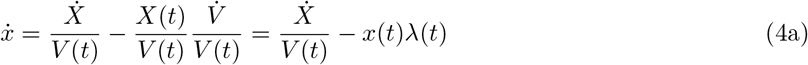

Substituting Eq (4a) in Eq (1a–2c) and rearranging yields:

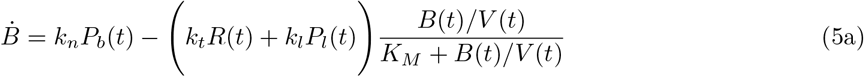

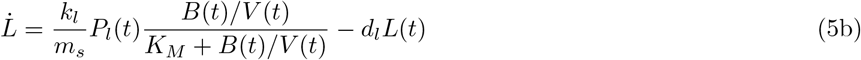

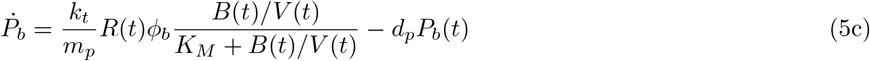

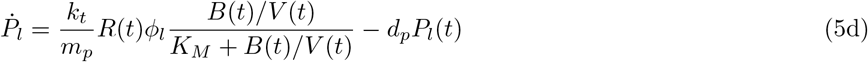

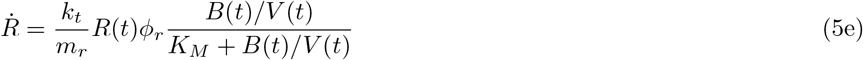

When cells are in a steady-state, all cellular components grow exponentially at a constant rate 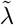 which is the steady-state growth rate. We use this property to find the steady-state of the dynamical system. More precisely, we have:

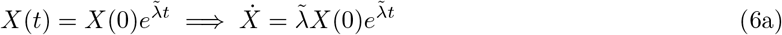

Note that 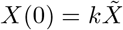, where 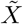 is abundance at cell division time when the cell is in the steady-state, and *k* is 1/2 if the abundance doubles over the life cycle. Substituting Eq 6a in the system (5a–5e) leads to:

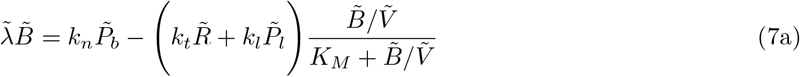

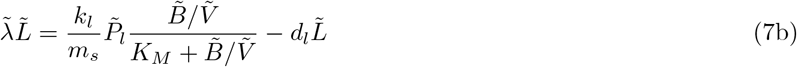

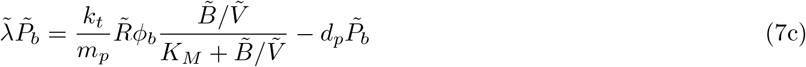

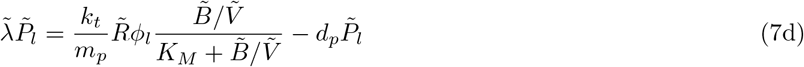

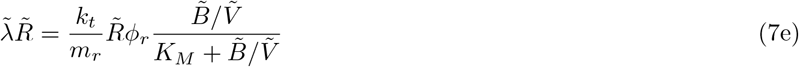

An intuitive interpretation of these equations is that the rate at which molecules are synthesized and degraded ultimately equals the rate at which they are diluted by growth. Note that 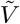 is the steady-state cell volume which is by definition the critical volume at which cell divides. To simplify downstream expressions, let *κ*_*n*_ = *k*_*n*_*/m*_*p*_, *κ*_*t*_ = *k*_*t*_*/m*_*r*_, *κ*_*l*_ = *k*_*l*_*/m*_*p*_. These are rate constants scaled by the size of a particular enzyme, capturing how fast an enzyme is relative to its size. Solving for molecular abundances yields:

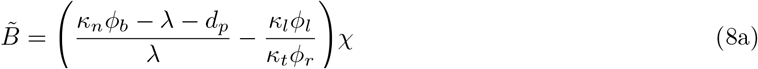

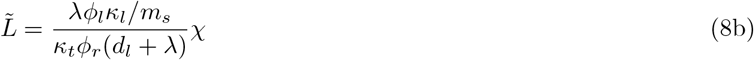

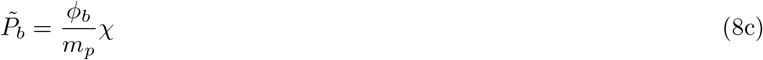

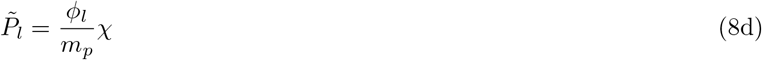

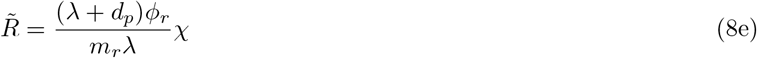

where

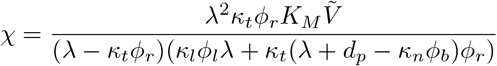

By definition, proteome mass fractions are:

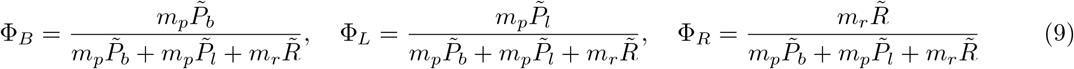

By replacing steady-state abundances in Eq 10 with solutions in Eq 8a–8e, we have:

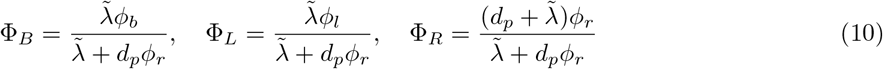

In the limit when there is no protein degradation (*d*_*p*_ = 0), the proteomic mass fractions are equal to the fraction of ribosomes allocated to the synthesis of a particular protein component. This result was originally obtained in [17]. However, we are still lacking the solution for the steady-state growth rate 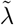.

Given that volume increases exponentially and from the definition of the cell volume (Eq 3b), we have:

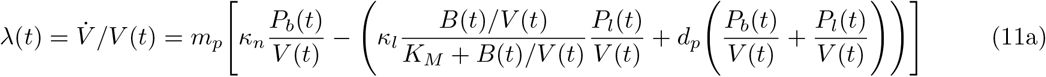

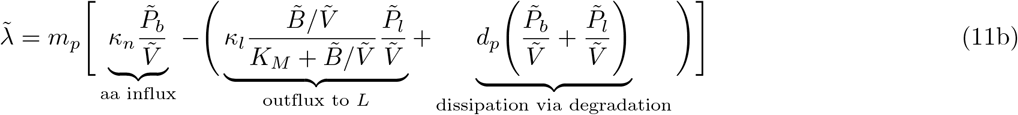

where Eq 11b has been obtained by applying 6a to molecular abundances in 11a. The steady-state growth rate is equal to the rate at which the building blocks are generated from the acquired nutrients minus the building blocks that are diverted into cell envelope synthesis and thus do not contribute to cytoplasmic volume production or are dissipated through protein degradation occurring at a rate *d*_*p*_. Note that diversion to lipids and other envelope constituents does not explicitly enter the growth rate because *L* does not contribute to the cytoplasmic volume. However, lipid production does affect the growth rate by altering the amount of available building blocks 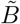, and by diverting proteome allocation to *P*_*l*_. Finally, substituting equations 8a–8e into 11b, we retrieve a cubic polynomial:

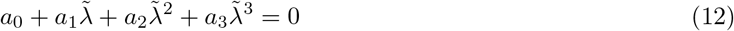

with the following coefficients:

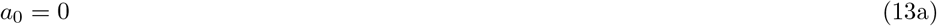

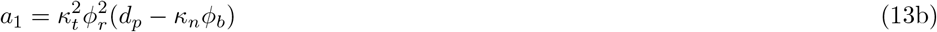

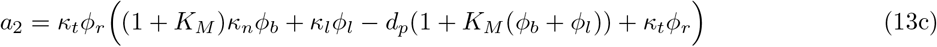

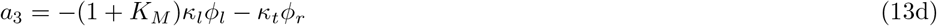

Eq 12 has two non-zero roots:

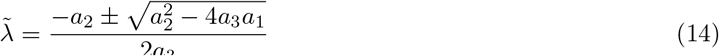

In the case of the positive branch, 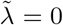 when *ϕ*_*r*_ = 0 and 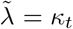 when *ϕ*_*r*_ = 1. This is clearly incorrect, given that 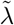 should be zero both when the cell does not have ribosomes (because there is nothing to build the cell) and when the cell contains only ribosomes without any other protein components (because there is nothing to deliver building blocks to the ribosomes). On the contrary, the negative branch attains values of zero both when *ϕ*_*r*_ = 0 and when *ϕ*_*r*_ = 1, so we take this solution as biologically meaningful. Finally, we obtain the steady-state abundances by substituting Eq 14 in equations 8a–8e.

The analytical solution for the steady-state growth rate 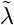 (Eq 14) is validated by comparison to numerically integrating equations 5a–5e (Fig 2; upper row). We assign initial abundances and integrate the system of ODEs until the cell volume reaches the critical division volume. Next, we reduce the initial abundances by the factor of two and restart the integration process. This emulates the process of cell division when cellular content is equipartitioned among the daughter cells. After a short out-of-equilibrium phase, the cell lineage settles into a steady-state that matches the analytical solution for 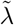 and the steady-state abundances.

**Figure 2.**
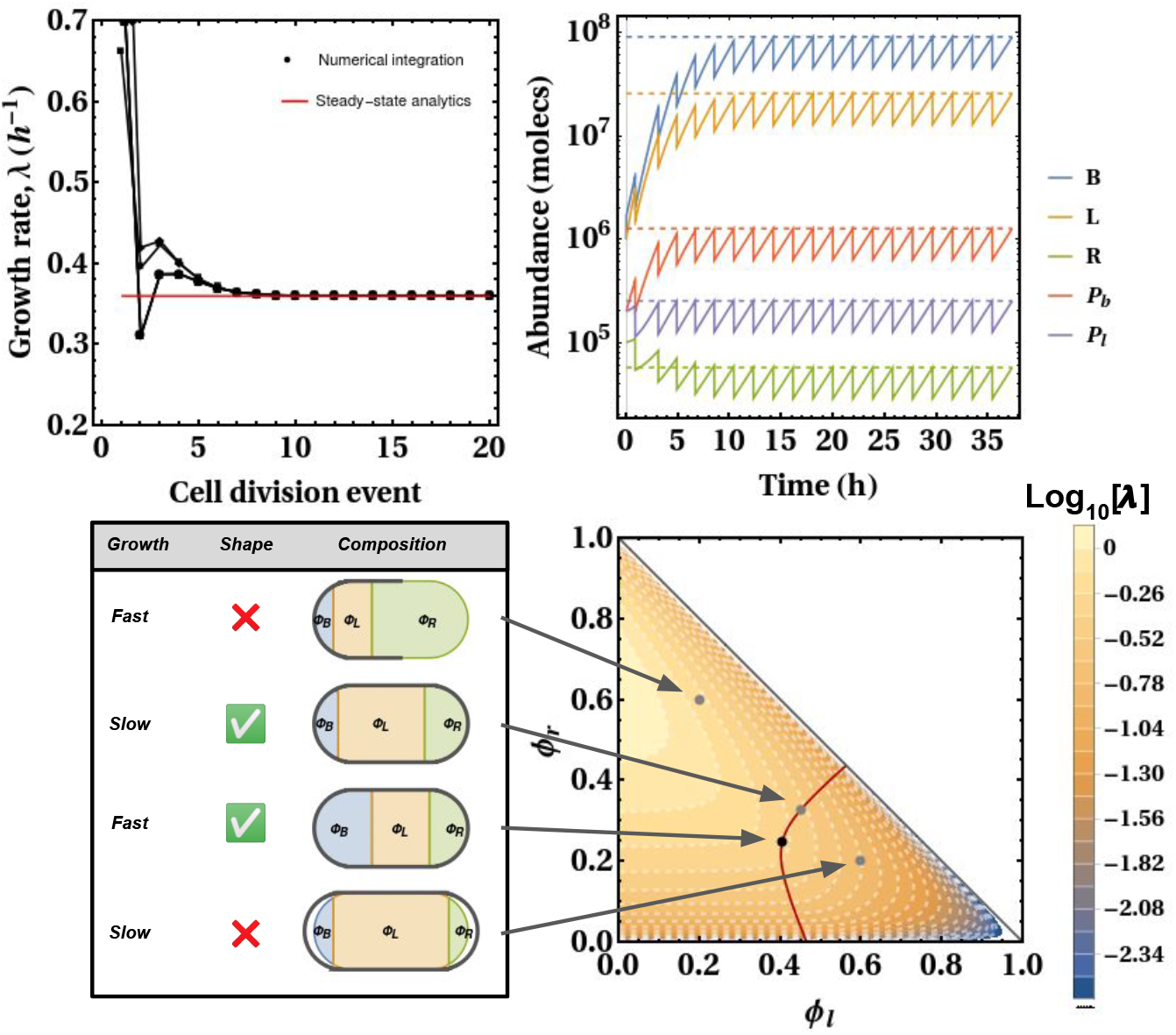
Comparison of the analytical and numerical solution for the cell in the steady-state. The upper left panel shows a close match of the steady-state growth rate (solid red line) relative to numerically integrated system of ODEs (black dots). Each set of points corresponds to a different set of initial molecular abundances. On the right, a similar match between analytics and numerics also exists for abundances of five molecular species; each grey dashed line represents the steady-state analytical solution given by Eq (8a–8e). The bottom row outlines the kind of problem that a cell has to solve. Parameters (bottom figures): *κ*_*l*_ = 4.32 h^−1^, *κ*_*t*_ = 2.59 h^−1^, *d*_*p*_ = *d*_*l*_ = 0 h^−1^, *K*_*M*_ = 0.1, *γ* = 9.75 ×10^8^ aa*/μm*^3^, *β* = 2 ×10^6^ lip*/μm*^2^, *m*_*s*_ = 314 aa, *V* = 3 ×10^9^ aa. Parameters (upper figures): *κ*_*l*_ = 50 h^−1^, *κ*_*t*_ = 1 h^−1^, *d*_*p*_ = *d*_*l*_ = 0.1 h^−1^, *K*_*M*_ = 0.01, *V* = 10^9^ aa, *ϕ*_*b*_ = 0.5, *ϕ*_*l*_ = 0.1, *ϕ*_*r*_ = 0.4.

Although the explicit solution for the maximal 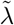 is prohibitively difficult to obtain, we can gain some insight by examining boundary cases when the cell does not have a membrane (*ϕ*_*l*_ = 0) and cellular processes are infinitely fast. When 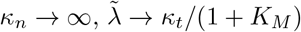, and, conversely, as *κ*_*t*_ → ∞ we have 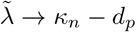. These two asymptotic results occur because the cell that instantaneously converts nutrients to building blocks will saturate the downstream translation machinery and the growth rate will be determined by the rate at which ribosomes operate. Conversely, when ribosomes are infinitely fast, the growth rate is set by how fast the building blocks are supplied to protein synthesis machinery.

### 3.2 Optimization problem of the cell

It is a working hypothesis that the steady-state growth rate 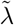 is a function of the ribosome partitioning parameters *ϕ*, and the cell has a task to find an optimal partitioning across different proteome components such that 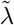 is maximized. This maximization is assumed to be achieved by homeostatic mechanisms that constantly take input from the environment and adjust proteome composition accordingly. Note, however, that the cell could attain maximal growth even if the partitioning parameters were hard-coded in the genome and thus cannot be actively adjusted (i.e., when homeostatic mechanisms are absent). In that case, the process of finding the peak is governed by natural selection and other evolutionary forces. Hence, our model holds regardless of whether the actual cells have regulatory mechanisms or not.

Growth rate maximization proceeds with two constraints. First, the fractions of ribosomes allocated to the production of three different proteome sectors have to sum to unity (c1). Second, the surface-to-volume ratio has to satisfy the constraint based on the shape of the cell (c2):

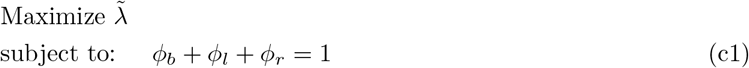

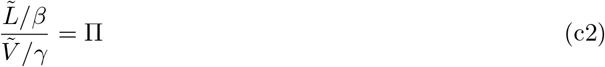

Note that *V* is the cytoplasmic volume, and thus the Π is the ratio of cell surface area to cytoplasmic volume. Parameters *γ* and *β* are unit conversion factors corresponding to the number of amino acids per unit of cell volume (molecs*/μm*^3^), and the number of lipids/envelope components per unit of cell surface area (molecs*/μm*^2^); These values are reported in Table 2. To intuitively understand the optimization problem, consider a landscape of ribosomal partitioning parameters and the resulting growth rates (Fig 2; bottom row). In the absence of geometric constraint (c2), the cell maximizes growth rate by (1) completely abolishing expression of the envelope-producing enzyme (when *ϕ*_*l*_ = 0), and (2) by optimally expressing ribosomes (when 0 *< ϕ*_*r*_ *<* 1); this is because envelope-producing enzyme acts as a burden that diverts resources from other proteome components, and because the cell has to balance the production of proteins (for which high ribosomal expression is required) and the production of building blocks that fuels the translation (for which low ribosomal expression is required). These two aspects explain why the growth rate is a monotonically decreasing function of *ϕ*_*l*_, and a non-monotonic function of *ϕ*_*r*_.

**Table 2.** Parameterization of the model. Standard errors were obtained using error propagation (see Section S1.9). † Surface-to-volume ratio represents the average across different dimensions of *E. coli* in our dataset. * Calculated in the text. All rate parameters are *Q*_10_-corrected as described in text. ° Rate constants estimated from the noted study.

| Symbol | Definition | Value | Units | Source |
| --- | --- | --- | --- | --- |
| $\kappa_n(\text{M63+gly})$ | Nutritional capacity in M63 with glycerol | $0.478 \pm 0.074$ | $h^{-1}$ | [17], $\circ$ |
| $\kappa_n(\text{M63+glc})$ | Nutritional capacity in M63 with glucose | $0.986 \pm 0.112$ | $h^{-1}$ | [17], $\circ$ |
| $\kappa_n(\text{cAA+gly})$ | Nutritional capacity in cAA and glycerol | $1.353 \pm 0.110$ | $h^{-1}$ | [17], $\circ$ |
| $\kappa_n(\text{cAA+glc})$ | Nutritional capacity in cAA and glucose | $1.866 \pm 0.236$ | $h^{-1}$ | [17], $\circ$ |
| $\kappa_n(\text{RDM+gly})$ | Nutritional capacity in RDM with glycerol | $2.325 \pm 0.285$ | $h^{-1}$ | [17], $\circ$ |
| $\kappa_n(\text{RDM+glc})$ | Nutritional capacity in RDM with glucose | $3.449 \pm 0.388$ | $h^{-1}$ | [17], $\circ$ |
| $\kappa_l$ | The envelope synthesis rate | $21.317 \pm 3.951$ | $h^{-1}$ | [29], $\circ$ |
| $\kappa_t$ | The protein synthesis rate | $2.579 \pm 0.175$ | $h^{-1}$ | [29], $\circ$ |
| $\epsilon$ | The cost of cell per unit of S/V | 0.11 | $\mu m$ | * |
| $\Pi$ | Surface-to-volume ratio | 5 | $\mu m^{-1}$ | † |
| $d_p$ | Protein degradation rate | 0.011 | $h^{-1}$ | [25] |
| $d_l$ | Cell envelope degradation rate | 0.211 | $h^{-1}$ | [27] |

Now suppose that the cell maximizes the growth rate, while at the same time having to maintain a surface-to-volume ratio dictated by the cell’s geometry; out of all partitioning parameters, only a subset satisfies this constraint and these parameters fall onto the red line in Fig 2. For example, a cell that has a low expression of the envelope-producing enzyme (the left-most grey point in the landscape) will have a high growth rate but won’t have enough envelope to cover its cytoplasm, thus making it a non-viable option. On the other hand, a cell that has a highly expressed envelope producer (the right-most grey point) will make too much membrane relative to its volume, thus causing the cell to wrinkle. However, a cell with intermediate values of *ϕ*_*l*_ will produce just enough envelope to cover its cytoplasm (the intermediate grey point), and the maximal growth rate is achieved by the adjustment of partitioning parameters along this line (black point in the landscape).

The developed model assumes that the S/V is constant across the cell cycle, which is wrong, given that the cell has to change the shape and size as it grows. Unfortunately, we currently cannot derive a more general model which accounts accurately for changes in shape as the individual cell evolves. However, the variation in S/V across the cell cycle is much smaller than the cross-species range of S/V that we focus on (Section S1.4), so it should not strongly affect the reported derivations.

### 3.3 Explicit solution for maximal growth under saturation kinetics

Although obtaining the analytical solution for maximal steady-state growth rate is prohibitively difficult, one can obtain this property for a special case when all enzymes are saturated, such that chemical reactions obey first-order mass-action kinetics. Indeed, most *E. coli* enzymes have *K*_*M*_ lower than their substrate’s concentrations [18]. This reduces a nonlinear system of Eq 7a–7e to a system of linear equations that can be readily solved for the partition parameters *ϕ*:

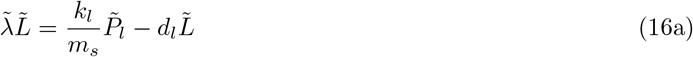

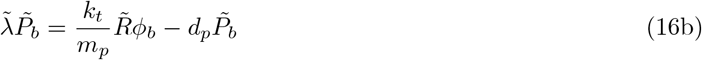

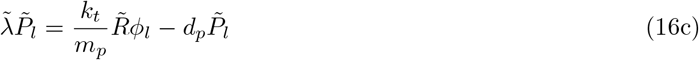

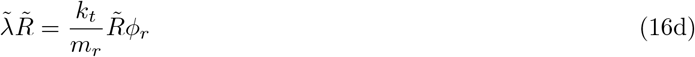

The steady-state growth is then obtained from Eq 11b by setting *K*_*M*_ = 0:

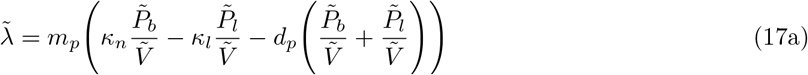

Substituting Eq 17a in Eq 16a–16d:

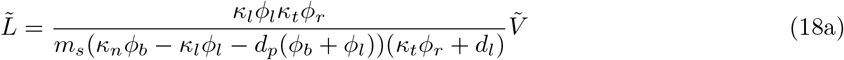

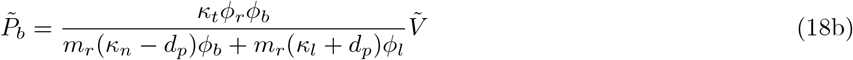

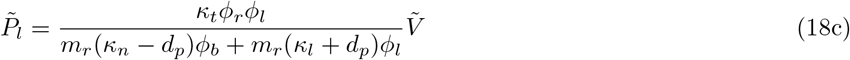

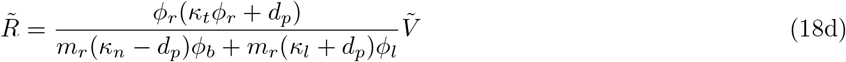

By substituting abundances in Eq 9 for Eq 18b–18d It immediately follows that the mass fractions of protein sectors are:

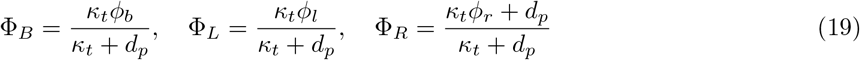

Note that substitution of Eq 18a–18d in Eq 17a yields 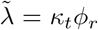, meaning that the growth rate is simply proportional to the fraction of ribosomes that are allocated to ribosome translation. The cell, however, cannot allocate the entirety of its ribosomes to this task because this would halt the production of other necessary protein components. Therefore, to find the optimal partitioning of ribosomes that maximizes the growth rate, one has to impose additional algebraic constraints. The first algebraic constraint reflects the fact that maximal growth is achieved when the building block influx matches the outflux. If influx is higher than the outflux, then buidling blocks unnecessarily accumulate in the cell, and if the outflux is higher, then the building block pool will become completely depleted and the chemical reactions will halt. Because *B* is being produced and consumed at matching rates, the building block pool does not grow over time, and we retrieve the constraint by setting the left-hand side of Eq 7a to zero and *K*_*M*_ = 0. The second expression imposes a constraint on the amount of resources that has to be diverted into cell envelope to ensure a proper cell shape:

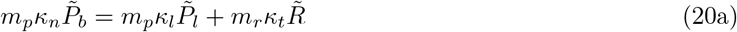

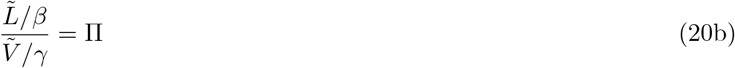

To obtain optimal partitioning parameters *ϕ* that satisfy these constraints, we substitute 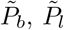, and 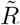 with solutions 18a–18d. While there are three *ϕ* parameters, there are only two degrees of freedom given that *ϕ*_*b*_ = 1 −*ϕ*_*r*_ −*ϕ*_*l*_. Thus, we have a system of two linear equations in two unknowns, which we solve to obtain the optimal partitioning of the proteome such that the flux of resources is balanced between catabolic and anabolic processes, and the cell has a proper shape given its size:

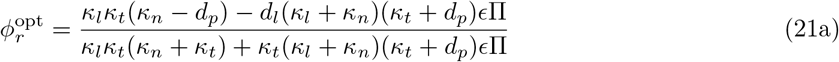

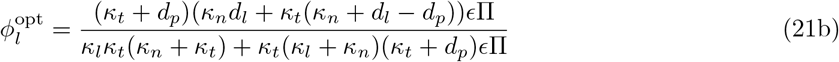

Substituting 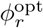 and 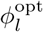 in Eq 18a–18d, one retrieves molecular abundances in terms of model parameters. Substitution of the same into Eq 19, allows one to obtain expressions for optimal proteome mass fractions. Finally, substitution in formula for 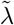 (Eq 17a) yields a maximal steady-state growth rate 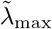:

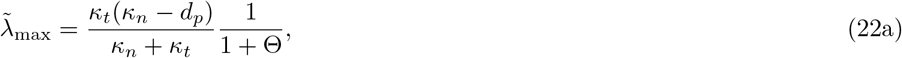

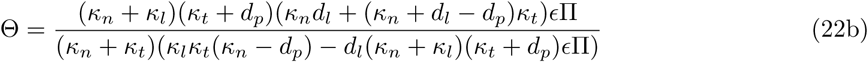

Where *ϵ* = *m*_*s*_*β/γ* is the resource cost of the unit S/V, or the resource investment in a unit of surface area per unit of cytoplasmic volume. For example, if *ϵ* = 2, then each added unit of the surface area requires twice as many amino acids relative to the added unit of cytoplasmic volume.

Intuitively, the first term in Eq 22a is the steady-state growth rate in the limit of no additional resource sink, and the second term represents a deviation from maximal achievable growth due to cell envelope production. The parameter Θ is the bioenergetic cost of cell envelope production, which depends not only on the actual amount of resources that go into this cellular feature (*ϵ*Π) but also on the rates of all cellular processes. Previous theoretical developments represent a special case of Eq 22a. In the limit of no investment into cell envelope (Π = 0), Eq 22a reduce to those reported in [15], and taking this further to the case without protein degradation (*d*_*p*_ = 0) yields the result in [17].

While the explicit solution for Θ appears complicated, some insight can be gained by looking at the special case when there is no degradation (*d*_*l*_ = *d*_*p*_ = 0):

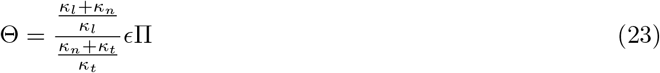

The part *ϵ*Π corresponds to the resource cost of producing the entire envelope structure. Parameter *ϵ* can be interpreted as the envelope cost of the cell with Π = 1 *μm*^−1^. This is because *γ* is the total number of amino acids per unit volume and *m*_*s*_*β* is the total number of amino acid equivalents required to build the unit of the cell surface. Given that Π is the surface-to-cytoplasmic volume ratio, the whole term *ϵ*Π is the cost of producing surface relative to the whole amino acid budget of the cytoplasmic volume. The fractional term in Eq 23 captures the cost of producing the enzyme machinery that builds the actual envelope. One cellular process supplies the building blocks at the per unit proteome rate *κ*_*n*_, and two other cellular processes are competing for this common pool at the per unit proteome rates of *κ*_*l*_ and *κ*_*t*_. For instance, if *κ*_*l*_ is decreased, the cell maximizes growth rate by overexpressing cell envelope-producing machinery in order to compensate for the low per per unit proteome rate, thus increasing the total costs of the envelope.

The expressions for proteomic mass fractions are retrieved by substituting partition parameters in Eq 19 for optimal partition that maximize the growth rate (Eq 21a and Eq 21b):

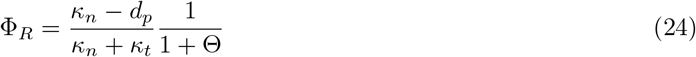

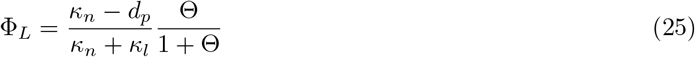

One can immediately see that ribosomal mass fraction is monotonically decreasing and envelope-producer mass fraction is monotonically increasing function of the surface-to-volume ratio, Π.

The analytical solution for 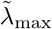, and optimal Φ_*R*_ and Φ_*L*_ in the saturation limit was cross-validated by comparison to the numerically maximized 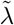 with algebraic constraints of the finite ribosomal pool (c1) and geometric constraint on the cell shape (c2), as described in Section 3.2, and optimization was performed using Nelder-Mead algorithm. To ensure that the optimizer obtains the global maximum, we repeated the optimization 20 times for each set of model parameters. Each iteration started by randomly seeding the points of the polytope. We take the highest 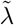 as the solution of the optimization problem. We generally find an excellent correspondence between analytics and numerics for both the growth rate and proteome composition (right panel in Fig 3). When enzymes in the model operate far from the saturation limit (i.e., when *K*_*M*_ 0), the analytics break down (see inset). Note that large *K*_*M*_ has a similar effect on scaling as a reduction in nutrient quality of the media. Therefore, although the analytic solution neglects the presence of the aqueous phase – metabolites and associated water molecules – inside the cell, the incorporation of this property affects the intercept but not the overall scaling pattern. For a fuller analysis of growth-impeding effects of cell envelope, see Section S1.1.

**Figure 3.**
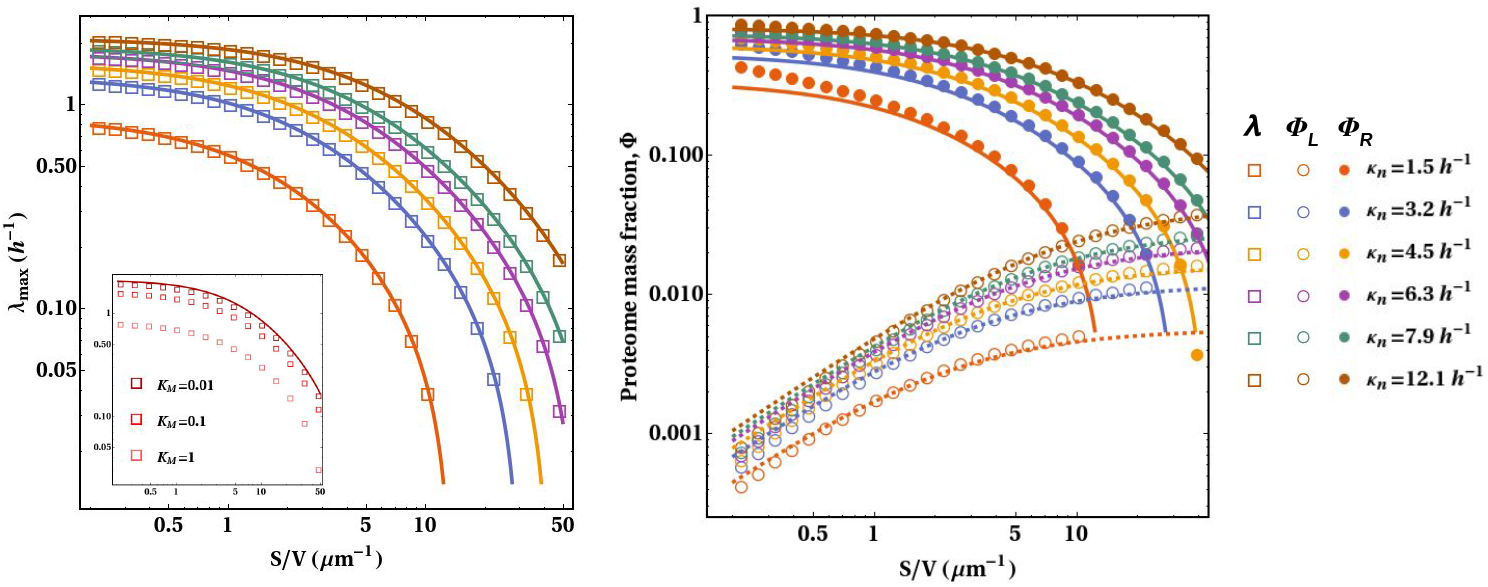
Maximal steady-state growth rate when all enzymes are saturated. Left panel: Growth rate scaling with S/V of the cell. Solid lines denote the analytical solution for 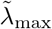 (Eq 22a), while squares represent numerically maximized growth rate (Eq 14). Different lines correspond to different values of *κ*_*n*_ (reported in the legend on the right), while all other model parameters are identical. Right panel: Proteome composition scaling with the cell shape. Colored lines signify the optimal ribosomal (solid lines) and envelope-producer (dashed lines) mass fractions as a function of Π. Full and open circles denote the same quantities that maximize growth rate in the numerical optimization problem. Other parameters: *κ*_*l*_ = 288 h^−1^, *κ*_*t*_ = 2.59 h^−1^, *d*_*p*_ = 0.14 h^−1^, *d*_*l*_ = 0.17 h^−1^, *K*_*M*_ = 0.001, *γ* = 9.75 × 10^8^ aa*/μm*^3^, *β* = 2 × 10^6^ lip*/μm*^2^, *m*_*s*_ = 314 aa.

### 3.4 Parameterization of the model

Eq 22a has six parameters (*κ*_*n*_, *κ*_*l*_, *κ*_*t*_, *ϵ, d*_*l*_, *d*_*p*_) and one variable (Π). Our goal was to constrain the rates of chemical reactions to the values occurring in *Escherichia coli*, and then ask how the growth rate would scale if the variable Π was altered. That is, we are interested in understanding how the growth rate would scale if all bacterial species were biochemically identical to an *E. coli* cell, and only differed in cell size.

We estimated the resource cost of the envelope in a cell with a unit S/V, *ϵ*, by assuming that resource investment is equal to the mass *M* of a structure. Given that our model consists of the whole envelope (lipids, peptides, and sugar attachments) with mass *M*_env_ and proteins of mass *M*_prot_, and that envelope and proteins account for roughly 30% and 55% of total cell mass *M*_*T*_ (see [19], and Chapter 2 in [20]), and Π = 5 (Table 2):

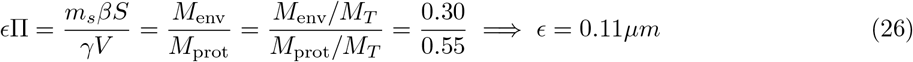

This is a crude estimate because it does not include the direct costs of envelope production and proteome components (i.e., resources needed to convert one chemical compound into another one), but these usually account for a small fraction of the total costs [21], and small changes in *ϵ* do not affect our conclusion.

The degradation rates are estimated separately from the literature. We assumed that the ribosomal degradation rate is zero, and we justify this assertion by three observations. First, ribosomal rRNA is remarkably stable in the exponential and stationary phases, and the degradation occurs only when the culture transitions between these two growth stages [16, 22]. Second, almost all ribosomal proteins have degradation rates close to zero [23]. Third, ribosomal proteins are mutually exchangeable when damaged, meaning that replacement is favored over degradation and re-synthesis [24]. The protein degradation rate was set to 0.05 per hour [25]. We also tried out estimates for various protein classes [26] but found little variation in the scaling pattern (Fig S1.7). The envelope degradation rate *d*_*l*_ is difficult to estimate exactly due to the diverse components that make up this structure, but was set to 1 per hour [27]. This study reports a hypermetric scaling of the peptidoglycan turnover and the growth rate (*d*_*l*_ = 0.7 × (Log[2]*/τ*)^1.38^). Assuming that an *E. coli* cell divides in 30 minutes, we have *d*_*l*_ *≃*1. Unfortunately, this is a very crude estimate based on the peptidoglycan degradation rate in *B. subtilis*, which constitutes a large part of its envelope. Peptidoglycan degradation rates are somewhat lower in *E. coli* [28] but on the other hand, that species has an outer membrane containing polysaccharides, and we do not know how fast this component is degraded.

The rate constants of chemical reactions (*κ*_*t*_, *κ*_*l*_, *κ*_*n*_) were inferred from *Escherichia coli* proteomic data across different growth conditions, leveraging growth laws derived in Section S1.2. We ignore the intercept of regressions (Eq S1.11a–S1.13b) and infer parameters strictly from the slopes of these relationships. The slope of the regression of the growth rate on the mass fraction of ribosomal proteins allows one to compute *κ*_*t*_ from Eq S1.11a. Similarly, the value of *κ*_*l*_ can be computed from the slope of the regression of the growth rate on the mass fraction of envelope-producing enzymes and Eq S1.11b. Our data for growth rates across different bacterial species are normalized to 20°C using *Q*_10_ correction with the coefficient of 2.5. Hence, we also normalized the inferred rate constants using the same method. Estimates of rate constants for *E. coli* are reported in Table 2.

Because *κ*_*n*_ captures both the intrinsic efficiency of metabolism to convert nutrients into building blocks and the nutrient state of the external environment, there will be one *κ*_*n*_ for every medium that *E. coli* is reared in. Intuitively, one would expect minimal media to have lower *κ*_*n*_ than rich media, as the latter contains better quality nutrients and thus leads to more amino acids being generated per unit time per molecule *P*_*b*_. The parameter *κ*_*n*_ is calculated from the slopes of ribosomal mass fraction across growth conditions with varying concentrations of translation inhibitor, after plugging in values of *κ*_*l*_, *d*_*p*_, and *ϵ*Π in Eq S1.12a. The values are computed for the following media: M63 with glycerol, M63 with glucose, casamino acids with glycerol, casamino acids with glucose, rich defined media with glycerol, and rich defined media with glucose.

One can intuitively explain the estimated rate constants by using logic outlined in [30]. It takes roughly 10^9^ glucose molecules to produce the entire carbon skeleton of an *E. coli* cell. Given that an amino acid has, on average, about the same number of carbon atoms as a single glucose molecule (5.35 C-atoms), one could say that the cell requires 10^9^ amino acid-equivalents for its construction. Therefore, with roughly 3 × 10^6^ copies of metabolic proteins in the cell, each building block producer making *k*_*n*_ = *κ*_*n*_×*m*_*p*_ = 2 325 amino acids per hour, it will take ~30 minutes for the metabolic proteins to generate enough amino acids to replace the cell. This is also the experimentally measured cell division time of an *E. coli* cell grown under favorable conditions. For the protein synthesis rate (i.e., the elongation rate) *κ*_*t*_, note that each ribosome converts *κ*_*t*_ × *m*_*r*_ amino acids into proteins (where the length of the ribosome *m*_*r*_ is 7336 amino acids). This means that the inferred translation rate *k*_*t*_ in our model is about 25 amino acids per ribosome per second, which is close (albeit slightly larger) to empirical values for an *E. coli* cell [20]. Length of metabolic protein was taken to be the median length of an *E. coli* protein (325 aa from [31]), and ribosome length is set to the total number of amino acids in ribosomal proteins (7336 aa from [32]).

## 4 Results

The model of the cell developed in section 3 purports to explain two phenomena: (1) the optimal partitioning of the proteome when the same bacterial species is reared under different growth conditions; and (2) the scaling of the growth rate and proteome composition across bacterial species of different shapes and sizes. Both sets of predictions are tested in the ensuing sections. Applying the developed theory to data on growth and proteome composition of the model organism *Escherichia coli*, we first show that our model yields a good qualitative description of physiological responses to changes in the environment. We then use this correspondence between theory and data to estimate the parameter values that yield a good quantitative fit as well (Section 4.1).

By parameterizing our model with values obtained from *E. coli*, one can address how the growth rate is supposed to scale with the shape and size of the cell (Section 4.2). This implicitly assumes that all bacterial species are biochemically identical to *E. coli*, such that the rates of all chemical reactions are the same. Lastly, in an attempt to simplify the expectation and remove dependence on parameters that are difficult to estimate, we look at the special case when degradation is absent, and the envelope is costly enough such that 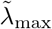 is inversely proportional to Π (Section 4.3).

### 4.1 Proteome reallocation across growth conditions

The internal homeostatic mechanisms allow the cell to allocate the proteome to different cellular tasks such that the growth rate is maximized. While *E. coli* cells achieve this via alarmone ppGpp [33], our model is agnostic of the exact mechanism and simply assumes that such resource-tuning strategy exists. One can express the proteome mass fraction of a particular component as a linear function of growth rate when the latter is altered via changes in growth conditions (derived in section S1.2). Given that there are three cellular processes in our representation of the cell (building block production, cell envelope synthesis, and protein synthesis), we ask how the proteome composition changes when each process is perturbed. More precisely, we are interested in how the proteome composition changes as the growth rate is modulated by changing the values of *κ*_*n*_, *κ*_*l*_, or *κ*_*t*_.

The first expectation is that the ribosomal mass fraction of the proteome Φ_*R*_ scales with the growth rate across different nutrient conditions 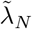 as:

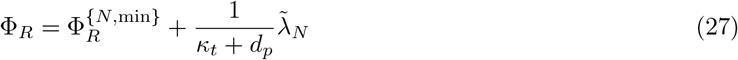

The cell re-balances building block supply and biosynthesis demand by allocating proteome to the limiting process, which is, in this case, protein synthesis. The proteomic data for *E. coli* qualitatively corroborate this expectation (Fig 4, upper left panel). Data from different studies appear to show a small amount of variation, but the overall trend is strong. The second expectation is that the envelope-producer fraction increases with the growth rate under nutrient perturbation:

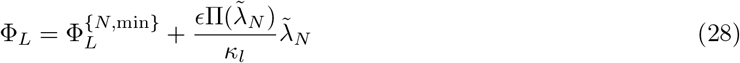

**Figure 4.**
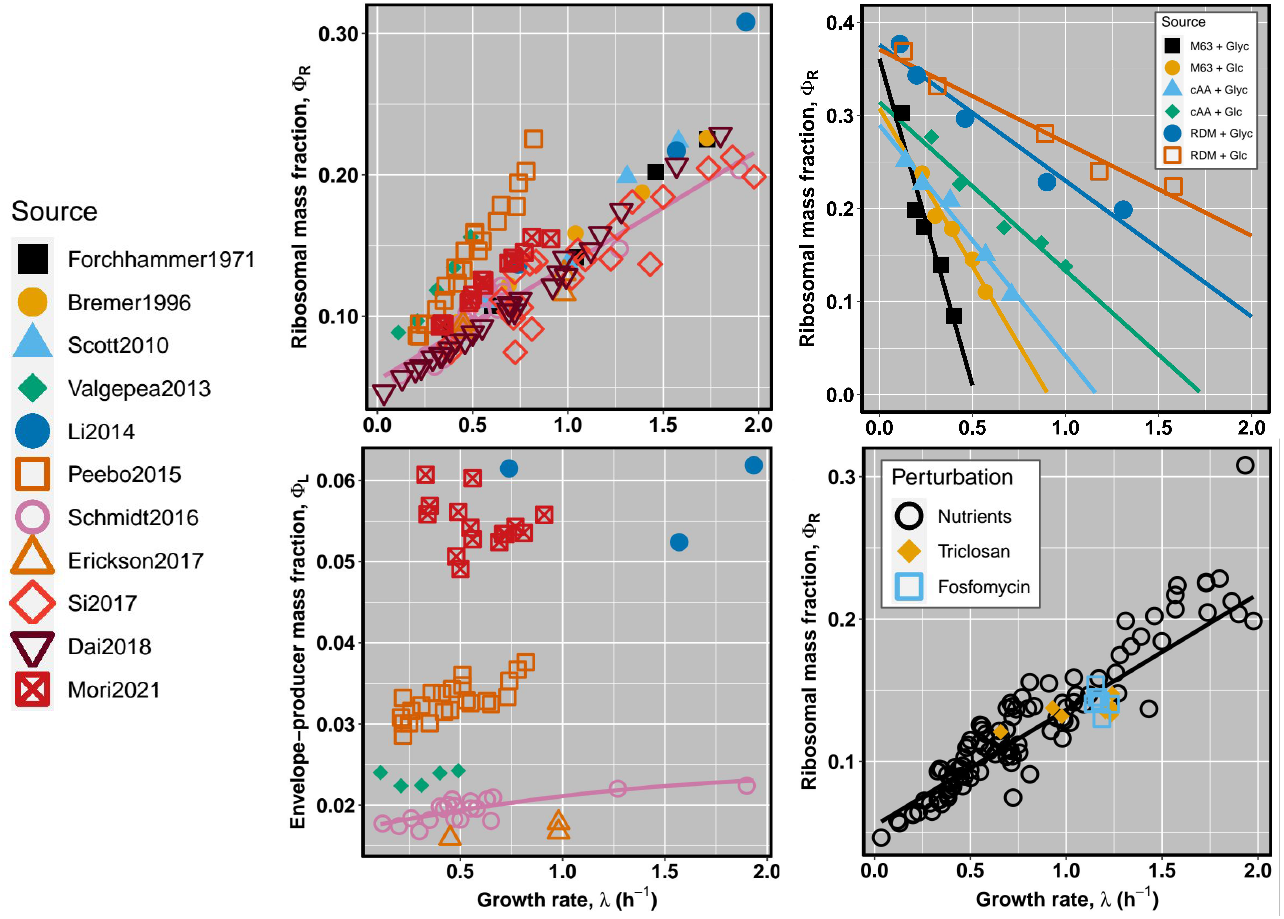
Comparison of empirical proteome composition changes with theoretical expectations. On the left, upper and lower panels represent Φ_*R*_ and Φ_*L*_ across various studies reported in the legend. Upper right panel are data on Φ_*R*_ under chloramphenicol treatment obtained from [17]. Each line corresponds to a different media as outlined in the corner legend. Lower right panel contrasts changes in Φ_*R*_ across nutrient conditions (orange points) with Φ_*R*_ changes under envelope producer inhibitors (black and grey dots), with data from [36]. Lines signify the ordinary least squares fit to the data from [17] (upper right panel), and [29] all other panels.

The slope captures the fact that faster growth implies that a cell has to produce its envelope faster. Given that each envelope-producer operates at a constant rate, the only way this constraint can be met is by allocating more enzymes to this task.

When inferring parameters, one must account for the fact that Π might depend on 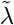. Π is largely independent of the growth conditions when the latter is varied with translation-inhibiting antibiotic (Fig S1.5). However, Π decreases with the growth rate when the bacteria are cultured with different carbon sources [34–36]. We account for this confounding factor by plugging the equation for the relationship 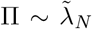 derived from empirical studies into Eq 28, and then infer *κ*_*l*_ using the non-linear least squares method (see Section S1.5 for details). While the proteomic data on envelope-producer component is less accurate compared to ribosomal proteins (probably owing to low abundance of envelope-producers), the predicted positive scaling still occurs in the two most comprehensive proteomic studies (pink circles and orange squares in lower left panel of Fig 4).

The third expectation is concerned with scaling of ribosomal mass fraction under translation inhibition:

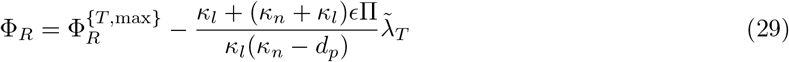

As the concentration of a translation-inhibiting drug is increased, so is the fraction of inhibited ribosomes. The cell compensates for this inhibition by overexpressing ribosomes, which causes a negative scaling between Φ_*R*_ and 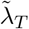. We see a good fit to data (upper right panel in Fig 4), and similar relationships can be obtained by using the alternative source of data from [36] (Fig S1.5).

The fourth prediction is that the response of Φ_*R*_ to nutrient quality perturbation is the same as the response to envelope-synthesis inhibition. This follows from the fact that the relationship

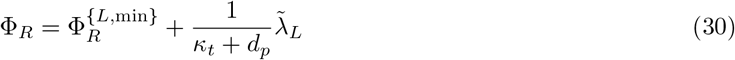

is identical to Eq 27. Thus, the prediction is that points from two types of perturbation experiments ought to fall onto the same line. Indeed, the treatment of *E. coli* culture with fosfomycin (peptidoglycan synthesis inhibitor) and triclosan (fatty acid synthesis inhibitor) reduces both the growth rate and the ribosomal mass fraction (lower-right panel in Fig 4). One can intuitively understand this behavior by noting that the cell responds to the envelope-synthesis inhibition by overexpressing envelope-producer to compensate for the poisoning of the enzymes, which reduces the resources available for ribosomal proteins thus causing the reduction in their expression.

### 4.2 Scaling behavior in a fully specified model

The theory is tested by comparing the expected to the observed scaling relationships using growth, cell shape, and proteomic data across heterotrophic bacteria. Details on data collection and normalization are reported in Section S1.6 and S1.7. This requires specifying model parameters. Because we are dealing with a somewhat metabolically homogeneous set of species (i.e., all are heterotrophs), we assume that all the cellular reaction rates are identical, and the only feature that varies across bacteria is Π. Furthermore, most of the maximal growth rates in our dataset were measured in the media containing yeast extract, so we assume that *κ*_*n*_ is fixed across species as well. The prerequisite for these assumptions to hold is that the type of metabolism of a bacterial species does not correlate with cell size and shape. That is, the variation in metabolism and external environment might affect dispersion around the scaling but does not affect the scaling itself.

The model is determined by six parameters: rate constants of three types of enzymes (*κ*_*n*_, *κ*_*l*_, *κ*_*t*_), degradation rates (*d*_*p*_, *d*_*l*_), and resource cost of the unit S/V (*ϵ*). The degradation rates of lipids, peptidoglycan, and proteins are set to values determined in pulse-chase experiments obtained from the literature (see Section 3.4). We take the energetic cost of the envelope, *ϵ*, from our earlier calculation (Eq 26). Rate constants are not only related to turnover numbers of respective enzyme classes but also depend on other cellular properties that are not explicitly accounted for in the model, such as the nutrient status of the environment, the number of enzymes belonging to a particular sector, the topology of a particular biochemical pathway, the amount of nucleic acid in the cell, and so on. Thus, by estimating rates from the empirical relationships we derived in Section 4.1, one can “collect” the effects of the explicitly unaccounted cellular properties into these three parameters.

The protein synthesis rate, *κ*_*t*_, can be estimated from the slope of Eq 27 if the degradation rate *d*_*p*_ is known. The envelope synthesis rate, *κ*_*l*_, is retrieved from the slope in Eq 30, if the envelope cost *ϵ*Π is known. And finally, the nutrient processing rate *κ*_*n*_ is inferred from slopes of changes in Φ_*R*_ as the growth rate is perturbed by the addition of translation-inhibiting drug chloramphenicol (Eq 29), after plugging in all of the previously-estimated parameters. Because *κ*_*n*_ also depends on the nutrient quality, the values were inferred for various media ranging from nutrient-depleted (M63 with glycerol) to nutrient-replete (RDM with glucose). All rate constants were estimated by fitting physiological scaling relationships to *E. coli* proteomic data. We first classified each protein into one of the three sectors based on its function (for details, see Section S1.7), and then pooled their individual masses to compute the total mass fractions Φ_*R*_ and Φ_*L*_. The surface-to-volume ratio was calculated from the linear dimensions of the cell (Section S1.3). Cytoplasmic volume was calculated by first subtracting 2× thickness of the bacterial envelope from the linear dimensions of the cell, where the latter was taken to be 30 nm [37].

We find that the curves parameterized with *κ*_*n*_ from minimal (black) to rich media (orange) capture most of the variation in growth rate and proteome composition data (Fig 5). Hence, in principle, one could attribute the scatter along the y-axis to differences in efficiencies in which bacteria take up and convert the nutrients into building blocks. Furthermore, Φ_*L*_ is expected to have a slightly positive and Φ_*R*_ a negative scaling with Π. The prediction is not unambiguously corroborated by the data. A large amount of variation in proteomic data around the expectation might reflect growth and temperature differences which we do not know how to correct for. Also, Φ_*L*_ is a low abundance proteome sector as it accounts for no more than a few percent of total proteome mass. These two effects might jointly make the data unreliable.

**Figure 5.**
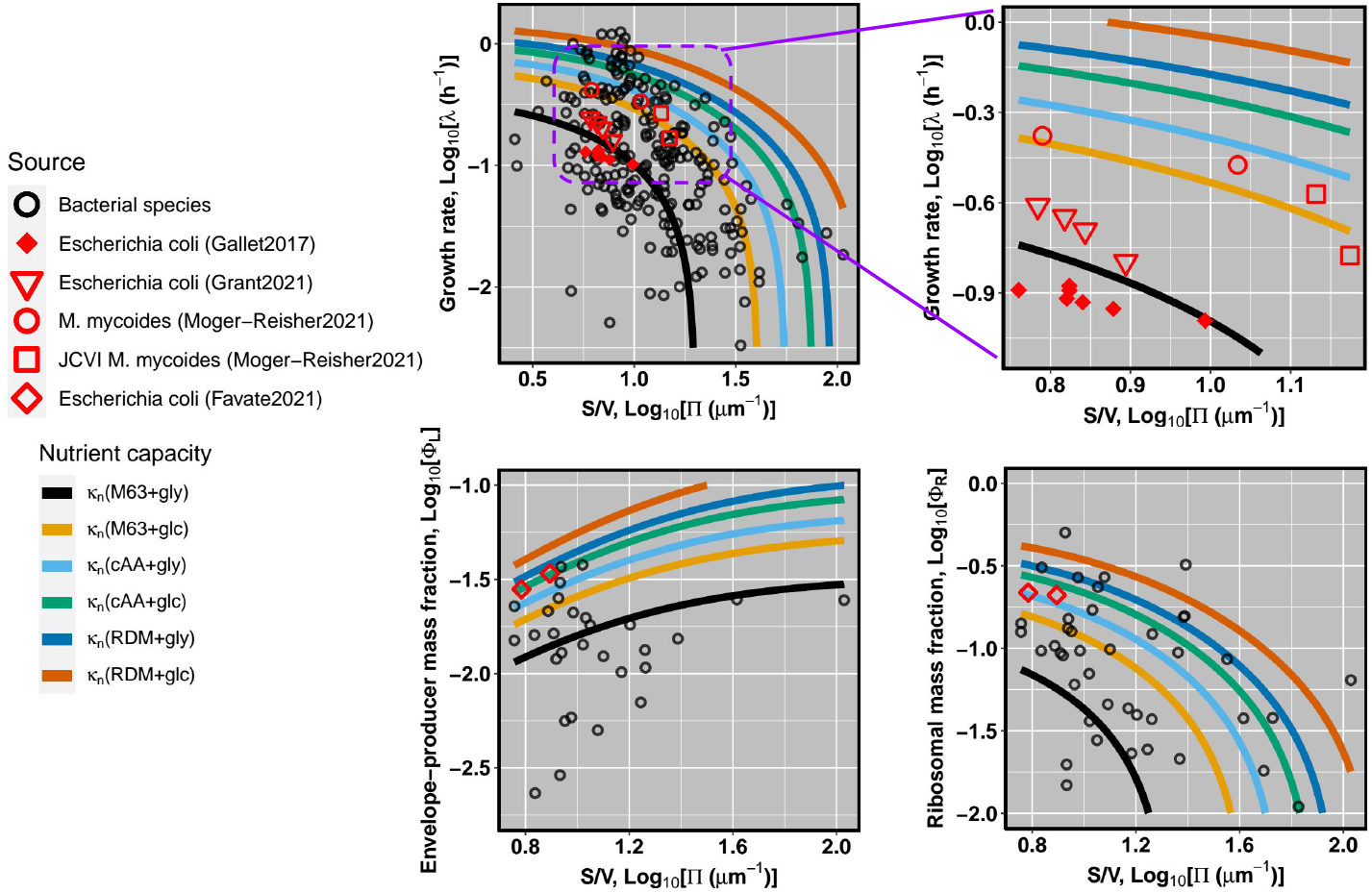
The comparison of expected and observed scaling relationships. Upper row: scaling of growth rate with S/V across bacterial species (left pane), and scaling in long-term evolution experiments (right pane); colored curves denote expected theoretical scaling given by Eq 22a and parameterized with *κ*_*n*_ estimated from various media, as reported in Table 2. Lower row: scaling of envelope-producer (left pane) and ribosomal mass fractions (right pane).

A large variation in growth rate data across bacteria might also stem from variation in morphology (e.g., motility structures), metabolism (e.g., aerobic or anaerobic capabilities), or environment (e.g., from oceanic sediments to digestive tract of mammals). Ideally, one would want to compare the growth rates across species with the same type of metabolism and the same environment and only differ in shape or size. This way, all of the morphological, metabolic, and environmental factors would be controlled, and the entirety of variation in growth can be attributed to variation in size. We attempted to control for these issues by using data from experimental evolution studies, where cells were kept in approximately exponential phase for a long period of time. In the wild, cell size and shape are determined by a myriad of ecological factors [38]. When these cells are taken out of their natural habitat and propagated in the nutrient-rich medium, the ecological factors dictating the size of the cell are removed. For instance, large cells might be deleterious *in situ* owing to size-selective predation but they do not suffer the same disadvantage when reared in the laboratory setting where predators are absent. Under laboratory selection for fast growth, a mutant with lower S/V has a growth advantage over the wild-type due to lower relative investment in envelope, and will spread and take over the population. In this situation, our theory predicts three outcomes. First, increasing growth rate should be accompanied by a reduction in S/V. Second, the mass fraction of envelope-producers should decrease because the envelope synthesis becomes a less of an obstacle with inflation of the cell size. Third, given a diminished need for synthesizing envelope-producers, a larger mass fraction of the proteome can be allocated to the ribosomal proteins.

To test the first prediction, we analyzed the scaling of the growth rate with S/V in Lenski’s long-term evolution experiment in *Escherichia coli* and with a more recent experimental evolution of *Mycoplasma mycoides*. First, we used S/V in combination with the relative fitness data for four LTEE time points spanning 50,000 generations [39]. Given that the relative fitness is the ratio of Malthusian parameters of evolved to ancestral line determined from the head-to-head competition (see [40]), we converted this metric to the growth rate by multiplying it with the growth rate of the ancestral strain (see *V*_max_ in [41]). However, because the competition experiment lasts for one day, competing strains enter into stationary phase after a few hours, so the competitive advantage of the evolved line might reflect not only differences in the growth rates but also the ability to survive in the stationary phase. Thus, we sought to cross-validate our approach by including data from [42], which reported the growth rates and cell volumes across LTEE. Cell volumes were converted to S/V using the empirical scaling *S* = 2*πV* ^2*/*3^ reported to hold across many bacterial species [43]. Second, we collected relative fitness and cell size of wild-type *Mycoplasma mycoides* and *M. mycoides* whose genome has been minimized by removal of non-essential genes [44]. Comparing LTEE data with our analytical model (Fig 5), we find that the observed scaling falls very near to the expected scaling for *E. coli* grown in the media with similar composition (M63) as that used in LTEE (Davis broth with glucose). Note that both indirect estimates (red points) and direct measurements (green points) of the growth rate fall onto the same line. A similar trend is observed in the experimental evolution of *M. mycoides*.

The latter two predictions were tested using data on the changes in the absolute transcript abundance over the course of the long-term experimental evolution in *E. coli* [45]. To convert transcript abundance into protein abundance, we first computed the number of proteins per transcript for each *E. coli* gene using data in [46], and then multiplied transcript abundance of each gene in long-term evolution experiment with the corresponding conversion factor. One can then obtain the mass fraction of a particular protein by multiplying the abundance with its molecular mass and normalizing by the total mass of the proteome. We find that the average ribosomal mass fraction slightly increases (0.210 ±0.02 vs. 0.218 ±0.01), while the average envelope-producer mass fraction decreases (0.034± 0.001 vs. 0.028 ±0.0008) during 50,000 generations of the experiment. This roughly coincides with the predicted scaling (orange and blue lines in Figure 5). It is currently unclear why the mass fractions do not adhere to the same line with the growth rate (black line); Perhaps this reflects differences between strains, given that Lenski used derivates of B strain while the expected scalings were plotted using parameters inferred from experiments with BW25113 and a strain derived from MG1655.

### 4.3 Scaling in the limit of large envelope costs and no degradation

Given the uncertainty of model parameters (especially envelope degradation), we attempted to simplify the theoretical expectation by focusing on the simplest possible case when degradation is absent, and the costs of the envelope are sufficiently high that Eq 22a can be approximated as 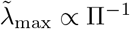, Eq 21a as Φ_*R*_ ∝ Π^−1^, and Eq 21b as Φ_*L*_∝ Π^0^. While this scenario might not be the most realistic, the benefit is that it does not require any further specification of model parameters because the prediction is that the slopes of the regression Log_10_ 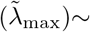 Log_10_(Π) and Log_10_(Φ_*R*_) ~Log_10_(Π) should be −1, and the slope of Log_10_(Φ_*L*_) ~Log_10_(Π) should be zero. Assuming that the shape is constant across bacteria, similar scalings can be obtained with cell volumes as the independent variable – given that Π ∝ *V* ^−1*/*3^. One expects Log_10_ 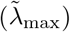~0.33 Log_10_(*V*), Log_10_(Φ_*R*_) ~0.33 Log_10_(*V*), and Φ_*L*_ retains independence as with S/V. We use these expectations as the null hypothesis for the slopes in the regression analysis. In addition to scaling expectations, we also wish to test whether surface-to-volume is a better predictor of the growth rate than the cell volume. To this end, we performed OLS regression with either S/V or V as independent variables, and the growth rate, or proteomic mass fractions as the dependent variable of various bacterial species (Table 3). While our theory relates the growth rate to the ratio of the surface area to cytoplasmic volume (S/V_cyt_), we also examined whether the dependent variables scale with surface area to the total cell volume (S/V_tot_). In total, the analysis included the growth rate data from 229 species, and ribosomal and envelope-producer mass fractions from 41 and 30 species, respectively.

**Table 3.** Observed and expected scaling exponent of the growth and proteome composition. Dependent variables are denoted in the first row, while independent variables are listed in the first column. Each table cell contains the observed slope with standard error, while the expected scaling exponent is reported in the parentheses. Blue color highlights the cases where the observed slope is not significantly different from the expected (i.e., where *p >* 0.05). Sample size: *n*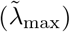 = 229, *n*(Φ_*R*_) = 41, *n*(Φ_*L*_) = 30.

| | $\tilde{\lambda}_{\max}$ | $\Phi_R$ | $\Phi_L$ |
| --- | --- | --- | --- |
| $S/V_{\text{cyt}}$ | $-1.012 \pm 0.115$ (−1) | $-0.50 \pm 0.20$ (−1) | $0.186 \pm 0.247$ (0) |
| $S/V_{\text{tot}}$ | $-1.247 \pm 0.149$ (−1) | $-0.927 \pm 0.252$ (−1) | $0.129 \pm 0.37$ (0) |
| $V_{\text{cyt}}$ | $0.067 \pm 0.059$ (0.33) | $0.064 \pm 0.105$ (0.33) | $-0.04 \pm 0.139$ (0) |
| $V_{\text{tot}}$ | $0.019 \pm 0.061$ (0.33) | $0.146 \pm 0.115$ (0.33) | $0 \pm 0.151$ (0) |

Five insights emerge. First, S/V_cyt_ is a moderate predictor of growth rate (adjusted *R*^2^ = 0.25), while cell volume appears neither to match the expected 0.33 scaling nor to show any correlation with the growth rate. Second, the observed scaling of growth with S/V is not significanly different from −1. Third, the scaling of proteomic mass fractions with S/V largely meets the expected slopes of −1, for Φ_*R*_, and 0 for Φ_*L*_. Fourth, both S/V_cyt_ and S/V_tot_ perform equally well, with S/V_cyt_ being a slightly better predictor of growth than S/V_tot_ (adjusted *R*^2^ of 0.25 vs. 0.23) and the scaling exponent being within one standard error of the estimated −1 in the case of the former. Fifth, Φ_*R*_ scales isometrically with S/V_tot_ but not with S/V_cyt_. Graphic representation of regression analysis is reported in Section S1.11.

We cross-validated these results by analyzing a recently-published study [47], which contains data on linear dimensions of the cell and the minimal doubling time. Although we do not find any correlation between cell volume and the growth rate, we find a very weak negative relationship (adjusted *R*^2^ = 0.08) between S/V and the growth rate (Fig S1.11) with a scaling exponent of −0.97 which is not significantly different from −1 (*p* = 0.92). Given that bacterial species with large S/V (i.e., small coccal or thin helical cells) are frequently parasitic, it is possible that the pattern is mainly driven by large-S/V species growing more slowly because of their natural habitat and not because of trade-offs in investments. If that is the case, then excluding parasitic species from the dataset would eliminate negative scaling. To control for this confounding factor, we separated data into free-living and host-associated species and then performed the regression analysis (see Section S1.12). We find a very weak negative relationship between S/V and the growth rate in both of these datasets and pooled data, indicating that preponderance of small-celled species in pathogens is unlikely to account for the scaling pattern.

Thus far, we have assumed that all species have an envelope that is ~30 nm thick, like that of an *E. coli* [37]. However, larger bacterial species may have thicker envelopes that require higher resource investments. If resource investments stay the same across the size range, then the variation in growth cannot be explained by the invariant S/V. To account for this possibility, we collected data on envelope thicknesses across 45 species and used these values to compute Π (see Section S1.10). We find that 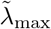 scales with Π with exponent − 1.15 (Fig S1.8) which is not significantly different from −1 (*p* = 0.422). Similarly, 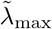 scales with *V*_cyt_ with exponent 0.37 (Table S1.6) which is not significantly different from 0.33 (*p* = 0.749). Therefore, the inclusion of envelope thickness data does not alter the conclusions reached by using fixed thickness.

Three conclusions are reached from the preceding analysis. First, Π is a moderate predictor of 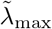, and it accounts for roughly a quarter of growth rate variation in the whole dataset. Second, S/V_cyt_ is slightly better predictor of growth rate than S/V_tot_. Third, the proteome composition qualitatively – but not quantitatively – fits the expected pattern. The ribosomal mass fraction of the proteome scales negatively with Π, albeit with a slope that is shallower than the expected −1, and the envelope producer mass fraction appears independent of Π, which is in accordance with theory.

### 4.4 Scaling of the metabolic rate with S/V

While the central trait in the preceding analysis is the growth rate, one can also obtain the expression for the metabolic rate, and the scaling of this feature can be further used to test our theory. The total metabolic rate of the cell that is in the steady-state, 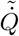, is the sum of the rates of all cellular reactions:

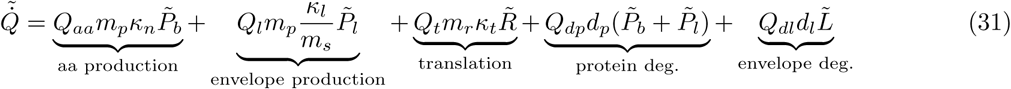

where *Q*_*x*_ corresponds to the number of ATPs hydrolyzed by the process *x*, which can be amino acid production, cell envelope prodution, translation, protein degradation, and degradation of cell envelope. Substituting 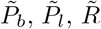, and 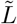 for optimal abundances yields the steady-state metabolic rate of the cell as a function of the rate constants and other model parameters. The full expression is cumbersome, and is only reported in supplemental *Mathematica* notebook. The steady-state metabolic rate 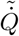 depends on the parameters reported in Table 2, and on additional parameters indicating the amount of ATP used by each reaction are needed. These are estimated in supplemental section S1.14, and reported in Table S1.9. The expected scaling of the steady-state metabolic rate is tested by using the measurements of the metabolic rate over the course of long-term experimental evolution in *E. coli* [48]. Note the cell volume measured in the original study was converted to S/V using the empirical scaling *S* = 2*πV* ^2*/*3^.

The analysis yields three insights. First, the observed scaling roughly matches the expected scaling relationship (Fig 6, left pane), meaning that one can predict the metabolic rate from our model without any additional fitting parameters. Second, both growth rate and metabolic rate exhibit scaling of an organism that was reared in the same medium. That is, growth and metabolic rate in LTEE fall close to the black line in Fig 5 and Fig 6, meaning that a single nutritional capacity parameter *κ*_*n*_ can account for the full behavior of the model. Third, the theory suggests the mechanism behind the scaling of metabolic rate. Namely, smaller ancestor has larger S/V and thus invests a larger portion of its metabolism to envelope synthesis, which is energetically less demanding process than the protein synthesis (middle pane). As the S/V decreases, the relative energy expenditure due to envelope synthesis decreases, and a greater portion of the metabolic rate is accounted for by the elevated rate of protein synthesis (right pane), which is a more energetically demanding process. In other words, with reduction of S/V, the cell shifts from a cheap to an energetically expensive process, resulting in an increase of overall energy consumption.

**Figure 6.**
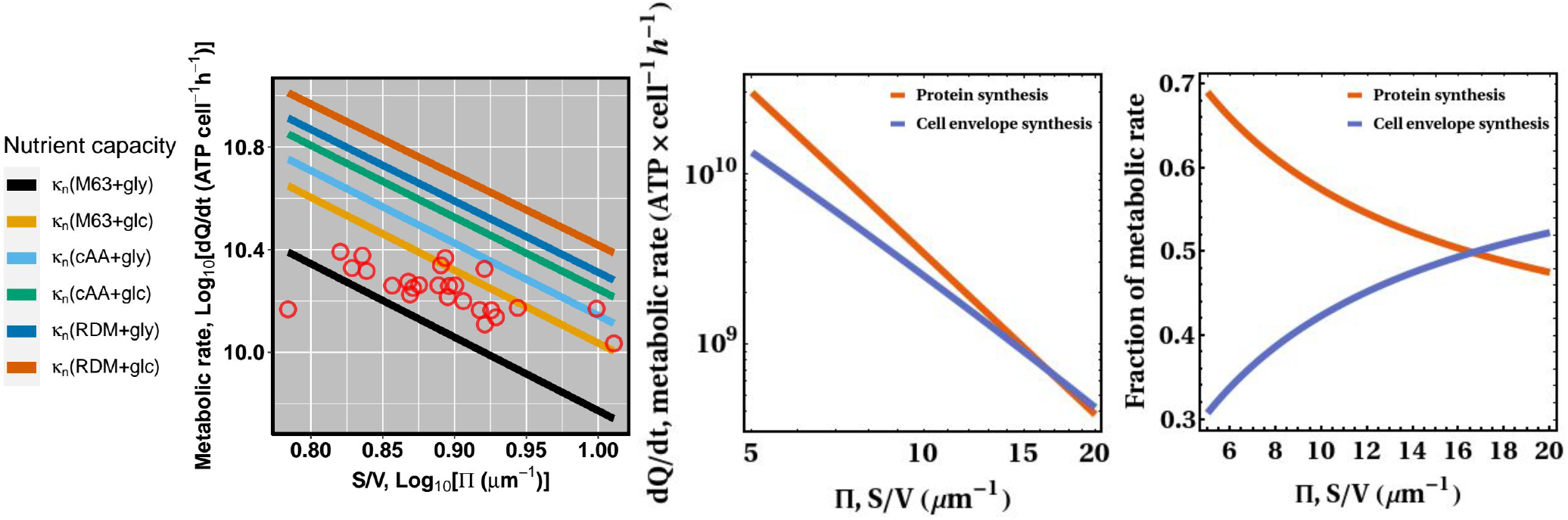
The scaling of the metabolic rate with S/V in the long-term experimental evolution of *E. coli*. Left pane: Comparison of measured metabolic rate and theoretical expectation (Eq 31) for different values of nutritional capacity *κ*_*n*_ inferred earlier. Points are metabolic rate measurements reported in [48]. Middle pane: the energy expenditure of protein and cell envelope synthesis. The contribution of these two processes to metabolic rate was obtained by setting *Q*_*l*_ = *Q*_*dp*_ = *Q*_*dl*_ = 0 (for protein synthesis), and *Q*_*t*_ = *Q*_*aa*_ = *Q*_*dp*_ = *Q*_*dl*_ = 0 (for cell envelope synthesis). Right pane: expected partitioning of the total metabolic rate among processes involved in protein and cell envelope synthesis.

Perhaps the good fit between the theory and measurements is surprising, given that many cellular processes that require energy – such as signaling, motility, synthesis of nucleic acids, and so on – are not explicitly accounted for in our model. Note, however, that we have parameterized Eq 31 using rate constants that were inferred from the changes in protemic composition with the growth rate (Section 3.4), and that in doing so, all of the aspects of the cell that were not explicitly accounted for in the model were “collected” in these three rate constants (namely, *κ*_*n*_, *κ*_*t*_, and *κ*_*l*_).

However, our theory cannot explain the scaling of metabolic rate across diverse prokaryotes, given that the cross-species metabolic rate scales with exponent of 2 with the cell volume [49, 50], which is much larger than the exponent of 1/3 observed in the long-term experimental evolution of E. coli. It is currently unclear what causes this discrepancy.

## 5 Discussion

Motivated by the observation that larger bacteria tend to grow faster, we proposed a theory that explains this relationship in terms of a trade-off between investment into surface features and biosynthetic machinery that builds the cell. By formalizing this verbal statement into a quantitative model, we obtained predictions that we sought to test with published data. Three conclusions are reached. First, the model recapitulates the previous physiological responses (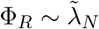, and 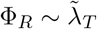), and uncovers new ones (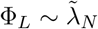, and 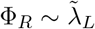). Second, it reveals that the natural variable that governs the scaling of the growth rate with bacterium size is the ratio of the cell surface area to volume and not the cell volume itself. And third, the model correctly predicts negative scaling of ribosomal content with S/V. On the other hand, the purported mild positive scaling of Φ_*L*_ is not unambiguously corroborated by proteomic data.

We attempted to control for factors that might affect the scaling. By cross-validating with an independently published data, we find that our conclusions are not an artifact of the dataset. By performing separate regressions with data on host-associated and free-living species, we excluded the possibility that the overall scaling is caused by small-celled species being dominated by organisms selected for slow growth to avoid excessively damaging their host. By including data on envelope thickness when calculating S/V_cyt_, we excluded the possibility that the envelope becomes thicker with cell size, thus increasing the relative investment into the envelope as cells become larger [51,52]. By comparing growth rates across experimentally evolved studies where fast growth is selected, we controlled for other factors – such as the environment that bacteria inhabit – that confound the scaling of the growth rate. Indeed, the scaling identical to cross-species comparisons holds as well.

Based on the preceding analysis, we propose the following scenario for the origins of the scaling of bacterial size and growth. Bacteria inhabit diverse environments that favor various cell shapes and sizes, due to size-selective predation, swimming efficiency in viscous environments, maximization of nutrient uptake rate, enhancing dispersal, and so on [38, 53]. The shape and size, in turn, set the maximal growth rate that a bacterium can achieve via investment trade-off between biosynthetic and envelope-producing processes. While our analysis was restricted to one particular theory, one might propose alternative explanations. These are now discussed.

### 5.1 Gene-repertoir hypothesis

It was proposed that the larger bacteria have larger genomes and a greater gene repertoire, which allows them to metabolize a more diverse set of nutrients (or use them more efficiently) compared to their smaller counterparts [3]. There are a few problems with this idea. First, it is difficult to test this hypothesis because it does not offer a quantitative prediction on how genome length or gene number scaling should translate to the scaling of the growth rate. For example, should one expect the growth rate to scale with an identical exponent as gene number? If so, data in [3] refutes this hypothesis, given that the growth rate scales with the power of 0.73 and gene number and genome size with the power of 0.35. On the other hand, if the growth rate scales with the square of the gene number, then their explanation seems more plausible. Second, even if the gene-repertoir hypothesis is true, it begs the question as to what causes genome size (or gene number) to scale with the power of 0.35 with cell mass instead of some other value. Third, while this hypothesis might in principle explain the interspecific growth scaling, it cannot explain scaling of growth in experimental evolution as all data points correspond to the same species with the identical gene repertoire.

### 5.2 Internal diffusion constraints hypothesis

An alternative hypothesis is that larger cells grow faster because their cytoplasm is less crowded, thus alleviating constraints from the internal diffusion of macromolecules [42]. According to this hypothesis, growth is proportional to the abundance of metabolic proteins, implying that faster growth can be only achieved by increasing their abundance. However, increased abundance of effectors can slow down intracellular diffusion and thus reduce the encounter rates between cellular components. To offset this problem, an increase in the total mass of these effectors has to be accompanied by a faster increase of volume, such that the density of the cell decreases. On the contrary, volume and mass scale proportionally in our data set (slope of 0.93 ± 0.084 with adjusted *R*^2^ = 0.88 is not significantly different from unity, *p* = 0.412), meaning that larger cells are not less dense. Furthermore, if selection optimizes the density of cytoplasm such that the rates of cellular reactions are maximized [54, 55], it is not clear why decrease in density would not increase the mean time required for two proteins to collide with one another and, thus, decrease the rate of cellular reactions.

### 5.3 A link between SMK law and cross-species growth rate scaling?

Finally, it is widely known that faster-growing cells tend to have larger volumes (colloquially referred to as SMK or Schaechter-Maaloe-Kjeldgaard law), e.g., when growth is altered by rearing bacteria in media of differing nutrient quality [56], and one might argue that the scaling of growth rate across bacteria and across time in lines selected for fast growth is driven by the same underlying mechanism. The correlation between size and growth within species is explained by the threshold initiator model, which assumes that the cell divides once a critical threshold of a particular division protein is reached [35]. According to this model, the cell has to balance between synthesis of division protein and ribosomes. When the cell is reared in nutrient-rich conditions, it allocates a greater proteome mass fraction to ribosomal proteins to support fast growth, and fewer resources are left for the division protein. Therefore, the cell produces division protein more slowly than it grows in size, causing the average cell to be larger [57].

It is possible to co-opt this model in an attempt to explain cross-species correlation between size and growth. Here, the idea is that fast-growing species have a more efficient metabolism than their slow-growing counterparts and thus allocate more resources toward ribosomes, leading to the reduction in the rate of synthesis of the division protein. There are two problems with this explanation. First, it is unclear why selection would make metabolism of some bacteria efficient but not others. Perhaps this might be attributed to differences in environments that they inhabit. For example, bacteria living in other organisms might not be selected for fast growth as a mechanism for preventing harm to the host, whereas selection in free-living species relentlesly promotes faster growth. We show that this is not so, and that the negative correlation between S/V and the growth rate holds both among host-associated bacteria, as well as among the free-living species (Section S1.12). To illustrate this point, consider *Streptococcus pyogenes*, inhabiting throat and dividing in 25 minutes, and *Treponema denticola*, inhabiting mouth and dividing in 20 hours. One is left wondering why selection in two species living in similar environments yields dissimilar metabolic efficiencies. Second, the SMK-based theory does not explain the difference in exponents between cross-species and within-species scaling. That is, the within-species scaling exponent of growth rate with S/V is −1.68 and not −1, as is the case for the cross-species scaling.

Perhaps one could argue that the observed cross-species scaling is caused by the fact that small-S/V species require richer media, so that the perceived higher growth rate is merely the consequence of correlation between nutrient concentration and S/V. However, most of the maximal growth rates in our data were measured in media of similar nutrient quality containing yeast extract and peptone. Furthermore, many of the large-S/V species – like spirochetes and mollicutes – require rich media with bovine serum albumin.

One may also use the SMK-based model to explain correlated changes in cell size with the growth rate in lines experimentally evolved for faster growth. Cells start with fairly inefficient metabolism, and over the course of experimental evolution, their metabolic capabilities are gradually improved. Over time, this causes an increasing re-allocation of the proteome from metabolic proteins and division protein to ribosomal proteins, thus inflating the cell. This hypothesis predicts that if a faster-growing evolved line is forced to divide more slowly such that it has the same growth rate as the ancestor, it would also have the same cell size as the ancestor. However, rearing an evolved population in a chemostat at lower dilution rate still yields larger cells relative to the ancestor [58], implying that increase in cell size cannot be explained only as a by-product of the physiological response to growth rate change.

### 5.4 Constraints on growth imposed by molecular stochasticity?

One can speculate about alternative explanations for the growth scaling pattern that have not been entertained in the literature. Two such scenarios come to mind. First, small-celled bacterial species have a smaller number of proteins in the cell, meaning that the noise in gene expression and partitioning of molecules at cell division time is exacerbated. This stochasticity can disbalance the optimal stoichiometry of biosynthetic pathways and, in turn, cause a reduction in the growth rate. Second, following the reasoning proposed in [59], fast-growing cells may be bigger because this reduces the noise in gene expression of metabolic proteins. The idea is that external ecological factors (such as availability of nutrients) determine the cell’s growth rate independent of the cell size. Faster-growing cells have a greater mass fraction allocated to ribosomal proteins, implying that a smaller mass fraction is allocated to other metabolic proteins. If the cell volume does not change, the faster-growing cell will experience a greater noise in the expression of metabolic proteins, owing to the low-copy number of this sector. Hence, fast growers might have evolved larger sizes to dampen this stochasticity. A problem with both of these hypotheses is that one would intuitively expect that the growth rate scales inversely with the volume of the cell. In contrast, judging from our preceding analysis, surface-to-volume seems to be a stronger predictor.

### 5.5 Relationship to previous studies

The theory developed here conflicts with three previously published conclusions. First, contrary to the assumption that the selection coefficient (i.e., the evolutionary cost) of a particular cellular feature is directly proportional to the fraction of total resources invested in the production of the trait [4, 60], here we show that this is not necessarily true, and that it can also depend on the rates of cellular reactions (Eq S1.2–S1.4). This is because the costs of the trait also have to include the costs of machinery that builds the trait (in this case, the envelope-producing enzyme), and the amount of enzymes produced depends on how fast it operates relative to other proteome components: If the enzyme is slow, then a large fraction of the proteome has to be allocated to it to achieve an appropriate flux. Second, although a recently published analysis [61] concluded that there is no correlation between cell diameter and growth rate across prokaryotes, our analysis reveals that growth rate correlates with S/V, at least when one focuses on bacteria for which both linear dimensions are available.

Third, it was argued that the observed scaling of various bacterial features necessarily imposes the upper limit on how large a bacterium can be [62]. According to this view, the ribosomes have to replicate themselves and all other proteins within the cell doubling time. So, if ribosomes, other proteins, and doubling time scale with different exponents, it is possible for a cell to reach a particular volume at which it does not have enough ribosomes to replicate the entire proteome within the doubling time inferred from a specific power function – an event deemed the “ribosome catastrophe”. However, given that many functions can fit the same data, it is unclear whether fitting pure power functions is adequate. Indeed, our model can explain the scaling of both proteome and growth rate without any fundamental cell size limit. Of course, there might be many reasons why bacteria cannot evolve extreme cell sizes, but inferring this limit from the scaling laws within extant bacterial species may be a questionable approach.

### 5.6 Consequences for the scaling of genomic features with cell size

Our model also offers a causal explanation for the correlation between genomic features associated with translation and growth rate. For example, faster-growing bacteria tend to have a greater number of rRNA operons [63]. According to our theory, bacteria grow slowly because they cannot allocate enough resources to ribosomes, given that other surface-related constraints have to be satisfied. Suppose the high copy number of rRNA genes is caused by the need to meet the high demand for ribosomes. In that case, species with large S/V will have no selective advantage in possessing additional gene copies, meaning that they will be purged by selection to reduce the costs of replicating the added DNA. The same explanation holds true for tRNA genes. Unfortunately, our theory does not yield the precise expectation for the scaling of genomic features because our model does not include genome as a resource component, so this hypothesis cannot be tested rigorously at the moment. Nonetheless, we find a negative log-log scaling between S/V of the species and its rRNA (*p <* 10^−9^, adjusted *R*^2^ = 0.18) and tRNA (*p <* 10^−13^, adjusted *R*^2^ = 0.26) gene number (see Section S1.13 in S1 Appendix for details).

In addition to explaining the scaling of growth rate across bacterial species, the developed theory may also explain an increase in cell size with an increasing growth rate in experimentally evolved lines. Mechanistically, a genotype with an increased cell size will reduce the metabolic burden of synthesizing the cell envelope and, hence, confer a higher growth rate. The cell size in long-term experimental evolution in *E. coli* exhibits a step-like trajectory, where relatively long intervals of cell volume constancy are interupted by short bursts of inflation of cell volume [64]. Interestingly, both relative fitness and cell size experience increase in roughly three bursts in first 1500 generations that appear to coincide with each other [65]. It is possible that the high correlation between size and fitness in early stages of the experiment are caused by the mechanism described above. This does not mean that other factors – such as enhanced nutrient import [66, 67] – do not improve the fitness, but rather that reduction in envelope-related costs might be one of them.

## 6 Conclusion

It is widely known that smaller eukaryotes tend to grow faster than larger ones. However, this trend does not hold in bacteria, where small-celled species grow slower. We propose that small bacteria – compared to their larger counterparts – have to invest a greater fraction of the total resources to the cell envelope owing to their large surface-to-volume ratio, leaving fewer resources for internal biosynthetic processes that build the cell. By representing the cell as being composed of proteins (that convert nutrients into biomass) and cell envelope, we find that cells with large surface-to-volume ratios grow more slowly because they have to invest more resources in the production of the cell envelope and the enzyme machinery that builds this structure, thus leaving fewer resources to ribosomes that replicate the cell. These predictions are corroborated by comparison with growth rate data across more than 200 bacterial species.

The model presented here is not universal, as it cannot account for the scaling of growth across the entire Tree of Life, most notably eukaryotes. We would expect the growth rate to monotonically increase as S/V decreases, but we know that this is not true because eukaryotes with much smaller S/V have growth rates that decrease with size [68]. Even large bacteria, like *Metabacterium polyspora* with a volume of 480 *μm*^3^, have doubling times measured in days [69]. Similarly, although our theory can explain size changes in *E. coli* lines selected for fast growth, it fails to account for size changes in eukaryotes under the same selective pressure. For example, propagation of *Kluyveromyces marxianus* in pH-auxostat leads to increase in S/V, which contradicts our theory [70]. Therefore, the scaling law proposed here breaks at some point, as other constraints become more dominant. Although advances in integrating multiple physical constraints acting at once have been made [71], the theory that mechanistically unifies them and derives expected scaling laws remains to be developed.

## 7 Data availability

All notebooks, data, and code used to derive the theoretical results and plot all the figures is available in Supporting information and at https://github.com/BogiTrick/growth_scaling_envelope.

## 8 Supporting information

**S1 Appendix. Robustness, corrections, and cross-validation**. Contains the analysis of the growth-impeding effects of cell envelope, derivation of physiological scaling laws, corrections for variation in S/V across cell cycle and growth conditions, data collection procedures, the sensitivity of the scaling to variation in protein degradation rates, correction for cell envelope thickness, cross-validation of the scaling trends with independent dataset. The method for obtaining errors of inferred parameters is described.

**S1 File. Notebooks and scripts**. Contains *Mathematica* notebooks for reproduction of entire theoretical derivation, spreadsheets with raw data, and R scripts used for data processing.

## 9 Acknowledgements

We are thankful to Paul Schavemaker, Jon Harrison, Jesse Taylor, and Kerry Geiler-Samerotte for the critical reading of the manuscript, and to Nkrumah Grant for sharing the cell size data in the long-term experimental evolution of *E. coli*.

## Supplementary Information – S1 Appendix

### S1.1 The growth-impeding effects of cell envelope

We are ultimately interested in how the growth rate 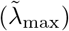 scales with the cell size across bacterial species with different shapes. We start by investigating the cell without degradation processes. In the limit of saturation kinetics, our model yields a closed-form solution for the maximal attainable growth given a particular cell size and shape. For simplicity, let us consider a case when degradation is absent (*d*_*p*_ = 0, *d*_*l*_ = 0), so that the Eq S1.9a simplifies to:

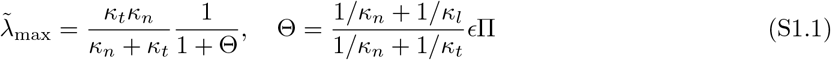

This expression has a simple, intuitive interpretation. The first term corresponds to Monod’s law [1], meaning that the growth rate is a hyperbolic function of the nutrient quality (*κ*_*n*_) and protein synthesis rate (*κ*_*t*_). As more nutrients are supplied (i.e., *κ*_*n*_ increased), the production of building blocks becomes less limiting than the downstream step of incorporating those components into the biomass. In the limit of infinite resource concentration (*κ*_*n*_ → ∞), the rate of growth will equal the rate at which those resources are converted into the proteins by ribosomes 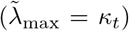. Likewise, in the limit of infinitely fast translation (*κ*_*t*_ → ∞) that instantaneously convert amino acids into biomass, the rate of growth will be identical to the rate at which nutrients are assimilated and supplied to ribosomes 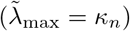.

The second term in Eq S1.1 captures the impediment of the growth rate caused by the production of the cell envelope, which acts as a resource sink. Here, two important insights emerge. First, the impediment term is a monotonically increasing function of Π, meaning that either smaller or elongated cells ought to have low growth rates. For example, two cells with identical cell volumes should have different growth rates if one is round and the other is elongated. Second, the growth rate impediment depends not only on the bioenergetic costs of the envelope – set by *ϵ*Π term but also on the rates of cellular reactions. The term Θ can be intuitively interpreted as the time required to produce a unit of surface area relative to the time needed for the production of a unit of proteome. Because *κ* is the rate of the biochemical step, 1*/κ* is the mean time to completion of that step. If the rate of the envelope-producing enzyme is low such that the time to build an envelope is large, then the cell has to reallocate more of its proteome toward envelope producers.

Therefore, the envelope has two kinds of growth-impeding effects, or costs: (1) structure-related, via allocation of resources to the envelope itself; and (2) machinery-related, via allocation of resources to enzymes that build the envelope. Both of these features divert resources away from processes that replicate the cell and thus impede the growth. The costs are computed by comparing the growth rates with and without a particular feature. This is simply the definition of a selection coefficient in reproduction occurs in the continuous-time [2]. For instance, the total growth impediment caused by envelope is:

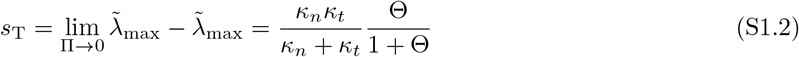

The structure-related cost is obtained by comparing growth rates of the cell lacking both structure and machinery (Π →0) to the one with structure and without machinery (*κ*_*l*_ → ∞). Intuitively, the proteome fraction allocated to envelope producers tends to zero when the envelope synthesis rate is infinitely fast. Likewise, the machinery-related cost is retrieved by contrasting the growth rates of the cell with structure and without machinery (*κ*_*l*_ → ∞), with the one that has both of these components (wild type growth rate). In the first case, the sole cause of the difference in growth rates is the presence of the structure, while in the second case, it is the presence of machinery:

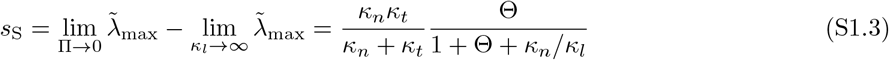

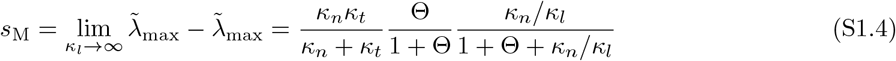

Three interesting consequences emerge from expressions S1.2–S1.4. First, the machinery costs decrease as envelope synthesis rate *κ*_*l*_ increases (because *∂s*_M_*/∂κ*_*l*_ *<* 0). The faster the envelope producer is, the fewer proteins are needed to achieve the same net rate. Second, the structure cost depends not only on the resources needed for the envelope as a structure (*E*Π) but also on nutrient processing and protein synthesis rates. Envelope synthesis competes with protein synthesis for the common building block pool, so even instantaneous production of the envelope diverts resources away from protein synthesis and thus slows down the growth. Third, the total cost increases with Π (because *∂s*_T_*/∂*Π *>* 0), and structure costs eventually come to dominate the total costs while machinery cost approaches zero (as Π → ∞, *s*_S_ → *s*_T_ and *s*_M_ → 0).

The inclusion of protein degradation in the model does not change the previously-reached conclusions significantly. Letting *d*_*l*_ = 0 and *d*_*p*_ *>* 0, in Eq S1.9a yields:

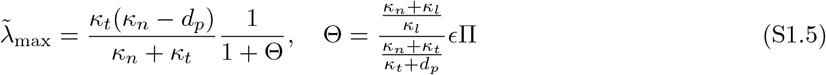

Relative to the no-degradation model, the growth rate is further impaired in two ways (solid and dashed lines in Fig S1.1). First, the maximal attainable growth rate in the absence of envelope is lowered because resources are now partly dissipated via protein degradation (purple lines). Second, the overall growth scaling is reduced because nutrient- and envelope-producing enzymes have to be overexpressed relative to the base scenario without degradation (orange lines) to compensate for lowered flux caused by the continual degradation of machinery. Finally, note that *κ*_*n*_ *> d*_*p*_ for the growth rate to be positive. If this condition is not met, the cell dissipates resources faster than it assimilates them, ultimately leading to complete destruction. Despite these quantitative differences, one still expects the growth rate to eventually approach inverse scaling with Π, as in the no-degradation case.

**Figure S1.1:**
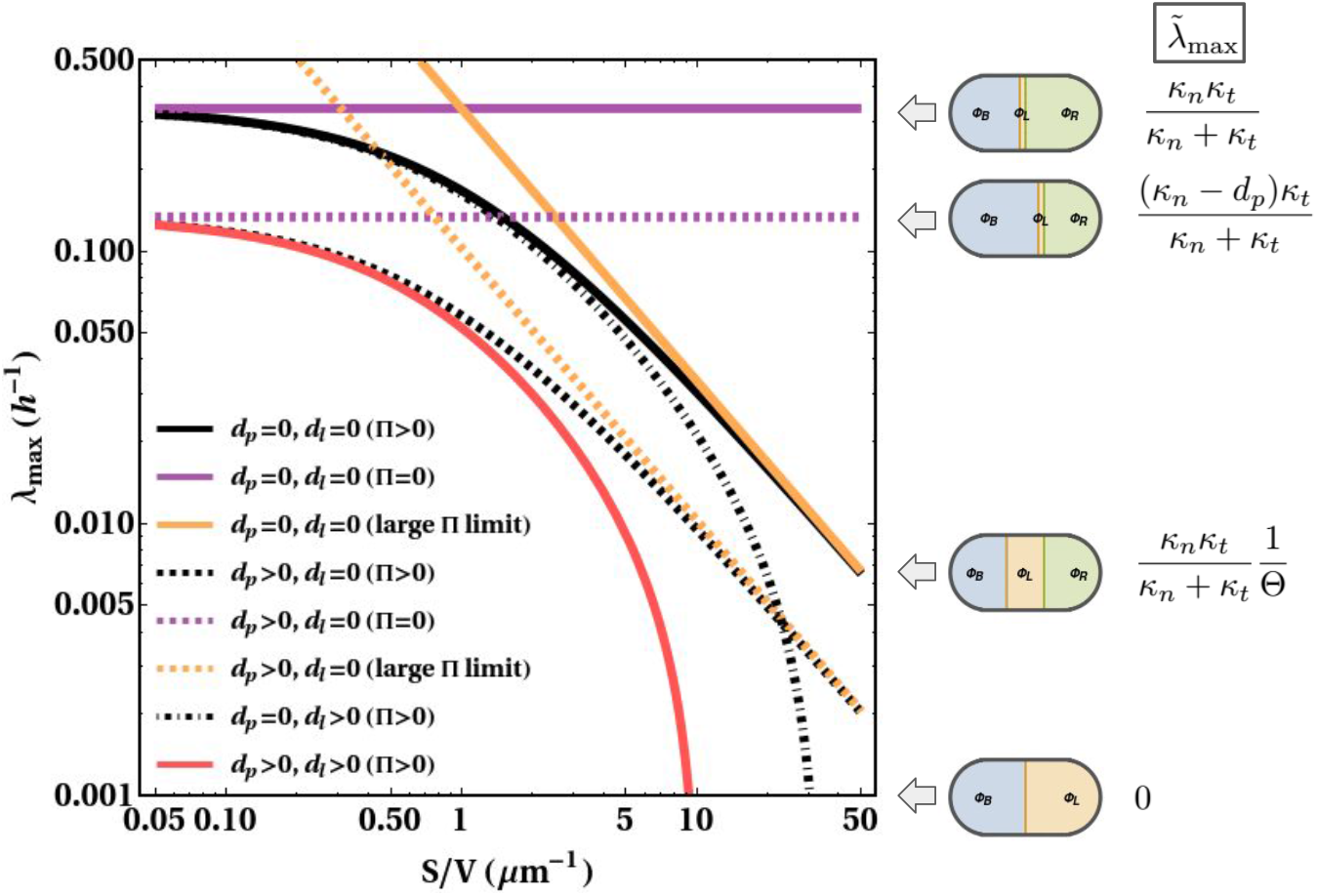
Analytics for the growth rate scaling. The general model is denoted with red dot-dashed curve. Special cases with absent or only one type of degradation are represented by black curves (see inset legend for details). Orange (purple) lines signify the growth rate scaling in the limit of large (small) S/V: Solid lines correspond to the case without protein degradation, while the dashed lines represent the case without the envelope degradation. Side cartoon outlines proteome composition and respective growth rates for solid purple, dashed purple, solid orange, and red solid lines. Parameters: *κ*_*l*_ = *κ*_*t*_ = *ϵ* = 1, *κ*_*n*_ = 0.5, *d*_*l*_ = 0.01, *d*_*p*_ = 0.3.

In the case of only envelope degradation (*d*_*l*_ *>* 0, *d*_*p*_ = 0), the expected scaling changes fundamentally (black dot-dashed line in Fig S1.1):

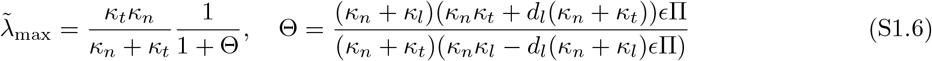

Two consequences are immediately clear. First, the asymptotic growth as cells become large (Π→ 0) is the same as in the no-degradation model. An infinitely large cell (i.e., Π → 0) still has to re-cycle proteins, but it does not have to re-cycle envelope because S/V approaches zero, implying that the asymptotic growth rate is larger in the case of an envelope-than in protein degradation case. Second, for every piece of an envelope that is added, envelope producers have to be overexpressed relative to the no-degradation case to meet an ever-increasing degradation demand. More formally from Eq S1.6, *∂*Θ*/∂*Π *>* 0 (i.e., costs increase with Π) and *∂*^2^Θ*/∂*Π^2^ *>* 0, (i.e., costs increase increasingly fast). This behavior ultimately leads to a critical surface-to-volume ratio Π_crit_ where the entire proteome is devoted to building, and re-cycling envelope and no resources are left for ribosomes. We find this point by setting 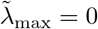 in Eq S1.6 and solving for Π:

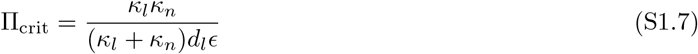

To prove that our interpretation is correct, set *d*_*p*_ = 0 in the expression for optimal fraction of ribosomes allocated to ribosome translation (Eq S1.8a), then substitute Π_crit_, and lastly simplify to zero. The value of Π_crit_ increases with nutrient quality *κ*_*n*_ and envelope synthesis rate *κ*_*l*_ because faster rates mean that fewer resources have to be allocated to the respective proteome component to achieve the same total flux. Moreover, it decreases with envelope degradation rate *d*_*l*_ and costs *E* as this requires heavier processing and synthesis machinery investment, thus shifting the critical point to smaller cells. Finally, allowing for both envelope and lipid degradation (Eq S1.9a) only exacerbates the envelope burden described here (red solid line).

### S1.2 Derivation of physiological scaling laws

In the main text, we derived the optimal partitioning of ribosomes:

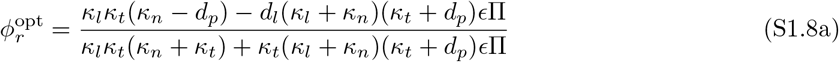

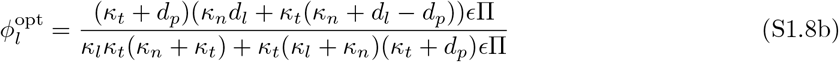

which gives the maximal growth rate:

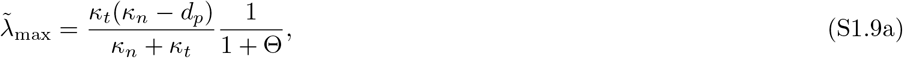

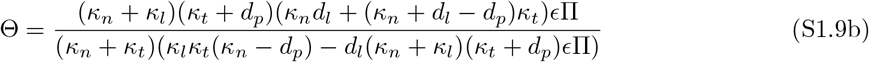

The optimal mass fractions of proteome are then obtained by replacing *ϕ* parameters in the following expressions:

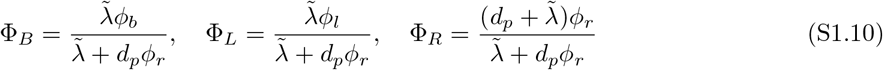

Now we have all the ingredients to derive relationships between the growth rate and proteome mass fractions when cellular processes are perturbed. These regularities are usually referred to in literature as growth laws.

The growth rate and proteome composition depend on the external environment in which the cell is reared. We obtain this dependence by modulating each of the three rate constants *κ*_*n*_, *κ*_*t*_, and *κ*_*l*_ and deriving the response that the cell elicits. We obtain the changes in proteome composition when the nutrient quality of the media is varied in three steps. First, Eq S1.9a is solved for *κ*_*n*_, which is now the function of the growth rate which is modulated by changing the amount of nutrients in the medium 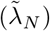. Second, this result is placed in Eq S1.8a and S1.8b to obtain optimal ribosomal allocation. Third and final, these expressions are plugged into formulae for the proteome mass fractions (Eq S1.10). After rearrangement:

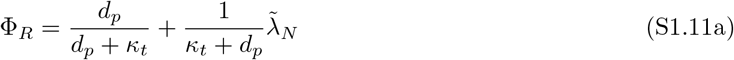

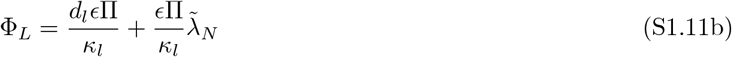

Growth rate can also be altered either by changing the concentration of translation inhibitor 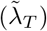. Thus, by solving Eq S1.9a for *κ*_*t*_ and following the same procedure outlined above, we get the optimal partitioning when the growth rate is varied by translation-inhibiting antibiotic:

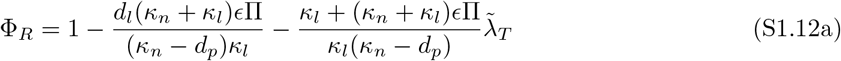

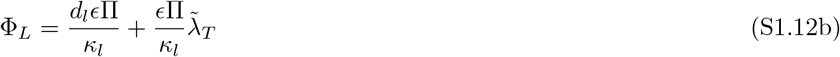

Finally, when the growth is perturbed via changes in concentration of envelope synthesis inhibitor 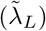, we solve Eq S1.9a for *κ*_*l*_ and obtain:

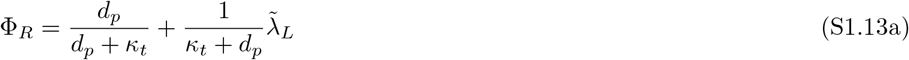

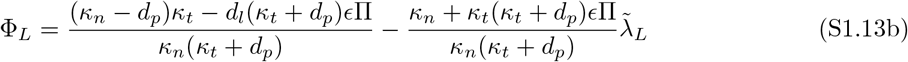

Thus, the mass fractions are a linear function of the growth rate, when the latter is perturbed either by changing the carbon source in the media 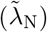, or by altering the concentration of antibiotics (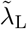 or 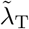). In the limit of no degradation, Eq S1.11a and S1.12a reduce to 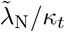 and 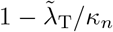, respectively, which are similar to the growth laws first reported in [3] with the exception that we did not account for minimal and maximal possible ribosomal content. One can intuitively interpret the intercept and the slope in Fig S1.2 the following way.

The intercept quantifies the basal expression of each proteomic component required to replenish constantly degraded components. As the exponential growth rate tends to zero, the ribosomal expression is completely abolished because protein synthesis is not needed in a non-dividing cell. However, when degradation is present, ribosomes are necessary even when the cell is not dividing in order for protein synthesis to balance the protein degradation. Hence, the non-zero intercept in Eq S1.11a. The same idea was recently proposed in [4], and presents the alternative to the idea that the excess ribosomes at slow growth represent the reserve that allows cells to quickly tune their growth to cycles of feast and famine [5]. A similar explanation holds for intercept in the Eq S1.11b. Absent lipid degradation, a non-dividing cell abolishes the expression of the envelope-producing enzyme because envelope synthesis only occurs during growth. However, if the envelope is being degraded, then even a non-dividing cell has to allocate some of its resources to an enzyme that will re-synthesize the degraded components. In general, the higher the degradation rates, the higher the intercept because more biosynthetic enzymes are needed to replenish a greater amount of degraded components.

The slope is the rate at which the mass fraction of a particular enzyme changes as the growth rate changes, and it is inversely related to the rate of that enzyme. The smaller the rate constant of an enzyme, the more limiting this step is in the process of growth. As the growth conditions improve, the cell has to allocate more of its resources to alleviate this limitation compared to the case when the enzyme is much faster. Take, for instance, changes in ribosomal content when growth is modulated by the nutrient quality of the media (Eq S1.11a): if *κ*_*t*_ is small, then modulating growth by rearing the cells with better carbon source requires a faster increase in ribosomal mass fraction relative to the case when protein synthesis rate is large. A similar interpretation holds for other growth laws: if the step is slow, then the cell has to allocate resources to it faster to meet the same increase in growth rate. There are generally three types of growth laws, illustrated in Fig S1.2: proteomic changes when nutrients are varied (Eq S1.11a, S1.11b), the translation rate is varied (Eq S1.12b, S1.12a), and the envelope synthesis rate is varied (Eq S1.13b, S1.13a).

**Figure S1.2:**
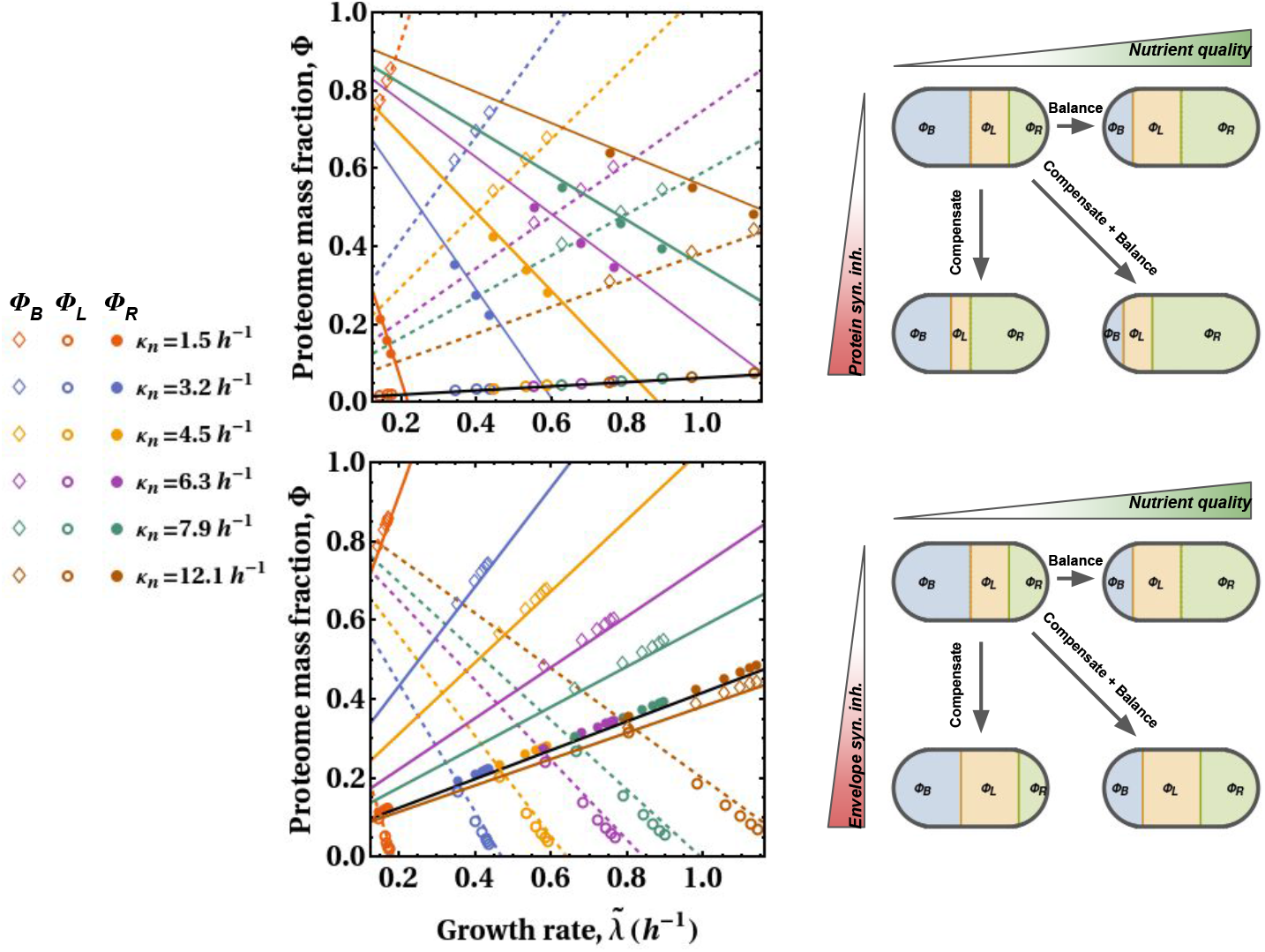
Changes in proteome composition across growth conditions. Panels on the left represent quantitative changes in all three proteome sectors, while the cartoons on the right give graphical interpretation of those changes. Upper row: Proteomic changes when nutrient quality and protein synthesis rate are modulated. Diamond and circle symbols represent numerically-computed optimal proteome mass fractions, and lines denote corresponding analytic solutions. Black line – Φ_*L*_ (Eq S1.12b); Colored solid lines – Φ_*R*_ (Eq S1.12a); Colored dashed lines – Φ_*B*_ (the rest of the proteome mass). Lower row: Proteomic changes when nutrient quality and envelope synthesis rate are modulated. Black line – Φ_*R*_ (Eq S1.13a); Colored dashed lines – Φ_*L*_ (Eq S1.13b); Colored solid lines – Φ_*B*_ (the rest of the proteome). Colors correspond to different *κ*_*n*_ values reported in the legend.

Consider optimal proteomic changes when cells are reared with translation inhibitor under poor nutrient conditions (e.g., blue points and lines in upper panel). Increasing the concentration of protein synthesis inhibitor increases the fraction of ribosomes that are inhibited, so the cell compensates for this inhibition by upregulating the ribosomes (note the negative scaling between Φ_*R*_ and 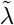). Given the finite proteome, this up-regulation is coupled to the down-regulation of envelope producer and building block producer. Now imagine that the cells were shifted to a richer media or a better carbon source (e.g., brown points and lines in upper panel). Two kinds of changes occur. First, the increased nutrient influx is balanced by an increase in the outflux of building blocks into the biomass by upregulating the ribosomal mass fraction. Second, the same amount of envelope has to be produced in a shorter time. Given that *κ*_*l*_ is fixed, this new goal can only be reached by increasing the Φ_*L*_. In short, rearing cells under better nutrient conditions leads to an increase in the expression of ribosomes and envelope producers.

An identical explanation holds when proteomic changes are induced by growing cells in the presence of envelope synthesis inhibitor (bottom panel). In this case, however, compensation for inhibition is achieved by overexpressing envelope producers (note the negative scaling between Φ_*L*_ and 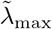), thus causing down-regulation of the two other proteome components. Similar to the previous up-shift experiment, cells reared under better nutrient conditions balance the increased nutrient influx by increasing the outflux of building blocks into the proteins (via up-regulation of ribosomes), and into the envelope (via up-regulation of envelope producer). Finally, note that there is a good agreement between numerically computed optimal proteomic mass fractions and analytical solutions under saturation kinetics (marker symbols and lines in Fig S1.2, respectively).

### S1.3 Formulas for surface area and volume of bacteria

In the main text, we use formula in Table S1.1 when calclating surface area and volume of the cell:

**Table S1.1:** Expressions used to compute surface area and volume in the main text from linear dimensions. D is diameter of the cell, and L is the pole-to-pole length of the cell.

|  | Surface area | Volume |
| --- | --- | --- |
| Capsule | $\pi DL$ | $\pi(D/2)^2(L - D/3)$ |
| Sphere | $\pi D^2$ | $\pi D^3/6$ |

To calculate surface area and volume of helical cells, we first calculate the total length of the helical cell, and then plug that length in the formula for capsule surface and volume. The actual lengh of the cell is 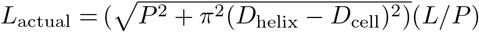, where *D*_helix_ and *D*_cell_ are diameter of helix and cell, respectively, while *P* is the pitch of the helix (i.e., the length of one turn of the helix), and *L* is the length of the helix without it being stretched out (see Eq 4 and 5 in [6]). The square root expression is the length of one turn of the helix, once the helix is fully stretched, and the *L/P* represents the number of turns that a helix has.

### S1.4 Justification for assuming fixed S/V across the cell cycle

The shape of bacterial cell has to change during the cell cycle as the bacterium grows. The variation in S:V across the cell cycle is much smaller than across the bacterial species. Most bacteria grow by elongating along the long axis and undergoing septation at an appropriate length. There are some examples of longitudinal bisection (e.g., *Spiroplasma poulsonii* [7] and *Thiosymbion* [8]) but these are peculiar rarities that do not represent the bulk of our data. This means that one can calculate the surface-to-volume ratio at cell division (Π_div_) and subtract this from the surface-to-volume ratio at cell birth (Π_birth_), yielding the difference in S/V, ΔΠ. This gives one an idea of how much S/V varies across the cell cycle (Fig S1.3).

**Figure S1.3:**
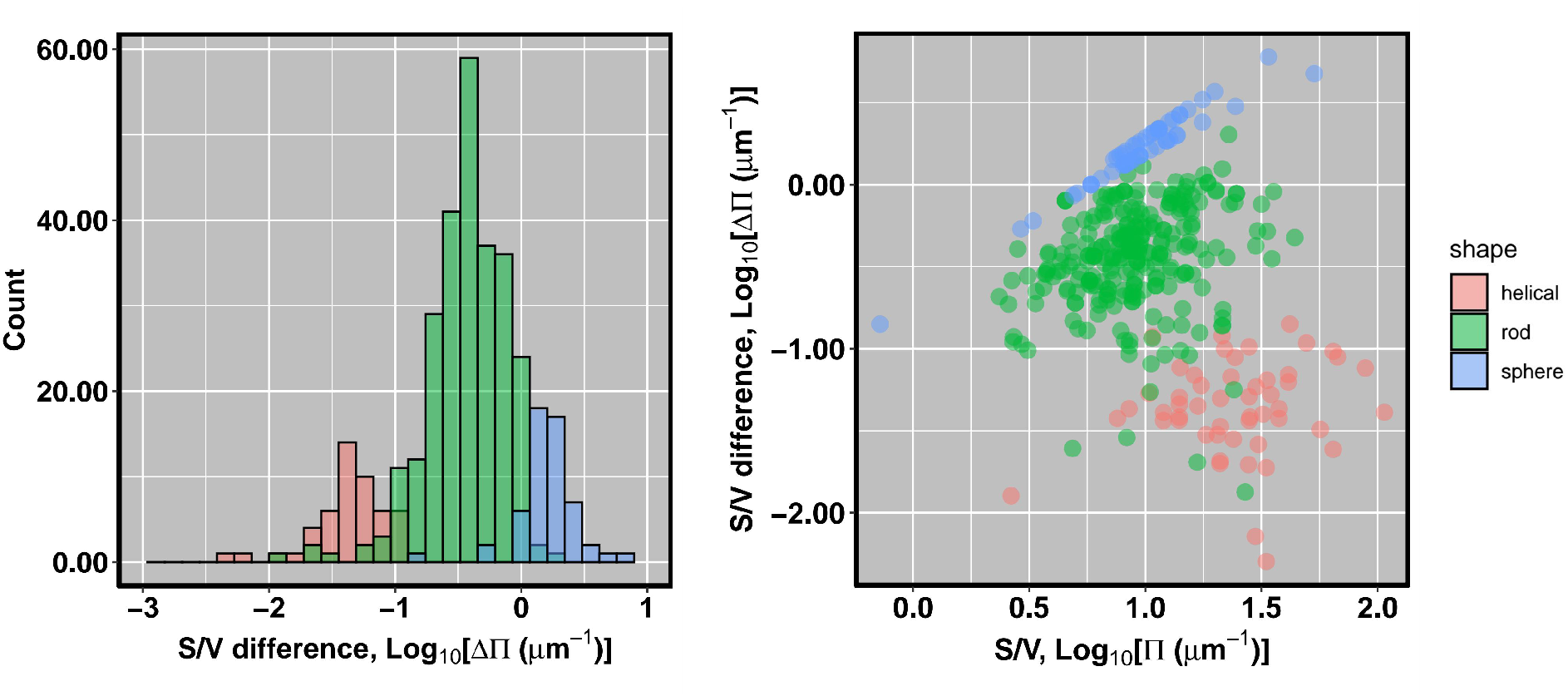
Variation in S/V across the cell cycle. Left pane represents the distribution of ΔΠ, while the right one depicts ΔΠ for each species in our dataset.

We see that ΔΠ is at least an order of magnitude smaller than Π across species with different. Therefore, although S/V varies across the cell cycle, this variation is much smaller than variation in S/V across bacterial species, and one can assume it fixed for a given species.

Another potential limitation of our model is that it does not include the synthesis of septum, which itself is a costly feature. However, the cost of producing septum scales with the square of diameter of the cell (*V* ^2*/*3^), while the total resource budget scales with the volume of the cell (*V* ^1^), implying that the relative cost (i.e., growth rate-penalty) of septum scales with *V* ^−1*/*3^. Thus, septum confers greater growth rate-penalty to small-celled organisms than to larger ones, and only makes the scaling more pronounced. This only strengthen, not weaken, our conclusion.

### S1.5 Correction for changes in S/V across growth conditions

The derived growth laws assume that Π of a given species is constant and not a subject to change. However, linear dimensions of the cell change across growth conditions. For example, cells tend to be larger and have smaller Π when reared in nutrient-rich media [9], and the cell size can change in non-intuitive ways when the culture is treated with various antibiotics [10–12]. Given that we are inferring capacity parameters (*κ*_*n*_, *κ*_*t*_, and *κ*_*l*_) from the slopes of the growth laws, it is possible that variable Π confounds are estimates. To account for this growth-dependancy, we establish empirical relationships between Π and 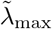 from published studies, and then use substitute it in the previously-derived growth laws.

The capacity parameters are inferred from three different equations. Parameters *κ*_*l*_ and *κ*_*n*_ are estimated from Eq 26 and 27 in the main text, and these depend on Π, while *κ*_*t*_ is inferred from Eq 25 and is independent of Π. Cells reared in rich media have smaller Π, and this pattern holds across three experimental studies [9, 10, 12]:

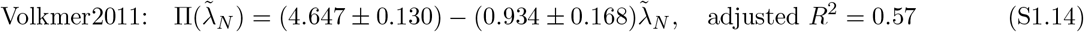

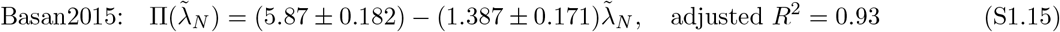

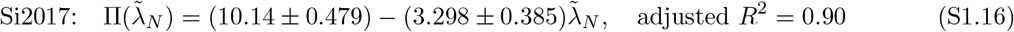

Substituting Π in Eq 28 of the main text for Eq S1.14–S1.16, one obtains the following expressions for Φ_*L*_ as the function of nutrient-modulated growth rate 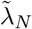 :

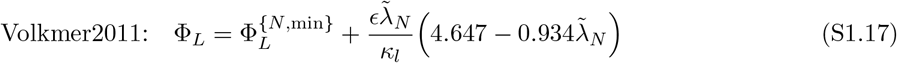

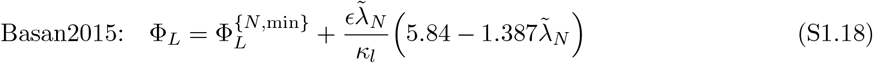

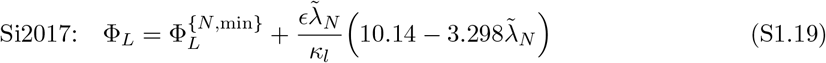

where we parameterize *d*_*l*_ and *ϵ*, and infer *κ*_*l*_ by non-linear least squares analysis using R package ‘nls’. The value of *κ*_*l*_ are similar regardless of the source of 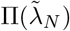 (Table S1.2), and the overall fit to data seems almost identical (Figure S1.4). In the main text, we use *κ*_*l*_ inferred from Volkmer’s study.

**Table S1.2:** Estimates and standard errors of the envelope-synthesis capacity, *κ*_*l*_, and minimal envelope-producer mass fraction, 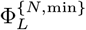, when growth is perturbed by manipulation of nutrient quality is performed with different carbon sources. Source of 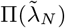 denoted in the first column.

| Source | $\kappa_l$ (estimate) | $\kappa_l$ (std. error) | $\Phi_L^{\{N,\min\}}$ (estimate) | $\Phi_L^{\{N,\min\}}$ (std. err) |
| --- | --- | --- | --- | --- |
| Volkmer2011 | 101.2 | 18.76 | 0.016 | $6 \times 10^{-4}$ |
| Basan2015 | 113.2 | 20.5 | 0.016 | $5 \times 10^{-4}$ |
| Si2017 | 141 | 25.41 | 0.017 | $5 \times 10^{-4}$ |

Second, it is possible that Π changes when growth is perturbed with translation-inhibiting antibiotic. To assess this, we collected data from three studies, and found that surface-to-volume is mainly independent of the growth rate, with only two out of 11 examined conditions resulting in correlation between Π and translation-perturbed growth rate. Thus, one can assume that Π is fixed when inferring *κ*_*n*_, which is the approach we take in the main text.

**Figure S1.4:**
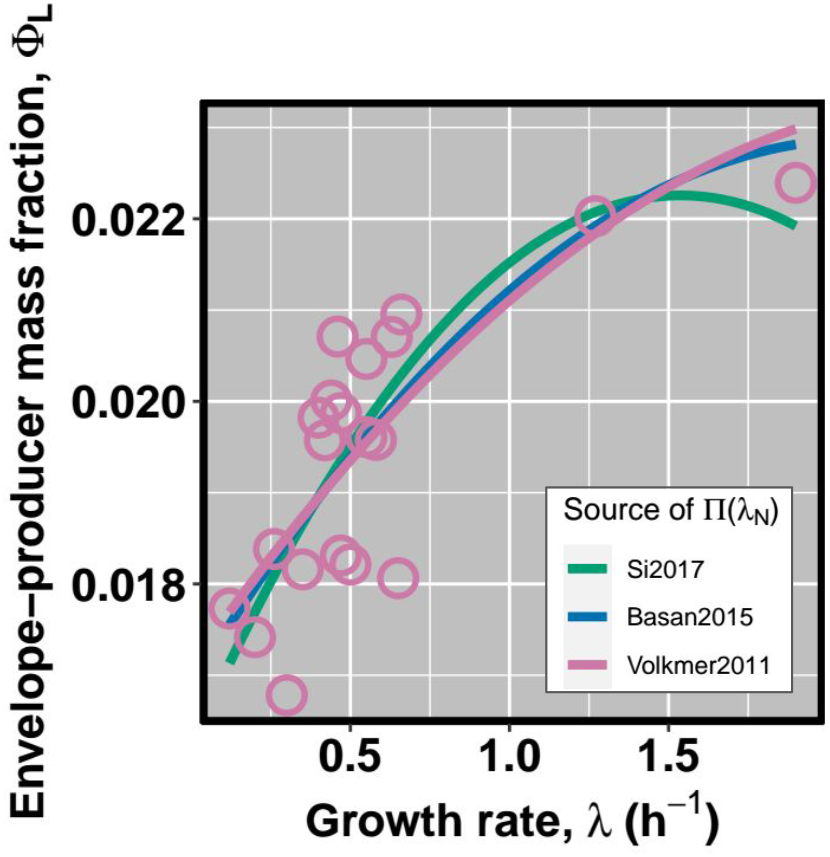
Scaling of the envelope-producer mass fraction Φ_*L*_ as the function of nutrient-modulated steady state growth rate 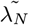. In the main text, we use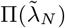 inferred from Volkmer2011 data, but note that using alternative datasets Basan2015 or Si2017 yields almost identical scaling. Pink circles denote data from [13].

However, if one wishes to account for these differences, this can be done readily by substituting Π in Eq 29 with the following expressions:

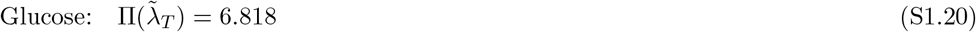

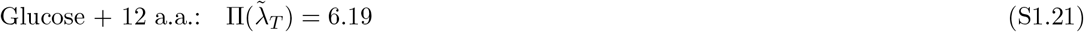

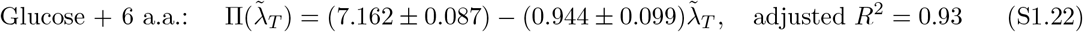

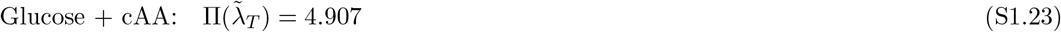

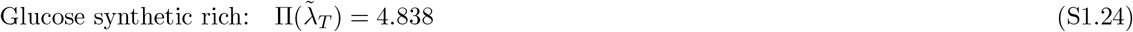

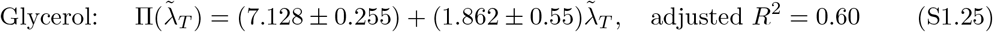

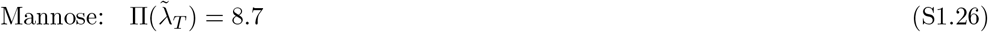

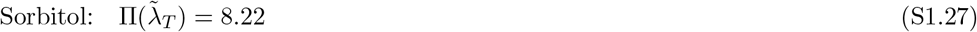

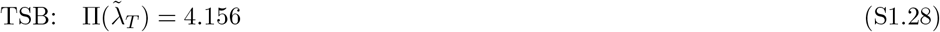

and then inferring *κ*_*n*_ by substituting Eq S1.20–S1.28 for Π in Eq 27, and then performing non-linear least squares analysis. The estimates of *κ*_*n*_ are given in Table S1.3. Parameter *κ*_*l*_ was inferred from Eq S1.19, as *κ*_*n*_ estimates also come from this dataset.

**Table S1.3:** Estimates and standard errors of the nutritional capacity, *κ*_*n*_, and maximal ribosomal mass fraction, 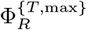, when translation-inhibition is performed with different carbon sources. Data from [12].

| | $\kappa_n$ (estimate) | $\kappa_n$ (std. error) | $\Phi_R^{\{T, \max\}}$ (estimate) | $\Phi_R^{\{T, \max\}}$ (std. err) |
| --- | --- | --- | --- | --- |
| Glucose | 7.956 | 0.769 | 0.351 | 0.01 |
| Glucose + 12 a.a. | 16.928 | 3.64 | 0.342 | 0.02 |
| Glucose + 6 a.a. | 17.469 | 2.327 | 0.274 | 0.01 |
| Glucose + cAA | 29.98 | 7.99 | 0.243 | 0.012 |
| Glucose synthetic rich | 12.993 | 1.844 | 0.424 | 0.019 |
| Glycerol | 10.829 | 3.26 | 0.262 | 0.025 |
| Mannose | 27.064 | 10.962 | 0.102 | 0.008 |
| Sorbitol | 51.83 | 29.833 | 0.118 | 0.01 |
| TSB | 17.176 | 2.394 | 0.355 | 0.017 |

**Figure S1.5:**
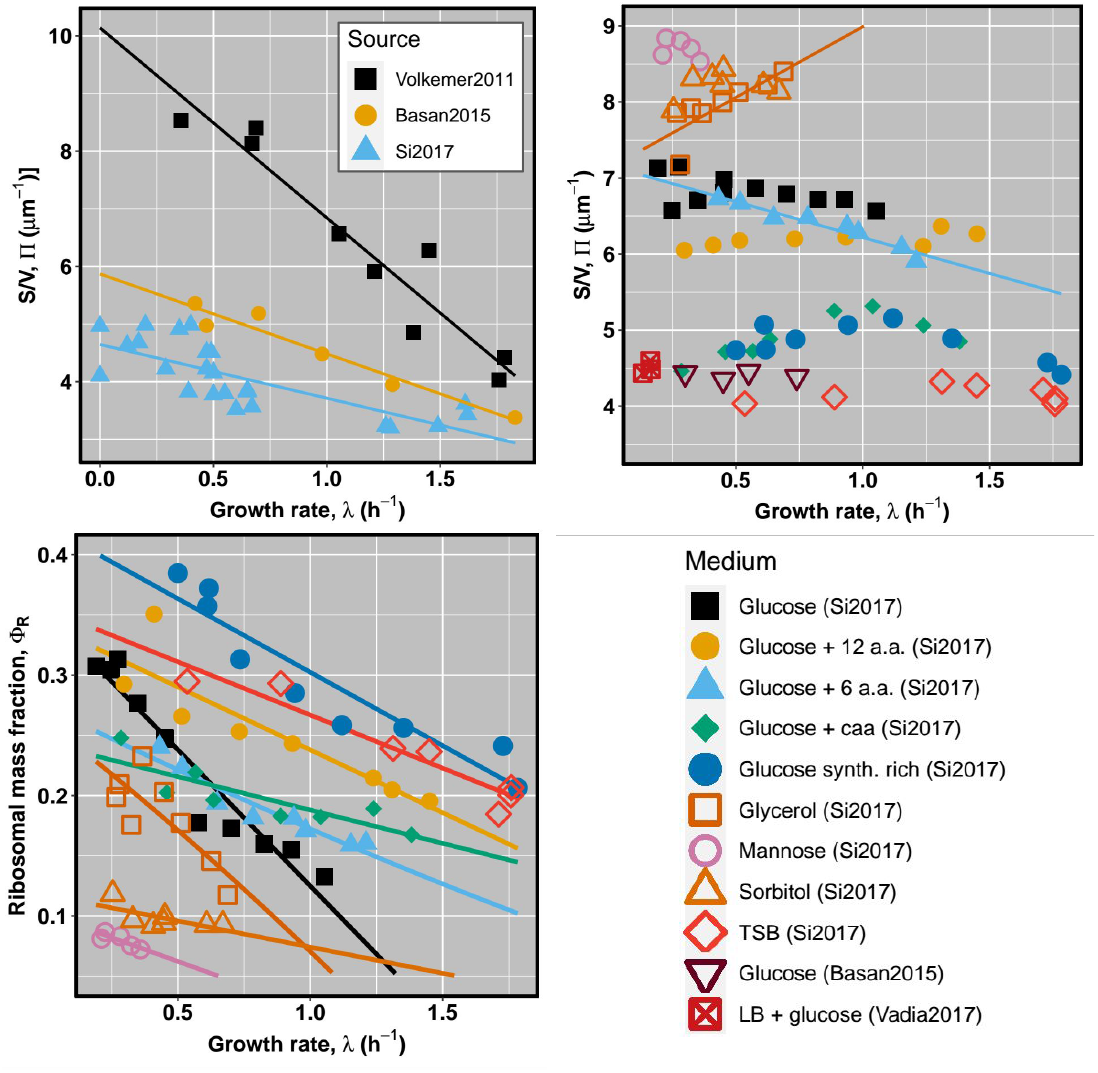
Changes in S/V with growth conditions. Upper row depicts the scaling of Π with nutrient-perturbed (left) and translation-perturbed (right) growth rate. Lower row: Relationship between ribosomal mass fraction and translation-perturbed growth rate. Legend denotes the nutrient used in translation-inhibition experiments. Solid lines signify OLS regressions, as described in the text above.

Parameterizing analytical solution for the maximal growth rate (Eq 22a in the main text) with parameters estimated by assuming growth-dependent S/V yields qualitativey similar scaling of growth and proteomic composition (Fig S1.6). However, because the slopes in Si2017 data are shallower than in Scott2010 data used in the main text, one necessarily obtaines higher *κ*_*n*_ values. This, in turn, implies that the scaling function will be shifted “upwards” toward higher growth rates, and higher Φ_*R*_ and Φ_*L*_.

**Figure S1.6:**
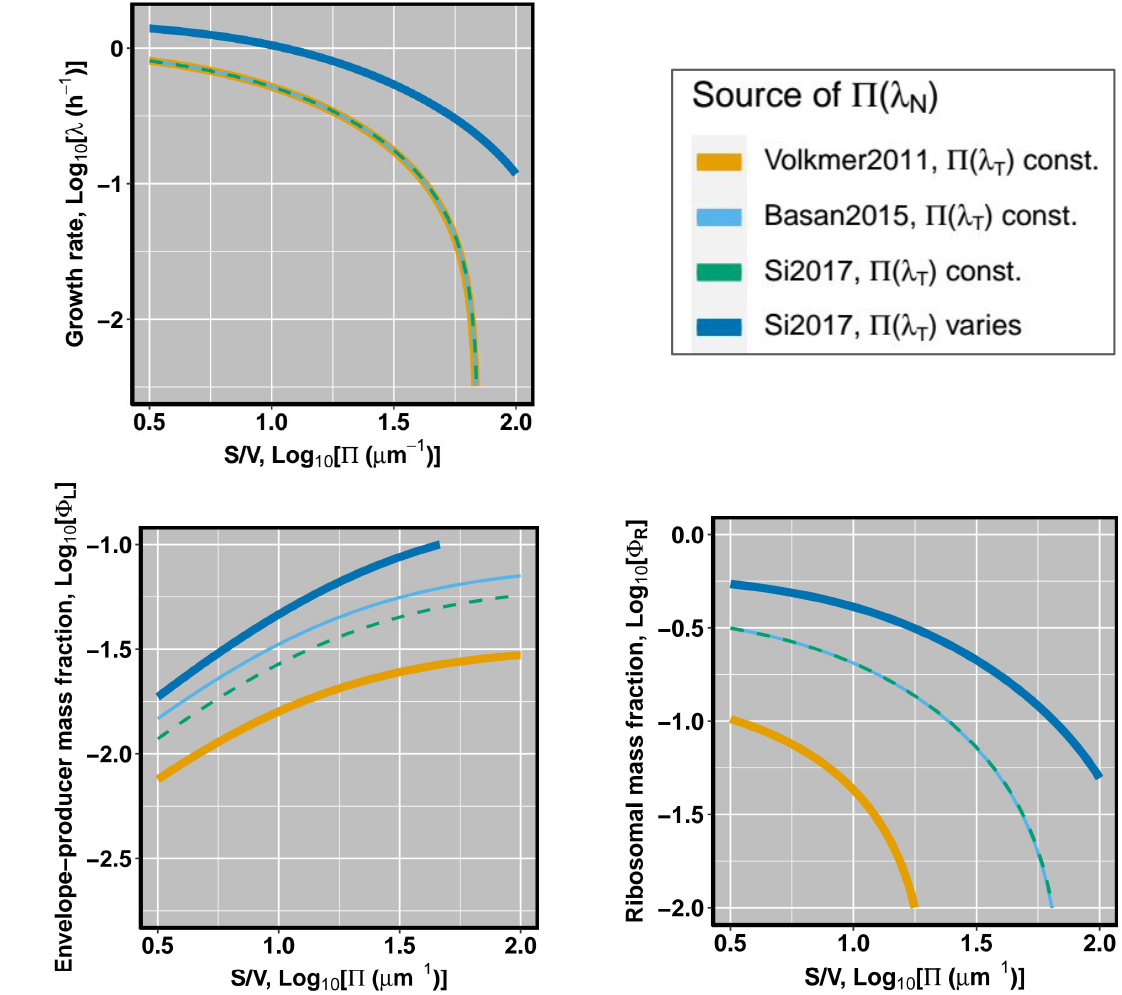
The effects of accounting for dependence of Π on growth conditions using different datasets on scaling patterns. Each line denotes *κ*_*n*_ and *κ*_*l*_ being inferred from a dataset in the legend in the upper right panel. Parameters *κ*_*l*_ and *κ*_*n*_ are inferred from 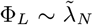 and 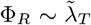, respecitvely, and we account for changes in Π across nutrient- or chloramphenicol-perturbed growth rate using data on cellular dimensions from [9, 10, 12]. Thick blue line denotes scaling using the mean of *κ*_*n*_ values across conditions in Table S1.3.

### S1.6 Data collection and normalization procedures

Growth rate and cell size data were obtained from the literature. Because most of the species were reared in nutrient-rich medium, often containing yeast extract, under optimal temperature, pH, and salinity, we assumed that the reported values correspond to the maximal growth that a species can attain. Given that different species are grown in disparate temperatures, we normalized growth rates to 20°C using Q_10_-correction with Q_10_ coefficient of 2.5, to exclude the potential confounding effect of temperature on growth scaling. The linear dimensions of the cell were also obtained from the literature, and were used to calculate cell volume and surface. Each cell is classified into one of the three categories based on its shape: spheres, rods, and helices.

Because we are interested in whether variation in growth rate can be solely explained in terms of variation in cell size and shape, we work with chemoorganoheterotrophic species. This ensures that variation in growth is not caused by differences in the type of metabolism that species have. However, even chemoorganoheterotrophs may generate energy in a variety of ways. One of the biggest differences is between oxidative phosphorylation and substrate phosphorylation (i.e., fermentation), the latter having lower energetic content. Hence, one can argue that obligate fermenters ought to grow slower owing to a slower rate of energy extraction from nutrients. In our model, this would translate to an organism having a lower *κ*_*n*_. If the growth rate of *E. coli* under anaerobic conditions is 63% of the growth rate under aerobic ones [14], one can assume that *κ*_*n*_ of anaerobes is also 63% of *κ*_*n*_ of aerobes. Formally, let *f* be the growth rate as the function of nutritional capacity *κ*_*n*_ (Eq S1.9a). Then the growth rate of an anaerobe 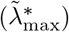 is going to be a function of the anaerobic nutritional capacity 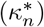. Let anaerobic *κ*_*n*_ be *α* = 0.63 of aerobic *κ*_*n*_:

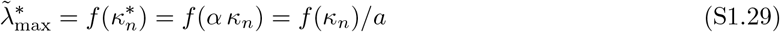

where *a* is conversion factor that depends on the model parameters and is given by:

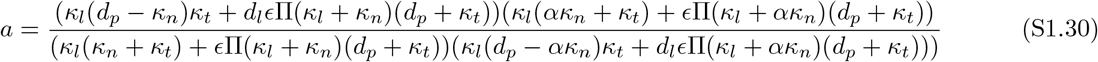

Therefore, to compare aerobes and anaerobes on the same footing, the growth rate of anaerobes obtained from the literature should be multiplied by the factor of *a*; The term *a* can be biologically interpreted as the fold-increase in growth rate that anaerobes would experience if they were to switch to oxidative phosphorylation as the means of generating energy. More precisely, *a* is obtained by taking the ratio of aerobic (with nutritional capacity being *κ*_*n*_) to anaerobic growth rate (with nutritional capacity being 0.63*κ*_*n*_). Each organism was determined whether it is an obligate fermenter by investigating its citrate cycle in KEGG PATHWAY database. A bacterium is deemed a fermenter if it does not have a total of 12 steps listed in the database, starting and ending with oxaloacetate. For species without biochemical annotation, we extracted information on their energy metabolism from the literature. Lastly, when we could not find data on the type of energy-generation pathways, we simply assumed that those species respire.

### S1.7 Proteomic data analysis

We use proteomic data to 1) infer the rate constants via physiological scaling laws, and 2) test the predictions on the scaling of the proteomic composition with the surface-to-volume ratio of the cell.

As outlined in the section S1.2, one needs to co-measure the growth rate across different conditions and the proteome mass fraction allocated to three sectors to infer capacities from the slope of regressions (Eq S1.11a–S1.13b). To this end, we use two types of studies: Those that directly quantified absolute abundances of each protein [13, 26–30], and those that indirectly measured ribosomal mass fraction from the total RNA-to-total protein ratio [3, 12, 31–33]. We use [13] as a primary source because it has the highest coverage of the *E. coli* proteome (in excess of 90% of the proteome is detected). To estimate mass fractions belonging to nutrient-processing, lipid-producing, or ribosomal protein, we classified the total mass (in units fg/cell) of each reported protein in one of the three possible groups based on its designation in KEGG BRITE database [34]. If a protein’s assigned function in the BRITE database had keywords “fatty acid biosynthesis”, “lipopolysaccharide biosynthesis”, “peptidoglycan biosynthesis”, we classified such protein as contributing to Φ_*L*_: If protein’s designation contained the keyword “ribosomal protein”, we classified it as contributing to Φ_*R*_. All other proteins were grouped in the Φ_*B*_ fraction. After binning, we calculate each mass fraction by dividing the total mass of proteins per cell in that class by the total mass of proteome per cell. Ribosomal mass fractions from indirect sources are obtained by multiplying the total RNA/total protein ratio with a conversion constant as reported in [33]. Some studies reported relative protein abundances, and we converted these into relative mass fractions by multiplying each entry with the molecular mass of the given protein.

To test the prediction on scaling of proteome composition with S/V, we collected quantified proteomes of bacterial species with S/V ranging from less than 5 *μm*^−1^ in *Lactococcus lactis* to more than 50 *μm*^−1^ in *Spiroplasma poulsonii*. In addition to proteomic studies, we also collected data for ribosome abundance and converted it to proteome mass fractions in the following way. First, the total mass of ribosomes was calculated as:

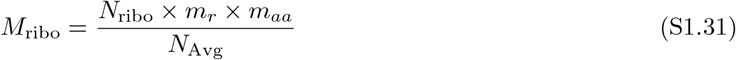

where *m*_*aa*_ = 110 g/mol is the molar mass of an average amino acid, *N*_Avg_ = 6.022 × 10^23^ molecules/mol, *m*_*r*_ = 7336 amino acids is the length of the protein component of a single ribosome (assumed to be constant across bacteria) and *N*_ribo_ is the ribosome abundance per cell. To calculate the mass of the total proteome, we first use the empirical scaling law (Eq S1.32a from prokaryote data in Fig 7.1 reported in Chapter 7 of [35]) that allows calculation of dry cell weight (in grams) from the cell volume *V* (in units of *μm*^3^). Then, knowing that the proteome of most bacterial cells accounts for 54% of the total dry weight (Fig 7.3 reported in Chapter 7 of [35]), the ribosomal mass fraction is calculated in three steps from cell volume:

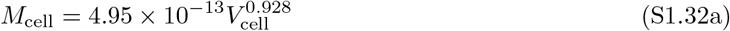

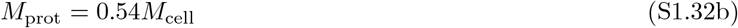

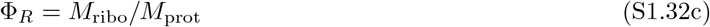

**Table S1.4:** List of sources of cross-species proteomic data.

| Species | Source |
| --- | --- |
| *Bacillus anthracis* | [15] |
| *Bacillus subtilis* | [15–17] |
| *Bacteroides thetaiotamicron* | [15] |
| *Bartonella henselae* | [15] |
| *Borrelia burgdorferi* | [18] |
| *Campylobacter jejuni* | [15, 19] |
| *Clostridium saccharolyticum* | [16] |
| *Dictyoglomus thermophilum* | [16] |
| *Escherichia coli* | [15, 17, 19] |
| *Granulicella tundricola* | [16] |
| *Helicobacter pylori* | [15] |
| *Lactococcus lactis* | [15] |
| *Legionella pneumophila* | [15, 17] |
| *Leptospira interrogans* | [15, 17] |
| *Listeria monocytogenes* | [15] |
| *Mesoplasma florum* | [20] |
| *Mycobacterium smegmatis* | [21] |
| *Mycobacterium tuberculosis* | [15, 17, 21] |
| *Mycoplasma pneumoniae* | [17] |
| Myojin amorphous bacteria | [21] |
| Myojin parakaryote | [22] |
| *Neisseria meningitidis* | [15] |
| *Odoribacter splanchnicus* | [16] |
| *Pelagibacter ubique* | [19] |
| *Persephonella marina* | [16] |
| *Polynucleobacter asymbioticus* | [23] |
| *Pseudomonas aeruginosa* | [15] |
| *Rickettsia prowazekii* | [17, 19] |
| *Salmonella typhimurium* | [15] |
| *Shewanella oneidensis* | [15] |
| *Sphingomonas* sp. RB2256 | [19] |
| *Sphingopyxis alaskensis* | [17] |
| *Spiroplasma melliferum* | [17, 19] |
| *Spiroplasma poulsonii* | [24] |
| *Staphylococcus aureus* | [15, 17] |
| *Streptococcus pyogenes* | [15] |
| *Thermotoga maritima* | [16] |
| *Treponema pallidum* | [25] |
| *Veillonella parvula* | [16] |
| WWE30OP11-OD1 | [19] |
| *Yersinia pestis* | [15] |

For cross-species data, we used proteomes from PaxDB [15], a recent compendium of proteomes across 100 species [16], a collection of ribosomal abundances from [17] and [19], and a number of additional quantitative proteomic studies not reported in above-mentioned databases: *Borrelia burgdorferi* [18], *Treponema pallidum* [25], *Polynucleobacter asymbioticus* [23], *Mesoplasma florum* [20], and *Spiroplasma poulsonii* [24]. In total, we have quantified Φ_*R*_ and Φ_*L*_ for 40 and 30 bacterial species, respectively.

### S1.8 Sensitivity of the growth rate scaling to variation in *d*_*p*_

Because protein degradation rates can vary depending on the protein class in question, we examined how the growth rate scales with Π under various estimates of *d*_*p*_ obtained from Table 2 in [36]. There is almost no variation in the scaling pattern. Degradation rates were calculated by dividing Log(2) with the half-life of a particular protein class S1.7. The yellow line denotes the degradation rates weighted by the corresponding fraction of total protein. All values standardized to 20°C using Q_10_-correction reported in the main text.

**Figure S1.7:**
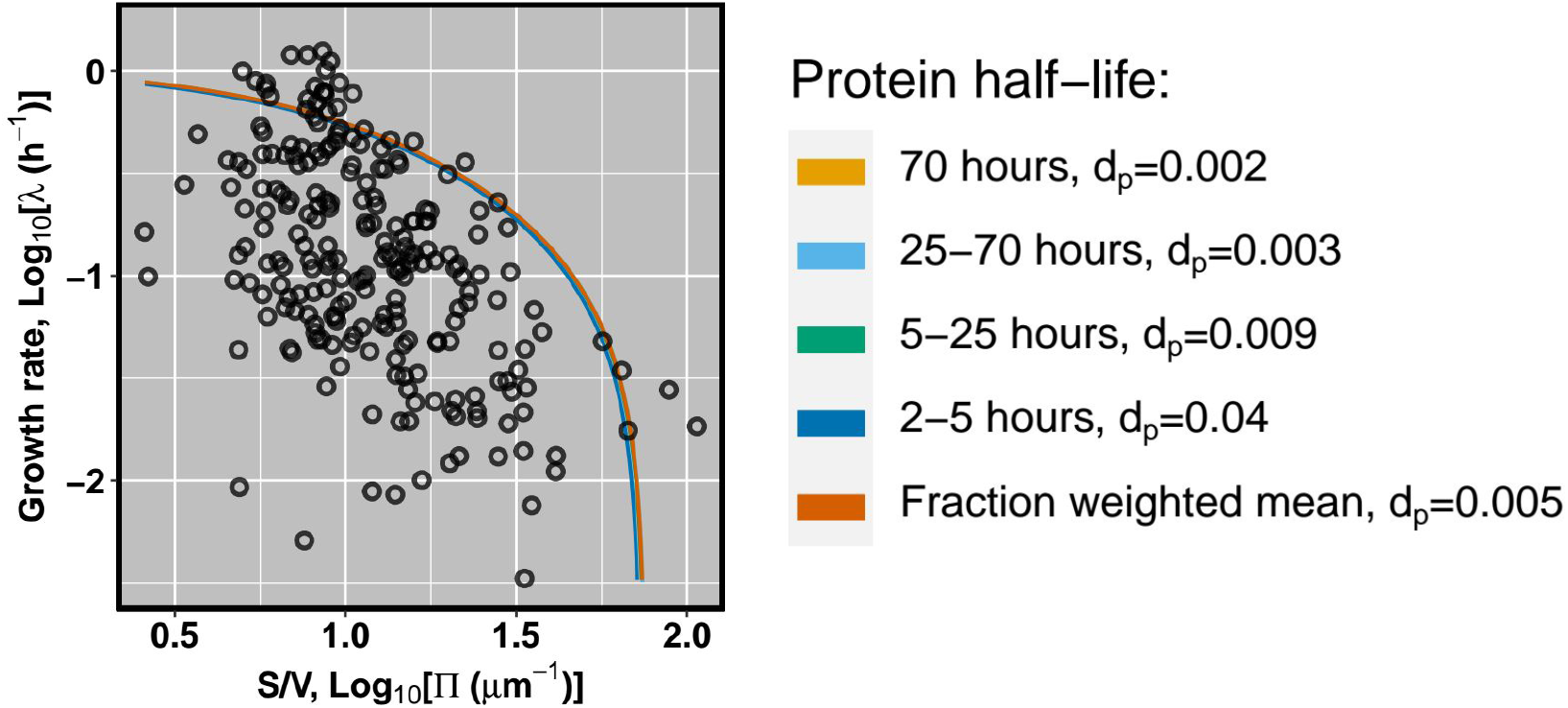
The growth rate scaling is robust to variation in protein degradation rates. All other rate constants as reported in the main text, and *κ*_*n*_ was taken for the medium with casamino acids and glucose.

### S1.9 Error estimates of inferred model parameters

The standard error of the rate constants inferred from the correlation between growth rate and proteomic mass fractions was calculated using the propagation of errors [37]. If *Z* is the function of *k* independent parameters (i.e., *Z* = *f* (*z*_1_, *z*_2_, …, *z*_*k*_)), then variance of *Z* can be expressed in terms of variances of *k* independent parameters *z* as:

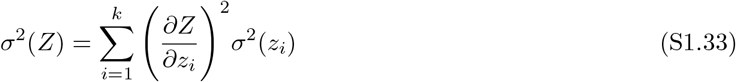

Let 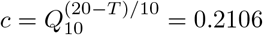 be the temperature correction for biological rates, with *Q*_10_ = 2.5 and *T* = 37°C. One can compute the variances of the inferred parameters from their respective formulae (Table S1.5), and the standard error of the mean is then 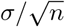, where *n* is the number of data points included in regression. Standard error of *κ*_*l*_ is reported directly from non-linear least squares used to obtain the estimate.

**Table S1.5:** Estimation of parameter variance. Columns two and three contain formulae used to compute the parameter and its variance from the slope *b* of the regression denoted in the last column. Growth rate is modulated either by alteration of carbon source (*λ*_*N*_), or the concentration of translation inhibitor (*λ*_*T*_).

| Parameter | Parameter estimate | Parameter variance | Regression |
| --- | --- | --- | --- |
| $\kappa_t$ | $c(1/b - d_p)$ | $c^2\sigma^2(b)/b^4$ | $\Phi_R \sim \lambda_N$ |
| $\kappa_n$ | $c \frac{\kappa_l(d_p b - \epsilon\Pi - 1)}{\kappa_l b + \epsilon\Pi}$ | $\frac{c^2\kappa_l^2(\kappa_l + \epsilon\Pi(d_p + \kappa_l))^2}{(b\kappa_l + \epsilon\Pi)^4}\sigma^2(b) + \frac{c^2\epsilon^2\Pi^2(1 - bd_p + \epsilon\Pi)^2}{(b\kappa_l + \epsilon\Pi)^4}\sigma^2(\kappa_l)$ | $\Phi_R \sim \lambda_T$ |

### S1.10 Correction for variation in cell envelope thickness

When calculating Π for each species in the dataset, all cell envelopes were assumed to be 30 nm-thick which is roughly the thickness of an *E. coli* envelope. However, the thickness might vary across species for two reasons. Firstly, some species might have thicker surfaces if they inhabit stressful environments. Secondly, larger-celled organisms might experience larger turgor pressure, so it has been suggested that cell walls have to become thicker to prevent an increase in stress on the wall. Therefore, the thickness of the cell envelope should scale inversely with the diameter of the cell, implying that one should observe 1/3 power scaling between thickness and cell volume. More precisely, stress *σ* exerted on the cell wall is:

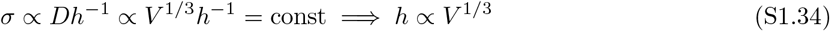

where *h* is envelope thickness, *V* is the volume of the cell [38]. Hence, as the cell volume increases, the benefit of producing less surface area relative to the cell’s volume may be canceled by that surface being thicker. Although we see a weak positive correlation, the scaling exponent is ~ 1*/*10 and thus much smaller than the expected 1/3 (left panel in Fig S1.8). This conclusion is not affected by the two outlier points with the thinnest envelopes, and the slope is almost identical when these are excluded.

**Figure S1.8:**
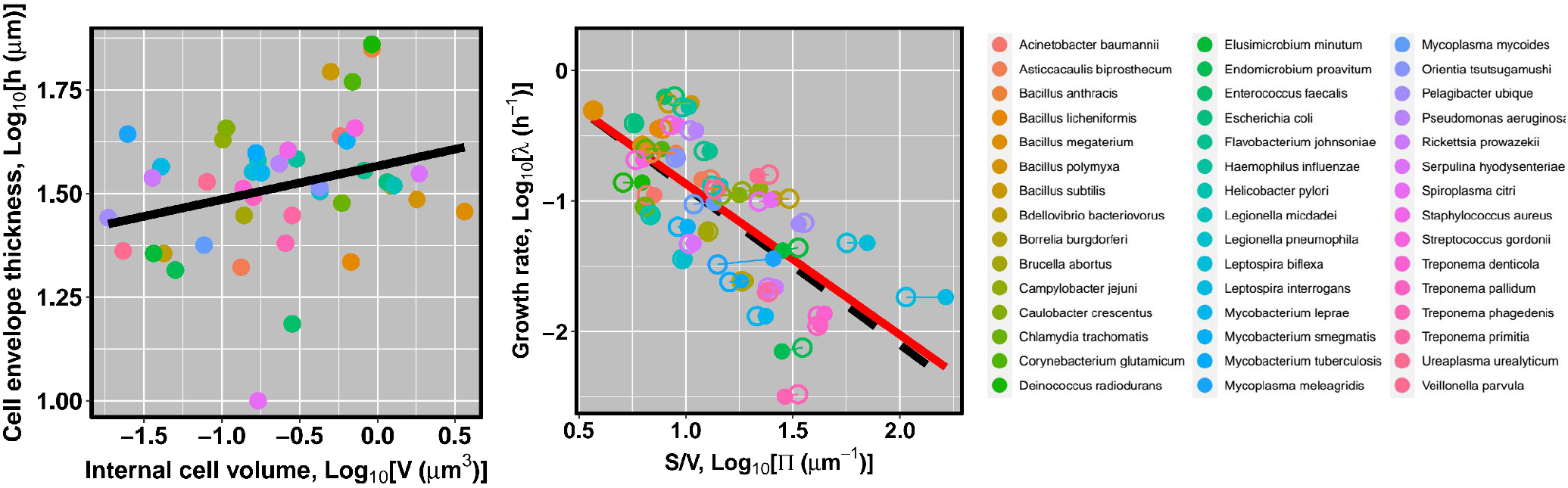
Variation in envelope thickness across bacteria. Left panel: Scaling of cell envelope thickness with internal volume of the cell. Regression equation is Log_10_[h]= (1.566 ± 0.034) + (0.081 ± 0.041)Log_10_[V] (*p* = 0.057, adjusted *R*^*2*^ = 0.06). Middle panel: Scaling of growth rate with surface-to-volume before (open circles) and after (filled circles) growth was corrected for cell envelope thickness. Black dashed and red solid lines denote regression equation for data before and after correction procedure, respectively. Right panel: Color code of plotted points. Sample size is 45 species.

Next, to correct for the difference in the cell envelope across species, we obtained the species-specific Π by first subtracting 2*h* from both width and length of the cell and then calculating the volume and the surface area; This gives us the S/V_cyt_ while accounting for the fact that cell envelope might differ from the assumed 30 nm. The correction shifts points toward larger (smaller) Π when the envelope is more (less) than 30 nm thick, as these cells ought to invest more (less) resources per unit of surface-to-volume. Note that correction also moves points along y-axis because of the correction for anaerobic metabolism depends on Π. We find that the transformation leaves points largely unaltered and still following the same trend expected by our theoretical expectations (Fig S1.8).

**Table S1.6:** Regression analysis of cell envelope-corrected data. Each element reports the regression coefficient and its standard error. Parentheses slope in the null hypothesis.

| | $\tilde{\lambda}_{\text{max}}$ (Generic thickness - 30 nm) | $\tilde{\lambda}_{\text{max}}$ (Species-specific thickness) |
| --- | --- | --- |
| $SV_{\text{cyt}}$ | $-1.205 \pm 0.189$ (-1) | $-1.155 \pm 0.192$ (-1) |
| $V_{\text{cyt}}$ | $0.376 \pm 0.136$ (0.33) | $0.374 \pm 0.137$ (0.33) |

### S1.11 Regression analysis in the limit of no degradation and large envelope costs

Figures S1.9 and S1.10 plot the regression lines reported in Table 3 in the main text.

**Figure S1.9:**
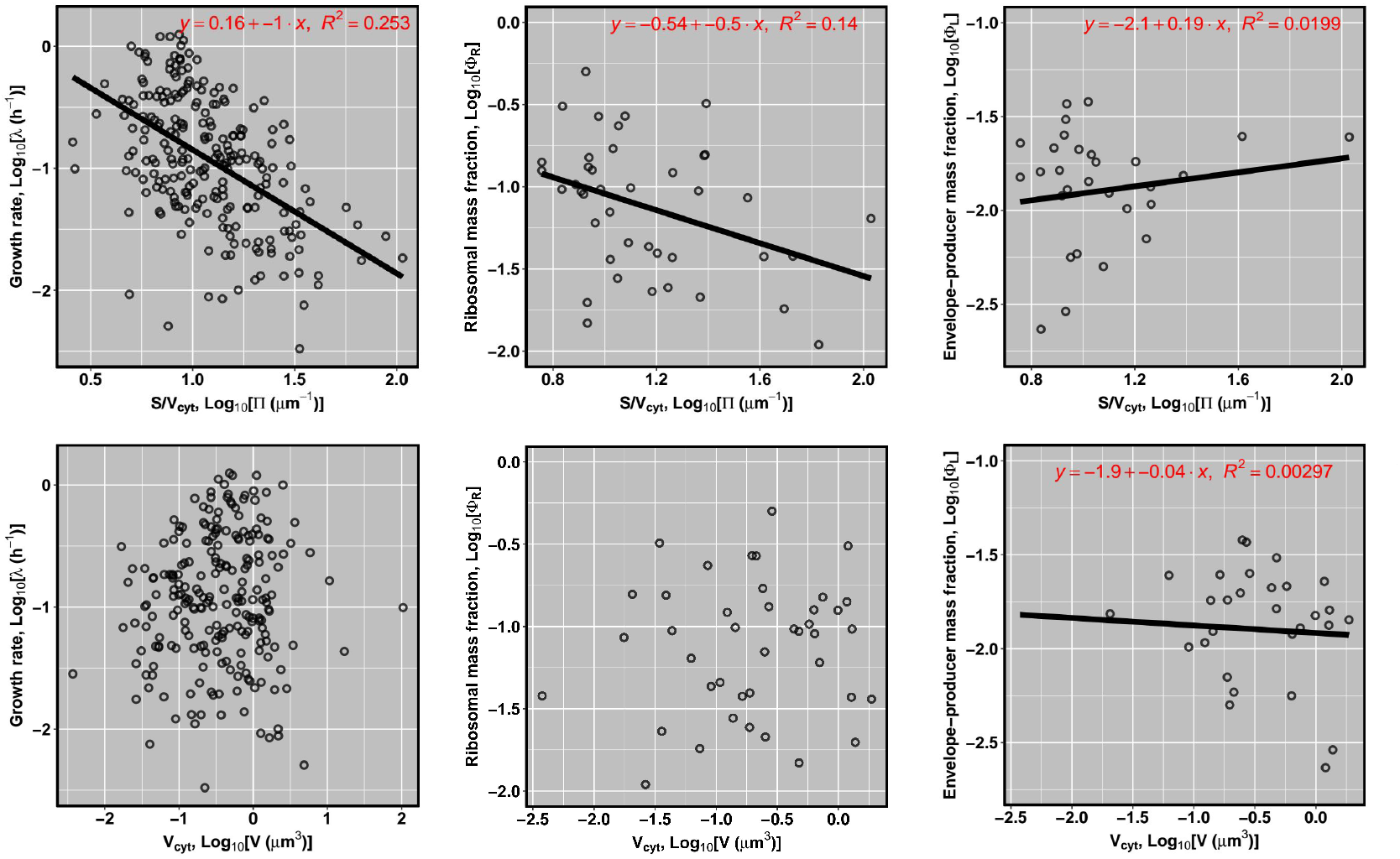
Regression analysis with S/V_cyt_ and V_cyt_ as independent variables.

**Figure S1.10:**
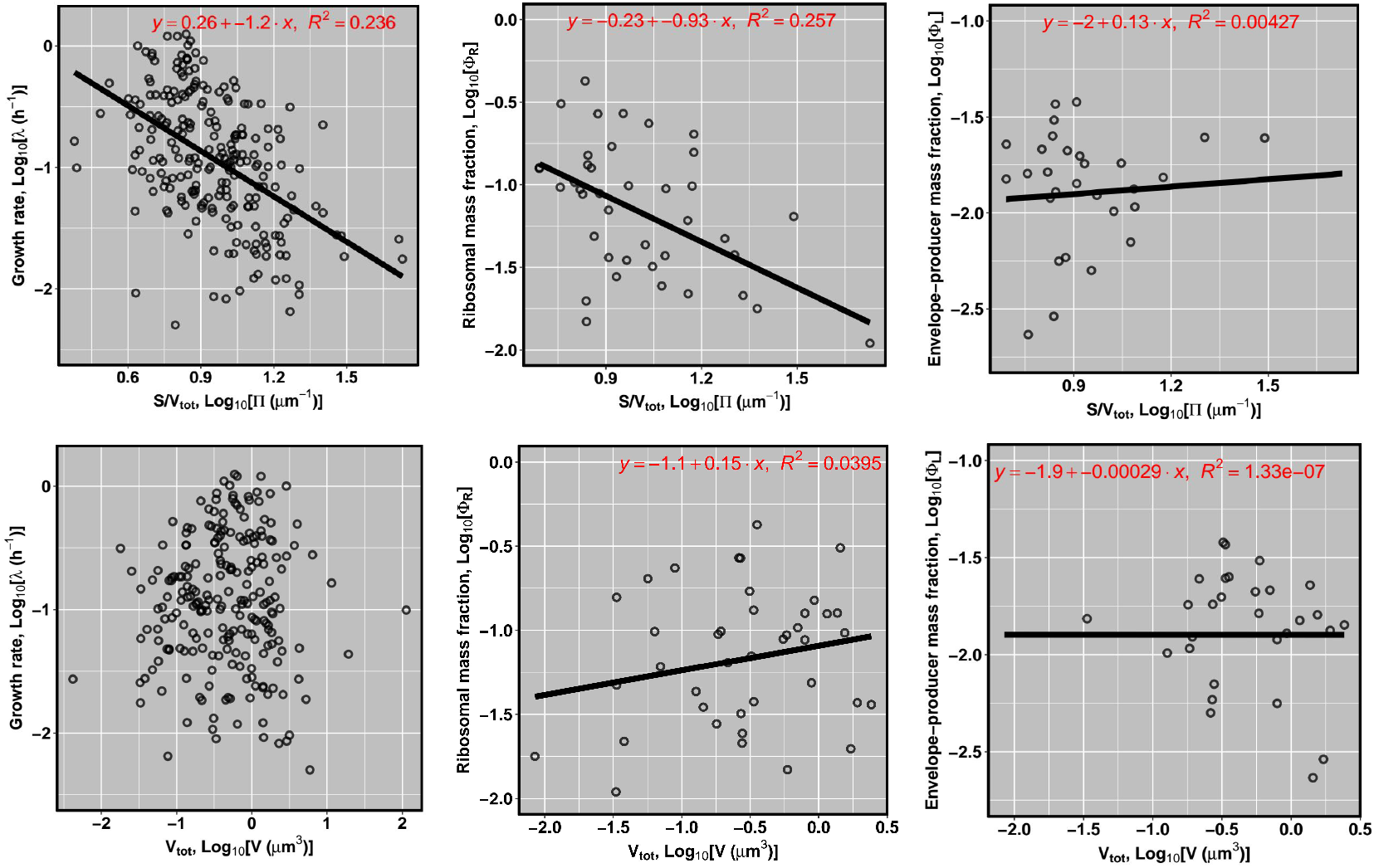
Regression analysis with S/V_tot_ and V_tot_ as independent variables.

### S1.12 Cross-validation of scaling patterns

Small spherical bacteria and elongated helical bacteria may be predominantly parasitic (*Mollicutes* and *Spirochaetes*), and that selection does not maximize growth to preserve the host. In that case, parasites might have lost the ability to achieve fast growth even outside of the host in a nutrient-replete environment. Although our data lacks environmental data, we looked at an independently collected sample that does have this information [39] and found a negative correlation between Π and the growth rate both in the pooled data as well as when the data was separated into free-living organisms and those that are host-associated (Fig S1.11). This scaling also holds if the regression analysis is separately applied to the free-living species (slope −0.87) and those that are associated with the host (slope −1.44), and both slopes are not significantly different from −1. The same conclusions hold if S:V_tot_ is used as an independent variable (see Table S1.7).

**Table S1.7:** Regression analysis of Madin et al. (2020) dataset. Description as in previous table. Sample size: all – 124, free-living – 85, host-associated – 39.

| | $\tilde{\lambda}_{\text{max}}$ (all bacteria) | $\tilde{\lambda}_{\text{max}}$ (free-living) | $\tilde{\lambda}_{\text{max}}$ (host-associated) |
| --- | --- | --- | --- |
| $SV_{\text{cyt}}$ | $-0.97 \pm 0.29$ ($-1$) | $-0.87 \pm 0.31$ ($-1$) | $-1.44 \pm 0.56$ ($-1$) |
| $SV_{\text{tot}}$ | $-1.02 \pm 0.32$ ($-1$) | $-0.92 \pm 0.33$ ($-1$) | $-1.52 \pm 0.61$ ($-1$) |

The scaling exponent is not significantly different from −1 (Table S1.7), indicating that the negative scaling is not because high-S:V region is dominated by host-associated organisms that might have been selected for slow growth.

**Figure S1.11:**
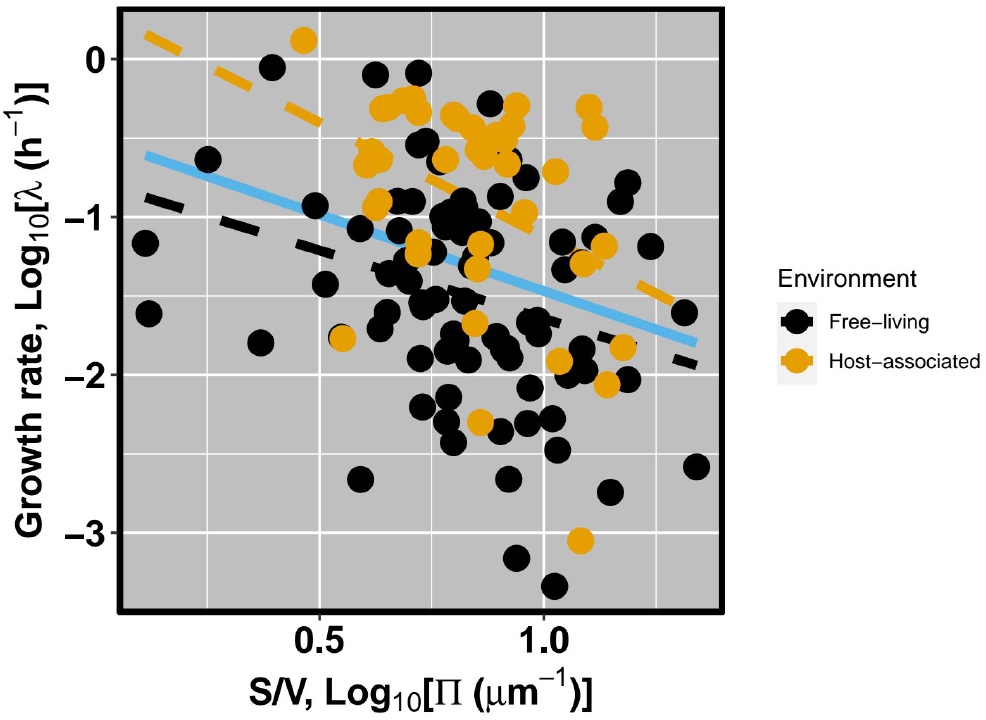
Scaling of the growth rate in Madin et al. (2020) dataset. Blue points – species with “host-” in isolation category. Red points – all other species. OLS regression denoted with lines: Solid black line – all species; Dashed black line – free-living species; Dashed grey line – host-associated species.

### S1.13 Scaling of translation-related genomic features

We obtained the number of genes coding for rRNA (197 species), and tRNA (198 species) from the NCBI Genome database [40]. We find a negative correlation between S:V_cyt_ and both tRNA and rRNA gene copy number and a positive correlation between the growth rate and the same genomic features (Fig S1.12).

**Figure S1.12:**
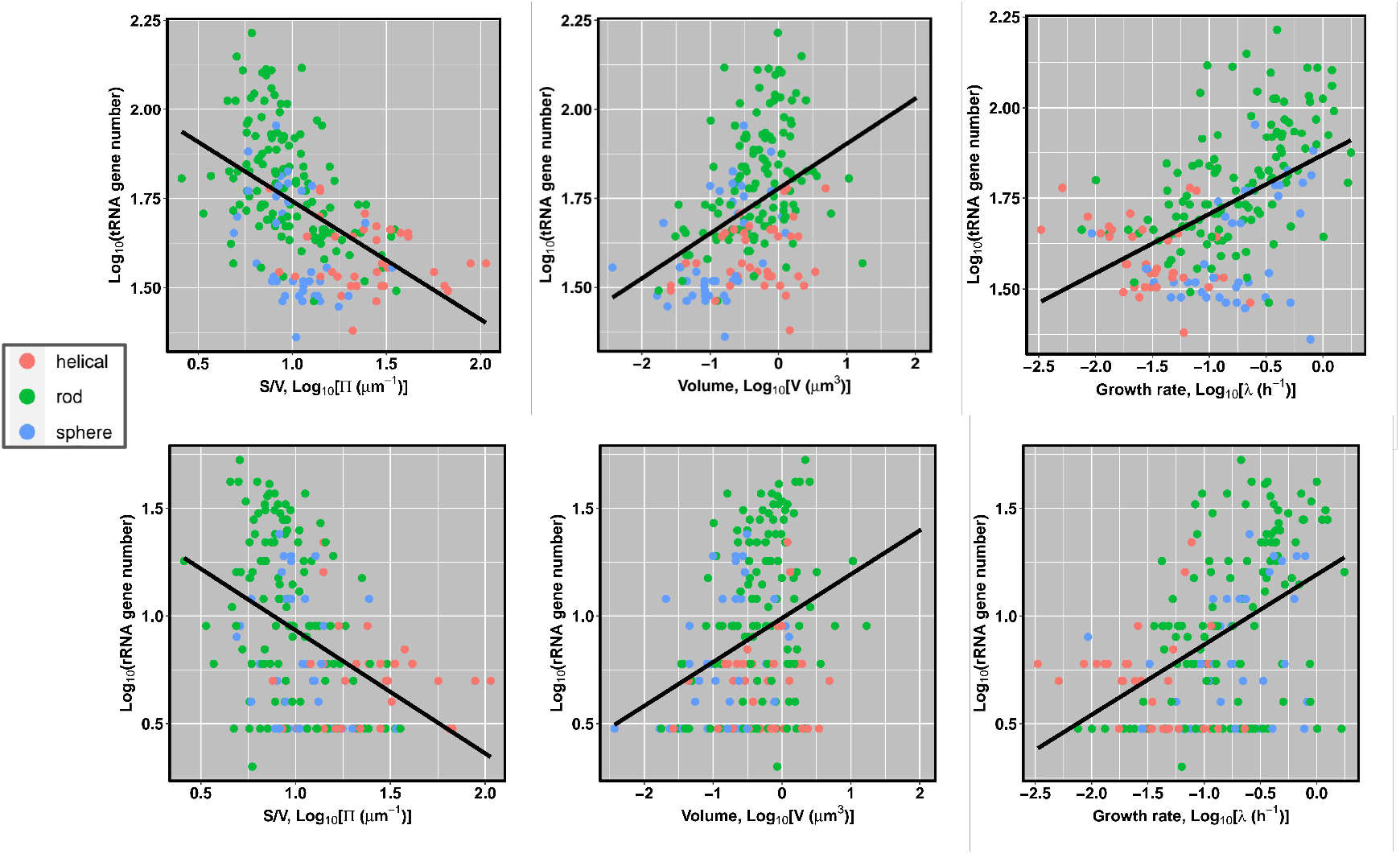
Size features as predictors of bacterial genomic features. Black line denotes OLS regression reported in Table S1.8.

**Table S1.8:** Predictors of tRNA and rRNA gene number. Dependent and independent variables are denoted in the first column, while the second column holds the estimated regression coefficient with standard errors. The adjusted R^2^ and p value are reported in the parentheses.

| Regression | Parameter estimate |
| --- | --- |
| $n(\text{tRNA}) \sim S:V$ | $-0.331 \pm 0.04, (0.26, 1.95 \times 10^{-14})$ |
| $n(\text{tRNA}) \sim V$ | $0.126 \pm 0.02, (0.16, 3.36 \times 10^{-9})$ |
| $n(\text{tRNA}) \sim \lambda$ | $0.163 \pm 0.02, (0.247, 5.82 \times 10^{-14})$ |
| $n(\text{rRNA}) \sim S:V$ | $-0.57 \pm 0.087, (0.177, 4.28 \times 10^{-10})$ |
| $n(\text{rRNA}) \sim V$ | $0.204 \pm 0.044, (0.096, 5.64 \times 10^{-6})$ |
| $n(\text{rRNA}) \sim \lambda$ | $0.325 \pm 0.042, (0.233, 4.39 \times 10^{-13})$ |

Three conclusions are reached. Firstly, we observe that the number of tRNA and rRNA gene copies increases with the growth rate, and both of these features decrease with S:V. Secondly, cytoplasmic volume *V* is a worse predictor of tRNA and rRNA gene repertoire than S:V. Thirdly and finally, S:V accounts for more variation in the number of tRNA genes than in the number of rRNA genes (Table S1.8).

### S1.14 Parameterization for the metabolic rate scaling

For parameterization of the metabolic rate equation, we use previously estimated model parameters (Table 2 in the main text), with an additional parameters retrieved from the literature (Table S1.9). Specifically, not only that one needs to know the rates of chemical reactions, but also how much ATP is consumed by these transformations. The cost is defined as the sum of the number of ATP molecules that has to be hydrolyzed to produce a particular conversion (e.g., from a building block into a unit of cell envelope), and the number of ATPs that could have been synthesized from NADH that was oxidized in the same process. This respectively corresponds to the direct costs and the opportunity cost of missed synthesis, as outlined in [41]. We assume that degradation of cell envelope constituents does not require energy, because we could not find data on ATP-dependence of cell wall hydrolases and other degradatory enzymes.

**Table S1.9:** Parameterization of the metabolic rate equation. † Estimated in the text. ° Assumed to be zero due to lack of data. * Total protein concentration (molecs/*μm*^*3*^) taken from [46], and the volume of *E. coli* taken as average across measurements in [9].

| Symbol | Definition | Value | Units | Source |
| --- | --- | --- | --- | --- |
| $N_l$ | Number of lipids in the cell | $22 \times 10^6$ | molecs | [42] |
| $N_{\text{pgn}}$ | Number of peptidoglycan disaccharide units | $3.5 \times 10^6$ | molecs | [42] |
| $N_l$ | Number of lipopolysaccharide molecules | $1.43 \times 10^6$ | molecs | [42] |
| $N_l$ | Number of Braun's lipoproteins | $10^6$ | molecs | [43] |
| $Q_{aa}$ | Direct costs of synthesizing an amino acid | 6 | ATP | [41] |
| $Q_t$ | Direct cost of elongating a polypeptide | 4 | ATP | [41] |
| $Q_l$ | Direct costs of synthesizing an envelope unit | 395 | ATP | † |
| $Q_{dp}$ | Direct costs of degrading a protein | $1 \times 325$ | ATP | [44, 45] |
| $Q_{dl}$ | Direct costs of degrading an envelope unit | 0 | ATP | ◦ |
| $m_s$ | Number of building blocks per envelope unit | 58 | molecs | † |
| $c_l$ | Cost of a lipid molecule | 74 | ATP | [43] |
| $c_{\text{pgn}}$ | Cost of a peptidoglycan disaccharide unit | 17 | ATP | [43] |
| $c_{\text{lps}}$ | Cost of a lipopolysaccharide molecule | 301 | ATP | [43] |
| $c_{aa}$ | Cost of incorporating an amino acid in protein | 10 | ATP | [41, 44] |
| $\gamma$ | Amino acids in the proteome per unit volume | $4.357 \times 10^9$ | molecs/$\mu\text{m}^3$ | [46]* |
| $S$ | Surface area of an *E. coli* cell | 4.32 | $\mu\text{m}^{-1}$ | [47] |

Parameter *m*_*s*_ is the total number of building block molecules required to produce a single unit of cell envelope, while *Q*_*l*_ is the number of ATPs used in the process of converting building blocks into an envelope unit. Because the surface area is originally expressed in terms of individual lipid molecules, the unit of cell envelope is composed of a lipid molecule and all the other contituents within the surface area of a single lipid molecule (i.e., peptidoglycan, lipoprotein, and lipopolysaccharide). Let *n*_*x*_ be the number of contituent *x* per unit area:

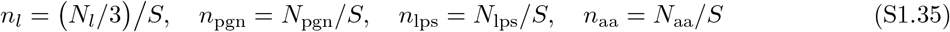

*N*_*l*_ was divided by 3 because the cell has three layers of lipids – two in inner membrane and one in outer membrane, given that the outer leaflet of the outer membrane contains lipopolysaccharide. Next, let *ρ*_*x*_ be the number of envelope constituent *x* per a single lipid molecule:

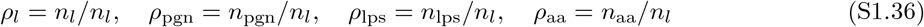

We can now calculate the number of ATP hydrolysis events required to convert building blocks into cell envelope unit:

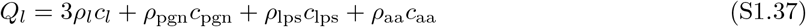

Lastly, the total number of building blocks needed for production of a single cell envelope unit is computed using costs of each constituent obtianed from [41, 43]:

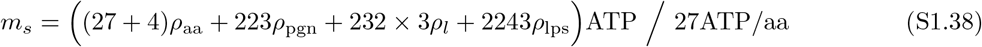

which is reported in Table S1.9.

