## Supplementary material for "Resource allocation to cell envelopes and the scaling of bacterial growth rate": Table_4_Combined Methylome, Transcriptome and Proteome Analyses Document Rapid Acclimatization of a Bacterium to Environmental Changes.pdf (from another version of this preprint: version 1, 2022-01-09)

**Supplementary Table S4.**

| Locus tag | Gene product | Log2 fold changes, 4ºC | p-value, 4ºC | Log2 fold changes, 26ºC* | p-value, 26ºC* |
|---|---|---|---|---|---|
| Pnuc_0763 | Citrate synthase | -0.3 | 0.0470334 | -0.32 | 0.0279037 |
| Pnuc_0942 | Aconitate hydratase B | 0.42 | 0.00586863 | 0.22 | 0.0122051 |
| Pnuc_0366 | Isocitrate dehydrogenase [NADP] | 0.41 | 0.000379697 | 0.39 | 7.12561E-05 |
| Pnuc_0840 | 2-oxoglutarate dehydrogenase, E1 subunit | -0.85 | 0.000238405 | -1.11 | 0.000146018 |
| Pnuc_0841 | 2-oxoglutarate dehydrogenase complex component E2 | -0.84 | 0.000496529 | -0.93 | 0.00022814 |
| Pnuc_0842 | Dihydrolipoyl dehydrogenase | -0.89 | 0.000745269 | -0.97 | 0.000568912 |
| Pnuc_1830 | Succinyl-CoA synthetase subunit alpha | -0.12 | 0.517745 | -0.2 | 0.202775 |
| Pnuc_1831 | Succinyl-CoA synthetase subunit beta | 0.069 | 0.680308 | -0.35 | 0.0459234 |
| Pnuc_0760 | Succinate dehydrogenase flavoprotein subunit | -0.32 | 0.0129625 | -0.49 | 8.07913E-05 |
| Pnuc_0761 | Succinate dehydrogenase subunit B | -0.66 | 0.000023 | -0.78 | 0.0006707 |
| Pnuc_1139 | Fumarate hydratase class I | -0.34 | 0.107649 | -0.35 | 0.0114142 |
| Pnuc_1618 | Fumarate hydratase class II | -0.05 | 0.814663 | 0.41 | 0.0339648 |
| Pnuc_1616 | Malate:quinone oxidoreductase | -3.05 | 0.18391 | -0.88 | 0.00848289 |
| Pnuc_0756 | Malate dehydrogenase | -0.44 | 0.00128864 | 0.0004 | 0.99599 |
| Pnuc_1043 | NADH-quinone oxidoreductase subunit I | -5.38 | 0.0208763 | -2.46 | 0.000255728 |
| Pnuc_1045 | NADH-quinone oxidoreductase subunit G | 0.47 | 0.0121389 | 0.28 | 0.014716 |
| Pnuc_1046 | NADH-quinone oxidoreductase subunit F | -0.55 | 0.0268552 | -0.75 | 0.00444586 |
| Pnuc_1048 | NADH-quinone oxidoreductase subunit D | -0.56 | 0.0039125 | -0.76 | 0.000154393 |
| Pnuc_1049 | NADH-quinone oxidoreductase subunit C | -0.59 | 0.00825663 | -0.37 | 0.018118 |
| Pnuc_1050 | NADH-quinone oxidoreductase subunit B | -1.85 | 0.456000 | 1.02 | 0.577000 |
| Pnuc_1936 | Cytochrome c oxidase, subunit III | -2.42 | 0.179170 | -0.67 | 0.014248 |
| Pnuc_1939 | Cytochrome c oxidase subunit 1 | -2.93 | 0.305045 | -0.71 | 0.000522 |
| Pnuc_1940 | Cytochrome c oxidase subunit 2 | -0.74 | 0.016616 | -0.84 | 0.000684 |
| Pnuc_0033 | Cytochrome c, class I | -1.00 | 0.002600 | -0.36 | 0.021062 |
| Pnuc_0022 | ATP synthase subunit b | -0.50 | 0.002802 | -0.50 | 0.002976 |
| Pnuc_0023 | ATP synthase subunit delta | -0.73 | 0.012487 | -0.73 | 0.001111 |
| Pnuc_0024 | ATP synthase subunit alpha | -0.40 | 0.002058 | -0.64 | 0.000566 |
| Pnuc_0025 | ATP synthase gamma chain | -0.93 | 0.018958 | -0.85 | 0.004344 |
| Pnuc_0026 | ATP synthase subunit beta | -0.50 | 0.021835 | -0.62 | 0.000378 |
| Pnuc_0027 | ATP synthase epsilon chain | -0.33 | 0.039365 | -0.35 | 0.007643 |

**Supplementary Table S4.** Tabulation of differentially expressed proteins obtained from 4ºC incubated / UV irradiated (26ºC*) grown cells participating in TCA cycle and respiratory chain.

**Supplementary Table S5.**

**UV treated sample (26ºC incubated)**

| Locus Tag | Abbreviation / Gene product | Function | References |
|---|---|---|---|
| Pnuc_0173 | Ahp: Alkyl hydroperoxide reductase (peroxiredoxin) | Cytosolic protein that can sense peroxide and protect other proteins by acting as an anti-oxidant. | Dubbs, J. M., and Mongkolsuk, S. (2007). Peroxiredoxins in bacterial antioxidant defense. *Subcell. Biochem*. 44, 143-193. |
| Pnuc_0196 | Obg | Stringent response GTPase that prevents ribosome assembly by binding to peptidyl transferase center of 50S subunit. | Feng, B., Mandava, C. S., Guo, Q., Wang, J., Cao, W., Li, N., et al. (2014). Structural and functional insights into the mode of action of a universally conserved Obg GTPase. *PLoS Biol*. 12, e1001866. |
| Pnuc_0254 | RfbD: dTDP-4-dehydro rhamnose reductase | Lipopolysaccharide core biosynthesis. | Jofré, E., Lagares, A., and Mori, G. (2004). Disruption of dTDP-rhamnose biosynthesis modifies lipopolysaccharide core, exopolysaccharide production, and root colonization in *Azospirillum brasilense*. *FEMS Microbiol. Lett*. 231, 267-275. |
| Pnuc_0257 | RfaC: Lipopolysaccharide heptosyltransferase I | Participates in outer membrane lipid core oligosaccharide building-up. | Kadrmas, J. L., Raetz, C. R. (1998). Enzymatic properties of lipopolysaccharide in *Escherichia coli*. Purification and properties of heptosyltransferase I. *J. Biol. Chem*. 273, 2799–2807. |
| Pnuc_0282 | KdsB: 3-deoxy-D-manno-octulosonate cytidylyltransferase | Lipopolysaccharide biosynthesis. | Kohlbrenner, W. E., Nuss, M. M., Fesik, S. W. (1987). $^{31}$P and $^{13}$C NMR studies of oxygen transfer during catalysis by 3-deoxy-D-manno-octulosonate cytidylyltransferase from *Escherichia coli*. *J. Biol. Chem*. 262, 4534-4537. |
| Pnuc_0284 | LpxK: Tetraacyldisaccharide 4'-kinase | Lipopolysaccharide core biosynthesis. | Garrett, T. A., Que, N. L., Raetz, C. R. (1998). Accumulation of a lipid A precursor lacking the 4'-phosphate following inactivation *of the Escherichia coli lpxK* gene. *J. Biol. Chem*. 273, 12457-12465 |

| | | | |
|---|---|---|---|
| Pnuc_0379 | PpiC: PpiC-type peptidyl-prolyl cis-trans isomerase | Catalyzes protein folding. Mutant carrying defective *ppiC* gene is sensitive to oxidative stress. | Rahfeld, J. U., Schierhorn, A., Mann, K., Fischer, G. (1994). A novel peptidyl-prolyl cis/trans isomerase from *Escherichia coli*. *FEBS Lett*. 343,65-69. |
| Pnuc_0429 | Ahp: Alkyl hydroperoxide reductase (peroxiredoxin) | Cytosolic enzyme that can detoxify various peroxides. | Parsonage, D., Karplus, P. A., Poole, L. B. (2008). Substrate specificity and redox potential of AhpC, a bacterial peroxiredoxin. *Proc. Natl. Acad. Sci. USA*. 105, 8209-8214. |
| Pnuc_0487 | Pirin | Interacts with the pyruvate dehydrogenase and subsequently inhibits its activity. | Soo, P. C., Horng, Y. T., Lai, M. J., Wei, J. R., Hsieh, S. C., Chang, Y. L., et al. (2007). Pirin regulates pyruvate catabolism by interacting with the pyruvate dehydrogenase E1 subunit and modulating pyruvate dehydrogenase activity. *J. Bacteriol*. 189, 109-118. |
| Pnuc_0610 | MsbA: Lipid A ABC exporter | Inner membrane lipid flippase. | Zhang, Y. M., Rock, C. O. (2008). Membrane lipid homeostasis in bacteria. *Nat. Rev. Microbiol*. 6, 222-233. |
| Pnuc_0611 | RNAse G | Endoribonuclease that participates in the maturation of tRNA, processing of rRNA and decay of mRNA. | Bernardini, A., Martínez, J. L. (2017). Genome-wide analysis shows that RNase G plays a global role in the stability of mRNAs in *Stenotrophomonas maltophilia*. *Sci. Rep*. 7, 16016. |
| Pnuc_0734<br>Pnuc_0735<br>Pnuc_0736 | PDH: Pyruvate dehydrogenase E1 component<br>PDH: Pyruvate dehydrogenase E2 component<br>PDH: Pyruvate dehydrogenase E3 component | Catalyzes the oxidative decarboxylation of pyruvate to form acetyl-CoA and $CO_2$. | Patel, M. S., Nemeria, N. S., Furey, W., and Jordan, F. (2014). The pyruvate dehydrogenase complexes: structure-based function and regulation. *J. Biol. Chem*. 289, 16615-16623. |
| Pnuc_0740 | IclR: Isocitrate lyase repressor | Represses the operon of *aceBAK* by binding directly to the promoter (aceB: malate synthase; aceA: isocitrate lyase monomer; aceK: isocitrate dehydrogenase kinase-phosphatase) | Pan, B., Unnikrishnan, I., LaPorte, D. C. (1996). The binding site of the IclR repressor protein overlaps the promoter of *aceBAK*. *J. Bacteriol*. 178, 3982-3984. |

| | | | |
|---|---|---|---|
| Pnuc_0786<br>Pnuc_0787 | CytC: Cytochrome C complex | Heme protein that transfers electron between complex II and complex IV in the electron transport chain. Iron ions are released from CytC during peroxide exposure that causes reactive ferryl-heme species based lipid peroxidation and reactive oxygen species cause DNA damage. | Kim, N. H., and Kang, J. H. (2006). Oxidative damage of DNA induced by the cytochrome C and hydrogen peroxide system. *J. Biochem. Mol. Biol*. 39, 452-456. |
| Pnuc_0918 | LplT: Lysophospholipid transporter | Flippase | Zhang, Y. M., and Rock, C. O. (2008). Membrane lipid homeostasis in bacteria. *Nat. Rev. Microbiol*. 6, 222-233. |
| Pnuc_0949 | Hsp33: 33 kDa heat shock protein | Chaperone function is activated during oxidative stress, senses oxidizing condition e.g. H2O2 and prevents protein aggregation. | Graf, P. C., and Jakob, U. (2002). Redox-regulated molecular chaperones. *Cell Mol. Life Sci.* 59, 1624-1631. |
| Pnuc_1044 | NDH: respiratory-chain NADH dehydrogenase subunit 1 | Respiratory chain enzyme that catalyzes the transfer of electron from NADH to coenzyme Q10. | Brandt, U. (2006). Energy converting NADH:quinone oxidoreductase (complex I). *Annu. Rev. Biochem*. 75, 69-92. |
| Pnuc_1131 | AlpA: Phage positive transcriptional regulator | DNA binding regulator that suppresses cellular sensitivity towards UV radiation in lon mutant cells. | Trempy, J. E., Kirby, J. E., and Gottesman, S. (1994). Alp suppression of Lon: dependence on the *slpA* gene. *J. Bacteriol*. 176, 2061-2067. |
| Pnuc_1263 | YggX: Fe(II) trafficking protein | Fe-S cluster protection protein. Mediates the repairing of iron-sulfur clusters effected by oxidative damage. | Velayudhan, J., Karlinsey, J. E., Frawley, E. R., Becker, L. A., Nartea, M., and Fang, F. C. (2014). Distinct roles of the *Salmonella enterica* serovar *Typhimurium* CyaY and YggX proteins in the biosynthesis and repair of iron-sulfur clusters. *Infect. Immun*. 82, 1390-1401. |
| Pnuc_1280 | MS: Malate synthase G | Glyoxylate shunt enzyme that catalyzes the condensation of acetyl-CoA and glyoxylate to yield malate and CoA. Response towards reactive oxygen species. | Slade, D., and Radman, M. (2011). Oxidative stress resistance in *Deinococcus radiodurans*. *Microbiol. Mol. Biol. Rev*. 75, 133-191 . |

| | | | |
|---|---|---|---|
| Pnuc_1376 | Rbr: Rubrerythrin | Cytosolic enzyme that is activated during oxidative stress. Converts harmful peroxide into water. | Lehmann, Y., Meile, L., and Teuber, M. (1996). Rubrerythrin from *Clostridium perfringens*: cloning of the gene, purification of the protein, and characterization of its superoxide dismutase function. *J. Bacteriol*. 178, 7152-7158. |
| Pnuc_1435 | DAP: Dipeptidyl aminopeptidase | Extracellular enzyme that catalyse the hydrolysis and sequential release of N-terminal dipeptides of polypeptides. | Gonzales, T., and Robert-Baudouy, J. (1996). Bacterial aminopeptidases: properties and functions. *FEMS Microbiol. Rev*. 18, 319-344. |
| Pnuc_1449 | UK: Uridylate kinase | Phosphorylation of UMP to UDP (that eventually forms thymine and cytosine via pyrimidine biosynthetic pathway). | Serina, L., Blondin, C., Krin E, Sismeiro, O., Danchin, A., Sakamoto, H., et al. (1995). *Escherichia coli* UMP-kinase, a member of the aspartokinase family, is a hexamer regulated by guanine nucleotides and UTP. *Biochemistry*. 34, 5066-5074. |
| Pnuc_1452 | Map: Methionine aminopeptidase | Catalyzes the hydrolysis of N-terminal methionine from newly translated polypeptide chain. | Xiao, Q., Zhang, F., Nacev, B. A., Liu, J. O., and Pei, D. (2010). Protein N-terminal processing: substrate specificity of *Escherichia coli* and human methionine aminopeptidases. *Biochemistry*. 49, 5588-5599. |
| Pnuc_1489 | HscA: Heat shock cognate 66-kDa | Chaperone protein involved in the maturation of Fe-S proteins. | Chandramouli, K., and Johnson, M. K. (2006). HscA and HscB stimulate [2Fe-2S] cluster transfer from IscU to apo-ferredoxin in an ATP-dependent reaction. *Biochemistry*. 45, 11087-11095. |
| Pnuc_1490 | HscB: Heat shock cognate 20-kDa | Co-chaperone protein involved in the maturation of Fe-S proteins. | Chandramouli, K., and Johnson, M. K. (2006). HscA and HscB stimulate [2Fe-2S] cluster transfer from IscU to apo-ferredoxin in an ATP-dependent reaction. *Biochemistry*. 45, 11087-11095. |
| Pnuc_1494 | IscR: Iron-Sulfur cluster regulator | Repressor of *isc* gene cluster (i.e. iron-sulfur biogenesis) and is induced during oxidative stress. | Crack, J. C., Green, J., Thomson, A. J., and Le Brun, N. E. (2012). Iron-sulfur cluster sensor-regulators. *Curr. Opin. Chem. Biol*. 16, 35-44 |
| Pnuc_1506 | ICL: Isocitrate lyase | Glyoxylate pathway enzyme. Catalyses cleavage of isocitrate into glyoxylate and succinate | Slade, D., and Radman, M. (2011). Oxidative stress resistance in *Deinococcus radiodurans*. *Microbiol. Mol. Biol. Rev*. 75, 133-191. |

| | | | |
|---|---|---|---|
| | | thereby bypassing TCA cycle. Response towards reactive oxygen species. | |
| Pnuc_1522 | WrbA: Tryptophan (W) repressor-binding flavodoxin-like protein | Defense against oxidative stress and has NAD(P)H :quinone oxidoreductases-like activity that allows quinone detoxification. | Patridge, E. V., and Ferry, J. G. (2006). WrbA from *Escherichia coli* and *Archaeoglobus fulgidus* is an NAD(P)H:quinone oxidoreductase. *J. Bacteriol*. 188, 3498-3506. |
| Pnuc_1534 | Ahp: Alkyl hydroperoxide reductase | Cytosolic enzyme that can reduce organic peroxides. | Poole, L. B., and Ellis, H. R. (1996). Flavin-dependent alkyl hydroperoxide reductase from *Salmonella typhimurium*. 1. purification and enzymatic activities of overexpressed AhpF and AhpC proteins. *Biochemistry*. 35, 56-64. |
| Pnuc_1626 | SOD: Superoxide dismutase, copper/zinc binding protein | Periplasmic enzyme (signal peptide predicted). Dismutation of superoxide into hydrogen peroxide and oxygen. Superoxides can otherwise attack DNA bases and Fe-S clusters of various enzymes. | De Groote, M. A., Ochsner, U. A., Shiloh, M. U., Nathan, C., McCord, J. M., Dinauer, M. C., et al. (1997). Periplasmic superoxide dismutase protects *Salmonella* from products of phagocyte NADPH-oxidase and nitric oxide synthase. *Proc. Natl. Acad. Sci. USA*. 94, 13997-14001 |
| Pnuc_1650 | S16: lon domain peptidase | Intracellular Lon ATP-dependent protease (related to MEROPS peptidase family S16) that degrades damaged poplypeptides. Lon protease increases tolerance towards UV sensitivity. | Tsilibaris, V., Maenhaut-Michel, G., and Van Melderen, L. (2006). Biological roles of the Lon ATP-dependent protease. *Res. Microbiol*. 157, 701-713. |
| Pnuc_1715 | UvrD: ATP-dependent DNA helicase | Transcription-coupled DNA repair enzyme. Directly rescues the replication fork blocked at transcription level by backtracking RNA polymerase. | Epshtein, V. (2015). UvrD helicase: an old dog with a new trick: how one step backward leads to many steps forward. *Bioessays*. 37, 12-19. |
| Pnuc_1732 | FkpB: Peptidyl-prolyl cis-trans isomerase | Cytosolic protein and has chaperone activity. | Jo, G. A., Lee, J. M., No, G., Kang, D. S., Kim, S. H., Ahn, S. H., et al. (2015). Isolation and characterization of a 17-kDa FKBP-type peptidyl-prolyl cis/trans isomerase from *Vibrio anguillarum*. *Prot. Express Purif*. 110, 130-137. |
| Pnuc_1769 | DnaK: Heat shock protein | Molecular chaperone that binds to the newly | Tomoyasu, T., Ogura, T., Tatsuta, T., and Bukau, B. (1998). Levels |

| | | | |
|---|---|---|---|
| | encoded by *dnaK* locus | synthesized protein and mediates its folding. Unfolded peptides are transferred to GroEL chaperone. | of DnaK and DnaJ provide tight control of heat shock gene expression and protein repair in *Escherichia coli*. *Mol. Microbiol*. 30, 567-581. |
| Pnuc_1805 | GroEL: Large polypeptide chaperone encoded by *groE* locus | Molecular chaperone. Mutation in *E. coli groL* gene makes them susceptible to UV-radiation. | Sargentini, N. J., Gularte, N. P., and Hudman, D. A. (2016). Screen for genes involved in radiation survival of *Escherichia coli* and construction of a reference database. *Mutat Res*. 793-794, 1-14. |
| Pnuc_1823 | Ste24p: MEROPS peptidase family M47 | Integral membrane endopeptidase. | Pryor, E. E. Jr., Horanyi, P. S., Clark, K. M., Fedoriw, N., Connelly, S. M., Koszelak-Rosenblum, M., et al. (2013). Structure of the integral membrane protein CAAX protease Ste24p. *Science*. 339, 1600-1604. |
| Pnuc_1869 | SurA: PpiC-type peptidyl-prolyl cis-trans isomerase | Periplasmic chaperone responsible for correct folding of outer membrane proteins. | Xu, X., Wang, S., Hu, Y. X., and McKay, D. B. (2007). The periplasmic bacterial molecular chaperone SurA adapts its structure to bind peptides in different conformations to assert a sequence preference for aromatic residues. *J. Mol. Biol*. 373, 367-381. |
| Pnuc_2008 | ExoIII: Exodeoxyribonuclease III | Involved in base excision repair using endonucleolytic activity at apurinic/ apyrimidinic site. *E. coli* mutant in this gene display sensitivity towards UV treatment. | Sammartano, L. J., Tuveson, R. W. (1983). *Escherichia coli xthA* mutants are sensitive to inactivation by broad-spectrum near-UV (300- to 400-nm) radiation. *J. Bacteriol*. 156, 904-906. |
| Pnuc_2021 | MreB: Rod shape-determining protein | Cytoskeletal protein that determines bacterial cell shape. Active movement of MreB creates fluidity in the cell membrane. | Strahl, H., Bürmann, F., Hamoen, L. W. (2014). The actin homologue MreB organizes the bacterial cell membrane. *Nat. Commun*. 5, 3442 |
| Pnuc_2053 | Cat: Catalase | Periplasmic enzyme (signal peptide predicted). Possesses catalase and peroxidase activity. Scavenges harmful peroxide at high concentration and converts it into water and oxygen. | Heimberger, A., Eisenstark, A. (1988). Compartmentalization of catalases in *Escherichia coli*. *Biochem. Biophys. Res. Commun*. 154, 392-397. |

| | | | |
|---|---|---|---|
| Pnuc_R0014 | tmRNA: transfer-messenger RNA | Ribonucleic acid with tRNA and mRNA properties. Rescues and recycles stalled translation complexes by tagging incompletely synthesized peptides for eventual proteolysis. | Janssen, B. D., and Hayes, C. S. (2012). The tmRNA ribosome rescue system. *Adv. Protein Chem. Struct. Biol*. 86, 151-191. |

**4ºC incubated sample**

| Locus Tag | Abbreviation / Gene product | Function | References |
|---|---|---|---|
| Pnuc_0032 | Des: Fatty acid desaturase | Polyunsaturated fatty acid biosynthesis enzyme that introduces double bond in saturated fatty acids. | Mansilla, M. C., Cybulski, L. E., Albanesi, D., and de Mendoza, D. (2004). Control of membrane lipid fluidity by molecular thermosensors. *J. Bacteriol*. 186, 6681-6688. |
| Pnuc_0132 | Phasin | Proteins surrounding PHA granules. Phasin has been shown to be expressed in other cold-adapted bacteria. | Ting, L., Williams, T. J., Cowley, M. J., Lauro, F. M., Guilhaus, M., Raftery, M. J., et al. (2010). Cold adaptation in the marine bacterium, *Sphingopyxis alaskensis*, assessed using quantitative proteomics. *Environ. Microbiol*. 12, 2658-2676. |
| Pnuc_0173 | Ahp: Alkyl hydroperoxide reductase (peroxiredoxin) | Cytosolic protein that can sense peroxide and protect other proteins by acting as an anti-oxidant. | Dubbs, J. M., and Mongkolsuk, S. (2007). Peroxiredoxins in bacterial antioxidant defense. *Subcell. Biochem*. 44, 143-93. |
| Pnuc_0429 | Ahp: Alkyl hydroperoxide reductase (peroxiredoxin) | Cytosolic enzyme that can detoxify various peroxides. | Parsonage, D., Karplus, P. A., and Poole, L. B. (2008). Substrate specificity and redox potential of AhpC, a bacterial peroxiredoxin. *Proc. Natl. Acad. Sci. USA*. 105, 8209-8214. |
| Pnuc_0586 | CSP: Cold-shock DNA-binding protein | Destabilizes secondary structures of RNA during cold-shock and acts as an RNA chaperone. | Jiang, W., Hou, Y., and Inouye, M. (1997). CspA, the major cold-shock protein of *Escherichia coli*, is an RNA chaperone. *J. Biol. Chem*. 272, 196-202. |

| | | | |
|---|---|---|---|
| Pnuc_0611 | RNAse G | Endoribonuclease that participates in the maturation of tRNA, processing of rRNA and decay of mRNA. | Ow, M. C., Perwez, T., and Kushner, S. R. (2003). RNase G of *Escherichia coli* exhibits only limited functional overlap with its essential homologue, RNase E. *Mol. Microbiol*. 49, 607-622. |
| Pnuc_0666 | Alanine dehydrogenase | Interconversion of L-alanine and pyruvate by reversible oxidative deamination activity / glyoxylate reductive aminase activity. | Giffin, M. M., Modesti, L., Raab, R. W., Wayne, L. G., and Sohaskey, C. D. (2012). *ald* of *Mycobacterium tuberculosis* encodes both the alanine dehydrogenase and the putative glycine dehydrogenase. *J. Bacteriol*. 194, 1045–1054. |
| Pnuc_0670 | Cyanophycin synthetase | Catalyzes the synthesis of biopolymer reserve multi-L-arginyl-poly-L-aspartic acid, i.e. cyanophycin. | Ziegler, K., Diener, A., Herpin, C., Richter, R., Deutzmann, R., and Lockau, W. (1998). Molecular characterization of cyanophycin synthetase, the enzyme catalyzing the biosynthesis of the cyanobacterial reserve material multi-L-arginyl-poly-L-aspartate (cyanophycin). *Eur. J. Biochem*. 254, 154-159. |
| Pnuc_0774<br>Pnuc_0913 | ACC: Acetyl-coenzyme A carboxylase carboxyl transferase subunit beta<br>ACC: Acetyl-CoA carboxylase, biotin carboxylase | Catalyzes the first step of long-chain fatty acid biosynthesis, i.e. conversion of acetyl-CoA into malonyl-CoA. | Zhang, Y-M., and Rock, C. O. (2008). Membrane lipid homeostasis in bacteria. *Nat. Rev. Microbiol*. 6, 222-233. |
| Pnuc_1489 | HscA: Heat shock cognate 66-kDa | Chaperone protein involved in the maturation of Fe-S proteins. Can be synthesized during cold-shock response. | Lelivelt, M. J., and Kawula, T. H. (1995). Hsc66, an Hsp70 homolog in *Escherichia coli*, is induced by cold shock but not by heat shock. *J. Bacteriol*. 177, 4900-4907. |
| Pnuc_1490 | HscB: Heat shock cognate 20-kDa | Co-chaperone protein involved in the maturation of Fe-S proteins. | Chandramouli, K., and Johnson, M. K. (2006). HscA and HscB stimulate [2Fe-2S] cluster transfer from IscU to apo-ferredoxin in an ATP-dependent reaction. *Biochemistry*. 45, 11087-11095. |
| Pnuc_1497 | PhaZ: Polyhydroxyalkanoate depolymerase (intracellular) | Increased polyhydroxyalkanoate (PHA) degradation for extracting energy and carbon. | Knoll, M., Hamm, T. M., Wagner, F., Martínez, V., and Pleiss, J. (2009). The PHA depolymerase engineering database: a systematic analysis tool for the diverse family of polyhydroxyalkanoate (PHA) depolymerases. *BMC Bioinform*. 10, 89. |

| | | | |
|---|---|---|---|
| Pnuc_1534 | Ahp: Alkyl hydroperoxide reductase | Cytosolic enzyme that can reduce organic peroxides. | Poole, L. B., and Ellis, H. R. (1996). Flavin-dependent alkyl hydroperoxide reductase from *Salmonella typhimurium*. 1. purification and enzymatic activities of overexpressed AhpF and AhpC proteins. *Biochemistry*. 35, 56-64. |
| Pnuc_1626 | SOD: Superoxide dismutase, copper/zinc binding protein | Periplasmic enzyme (signal peptide predicted). Dismutation of superoxide into hydrogen peroxide and oxygen. Superoxides can otherwise attack DNA bases and Fe-S clusters of various enzymes. | De Groote, M. A., Ochsner, U. A., Shiloh, M. U., Nathan, C., McCord, J.M., Dinauer, M. C., et al. (1997). Periplasmic superoxide dismutase protects *Salmonella* from products of phagocyte NADPH-oxidase and nitric oxide synthase. *Proc. Natl. Acad. Sci. USA*. 94, 13997-14001. |
| Pnuc_1732 | FkpB: Peptidyl-prolyl cis-trans isomerase | Cytosolic protein and has chaperone activity. Possibly involved in cold adaptation. | Suzuki, Y., Haruki, M., Takano, K., Morikawa, M., and Kanaya, S. (2004). Possible involvement of an FKBP family member protein from a psychrotrophic bacterium *Shewanella* sp. SIB1 in cold-adaptation. *Eur. J. Biochem*. 271, 1372-1381. |
| Pnuc_1811 | Sua5: Translation factor | Required for threonylcarbamoyl group modification on adenine (t6A37) of the tRNA to recognize ANN codons thereby helping in accurate translation by stabilizing tRNA codon interactions with the ribosome. | El Yacoubi, B., Lyons, B., Cruz, Y., Reddy, R., Nordin, B., Agnelli, F., et al. (2009). The universal YrdC/Sua5 family is required for the formation of threonylcarbamoyladenosine in tRNA. *Nucleic Acids Res*. 37, 2894–2909. |
| Pnuc_1869 | SurA: PpiC-type peptidyl-prolyl cis-trans isomerase | Periplasmic chaperone responsible for correct folding of outer membrane proteins. | Xu, X., Wang, S., Hu, Y. X., and McKay, D. B. (2007). The periplasmic bacterial molecular chaperone SurA adapts its structure to bind peptides in different conformations to assert a sequence preference for aromatic residues. *J. Mol. Biol*. 373, 367-381. |
| Pnuc_1961 | LS: Lipoyl synthase | Cofactor lipoic acid biosynthetic enzyme. Lipoic acid has been reported as a key contributor in defending against oxidative stress in bacteria. | Bryk, R., Lima, C. D., Erdjument-Bromage, H., Tempst, P., and Nathan, C. (2002). Metabolic enzymes of mycobacteria linked to antioxidant defense by a thioredoxin-like protein. *Science*. 295, 1073-1077. |

| | | | |
|---|---|---|---|
| Pnuc_2021 | MreB: Rod shape-determining protein | Cytoskeletal protein that determines bacterial cell shape. MreB was upregulated during cold-shock treatment of *Vibrio parahaemolyticus*. MreB-ADP bound state may save ATP for cellular activities. | Chiu, S. W., Chen, S. Y., and Wong, H. C. (2008). Localization and expression of MreB in *Vibrio parahaemolyticus* under different stresses. *Appl. Environ. Microbiol*. 74, 7016-7022. |
| Pnuc_2030 | CA: Carbonate dehydratase | This gene encodes for a cytosolic enzyme that catalyzes the reversible hydration of carbon dioxide. | Smith, K. S., and Ferry, J. G. (2000). Prokaryotic carbonic anhydrases. *FEMS Microbiol. Rev*. 24, 335-366. |

**26ºC incubated sample**

| Locus Tag | Abbreviation / Gene product | Function | References |
|---|---|---|---|
| Pnuc_0161 | FtsI: Peptidoglycan synthetase | Cell wall biosynthesis and cell division proteins. | Typas, A., Banzhaf, M., Gross, C. A., and Vollmer, W. (2012). From the regulationof peptidoglycan synthesis to bacterial growth and morphology. *Nat. Rev. Microbiol*. 10, 123-136. O'Donnell, M., Langston, L., and Stillman, B. (2013). Principles and concepts of DNA replication in bacteria, archaea, and eukarya. *Cold Spring Harb. Perspect. Biol*. 5,a010108. |
| Pnuc_0163 | MurF: UDP-N-acetylmuramoy lalanyl-D-glutamyl-2, 6-diaminopimelate--D-alanyl-D-alanyl ligase | | |
| Pnuc_0164 | MraY: PhosphoN-acetylmuramoyl-pentapeptide-transferase | | |
| Pnuc_0166 | FtsW: Cell cycle protein | | |
| Pnuc_0168 | MurC: UDP-N-acetylmuramate--L-alanine ligase | | |
| Pnuc_0171 | FtsA: Cell division protein | | |
| Pnuc_0172 | FtsZ: Cell division protein | | |
| Pnuc_0173 | Ahp: Alkyl hydroperoxide reductase (peroxiredoxin) | Cytosolic protein that can sense peroxide and protect other proteins by acting as an anti-oxidant. | Dubbs, J. M., and Mongkolsuk, S. (2007). Peroxiredoxins in bacterial antioxidant defense. *Subcell. Biochem*. 44, 143-93. |

| | | | |
|---|---|---|---|
| Pnuc_0289 | MurB: UDP-N-acetylenolpyruvoyl glucosamine reductase | Cell wall biosynthesis | Typas, A., Banzhaf, M., Gross, C. A., and Vollmer, W. (2012). From the regulationof peptidoglycan synthesis to bacterial growth and morphology. *Nat. Rev. Microbiol*. 10, 123-136. |
| Pnuc_0292 | XerD: Recombinase | Catalyzes recombination event and keeps DNA molecule in monomeric state. | Wang, X., Llopis, P. M., and Rudner, D. Z. (2013). Organization and segregation of bacterial chromosomes. *Nat. Rev. Genet*. 14, 191-203. |
| Pnuc_0403 | Do: Protease | Periplasmic serine protease that degrades cellular proteins and is important in cell survival at elevated temperature. | Seol, J. H., Woo, S. K., Jung, E. M., Yoo, S. J., Lee, C.S., Kim, K. J., et al. (1991). Protease Do is essential for survival of *Escherichia coli* at high temperatures: its identity with the htrA gene product. *Biochem. Biophys. Res. Commun*. 176, 730–736. |
| Pnuc_0404 | EF4: GTP-binding protein LepA | EF4 can back-translocate the ribosome during stalling of protein synthesis under stress scenario. | Starosta, A. L., Lassak, J., Jung, K., and Wilson, D. N. (2014). The bacterial translation stress response. *FEMS Microbiol. Rev*. 38, 1172-1201. |
| Pnuc_0424 | DnaB: Helicase | Catalyzes the opening and propagation of replication fork. | Biswas, E. E., and Biswas, S. B. (1999). Mechanism of DnaB helicase of *Escherichia coli*: structural domains involved in ATP hydrolysis, DNA binding, and oligomerization. *Biochemistry*. 38, 10919-10928. |
| Pnuc_0429 | Ahp: Alkyl hydroperoxide reductase (peroxiredoxin) | Cytosolic enzyme that can detoxify various peroxides. | Parsonage, D., Karplus, P. A., and Poole, L. B. (2008). Substrate specificity and redox potential of AhpC, a bacterial peroxiredoxin. *Proc. Natl. Acad. Sci. USA*. 105, 8209-8214. |
| Pnuc_0509 | MltB: Lytic murein trans-glycosylase B | Cell wall recycling protein. | Suvorov, M., Lee, M., Hesek, D., Boggess, B., and Mobashery, S. (2008). Lytic transglycosylase MltB of *Escherichia coli* and its role in recycling of peptidoglycan strands of bacterial cell wall. *J. Am. Chem Soc*. 130, 11878-11879. |
| Pnuc_0528 | DctP: TRAP dicarboxylate transporter subunit | Tripartite ATP-independent periplasmic carrier that transports C4- dicarboxylates, e.g. malate, | Mulligan, C., Fischer, M., and Thomas, G. H. (2011). Tripartite ATP-independent periplasmic (TRAP) transporters in bacteria |

| | | | |
|---|---|---|---|
| | | succinate and fumarate. | and archaea. *FEMS Microbiol. Rev*. 35, 68-86. |
| Pnuc_0624 | PolIII: DNA polymerase III chi subunit, HolC | DNA replication. | Witte, G., Urbanke, C., and Curth, U. (2003). DNA polymerase III chi subunit ties single-stranded DNA binding protein to the bacterial replication machinery. *Nucleic Acids Res*. 31, 4434-4440 |
| Pnuc_0645 | ScpA: Chromosome segregation and condensation protein | Interacts with SMC and participates in the partitioning of chromosome during cell division. Mutation in *scpA* gene shows temperature-sensitive growth in *Bacillus subtilis*. | Soppa, J., Kobayashi, K., Noirot-Gros, M. F., Oesterhelt, D., Ehrlich, S. D., Dervyn, E., et al. (2002). Discovery of two novel families of proteins that are proposed to interact with prokaryotic SMC proteins, and characterization of the *Bacillus subtilis* family members ScpA and ScpB. *Mol. Microbiol*. 45, 59-71. |
| Pnuc_0690 | FtsK: DNA translocase | Motor protein located at the cell division septum and moves double strand DNA towards recombination site. | Crozat, E., and Grainge, I. (2010). FtsK DNA translocase: the fast motor that knows where it's going. *Chembiochem*. 11, 2232-2243 |
| Pnuc_0747 | DsbA: DSBA oxidoreductase | Periplasmic enzyme that catalyzes the disulfide bond formation and helps in correct protein folding. | Paxman, J. J., Borg, N. A., Horne, J., Thompson, P. E., Chin, Y., Sharma, P., et al. (2009). The structure of the bacterial oxidoreductase enzyme DsbA in complex with a peptide reveals a basis for substrate specificity in the catalytic cycle of DsbA enzymes. *J. Biol. Chem*. 284, 17835-17845. |
| Pnuc_0900<br>Pnuc_0901 | DNA topoisomerase IV subunit B<br>DNA topoisomerase IV subunit A | Disentangles the intertwined DNA after replication process, by creating double-stranded break. | Rawdon, E. J., Dorier, J., Racko, D., Millett, K. C., and Stasiak, A. (2016). How topoisomerase IV can efficiently unknot and decatenate negatively supercoiled DNA molecules without causing their torsional relaxation. *Nucleic Acids Res*. 44, 4528-4538 |
| Pnuc_1064 | σ32: RNA polymerase, sigma 32 subunit, RpoH | Sigma subunit of RNA polymerase required for the expression of genes during heat shock response. | Grainger, D. C., and Busby, S. J. (2008). Global regulators of transcription in *Escherichia coli*: mechanisms of action and methods for study. *Adv. Appl. Microbiol*. 65, 93-113. |

| | | | |
|---|---|---|---|
| Pnuc_1080 | Ribonuclease PH | Exoribonuclease that participates in the maturation of 3' side of tRNA by removing -CCA sequence. | Kelly, K. O, and Deutscher, M. P. (1992). Characterization of *Eschericia coli* RNase PH. *J. Biol. Chem*. 267, 17153-17158. |
| Pnuc_1344 | AhpC: alkyl hydroperoxide reductase (peroxiredoxin) | Periplasm residing and H2O2 scavenging enzyme. | LeBlanc, J. J., Davidson, R. J., and Hoffman, P. S. (2006). Compensatory functions of two alkyl hydroperoxide reductases in the oxidative defense system of *Legionella pneumophila*. *J. Bacteriol*. 188, 6235-6244. |
| Pnuc_1376 | Rbr: Rubrerythrin | Cytosolic enzyme that is activated during oxidative stress. Converts harmful peroxide into water. | Lehmann, Y., Meile, L., and Teuber, M. (1996). Rubrerythrin from *Clostridium perfringens*: cloning of the gene, purification of the protein, and characterization of its superoxide dismutase function. *J. Bacteriol*. 178, 7152-7158. |
| Pnuc_1385 | SOD: Manganese and iron superoxide dismutase | Cytosolic enzyme that catalyzes the conversion of superoxide radicals into oxygen. | Miller, A-F. (2012). Superoxide dismutases: Ancient enzymes and new insights. *FEBS Lett*. 586, 585-595. |
| Pnuc_1415 | LdcA: Murein tetrapeptidase LD-carboxypeptidase | Peptidoglycan recycling enzyme. Catabolize GlcNAc-MurNAc(anhydro)-tetrapeptide to tripeptide.* | Das, D., Hervé, M., Elsliger, M. A., Kadam, R. U., Grant, J. C., Chiu, H. J., et al. (2013). Structure and function of a novel LD-carboxypeptidase a involved in peptidoglycan recycling. *J. Bacteriol*. 195, 5555-5566. |
| Pnuc_1457 | SMC: Structural maintenance of chromosomes protein | Protein involved in compacting replicated DNA. | Graumann, P. L. (2001). SMC proteins in bacteria: condensation motors for chromosome segregation? *Biochimie*. 83, 53-59. |
| Pnuc_1534 | Ahp: Alkyl hydroperoxide reductase | Cytosolic enzyme that can reduce organic peroxides. | Poole, L. B., and Ellis, H. R. (1996). Flavin-dependent alkyl hydroperoxide reductase from *Salmonella typhimurium*. 1. purification and enzymatic activities of overexpressed AhpF and AhpC proteins. *Biochemistry*. 35, 56-64. |
| Pnuc_1699 | DnaG: DNA primase | Synthesizes the RNA primer that is used by DNA polymerase during replication process. | Rowen, L., and Kornberg, A. (1978). Primase, the *dnaG* protein of *Escherichia coli*. An enzyme which starts DNA chains. *J. Biol. Chem*. 253, 758-764. |

| | | | |
|---|---|---|---|
| Pnuc_1711 | PolI: DNA polymerase I | Prokaryotic DNA replication enzyme (polymerase, proofreading and nick translation properties). | Patel, P. H., Suzuki, M., Adman, E., Shinkai, A., and Loeb, L. A. (2001). Prokaryotic DNA polymerase I: evolution, structure, and "base flipping" mechanism for nucleotide selection. *J. Mol. Biol.* 308, 823-837. |
| Pnuc_1757 | MurJ: Lipid II flippase | Peptidoglycan biosynthesis. | Sham, L. T., Butler, E. K., Lebar, M. D., Kahne, D., Bernhardt, T. G., and Ruiz, N. (2014). MurJ is the flippase of lipid-linked precursors for peptidoglycan biogenesis. *Science*. 345, 220-222. |
| Pnuc_1787 | RecG: ATP-dependent DNA helicase | Promotes double-strand break repair and branch migration in homologous recombination event. | Toseland, C. P., Powell, B., and Webb, M. R. (2012). ATPase cycle and DNA unwinding kinetics of RecG helicase. *PLoS One*. 7, e38270. |
| Pnuc_1880 | RuvB: Holliday junction DNA (ruv mutant/recombinant sensitive to UV) | Promotes branch migration of the Holliday junction during genetic recombination event. | Smith, G. R. (1988). Homologous recombination in procaryotes. *Microbiol. Rev*. 52, 1-28. |
| Pnuc_1929 | σ24: RNA polymerase, sigma-24 subunit, RpoE | Sigma subunit of RNA polymerase required for the expression of genes during heat shock response and in response to misfolded proteins. | Grainger, D. C., and Busby, S. J. (2008). Global regulators of transcription in *Escherichia coli*: mechanisms of action and methods for study. *Adv. Appl. Microbiol*. 65, 93-113. |
| Pnuc_2021 | MreB: Rod shape-determining protein | | |
| Pnuc_2022 | MreC: Cell shape-determining protein | Cell shape maintenance proteins. | Typas, A., Banzhaf, M., Gross, C. A., and Vollmer, W. (2012). From the regulationof peptidoglycan synthesis to bacterial growth and morphology. *Nat. Rev. Microbiol*. 10, 123-136. |
| Pnuc_2024 | MrdA: Cell elongation-specific peptidoglycan D,D-transpeptidase | | |
| Pnuc_R0042 | RNase P | Endonucleolytic ribozyme that participates in the maturation of 5' side of tRNA. | Kirsebom, L. A. (2002). RNase P RNA-mediated catalysis. *Biochem. Soc. Trans*. 30, 1153-1158. |

**Supplementary Table S5.** Tabulation and explanation of abbreviations used in Fig. 5. Transcriptome and proteome data were used to prepare Fig. 5.

Upregulated genes or proteins in black font. Red font depicts downregulated genes or proteins.
*MurNAc: N- acetylmuramic acid, GlcNAc: N-acetylglucosamine.

### Supplementary Table S6.

| 4°C | | | | | |
|---|---|---|---|---|---|
| **Methylation** | **Location** | **Strand** | **Motif** | **Gene / Intergenic region** | **Count** |
| m6A | 1453 | + | | Pnuc_0001, chromosomal replication initiator protein DnaA | 652(+) |
| m6A | 2464 | - | | upstream Pnuc_2088, LSU ribosomal protein L34P | 621(-) |
| m6A | 7722 | + | | Pnuc_0008, amino acid/amide ABC transporter substrate-binding protein, HAAT family SIG | **Total** = 1273 (m6A + m4C) |
| m6A | 10210 | + | | Pnuc_0010, amino acid/amide ABC transporter membrane protein 2, HAAT family | |
| m4C | 10734 | + | | upstream Pnuc_0014, protein of unknown function DUF81 | 466(+) |
| m4C | 13299 | + | | upstream Pnuc_0014, protein of unknown function DUF81 | 433(-) |
| m4C | 13306 | - | | Pnuc_0013, acyl-CoA dehydrogenase domain protein | **Total** = 899 (m6A) |
| m6A | 13367 | - | | Pnuc_0013, acyl-CoA dehydrogenase domain protein | |
| m6A | 13470 | - | | Pnuc_0013, acyl-CoA dehydrogenase domain protein | 186(+) |
| m6A | 14643 | + | | upstream Pnuc_0015, glucose inhibited division protein A | 188(-) |
| m6A | 17573 | + | | Pnuc_0017, chromosome segregation ATPase | **Total** = 374 (m4C) |
| m6A | 18130 | + | | Pnuc_0018, chromosome segregation DNA-binding protein | |
| m4C | 18360 | - | | upstream Pnuc_0013, acyl-CoA dehydrogenase domain protein | Methylated genes = 438 (20.97%) |
| m4C | 18392 | + | | Pnuc_0018, chromosome segregation DNA-binding protein | |
| m6A | 18941 | + | | Pnuc_0018, chromosome segregation DNA-binding protein | |
| m6A | 19246 | + | | Pnuc_0019, putative ATP synthase protein I AtpI | |
| m6A | 20464 | - | | upstream Pnuc_0013, acyl-CoA dehydrogenase domain protein | |
| m4C | 21005 | + | | Pnuc_0022, ATP synthase F0 subcomplex B subunit | |
| m4C | 21561 | - | | upstream Pnuc_0013, acyl-CoA dehydrogenase domain protein | |
| m4C | 23579 | + | | Pnuc_0025, ATP synthase F1 subcomplex gamma subunit | |
| m6A | 24745 | - | | upstream Pnuc_0013, acyl-CoA dehydrogenase domain protein | |
| m6A | 28723 | + | | Pnuc_0029, replication restart DNA helicase PriA | |
| m4C | 28814 | + | | Pnuc_0029, replication restart DNA helicase PriA | |
| m4C | 28879 | + | | Pnuc_0029, replication restart DNA helicase PriA | |
| m4C | 32583 | - | | upstream Pnuc_R0001, tRNA-Arg | |
| m4C | 32592 | - | | upstream Pnuc_R0001, tRNA-Arg | |
| m4C | 36190 | - | | Pnuc_0038, GCN5-related N-acetyltransferase | |
| m6A | 36462 | + | | upstream Pnuc_R0002, 16S ribosomal RNA | |
| m6A | 37807 | + | CTAYNNNNNNNTRTC | Pnuc_R0002, 16S ribosomal RNA | |
| m6A | 37817 | - | GAYANNNNNNNRTAG | upstream Pnuc_0038, GCN5-related N-acetyltransferase | |
| m4C | 38568 | - | | upstream Pnuc_0038, GCN5-related N-acetyltransferase | |
| m6A | 41947 | + | | Pnuc_R0006, 5S ribosomal RNA | |
| m6A | 46043 | + | | upstream Pnuc_0044, LSU ribosomal protein L10P | |
| m4C | 47540 | - | | upstream Pnuc_0038, GCN5-related N-acetyltransferase | |
| m4C | 48179 | - | | upstream Pnuc_0038, GCN5-related N-acetyltransferase | |
| m4C | 49107 | - | | upstream Pnuc_0038, GCN5-related N-acetyltransferase | |
| m6A | 50868 | + | | Pnuc_0046, DNA-directed RNA polymerase subunit beta | |
| m6A | 52629 | - | | upstream Pnuc_0038, GCN5-related N-acetyltransferase | |
| m4C | 52724 | + | | Pnuc_0047, DNA-directed RNA polymerase subunit beta | |

| | | | | |
|---|---|---|---|---|
| m4C | 54355 | + | | Pnuc_0047, DNA-directed RNA polymerase subunit beta |
| m6A | 54908 | + | | Pnuc_0047, DNA-directed RNA polymerase subunit beta |
| m4C | 54915 | - | | upstream Pnuc_0038, GCN5-related N-acetyltransferase |
| m4C | 54966 | - | | upstream Pnuc_0038, GCN5-related N-acetyltransferase |
| m6A | 55842 | + | | upstream Pnuc_0048, SSU ribosomal protein S12P |
| m6A | 56187 | + | | Pnuc_0048, SSU ribosomal protein S12P |
| m6A | 57571 | + | | Pnuc_0050, translation elongation factor 2 (EF-2/EF-G) |
| m4C | 57861 | - | | upstream Pnuc_0038, GCN5-related N-acetyltransferase |
| m4C | 65434 | + | | Pnuc_0061, LSU ribosomal protein L29P |
| m4C | 68155 | - | | upstream Pnuc_0038, GCN5-related N-acetyltransferase |
| m6A | 68815 | + | | Pnuc_0069, LSU ribosomal protein L18P |
| m4C | 71592 | - | | upstream Pnuc_0038, GCN5-related N-acetyltransferase |
| m4C | 74844 | + | | upstream Pnuc_0080, CutA1 divalent ion tolerance protein |
| m6A | 75131 | - | | upstream Pnuc_0038, GCN5-related N-acetyltransferase |
| m4C | 76577 | - | | upstream Pnuc_0038, GCN5-related N-acetyltransferase |
| m4C | 77241 | - | | upstream Pnuc_0038, GCN5-related N-acetyltransferase |
| m6A | 79279 | - | | upstream Pnuc_0084, cell division checkpoint GTPase YihA |
| m4C | 82329 | - | | Pnuc_0088, 3-dehydroquinate synthase |
| m6A | 83003 | + | GAYANNNNNNNRTAG | upstream Pnuc_0092, penicillin-binding protein, 1A family |
| m6A | 83013 | - | CTAYNNNNNNNTRTC | Pnuc_0088, 3-dehydroquinate synthase |
| m6A | 84386 | + | CTAYNNNNNNNTRTC | upstream Pnuc_0092, penicillin-binding protein, 1A family |
| m6A | 84396 | - | GAYANNNNNNNRTAG | Pnuc_0090, type II and III secretion system protein |
| m6A | 84996 | - | | Pnuc_0091, hypothetical protein |
| m4C | 86659 | - | | upstream Pnuc_0091, hypothetical protein |
| m4C | 87028 | + | | Pnuc_0092, penicillin-binding protein, 1A family |
| m4C | 87343 | + | | Pnuc_0092, penicillin-binding protein, 1A family |
| m6A | 87781 | - | | upstream Pnuc_0091, hypothetical protein |
| m6A | 93455 | - | | upstream Pnuc_0091, hypothetical protein |
| m4C | 95506 | - | | upstream Pnuc_0091, hypothetical protein |
| m4C | 97538 | + | | Pnuc_0099, VacJ family lipoprotein |
| m6A | 97671 | - | | upstream Pnuc_0091, hypothetical protein |
| m4C | 97977 | - | | upstream Pnuc_0091, hypothetical protein |
| m4C | 99225 | - | | upstream Pnuc_0091, hypothetical protein |
| m6A | 101158 | - | | upstream Pnuc_0091, hypothetical protein |
| m6A | 102779 | + | GAYANNNNNNNRTAG | Pnuc_0107, histidinol dehydrogenase |
| m6A | 102789 | - | CTAYNNNNNNNTRTC | upstream Pnuc_0091, hypothetical protein |
| m6A | 103179 | + | CTAYNNNNNNNTRTC | Pnuc_0107, histidinol dehydrogenase |
| m6A | 103189 | - | GAYANNNNNNNRTAG | upstream Pnuc_0091, hypothetical protein |
| m6A | 106752 | + | | Pnuc_0111, 1-(5-phosphoribosyl)-5-[(5-phosphoribosylamino)methylideneamino]imidazole-4-carboxamide isomerase |
| m6A | 108116 | - | | upstream Pnuc_0108, histidinol phosphate aminotransferase apoenzyme |
| m6A | 111954 | - | | Pnuc_0119, pseudo |
| m4C | 112192 | - | | Pnuc_0119, pseudo |
| m6A | 112314 | + | | upstream Pnuc_0120, porin, Gram-negative type |
| m6A | 113863 | - | | upstream Pnuc_0119, pseudo |

| | | | | |
|---|---|---|---|---|
| m4C | 114745 | + | | Pnuc_0121, TPR repeat-containing protein |
| m6A | 115995 | - | | upstream Pnuc_0119, pseudo |
| m6A | 117881 | - | | Pnuc_0122, Tetratricopeptide TPR_2 repeat protein |
| m4C | 117995 | - | | Pnuc_0122, Tetratricopeptide TPR_2 repeat protein |
| m6A | 118329 | - | | Pnuc_0123, peptidase S1 and S6, chymotrypsin/Hap |
| m6A | 119609 | + | | upstream Pnuc_0125, ubiquinol-cytochrome c reductase, iron-sulfur subunit |
| m4C | 119836 | + | | upstream Pnuc_0125, ubiquinol-cytochrome c reductase, iron-sulfur subunit |
| m4C | 119892 | - | | upstream Pnuc_0124, large conductance mechanosensitive channel protein |
| m6A | 119969 | + | | Pnuc_0125, ubiquinol-cytochrome c reductase, iron-sulfur subunit |
| m6A | 122705 | + | CTAYNNNNNNNNTRTC | Pnuc_0127, cytochrome c1 |
| m6A | 122715 | - | GAYANNNNNNNNRTAG | upstream Pnuc_0124, large conductance mechanosensitive channel protein |
| m6A | 123799 | + | | Pnuc_0129, Stringent starvation protein B |
| m6A | 130121 | + | GAYANNNNNNNNRTAG | upstream Pnuc_0142, GTP-binding protein YchF |
| m6A | 130131 | - | CTAYNNNNNNNNTRTC | Pnuc_0136, bacterial peptide chain release factor 1(bRF-1) |
| m4C | 131167 | - | | Pnuc_0137, glutamyl-tRNA reductase |
| m4C | 135156 | + | | upstream Pnuc_0142, GTP-binding protein YchF |
| m6A | 145576 | - | | Pnuc_0152, amidohydrolase |
| m6A | 145707 | + | | upstream Pnuc_0156, biotin synthase |
| m6A | 147876 | + | | upstream Pnuc_0156, biotin synthase |
| m6A | 149067 | + | CTAYNNNNNNNNTRTC | upstream Pnuc_0156, biotin synthase |
| m4C | 149837 | + | | upstream Pnuc_0156, biotin synthase |
| m4C | 154466 | - | | upstream Pnuc_0155, aminotransferase |
| m6A | 157993 | + | GAYANNNNNNNNRTAG | Pnuc_0163, UDP-N-acetylmuramoylalanyl-D-glutamyl-2,6-diaminopimelate-D-alanyl-D-alanyl ligase |
| m6A | 158003 | - | CTAYNNNNNNNNTRTC | upstream Pnuc_0155, aminotransferase |
| m6A | 159967 | + | | Pnuc_0165, UDP-N-acetylmuramoylalanine--D-glutamate ligase |
| m6A | 162336 | - | | upstream Pnuc_0155, aminotransferase |
| m4C | 164580 | - | | upstream Pnuc_0155, aminotransferase |
| m6A | 167650 | + | | Pnuc_0171, cell division protein FtsA |
| m4C | 168631 | + | | Pnuc_0172, cell division protein FtsZ |
| m6A | 170853 | + | | Pnuc_0174, UDP-3-O-[3-hydroxymyristoyl] N-acetylglucosamine deacetylase |
| m4C | 170901 | + | | Pnuc_0174, UDP-3-O-[3-hydroxymyristoyl] N-acetylglucosamine deacetylase |
| m6A | 171824 | + | | Pnuc_0176, protein translocase subunit secA |
| m6A | 175537 | + | | Pnuc_0178, aldehyde oxidase and xanthine dehydrogenase,molybdopterin binding protein |
| m4C | 179581 | - | | upstream Pnuc_0175, hypothetical protein |
| m6A | 180449 | - | | Pnuc_0183, dephospho-CoA kinase |
| m4C | 180979 | + | | upstream Pnuc_0188, methylated-DNA--protein-cysteine methyltransferase |
| m6A | 183387 | + | | upstream Pnuc_0188, methylated-DNA--protein-cysteine methyltransferase |
| m6A | 190277 | + | | upstream Pnuc_0194, LSU ribosomal protein L21P |
| m4C | 190717 | + | | upstream Pnuc_0194, LSU ribosomal protein L21P |
| m4C | 192640 | - | | upstream Pnuc_0193, Farnesyltranstransferase |
| m6A | 196749 | - | | upstream Pnuc_0199, NUDIX hydrolase, downstream Pnuc_0201, signal recognition particle subunit FFH/SRP54 (srp54) |
| m6A | 196752 | + | | upstream Pnuc_0202, cytochrome c assembly protein |
| m6A | 197824 | + | | upstream Pnuc_0202, cytochrome c assembly protein |
| m6A | 200762 | + | CTAYNNNNNNNNTRTC | Pnuc_0204, ribonucleoside-diphosphate reductase, alpha subunit |

| | | | | |
|---|---|---|---|---|
| m6A | 200772 | - | GAYANNNNNNNRTAG | upstream Pnuc_0201, signal recognition particle subunit FFH/SRP54 (srp54) |
| m6A | 200911 | - | | upstream Pnuc_0201, signal recognition particle subunit FFH/SRP54 (srp54) |
| m6A | 201518 | - | | upstream Pnuc_0201, signal recognition particle subunit FFH/SRP54 (srp54) |
| m4C | 202691 | - | | upstream Pnuc_0201, signal recognition particle subunit FFH/SRP54 (srp54) |
| m6A | 202867 | - | | upstream Pnuc_0201, signal recognition particle subunit FFH/SRP54 (srp54) |
| m6A | 204236 | + | | Pnuc_0205, Ribonucleoside-diphosphate reductase |
| m6A | 204565 | + | | upstream Pnuc_0206, hypothetical protein |
| m6A | 204937 | + | | Pnuc_0206, hypothetical protein |
| m6A | 207217 | - | | Pnuc_0208, [LSU ribosomal protein L11P]-lysine N-methyltransferase |
| m6A | 209668 | + | GAYANNNNNNNRTAG | upstream Pnuc_0213, UDP-N-acetylmuramate |
| m6A | 209678 | - | CTAYNNNNNNNTRTC | Pnuc_0211, 3-dehydroquinate dehydratase |
| m6A | 210167 | + | | upstream Pnuc_0213, UDP-N-acetylmuramate |
| m6A | 210170 | + | | upstream Pnuc_0213, UDP-N-acetylmuramate |
| m4C | 210233 | - | | upstream Pnuc_0211, 3-dehydroquinate dehydratase |
| m4C | 210234 | - | | upstream Pnuc_0211, 3-dehydroquinate dehydratase |
| m6A | 210241 | + | | upstream Pnuc_0213, UDP-N-acetylmuramate |
| m6A | 210710 | - | | Pnuc_0212, alkyl hydroperoxide reductase/ Thiol specific antioxidant/ Mal allergen |
| m6A | 213423 | + | | Pnuc_0215, ribonuclease II |
| m6A | 215142 | - | | upstream Pnuc_0212, alkyl hydroperoxide reductase/ Thiol specific antioxidant/ Mal allergen |
| m6A | 215145 | - | | upstream Pnuc_0212, alkyl hydroperoxide reductase/ Thiol specific antioxidant/ Mal allergen |
| m6A | 215821 | - | | upstream Pnuc_0212, alkyl hydroperoxide reductase/ Thiol specific antioxidant/ Mal allergen |
| m4C | 216905 | - | | upstream Pnuc_0212, alkyl hydroperoxide reductase/ Thiol specific antioxidant/ Mal allergen |
| m4C | 218122 | + | | upstream Pnuc_0220, magnesium and cobalt transport protein CorA |
| m6A | 220559 | + | | upstream Pnuc_0224, Malate dehydrogenase (oxaloacetate-decarboxylating) (NADP(+))., Phosphate acetyltransferase |
| m6A | 220716 | - | | Pnuc_0223, thiamine-phosphate kinase |
| m6A | 220989 | - | | Pnuc_0223, thiamine-phosphate kinase |
| m6A | 224087 | - | | upstream Pnuc_0223, thiamine-phosphate kinase |
| m6A | 227606 | + | | Pnuc_0228, glyceraldehyde-3-phosphate dehydrogenase |
| m6A | 227673 | + | GAYANNNNNNNRTAG | Pnuc_0228, glyceraldehyde-3-phosphate dehydrogenase |
| m6A | 227683 | - | CTAYNNNNNNNTRTC | upstream Pnuc_0226, protein of unknown function DUF558 |
| m6A | 227895 | + | | Pnuc_0228, glyceraldehyde-3-phosphate dehydrogenase |
| m6A | 228520 | + | | Pnuc_0228, glyceraldehyde-3-phosphate dehydrogenase |
| m6A | 228630 | - | | upstream Pnuc_0226, protein of unknown function DUF558 |
| m4C | 228649 | - | | upstream Pnuc_0226, protein of unknown function DUF558 |
| m4C | 231697 | + | | Pnuc_0232, leucyl-tRNA synthetase |
| m6A | 231802 | + | | Pnuc_0232, leucyl-tRNA synthetase |
| m4C | 232073 | - | | upstream Pnuc_0229, ferric uptake regulator, Fur family |
| m6A | 238728 | + | | Pnuc_0239, glutamate-1-semialdehyde 2,1-aminomutase |
| m4C | 238745 | - | | upstream Pnuc_0229, ferric uptake regulator, Fur family |
| m4C | 240934 | + | | upstream Pnuc_0242, protein of unknown function DUF218 |
| m6A | 242771 | + | | Pnuc_0242, protein of unknown function DUF218 |
| m4C | 245419 | - | | upstream Pnuc_0241, deoxyribodipyrimidine photo-lyase type I |
| m6A | 248429 | + | CTAYNNNNNNNTRTC | Pnuc_0249, glycosyl transferase, family 25 |
| m6A | 248439 | - | GAYANNNNNNNRTAG | upstream Pnuc_0241, deoxyribodipyrimidine photo-lyase type I |

| | | | | |
|---|---|---|---|---|
| m4C | 250123 | - | | Pnuc_0251, hypothetical protein |
| m6A | 250899 | + | CTAYNNNNNNNNTRTC | Pnuc_0252, hypothetical protein |
| m6A | 250909 | - | GAYANNNNNNNNRTAG | upstream Pnuc_0251, hypothetical protein |
| m6A | 251656 | + | GAYANNNNNNNNRTAG | upstream Pnuc_0253, dTDP-glucose 4,6-dehydratase |
| m6A | 251666 | - | CTAYNNNNNNNNTRTC | upstream Pnuc_0251, hypothetical protein |
| m6A | 263341 | + | | upstream Pnuc_0267, hypothetical protein |
| m4C | 263605 | + | | upstream Pnuc_0267, hypothetical protein |
| m4C | 265118 | - | | upstream Pnuc_0266, 3,4-dihydroxy-2-butanone 4-phosphate synthase |
| m6A | 269370 | + | GAYANNNNNNNNRTAG | upstream Pnuc_0279, serine hydroxymethyltransferase |
| m6A | 269380 | - | CTAYNNNNNNNNTRTC | Pnuc_0272 (start codon), preQ(0) biosynthesis protein QueC |
| m4C | 270928 | - | | Pnuc_0275, TolB, N-terminal domain protein |
| m6A | 271013 | + | GAYANNNNNNNNRTAG | upstream Pnuc_0279, serine hydroxymethyltransferase |
| m6A | 271023 | - | CTAYNNNNNNNNTRTC | Pnuc_0275, TolB, N-terminal domain protein |
| m6A | 271901 | + | | upstream Pnuc_0279, serine hydroxymethyltransferase |
| m6A | 271994 | + | | upstream Pnuc_0279, serine hydroxymethyltransferase |
| m6A | 272665 | - | | Pnuc_0276, Cell division and transport-associated protein TolA |
| m6A | 272704 | - | | Pnuc_0276, Cell division and transport-associated protein TolA |
| m6A | 276373 | + | | upstream Pnuc_0287, Exodeoxyribonuclease VII large subunit |
| m6A | 277009 | + | CTAYNNNNNNNNTRTC | upstream Pnuc_0287, Exodeoxyribonuclease VII large subunit |
| m6A | 277019 | - | GAYANNNNNNNNRTAG | Pnuc_0282, 3-deoxy-D-manno-octulosonate cytidylyltransferase |
| m4C | 278708 | + | | upstream Pnuc_0287, Exodeoxyribonuclease VII large subunit |
| m6A | 283189 | + | GAYANNNNNNNNRTAG | upstream Pnuc_0290, protein of unknown function DUF520 |
| m6A | 283199 | - | CTAYNNNNNNNNTRTC | Pnuc_0289, UDP-N-acetylmuramate dehydrogenase |
| m6A | 287569 | + | | upstream Pnuc_0296, High potential iron-sulfur protein |
| m6A | 287750 | - | | upstream Pnuc_0289, UDP-N-acetylmuramate dehydrogenase |
| m6A | 297004 | + | | Pnuc_0306, glycosyl transferase, family 2 |
| m4C | 299064 | + | | upstream Pnuc_0311, polysaccharide biosynthesis protein |
| m6A | 305645 | + | | upstream Pnuc_0314, transferase hexapeptide repeat containing protein |
| m6A | 310366 | + | CTAYNNNNNNNNTRTC | Pnuc_0318, ABC-2 type transporter |
| m6A | 310376 | - | GAYANNNNNNNNRTAG | upstream Pnuc_0316, glycosyl transferase, family 4 |
| m6A | 312933 | + | GAYANNNNNNNNRTAG | Pnuc_0320, Methyltransferase type 11 |
| m6A | 312943 | - | CTAYNNNNNNNNTRTC | upstream Pnuc_0316, glycosyl transferase, family 4 |
| m6A | 313175 | + | GAYANNNNNNNNRTAG | Pnuc_0320, Methyltransferase type 11 |
| m6A | 313185 | - | CTAYNNNNNNNNTRTC | upstream Pnuc_0316, glycosyl transferase, family 4 |
| m6A | 317235 | + | | Pnuc_0322, hypothetical protein |
| m6A | 318999 | + | CTAYNNNNNNNNTRTC | upstream Pnuc_0324, NAD-dependent epimerase/dehydratase |
| m6A | 319009 | - | GAYANNNNNNNNRTAG | upstream Pnuc_0316, glycosyl transferase, family 4 |
| m6A | 320925 | + | GAYANNNNNNNNRTAG | Pnuc_0325, amine oxidase |
| m6A | 320935 | - | CTAYNNNNNNNNTRTC | upstream Pnuc_0316, glycosyl transferase, family 4 |
| m6A | 321154 | + | CTAYNNNNNNNNTRTC | Pnuc_0325, amine oxidase |
| m6A | 321164 | - | GAYANNNNNNNNRTAG | upstream Pnuc_0316, glycosyl transferase, family 4 |
| m6A | 324292 | + | GAYANNNNNNNNRTAG | Pnuc_0329, UbiA prenyltransferase |
| m6A | 324302 | - | CTAYNNNNNNNNTRTC | upstream Pnuc_0328, FAD linked oxidase domain protein |
| m4C | 325168 | - | | upstream Pnuc_0328, FAD linked oxidase domain protein |

| | | | | |
|---|---|---|---|---|
| m6A | 331725 | + | | upstream Pnuc_0337, acyltransferase 3 |
| m4C | 331986 | + | | upstream Pnuc_0337, acyltransferase 3 |
| m6A | 335858 | + | CTAYNNNNNNNNTRTC | Pnuc_0338, hypothetical protein |
| m6A | 335868 | - | GAYANNNNNNNNRTAG | upstream Pnuc_0336, hypothetical protein |
| m6A | 345459 | + | GAYANNNNNNNNRTAG | upstream Pnuc_0351, acyltransferase 3 |
| m6A | 345469 | - | CTAYNNNNNNNNTRTC | upstream Pnuc_0349, hypothetical protein |
| m6A | 347772 | + | GAYANNNNNNNNRTAG | Pnuc_0353, Choline dehydrogenase |
| m6A | 347782 | - | CTAYNNNNNNNNTRTC | upstream Pnuc_0352, protein of unknown function DUF615 |
| m6A | 349550 | + | | Pnuc_0354, protein of unknown function DUF6, transmembrane |
| m4C | 351661 | - | | upstream Pnuc_0352, protein of unknown function DUF615 |
| m6A | 372082 | + | | Pnuc_0379, PpiC-type peptidyl-prolyl cis-trans isomerase |
| m6A | 374721 | + | | upstream Pnuc_0384, D-isomer specific 2-hydroxyacid dehydrogenase, NAD-binding protein |
| m6A | 375651 | + | | upstream Pnuc_0384, D-isomer specific 2-hydroxyacid dehydrogenase, NAD-binding protein |
| m4C | 376339 | - | | upstream Pnuc_0383, 4Fe-4S ferredoxin, iron-sulfur binding domain protein |
| m4C | 381037 | - | | Pnuc_0389, RNAse E |
| m6A | 381129 | - | | Pnuc_0389, RNAse E |
| m6A | 382455 | - | | Pnuc_0389, RNAse E |
| m6A | 382815 | + | | upstream Pnuc_0390, ribosomal large subunit pseudouridine synthase C |
| m6A | 383532 | + | | Pnuc_0390, ribosomal large subunit pseudouridine synthase C |
| m4C | 387658 | - | | upstream Pnuc_0394, maf protein |
| m4C | 387721 | + | | Pnuc_0395, protein of unknown function DUF177 |
| m4C | 387959 | - | | upstream Pnuc_0394, maf protein |
| m6A | 389530 | + | | Pnuc_0398, 3-oxoacyl-[acyl-carrier-protein] synthase III |
| m6A | 389684 | + | | Pnuc_0398, 3-oxoacyl-[acyl-carrier-protein] synthase III |
| m6A | 396752 | + | | Pnuc_0404, GTP-binding protein LepA |
| m4C | 396765 | + | | Pnuc_0404, GTP-binding protein LepA |
| m4C | 396769 | + | | Pnuc_0404, GTP-binding protein LepA |
| m4C | 396777 | + | | Pnuc_0404, GTP-binding protein LepA |
| m4C | 396778 | + | | Pnuc_0404, GTP-binding protein LepA |
| m6A | 397041 | + | CTAYNNNNNNNNTRTC | Pnuc_0404, GTP-binding protein LepA |
| m6A | 397051 | - | GAYANNNNNNNNRTAG | upstream Pnuc_0394, maf protein |
| m4C | 398437 | + | | Pnuc_0406, RNAse III |
| m6A | 398446 | + | | Pnuc_0406, RNAse III |
| m6A | 401086 | + | CTAYNNNNNNNNTRTC | Pnuc_0409, pyridoxine 5'-phosphate synthase |
| m6A | 401096 | - | GAYANNNNNNNNRTAG | upstream Pnuc_0394, maf protein |
| m6A | 401232 | + | GAYANNNNNNNNRTAG | Pnuc_0410, holo-acyl-carrier-protein synthase |
| m6A | 401242 | - | CTAYNNNNNNNNTRTC | upstream Pnuc_0394, maf protein |
| m4C | 402615 | - | | upstream Pnuc_0394, maf protein |
| m4C | 404201 | - | | upstream Pnuc_0413, translation elongation factor P (EF-P) |
| m4C | 407370 | - | | upstream Pnuc_0414, conserved hypothetical protein |
| m4C | 407402 | - | | upstream Pnuc_0414, conserved hypothetical protein |
| m6A | 407449 | - | | upstream Pnuc_0414, conserved hypothetical protein |
| m6A | 407712 | - | | upstream Pnuc_0414, conserved hypothetical protein |
| m6A | 415368 | - | | Pnuc_0426, MscS Mechanosensitive ion channel |

| | | | | |
|---|---|---|---|---|
| m6A | 418091 | + | | upstream Pnuc_0437, aminotransferase |
| m6A | 420341 | + | | upstream Pnuc_0437, aminotransferase |
| m6A | 421455 | + | GAYANNNNNNNRTAG | upstream Pnuc_0437, aminotransferase |
| m6A | 421465 | - | CTAYNNNNNNNTRTC | Pnuc_0432, formyl transferase domain protein |
| m6A | 422836 | + | CTAYNNNNNNNTRTC | upstream Pnuc_0437, aminotransferase |
| m6A | 422846 | - | GAYANNNNNNNRTAG | Pnuc_0434, DegT/DnrJ/EryC1/StrS aminotransferase |
| m6A | 424440 | + | | upstream Pnuc_0437, aminotransferase |
| m6A | 428162 | - | | upstream Pnuc_0436, protein of unknown function DUF498 |
| m4C | 433725 | + | | Pnuc_0443, Carboxylesterase, type B |
| m4C | 433790 | - | | upstream Pnuc_0436, protein of unknown function DUF498 |
| m4C | 436964 | + | | upstream Pnuc_0447, ATPase AAA-2 domain protein |
| m6A | 439040 | + | GAYANNNNNNNRTAG | Pnuc_0447, ATPase AAA-2 domain protein |
| m6A | 439050 | - | CTAYNNNNNNNTRTC | upstream Pnuc_0446, Excinuclease ABC, C subunit domain protein |
| m6A | 440012 | + | GAYANNNNNNNRTAG | Pnuc_0447, ATPase AAA-2 domain protein |
| m6A | 440022 | - | CTAYNNNNNNNTRTC | upstream Pnuc_0446, Excinuclease ABC, C subunit domain protein |
| m6A | 442569 | + | GAYANNNNNNNRTAG | Pnuc_0450, major facilitator superfamily MFS_1 |
| m6A | 442579 | - | CTAYNNNNNNNTRTC | upstream Pnuc_0449, Uncharacterized protein UPF0065 |
| m4C | 442618 | - | | upstream Pnuc_0449, Uncharacterized protein UPF0065 |
| m6A | 443389 | - | | upstream Pnuc_0449, Uncharacterized protein UPF0065 |
| m6A | 445114 | + | | Pnuc_0451, heavy metal translocating P-type ATPase |
| m6A | 445942 | + | CTAYNNNNNNNTRTC | upstream Pnuc_0453, cytochrome c oxidase, cbb3-type, subunit I |
| m6A | 445952 | - | GAYANNNNNNNRTAG | upstream Pnuc_0449, Uncharacterized protein UPF0065 |
| m6A | 445992 | - | | upstream Pnuc_0449, Uncharacterized protein UPF0065 |
| m4C | 447945 | - | | upstream Pnuc_0449, Uncharacterized protein UPF0065 |
| m6A | 450432 | + | | Pnuc_0457, 4Fe-4S ferredoxin, iron-sulfur binding domain protein |
| m6A | 452613 | - | | upstream Pnuc_0460, putative transcriptional regulator, Crp/Fnr family |
| m6A | 453512 | + | CTAYNNNNNNNTRTC | Pnuc_0463, UspA domain protein |
| m6A | 453522 | - | GAYANNNNNNNRTAG | upstream Pnuc_0462, conserved hypothetical protein |
| m6A | 456340 | + | | upstream Pnuc_0466, Mandelate racemase/muconate lactonizing enzyme, C-terminal domain protein |
| m6A | 456402 | - | | upstream Pnuc_0465, 3-hydroxyacyl-CoA dehydrogenase |
| m6A | 457988 | - | | Pnuc_0467, TRAP transporter, 4TM/12TM fusion protein |
| m6A | 460630 | - | | upstream Pnuc_0468, TRAP transporter solute receptor, TAXI family |
| m4C | 460892 | - | | Pnuc_0469, dihydrodipicolinate synthetase |
| m6A | 461835 | + | GAYANNNNNNNRTAG | upstream Pnuc_0471, 3-hydroxyacyl-CoA dehydrogenase |
| m6A | 461845 | - | CTAYNNNNNNNTRTC | Pnuc_0470, D-isomer specific 2-hydroxyacid dehydrogenase, NAD-binding protein |
| m6A | 462740 | - | | upstream Pnuc_0470, D-isomer specific 2-hydroxyacid dehydrogenase, NAD-binding protein |
| m4C | 463424 | - | | upstream Pnuc_0470, D-isomer specific 2-hydroxyacid dehydrogenase, NAD-binding protein |
| m6A | 465489 | - | | Pnuc_0474, cytochrome c, class II |
| m4C | 466672 | + | | upstream Pnuc_0476, Extradiol ring-cleavage dioxygenase, class III enzyme, subunit B |
| m4C | 467171 | + | | Pnuc_0476, Extradiol ring-cleavage dioxygenase, class III enzyme, subunit B |
| m6A | 474614 | + | CTAYNNNNNNNTRTC | upstream Pnuc_0485, Uncharacterized protein UPF0065 |
| m6A | 474624 | - | GAYANNNNNNNRTAG | upstream Pnuc_0484, Rhodanese domain protein |
| m4C | 478450 | + | | Pnuc_0489, TPR repeat-containing protein |
| m6A | 482125 | - | | Pnuc_0492, OmpA/MotB domain protein |

| | | | | |
|---|---|---|---|---|
| m4C | 483340 | - | | upstream Pnuc_0492, OmpA/MotB domain protein |
| m6A | 487192 | + | | Pnuc_0495, chorismate mutase |
| m4C | 487707 | - | | upstream Pnuc_0492, OmpA/MotB domain protein |
| m4C | 488286 | - | | upstream Pnuc_0492, OmpA/MotB domain protein |
| m4C | 489439 | - | | upstream Pnuc_0492, OmpA/MotB domain protein |
| m6A | 493196 | + | | Pnuc_0500, SSU ribosomal protein S1P |
| m6A | 495425 | - | | upstream Pnuc_0492, OmpA/MotB domain protein |
| m6A | 499450 | + | | upstream Pnuc_0507, helix-hairpin-helix motif protein |
| m4C | 499573 | + | | Pnuc_0507, helix-hairpin-helix motif protein |
| m4C | 499947 | - | | upstream Pnuc_0492, OmpA/MotB domain protein |
| m4C | 502073 | - | | Pnuc_0509, lytic murein transglycosylase B |
| m4C | 507808 | + | | Pnuc_0514, acyl-CoA dehydrogenase domain protein |
| m4C | 508502 | - | | upstream Pnuc_0509, lytic murein transglycosylase B |
| m6A | 508522 | - | | upstream Pnuc_0509, lytic murein transglycosylase B |
| m4C | 508567 | + | | upstream Pnuc_0517, SSU ribosomal protein S16P |
| m6A | 511656 | + | GAYANNNNNNNRTAG | Pnuc_0520, LSU ribosomal protein L19P |
| m6A | 511666 | - | CTAYNNNNNNNTRTC | upstream Pnuc_0516, protein of unknown function DUF306, Meta and HslJ |
| m6A | 512017 | + | | upstream Pnuc_0521, NUDIX hydrolase |
| m6A | 512339 | + | | Pnuc_0521, NUDIX hydrolase |
| m6A | 513793 | + | CTAYNNNNNNNTRTC | upstream Pnuc_0525, Exonuclease, RNase T and DNA polymerase III |
| m6A | 513803 | - | GAYANNNNNNNRTAG | Pnuc_0523, ribosome small subunit-dependent GTPase A |
| m6A | 515484 | + | | upstream Pnuc_0525, Exonuclease, RNase T and DNA polymerase III |
| m6A | 515507 | + | | upstream Pnuc_0525, Exonuclease, RNase T and DNA polymerase III |
| m4C | 515722 | + | | upstream Pnuc_0525, Exonuclease, RNase T and DNA polymerase III |
| m6A | 516835 | + | GAYANNNNNNNRTAG | upstream Pnuc_0527, microcin-processing peptidase 1, Unknown type peptidase, MEROPS family U62 |
| m6A | 516845 | - | CTAYNNNNNNNTRTC | Pnuc_0526, molybdopterin adenylyltransferase |
| m4C | 518061 | - | | upstream Pnuc_0526, molybdopterin adenylyltransferase |
| m6A | 529410 | + | GAYANNNNNNNRTAG | upstream Pnuc_0541, Glutathione S-transferase, N-terminal domain protein |
| m6A | 529420 | - | CTAYNNNNNNNTRTC | upstream Pnuc_0538, hypothetical protein |
| m6A | 539761 | - | | upstream Pnuc_0547, putative phytochelatin synthase |
| m6A | 540300 | - | | upstream upstream Pnuc_0547, putative phytochelatin synthase |
| m4C | 548217 | + | | Pnuc_0556, DoxX family protein |
| m6A | 553023 | + | CTAYNNNNNNNTRTC | upstream Pnuc_0565, hypothetical protein |
| m6A | 553033 | - | GAYANNNNNNNRTAG | Pnuc_0563, integral membrane sensor hybrid histidine kinase |
| m6A | 555985 | + | | Pnuc_0566, hypothetical protein |
| m6A | 556714 | + | | upstream Pnuc_0570, cation diffusion facilitator family transporter |
| m6A | 557691 | + | | upstream Pnuc_0570, cation diffusion facilitator family transporter |
| m4C | 558569 | - | | Pnuc_0569, GCN5-related N-acetyltransferase |
| m6A | 560162 | + | GAYANNNNNNNRTAG | Pnuc_0570, cation diffusion facilitator family transporter |
| m6A | 560172 | - | CTAYNNNNNNNTRTC | upstream Pnuc_0569, GCN5-related N-acetyltransferase |
| m6A | 561977 | - | | upstream Pnuc_0572, Methyltransferase type 11 |
| m6A | 563811 | - | | upstream Pnuc_0572, Methyltransferase type 12 |
| m6A | 563878 | + | | Pnuc_0574, pseudo |
| m4C | 570856 | + | | upstream Pnuc_0583, hypothetical protein |

| | | | | |
|---|---|---|---|---|
| m4C | 570932 | - | | Pnuc_0582, hypothetical protein |
| m4C | 570962 | + | | upstream Pnuc_0583, hypothetical protein |
| m6A | 571162 | + | | Pnuc_0583, hypothetical protein |
| m6A | 573053 | - | | upstream Pnuc_0582, hypothetical protein |
| m4C | 576206 | - | | Pnuc_0591, conserved hypothetical protein |
| m6A | 578571 | - | | Pnuc_0593, 2-keto-3-deoxygluconate permease |
| m6A | 587229 | + | GAYANNNNNNNNRTAG | upstream Pnuc_0602, Serine--glyoxylate transaminase |
| m6A | 587239 | - | CTAYNNNNNNNNTRTC | Pnuc_0600, FAD linked oxidase domain protein |
| m6A | 587602 | + | | upstream Pnuc_0602, Serine--glyoxylate transaminase |
| m6A | 588730 | - | | upstream Pnuc_0601, transcriptional regulator, GntR family |
| m6A | 590985 | + | | Pnuc_0604, DEAD/DEAH box helicase domain protein |
| m6A | 597730 | + | | upstream Pnuc_0610, lipid A ABC exporter, fused ATPase and inner membrane subunits MsbA |
| m6A | 598039 | + | | upstream Pnuc_0610, lipid A ABC exporter, fused ATPase and inner membrane subunits MsbA |
| m6A | 598315 | + | | upstream Pnuc_0610, lipid A ABC exporter, fused ATPase and inner membrane subunits MsbA |
| m6A | 598932 | + | GAYANNNNNNNNRTAG | upstream Pnuc_0610, lipid A ABC exporter, fused ATPase and inner membrane subunits MsbA |
| m6A | 598942 | - | CTAYNNNNNNNNTRTC | Pnuc_0609, glycosyl transferase, family 2 |
| m6A | 601334 | + | | upstream Pnuc_R0016, tRNA-Leu |
| m4C | 602184 | - | | Pnuc_0611, RNAse G |
| m6A | 606107 | - | | Pnuc_0617, phosphoribosylamine-glycine ligase |
| m6A | 606694 | + | | upstream Pnuc_R0016, tRNA-Leu |
| m6A | 607185 | + | | upstream Pnuc_R0016, tRNA-Leu |
| m6A | 607706 | + | | upstream Pnuc_R0016, tRNA-Leu |
| m6A | 610532 | - | | Pnuc_0621, permease YjgP/YjgQ family protein |
| m6A | 610775 | - | | Pnuc_0621, permease YjgP/YjgQ family protein |
| m4C | 612570 | - | | upstream Pnuc_0622, permease YjgP/YjgQ family protein |
| m6A | 613102 | + | | Pnuc_0623, aminopeptidase A, Metallo peptidase, MEROPS family M17 |
| m4C | 616738 | - | | upstream Pnuc_0622, permease YjgP/YjgQ family protein |
| m6A | 616740 | - | | upstream Pnuc_0622, permease YjgP/YjgQ family protein |
| m6A | 618648 | + | GAYANNNNNNNNRTAG | Pnuc_0628, TRAP dicarboxylate transporter- DctP subunit |
| m6A | 618658 | - | CTAYNNNNNNNNTRTC | upstream Pnuc_0622, permease YjgP/YjgQ family protein |
| m6A | 621941 | + | GAYANNNNNNNNRTAG | Pnuc_0631, AMP-dependent synthetase and ligase |
| m6A | 621951 | - | CTAYNNNNNNNNTRTC | upstream Pnuc_0622, permease YjgP/YjgQ family protein |
| m6A | 624339 | - | | Pnuc_0635, TonB-dependent receptor, plug |
| m6A | 626322 | - | | adenosylcobalamin binding moiety (RF00174) |
| m4C | 626581 | - | | upstream Pnuc_0635, TonB-dependent receptor, plug |
| m6A | 627186 | - | | upstream Pnuc_0635, TonB-dependent receptor, plug |
| m4C | 628238 | - | | upstream Pnuc_0635, TonB-dependent receptor, plug |
| m6A | 632692 | + | | Pnuc_0641, Methionyl-tRNA formyltransferase |
| m6A | 633870 | + | | Pnuc_0643, penicillin amidase, Cysteine peptidase, MEROPS family C59 |
| m4C | 638228 | - | | Pnuc_0647, methionyl-tRNA synthetase |
| m6A | 639054 | + | GAYANNNNNNNNRTAG | Pnuc_0648, formate dehydrogenase, subunit FdhD |
| m6A | 639064 | - | CTAYNNNNNNNNTRTC | upstream Pnuc_0647, methionyl-tRNA synthetase |
| m6A | 639443 | + | | Pnuc_0648, formate dehydrogenase, subunit FdhD |
| m6A | 640505 | - | | upstream Pnuc_0647, methionyl-tRNA synthetase |

| | | | | |
|---|---|---|---|---|
| m6A | 640962 | - | | upstream Pnuc_0647, methionyl-tRNA synthetase |
| m4C | 641546 | - | | upstream Pnuc_0647, methionyl-tRNA synthetase |
| m6A | 642290 | + | CTAYNNNNNNNNTRTC | Pnuc_0651, Lysine decarboxylase |
| m6A | 642300 | - | GAYANNNNNNNNRTAG | upstream Pnuc_0647, methionyl-tRNA synthetase |
| m4C | 648813 | + | | Pnuc_0657, Uroporphyrinogen III synthase HEM4 |
| m4C | 651326 | - | | Pnuc_0659, DEAD/DEAH box helicase domain protein |
| m4C | 653254 | + | | upstream Pnuc_0662, peptide methionine sulfoxide reductase |
| m6A | 653833 | + | | Pnuc_0662, peptide methionine sulfoxide reductase |
| m6A | 656037 | + | | upstream Pnuc_0665, amino acid/polyamine/organocation transporter, APC superfamily |
| m6A | 659995 | - | | Pnuc_0663, Pyridoxamine 5'-phosphate oxidase |
| m6A | 664401 | + | CTAYNNNNNNNNTRTC | upstream Pnuc_0672, ABC transporter related protein |
| m6A | 664411 | - | GAYANNNNNNNNRTAG | Pnuc_0670, cyanophycin synthetase |
| m6A | 671237 | - | | upstream Pnuc_0671, cyanophycin synthetase |
| m6A | 671297 | - | | upstream Pnuc_0671, cyanophycin synthetase |
| m6A | 673790 | + | | Pnuc_0676, TonB-dependent receptor |
| m4C | 678877 | - | | Pnuc_0680, dihydroxyacid dehydratase |
| m4C | 681991 | + | | upstream Pnuc_0686, V-type H(+)-translocating pyrophosphatase |
| m6A | 685642 | + | CTAYNNNNNNNNTRTC | Pnuc_0686, V-type H(+)-translocating pyrophosphatase |
| m6A | 685652 | - | GAYANNNNNNNNRTAG | upstream Pnuc_0685, Inorganic diphosphatase |
| m6A | 686516 | + | | upstream Pnuc_0687, conserved hypothetical protein |
| m4C | 688473 | + | | upstream Pnuc_0690, DNA translocase FtsK |
| m6A | 688785 | + | | Pnuc_0690, DNA translocase FtsK |
| m6A | 688807 | - | | upstream Pnuc_0689, thioredoxin reductase |
| m6A | 693622 | + | | upstream Pnuc_R0018, tRNA-Ser |
| m6A | 696582 | - | | upstream Pnuc_0694, transcriptional regulator/antitoxin, MazE |
| m4C | 699999 | - | | upstream Pnuc_0699, Exonuclease, RNase T and DNA polymerase III |
| m4C | 702532 | - | | upstream Pnuc_R0019, SRP RNA; RNA component of signal recognition particle |
| m6A | 706399 | + | | Pnuc_0705, DNA-directed DNA polymerase |
| m6A | 706438 | + |  | upstream Pnuc_0706, Ferric reductase domain protein protein transmembrane component, N-terminal domain protein |
| m6A | 706486 | - | | upstream Pnuc_R0019, SRP RNA; RNA component of signal recognition particle |
| m6A | 708158 | - | | Pnuc_0708, hypothetical protein |
| m6A | 708630 | + | | upstream Pnuc_0709, protein of unknown function DUF465 |
| m6A | 709971 | + | | Pnuc_0711, AsmA family protein |
| m6A | 716204 | - | | upstream Pnuc_0716, YaeQ family protein |
| m6A | 716815 | + | CTAYNNNNNNNNTRTC | Pnuc_0718, coproporphyrinogen III oxidase, anaerobic |
| m6A | 716825 | - | GAYANNNNNNNNRTAG | upstream Pnuc_0716, YaeQ family protein |
| m6A | 717198 | + | CTAYNNNNNNNNTRTC | upstream Pnuc_0719, OmpW family protein |
| m6A | 717208 | - | GAYANNNNNNNNRTAG | upstream Pnuc_0716, YaeQ family protein |
| m6A | 720208 | + | | Pnuc_0721, ABC transporter related protein |
| m4C | 720366 | + | | Pnuc_0721, ABC transporter related protein |
| m6A | 723879 | + | | upstream Pnuc_0726, TRAP transporter solute receptor, TAXI family |
| m6A | 729769 | - | | Pnuc_0730, oligopeptidase A, Metallo peptidase, MEROPS family M03A |
| m4C | 729810 | + | | upstream Pnuc_0734, 2-oxo-acid dehydrogenase E1 subunit, homodimeric type |
| m4C | 729823 | + | | upstream Pnuc_0734, 2-oxo-acid dehydrogenase E1 subunit, homodimeric type |

| | | | | |
|---|---|---|---|---|
| m4C | 730022 | - | | Pnuc_0730, oligopeptidase A, Metallo peptidase, MEROPS family M03A |
| m6A | 730423 | + | CTAYNNNNNNNNTRTC | upstream Pnuc_0734, 2-oxo-acid dehydrogenase E1 subunit, homodimeric type |
| m6A | 730433 | - | GAYANNNNNNNNRTAG | Pnuc_0731, methenyltetrahydrofolate cyclohydrolase / 5,10-methylenetetrahydrofolate dehydrogenase (NADP+) |
| m6A | 732539 | - | | Pnuc_0733, PAS/PAC sensor signal transduction histidine kinase |
| m6A | 732949 | - | | Pnuc_0733, PAS/PAC sensor signal transduction histidine kinase |
| m4C | 733221 | + | | upstream Pnuc_0734, 2-oxo-acid dehydrogenase E1 subunit, homodimeric type |
| m4C | 733398 | + | | upstream Pnuc_0734, 2-oxo-acid dehydrogenase E1 subunit, homodimeric type |
| m6A | 734167 | + | | upstream Pnuc_0734, 2-oxo-acid dehydrogenase E1 subunit, homodimeric type |
| m4C | 734248 | - | | Pnuc_0733, PAS/PAC sensor signal transduction histidine kinase |
| m6A | 739406 | + | CTAYNNNNNNNNTRTC | Pnuc_0736, dihydrolipoamide dehydrogenase |
| m6A | 739416 | - | GAYANNNNNNNNRTAG | upstream Pnuc_0733, PAS/PAC sensor signal transduction histidine kinase |
| m6A | 740234 | - | | upstream Pnuc_0733, PAS/PAC sensor signal transduction histidine kinase |
| m4C | 741368 | + | | upstream Pnuc_0738, protein of unknown function DUF6, transmembrane |
| m6A | 741643 | + | CTAYNNNNNNNNTRTC | Pnuc_0738, protein of unknown function DUF6, transmembrane |
| m6A | 741653 | - | GAYANNNNNNNNRTAG | upstream Pnuc_0737, phasin family protein |
| m6A | 744485 | + | | upstream Pnuc_0753, protein of unknown function DUF328 |
| m4C | 745336 | - | | Pnuc_0741, dihydroorotate oxidase A |
| m6A | 749953 | + | | upstream Pnuc_0753, protein of unknown function DUF328 |
| m6A | 750406 | + | GAYANNNNNNNNRTAG | upstream Pnuc_0753, protein of unknown function DUF328 |
| m6A | 750416 | - | CTAYNNNNNNNNTRTC | Pnuc_0747, DSBA oxidoreductase |
| m6A | 751528 | + | | upstream Pnuc_0753, protein of unknown function DUF328 |
| m6A | 754281 | + | | upstream Pnuc_0753, protein of unknown function DUF328 |
| m6A | 755475 | + | | Pnuc_0753, protein of unknown function DUF328 |
| m6A | 759180 | + | | Pnuc_0757,transcriptional regulator, GntR family |
| m6A | 762452 | + | CTAYNNNNNNNNTRTC | Pnuc_0761, succinate dehydrogenase subunit B |
| m6A | 762462 | - | GAYANNNNNNNNRTAG | upstream Pnuc_0756, malate dehydrogenase (NAD) |
| m6A | 762693 | - | | upstream Pnuc_0756, malate dehydrogenase (NAD) |
| m6A | 766548 | - | | upstream Pnuc_0756, malate dehydrogenase (NAD) |
| m6A | 768396 | + | | Pnuc_0768, aspartate semialdehyde dehydrogenase |
| m6A | 768513 | + | CTAYNNNNNNNNTRTC | Pnuc_0768, aspartate semialdehyde dehydrogenase |
| m6A | 768523 | - | GAYANNNNNNNNRTAG | upstream Pnuc_0756, malate dehydrogenase (NAD) |
| m6A | 770717 | + | | Pnuc_0769, Tfp pilus assembly protein FimV-like protein |
| m6A | 774540 | + | CTAYNNNNNNNNTRTC | Pnuc_0774, acetyl-CoA carboxylase carboxyltransferase subunit alpha |
| m6A | 774550 | - | GAYANNNNNNNNRTAG | upstream Pnuc_0756, malate dehydrogenase (NAD) |
| m6A | 777961 | - | | upstream Pnuc_0756, malate dehydrogenase (NAD) |
| m6A | 783973 | + | GAYANNNNNNNNRTAG | Pnuc_0784, OmpA/MotB domain protein |
| m6A | 783983 | - | CTAYNNNNNNNNTRTC | upstream Pnuc_0782, molybdate ABC transporter, inner membrane subunit |
| m6A | 786874 | + | GAYANNNNNNNNRTAG | upstream Pnuc_0788, protein of unknown function DUF395, YeeE/YedE |
| m6A | 786884 | - | CTAYNNNNNNNNTRTC | Pnuc_0787, cytochrome c family protein |
| m4C | 787697 | - | | upstream Pnuc_0787, cytochrome c family protein |
| m6A | 789389 | + | GAYANNNNNNNNRTAG | Pnuc_0792,Rhodanese domain protein |
| m6A | 789399 | - | CTAYNNNNNNNNTRTC | upstream Pnuc_0790, hypothetical protein |
| m6A | 793524 | - | | Pnuc_0798, protein of unknown function DUF81 |
| m6A | 793988 | + | CTAYNNNNNNNNTRTC | upstream Pnuc_0800, beta-lactamase domain protein |

| | | | | |
|---|---|---|---|---|
| m6A | 793998 | - | GAYANNNNNNNRTAG | Pnuc_0799, FAD-dependent pyridine nucleotide-disulfide oxidoreductase |
| m6A | 795260 | - | | upstream Pnuc_0799, FAD-dependent pyridine nucleotide-disulfide oxidoreductase |
| m4C | 796423 | - | | upstream Pnuc_0799, FAD-dependent pyridine nucleotide-disulfide oxidoreductase |
| m6A | 797960 | + | | Pnuc_0802, sulfur dehydrogenase subunit SoxD |
| m6A | 799290 | + | | Pnuc_0802, sulfur dehydrogenase subunit SoxD |
| m6A | 800576 | - | | upstream Pnuc_0799, FAD-dependent pyridine nucleotide-disulfide oxidoreductase |
| m4C | 801384 | - | | upstream Pnuc_0799, FAD-dependent pyridine nucleotide-disulfide oxidoreductase |
| m6A | 802163 | + | GAYANNNNNNNRTAG | Pnuc_0807, sulfate thiol esterase SoxB |
| m6A | 802173 | - | CTAYNNNNNNNTRTC | upstream Pnuc_0799, FAD-dependent pyridine nucleotide-disulfide oxidoreductase |
| m4C | 804728 | - | | upstream Pnuc_0799, FAD-dependent pyridine nucleotide-disulfide oxidoreductase |
| m6A | 807320 | - | | upstream Pnuc_0799, FAD-dependent pyridine nucleotide-disulfide oxidoreductase |
| m4C | 810697 | + | | upstream Pnuc_0819, conserved hypothetical integral membrane protein |
| m6A | 811058 | - | | Pnuc_0818, acyl-CoA dehydrogenase domain protein |
| m4C | 811995 | - | | upstream Pnuc_0818, acyl-CoA dehydrogenase domain protein |
| m6A | 815395 | + | GAYANNNNNNNRTAG | upstream Pnuc_0825, 3-oxoacid CoA-transferase, A subunit |
| m6A | 815405 | - | CTAYNNNNNNNTRTC | Pnuc_0822, protein of unknown function DUF482 |
| m6A | 819321 | - | | upstream Pnuc_0824, Electron-transferring-flavoprotein dehydrogenase |
| m6A | 820903 | + | CTAYNNNNNNNTRTC | Pnuc_0827, cation diffusion facilitator family transporter |
| m6A | 820913 | - | GAYANNNNNNNRTAG | upstream Pnuc_0824, Electron-transferring-flavoprotein dehydrogenase |
| m4C | 821539 | + | | Pnuc_0828, (p)ppGpp synthetase I, SpoT/RelA |
| m4C | 822105 | + | | Pnuc_0828, (p)ppGpp synthetase I, SpoT/RelA |
| m6A | 822198 | + | | Pnuc_0828, (p)ppGpp synthetase I, SpoT/RelA |
| m6A | 823095 | + | | Pnuc_0828, (p)ppGpp synthetase I, SpoT/RelA |
| m4C | 823200 | + | | Pnuc_0828, (p)ppGpp synthetase I, SpoT/RelA |
| m6A | 829107 | + | CTAYNNNNNNNTRTC | Pnuc_0834, phenylalanyl-tRNA synthetase beta subunit |
| m6A | 829117 | - | GAYANNNNNNNRTAG | upstream Pnuc_0824, Electron-transferring-flavoprotein dehydrogenase |
| m6A | 834695 | - | | upstream Pnuc_0839, DNA binding domain, excisionase family |
| m6A | 837877 | + | GAYANNNNNNNRTAG | Pnuc_0842, dihydrolipoamide dehydrogenase |
| m6A | 837887 | - | CTAYNNNNNNNTRTC | upstream Pnuc_0839, DNA binding domain, excisionase family |
| m4C | 840269 | + | | Pnuc_0844, PHP C-terminal domain protein |
| m4C | 845973 | - | | Pnuc_0850, Cupin 4 family protein |
| m6A | 856210 | + | GAYANNNNNNNRTAG | Pnuc_0862, aspartate kinase |
| m6A | 856220 | - | CTAYNNNNNNNTRTC | upstream Pnuc_0857, Tetratricopeptide TPR_2 repeat protein |
| m6A | 857075 | - | | upstream Pnuc_0857, Tetratricopeptide TPR_2 repeat protein |
| m6A | 857209 | - | | upstream Pnuc_0857, Tetratricopeptide TPR_2 repeat protein |
| m4C | 857909 | + | | Pnuc_0863, metallophosphoesterase |
| m6A | 857935 | - | | upstream Pnuc_0857, Tetratricopeptide TPR_2 repeat protein |
| m6A | 864766 | + | CTAYNNNNNNNTRTC | Pnuc_0870, transporter, hydrophobe/amphiphile efflux-1(HAE1) family |
| m6A | 864776 | - | GAYANNNNNNNRTAG | upstream Pnuc_0865, transcriptional regulator, XRE family |
| m6A | 865018 | + | CTAYNNNNNNNTRTC | Pnuc_0870, transporter, hydrophobe/amphiphile efflux-1(HAE1) family |
| m6A | 865028 | - | GAYANNNNNNNRTAG | upstream Pnuc_0865, transcriptional regulator, XRE family |
| m6A | 865032 | - | | upstream Pnuc_0865, transcriptional regulator, XRE family |
| m6A | 865350 | - | | upstream Pnuc_0865, transcriptional regulator, XRE family |
| m6A | 868410 | + | GAYANNNNNNNRTAG | Pnuc_0872, Formyl-CoA transferase |

| | | | | |
|---|---|---|---|---|
| m6A | 868420 | - | CTAYNNNNNNNTRTC | upstream Pnuc_0865, transcriptional regulator, XRE family |
| m4C | 868513 | - | | upstream Pnuc_0865, transcriptional regulator, XRE family |
| m4C | 871319 | - | | Pnuc_0875, hypothetical protein |
| m4C | 871406 | + | | upstream Pnuc_0876, hypothetical protein |
| m6A | 873168 | + | CTAYNNNNNNNTRTC | Pnuc_0879, putative transmembrane protein |
| m6A | 873178 | - | GAYANNNNNNNRTAG | upstream Pnuc_0875, hypothetical protein |
| m4C | 873322 | + | | Pnuc_0880, hypothetical protein |
| m4C | 874847 | + | | Pnuc_0883, AMP-dependent synthetase and ligase |
| m4C | 875451 | + | | Pnuc_0883, AMP-dependent synthetase and ligase |
| m4C | 878708 | + | | upstream Pnuc_0888, hypothetical protein |
| m6A | 881884 | - | | Pnuc_R0023, tRNA-Asn |
| m6A | 882224 | - | CTAYNNNNNNNTRTC | upstream Pnuc_R0023, tRNA-Asn |
| m6A | 883131 | - | | upstream Pnuc_R0023, tRNA-Asn |
| m6A | 891700 | - | | Pnuc_0899, Formyl-CoA transferase |
| m4C | 891750 | - | | Pnuc_0899, Formyl-CoA transferase |
| m6A | 894883 | + | GAYANNNNNNNRTAG | Pnuc_0901, DNA topoisomerase IV subunit A |
| m6A | 894893 | - | CTAYNNNNNNNTRTC | upstream Pnuc_0899, Formyl-CoA transferase |
| m6A | 895617 | + | | Pnuc_0901, DNA topoisomerase IV subunit A |
| m4C | 896715 | + | | Pnuc_0901, DNA topoisomerase IV subunit A |
| m6A | 898234 | + | CTAYNNNNNNNTRTC | upstream Pnuc_0903, conserved hypothetical protein |
| m6A | 898244 | - | GAYANNNNNNNRTAG | upstream Pnuc_0902, predicted sulfurylase subunit, molybdopterin cytosine dinucleotide biosynthesis |
| m6A | 898893 | - | | upstream Pnuc_0902, predicted sulfurylase subunit, molybdopterin cytosine dinucleotide biosynthesis |
| m6A | 903152 | + | GAYANNNNNNNRTAG | upstream Pnuc_0910, methylmalonyl-CoA mutase |
| m6A | 903162 | - | CTAYNNNNNNNTRTC | upstream Pnuc_0909, transcriptional regulator, GntR family |
| m4C | 905948 | + | | Pnuc_0911, LAO/AO transport system ATPase |
| m4C | 906214 | + | | Pnuc_0911, LAO/AO transport system ATPase |
| m6A | 908367 | + | | Pnuc_0913, acetyl-CoA carboxylase, biotin carboxylase |
| m6A | 908555 | + | | Pnuc_0913, acetyl-CoA carboxylase, biotin carboxylase |
| m6A | 909186 | - | | upstream Pnuc_0909, transcriptional regulator, GntR family |
| m6A | 909439 | + | CTAYNNNNNNNTRTC | Pnuc_0913, acetyl-CoA carboxylase, biotin carboxylase |
| m6A | 909449 | - | GAYANNNNNNNRTAG | upstream Pnuc_0909, transcriptional regulator, GntR family |
| m4C | 920817 | + | | Pnuc_0925, polyhydroxyalkonate synthesis repressor, PhaR |
| m4C | 926448 | - | | Pnuc_0929, transcription-repair coupling factor |
| m6A | 927815 | + | | upstream Pnuc_0930, 2-C-methyl-D-erythritol 4-phosphate cytidylyltransferase |
| m4C | 927861 | - | | Pnuc_0929, transcription-repair coupling factor |
| m6A | 931352 | - | | upstream Pnuc_0929, transcription-repair coupling factor |
| m6A | 933367 | + | | Pnuc_0935, ATP-dependent Clp protease ATP-binding subunit ClpX |
| m4C | 933464 | + | | Pnuc_0935, ATP-dependent Clp protease ATP-binding subunit ClpX |
| m6A | 935451 | + | | Pnuc_0936, ATP-dependent proteinase, Serine peptidase, MEROPS family S16 |
| m4C | 936279 | - | | upstream Pnuc_0929, transcription-repair coupling factor |
| m6A | 936991 | + | CTAYNNNNNNNTRTC | Pnuc_0937, hypothetical protein |
| m6A | 937001 | - | GAYANNNNNNNRTAG | upstream Pnuc_0929, transcription-repair coupling factor |
| m6A | 939732 | + | | Pnuc_0940, phosphoribosylformylglycinamidine synthase |
| m6A | 943496 | + | | Pnuc_0941, conserved hypothetical protein |

| | | | | |
|---|---|---|---|---|
| m6A | 946091 | + | GAYANNNNNNNRTAG | upstream Pnuc_0945, CTP synthase |
| m6A | 946101 | - | CTAYNNNNNNNTRTC | Pnuc_0942, aconitase |
| m4C | 946664 | + | | upstream Pnuc_0945, CTP synthase |
| m4C | 946768 | + | | upstream Pnuc_0945, CTP synthase |
| m4C | 954511 | - | | upstream Pnuc_0949, Hsp33 protein |
| m6A | 959740 | + | GAYANNNNNNNRTAG | Pnuc_0958, 2-hydroxy-3-oxopropionate reductase |
| m6A | 959750 | - | CTAYNNNNNNNTRTC | upstream Pnuc_0957, hypothetical protein |
| m6A | 959916 | + | | Pnuc_0958, 2-hydroxy-3-oxopropionate reductase |
| m6A | 959987 | - | | upstream Pnuc_0957, hypothetical protein |
| m4C | 962782 | + | | upstream Pnuc_0968, glycine cleavage T protein (aminomethyl transferase) |
| m6A | 964248 | + | | upstream Pnuc_0968, glycine cleavage T protein (aminomethyl transferase) |
| m6A | 964737 | + | | upstream Pnuc_0968, glycine cleavage T protein (aminomethyl transferase) |
| m6A | 966942 | + | CTAYNNNNNNNTRTC | upstream Pnuc_0968, glycine cleavage T protein (aminomethyl transferase) |
| m6A | 966952 | - | GAYANNNNNNNRTAG | Pnuc_0967, aminodeoxychorismate lyase |
| m6A | 967979 | + | CTAYNNNNNNNTRTC | Pnuc_0968, glycine cleavage T protein (aminomethyl transferase) |
| m6A | 967989 | - | GAYANNNNNNNRTAG | upstream Pnuc_0967, aminodeoxychorismate lyase |
| m6A | 974583 | + | | upstream Pnuc_0980, flavodoxin/nitric oxide synthase |
| m6A | 975403 | - | | Pnuc_0977, chorismate synthase |
| m6A | 975704 | + | | upstream Pnuc_0980, flavodoxin/nitric oxide synthase |
| m4C | 979951 | - | | upstream Pnuc_0979, tRNA-processing RNAse BN |
| m6A | 980984 | + | GAYANNNNNNNRTAG | upstream Pnuc_0999, ABC-type transporter, periplasmic component, NitT family |
| m6A | 980994 | - | CTAYNNNNNNNTRTC | Pnuc_0983, alpha/beta hydrolase fold protein |
| m6A | 982286 | + | CTAYNNNNNNNTRTC | upstream Pnuc_0999, ABC-type transporter, periplasmic component, NitT family |
| m6A | 982296 | - | GAYANNNNNNNRTAG | Pnuc_0986, hypothetical protein |
| m6A | 988136 | + | GAYANNNNNNNRTAG | upstream Pnuc_0999, ABC-type transporter, periplasmic component, NitT family |
| m6A | 988146 | - | CTAYNNNNNNNTRTC | Pnuc_0992, Choloylglycine hydrolase |
| m4C | 993458 | - | | upstream Pnuc_0997, ABC transporter nitrate-binding protein |
| m6A | 995404 | + | CTAYNNNNNNNTRTC | Pnuc_0999, ABC-type transporter, periplasmic component, NitT family |
| m6A | 995414 | - | GAYANNNNNNNRTAG | upstream Pnuc_0998, uroporphyrin-III C-methyltransferase |
| m6A | 997866 | + | | Pnuc_1000, assimilatory nitrite reductase (NAD(P)H) large subunit precursor |
| m6A | 999078 | - | | upstream Pnuc_0998, uroporphyrin-III C-methyltransferase |
| m6A | 1000022 | + | | Pnuc_1002, assimilatory nitrate reductase (NADH) alpha subunit apoprotein |
| m6A | 1000889 | + | | Pnuc_1002, assimilatory nitrate reductase (NADH) alpha subunit apoprotein |
| m4C | 1001341 | - | | upstream Pnuc_0998, uroporphyrin-III C-methyltransferase |
| m6A | 1001443 | - | | upstream Pnuc_0998, uroporphyrin-III C-methyltransferase |
| m4C | 1003514 | + | | upstream Pnuc_1007, protein of unknown function DUF1348 |
| m4C | 1005364 | + | | Pnuc_1008, nitroreductase |
| m6A | 1008651 | + | CTAYNNNNNNNTRTC | Pnuc_1010, TonB-dependent receptor |
| m6A | 1008661 | - | GAYANNNNNNNRTAG | upstream Pnuc_R0031, tRNA-Pro2 |
| m4C | 1009132 | + | | upstream Pnuc_1015, protein of unknown function UPF0044 |
| m6A | 1010336 | + | | upstream Pnuc_1015, protein of unknown function UPF0044 |
| m6A | 1013228 | + | | upstream Pnuc_1015, protein of unknown function UPF0044 |
| m4C | 1016761 | + | | upstream Pnuc_1023, Methyltransferase type 11 |
| m6A | 1019053 | + | CTAYNNNNNNNTRTC | upstream Pnuc_1023, Methyltransferase type 11 |

| | | | | |
|---|---|---|---|---|
| m6A | 1019063 | - | GAYANNNNNNNRTAG | Pnuc_1019, carbamoyl-phosphate synthase small subunit |
| m6A | 1019396 | - | | Pnuc_1019, carbamoyl-phosphate synthase small subunit |
| m6A | 1020266 | - | GAYANNNNNNNRTAG | upstream Pnuc_1019, carbamoyl-phosphate synthase small subunit |
| m4C | 1021239 | + | | upstream Pnuc_1023, Methyltransferase type 11 |
| m6A | 1021303 | + | CTAYNNNNNNNTRTC | upstream Pnuc_1023, Methyltransferase type 11 |
| m6A | 1021313 | - | GAYANNNNNNNRTAG | Pnuc_1020, propionyl-CoA synthetase |
| m4C | 1021746 | + | | upstream Pnuc_1023, Methyltransferase type 11 |
| m6A | 1027364 | + | | Pnuc_1026, acriflavin resistance protein |
| m6A | 1027462 | + | GAYANNNNNNNRTAG | Pnuc_1026, acriflavin resistance protein |
| m6A | 1027472 | - | CTAYNNNNNNNTRTC | upstream Pnuc_1022, Hydroxyacylglutathione hydrolase |
| m4C | 1030880 | + | | Pnuc_1027, DNA polymerase III, epsilon subunit |
| m6A | 1032141 | + | | upstream Pnuc_1032, Amidase |
| m4C | 1033093 | - | | Pnuc_1030, hypothetical protein |
| m6A | 1033357 | + | | upstream Pnuc_1032, Amidase |
| m6A | 1034059 | - | | Pnuc_1031, hypothetical protein |
| m6A | 1034060 | + | CTAYNNNNNNNTRTC | upstream Pnuc_1032, Amidase |
| m6A | 1034070 | - | GAYANNNNNNNRTAG | Pnuc_1031, hypothetical protein |
| m4C | 1036386 | - | | Pnuc_1033, Uncharacterized protein UPF0065 |
| m6A | 1038226 | + | | upstream Pnuc_1035, methylmalonate-semialdehyde dehydrogenase (acylating) |
| m6A | 1042338 | + | | upstream Pnuc_1057, putative lipoprotein |
| m4C | 1043542 | + | | upstream Pnuc_1057, putative lipoprotein |
| m4C | 1046690 | + | | upstream Pnuc_1057, putative lipoprotein |
| m6A | 1050989 | + | | upstream Pnuc_1057, putative lipoprotein |
| m6A | 1055538 | + | GAYANNNNNNNRTAG | upstream Pnuc_1057, putative lipoprotein |
| m6A | 1055548 | - | CTAYNNNNNNNTRTC | Pnuc_1051, NADH dehydrogenase subunit A |
| m4C | 1055615 | - | | Pnuc_1051, NADH dehydrogenase subunit A |
| m6A | 1061186 | - | | upstream Pnuc_1056, SSU ribosomal protein S15P |
| m6A | 1061342 | - | | upstream Pnuc_1056, SSU ribosomal protein S15P |
| m4C | 1064577 | + | | upstream Pnuc_1064, RNA polymerase, sigma-24 subunit, RpoE |
| m4C | 1068864 | + | | Pnuc_1064, RNA polymerase, sigma-24 subunit, RpoE |
| m4C | 1068952 | - | | upstream Pnuc_1063, acetolactate synthase, large subunit |
| m4C | 1068979 | + | | Pnuc_1064, RNA polymerase, sigma-24 subunit, RpoE |
| m4C | 1071149 | - | | Pnuc_1068, protein of unknown function DUF214 |
| m4C | 1071207 | + | | upstream Pnuc_1075, transcription elongation factor GreB |
| m6A | 1073342 | + | | upstream Pnuc_1075, transcription elongation factor GreB |
| m6A | 1074889 | - | | Pnuc_1070, Uncharacterized protein UPF0065 |
| m4C | 1075611 | + | | upstream Pnuc_1075, transcription elongation factor GreB |
| m6A | 1075780 | + | | upstream Pnuc_1075, transcription elongation factor GreB |
| m6A | 1075790 | - | | Pnuc_1071, hypothetical protein |
| m6A | 1076951 | + | CTAYNNNNNNNTRTC | upstream Pnuc_1075, transcription elongation factor GreB |
| m6A | 1076961 | - | GAYANNNNNNNRTAG | Pnuc_1073, ABC transporter related protein |
| m6A | 1077434 | + | | upstream Pnuc_1075, transcription elongation factor GreB |
| m6A | 1082384 | + | | upstream Pnuc_1080, RNAse PH |
| m6A | 1082427 | + | | upstream Pnuc_1080, RNAse PH |

| | | | | |
|---|---|---|---|---|
| m6A | 1088310 | - | | Pnuc_1084, beta-lactamase domain protein |
| m6A | 1090450 | + | | upstream Pnuc_1086, Patatin |
| m6A | 1093084 | + | GAYANNNNNNNRTAG | Pnuc_1087, hypothetical protein |
| m6A | 1093094 | - | CTAYNNNNNNNTRTC | upstream Pnuc_1085, integral membrane sensor signal transduction histidine kinase |
| m6A | 1095712 | + | CTAYNNNNNNNTRTC | upstream Pnuc_1091, glucose-6-phosphate isomerase |
| m6A | 1095722 | - | GAYANNNNNNNRTAG | upstream Pnuc_1089, hypothetical protein |
| m6A | 1101440 | + | | upstream Pnuc_1097, hypothetical protein |
| m6A | 1108023 | + | GAYANNNNNNNRTAG | upstream Pnuc_1097, hypothetical protein |
| m6A | 1108033 | - | CTAYNNNNNNNTRTC | Pnuc_1095, outer membrane autotransporter barrel domain protein |
| m6A | 1109515 | + | CTAYNNNNNNNTRTC | upstream Pnuc_1097, hypothetical protein |
| m6A | 1109525 | - | GAYANNNNNNNRTAG | Pnuc_1095, outer membrane autotransporter barrel domain protein |
| m6A | 1111707 | + | GAYANNNNNNNRTAG | upstream Pnuc_1097, hypothetical protein |
| m6A | 1111717 | - | CTAYNNNNNNNTRTC | Pnuc_1095, outer membrane autotransporter barrel domain protein |
| m6A | 1111938 | + | | upstream Pnuc_1097, hypothetical protein |
| m4C | 1112185 | - | | Pnuc_1095, outer membrane autotransporter barrel domain protein |
| m6A | 1112942 | + | | upstream Pnuc_1097, hypothetical protein |
| m4C | 1114862 | + | | upstream Pnuc_1097, hypothetical protein |
| m6A | 1115655 | - | | Pnuc_1095, outer membrane autotransporter barrel domain protein |
| m6A | 1115916 | + | GAYANNNNNNNRTAG | upstream Pnuc_1097, hypothetical protein |
| m6A | 1115926 | - | CTAYNNNNNNNTRTC | Pnuc_1095, outer membrane autotransporter barrel domain protein |
| m4C | 1116418 | + | | upstream Pnuc_1097, hypothetical protein |
| m6A | 1117112 | + | | upstream Pnuc_1097, hypothetical protein |
| m4C | 1117990 | + | | upstream Pnuc_1097, hypothetical protein |
| m6A | 1120412 | + | | upstream Pnuc_1097, hypothetical protein |
| m6A | 1121807 | + | | upstream Pnuc_1097, hypothetical protein |
| m6A | 1122497 | - | | Pnuc_1095, outer membrane autotransporter barrel domain protein |
| m6A | 1124061 | - | | Pnuc_1095, outer membrane autotransporter barrel domain protein |
| m6A | 1124249 | + | | upstream Pnuc_1097, hypothetical protein |
| m6A | 1125273 | + | | upstream Pnuc_1097, hypothetical protein |
| m6A | 1129263 | - | | Pnuc_1095, outer membrane autotransporter barrel domain protein |
| m6A | 1129364 | - | | Pnuc_1095, outer membrane autotransporter barrel domain protein |
| m6A | 1130061 | + | | upstream Pnuc_1097, hypothetical protein |
| m4C | 1138027 | - | | Pnuc_1096, sulfotransferase |
| m6A | 1138538 | - | | Pnuc_1096, sulfotransferase |
| m6A | 1139780 | - | | upstream Pnuc_1096, sulfotransferase |
| m6A | 1145089 | + | | Pnuc_1103, quinolinate synthetase A |
| m6A | 1150445 | + | GAYANNNNNNNRTAG | upstream Pnuc_1112, protein of unknown function DUF1080 |
| m6A | 1150455 | - | CTAYNNNNNNNTRTC | Pnuc_1107, transcriptional modulator of MazE/toxin, MazF |
| m6A | 1151487 | + | GAYANNNNNNNRTAG | upstream Pnuc_1112, protein of unknown function DUF1080 |
| m6A | 1151497 | - | CTAYNNNNNNNTRTC | Pnuc_1109, NAD-dependent formate dehydrogenase iron-sulfur protein (catalytic activity) |
| m6A | 1156346 | + | CTAYNNNNNNNTRTC | upstream Pnuc_1112, protein of unknown function DUF1080 |
| m6A | 1156356 | - | GAYANNNNNNNRTAG | Pnuc_1111, protein of unknown function DUF748 |
| m6A | 1157974 | - | | Pnuc_1111, protein of unknown function DUF748 |
| m6A | 1159304 | + | GAYANNNNNNNRTAG | Pnuc_1112, protein of unknown function DUF1080 |

| | | | | |
|---|---|---|---|---|
| m6A | 1159314 | - | CTAYNNNNNNNTRTC | upstream Pnuc_1111, protein of unknown function DUF748 |
| m6A | 1159365 | + | | Pnuc_1112, protein of unknown function DUF1080 |
| m6A | 1160589 | - | | upstream Pnuc_1115, YCII-related protein |
| m6A | 1160818 | + | | Pnuc_1116, phage integrase family protein |
| m6A | 1162259 | - | | Pnuc_1117, hypothetical protein |
| m6A | 1164942 | - | | upstream Pnuc_1119, hypothetical protein |
| m6A | 1167044 | + | CTAYNNNNNNNTRTC | Pnuc_1122, Exonuclease, RNase T and DNA polymerase III |
| m6A | 1167054 | - | GAYANNNNNNNRTAG | upstream Pnuc_1121, hypothetical protein |
| m6A | 1170355 | - | | Pnuc_1124, hypothetical protein |
| m4C | 1173713 | + | | upstream Pnuc_1127, hypothetical protein |
| m6A | 1173977 | + | GAYANNNNNNNRTAG | Pnuc_1127, hypothetical protein |
| m6A | 1173987 | - | CTAYNNNNNNNTRTC | upstream Pnuc_1126, hypothetical protein |
| m4C | 1176445 | - | | upstream Pnuc_1129, conserved hypothetical protein |
| m6A | 1177319 | + | GAYANNNNNNNRTAG | upstream Pnuc_1131, phage transcriptional regulator, AlpA |
| m6A | 1177329 | - | CTAYNNNNNNNTRTC | Pnuc_1130, protein of unknown function DUF1376 |
| m6A | 1177333 | - | | Pnuc_1130, protein of unknown function DUF1376 |
| m4C | 1177393 | + | | upstream Pnuc_1131, phage transcriptional regulator, AlpA |
| m6A | 1180795 | - | | Pnuc_1133, N-6 DNA methylase |
| m4C | 1187078 | + | | upstream Pnuc_1139, fumarase |
| m4C | 1188896 | + | | upstream Pnuc_1139, fumarase |
| m6A | 1189256 | + | | Pnuc_1139, fumarase |
| m6A | 1191165 | + | CTAYNNNNNNNTRTC | Pnuc_1140, glutamate racemase |
| m6A | 1191175 | - | GAYANNNNNNNRTAG | upstream Pnuc_1138, acetyl-coenzyme A synthetase |
| m4C | 1197227 | + | | upstream Pnuc_1150, Penicillin amidase |
| m6A | 1199103 | + | | upstream Pnuc_1150, Penicillin amidase |
| m4C | 1199796 | + | | upstream Pnuc_1150, Penicillin amidase |
| m6A | 1199870 | + | GAYANNNNNNNRTAG | upstream Pnuc_1150, Penicillin amidase |
| m6A | 1199880 | - | CTAYNNNNNNNTRTC | Pnuc_1149, filamentous hemagglutinin family outer membrane protein |
| m6A | 1201836 | - | | Pnuc_1149, filamentous hemagglutinin family outer membrane protein |
| m4C | 1204187 | + | | upstream Pnuc_1150, Penicillin amidase |
| m6A | 1205143 | - | | upstream Pnuc_R0036, tRNA-Leu |
| m6A | 1206854 | + | CTAYNNNNNNNTRTC | Pnuc_1150, Penicillin amidase |
| m6A | 1206864 | - | GAYANNNNNNNRTAG | upstream Pnuc_R0036, tRNA-Leu |
| m6A | 1211099 | - | | Pnuc_1158, aldehyde oxidase and xanthine dehydrogenase, molybdopterin binding protein |
| m6A | 1218238 | + | CTAYNNNNNNNTRTC | upstream Pnuc_1167, Cellulase |
| m6A | 1218248 | - | GAYANNNNNNNRTAG | Pnuc_1164, conserved hypothetical protein |
| m6A | 1221438 | + | | upstream Pnuc_1167, Cellulase |
| m4C | 1221658 | - | | Pnuc_1166, metal dependent phosphohydrolase |
| m6A | 1224126 | + | | Pnuc_1168, conserved hypothetical protein |
| m6A | 1227335 | + | | upstream Pnuc_1176, hypothetical protein |
| m6A | 1228000 | + | GAYANNNNNNNRTAG | upstream Pnuc_1176, hypothetical protein |
| m6A | 1228010 | - | CTAYNNNNNNNTRTC | Pnuc_1170, cellulose synthase operon C domain protein |
| m6A | 1228683 | + | GAYANNNNNNNRTAG | upstream Pnuc_1176, hypothetical protein |
| m6A | 1228693 | - | CTAYNNNNNNNTRTC | Pnuc_1170, cellulose synthase operon C domain protein |

| | | | | |
|---|---|---|---|---|
| m6A | 1228990 | + | CTAYNNNNNNNTRTC | upstream Pnuc_1176, hypothetical protein |
| m6A | 1229000 | - | GAYANNNNNNNRTAG | Pnuc_1171, Cellulose synthase (UDP-forming) |
| m6A | 1229564 | + | | upstream Pnuc_1176, hypothetical protein |
| m6A | 1235057 | + | | Pnuc_1176, hypothetical protein |
| m6A | 1236712 | + | CTAYNNNNNNNTRTC | upstream Pnuc_1184, Amidase |
| m6A | 1239968 | + | GAYANNNNNNNRTAG | upstream Pnuc_1184, Amidase |
| m6A | 1239978 | - | CTAYNNNNNNNTRTC | Pnuc_1180, putative ABC transporter, periplasmic protein |
| m6A | 1240127 | + | GAYANNNNNNNRTAG | upstream Pnuc_1184, Amidase |
| m6A | 1240137 | - | CTAYNNNNNNNTRTC | Pnuc_1180, putative ABC transporter, periplasmic protein |
| m4C | 1244461 | + | | Pnuc_1185, ABC transporter, substrate binding protein |
| m6A | 1245022 | + | | Pnuc_1186, binding-protein-dependent transport systems inner membrane component |
| m6A | 1247136 | + | | upstream Pnuc_1210, Glutamate synthase (NADPH) |
| m6A | 1247968 | + | CTAYNNNNNNNTRTC | upstream Pnuc_1210, Glutamate synthase (NADPH) |
| m6A | 1247978 | - | GAYANNNNNNNRTAG | Pnuc_1190, urease accessory protein UreG |
| m4C | 1248316 | - | | Pnuc_1191, Urease accessory protein UreF |
| m6A | 1252367 | + | | upstream Pnuc_1210, Glutamate synthase (NADPH) |
| m6A | 1252457 | + | GAYANNNNNNNRTAG | upstream Pnuc_1210, Glutamate synthase (NADPH) |
| m6A | 1252467 | - | CTAYNNNNNNNTRTC | Pnuc_1196, urease, gamma subunit |
| m4C | 1253909 | - | | Pnuc_1198, amino acid/amide ABC transporter ATP-binding protein 2, HAAT family |
| m4C | 1262269 | - | | Pnuc_1205, protein of unknown function DUF1501 |
| m6A | 1262425 | + | CTAYNNNNNNNTRTC | upstream Pnuc_1210, Glutamate synthase (NADPH) |
| m6A | 1262435 | - | GAYANNNNNNNRTAG | Pnuc_1205, protein of unknown function DUF1501 |
| m4C | 1262636 | - | | Pnuc_1205, protein of unknown function DUF1501 |
| m4C | 1267728 | - | | Pnuc_1211, diguanylate cyclase |
| m4C | 1273950 | - | | upstream Pnuc_1217, Glutathione S-transferase, N-terminal domain protein |
| m6A | 1278338 | - | | upstream Pnuc_1217, Glutathione S-transferase, N-terminal domain protein |
| m6A | 1280531 | - | | Pnuc_1222, RNA binding S1 domain protein |
| m4C | 1282187 | - | | Pnuc_1222, RNA binding S1 domain protein |
| m6A | 1285244 | - | | Pnuc_1224, GTP-binding protein TypA |
| m6A | 1285533 | - | | Pnuc_1224, GTP-binding protein TypA |
| m6A | 1286269 | + | | upstream Pnuc_1232, Patatin |
| m6A | 1286462 | + | | upstream Pnuc_1232, Patatin |
| m6A | 1289563 | + | | upstream Pnuc_1232, Patatin |
| m4C | 1290028 | + | | upstream Pnuc_1232, Patatin |
| m6A | 1291635 | - | | Pnuc_1229, protein of unknown function DUF150 |
| m6A | 1296429 | + | CTAYNNNNNNNTRTC | upstream Pnuc_1238, Enoyl-[acyl-carrier-protein] reductase (NADH) |
| m6A | 1296439 | - | GAYANNNNNNNRTAG | upstream Pnuc_1233, NLP/P60 protein |
| m6A | 1297321 | + | | upstream Pnuc_1238, Enoyl-[acyl-carrier-protein] reductase (NADH) |
| m6A | 1300658 | + | | upstream Pnuc_1238, Enoyl-[acyl-carrier-protein] reductase (NADH) |
| m6A | 1304342 | - | | upstream Pnuc_1237, extracellular solute-binding protein, family 5 |
| m6A | 1306971 | + | | Pnuc_1242, MmgE/PrpD family protein |
| m6A | 1307149 | + | GAYANNNNNNNRTAG | Pnuc_1242, MmgE/PrpD family protein |
| m6A | 1307159 | - | CTAYNNNNNNNTRTC | upstream Pnuc_1237, extracellular solute-binding protein, family 5 |
| m6A | 1307337 | + | | Pnuc_1242, MmgE/PrpD family protein |

| | | | | |
|---|---|---|---|---|
| m6A | 1311414 | + | CTAYNNNNNNNTRTC | upstream Pnuc_1249, integrase, catalytic region |
| m6A | 1311424 | - | GAYANNNNNNNRTAG | Pnuc_1246, BNR/Asp-box repeat protein |
| m4C | 1313449 | + | | upstream Pnuc_1249, integrase, catalytic region |
| m6A | 1317420 | + | CTAYNNNNNNNTRTC | upstream Pnuc_1262, N-acetylglutamate synthase |
| m6A | 1317430 | - | GAYANNNNNNNRTAG | Pnuc_1252, Exodeoxyribonuclease III |
| m6A | 1319635 | - | | Pnuc_1254, signal transduction histidine kinase, nitrogen specific, NtrB |
| m4C | 1322023 | - | | upstream Pnuc_1255, L-glutamine synthetase |
| m6A | 1328635 | - | | Pnuc_1261, ATP-dependent helicase HrpA |
| m6A | 1330744 | - | | upstream Pnuc_1261, ATP-dependent helicase HrpA |
| m6A | 1338830 | + | | upstream Pnuc_R0038, tRNA-Leu |
| m6A | 1339251 | + | | upstream Pnuc_R0038, tRNA-Leu |
| m6A | 1343384 | + | | Pnuc_1272, amino acid permease-associated region |
| m4C | 1346506 | - | | Pnuc_1276, Carboxymethylenebutenolidase |
| m6A | 1347380 | - | | Pnuc_1277, major facilitator superfamily MFS_1 |
| m6A | 1347471 | + | | upstream Pnuc_1278, hypothetical protein |
| m4C | 1351779 | - | | Pnuc_1281, Adenylosuccinate synthetase |
| m6A | 1353636 | + | | upstream Pnuc_1294, protein of unknown function UPF0005 |
| m4C | 1359614 | + | | upstream Pnuc_1294, protein of unknown function UPF0005 |
| m6A | 1362325 | + | | upstream Pnuc_1294, protein of unknown function UPF0005 |
| m6A | 1362988 | + | | upstream Pnuc_1294, protein of unknown function UPF0005 |
| m6A | 1365158 | + | | upstream Pnuc_1294, protein of unknown function UPF0005 |
| m4C | 1365230 | + | | upstream Pnuc_1294, protein of unknown function UPF0005 |
| m4C | 1367524 | - | | Pnuc_1295, 23S rRNA m(5)U-1939 methyltransferase |
| m6A | 1373200 | + | | upstream Pnuc_1307, thioredoxin |
| m4C | 1373904 | - | | Pnuc_1300, heavy metal efflux pump, CzcA family |
| m4C | 1374186 | - | | Pnuc_1300, heavy metal efflux pump, CzcA family |
| m6A | 1375845 | - | | Pnuc_1302, outer membrane efflux protein |
| m4C | 1377231 | + | | upstream Pnuc_1307, thioredoxin |
| m6A | 1377446 | - | | Pnuc_1302, outer membrane efflux protein |
| m4C | 1378511 | - | | Pnuc_1304, Tetratricopeptide TPR_2 repeat protein |
| m4C | 1380280 | + | | upstream Pnuc_1307, thioredoxin |
| m4C | 1382305 | + | | upstream Pnuc_1307, thioredoxin |
| m6A | 1383404 | + | | upstream Pnuc_1307, thioredoxin |
| m6A | 1383845 | + | | upstream Pnuc_1307, thioredoxin |
| m4C | 1389663 | + | | Pnuc_1310, putative inner membrane transmembrane protein |
| m6A | 1389671 | - | | upstream Pnuc_1306, DNA helicase/exodeoxyribonuclease V, subunit B |
| m6A | 1399354 | + | | upstream Pnuc_1323, Uncharacterized protein UPF0065 |
| m6A | 1408563 | - | | upstream Pnuc_1330, FeoA family protein |
| m6A | 1408639 | + | | Pnuc_1331, Carbohydrate-selective porin OprB |
| m6A | 1410626 | + | GAYANNNNNNNRTAG | Pnuc_1332, negative transcriptional regulator |
| m6A | 1410636 | - | CTAYNNNNNNNTRTC | upstream Pnuc_1330, FeoA family protein |
| m6A | 1412582 | + | | upstream Pnuc_1336, putative sulfate transport system substrate-binding protein |
| m6A | 1413827 | + | | upstream Pnuc_1336, putative sulfate transport system substrate-binding protein |
| m4C | 1413951 | - | | Pnuc_1335, transcriptional regulator of molybdate metabolism, LysR family |

| | | | | |
|---|---|---|---|---|
| m6A | 1413960 | - | | Pnuc_1335, transcriptional regulator of molybdate metabolism, LysR family |
| m6A | 1415967 | - | | Pnuc_1338, hypothetical protein |
| m6A | 1416772 | - | GAYANNNNNNNRTAG | upstream Pnuc_1338, hypothetical protein |
| m6A | 1417579 | + | GAYANNNNNNNRTAG | Pnuc_1341, conserved hypothetical protein |
| m6A | 1417589 | - | CTAYNNNNNNNTRTC | upstream Pnuc_1338, hypothetical protein |
| m6A | 1419263 | + | CTAYNNNNNNNTRTC | Pnuc_1343, transglutaminase domain protein |
| m6A | 1422263 | + | CTAYNNNNNNNTRTC | upstream Pnuc_1347, protein of unknown function DUF1330 |
| m6A | 1422273 | - | GAYANNNNNNNRTAG | upstream Pnuc_1346, conserved hypothetical protein |
| m4C | 1425242 | + | | Pnuc_1350, Rhodanese domain protein |
| m4C | 1427115 | + | | Pnuc_1352, hypothetical protein |
| m4C | 1432162 | - | | upstream Pnuc_1359, conserved hypothetical protein 730 |
| m6A | 1434445 | + | CTAYNNNNNNNTRTC | upstream Pnuc_1363, carbohydrate kinase, YjeF related protein |
| m6A | 1434455 | - | GAYANNNNNNNRTAG | Pnuc_1362, RNA-metabolising metallo-beta-lactamase |
| m4C | 1435865 | - | | upstream Pnuc_1362, RNA-metabolising metallo-beta-lactamase |
| m6A | 1437655 | - | | Pnuc_1367, aminotransferase, class V |
| m6A | 1438918 | - | | Pnuc_1368, Uncharacterized protein UPF0065 |
| m6A | 1439629 | - | | Pnuc_1368, Uncharacterized protein UPF0065 |
| m6A | 1439659 | - | | Pnuc_1368, Uncharacterized protein UPF0065 |
| m4C | 1440473 | - | | Pnuc_1369, 5-carboxymethyl-2-hydroxymuconate Delta-isomerase |
| m4C | 1444140 | - | | upstream Pnuc_1370, protein of unknown function DUF6, transmembrane |
| m4C | 1446510 | - | | Pnuc_1375, transcriptional regulator |
| m4C | 1447384 | + | | Pnuc_1376, Rubrerythrin |
| m4C | 1447784 | - | | upstream Pnuc_1375, transcriptional regulator |
| m6A | 1448329 | + | | Pnuc_1378, FAD-dependent pyridine nucleotide-disulfide oxidoreductase |
| m6A | 1451338 | + | | upstream Pnuc_1382, DSBA oxidoreductase |
| m6A | 1453284 | + | | upstream Pnuc_1382, DSBA oxidoreductase |
| m6A | 1461662 | - | | Pnuc_1390, cardiolipin synthetase 2 |
| m6A | 1464729 | + | GAYANNNNNNNRTAG | upstream Pnuc_1407, heavy metal translocating P-type ATPase |
| m6A | 1464739 | - | CTAYNNNNNNNTRTC | Pnuc_1395, formate dehydrogenase gamma subunit |
| m4C | 1466782 | + | | upstream Pnuc_1407, heavy metal translocating P-type ATPase |
| m6A | 1467035 | + | | upstream Pnuc_1407, heavy metal translocating P-type ATPase |
| m6A | 1473645 | + | GAYANNNNNNNRTAG | upstream Pnuc_1407, heavy metal translocating P-type ATPase |
| m6A | 1473655 | - | CTAYNNNNNNNTRTC | Pnuc_1402, conserved hypothetical protein |
| m6A | 1473786 | + | | upstream Pnuc_1407, heavy metal translocating P-type ATPase |
| m4C | 1474986 | - | | upstream Pnuc_1403, protein of unknown function DUF59 |
| m4C | 1475177 | + | | upstream Pnuc_1407, heavy metal translocating P-type ATPase |
| m4C | 1478223 | - | | upstream Pnuc_1406, Heavy metal transport/detoxification protein |
| m4C | 1478440 | + | | Pnuc_1407, heavy metal translocating P-type ATPase |
| m4C | 1480546 | + | | Pnuc_1409, 4-amino-4-deoxy-L-arabinose transferase and related glycosyltransferase of PMT family-like protein |
| m6A | 1483806 | - | | upstream Pnuc_1408, conserved hypothetical protein |
| m6A | 1484850 | - | | upstream Pnuc_1408, conserved hypothetical protein |
| m6A | 1487423 | - | | Pnuc_1416, CMP/dCMP deaminase, zinc-binding protein |
| m6A | 1488814 | + | CTAYNNNNNNNTRTC | upstream Pnuc_1426, SsrA-binding protein |
| m6A | 1488824 | - | GAYANNNNNNNRTAG | Pnuc_1418, Radical SAM domain protein |

| | | | | |
|---|---|---|---|---|
| m6A | 1489493 | + | | upstream Pnuc_1426, SsrA-binding protein |
| m4C | 1490700 | + | | upstream Pnuc_1426, SsrA-binding protein |
| m6A | 1494931 | - | | Pnuc_1423, inosine-5'-monophosphate dehydrogenase |
| m4C | 1496081 | + | | upstream Pnuc_1426, SsrA-binding protein |
| m6A | 1496168 | + | CTAYNNNNNNNNTRTC | upstream Pnuc_1426, SsrA-binding protein |
| m6A | 1496178 | - | GAYANNNNNNNNRTAG | Pnuc_1425, cyclase/dehydrase |
| m6A | 1496760 | + | | Pnuc_1426, SsrA-binding protein |
| m4C | 1496925 | + | | upstream Pnuc_1427, protein of unknown function DUF404 |
| m6A | 1498558 | + | | Pnuc_1428, protein of unknown function DUF403 |
| m4C | 1498741 | + | | Pnuc_1428, protein of unknown function DUF403 |
| m6A | 1499062 | - | | upstream Pnuc_1425, cyclase/dehydrase |
| m6A | 1504352 | - | | Pnuc_1433, phosphoenolpyruvate synthase |
| m6A | 1505598 | + | CTAYNNNNNNNNTRTC | Pnuc_1434, protein of unknown function DUF299 |
| m6A | 1505608 | - | GAYANNNNNNNNRTAG | Pnuc_1433, phosphoenolpyruvate synthase |
| m4C | 1505738 | + | | Pnuc_1434, protein of unknown function DUF299 |
| m4C | 1507547 | + | | upstream Pnuc_1452, methionine aminopeptidase, type I |
| m4C | 1508926 | - | | Pnuc_1438, lipid-A-disaccharide synthase |
| m6A | 1517912 | + | | upstream Pnuc_1452, methionine aminopeptidase, type I |
| m6A | 1519356 | + | CTAYNNNNNNNNTRTC | upstream Pnuc_1452, methionine aminopeptidase, type I |
| m6A | 1519366 | - | GAYANNNNNNNNRTAG | Pnuc_1448, ribosome recycling factor |
| m6A | 1520796 | - | | Pnuc_1450, translation elongation factor Ts (EF-Ts) |
| m6A | 1520804 | + | | upstream Pnuc_1452, methionine aminopeptidase, type I |
| m4C | 1523178 | + | | Pnuc_1452, methionine aminopeptidase, type I |
| m6A | 1526171 | + | | upstream Pnuc_1458, 2,3,4,5-tetrahydropyridine-2,6-dicarboxylate N-succinyltransferase |
| m6A | 1526426 | + | | upstream Pnuc_1458, 2,3,4,5-tetrahydropyridine-2,6-dicarboxylate N-succinyltransferase |
| m6A | 1528113 | - | | Pnuc_1455, DNA ligase, NAD-dependent |
| m4C | 1528219 | + | | upstream Pnuc_1458, 2,3,4,5-tetrahydropyridine-2,6-dicarboxylate N-succinyltransferase |
| m6A | 1530405 | - | | Pnuc_1457, chromosome segregation protein SMC |
| m6A | 1548614 | - | | upstream Pnuc_1468, lipoprotein releasing system, transmembrane protein, LolC/E family |
| m6A | 1548948 | - | | upstream Pnuc_1468, lipoprotein releasing system, transmembrane protein, LolC/E family |
| m6A | 1552414 | + | GAYANNNNNNNNRTAG | upstream Pnuc_1477, peptide deformylase |
| m6A | 1552424 | - | CTAYNNNNNNNNTRTC | Pnuc_1473, hypothetical protein |
| m4C | 1554275 | - | | Pnuc_1476, cysteine synthase A |
| m4C | 1555954 | + | | upstream Pnuc_1479, Ferredoxin--NADP(+) reductase |
| m6A | 1557612 | + | CTAYNNNNNNNNTRTC | Pnuc_1480, protein of unknown function DUF81 |
| m6A | 1557622 | - | GAYANNNNNNNNRTAG | upstream Pnuc_1478, disulfide bond formation protein DsbB |
| m6A | 1558540 | - | | upstream Pnuc_1478, disulfide bond formation protein DsbB |
| m6A | 1560951 | + | | Pnuc_1483, sulfate adenylyltransferase subunit 1 |
| m6A | 1561444 | + | | Pnuc_1484, hypothetical protein |
| m6A | 1561655 | + | | Pnuc_1484, hypothetical protein |
| m6A | 1564637 | + | | upstream Pnuc_1496, aminotransferase |
| m6A | 1568411 | + | | upstream Pnuc_1496, aminotransferase |
| m6A | 1570108 | + | | upstream Pnuc_1496, aminotransferase |
| m6A | 1570379 | - | | Pnuc_1495, Excinuclease ABC subunit B |

| | | | | |
|---|---|---|---|---|
| m4C | 1570421 | - | | Pnuc_1495, Excinuclease ABC subunit B |
| m6A | 1570842 | + | CTAYNNNNNNNTRTC | upstream Pnuc_1496, aminotransferase |
| m6A | 1570852 | - | GAYANNNNNNNRTAG | Pnuc_1495, Excinuclease ABC subunit B |
| m4C | 1572033 | - | | Pnuc_1495, Excinuclease ABC subunit B |
| m6A | 1572464 | - | | upstream Pnuc_1495, Excinuclease ABC subunit B |
| m4C | 1572536 | + | | upstream Pnuc_1496, aminotransferase |
| m4C | 1572718 | - | | upstream Pnuc_1495, Excinuclease ABC subunit B |
| m4C | 1575927 | - | | upstream Pnuc_1495, Excinuclease ABC subunit B |
| m4C | 1581403 | - | | Pnuc_1506, isocitrate lyase |
| m6A | 1582960 | + | GAYANNNNNNNRTAG | upstream Pnuc_1528, conserved hypothetical protein |
| m6A | 1582970 | - | CTAYNNNNNNNTRTC | Pnuc_1507, major facilitator superfamily MFS_1 |
| m6A | 1591897 | + | CTAYNNNNNNNTRTC | upstream Pnuc_1528, conserved hypothetical protein |
| m6A | 1591907 | - | GAYANNNNNNNRTAG | Pnuc_1518, heme exporter protein CcmA |
| m4C | 1595424 | + | | upstream Pnuc_1528, conserved hypothetical protein |
| m6A | 1599496 | + | CTAYNNNNNNNTRTC | Pnuc_1529, histone deacetylase superfamily |
| m6A | 1599506 | - | GAYANNNNNNNRTAG | upstream Pnuc_1527, hypothetical protein |
| m6A | 1601203 | + | GAYANNNNNNNRTAG | upstream Pnuc_1534, alkyl hydroperoxide reductase/ Thiol specific antioxidant/ Mal allergen |
| m6A | 1601213 | - | CTAYNNNNNNNTRTC | Pnuc_1531, putative lipoprotein |
| m6A | 1602159 | + | | upstream Pnuc_1534, alkyl hydroperoxide reductase/ Thiol specific antioxidant/ Mal allergen |
| m6A | 1602318 | - | | Pnuc_1533, hypothetical protein |
| m6A | 1606433 | + | GAYANNNNNNNRTAG | upstream Pnuc_1544, transcriptional regulator, GntR family |
| m6A | 1606443 | - | CTAYNNNNNNNTRTC | Pnuc_1538, TRAP C4-dicarboxylate transport system permease DctM subunit |
| m6A | 1609714 | - | | Pnuc_1540, short-chain dehydrogenase/reductase SDR |
| m4C | 1613302 | + | | upstream Pnuc_1544, transcriptional regulator, GntR family |
| m6A | 1616212 | - | | upstream Pnuc_1545, class II aldolase/adducin family protein |
| m6A | 1617376 | - | | upstream Pnuc_1545, class II aldolase/adducin family protein |
| m6A | 1617388 | + | GAYANNNNNNNRTAG | Pnuc_1547, protein of unknown function DUF6, transmembrane |
| m6A | 1617398 | - | CTAYNNNNNNNTRTC | upstream Pnuc_1545, class II aldolase/adducin family protein |
| m6A | 1620862 | + | | upstream Pnuc_1553, transcriptional regulator, AsnC family |
| m4C | 1623161 | + | | upstream Pnuc_1553, transcriptional regulator, AsnC family |
| m6A | 1625171 | + | | upstream Pnuc_1553, transcriptional regulator, AsnC family |
| m6A | 1628517 | + | GAYANNNNNNNRTAG | upstream Pnuc_1560, transcriptional regulator, LysR family |
| m6A | 1628527 | - | CTAYNNNNNNNTRTC | Pnuc_1556, Enoyl-CoA hydratase/isomerase |
| m6A | 1638119 | + | CTAYNNNNNNNTRTC | Pnuc_1564, FMN-binding domain protein |
| m6A | 1638129 | - | GAYANNNNNNNRTAG | upstream Pnuc_1563, conserved hypothetical protein |
| m6A | 1640591 | - | | upstream Pnuc_1567, Hydroxypyruvate isomerase |
| m6A | 1647158 | + | | upstream Pnuc_1581, hypothetical protein |
| m6A | 1647640 | - | | Pnuc_1575, anti-ECFsigma factor, ChrR |
| m6A | 1651758 | - | | Pnuc_1579, protein tyrosine phosphatase |
| m6A | 1652331 | + | | upstream Pnuc_1581, hypothetical protein |
| m6A | 1654135 | - | | Pnuc_1583, protein of unknown function DUF344 |
| m6A | 1654835 | - | | Pnuc_1584, lyso-ornithine lipid acyltransferase |
| m6A | 1657698 | + | | Pnuc_1588, ornithine-acyl[acyl carrier protein]N-acyltransferase |
| m6A | 1658012 | + | GAYANNNNNNNRTAG | Pnuc_1588, ornithine-acyl[acyl carrier protein]N-acyltransferase |

| | | | | |
|---|---|---|---|---|
| m6A | 1658022 | - | CTAYNNNNNNNTRTC | upstream Pnuc_1584, lyso-ornithine lipid acyltransferase |
| m6A | 1660810 | + | CTAYNNNNNNNTRTC | upstream Pnuc_1595, MgtC/SapB transporter |
| m6A | 1660820 | - | GAYANNNNNNNRTAG | Pnuc_1591, MgtC/SapB transporter |
| m6A | 1667562 | + | | upstream Pnuc_1606, short-chain dehydrogenase/reductase SDR |
| m4C | 1668273 | + | | upstream Pnuc_1606, short-chain dehydrogenase/reductase SDR |
| m4C | 1668542 | + | | upstream Pnuc_1606, short-chain dehydrogenase/reductase SDR |
| m6A | 1670909 | + | | upstream Pnuc_1606, short-chain dehydrogenase/reductase SDR |
| m4C | 1673647 | - | | Pnuc_1604, conserved hypothetical protein |
| m6A | 1676774 | - | | Pnuc_1608, deoxyribodipyrimidine photolyase-related protein |
| m6A | 1677038 | - | | Pnuc_1609, conserved hypothetical protein |
| m4C | 1678665 | + | | upstream Pnuc_1622, hypothetical protein |
| m6A | 1690825 | - | | upstream Pnuc_1621, Mg2+ transporter protein, CorA family protein |
| m4C | 1691114 | + | | upstream Pnuc_1627, putative phosphohistidine phosphatase, SixA |
| m4C | 1697565 | + | | upstream Pnuc_1636, transcriptional regulator, GntR family |
| m4C | 1698510 | - | | Pnuc_1631, Hydroxypyruvate isomerase |
| m6A | 1699838 | + | CTAYNNNNNNNTRTC | upstream Pnuc_1636, transcriptional regulator, GntR family |
| m6A | 1699848 | - | GAYANNNNNNNRTAG | Pnuc_1633, type III effector Hrp-dependent outer domain protein |
| m6A | 1701867 | + | | upstream Pnuc_1636, transcriptional regulator, GntR family |
| m6A | 1702881 | + | | upstream Pnuc_1636, transcriptional regulator, GntR family |
| m4C | 1705118 | - | | upstream Pnuc_1637, NAD-dependent epimerase/dehydratase |
| m6A | 1705954 | + | CTAYNNNNNNNTRTC | Pnuc_1638, protein of unknown function DUF6, transmembrane |
| m6A | 1705964 | - | GAYANNNNNNNRTAG | upstream Pnuc_1637, NAD-dependent epimerase/dehydratase |
| m6A | 1709911 | - | | upstream Pnuc_1641, conserved hypothetical protein |
| m6A | 1712078 | + | CTAYNNNNNNNTRTC | upstream Pnuc_1649, hypothetical protein |
| m6A | 1712822 | + | CTAYNNNNNNNTRTC | upstream Pnuc_1649, hypothetical protein |
| m6A | 1712832 | - | GAYANNNNNNNRTAG | Pnuc_1647, DNA polymerase III, epsilon subunit |
| m6A | 1722847 | + | | Pnuc_1660, putative uncharacterized conserved protein |
| m4C | 1725615 | + | | upstream Pnuc_1664, spermine/spermidine N-acetyltransferase |
| m6A | 1726717 | + | | upstream Pnuc_1664, spermine/spermidine N-acetyltransferase |
| m6A | 1729661 | - | | Pnuc_1666, two component transcriptional regulator, winged helix family |
| m6A | 1730416 | - | | Pnuc_1667, osmosensitive K+ channel signal transduction histidine kinase |
| m4C | 1731262 | - | | Pnuc_1667, osmosensitive K+ channel signal transduction histidine kinase |
| m6A | 1736729 | + | | upstream Pnuc_1678, putative transcriptional regulator, MerR family |
| m6A | 1736959 | + | GAYANNNNNNNRTAG | upstream Pnuc_1678, putative transcriptional regulator, MerR family |
| m6A | 1736969 | - | CTAYNNNNNNNTRTC | Pnuc_1670, potassium-transporting ATPase, A subunit |
| m6A | 1737694 | - | | Pnuc_1670, potassium-transporting ATPase, A subunit |
| m4C | 1737703 | + | | upstream Pnuc_1678, putative transcriptional regulator, MerR family |
| m6A | 1738403 | + | CTAYNNNNNNNTRTC | upstream Pnuc_1678, putative transcriptional regulator, MerR family |
| m6A | 1738413 | - | GAYANNNNNNNRTAG | Pnuc_1672, Heavy metal transport/detoxification protein |
| m6A | 1740542 | - | | Pnuc_1676, conserved secreted protein with internal repeats |
| m6A | 1741194 | - | | Pnuc_1677, heavy metal translocating P-type ATPase |
| m4C | 1748197 | + | | upstream Pnuc_1688, hypothetical protein |
| m6A | 1748827 | + | GAYANNNNNNNRTAG | upstream Pnuc_1688, hypothetical protein |
| m6A | 1748837 | - | CTAYNNNNNNNTRTC | Pnuc_1681, efflux transporter, RND family, MFP subunit |

| | | | | |
|---|---|---|---|---|
| m4C | 1749518 | + | | upstream Pnuc_1688, hypothetical protein |
| m6A | 1749645 | + | | upstream Pnuc_1688, hypothetical protein |
| m6A | 1750197 | + | | upstream Pnuc_1688, hypothetical protein |
| m6A | 1756678 | + | | Pnuc_1692, hypothetical protein |
| m6A | 1758014 | + | GAYANNNNNNNNRTAG | upstream Pnuc_1702, O-sialoglycoprotein endopeptidase |
| m6A | 1758024 | - | CTAYNNNNNNNNTRTC | Pnuc_1694, phage integrase family protein |
| m4C | 1761275 | + | | upstream Pnuc_1702, O-sialoglycoprotein endopeptidase |
| m4C | 1764648 | - | | Pnuc_1698, RNA polymerase, sigma 70 subunit, RpoD |
| m6A | 1765462 | - | | upstream Pnuc_1698, RNA polymerase, sigma 70 subunit, RpoD |
| m6A | 1765623 | - | | upstream Pnuc_1698, RNA polymerase, sigma 70 subunit, RpoD |
| m6A | 1765631 | - | | upstream Pnuc_1698, RNA polymerase, sigma 70 subunit, RpoD |
| m4C | 1765714 | - | | Pnuc_1699, DNA primase |
| m4C | 1768466 | - | | upstream Pnuc_1701, SSU ribosomal protein S21P |
| m4C | 1768677 | - | | upstream Pnuc_1701, SSU ribosomal protein S21P |
| m6A | 1774025 | + | | Pnuc_1707, Rieske (2Fe-2S) domain protein |
| m4C | 1774605 | - | | upstream Pnuc_1706, Exonuclease VII, small subunit |
| m4C | 1776410 | + | | upstream Pnuc_1712, putative transmembrane protein |
| m6A | 1782114 | - | | upstream Pnuc_1711, DNA polymerase I |
| m4C | 1782990 | - | | Pnuc_1715, ATP-dependent DNA helicase UvrD |
| m4C | 1783268 | - | | Pnuc_1715, ATP-dependent DNA helicase UvrD |
| m4C | 1784851 | + | | Pnuc_1716, valyl-tRNA synthetase |
| m6A | 1784984 | + | | Pnuc_1716, valyl-tRNA synthetase |
| m4C | 1786759 | - | | upstream Pnuc_1715, ATP-dependent DNA helicase UvrD |
| m6A | 1787865 | + | | Pnuc_1717, UDP-glucose pyrophosphorylase |
| m6A | 1787955 | - | | upstream Pnuc_1715, ATP-dependent DNA helicase UvrD |
| m6A | 1788038 | - | | upstream Pnuc_1715, ATP-dependent DNA helicase UvrD |
| m4C | 1788326 | - | | upstream Pnuc_1715, ATP-dependent DNA helicase UvrD |
| m4C | 1789887 | - | | Pnuc_1719, alanyl-tRNA synthetase |
| m6A | 1792308 | - | | upstream Pnuc_1719, alanyl-tRNA synthetase |
| m6A | 1793894 | + | | Pnuc_1722, glutaminyl-tRNA synthetase |
| m6A | 1797750 | + | CTAYNNNNNNNNTRTC | Pnuc_1725, acetylornithine aminotransferase apoenzyme |
| m6A | 1797760 | - | GAYANNNNNNNNRTAG | upstream Pnuc_1723, NAD-dependent epimerase/dehydratase |
| m4C | 1799103 | + | | upstream Pnuc_1733, DNA replication and repair protein RadC |
| m4C | 1799398 | - | | Pnuc_1726, amino acid/amide ABC transporter ATP-binding protein 2, HAAT family |
| m6A | 1803154 | - | | Pnuc_1730, amino acid/amide ABC transporter substrate-binding protein, HAAT family |
| m4C | 1805662 | - | | upstream Pnuc_1732, peptidylprolyl isomerase, FKBP-type |
| m4C | 1806580 | + | | upstream Pnuc_1739, FMN adenylyltransferase / riboflavin kinase |
| m6A | 1807138 | - | | Pnuc_1736, fatty acid desaturase |
| m6A | 1807884 | - | | Pnuc_1737, Fmu (Sun) domain protein |
| m6A | 1808749 | - | | Pnuc_1737, Fmu (Sun) domain protein |
| m6A | 1808930 | - | | upstream Pnuc_1737, Fmu (Sun) domain protein |
| m6A | 1809803 | + | | Pnuc_1739, FMN adenylyltransferase / riboflavin kinase |
| m6A | 1815162 | - | | upstream Pnuc_1738, formyltetrahydrofolate-dependent phosphoribosylglycinamide formyltransferase |
| m4C | 1815678 | - | | upstream Pnuc_1738, formyltetrahydrofolate-dependent phosphoribosylglycinamide formyltransferase |

| | | | | |
|---|---|---|---|---|
| m4C | 1816159 | - | | upstream Pnuc_1738, formyltetrahydrofolate-dependent phosphoribosylglycinamide formyltransferase |
| m6A | 1819565 | - | | upstream Pnuc_1747, hypothetical protein |
| m6A | 1821583 | - | | Pnuc_1748, ATP-dependent Clp protease ATP-binding subunit ClpA |
| m4C | 1822297 | + | | upstream Pnuc_1750, cold-shock DNA-binding protein family |
| m4C | 1825922 | - | | Pnuc_1754, ornithine carbamoyltransferase |
| m6A | 1826804 | + | | upstream Pnuc_1757, integral membrane protein MviN |
| m4C | 1827004 | - | | upstream Pnuc_1756, SSU ribosomal protein S20P |
| m4C | 1827065 | - | | upstream Pnuc_1756, SSU ribosomal protein S20P |
| m4C | 1827074 | - | | upstream Pnuc_1756, SSU ribosomal protein S20P |
| m4C | 1827283 | - | | upstream Pnuc_1756, SSU ribosomal protein S20P |
| m4C | 1827964 | + | | Pnuc_1757, integral membrane protein MviN |
| m6A | 1828079 | - | | upstream Pnuc_1756, SSU ribosomal protein S20P |
| m4C | 1830997 | + | | upstream Pnuc_1762, protein of unknown function UPF0118 |
| m6A | 1832665 | + | CTAYNNNNNNNNTRTC | Pnuc_1762, protein of unknown function UPF0118 |
| m6A | 1832675 | - | GAYANNNNNNNNRTAG | upstream Pnuc_1761, phosphoribosylformylglycinamidine cyclo-ligase |
| m4C | 1833663 | + | | Pnuc_1763, regulatory inactivation of DnaA Hda protein |
| m6A | 1836536 | + | | Pnuc_1766, 2-amino-4-hydroxy-6-hydroxymethyldihydropteridine pyrophosphokinase |
| m4C | 1837776 | - | | Pnuc_1768, chaperone protein DnaJ |
| m4C | 1837887 | - | | Pnuc_1768, chaperone protein DnaJ |
| m6A | 1840618 | + | | upstream Pnuc_1773, NAD(+) kinase |
| m4C | 1840727 | - | | upstream Pnuc_1769, chaperone protein DnaK |
| m6A | 1843285 | + | GAYANNNNNNNNRTAG | upstream Pnuc_1773, NAD(+) kinase |
| m6A | 1843295 | - | CTAYNNNNNNNNTRTC | Pnuc_1772, heat-inducible transcription repressor HrcA |
| m6A | 1843408 | - | | Pnuc_1772, heat-inducible transcription repressor HrcA |
| m6A | 1844781 | + | | Pnuc_1774, DNA replication and repair protein RecN |
| m4C | 1846378 | + | | upstream Pnuc_1776, membrane protein-like protein |
| m6A | 1850256 | + | | upstream Pnuc_1776, membrane protein-like protein |
| m4C | 1850611 | - | | upstream Pnuc_1775, (Glutamate--ammonia-ligase) adenylyltransferase |
| m4C | 1852602 | - | | upstream Pnuc_1775, (Glutamate--ammonia-ligase) adenylyltransferase |
| m6A | 1853226 | + | CTAYNNNNNNNNTRTC | Pnuc_1777, Nitrilase/cyanide hydratase and apolipoprotein N-acyltransferase |
| m6A | 1853236 | - | GAYANNNNNNNNRTAG | upstream Pnuc_1775, (Glutamate-ammonia-ligase) adenylyltransferase |
| m4C | 1855317 | + | | Pnuc_1778, microcin-processing peptidase 2, Unknown type peptidase, MEROPS family U62 |
| m6A | 1857619 | - | | upstream Pnuc_1780, ATP:cob(I)alamin adenosyltransferase |
| m4C | 1860209 | - | | upstream Pnuc_1780, ATP:cob(I)alamin adenosyltransferase |
| m6A | 1861824 | + | GAYANNNNNNNNRTAG | Pnuc_1784, alanine racemase domain protein |
| m6A | 1861834 | - | CTAYNNNNNNNNTRTC | upstream Pnuc_1780, ATP:cob(I)alamin adenosyltransferase |
| m6A | 1862840 | - | | upstream Pnuc_1780, ATP:cob(I)alamin adenosyltransferase |
| m6A | 1863024 | - | | Pnuc_1786, 4-hydroxybenzoate octaprenyltransferase |
| m4C | 1863093 | + | | upstream Pnuc_1788, S-adenosylmethionine-tRNA-ribosyltransferase-isomerase |
| m6A | 1865372 | + | GAYANNNNNNNNRTAG | upstream Pnuc_1788, S-adenosylmethionine-tRNA-ribosyltransferase-isomerase |
| m6A | 1865382 | - | CTAYNNNNNNNNTRTC | Pnuc_1787, ATP-dependent DNA helicase RecG |
| m4C | 1865556 | - | | Pnuc_1787, ATP-dependent DNA helicase RecG |
| m4C | 1865946 | + | | Pnuc_1788, S-adenosylmethionine-tRNA-ribosyltransferase-isomerase |
| m6A | 1866093 | + | | Pnuc_1788, S-adenosylmethionine-tRNA-ribosyltransferase-isomerase |

| | | | | |
|---|---|---|---|---|
| m4C | 1867492 | - | | upstream Pnuc_1787, ATP-dependent DNA helicase RecG |
| m6A | 1867542 | + | CTAYNNNNNNNNTRTC | Pnuc_1789, tRNA-guanine transglycosylase |
| m6A | 1867552 | - | GAYANNNNNNNNRTAG | upstream Pnuc_1787, ATP-dependent DNA helicase RecG |
| m6A | 1868132 | + | | Pnuc_1790, protein translocase subunit yajC |
| m4C | 1872268 | + | | upstream Pnuc_1795, Glutathione S-transferase, N-terminal domain protein |
| m6A | 1874872 | + | GAYANNNNNNNNRTAG | upstream Pnuc_1797, alanine dehydrogenase/PNT domain protein |
| m6A | 1874879 | + | | upstream Pnuc_1797, alanine dehydrogenase/PNT domain protein |
| m6A | 1874882 | - | CTAYNNNNNNNNTRTC | Pnuc_1796, tRNA (5-methylaminomethyl-2-thiouridylate)-methyltransferase |
| m4C | 1875999 | + | | upstream Pnuc_1797, alanine dehydrogenase/PNT domain protein |
| m6A | 1876112 | + | | upstream Pnuc_1797, alanine dehydrogenase/PNT domain protein |
| m4C | 1878427 | + | | Pnuc_1799, NAD(P) transhydrogenase, beta subunit |
| m6A | 1879092 | - | | upstream Pnuc_1796, tRNA (5-methylaminomethyl-2-thiouridylate)-methyltransferase |
| m6A | 1880808 | + | GAYANNNNNNNNRTAG | upstream Pnuc_1802, hypothetical protein |
| m6A | 1880818 | - | CTAYNNNNNNNNTRTC | Pnuc_1801, glycosyl transferase, family 11 |
| m6A | 1881762 | + | CTAYNNNNNNNNTRTC | Pnuc_1802, hypothetical protein |
| m6A | 1881772 | - | GAYANNNNNNNNRTAG | upstream Pnuc_1801, glycosyl transferase, family 11 |
| m6A | 1882788 | - | | Pnuc_1804, conserved hypothetical protein |
| m6A | 1884638 | - | | Pnuc_1805, chaperonin GroEL |
| m6A | 1887649 | + | | Pnuc_1809, amino acid-binding domain sensor hybrid histidine kinase |
| m6A | 1887784 | + | | Pnuc_1809, amino acid-binding domain sensor hybrid histidine kinase |
| m6A | 1892173 | - | | Pnuc_1811, translation factor SUA5 |
| m6A | 1894654 | - | | Pnuc_1814, phosphoribosylaminoimidazole-succinocarboxamide synthase |
| m6A | 1894804 | + | GAYANNNNNNNNRTAG | upstream Pnuc_1818, branched chain amino acid aminotransferase apoenzyme |
| m6A | 1894814 | - | CTAYNNNNNNNNTRTC | Pnuc_1814, phosphoribosylaminoimidazole-succinocarboxamide synthase |
| m4C | 1895131 | - | | Pnuc_1815, fructose-bisphosphate aldolase |
| m4C | 1895880 | + | | upstream Pnuc_1818, branched chain amino acid aminotransferase apoenzyme |
| m4C | 1895881 | + | | upstream Pnuc_1818, branched chain amino acid aminotransferase apoenzyme |
| m4C | 1896166 | + | | upstream Pnuc_1818, branched chain amino acid aminotransferase apoenzyme |
| m6A | 1897475 | - | | Pnuc_1817, phosphoglycerate kinase |
| m6A | 1899165 | + | CTAYNNNNNNNNTRTC | Pnuc_1818, branched chain amino acid aminotransferase apoenzyme |
| m6A | 1899175 | - | GAYANNNNNNNNRTAG | upstream Pnuc_1817, phosphoglycerate kinase |
| m6A | 1902175 | - | | upstream Pnuc_1817, phosphoglycerate kinase |
| m4C | 1903822 | + | | upstream Pnuc_1824, GTP cyclohydrolase subunit MoaC |
| m6A | 1904068 | + | | Pnuc_1824, GTP cyclohydrolase subunit MoaC |
| m4C | 1914844 | + | | upstream Pnuc_1835, hypothetical protein |
| m6A | 1914998 | + | | upstream Pnuc_1835, hypothetical protein |
| m6A | 1916434 | + | GAYANNNNNNNNRTAG | Pnuc_1835, hypothetical protein |
| m6A | 1916444 | - | CTAYNNNNNNNNTRTC | upstream Pnuc_1834, conserved hypothetical protein |
| m6A | 1918050 | + | GAYANNNNNNNNRTAG | Pnuc_1838, hypothetical protein |
| m6A | 1918060 | - | CTAYNNNNNNNNTRTC | upstream Pnuc_1834, conserved hypothetical protein |
| m6A | 1921973 | + | GAYANNNNNNNNRTAG | upstream Pnuc_1847, secreted protein |
| m6A | 1921983 | - | CTAYNNNNNNNNTRTC | upstream Pnuc_1843, hypothetical protein |
| m6A | 1923748 | + | | upstream Pnuc_1847, secreted protein |
| m6A | 1924410 | + | | upstream Pnuc_1847, secreted protein |

| | | | | |
|---|---|---|---|---|
| m4C | 1927470 | + | | upstream Pnuc_1851, maleylacetoacetate isomerase |
| m6A | 1929095 | - | | Pnuc_1852, transferase hexapeptide repeat protein |
| m6A | 1931420 | - | | upstream Pnuc_1857, Rhodanese domain protein |
| m6A | 1934625 | - | | upstream Pnuc_1857, Rhodanese domain protein |
| m6A | 1935097 | + | GAYANNNNNNNRTAG | Pnuc_1861, CBS domain containing protein |
| m6A | 1935107 | - | CTAYNNNNNNNTRTC | upstream Pnuc_1857, Rhodanese domain protein |
| m6A | 1936096 | - | | upstream Pnuc_1857, Rhodanese domain protein |
| m6A | 1936488 | - | | upstream Pnuc_1857, Rhodanese domain protein |
| m6A | 1936868 | - | | upstream Pnuc_1857, Rhodanese domain protein |
| m6A | 1937416 | + | | Pnuc_1863, glycyl-tRNA synthetase alpha chain |
| m4C | 1937691 | + | | Pnuc_1863, glycyl-tRNA synthetase alpha chain |
| m6A | 1940032 | + | CTAYNNNNNNNTRTC | Pnuc_1865, D-alpha,beta-D-heptose 1,7-bisphosphate phosphatase |
| m6A | 1940042 | - | GAYANNNNNNNRTAG | upstream Pnuc_1857, Rhodanese domain protein |
| m6A | 1943461 | + | | upstream Pnuc_1871, aminoglycoside phosphotransferase |
| m6A | 1954638 | - | | upstream Pnuc_1870, Organic solvent tolerance protein |
| m6A | 1954913 | - | | upstream Pnuc_1870, Organic solvent tolerance protein |
| m6A | 1956898 | - | | Pnuc_1881, tyrosyl-tRNA synthetase |
| m4C | 1962595 | - | | upstream Pnuc_R0042, RNase P |
| m6A | 1968003 | + | CTAYNNNNNNNTRTC | upstream Pnuc_1900, L-threonine ammonia-lyase |
| m6A | 1968013 | - | GAYANNNNNNNRTAG | Pnuc_1892, aspartyl-tRNA synthetase |
| m6A | 1968185 | + | GAYANNNNNNNRTAG | upstream Pnuc_1900, L-threonine ammonia-lyase |
| m6A | 1968195 | - | CTAYNNNNNNNTRTC | Pnuc_1892, aspartyl-tRNA synthetase |
| m4C | 1968481 | + | | upstream Pnuc_1900, L-threonine ammonia-lyase |
| m6A | 1968822 | + | | upstream Pnuc_1900, L-threonine ammonia-lyase |
| m4C | 1969703 | + | | upstream Pnuc_1900, L-threonine ammonia-lyase |
| m6A | 1974979 | + | GAYANNNNNNNRTAG | upstream Pnuc_1900, L-threonine ammonia-lyase |
| m6A | 1974989 | - | CTAYNNNNNNNTRTC | Pnuc_1899, FAD linked oxidase domain protein |
| m6A | 1981269 | - | | upstream Pnuc_1899, FAD linked oxidase domain protein |
| m4C | 1982013 | - | | Pnuc_1903, single-strand binding protein |
| m6A | 1982693 | + | | upstream Pnuc_1905, Excinuclease ABC subunit A |
| m4C | 1983946 | - | | upstream Pnuc_1904, major facilitator superfamily MFS_1 |
| m6A | 1985406 | - | | upstream Pnuc_1904, major facilitator superfamily MFS_2 |
| m6A | 1988342 | + | | Pnuc_1907, KpsF/GutQ family protein |
| m4C | 1990900 | - | | upstream Pnuc_1906, Kef-type potassium/proton antiporter, CPA2 family |
| m4C | 1994023 | - | | upstream Pnuc_1906, Kef-type potassium/proton antiporter, CPA2 family |
| m6A | 1994031 | + | | Pnuc_1915, Uncharacterized P-loop ATPase protein UPF0042 |
| m4C | 1998579 | - | | upstream Pnuc_1917, DNA-(apurinic or apyrimidinic site) lyase |
| m6A | 1999753 | + | GAYANNNNNNNRTAG | Pnuc_1920, ribose-phosphate pyrophosphokinase |
| m6A | 1999763 | - | CTAYNNNNNNNTRTC | upstream Pnuc_1917, DNA-(apurinic or apyrimidinic site) lyase |
| m4C | 1999808 | + | | Pnuc_1920, ribose-phosphate pyrophosphokinase |
| m4C | 2000189 | + | | upstream Pnuc_1921, LSU ribosomal protein L25P |
| m6A | 2000287 | - | | upstream Pnuc_1917, DNA-(apurinic or apyrimidinic site) lyase |
| m6A | 2000847 | + | | Pnuc_1921, LSU ribosomal protein L25P |
| m4C | 2002810 | - | | Pnuc_1925, putative methyltransferase |

| | | | | |
|---|---|---|---|---|
| m4C | 2004211 | - | | Pnuc_1926, peptidase M16 domain protein |
| m6A | 2006000 | + | | Pnuc_1928, signal recognition particle-docking protein FtsY |
| m6A | 2008414 | + | GAYANNNNNNNRTAG | upstream Pnuc_1935, conserved hypothetical protein |
| m6A | 2008424 | - | CTAYNNNNNNNTRTC | Pnuc_1931, protoheme IX farnesyltransferase |
| m6A | 2008780 | + | | upstream Pnuc_1935, conserved hypothetical protein |
| m6A | 2010428 | - | | Pnuc_1933, putative transmembrane protein |
| m6A | 2012994 | - | | upstream Pnuc_1936, cytochrome c oxidase, subunit III |
| m4C | 2013806 | - | | upstream Pnuc_1938, cytochrome c oxidase assembly protein CtaG/Cox11 |
| m6A | 2014025 | - | | Pnuc_1939, Cytochrome-c oxidase |
| m6A | 2014353 | - | | Pnuc_1939, Cytochrome-c oxidase |
| m6A | 2014371 | + | CTAYNNNNNNNTRTC | upstream Pnuc_1942, phosphoribosyltransferase |
| m6A | 2014381 | - | GAYANNNNNNNRTAG | Pnuc_1939, Cytochrome-c oxidase |
| m6A | 2014641 | + | GAYANNNNNNNRTAG | upstream Pnuc_1942, phosphoribosyltransferase |
| m6A | 2014651 | - | CTAYNNNNNNNTRTC | Pnuc_1939, Cytochrome-c oxidase |
| m6A | 2019976 | + | | upstream Pnuc_1948, phosphoglycerate mutase |
| m4C | 2020099 | + | | upstream Pnuc_1948, phosphoglycerate mutase |
| m4C | 2020195 | + | | upstream Pnuc_1948, phosphoglycerate mutase |
| m6A | 2021719 | + | | Pnuc_1948, phosphoglycerate mutase |
| m4C | 2023023 | + | | Pnuc_1949, carboxyl-terminal protease |
| m4C | 2023875 | + | | Pnuc_1950, [sulfur carrier protein ThiS]adenylyltransferase |
| m6A | 2035852 | + | | upstream Pnuc_1967, biotin--acetyl-CoA-carboxylase ligase |
| m4C | 2039264 | - | | upstream Pnuc_1969, rfaE bifunctional protein |
| m4C | 2040106 | - | | upstream Pnuc_1969, rfaE bifunctional protein |
| m4C | 2045187 | + | | upstream Pnuc_1978, conserved hypothetical protein |
| m4C | 2046520 | + | | upstream Pnuc_1978, conserved hypothetical protein |
| m6A | 2050688 | + | | upstream Pnuc_1981 |
| m6A | 2054081 | - | | upstream Pnuc_1984, Alcohol dehydrogenase, zinc-binding domain protein |
| m6A | 2054201 | + | CTAYNNNNNNNTRTC | Pnuc_1985, hypothetical protein |
| m4C | 2055912 | + | | upstream Pnuc_1996, protein of unknown function DUF185 |
| m4C | 2057298 | - | | Pnuc_1988, acriflavin resistance protein |
| m4C | 2060192 | - | | Pnuc_1990, RND efflux system, outer membrane lipoprotein, NodT family |
| m4C | 2060483 | + | | upstream Pnuc_1996, protein of unknown function DUF185 |
| m6A | 2063777 | + | CTAYNNNNNNNTRTC | upstream Pnuc_1996, protein of unknown function DUF185 |
| m6A | 2063787 | - | GAYANNNNNNNRTAG | Pnuc_1992, UDP-N-acetylglucosamine pyrophosphorylase / glucosamine-1-phosphate N-acetyltransferase |
| m6A | 2066312 | + | GAYANNNNNNNRTAG | upstream Pnuc_1996, protein of unknown function DUF185 |
| m6A | 2066322 | - | CTAYNNNNNNNTRTC | Pnuc_1994, dihydroneopterin aldolase |
| m6A | 2072803 | - | | upstream Pnuc_1999, Lytic transglycosylase, catalytic |
| m6A | 2074411 | - | | Pnuc_2002, adenosylhomocysteinase |
| m6A | 2077839 | - | | upstream Pnuc_2003, methionine adenosyltransferase |
| m4C | 2081025 | + | | Pnuc_2008, exodeoxyribonuclease III Xth |
| m6A | 2081485 | + | | upstream Pnuc_2021, rod shape-determining protein MreB |
| m6A | 2082597 | + | | upstream Pnuc_2021, rod shape-determining protein MreB |
| m6A | 2085911 | - | | upstream Pnuc_2013, N-acetylglutamate kinase |
| m6A | 2085972 | + | GAYANNNNNNNRTAG | upstream Pnuc_2021, rod shape-determining protein MreB |

| | | | | |
|---|---|---|---|---|
| m6A | 2085982 | - | CTAYNNNNNNNTRTC | Pnuc_2014, transcriptional regulator, TraR/DksA family |
| m4C | 2089737 | + | | upstream Pnuc_2021, rod shape-determining protein MreB |
| m4C | 2091198 | - | | Pnuc_2019, aspartyl/glutamyl-tRNA(Asn/Gln) amidotransferase subunit A |
| m4C | 2092910 | + | | Pnuc_2021, rod shape-determining protein MreB |
| m6A | 2093215 | - | | upstream Pnuc_2020, aspartyl/glutamyl-tRNA(Asn/Gln) amidotransferase subunit C |
| m4C | 2094451 | + | | Pnuc_2022, rod shape-determining protein MreC |
| m4C | 2094517 | + | | Pnuc_2022, rod shape-determining protein MreC |
| m4C | 2095554 | + | | Pnuc_2024, cell elongation-specific peptidoglycan D,D-transpeptidase |
| m6A | 2098007 | + | | Pnuc_2025, cell elongation-specific peptidoglycan biosynthesis regulator RodA |
| m6A | 2098082 | - | | upstream Pnuc_2020, aspartyl/glutamyl-tRNA(Asn/Gln) amidotransferase subunit C |
| m4C | 2098226 | + | | Pnuc_2025, cell elongation-specific peptidoglycan biosynthesis regulator RodA |
| m4C | 2098267 | + | | upstream Pnuc_2028, Aspartyl/Asparaginyl beta-hydroxylase |
| m6A | 2100138 | - | | upstream Pnuc_2027, conserved hypothetical protein |
| m6A | 2100782 | - | | upstream Pnuc_2029, protein of unknown function DUF167 |
| m4C | 2101468 | + | | Pnuc_2031, lipid A biosynthesis acyltransferase |
| m6A | 2102026 | + | CTAYNNNNNNNTRTC | Pnuc_2031, lipid A biosynthesis acyltransferase |
| m6A | 2102036 | - | GAYANNNNNNNRTAG | upstream Pnuc_2029, protein of unknown function DUF167 |
| m4C | 2102300 | - | | upstream Pnuc_2029, protein of unknown function DUF167 |
| m6A | 2104651 | + | | upstream Pnuc_2035, ribosomal large subunit pseudouridine synthase F |
| m4C | 2108072 | - | | upstream Pnuc_2034, sodium:dicarboxylate symporter |
| m4C | 2108776 | + | | upstream Pnuc_2038, GCN5-related N-acetyltransferase |
| m6A | 2109837 | - | | upstream Pnuc_2037, conserved hypothetical protein |
| m6A | 2110796 | + | | Pnuc_2040, MltA-interacting MipA family protein |
| m4C | 2118082 | + | | upstream Pnuc_2056, protein of unknown function DUF897 |
| m6A | 2118474 | + | GAYANNNNNNNRTAG | upstream Pnuc_2056, protein of unknown function DUF897 |
| m6A | 2118484 | - | CTAYNNNNNNNTRTC | Pnuc_2049, hypothetical protein |
| m6A | 2121455 | + | | upstream Pnuc_2056, protein of unknown function DUF897 |
| m6A | 2122486 | - | | Pnuc_2054, CHRD domain containing protein |
| m6A | 2127286 | - | | Pnuc_2060, Patatin |
| m4C | 2128344 | - | | Pnuc_2060, Patatin |
| m4C | 2128565 | + | | upstream Pnuc_2062, hypothetical protein |
| m6A | 2128606 | + | | upstream Pnuc_2062, hypothetical protein |
| m6A | 2129286 | - | | upstream Pnuc_2060, Patatin |
| m6A | 2133944 | + | GAYANNNNNNNRTAG | Pnuc_2065, hypothetical protein |
| m6A | 2133954 | - | CTAYNNNNNNNTRTC | upstream Pnuc_2064, hypothetical protein |
| m6A | 2141211 | - | | Pnuc_2073, protein of unknown function DUF181 |
| m6A | 2141500 | + | | upstream Pnuc_2080, DNA protecting protein DprA |
| m6A | 2141510 | - | | Pnuc_2073, protein of unknown function DUF181 |
| m6A | 2144251 | + | | upstream Pnuc_2080, DNA protecting protein DprA |
| m6A | 2145421 | - | | Pnuc_2076, putative periplasmic ligand-binding sensor protein |
| m6A | 2146256 | + | CTAYNNNNNNNTRTC | upstream Pnuc_2080, DNA protecting protein DprA |
| m6A | 2146266 | - | GAYANNNNNNNRTAG | Pnuc_2077, sun protein |
| m4C | 2149663 | - | | upstream Pnuc_2079, peptide deformylase |
| m4C | 2151298 | + | | Pnuc_2081, DNA topoisomerase III |

| | | | | |
|---|---|---|---|---|
| m4C | 2152087 | - | | upstream Pnuc_2079, peptide deformylase |
| m4C | 2155660 | - | | Pnuc_2084, tRNA modification GTPase trmE |

| 26°C | | | | | |
|---|---|---|---|---|---|
| **Methylation** | **Location** | **Strand** | **Motif** | **Gene / Intergenic region** | **Count** |
| m6A | 901 | - | | upstream Pnuc_2088, LSU ribosomal protein L34P | 769(+) |
| m4C | 1132 | + | | Pnuc_0001,chromosomal replication initiator protein DnaA | 735(-) |
| m4C | 6952 | + | | Pnuc_0007, amino acid/amide ABC transporter ATP-binding protein 2, HAAT family | **Total** = 1504 (m6A + m4C) |
| m4C | 7696 | - | | upstream Pnuc_2088, LSU ribosomal protein L34P | |
| m4C | 14758 | + | | Pnuc_0015, glucose inhibited division protein A | 341(+) |
| m4C | 19205 | - | | upstream Pnuc_0013, acyl-CoA dehydrogenase domain protein | 343(-) |
| m4C | 22089 | - | | upstream Pnuc_0013, acyl-CoA dehydrogenase domain protein | **Total** = 684 (m6A) |
| m4C | 22150 | - | | upstream Pnuc_0013, acyl-CoA dehydrogenase domain protein | |
| m4C | 26361 | + | | Pnuc_0028, uroporphyrinogen decarboxylase | 428(+) |
| m6A | 26488 | + | | Pnuc_0028, uroporphyrinogen decarboxylase | 392(-) |
| m6A | 28723 | + | | Pnuc_0029, replication restart DNA helicase PriA | **Total** = 820 (m4C) |
| m4C | 31637 | - | | Pnuc_0032, fatty acid desaturase | |
| m6A | 32663 | + | | Pnuc_0033, cytochrome c, class I | Methylated genes = 512 (24.52%) |
| m4C | 33233 | + | | upstream Pnuc_0035, phosphoheptose isomerase | |
| m6A | 35133 | + | | Pnuc_0036, transport-associated protein | |
| m4C | 37437 | - | | upstream Pnuc_0038, GCN5-related N-acetyltransferase | |
| m6A | 37807 | + | CTAYNNNNNNNNTRTC | Pnuc_R0002, 16S ribosomal RNA | |
| m6A | 37817 | - | GAYANNNNNNNNRTAG | upstream Pnuc_0038, GCN5-related N-acetyltransferase | |
| m4C | 39095 | - | | upstream Pnuc_0038, GCN5-related N-acetyltransferase | |
| m4C | 40789 | - | | upstream Pnuc_0038, GCN5-related N-acetyltransferase | |
| m4C | 44092 | + | | Pnuc_0041, transcription antitermination protein nusG | |
| m4C | 44677 | - | | upstream Pnuc_0038, GCN5-related N-acetyltransferase | |
| m4C | 47634 | + | | Pnuc_0046, DNA-directed RNA polymerase subunit beta | |
| m4C | 47980 | - | | upstream Pnuc_0038, GCN5-related N-acetyltransferase | |
| m4C | 47982 | - | | upstream Pnuc_0038, GCN5-related N-acetyltransferase | |
| m4C | 49472 | - | | upstream Pnuc_0038, GCN5-related N-acetyltransferase | |
| m4C | 52984 | + | | Pnuc_0047, DNA-directed RNA polymerase subunit beta | |
| m4C | 53025 | + | | Pnuc_0047, DNA-directed RNA polymerase subunit beta | |
| m4C | 54887 | + | | Pnuc_0047, DNA-directed RNA polymerase subunit beta | |
| m4C | 57126 | - | | upstream Pnuc_0038, GCN5-related N-acetyltransferase | |
| m4C | 57143 | - | | upstream Pnuc_0038, GCN5-related N-acetyltransferase | |
| m4C | 58975 | + | | Pnuc_0050, translation elongation factor 2 (EF-2/EF-G) | |
| m4C | 59477 | - | | upstream Pnuc_0038, GCN5-related N-acetyltransferase | |
| m4C | 64043 | - | | upstream Pnuc_0038, GCN5-related N-acetyltransferase | |
| m4C | 65373 | + | | Pnuc_0061, LSU ribosomal protein L29P | |
| m4C | 65930 | - | | upstream Pnuc_0038, GCN5-related N-acetyltransferase | |
| m4C | 65949 | - | | upstream Pnuc_0038, GCN5-related N-acetyltransferase | |
| m4C | 66764 | + | | Pnuc_0065, LSU ribosomal protein L5P | |

| | | | | |
|---|---|---|---|---|
| m4C | 68018 | + | | Pnuc_0068, LSU ribosomal protein L6P |
| m6A | 68025 | + | | Pnuc_0068, LSU ribosomal protein L6P |
| m4C | 70249 | + | | Pnuc_0073, protein translocase subunit secY/sec61 alpha |
| m4C | 71180 | - | | upstream Pnuc_0038, GCN5-related N-acetyltransferase |
| m4C | 71592 | - | | upstream Pnuc_0038, GCN5-related N-acetyltransferase |
| m4C | 76642 | + | | Pnuc_0081, Protein-disulfide reductase |
| m4C | 76864 | - | | upstream Pnuc_0038, GCN5-related N-acetyltransferase |
| m4C | 80624 | - | | Pnuc_0086, diaminopimelate decarboxylase |
| m6A | 82350 | - | | Pnuc_0088, 3-dehydroquinate synthase |
| m6A | 83003 | + | GAYANNNNNNNRTAG | upstream Pnuc_0092, penicillin-binding protein, 1A family |
| m6A | 83013 | - | CTAYNNNNNNNTRTC | Pnuc_0088, 3-dehydroquinate synthase |
| m6A | 83216 | + | | upstream Pnuc_0092, penicillin-binding protein, 1A family |
| m6A | 83832 | - | | Pnuc_0090, type II and III secretion system protein |
| m6A | 84386 | + | CTAYNNNNNNNTRTC | upstream Pnuc_0092, penicillin-binding protein, 1A family |
| m6A | 84396 | - | GAYANNNNNNNRTAG | Pnuc_0090, type II and III secretion system protein |
| m4C | 84866 | + | | upstream Pnuc_0092, penicillin-binding protein, 1A family |
| m6A | 85182 | - | | Pnuc_0091, hypothetical protein |
| m6A | 87597 | + | | Pnuc_0092, penicillin-binding protein, 1A family |
| m4C | 88649 | - | | upstream Pnuc_0091, hypothetical protein |
| m4C | 90112 | + | | Pnuc_0094, glutamate synthase (NADH) large subunit |
| m4C | 96037 | + | | Pnuc_0097, protein of unknown function DUF140 |
| m4C | 96490 | + | | Pnuc_0097, protein of unknown function DUF140 |
| m4C | 96752 | - | | upstream Pnuc_0091, hypothetical protein |
| m4C | 98127 | + | | Pnuc_0100, toluene tolerance family protein |
| m4C | 100046 | + | | Pnuc_0103, ABC-2 type transporter |
| m4C | 100149 | - | | upstream Pnuc_0091, hypothetical protein |
| m6A | 102779 | + | GAYANNNNNNNRTAG | Pnuc_0107, histidinol dehydrogenase |
| m6A | 102789 | - | CTAYNNNNNNNTRTC | upstream Pnuc_0091, hypothetical protein |
| m6A | 103179 | + | CTAYNNNNNNNTRTC | Pnuc_0107, histidinol dehydrogenase |
| m6A | 103189 | - | GAYANNNNNNNRTAG | upstream Pnuc_0091, hypothetical protein |
| m6A | 105053 | - | | Pnuc_0108, histidinol phosphate aminotransferase apoenzyme |
| m4C | 112062 | + | | upstream Pnuc_0120, porin, Gram-negative type |
| m6A | 112153 | - | | Pnuc_0119, pseudo |
| m4C | 112352 | - | | Pnuc_0119, pseudo |
| m4C | 112487 | + | | upstream Pnuc_0120, porin, Gram-negative type |
| m4C | 112495 | - | | upstream Pnuc_0119, pseudo |
| m6A | 113017 | - | | upstream Pnuc_0119, pseudo |
| m4C | 113636 | - | | upstream Pnuc_0119, pseudo |
| m4C | 114280 | + | | Pnuc_0121, TPR repeat-containing protein |
| m4C | 115403 | - | | upstream Pnuc_0119, pseudo |
| m4C | 117662 | - | | Pnuc_0122, Tetratricopeptide TPR_2 repeat protein |
| m4C | 117690 | + | | upstream Pnuc_0125, ubiquinol-cytochrome c reductase, iron-sulfur subunit |
| m4C | 119521 | + | | upstream Pnuc_0125, ubiquinol-cytochrome c reductase, iron-sulfur subunit |
| m6A | 119660 | + | | upstream Pnuc_0125, ubiquinol-cytochrome c reductase, iron-sulfur subunit |

| | | | | |
|---|---|---|---|---|
| m6A | 122705 | + | CTAYNNNNNNNTRTC | Pnuc_0127, cytochrome c1 |
| m6A | 122715 | - | GAYANNNNNNNRTAG | upstream Pnuc_0124, large conductance mechanosensitive channel protein |
| m4C | 128274 | - | | Pnuc_0134, glutaredoxin-like protein |
| m4C | 128600 | + | | upstream Pnuc_0142, GTP-binding protein YchF |
| m6A | 130121 | + | GAYANNNNNNNRTAG | upstream Pnuc_0142, GTP-binding protein YchF |
| m6A | 130131 | - | CTAYNNNNNNNTRTC | Pnuc_0136, bacterial peptide chain release factor 1(bRF-1) |
| m4C | 130852 | + | | upstream Pnuc_0142, GTP-binding protein YchF |
| m6A | 131767 | + | | upstream Pnuc_0142, GTP-binding protein YchF |
| m4C | 134617 | + | | upstream Pnuc_0142, GTP-binding protein YchF |
| m4C | 135742 | + | | upstream Pnuc_0142, GTP-binding protein YchF |
| m4C | 136496 | - | | Pnuc_0141, Ubiquinone biosynthesis hydroxylase, UbiH/UbiF/VisC/COQ6 family |
| m6A | 137333 | + | | Pnuc_0142, GTP-binding protein YchF |
| m4C | 137852 | + | | Pnuc_0142, GTP-binding protein YchF |
| m6A | 139555 | - | | Pnuc_0145, indole-3-glycerol phosphate synthase |
| m4C | 142160 | + | | upstream Pnuc_0150, ApaG domain protein |
| m4C | 144075 | - | | upstream Pnuc_0149, ribulose-5-phosphate 3-epimerase |
| m6A | 149067 | + | CTAYNNNNNNNTRTC | upstream Pnuc_0156, biotin synthase |
| m6A | 149077 | - | GAYANNNNNNNRTAG | Pnuc_0155, aminotransferase |
| m4C | 149940 | + | | Pnuc_0156, biotin synthase |
| m4C | 151549 | - | | upstream Pnuc_0155, aminotransferase |
| m4C | 154698 | + | | Pnuc_0161, peptidoglycan synthetase FtsI |
| m6A | 157261 | - | | upstream Pnuc_0155, aminotransferase |
| m6A | 157993 | + | GAYANNNNNNNRTAG | Pnuc_0163, UDP-N-acetylmuramoylalanyl-D-glutamyl-2,6-diaminopimelate--D-alanyl-D-alanyl ligase |
| m6A | 158003 | - | CTAYNNNNNNNTRTC | upstream Pnuc_0155, aminotransferase |
| m4C | 160218 | - | | upstream Pnuc_0155, aminotransferase |
| m6A | 163647 | + | | Pnuc_0168, UDP-N-acetylmuramate--L-alanine ligase |
| m4C | 165541 | + | | Pnuc_0169, D-alanine--D-alanine ligase |
| m4C | 165703 | - | | upstream Pnuc_0155, aminotransferase |
| m4C | 165797 | - | | upstream Pnuc_0155, aminotransferase |
| m4C | 168341 | - | | upstream Pnuc_0155, aminotransferase |
| m6A | 171550 | + | | Pnuc_0176, protein translocase subunit secA |
| m4C | 173940 | - | | upstream Pnuc_0175, hypothetical protein |
| m6A | 174286 | + | | upstream Pnuc_0177, (2Fe-2S)-binding domain protein |
| m4C | 175181 | - | | upstream Pnuc_0175, hypothetical protein |
| m4C | 175458 | - | | upstream Pnuc_0175, hypothetical protein |
| m6A | 175924 | - | | upstream Pnuc_0175, hypothetical protein |
| m4C | 176761 | - | | upstream Pnuc_0175, hypothetical protein |
| m4C | 177012 | + | | Pnuc_0178, aldehyde oxidase and xanthine dehydrogenase,molybdopterin binding protein |
| m6A | 177242 | - | | upstream Pnuc_0175, hypothetical protein |
| m4C | 179707 | - | | Pnuc_0182, protein of unknown function DUF1342 |
| m4C | 179903 | - | | Pnuc_0182, protein of unknown function DUF1342 |
| m4C | 180511 | - | | Pnuc_0183, dephospho-CoA kinase |
| m4C | 181009 | + | | upstream Pnuc_0188, methylated-DNA--protein-cysteine methyltransferase |
| m4C | 181048 | + | | upstream Pnuc_0188, methylated-DNA--protein-cysteine methyltransferase |

| | | | | |
|---|---|---|---|---|
| m4C | 182559 | - | | Pnuc_0185, type II secretion system protein |
| m6A | 183048 | + | | upstream Pnuc_0188, methylated-DNA--protein-cysteine methyltransferase |
| m4C | 188018 | + | | Pnuc_0190, Na+/solute symporter |
| m6A | 190987 | - | | Pnuc_0193, Farnesyltranstransferase |
| m4C | 191212 | + | | Pnuc_0194, LSU ribosomal protein L21P |
| m4C | 193735 | + | | Pnuc_0197, glutamate 5-kinase |
| m4C | 196593 | - | | upstream Pnuc_0199, NUDIX hydrolase |
| m4C | 197657 | + | | upstream Pnuc_0202, cytochrome c assembly protein |
| m6A | 199926 | + | | upstream Pnuc_0204, ribonucleoside-diphosphate reductase, alpha subunit |
| m6A | 200409 | - | | upstream Pnuc_0201, signal recognition particle subunit FFH/SRP54 (srp54) |
| m6A | 200762 | + | CTAYNNNNNNNNTRTC | Pnuc_0204, ribonucleoside-diphosphate reductase, alpha subunit |
| m6A | 200772 | - | GAYANNNNNNNNRTAG | upstream Pnuc_0201, signal recognition particle subunit FFH/SRP54 (srp54) |
| m4C | 201940 | - | | upstream Pnuc_0201, signal recognition particle subunit FFH/SRP54 (srp54) |
| m4C | 202708 | - | | upstream Pnuc_0201, signal recognition particle subunit FFH/SRP54 (srp54) |
| m4C | 202934 | + | | Pnuc_0204, ribonucleoside-diphosphate reductase, alpha subunit |
| m6A | 203706 | - | | upstream Pnuc_0201, signal recognition particle subunit FFH/SRP54 (srp54) |
| m4C | 204062 | - | | upstream Pnuc_0201, signal recognition particle subunit FFH/SRP54 (srp54) |
| m4C | 205116 | - | | upstream Pnuc_0201, signal recognition particle subunit FFH/SRP54 (srp54) |
| m4C | 205165 | + | | upstream Pnuc_0213, UDP-N-acetylmuramate |
| m6A | 208466 | + | | upstream Pnuc_0213, UDP-N-acetylmuramate |
| m6A | 209668 | + | GAYANNNNNNNNRTAG | upstream Pnuc_0213, UDP-N-acetylmuramate |
| m6A | 209678 | - | CTAYNNNNNNNNTRTC | Pnuc_0211, 3-dehydroquinate dehydratase |
| m6A | 210232 | + | | upstream Pnuc_0213, UDP-N-acetylmuramate |
| m4C | 211553 | + | | Pnuc_0213, UDP-N-acetylmuramate |
| m6A | 216842 | - | | upstream Pnuc_0212, alkyl hydroperoxide reductase/ Thiol specific antioxidant/ Mal allergen |
| m4C | 217176 | - | | upstream Pnuc_0212, alkyl hydroperoxide reductase/ Thiol specific antioxidant/ Mal allergen |
| m4C | 218942 | + | | Pnuc_0220, magnesium and cobalt transport protein CorA |
| m4C | 219189 | + | | Pnuc_0220, magnesium and cobalt transport protein CorA |
| m4C | 221772 | - | | upstream Pnuc_0223, thiamine-phosphate kinase |
| m4C | 224128 | - | | upstream Pnuc_0223, thiamine-phosphate kinase |
| m4C | 224476 | - | | upstream Pnuc_0223, thiamine-phosphate kinase |
| m4C | 226644 | + | | Pnuc_0227, transketolase |
| m4C | 227034 | - | | upstream Pnuc_0226, protein of unknown function DUF558 |
| m6A | 227673 | + | GAYANNNNNNNNRTAG | Pnuc_0228, glyceraldehyde-3-phosphate dehydrogenase |
| m6A | 227683 | - | CTAYNNNNNNNNTRTC | upstream Pnuc_0226, protein of unknown function DUF558 |
| m6A | 228520 | + | | Pnuc_0228, glyceraldehyde-3-phosphate dehydrogenase |
| m6A | 228733 | - | | Pnuc_0229, ferric uptake regulator, Fur family |
| m4C | 230196 | + | | Pnuc_0231, dihydrodipicolinate reductase |
| m6A | 236488 | + | | Pnuc_0235, glutamate-5-semialdehyde dehydrogenase |
| m4C | 236619 | - | | upstream Pnuc_0229, ferric uptake regulator, Fur family |
| m6A | 238728 | + | | Pnuc_0239, glutamate-1-semialdehyde 2,1-aminomutase |
| m4C | 238745 | - | | upstream Pnuc_0229, ferric uptake regulator, Fur family |
| m4C | 239533 | + | | Pnuc_0239, glutamate-1-semialdehyde 2,1-aminomutase |
| m4C | 241062 | + | | upstream Pnuc_0242, protein of unknown function DUF218 |

| | | | | |
|---|---|---|---|---|
| m4C | 241500 | + | | upstream Pnuc_0242, protein of unknown function DUF218 |
| m4C | 243937 | + | | Pnuc_0244, Holliday junction resolvase YqgF |
| m4C | 243991 | + | | Pnuc_0244, Holliday junction resolvase YqgF |
| m4C | 244876 | - | | upstream Pnuc_0241, deoxyribodipyrimidine photo-lyase type I |
| m6A | 246363 | - | | upstream Pnuc_0241, deoxyribodipyrimidine photo-lyase type I |
| m6A | 246364 | - | | upstream Pnuc_0241, deoxyribodipyrimidine photo-lyase type I |
| m4C | 247712 | - | | upstream Pnuc_0241, deoxyribodipyrimidine photo-lyase type I |
| m6A | 248439 | - | GAYANNNNNNNRTAG | upstream Pnuc_0241, deoxyribodipyrimidine photo-lyase type I |
| m4C | 250708 | - | | Pnuc_0251, hypothetical protein |
| m6A | 250899 | + | CTAYNNNNNNNTRTC | Pnuc_0252, hypothetical protein |
| m6A | 250909 | - | GAYANNNNNNNRTAG | upstream Pnuc_0251, hypothetical protein |
| m6A | 251656 | + | GAYANNNNNNNRTAG | upstream Pnuc_0253, dTDP-glucose 4,6-dehydratase |
| m6A | 251666 | - | CTAYNNNNNNNTRTC | upstream Pnuc_0251, hypothetical protein |
| m4C | 255335 | - | | upstream Pnuc_0251, hypothetical protein |
| m4C | 256362 | - | | Pnuc_0258, type I secretion outer membrane protein, TolC family |
| m4C | 256533 | + | | upstream Pnuc_0259, Undecaprenyl-diphosphatase |
| m4C | 257450 | - | | Pnuc_0258, type I secretion outer membrane protein, TolC family |
| m4C | 258503 | + | | Pnuc_0259, Undecaprenyl-diphosphatase |
| m4C | 262450 | + | | upstream Pnuc_0267, hypothetical protein |
| m6A | 264991 | + | | upstream Pnuc_0267, hypothetical protein |
| m4C | 268392 | - | | upstream Pnuc_0266, 3,4-dihydroxy-2-butanone 4-phosphate synthase |
| m4C | 269161 | - | | Pnuc_0272, preQ(0) biosynthesis protein QueC |
| m6A | 269370 | + | GAYANNNNNNNRTAG | upstream Pnuc_0279, serine hydroxymethyltransferase |
| m6A | 269380 | - | CTAYNNNNNNNTRTC | Pnuc_0272 (start codon), preQ(0) biosynthesis protein QueC |
| m4C | 270060 | + | | upstream Pnuc_0279, serine hydroxymethyltransferase |
| m4C | 270570 | + | | upstream Pnuc_0279, serine hydroxymethyltransferase |
| m6A | 271023 | - | CTAYNNNNNNNTRTC | Pnuc_0275, TolB, N-terminal domain protein |
| m4C | 271970 | + | | upstream Pnuc_0279, serine hydroxymethyltransferase |
| m4C | 272119 | - | | Pnuc_0276, Cell division and transport-associated protein TolA |
| m6A | 277009 | + | CTAYNNNNNNNTRTC | upstream Pnuc_0287, Exodeoxyribonuclease VII large subunit |
| m6A | 277019 | - | GAYANNNNNNNRTAG | Pnuc_0282, 3-deoxy-D-manno-octulosonate cytidylyltransferase |
| m6A | 277383 | + | | upstream Pnuc_0287, Exodeoxyribonuclease VII large subunit |
| m4C | 279494 | + | | upstream Pnuc_0287, Exodeoxyribonuclease VII large subunit |
| m4C | 280973 | - | | upstream Pnuc_0286, MotA/TolQ/ExbB proton channel |
| m4C | 281999 | + | | Pnuc_0288, conserved hypothetical protein |
| m4C | 282108 | - | | upstream Pnuc_0286, MotA/TolQ/ExbB proton channel |
| m6A | 283189 | + | GAYANNNNNNNRTAG | upstream Pnuc_0290, protein of unknown function DUF520 |
| m6A | 283199 | - | CTAYNNNNNNNTRTC | Pnuc_0289, UDP-N-acetylmuramate dehydrogenase |
| m4C | 283507 | + | | upstream Pnuc_0290, protein of unknown function DUF520 |
| m4C | 284893 | - | | upstream Pnuc_0289, UDP-N-acetylmuramate dehydrogenase |
| m4C | 285687 | - | | upstream Pnuc_0289, UDP-N-acetylmuramate dehydrogenase |
| m4C | 295143 | - | | upstream Pnuc_0300, hypothetical protein |
| m4C | 300803 | + | | upstream Pnuc_0311, polysaccharide biosynthesis protein |
| m4C | 301640 | + | | upstream Pnuc_0311, polysaccharide biosynthesis protein |

| | | | | |
|---|---|---|---|---|
| m6A | 310366 | + | CTAYNNNNNNNTRTC | Pnuc_0318, ABC-2 type transporter |
| m6A | 310376 | - | GAYANNNNNNNRTAG | upstream Pnuc_0316, glycosyl transferase, family 4 |
| m6A | 312933 | + | GAYANNNNNNNRTAG | Pnuc_0320, Methyltransferase type 11 |
| m6A | 312943 | - | CTAYNNNNNNNTRTC | upstream Pnuc_0316, glycosyl transferase, family 4 |
| m6A | 313175 | + | GAYANNNNNNNRTAG | Pnuc_0320, Methyltransferase type 11 |
| m6A | 313185 | - | CTAYNNNNNNNTRTC | upstream Pnuc_0316, glycosyl transferase, family 4 |
| m4C | 318996 | + | | Pnuc_0323, Phytanoyl-CoA dioxygenase |
| m6A | 319009 | - | GAYANNNNNNNRTAG | upstream Pnuc_0316, glycosyl transferase, family 4 |
| m4C | 320324 | + | | Pnuc_0325, amine oxidase |
| m4C | 320845 | + | | Pnuc_0325, amine oxidase |
| m6A | 320925 | + | GAYANNNNNNNRTAG | Pnuc_0325, amine oxidase |
| m6A | 320935 | - | CTAYNNNNNNNTRTC | upstream Pnuc_0316, glycosyl transferase, family 4 |
| m6A | 321154 | + | CTAYNNNNNNNTRTC | Pnuc_0325, amine oxidase |
| m6A | 321164 | - | GAYANNNNNNNRTAG | upstream Pnuc_0316, glycosyl transferase, family 4 |
| m4C | 322076 | + | | upstream Pnuc_0329, UbiA prenyltransferase |
| m6A | 322610 | + | | upstream Pnuc_0329, UbiA prenyltransferase |
| m6A | 324282 | + | | Pnuc_0329, UbiA prenyltransferase |
| m6A | 324292 | + | GAYANNNNNNNRTAG | Pnuc_0329, UbiA prenyltransferase |
| m6A | 324302 | - | CTAYNNNNNNNTRTC | upstream Pnuc_0328, FAD linked oxidase domain protein |
| m4C | 328401 | - | | upstream Pnuc_0332, hypothetical protein |
| m6A | 329369 | - | | Pnuc_0334, Adenylyl-sulfate kinase |
| m4C | 329501 | - | | upstream Pnuc_0334, Adenylyl-sulfate kinase |
| m4C | 329877 | + | | Pnuc_0335, glycosyl transferase, family 2 |
| m6A | 332882 | - | | upstream Pnuc_0336, hypothetical protein |
| m4C | 334253 | + | | Pnuc_0337, acyltransferase 3 |
| m6A | 335211 | - | | upstream Pnuc_0336, hypothetical protein |
| m6A | 335858 | + | CTAYNNNNNNNTRTC | Pnuc_0338, hypothetical protein |
| m6A | 335868 | - | GAYANNNNNNNRTAG | upstream Pnuc_0336, hypothetical protein |
| m4C | 336301 | + | | upstream Pnuc_0340, cobalamin (vitamin B12) biosynthesis CbiX protein |
| m4C | 338125 | + | | Pnuc_0342, transcriptional regulator, ArsR family |
| m6A | 345459 | + | GAYANNNNNNNRTAG | upstream Pnuc_0351, acyltransferase 3 |
| m6A | 345469 | - | CTAYNNNNNNNTRTC | upstream Pnuc_0349, hypothetical protein |
| m6A | 347772 | + | GAYANNNNNNNRTAG | Pnuc_0353, Choline dehydrogenase |
| m6A | 347782 | - | CTAYNNNNNNNTRTC | upstream Pnuc_0352, protein of unknown function DUF615 |
| m4C | 349093 | + | | Pnuc_0354, protein of unknown function DUF6, transmembrane |
| m4C | 349184 | - | | upstream Pnuc_0352, protein of unknown function DUF615 |
| m4C | 354187 | + | | Pnuc_0358, amino acid/amide ABC transporter ATP-binding protein 2, HAAT family |
| m4C | 354926 | + | | upstream Pnuc_0362, conserved hypothetical protein |
| m6A | 357990 | + | | upstream Pnuc_0364, Lytic transglycosylase, catalytic |
| m6A | 358119 | - | | Pnuc_0363, DNA/RNA non-specific endonuclease |
| m4C | 363047 | + | | upstream Pnuc_0367, BLUF domain protein |
| m4C | 367208 | + | | Pnuc_0371, transcriptional regulator, TetR family |
| m4C | 367793 | + | | Pnuc_0372, isochorismatase hydrolase |
| m6A | 371430 | + | | upstream Pnuc_0379, PpiC-type peptidyl-prolyl cis-trans isomerase |

| | | | | |
|---|---|---|---|---|
| m4C | 373857 | + | | upstream Pnuc_0384, D-isomer specific 2-hydroxyacid dehydrogenase, NAD-binding protein |
| m6A | 374462 | + | | upstream Pnuc_0384, D-isomer specific 2-hydroxyacid dehydrogenase, NAD-binding protein |
| m4C | 375418 | - | | Pnuc_0383, 4Fe-4S ferredoxin, iron-sulfur binding domain protein |
| m4C | 376773 | - | | Pnuc_0385, conserved hypothetical protein |
| m4C | 378167 | + | | upstream Pnuc_0390, ribosomal large subunit pseudouridine synthase C |
| m4C | 379643 | + | | upstream Pnuc_0390, ribosomal large subunit pseudouridine synthase C |
| m4C | 380414 | + | | upstream Pnuc_0390, ribosomal large subunit pseudouridine synthase C |
| m6A | 381442 | - | | Pnuc_0389, RNAse E |
| m6A | 382815 | + | | upstream Pnuc_0390, ribosomal large subunit pseudouridine synthase C |
| m6A | 387862 | - | | upstream Pnuc_0394, maf protein |
| m4C | 387959 | - | | upstream Pnuc_0394, maf protein |
| m4C | 388854 | - | | upstream Pnuc_0394, maf protein |
| m4C | 391943 | + | | Pnuc_0400, 3-oxoacyl-[acyl-carrier-protein] reductase |
| m4C | 392231 | - | | upstream Pnuc_0394, maf protein |
| m4C | 395248 | - | | upstream Pnuc_0394, maf protein |
| m6A | 395250 | - | | upstream Pnuc_0394, maf protein |
| m6A | 397041 | + | CTAYNNNNNNNNTRTC | Pnuc_0404, GTP-binding protein LepA |
| m6A | 397051 | - | GAYANNNNNNNNRTAG | upstream Pnuc_0394, maf protein |
| m4C | 398900 | - | | upstream Pnuc_0394, maf protein |
| m6A | 400511 | - | | upstream Pnuc_0394, maf protein |
| m6A | 401086 | + | CTAYNNNNNNNNTRTC | Pnuc_0409, pyridoxine 5'-phosphate synthase |
| m6A | 401096 | - | GAYANNNNNNNNRTAG | upstream Pnuc_0394, maf protein |
| m6A | 401232 | + | GAYANNNNNNNNRTAG | Pnuc_0410, holo-acyl-carrier-protein synthase |
| m6A | 401242 | - | CTAYNNNNNNNNTRTC | upstream Pnuc_0394, maf protein |
| m6A | 401354 | + | | Pnuc_0410, holo-acyl-carrier-protein synthase |
| m4C | 407582 | + | | Pnuc_0416, CDP-diacylglycerol-glycerol-3-phosphate-3-phosphatidyltransferase |
| m4C | 408512 | - | | upstream Pnuc_0414, conserved hypothetical protein |
| m4C | 410693 | + | | upstream Pnuc_0420, SSU ribosomal protein S6P |
| m6A | 413274 | + | | Pnuc_0424, DnaB domain protein helicase, C-terminal domain protein |
| m4C | 414298 | + | | Pnuc_0424, DnaB domain protein helicase, C-terminal domain protein |
| m4C | 417659 | - | | Pnuc_0428, PhoH family protein |
| m4C | 418723 | + | | upstream Pnuc_0437, aminotransferase |
| m6A | 419827 | - | | upstream Pnuc_0430, polysaccharide deacetylase |
| m6A | 420798 | - | | Pnuc_0431, NAD-dependent epimerase/dehydratase |
| m4C | 421451 | + | | upstream Pnuc_0437, aminotransferase |
| m6A | 421455 | + | GAYANNNNNNNNRTAG | upstream Pnuc_0437, aminotransferase |
| m6A | 421465 | - | CTAYNNNNNNNNTRTC | Pnuc_0432, formyl transferase domain protein |
| m4C | 422526 | - | | Pnuc_0433, glycosyl transferase, family 2 |
| m6A | 422836 | + | CTAYNNNNNNNNTRTC | upstream Pnuc_0437, aminotransferase |
| m6A | 422846 | - | GAYANNNNNNNNRTAG | Pnuc_0434, DegT/DnrJ/EryC1/StrS aminotransferase |
| m4C | 425783 | - | | Pnuc_0436, protein of unknown function DUF498 |
| m4C | 425791 | - | | Pnuc_0436, protein of unknown function DUF499 |
| m4C | 434948 | + | | Pnuc_0444, major facilitator superfamily MFS_1 |
| m4C | 436608 | + | | upstream Pnuc_0447, ATPase AAA-2 domain protein |

| | | | | |
|---|---|---|---|---|
| m4C | 438249 | + | | Pnuc_0447, ATPase AAA-2 domain protein |
| m6A | 438837 | + | | Pnuc_0447, ATPase AAA-2 domain protein |
| m6A | 439040 | + | GAYANNNNNNNRTAG | Pnuc_0447, ATPase AAA-2 domain protein |
| m6A | 439050 | - | CTAYNNNNNNNTRTC | upstream Pnuc_0446, Excinuclease ABC, C subunit domain protein |
| m4C | 439112 | + | | Pnuc_0447, ATPase AAA-2 domain protein |
| m6A | 440012 | + | GAYANNNNNNNRTAG | Pnuc_0447, ATPase AAA-2 domain protein |
| m6A | 440022 | - | CTAYNNNNNNNTRTC | upstream Pnuc_0446, Excinuclease ABC, C subunit domain protein |
| m6A | 442569 | + | GAYANNNNNNNRTAG | Pnuc_0450, major facilitator superfamily MFS_1 |
| m6A | 442579 | - | CTAYNNNNNNNTRTC | upstream Pnuc_0449, Uncharacterized protein UPF0065 |
| m4C | 445233 | - | | upstream Pnuc_0449, Uncharacterized protein UPF0065 |
| m6A | 445942 | + | CTAYNNNNNNNTRTC | upstream Pnuc_0453, cytochrome c oxidase, cbb3-type, subunit I |
| m6A | 445952 | - | GAYANNNNNNNRTAG | upstream Pnuc_0449, Uncharacterized protein UPF0065 |
| m4C | 449523 | - | | upstream Pnuc_0449, Uncharacterized protein UPF0066 |
| m4C | 451434 | + | | upstream Pnuc_0461, putative transmembrane protein |
| m6A | 453512 | + | CTAYNNNNNNNTRTC | Pnuc_0463, UspA domain protein |
| m6A | 453522 | - | GAYANNNNNNNRTAG | upstream Pnuc_0462, conserved hypothetical protein |
| m6A | 454717 | + | | upstream Pnuc_0466, Mandelate racemase/muconate lactonizing enzyme, C-terminal domain protein |
| m4C | 457971 | + | | upstream Pnuc_0471, 3-hydroxyacyl-CoA dehydrogenase |
| m4C | 458223 | + | | upstream Pnuc_0471, 3-hydroxyacyl-CoA dehydrogenase |
| m4C | 461032 | + | | upstream Pnuc_0471, 3-hydroxyacyl-CoA dehydrogenase |
| m6A | 461835 | + | GAYANNNNNNNRTAG | upstream Pnuc_0471, 3-hydroxyacyl-CoA dehydrogenase |
| m6A | 461845 | - | CTAYNNNNNNNTRTC | Pnuc_0470, D-isomer specific 2-hydroxyacid dehydrogenase, NAD-binding protein |
| m4C | 465062 | - | | upstream Pnuc_0472, major facilitator superfamily MFS_1 |
| m4C | 465452 | + | | Pnuc_0473, cytochrome B561 |
| m4C | 472152 | + | | upstream Pnuc_0485, Uncharacterized protein UPF0065 |
| m6A | 472803 | + | | upstream Pnuc_0485, Uncharacterized protein UPF0066 |
| m4C | 474531 | + | | upstream Pnuc_0485, Uncharacterized protein UPF0067 |
| m6A | 474614 | + | CTAYNNNNNNNTRTC | upstream Pnuc_0485, Uncharacterized protein UPF0068 |
| m4C | 474619 | - | | upstream Pnuc_0484, Rhodanese domain protein |
| m6A | 474624 | - | GAYANNNNNNNRTAG | upstream Pnuc_0484, Rhodanese domain protein |
| m4C | 477739 | - | | upstream Pnuc_0484, Rhodanese domain protein |
| m4C | 479258 | - | | upstream Pnuc_0484, Rhodanese domain protein |
| m4C | 481144 | + | | upstream Pnuc_0493, DNA gyrase subunit A |
| m4C | 482087 | - | | upstream Pnuc_0491, 3-demethylubiquinone-9 3-methyltransferase |
| m4C | 489516 | + | | Pnuc_0497, prephenate dehydrogenase |
| m4C | 489659 | + | | Pnuc_0497, prephenate dehydrogenase |
| m4C | 489973 | - | | upstream Pnuc_0492, OmpA/MotB domain protein |
| m6A | 490233 | + | | Pnuc_0498, 3-phosphoshikimate 1-carboxyvinyltransferase |
| m4C | 491263 | + | | Pnuc_0499, cytidylate kinase |
| m4C | 491303 | - | | upstream Pnuc_0492, OmpA/MotB domain protein |
| m6A | 491847 | + | | upstream Pnuc_0500, SSU ribosomal protein S1P |
| m4C | 492724 | - | | upstream Pnuc_0492, OmpA/MotB domain protein |
| m4C | 496985 | + | | Pnuc_0504, D-alpha,beta-D-heptose 7-phosphate 1-kinase |
| m4C | 501868 | + | | upstream Pnuc_0510, histone deacetylase superfamily |

| | | | | |
|---|---|---|---|---|
| m4C | 504222 | + | | Pnuc_0511, AMP-dependent synthetase and ligase |
| m6A | 506888 | - | | upstream Pnuc_0509, lytic murein transglycosylase B |
| m4C | 508533 | - | | upstream Pnuc_0509, lytic murein transglycosylase B |
| m6A | 511656 | + | GAYANNNNNNNRTAG | Pnuc_0520, LSU ribosomal protein L19P |
| m6A | 511666 | - | CTAYNNNNNNNTRTC | upstream Pnuc_0516, protein of unknown function DUF306, Meta and HslJ |
| m6A | 513793 | + | CTAYNNNNNNNTRTC | upstream Pnuc_0525, Exonuclease, RNase T and DNA polymerase III |
| m6A | 513803 | - | GAYANNNNNNNRTAG | Pnuc_0523, ribosome small subunit-dependent GTPase A |
| m4C | 515473 | + | | upstream Pnuc_0525, Exonuclease, RNase T and DNA polymerase III |
| m4C | 515660 | + | | upstream Pnuc_0525, Exonuclease, RNase T and DNA polymerase III |
| m4C | 515722 | + | | upstream Pnuc_0525, Exonuclease, RNase T and DNA polymerase III |
| m4C | 516276 | + | | Pnuc_0525, Exonuclease, RNase T and DNA polymerase III |
| m4C | 516409 | + | | Pnuc_0525, Exonuclease, RNase T and DNA polymerase III |
| m6A | 516835 | + | GAYANNNNNNNRTAG | upstream Pnuc_0527, microcin-processing peptidase 1, Unknown type peptidase, MEROPS family U62 |
| m6A | 516845 | - | CTAYNNNNNNNTRTC | Pnuc_0526, molybdopterin adenylyltransferase |
| m6A | 517316 | - | | upstream Pnuc_0526, molybdopterin adenylyltransferase |
| m4C | 523583 | + | | upstream Pnuc_0534, protein of unknown function DUF485 |
| m4C | 524625 | - | | upstream Pnuc_0531, phospholipase/Carboxylesterase |
| m6A | 525416 | - | | upstream Pnuc_0531, phospholipase/Carboxylesterase |
| m4C | 525674 | + | | Pnuc_0535, SSS sodium solute transporter superfamily |
| m4C | 525690 | - | | upstream Pnuc_0531, phospholipase/Carboxylesterase |
| m6A | 529410 | + | GAYANNNNNNNRTAG | upstream Pnuc_0541, Glutathione S-transferase, N-terminal domain protein |
| m6A | 529420 | - | CTAYNNNNNNNTRTC | upstream Pnuc_0538, hypothetical protein |
| m4C | 529465 | + | | upstream Pnuc_0541, Glutathione S-transferase, N-terminal domain protein |
| m4C | 532022 | + | | upstream Pnuc_0541, Glutathione S-transferase, N-terminal domain protein |
| m4C | 534332 | - | | upstream Pnuc_0540, GCN5-related N-acetyltransferase |
| m4C | 535051 | - | | upstream Pnuc_0540, GCN5-related N-acetyltransferase |
| m6A | 540218 | - | | upstream Pnuc_0547, putative phytochelatin synthase |
| m4C | 541005 | + | | Pnuc_0550, secretory lipase |
| m4C | 543554 | + | | Pnuc_0552, Cl- channel, voltage-gated family protein |
| m4C | 548812 | - | | upstream Pnuc_0555, cytochrome c, class I |
| m6A | 549517 | + | | upstream Pnuc_0559, putative transmembrane protein |
| m6A | 553023 | + | CTAYNNNNNNNTRTC | upstream Pnuc_0565, hypothetical protein |
| m6A | 553033 | - | GAYANNNNNNNRTAG | Pnuc_0563, integral membrane sensor hybrid histidine kinase |
| m6A | 555606 | - | | upstream Pnuc_0564, two component transcriptional regulator, LuxR family |
| m4C | 556422 | + | | upstream Pnuc_0570, cation diffusion facilitator family transporter |
| m4C | 557356 | + | | upstream Pnuc_0570, cation diffusion facilitator family transporter |
| m6A | 560162 | + | GAYANNNNNNNRTAG | Pnuc_0570, cation diffusion facilitator family transporter |
| m6A | 560172 | - | CTAYNNNNNNNTRTC | upstream Pnuc_0569, GCN5-related N-acetyltransferase |
| m6A | 561102 | - | | Pnuc_0571, beta-lactamase domain protein |
| m4C | 572159 | - | | upstream Pnuc_0582, hypothetical protein |
| m4C | 572808 | + | | upstream Pnuc_0586, cold-shock DNA-binding protein family |
| m6A | 580739 | + | | upstream Pnuc_0597, putative iron-sulfur cluster binding protein |
| m6A | 584889 | - | | Pnuc_0600, FAD linked oxidase domain protein |
| m6A | 587239 | - | CTAYNNNNNNNTRTC | Pnuc_0600, FAD linked oxidase domain protein |

| | | | | |
|---|---|---|---|---|
| m4C | 591454 | - | | upstream Pnuc_0601, transcriptional regulator, GntR family |
| m4C | 597394 | - | | Pnuc_0607, glycosyl transferase, family 9 |
| m6A | 598039 | + | | upstream Pnuc_0610, lipid A ABC exporter, fused ATPase and inner membrane subunits MsbA |
| m4C | 598167 | - | | Pnuc_0608, glycosyl transferase, family 2 |
| m4C | 598618 | + | | upstream Pnuc_0610, lipid A ABC exporter, fused ATPase and inner membrane subunits MsbA |
| m6A | 598932 | + | GAYANNNNNNNRTAG | upstream Pnuc_0610, lipid A ABC exporter, fused ATPase and inner membrane subunits MsbA |
| m6A | 598942 | - | CTAYNNNNNNNTRTC | Pnuc_0609, glycosyl transferase, family 2 |
| m6A | 599429 | + | | Pnuc_0610, lipid A ABC exporter, fused ATPase and inner membrane subunits MsbA |
| m4C | 599511 | + | | Pnuc_0610, lipid A ABC exporter, fused ATPase and inner membrane subunits MsbA |
| m4C | 602158 | - | | Pnuc_0611, RNAse G |
| m6A | 604916 | + | | upstream Pnuc_R0016, tRNA-Leu |
| m6A | 605773 | - | | Pnuc_0617, phosphoribosylamine-glycine ligase |
| m6A | 606385 | - | | Pnuc_0617, phosphoribosylamine-glycine ligase |
| m4C | 612590 | - | | upstream Pnuc_0622, permease YjgP/YjgQ family protein |
| m6A | 616740 | - | | upstream Pnuc_0622, permease YjgP/YjgQ family protein |
| m4C | 616828 | + | | Pnuc_0627, Malonyl-CoA decarboxylase |
| m6A | 618648 | + | GAYANNNNNNNRTAG | Pnuc_0628, TRAP dicarboxylate transporter- DctP subunit |
| m6A | 618658 | - | CTAYNNNNNNNTRTC | upstream Pnuc_0622, permease YjgP/YjgQ family protein |
| m4C | 618674 | + | | Pnuc_0628, TRAP dicarboxylate transporter- DctP subunit |
| m4C | 620884 | + | | Pnuc_0631, AMP-dependent synthetase and ligase |
| m6A | 621941 | + | GAYANNNNNNNRTAG | Pnuc_0631, AMP-dependent synthetase and ligase |
| m6A | 621951 | - | CTAYNNNNNNNTRTC | upstream Pnuc_0622, permease YjgP/YjgQ family protein |
| m6A | 624339 | - | | Pnuc_0635, TonB-dependent receptor, plug |
| m6A | 629963 | + | | Pnuc_0638, L-carnitine dehydratase/bile acid-inducible protein F |
| m4C | 631605 | + | | Pnuc_0640, L-carnitine dehydratase/bile acid-inducible protein F |
| m6A | 632692 | + | | Pnuc_0641, Methionyl-tRNA formyltransferase |
| m4C | 634706 | - | | upstream Pnuc_0642, hypothetical protein |
| m4C | 634852 | - | | upstream Pnuc_0642, hypothetical protein |
| m4C | 636470 | + | | upstream Pnuc_0648, formate dehydrogenase, subunit FdhD |
| m6A | 637029 | + | | upstream Pnuc_0648, formate dehydrogenase, subunit FdhD |
| m4C | 637138 | + | | upstream Pnuc_0648, formate dehydrogenase, subunit FdhD |
| m4C | 637356 | - | | Pnuc_0647, methionyl-tRNA synthetase |
| m4C | 637984 | - | | Pnuc_0647, methionyl-tRNA synthetase |
| m6A | 638441 | + | | upstream Pnuc_0648, formate dehydrogenase, subunit FdhD |
| m6A | 639054 | + | GAYANNNNNNNRTAG | Pnuc_0648, formate dehydrogenase, subunit FdhD |
| m6A | 639064 | - | CTAYNNNNNNNTRTC | upstream Pnuc_0647, methionyl-tRNA synthetase |
| m4C | 639945 | + | | Pnuc_0649, dCTP deaminase |
| m4C | 640774 | - | | upstream Pnuc_0647, methionyl-tRNA synthetase |
| m6A | 642290 | + | CTAYNNNNNNNTRTC | Pnuc_0651, Lysine decarboxylase |
| m6A | 642300 | - | GAYANNNNNNNRTAG | upstream Pnuc_0647, methionyl-tRNA synthetase |
| m4C | 644088 | - | | Pnuc_0653, TRAP dicarboxylate transporter, DctM subunit |
| m4C | 645924 | - | | Pnuc_0654, Tripartite ATP-independent periplasmic transporter, DctQ component |
| m4C | 648447 | + | | Pnuc_0656, porphobilinogen deaminase |
| m4C | 650385 | + | | upstream Pnuc_0660, Lysine exporter protein (LYSE/YGGA) |

| | | | | |
|---|---|---|---|---|
| m4C | 650696 | + | | upstream Pnuc_0660, Lysine exporter protein (LYSE/YGGA) |
| m4C | 653141 | + | | upstream Pnuc_0662, peptide methionine sulfoxide reductase |
| m4C | 653204 | + | | upstream Pnuc_0662, peptide methionine sulfoxide reductase |
| m4C | 654796 | - | | Pnuc_0663, Pyridoxamine 5'-phosphate oxidase |
| m6A | 656037 | + | | upstream Pnuc_0665, amino acid/polyamine/organocation transporter, APC superfamily |
| m6A | 656304 | - | | upstream Pnuc_0663, Pyridoxamine 5'-phosphate oxidase |
| m4C | 656646 | - | | upstream Pnuc_0663, Pyridoxamine 5'-phosphate oxidase |
| m4C | 661291 | + | | Pnuc_0668, aminopeptidase N |
| m4C | 663672 | + | | upstream Pnuc_0672, ABC transporter related protein |
| m4C | 664099 | + | | upstream Pnuc_0672, ABC transporter related protein |
| m6A | 664401 | + | CTAYNNNNNNNNTRTC | upstream Pnuc_0672, ABC transporter related protein |
| m4C | 664610 | - | | Pnuc_0670, cyanophycin synthetase |
| m4C | 668365 | - | | upstream Pnuc_0671, cyanophycin synthetase |
| m4C | 669887 | + | | Pnuc_0672, ABC transporter related protein |
| m6A | 670881 | - | | upstream Pnuc_0671, cyanophycin synthetase |
| m4C | 673061 | - | | upstream Pnuc_0671, cyanophycin synthetase |
| m4C | 673851 | - | | upstream Pnuc_0671, cyanophycin synthetase |
| m4C | 676557 | - | | upstream Pnuc_0671, cyanophycin synthetase |
| m6A | 679130 | + | | upstream Pnuc_0681, response regulator receiver protein |
| m4C | 683590 | + | | upstream Pnuc_0686, V-type H(+)-translocating pyrophosphatase |
| m4C | 683682 | - | | upstream Pnuc_0684, NAD+ synthetase |
| m6A | 685642 | + | CTAYNNNNNNNNTRTC | Pnuc_0686, V-type H(+)-translocating pyrophosphatase |
| m6A | 685652 | - | GAYANNNNNNNNRTAG | upstream Pnuc_0685, Inorganic diphosphatase |
| m4C | 688295 | + | | upstream Pnuc_0690, DNA translocase FtsK |
| m4C | 689350 | + | | Pnuc_0690, DNA translocase FtsK |
| m4C | 690307 | - | | upstream Pnuc_0689, thioredoxin reductase |
| m4C | 692254 | - | | upstream Pnuc_0689, thioredoxin reductase |
| m6A | 694070 | - | | upstream Pnuc_0694, transcriptional regulator/antitoxin, MazE |
| m4C | 695098 | + | | Pnuc_0695, ATP-dependent DNA helicase RecQ |
| m4C | 702329 | - | | upstream Pnuc_R0019, SRP RNA; RNA component of signal recognition particle |
| m4C | 702948 | - | | upstream Pnuc_R0019, SRP RNA; RNA component of signal recognition particle |
| m4C | 704404 | + | | Pnuc_0703, Carbohydrate-selective porin OprB |
| m4C | 704548 | + | | upstream Pnuc_0704, SOS response UmuD protein, Serine peptidase,MEROPS family S24 |
| m4C | 706374 | - | | upstream Pnuc_R0019, SRP RNA; RNA component of signal recognition particle |
| m6A | 706486 | - | | upstream Pnuc_R0019, SRP RNA; RNA component of signal recognition particle |
| m6A | 708630 | + | | upstream Pnuc_0709, protein of unknown function DUF465 |
| m6A | 711386 | - | | Pnuc_0712, membrane protein of unknown function |
| m4C | 712379 | - | | upstream Pnuc_0713, SlyX family protein |
| m4C | 712948 | + | | Pnuc_0714, luciferase family protein |
| m4C | 713493 | - | | Pnuc_0716, YaeQ family protein |
| m4C | 714147 | - | | upstream Pnuc_0716, YaeQ family protein |
| m6A | 715576 | - | | upstream Pnuc_0716, YaeQ family protein |
| m6A | 716042 | + | | Pnuc_0718, coproporphyrinogen III oxidase, anaerobic |
| m6A | 716815 | + | CTAYNNNNNNNNTRTC | Pnuc_0718, coproporphyrinogen III oxidase, anaerobic |

| | | | | |
|---|---|---|---|---|
| m6A | 716825 | - | GAYANNNNNNNRTAG | upstream Pnuc_0716, YaeQ family protein |
| m6A | 717198 | + | CTAYNNNNNNNTRTC | upstream Pnuc_0719, OmpW family protein |
| m6A | 717208 | - | GAYANNNNNNNRTAG | upstream Pnuc_0716, YaeQ family protein |
| m4C | 719965 | + | | Pnuc_0721, ABC transporter related protein |
| m4C | 720667 | - | | Pnuc_0722, conserved hypothetical protein |
| m4C | 724646 | + | | Pnuc_0726, TRAP transporter solute receptor, TAXI family |
| m6A | 725043 | - | | upstream Pnuc_0725, Pirin domain protein |
| m6A | 726854 | - | | Pnuc_0728, sulfide dehydrogenase (flavocytochrome), flavoprotein subunit |
| m4C | 727058 | + | | upstream Pnuc_0734, 2-oxo-acid dehydrogenase E1 subunit, homodimeric type |
| m4C | 727350 | + | | upstream Pnuc_0734, 2-oxo-acid dehydrogenase E1 subunit, homodimeric type |
| m4C | 727751 | + | | upstream Pnuc_0734, 2-oxo-acid dehydrogenase E1 subunit, homodimeric type |
| m4C | 728120 | + | | upstream Pnuc_0734, 2-oxo-acid dehydrogenase E1 subunit, homodimeric type |
| m4C | 729810 | + | | upstream Pnuc_0734, 2-oxo-acid dehydrogenase E1 subunit, homodimeric type |
| m6A | 730423 | + | CTAYNNNNNNNTRTC | upstream Pnuc_0734, 2-oxo-acid dehydrogenase E1 subunit, homodimeric type |
| m6A | 730433 | - | GAYANNNNNNNRTAG | Pnuc_0731, methenyltetrahydrofolate cyclohydrolase / 5,10-methylenetetrahydrofolate dehydrogenase (NADP+) |
| m4C | 734318 | - | | upstream Pnuc_0733, PAS/PAC sensor signal transduction histidine kinase |
| m4C | 738410 | + | | Pnuc_0734, 2-oxo-acid dehydrogenase E1 subunit, homodimeric type |
| m6A | 739406 | + | CTAYNNNNNNNTRTC | Pnuc_0736, dihydrolipoamide dehydrogenase |
| m6A | 739416 | - | GAYANNNNNNNRTAG | upstream Pnuc_0733, PAS/PAC sensor signal transduction histidine kinase |
| m4C | 739572 | - | | upstream Pnuc_0733, PAS/PAC sensor signal transduction histidine kinase |
| m6A | 741643 | + | CTAYNNNNNNNTRTC | Pnuc_0738, protein of unknown function DUF6, transmembrane |
| m6A | 741653 | - | GAYANNNNNNNRTAG | upstream Pnuc_0737, phasin family protein |
| m4C | 742273 | + | | Pnuc_0738, protein of unknown function DUF6, transmembrane |
| m4C | 742983 | + | | Pnuc_0739, murein-DD-endopeptidase, Serine peptidase,MEROPS family S11 |
| m4C | 743138 | - | | upstream Pnuc_0737, phasin family protein |
| m4C | 746050 | - | | Pnuc_0742, Arginyltransferase |
| m4C | 746222 | - | | Pnuc_0743, Leucyltransferase |
| m6A | 746554 | + | | upstream Pnuc_0753, protein of unknown function DUF328 |
| m4C | 746934 | - | | Pnuc_0744, NUDIX hydrolase |
| m4C | 747724 | - | | Pnuc_0745, Anthranilate synthase component I and chorismate binding protein |
| m4C | 749359 | + | | upstream Pnuc_0753, protein of unknown function DUF328 |
| m4C | 749722 | - | | Pnuc_0746, peptidyl-prolyl cis-trans isomerase, cyclophilin type |
| m6A | 750406 | + | GAYANNNNNNNRTAG | upstream Pnuc_0753, protein of unknown function DUF328 |
| m6A | 750416 | - | CTAYNNNNNNNTRTC | Pnuc_0747, DSBA oxidoreductase |
| m6A | 751830 | + | | upstream Pnuc_0753, protein of unknown function DUF328 |
| m6A | 762452 | + | CTAYNNNNNNNTRTC | Pnuc_0761, succinate dehydrogenase subunit B |
| m6A | 762462 | - | GAYANNNNNNNRTAG | upstream Pnuc_0756, malate dehydrogenase (NAD) |
| m6A | 763654 | + | | Pnuc_0763, citrate synthase |
| m4C | 763691 | - | | upstream Pnuc_0756, malate dehydrogenase (NAD) |
| m4C | 763891 | - | | upstream Pnuc_0756, malate dehydrogenase (NAD) |
| m4C | 763981 | - | | upstream Pnuc_0756, malate dehydrogenase (NAD) |
| m4C | 764280 | + | | Pnuc_0763, citrate synthase |
| m4C | 764981 | + | | Pnuc_0764, 3-isopropylmalate dehydratase, large subunit |
| m6A | 767120 | + | | upstream Pnuc_0767, 3-isopropylmalate dehydrogenase |

| | | | | |
|---|---|---|---|---|
| m4C | 767735 | + | | Pnuc_0767, 3-isopropylmalate dehydrogenase |
| m6A | 768513 | + | CTAYNNNNNNNTRTC | Pnuc_0768, aspartate semialdehyde dehydrogenase |
| m6A | 768523 | - | GAYANNNNNNNRTAG | upstream Pnuc_0756, malate dehydrogenase (NAD) |
| m4C | 768661 | + | | Pnuc_0768, aspartate semialdehyde dehydrogenase |
| m4C | 768983 | + | | Pnuc_0768, aspartate semialdehyde dehydrogenase |
| m6A | 771107 | + | | Pnuc_0770, tRNA pseudouridine synthase A |
| m4C | 771705 | - | | upstream Pnuc_0756, malate dehydrogenase (NAD) |
| m4C | 774306 | - | | upstream Pnuc_0756, malate dehydrogenase (NAD) |
| m4C | 774401 | - | | upstream Pnuc_0756, malate dehydrogenase (NAD) |
| m6A | 774540 | + | CTAYNNNNNNNTRTC | Pnuc_0774, acetyl-CoA carboxylase carboxyltransferase subunit alpha |
| m6A | 774550 | - | GAYANNNNNNNRTAG | upstream Pnuc_0756, malate dehydrogenase (NAD) |
| m4C | 775167 | - | | upstream Pnuc_0756, malate dehydrogenase (NAD) |
| m4C | 775467 | - | | upstream Pnuc_0756, malate dehydrogenase (NAD) |
| m4C | 775663 | - | | upstream Pnuc_0756, malate dehydrogenase (NAD) |
| m4C | 775894 | + | | Pnuc_0775, FolC bifunctional protein |
| m4C | 776520 | - | | upstream Pnuc_0756, malate dehydrogenase (NAD) |
| m4C | 779460 | - | | upstream Pnuc_0756, malate dehydrogenase (NAD) |
| m6A | 783630 | - | | upstream Pnuc_0782, molybdate ABC transporter, inner membrane subunit |
| m6A | 783973 | + | GAYANNNNNNNRTAG | Pnuc_0784, OmpA/MotB domain protein |
| m6A | 783983 | - | CTAYNNNNNNNTRTC | upstream Pnuc_0782, molybdate ABC transporter, inner membrane subunit |
| m6A | 784667 | - | | upstream Pnuc_0782, molybdate ABC transporter, inner membrane subunit |
| m4C | 786811 | - | | Pnuc_0787, cytochrome c family protein |
| m6A | 786874 | + | GAYANNNNNNNRTAG | upstream Pnuc_0788, protein of unknown function DUF395, YeeE/YedE |
| m6A | 786884 | - | CTAYNNNNNNNTRTC | Pnuc_0787, cytochrome c family protein |
| m6A | 789389 | + | GAYANNNNNNNRTAG | Pnuc_0792, Rhodanese domain protein |
| m6A | 789399 | - | CTAYNNNNNNNTRTC | upstream Pnuc_0790, hypothetical protein |
| m4C | 792064 | + | | Pnuc_0796, cyclic nucleotide-binding protein |
| m6A | 793322 | + | | upstream Pnuc_0800, beta-lactamase domain protein |
| m6A | 793988 | + | CTAYNNNNNNNTRTC | upstream Pnuc_0800, beta-lactamase domain protein |
| m6A | 793998 | - | GAYANNNNNNNRTAG | Pnuc_0799, FAD-dependent pyridine nucleotide-disulfide oxidoreductase |
| m6A | 794087 | + | | upstream Pnuc_0800, beta-lactamase domain protein |
| m4C | 801055 | + | | Pnuc_0807, sulfate thiol esterase SoxB |
| m4C | 801332 | - | | upstream Pnuc_0799, FAD-dependent pyridine nucleotide-disulfide oxidoreductase |
| m6A | 802163 | + | GAYANNNNNNNRTAG | Pnuc_0807, sulfate thiol esterase SoxB |
| m6A | 802173 | - | CTAYNNNNNNNTRTC | upstream Pnuc_0799, FAD-dependent pyridine nucleotide-disulfide oxidoreductase |
| m4C | 804214 | - | | upstream Pnuc_0799, FAD-dependent pyridine nucleotide-disulfide oxidoreductase |
| m6A | 811058 | - | | Pnuc_0818, acyl-CoA dehydrogenase domain protein |
| m4C | 812378 | - | | upstream Pnuc_0818, acyl-CoA dehydrogenase domain protein |
| m4C | 812493 | - | | upstream Pnuc_0818, acyl-CoA dehydrogenase domain protein |
| m6A | 815395 | + | GAYANNNNNNNRTAG | upstream Pnuc_0825, 3-oxoacid CoA-transferase, A subunit |
| m6A | 815405 | - | CTAYNNNNNNNTRTC | Pnuc_0822, protein of unknown function DUF482 |
| m4C | 818014 | + | | Pnuc_0825, 3-oxoacid CoA-transferase, A subunit |
| m6A | 820903 | + | CTAYNNNNNNNTRTC | Pnuc_0827, cation diffusion facilitator family transporter |
| m6A | 820913 | - | GAYANNNNNNNRTAG | upstream Pnuc_0824, Electron-transferring-flavoprotein dehydrogenase |

| | | | | |
|---|---|---|---|---|
| m4C | 822831 | - | | upstream Pnuc_0824, Electron-transferring-flavoprotein dehydrogenase |
| m4C | 825735 | - | | upstream Pnuc_0824, Electron-transferring-flavoprotein dehydrogenase |
| m6A | 829107 | + | CTAYNNNNNNNTRTC | Pnuc_0834, phenylalanyl-tRNA synthetase beta subunit |
| m6A | 829117 | - | GAYANNNNNNNRTAG | upstream Pnuc_0824, Electron-transferring-flavoprotein dehydrogenase |
| m6A | 829240 | - | | upstream Pnuc_0824, Electron-transferring-flavoprotein dehydrogenase |
| m4C | 831183 | + | | Pnuc_0837, thioesterase superfamily protein |
| m4C | 832785 | + | | upstream Pnuc_0840, 2-oxoglutarate dehydrogenase, E1 subunit |
| m6A | 837877 | + | GAYANNNNNNNRTAG | Pnuc_0842, dihydrolipoamide dehydrogenase |
| m6A | 837887 | - | CTAYNNNNNNNTRTC | upstream Pnuc_0839, DNA binding domain, excisionase family |
| m4C | 839527 | + | | Pnuc_0843, AFG1-family ATPase |
| m4C | 839567 | + | | Pnuc_0843, AFG1-family ATPase |
| m4C | 840271 | - | | upstream Pnuc_0839, DNA binding domain, excisionase family |
| m4C | 840802 | - | | upstream Pnuc_0839, DNA binding domain, excisionase family |
| m4C | 841897 | - | | upstream Pnuc_0839, DNA binding domain, excisionase family |
| m6A | 842339 | - | | upstream Pnuc_0839, DNA binding domain, excisionase family |
| m4C | 843787 | + | | Pnuc_0848, NlpBDapX family lipoprotein |
| m4C | 846473 | - | | upstream Pnuc_0850, Cupin 4 family protein |
| m6A | 846578 | + | | Pnuc_0851, putative FkbP-type peptidyl-prolyl cis-trans isomerase |
| m4C | 850566 | - | | Pnuc_0857, Tetratricopeptide TPR_2 repeat protein |
| m6A | 852895 | - | | upstream Pnuc_0857, Tetratricopeptide TPR_2 repeat protein |
| m4C | 855639 | + | | Pnuc_0861, tRNA(Ile)-lysidine synthetase |
| m6A | 856043 | + | | Pnuc_0862, aspartate kinase |
| m6A | 856210 | + | GAYANNNNNNNRTAG | Pnuc_0862, aspartate kinase |
| m6A | 856220 | - | CTAYNNNNNNNTRTC | upstream Pnuc_0857, Tetratricopeptide TPR_2 repeat protein |
| m4C | 856296 | + | | Pnuc_0862, aspartate kinase |
| m4C | 856710 | + | | Pnuc_0862, aspartate kinase |
| m4C | 859723 | - | | Pnuc_0865, transcriptional regulator, XRE family |
| m6A | 860021 | - | | upstream Pnuc_0865, transcriptional regulator, XRE family |
| m4C | 860044 | - | | upstream Pnuc_0865, transcriptional regulator, XRE family |
| m4C | 862259 | - | | upstream Pnuc_0865, transcriptional regulator, XRE family |
| m4C | 862685 | - | | upstream Pnuc_0865, transcriptional regulator, XRE family |
| m4C | 863254 | - | | upstream Pnuc_0865, transcriptional regulator, XRE family |
| m4C | 864375 | + | | Pnuc_0870, transporter, hydrophobe/amphiphile efflux-1(HAE1) family |
| m6A | 864766 | + | CTAYNNNNNNNTRTC | Pnuc_0870, transporter, hydrophobe/amphiphile efflux-1(HAE1) family |
| m6A | 864776 | - | GAYANNNNNNNRTAG | upstream Pnuc_0865, transcriptional regulator, XRE family |
| m6A | 865018 | + | CTAYNNNNNNNTRTC | Pnuc_0870, transporter, hydrophobe/amphiphile efflux-1(HAE1) family |
| m6A | 865028 | - | GAYANNNNNNNRTAG | upstream Pnuc_0865, transcriptional regulator, XRE family |
| m4C | 865461 | - | | upstream Pnuc_0865, transcriptional regulator, XRE family |
| m6A | 868410 | + | GAYANNNNNNNRTAG | Pnuc_0872, Formyl-CoA transferase |
| m6A | 868420 | - | CTAYNNNNNNNTRTC | upstream Pnuc_0865, transcriptional regulator, XRE family |
| m4C | 868581 | - | | upstream Pnuc_0865, transcriptional regulator, XRE family |
| m4C | 870431 | + | | Pnuc_0874, hypothetical protein |
| m4C | 870709 | - | | upstream Pnuc_0865, transcriptional regulator, XRE family |
| m4C | 871483 | - | | upstream Pnuc_0875, hypothetical protein |

| | | | | |
|---|---|---|---|---|
| m6A | 872853 | - | | upstream Pnuc_0875, hypothetical protein |
| m6A | 873168 | + | CTAYNNNNNNNTRTC | Pnuc_0879, putative transmembrane protein |
| m6A | 873178 | - | GAYANNNNNNNRTAG | upstream Pnuc_0875, hypothetical protein |
| m4C | 877940 | + | | upstream Pnuc_0888, hypothetical protein |
| m4C | 879366 | - | | Pnuc_0886, conserved hypothetical protein |
| m4C | 880969 | + | | upstream Pnuc_0888, hypothetical protein |
| m6A | 882214 | + | GAYANNNNNNNRTAG | Pnuc_0889, phosphomethylpyrimidine kinase |
| m6A | 882224 | - | CTAYNNNNNNNTRTC | upstream Pnuc_R0023, tRNA-Asn |
| m4C | 882503 | + | | Pnuc_0889, phosphomethylpyrimidine kinase |
| m4C | 887087 | + | | upstream Pnuc_0900, DNA topoisomerase IV subunit B |
| m4C | 887655 | + | | upstream Pnuc_0900, DNA topoisomerase IV subunit B |
| m4C | 889137 | - | | Pnuc_0897, NUDIX hydrolase |
| m6A | 889846 | - | | upstream Pnuc_0897, NUDIX hydrolase |
| m6A | 891433 | + | | upstream Pnuc_0900, DNA topoisomerase IV subunit B |
| m4C | 894196 | + | | Pnuc_0900, DNA topoisomerase IV subunit B |
| m6A | 894883 | + | GAYANNNNNNNRTAG | Pnuc_0901, DNA topoisomerase IV subunit A |
| m6A | 894893 | - | CTAYNNNNNNNTRTC | upstream Pnuc_0899, Formyl-CoA transferase |
| m6A | 895256 | - | | upstream Pnuc_0899, Formyl-CoA transferase |
| m4C | 896466 | + | | Pnuc_0901, DNA topoisomerase IV subunit A |
| m6A | 896992 | - | | upstream Pnuc_0899, Formyl-CoA transferase |
| m6A | 898234 | + | CTAYNNNNNNNTRTC | upstream Pnuc_0903, conserved hypothetical protein |
| m6A | 898244 | - | GAYANNNNNNNRTAG | upstream Pnuc_0902, predicted sulfurylase subunit, molybdopterin cytosine dinucleotide biosynthesis |
| m6A | 901432 | + | | upstream Pnuc_0907, putative periplasmic cytochrome type-c oxidoreductase signal peptide protein |
| m6A | 903152 | + | GAYANNNNNNNRTAG | upstream Pnuc_0910, methylmalonyl-CoA mutase |
| m6A | 903162 | - | CTAYNNNNNNNTRTC | upstream Pnuc_0909, transcriptional regulator, GntR family |
| m6A | 905160 | + | | Pnuc_0910, methylmalonyl-CoA mutase |
| m6A | 909439 | + | CTAYNNNNNNNTRTC | Pnuc_0913, acetyl-CoA carboxylase, biotin carboxylase |
| m6A | 909449 | - | GAYANNNNNNNRTAG | upstream Pnuc_0909, transcriptional regulator, GntR family |
| m4C | 911262 | + | | Pnuc_0915, peptidase M22, glycoprotease |
| m4C | 912107 | + | | Pnuc_0917, hypothetical protein |
| m4C | 913241 | + | | upstream Pnuc_0919, alanine racemase |
| m4C | 917021 | - | | upstream Pnuc_0920, putative transmembrane protein |
| m6A | 918248 | + | | Pnuc_0923, poly(R)-hydroxyalkanoic acid synthase, class I |
| m6A | 921978 | + | | Pnuc_0926, SSU ribosomal protein S12P methylthiotransferase |
| m4C | 925623 | - | | Pnuc_0929, transcription-repair coupling factor |
| m4C | 926986 | - | | Pnuc_0929, transcription-repair coupling factor |
| m4C | 929165 | - | | upstream Pnuc_0929, transcription-repair coupling factor |
| m4C | 929591 | + | | Pnuc_0932, nitroreductase |
| m4C | 932124 | - | | upstream Pnuc_0929, transcription-repair coupling factor |
| m4C | 933953 | + | | Pnuc_0936, ATP-dependent proteinase, Serine peptidase,MEROPS family S16 |
| m6A | 936437 | - | | upstream Pnuc_0929, transcription-repair coupling factor |
| m6A | 936991 | + | CTAYNNNNNNNTRTC | Pnuc_0937, hypothetical protein |
| m6A | 937001 | - | GAYANNNNNNNRTAG | upstream Pnuc_0929, transcription-repair coupling factor |
| m4C | 937618 | - | | upstream Pnuc_0929, transcription-repair coupling factor |

| | | | | |
|---|---|---|---|---|
| m4C | 940174 | + | | Pnuc_0940, phosphoribosylformylglycinamidine synthase |
| m6A | 945908 | + | | upstream Pnuc_0945, CTP synthase |
| m6A | 946091 | + | GAYANNNNNNNRTAG | upstream Pnuc_0945, CTP synthase |
| m6A | 946101 | - | CTAYNNNNNNNTRTC | Pnuc_0942, aconitase |
| m4C | 946523 | - | | Pnuc_0942, aconitase |
| m4C | 946770 | + | | upstream Pnuc_0945, CTP synthase |
| m4C | 948299 | + | | Pnuc_0945, CTP synthase |
| m4C | 949510 | + | | Pnuc_0946, 2-dehydro-3-deoxyphosphooctonate aldolase |
| m4C | 951235 | - | | upstream Pnuc_0944, conserved hypothetical protein |
| m4C | 954331 | + | | Pnuc_0950, glutamyl-tRNA synthetase |
| m4C | 959380 | + | | upstream Pnuc_0958, 2-hydroxy-3-oxopropionate reductase |
| m6A | 959740 | + | GAYANNNNNNNRTAG | Pnuc_0958, 2-hydroxy-3-oxopropionate reductase |
| m6A | 959750 | - | CTAYNNNNNNNTRTC | upstream Pnuc_0957, hypothetical protein |
| m4C | 960743 | + | | Pnuc_0959, protein of unknown function UPF0227 |
| m6A | 962004 | + | | Pnuc_0961, hypothetical protein |
| m4C | 963866 | + | | upstream Pnuc_0968, glycine cleavage T protein (aminomethyl transferase) |
| m4C | 964205 | - | | Pnuc_0964, hydrolase, TatD family |
| m4C | 965166 | - | | Pnuc_0965, DNA polymerase III, delta prime subunit |
| m6A | 966942 | + | CTAYNNNNNNNTRTC | upstream Pnuc_0968, glycine cleavage T protein (aminomethyl transferase) |
| m6A | 966952 | - | GAYANNNNNNNRTAG | Pnuc_0967, aminodeoxychorismate lyase |
| m6A | 967979 | + | CTAYNNNNNNNTRTC | Pnuc_0968, glycine cleavage T protein (aminomethyl transferase) |
| m6A | 967989 | - | GAYANNNNNNNRTAG | upstream Pnuc_0967, aminodeoxychorismate lyase |
| m6A | 968424 | + | | Pnuc_0968, glycine cleavage T protein (aminomethyl transferase) |
| m4C | 969819 | - | | Pnuc_0970, thioesterase superfamily protein |
| m6A | 971675 | - | | upstream Pnuc_0970, thioesterase superfamily protein |
| m4C | 975518 | + | | upstream Pnuc_0980, flavodoxin/nitric oxide synthase |
| m4C | 976238 | - | | Pnuc_0978, CBS domain containing protein |
| m6A | 976646 | - | | Pnuc_0979, tRNA-processing RNAse BN |
| m6A | 979842 | - | | upstream Pnuc_0979, tRNA-processing RNAse BN |
| m6A | 980618 | + | | upstream Pnuc_0999, ABC-type transporter, periplasmic component, NitT family |
| m6A | 980984 | + | GAYANNNNNNNRTAG | upstream Pnuc_0999, ABC-type transporter, periplasmic component, NitT family |
| m6A | 980994 | - | CTAYNNNNNNNTRTC | Pnuc_0983, alpha/beta hydrolase fold protein |
| m6A | 982286 | + | CTAYNNNNNNNTRTC | upstream Pnuc_0999, ABC-type transporter, periplasmic component, NitT family |
| m6A | 982296 | - | GAYANNNNNNNRTAG | Pnuc_0986, hypothetical protein |
| m6A | 984361 | + | | upstream Pnuc_0999, ABC-type transporter, periplasmic component, NitT family |
| m6A | 984510 | + | | upstream Pnuc_0999, ABC-type transporter, periplasmic component, NitT family |
| m4C | 986213 | - | | Pnuc_0991, NADH dehydrogenase |
| m6A | 988136 | + | GAYANNNNNNNRTAG | upstream Pnuc_0999, ABC-type transporter, periplasmic component, NitT family |
| m6A | 988146 | - | CTAYNNNNNNNTRTC | Pnuc_0992, Choloylglycine hydrolase |
| m6A | 990429 | - | | Pnuc_0995, nitrate ABC transporter, ATPase subunits C and D |
| m4C | 993131 | + | | upstream Pnuc_0999, ABC-type transporter, periplasmic component, NitT family |
| m4C | 994692 | - | | upstream Pnuc_0998, uroporphyrin-III C-methyltransferase |
| m6A | 995404 | + | CTAYNNNNNNNTRTC | Pnuc_0999, ABC-type transporter, periplasmic component, NitT family |
| m4C | 995853 | - | | upstream Pnuc_0998, uroporphyrin-III C-methyltransferase |

| | | | | |
|---|---|---|---|---|
| m6A | 997866 | + | | Pnuc_1000, assimilatory nitrite reductase (NAD(P)H) large subunit precursor |
| m6A | 1006842 | - | | upstream Pnuc_R0031, tRNA-Pro |
| m6A | 1008651 | + | CTAYNNNNNNNTRTC | Pnuc_1010, TonB-dependent receptor |
| m6A | 1008661 | - | GAYANNNNNNNRTAG | upstream Pnuc_R0031, tRNA-Pro2 |
| m6A | 1009080 | + | | upstream Pnuc_1015, protein of unknown function UPF0044 |
| m4C | 1009251 | - | | Pnuc_1011, phosphoglucosamine mutase |
| m4C | 1009306 | + | | upstream Pnuc_1015, protein of unknown function UPF0044 |
| m4C | 1010615 | - | | Pnuc_1012, Dihydropteroate synthase |
| m4C | 1011804 | + | | upstream Pnuc_1015, protein of unknown function UPF0044 |
| m4C | 1013826 | - | | upstream Pnuc_1014, 23S rRNA Um-2552 2'-O-methyltransferase |
| m4C | 1015521 | - | | upstream Pnuc_1017, transcription elongation factor GreA |
| m6A | 1019053 | + | CTAYNNNNNNNTRTC | upstream Pnuc_1023, Methyltransferase type 11 |
| m4C | 1019054 | + | | upstream Pnuc_1023, Methyltransferase type 11 |
| m6A | 1019063 | - | GAYANNNNNNNRTAG | Pnuc_1019, carbamoyl-phosphate synthase small subunit |
| m4C | 1019978 | + | | upstream Pnuc_1023, Methyltransferase type 11 |
| m6A | 1020256 | + | CTAYNNNNNNNTRTC | upstream Pnuc_1023, Methyltransferase type 11 |
| m6A | 1020266 | - | GAYANNNNNNNRTAG | upstream Pnuc_1019, carbamoyl-phosphate synthase small subunit |
| m4C | 1020270 | - | | upstream Pnuc_1019, carbamoyl-phosphate synthase small subunit |
| m4C | 1020860 | - | | Pnuc_1020, propionyl-CoA synthetase |
| m4C | 1020960 | - | | Pnuc_1020, propionyl-CoA synthetase |
| m6A | 1021303 | + | CTAYNNNNNNNTRTC | upstream Pnuc_1023, Methyltransferase type 11 |
| m6A | 1021313 | - | GAYANNNNNNNRTAG | Pnuc_1020, propionyl-CoA synthetase |
| m4C | 1021990 | - | | Pnuc_1020, propionyl-CoA synthetase |
| m6A | 1027462 | + | GAYANNNNNNNRTAG | Pnuc_1026, acriflavin resistance protein |
| m6A | 1027472 | - | CTAYNNNNNNNTRTC | upstream Pnuc_1022, Hydroxyacylglutathione hydrolase |
| m6A | 1027824 | - | | upstream Pnuc_1022, Hydroxyacylglutathione hydrolase |
| m4C | 1030879 | - | | upstream Pnuc_1022, Hydroxyacylglutathione hydrolase |
| m4C | 1032865 | - | | Pnuc_1029, hypothetical protein |
| m6A | 1033357 | + | | upstream Pnuc_1032, Amidase |
| m6A | 1034059 | - | | Pnuc_1031, hypothetical protein |
| m6A | 1034060 | + | CTAYNNNNNNNTRTC | upstream Pnuc_1032, Amidase |
| m6A | 1034070 | - | GAYANNNNNNNRTAG | Pnuc_1031, hypothetical protein |
| m4C | 1035106 | - | | upstream Pnuc_1031, hypothetical protein |
| m4C | 1036445 | + | | upstream Pnuc_1035, methylmalonate-semialdehyde dehydrogenase(acylating) |
| m4C | 1037210 | + | | upstream Pnuc_1035, methylmalonate-semialdehyde dehydrogenase(acylating) |
| m4C | 1037402 | - | | Pnuc_1034, NADH:flavin oxidoreductase/NADH oxidase |
| m4C | 1037699 | - | | Pnuc_1034, NADH:flavin oxidoreductase/NADH oxidase |
| m4C | 1039663 | + | | Pnuc_1035, methylmalonate-semialdehyde dehydrogenase(acylating) |
| m6A | 1039882 | - | | upstream Pnuc_1034, NADH:flavin oxidoreductase/NADH oxidase |
| m4C | 1040324 | + | | upstream Pnuc_1057, putative lipoprotein |
| m4C | 1042147 | + | | upstream Pnuc_1057, putative lipoprotein |
| m4C | 1042450 | - | | Pnuc_1038, NADH dehydrogenase subunit N |
| m4C | 1042803 | + | | upstream Pnuc_1057, putative lipoprotein |
| m4C | 1043974 | - | | Pnuc_1039, NADH dehydrogenase subunit M |

| | | | | |
|---|---|---|---|---|
| m4C | 1044521 | - | | Pnuc_1040, NADH dehydrogenase subunit L |
| m4C | 1045576 | - | | Pnuc_1040, NADH dehydrogenase subunit L |
| m4C | 1047364 | - | | Pnuc_1043, NADH dehydrogenase subunit I |
| m4C | 1050439 | + | | upstream Pnuc_1057, putative lipoprotein |
| m6A | 1050989 | + | | upstream Pnuc_1057, putative lipoprotein |
| m4C | 1054222 | - | | Pnuc_1048, NADH dehydrogenase subunit D |
| m6A | 1055538 | + | GAYANNNNNNNRTAG | upstream Pnuc_1057, putative lipoprotein |
| m6A | 1055548 | - | CTAYNNNNNNNTRTC | Pnuc_1051, NADH dehydrogenase subunit A |
| m4C | 1057480 | + | | upstream Pnuc_1057, putative lipoprotein |
| m4C | 1060960 | + | | upstream Pnuc_1057, putative lipoprotein |
| m4C | 1062053 | + | | upstream Pnuc_1064, RNA polymerase, sigma-24 subunit, RpoE |
| m4C | 1062431 | + | | upstream Pnuc_1064, RNA polymerase, sigma-24 subunit, RpoE |
| m4C | 1062602 | + | | upstream Pnuc_1064, RNA polymerase, sigma-24 subunit, RpoE |
| m4C | 1063494 | + | | upstream Pnuc_1064, RNA polymerase, sigma-24 subunit, RpoE |
| m4C | 1063802 | + | | upstream Pnuc_1064, RNA polymerase, sigma-24 subunit, RpoE |
| m4C | 1066303 | - | | Pnuc_1062, acetolactate synthase, small subunit |
| m4C | 1067873 | + | | upstream Pnuc_1064, RNA polymerase, sigma-24 subunit, RpoE |
| m4C | 1068154 | + | | upstream Pnuc_1064, RNA polymerase, sigma-24 subunit, RpoE |
| m4C | 1068289 | + | | upstream Pnuc_1064, RNA polymerase, sigma-24 subunit, RpoE |
| m6A | 1069023 | - | | upstream Pnuc_1063, acetolactate synthase, large subunit |
| m4C | 1069538 | - | | upstream Pnuc_1063, acetolactate synthase, large subunit |
| m4C | 1069654 | + | | upstream Pnuc_1066, hypothetical protein |
| m4C | 1072487 | - | | Pnuc_1068, protein of unknown function DUF214 |
| m6A | 1075780 | + | GAYANNNNNNNRTAG | upstream Pnuc_1075, transcription elongation factor GreB |
| m6A | 1075790 | - | CTAYNNNNNNNTRTC | Pnuc_1071, hypothetical protein |
| m6A | 1076951 | + | CTAYNNNNNNNTRTC | upstream Pnuc_1075, transcription elongation factor GreB |
| m6A | 1076961 | - | GAYANNNNNNNRTAG | Pnuc_1073, ABC transporter related protein |
| m4C | 1080829 | - | | Pnuc_1074, protein of unknown function DUF140 |
| m6A | 1092756 | - | | upstream Pnuc_1085, integral membrane sensor signal transduction histidine kinase |
| m6A | 1093084 | + | GAYANNNNNNNRTAG | Pnuc_1087, hypothetical protein |
| m6A | 1093094 | - | CTAYNNNNNNNTRTC | upstream Pnuc_1085, integral membrane sensor signal transduction histidine kinase |
| m6A | 1095712 | + | CTAYNNNNNNNTRTC | upstream Pnuc_1091, glucose-6-phosphate isomerase |
| m6A | 1095722 | - | GAYANNNNNNNRTAG | upstream Pnuc_1089, hypothetical protein |
| m6A | 1096119 | - | | upstream Pnuc_1089, hypothetical protein |
| m6A | 1096606 | - | | Pnuc_1090, FAD dependent oxidoreductase |
| m4C | 1106620 | + | | upstream Pnuc_1097, hypothetical protein |
| m4C | 1107202 | + | | upstream Pnuc_1097, hypothetical protein |
| m6A | 1108023 | + | GAYANNNNNNNRTAG | upstream Pnuc_1097, hypothetical protein |
| m6A | 1108033 | - | CTAYNNNNNNNTRTC | Pnuc_1095, outer membrane autotransporter barrel domain protein |
| m4C | 1108297 | + | | upstream Pnuc_1097, hypothetical protein |
| m6A | 1108814 | - | | Pnuc_1095, outer membrane autotransporter barrel domain protein |
| m6A | 1109515 | + | CTAYNNNNNNNTRTC | upstream Pnuc_1097, hypothetical protein |
| m6A | 1109525 | - | GAYANNNNNNNRTAG | Pnuc_1095, outer membrane autotransporter barrel domain protein |
| m4C | 1111018 | + | | upstream Pnuc_1097, hypothetical protein |

| | | | | |
|---|---|---|---|---|
| m6A | 1111532 | + | | upstream Pnuc_1097, hypothetical protein |
| m6A | 1111707 | + | GAYANNNNNNNRTAG | upstream Pnuc_1097, hypothetical protein |
| m6A | 1111717 | - | CTAYNNNNNNNTRTC | Pnuc_1095, outer membrane autotransporter barrel domain protein |
| m6A | 1111938 | + | | upstream Pnuc_1097, hypothetical protein |
| m6A | 1115916 | + | GAYANNNNNNNRTAG | upstream Pnuc_1097, hypothetical protein |
| m6A | 1115926 | - | CTAYNNNNNNNTRTC | Pnuc_1095, outer membrane autotransporter barrel domain protein |
| m4C | 1118871 | - | | Pnuc_1095, outer membrane autotransporter barrel domain protein |
| m6A | 1118964 | - | | Pnuc_1095, outer membrane autotransporter barrel domain protein |
| m4C | 1119946 | + | | upstream Pnuc_1097, hypothetical protein |
| m6A | 1120460 | + | | upstream Pnuc_1097, hypothetical protein |
| m4C | 1120603 | + | | upstream Pnuc_1097, hypothetical protein |
| m6A | 1124249 | + | | upstream Pnuc_1097, hypothetical protein |
| m6A | 1124256 | - | | Pnuc_1095, outer membrane autotransporter barrel domain protein |
| m4C | 1126657 | - | | Pnuc_1095, outer membrane autotransporter barrel domain protein |
| m4C | 1131493 | + | | upstream Pnuc_1097, hypothetical protein |
| m6A | 1131887 | + | | upstream Pnuc_1097, hypothetical protein |
| m6A | 1134204 | - | | upstream Pnuc_1095, outer membrane autotransporter barrel domain protein |
| m4C | 1141529 | + | | upstream Pnuc_1101, protein of unknown function DUF583 |
| m6A | 1141757 | - | | Pnuc_1098, methyltransferase FkbM family |
| m4C | 1144057 | + | | Pnuc_1101, protein of unknown function DUF583 |
| m6A | 1146350 | + | | Pnuc_1104, nicotinate-nucleotide pyrophosphorylase(carboxylating) |
| m6A | 1147304 | - | | Pnuc_1105, TRAP transporter solute receptor, TAXI family |
| m6A | 1150445 | + | GAYANNNNNNNRTAG | upstream Pnuc_1112, protein of unknown function DUF1080 |
| m6A | 1150455 | - | CTAYNNNNNNNTRTC | Pnuc_1107, transcriptional modulator of MazE/toxin, MazF |
| m6A | 1151487 | + | GAYANNNNNNNRTAG | upstream Pnuc_1112, protein of unknown function DUF1080 |
| m6A | 1151497 | - | CTAYNNNNNNNTRTC | Pnuc_1109, NAD-dependent formate dehydrogenase iron-sulfur protein (catalytic activity) |
| m4C | 1152128 | - | | Pnuc_1109, NAD-dependent formate dehydrogenase iron-sulfur protein (catalytic activity) |
| m6A | 1156346 | + | CTAYNNNNNNNTRTC | upstream Pnuc_1112, protein of unknown function DUF1080 |
| m6A | 1156356 | - | GAYANNNNNNNRTAG | Pnuc_1111, protein of unknown function DUF748 |
| m6A | 1156357 | + | | upstream Pnuc_1112, protein of unknown function DUF1080 |
| m6A | 1156484 | - | | Pnuc_1111, protein of unknown function DUF748 |
| m4C | 1156774 | - | | Pnuc_1111, protein of unknown function DUF749 |
| m6A | 1159304 | + | GAYANNNNNNNRTAG | Pnuc_1112, protein of unknown function DUF1080 |
| m6A | 1159314 | - | CTAYNNNNNNNTRTC | upstream Pnuc_1111, protein of unknown function DUF748 |
| m6A | 1159459 | + | | upstream Pnuc_1116, phage integrase family protein |
| m6A | 1163127 | - | | Pnuc_1118, SMC domain protein |
| m4C | 1164252 | - | | upstream Pnuc_1118, SMC domain protein |
| m6A | 1165285 | - | | Pnuc_1120, hypothetical protein |
| m6A | 1167044 | + | CTAYNNNNNNNTRTC | Pnuc_1122, Exonuclease, RNase T and DNA polymerase III |
| m6A | 1167054 | - | GAYANNNNNNNRTAG | upstream Pnuc_1121, hypothetical protein |
| m4C | 1169095 | - | | upstream Pnuc_1121, hypothetical protein |
| m4C | 1169873 | + | | upstream Pnuc_1127, hypothetical protein |
| m4C | 1170203 | + | | upstream Pnuc_1127, hypothetical protein |
| m4C | 1173732 | + | | upstream Pnuc_1127, hypothetical protein |

| | | | | |
|---|---|---|---|---|
| m6A | 1173977 | + | GAYANNNNNNNRTAG | Pnuc_1127, hypothetical protein |
| m6A | 1173987 | - | CTAYNNNNNNNTRTC | upstream Pnuc_1126, hypothetical protein |
| m6A | 1177319 | + | GAYANNNNNNNRTAG | upstream Pnuc_1131, phage transcriptional regulator, AlpA |
| m6A | 1177329 | - | CTAYNNNNNNNTRTC | Pnuc_1130, protein of unknown function DUF1376 |
| m4C | 1177539 | + | | upstream Pnuc_1131, phage transcriptional regulator, AlpA |
| m6A | 1178648 | - | | Pnuc_1132, restriction modification system DNA specificity domain protein |
| m4C | 1188747 | + | | upstream Pnuc_1139, fumarase |
| m4C | 1189031 | + | | upstream Pnuc_1139, fumarase |
| m6A | 1189221 | - | | upstream Pnuc_1138, acetyl-coenzyme A synthetase |
| m4C | 1189269 | + | | Pnuc_1139, fumarase |
| m4C | 1190609 | + | | Pnuc_1139, fumarase |
| m6A | 1191165 | + | CTAYNNNNNNNTRTC | Pnuc_1140, glutamate racemase |
| m6A | 1191175 | - | GAYANNNNNNNRTAG | upstream Pnuc_1138, acetyl-coenzyme A synthetase |
| m4C | 1193460 | + | | upstream Pnuc_1150, Penicillin amidase |
| m4C | 1194068 | + | | upstream Pnuc_1150, Penicillin amidase |
| m4C | 1195742 | - | | Pnuc_1146, phosphate ABC transporter substrate-binding protein, PhoT family |
| m6A | 1199870 | + | GAYANNNNNNNRTAG | upstream Pnuc_1150, Penicillin amidase |
| m6A | 1199880 | - | CTAYNNNNNNNTRTC | Pnuc_1149, filamentous hemagglutinin family outer membrane protein |
| m4C | 1200717 | + | | upstream Pnuc_1150, Penicillin amidase |
| m4C | 1202402 | + | | upstream Pnuc_1150, Penicillin amidase |
| m4C | 1204358 | - | | Pnuc_R0036, tRNA-Leu |
| m4C | 1204364 | - | | Pnuc_R0036, tRNA-Leu |
| m6A | 1206854 | + | CTAYNNNNNNNTRTC | Pnuc_1150, Penicillin amidase |
| m6A | 1206864 | - | GAYANNNNNNNRTAG | upstream Pnuc_R0036, tRNA-Leu |
| m4C | 1209983 | - | | Pnuc_1156, hypothetical protein |
| m4C | 1214543 | + | | Pnuc_1161, LrgB family protein |
| m4C | 1216992 | + | | Pnuc_1163, gamma-glutamyltransferase 2, Threonine peptidase, MEROPS family T03 |
| m6A | 1218238 | + | CTAYNNNNNNNTRTC | upstream Pnuc_1167, Cellulase |
| m6A | 1218248 | - | GAYANNNNNNNRTAG | Pnuc_1164, conserved hypothetical protein |
| m4C | 1223679 | + | | Pnuc_1168, conserved hypothetical protein |
| m6A | 1223697 | + | | Pnuc_1168, conserved hypothetical protein |
| m4C | 1225020 | + | | upstream Pnuc_1176, hypothetical protein |
| m4C | 1225259 | + | | upstream Pnuc_1176, hypothetical protein |
| m4C | 1225329 | - | | Pnuc_1170, cellulose synthase operon C domain protein |
| m4C | 1225578 | + | | upstream Pnuc_1176, hypothetical protein |
| m6A | 1228000 | + | GAYANNNNNNNRTAG | upstream Pnuc_1176, hypothetical protein |
| m6A | 1228010 | - | CTAYNNNNNNNTRTC | Pnuc_1170, cellulose synthase operon C domain protein |
| m6A | 1228683 | + | GAYANNNNNNNRTAG | upstream Pnuc_1176, hypothetical protein |
| m6A | 1228693 | - | CTAYNNNNNNNTRTC | Pnuc_1170, cellulose synthase operon C domain protein |
| m6A | 1228990 | + | CTAYNNNNNNNTRTC | upstream Pnuc_1176, hypothetical protein |
| m6A | 1229362 | - | | Pnuc_1171, Cellulose synthase (UDP-forming) |
| m4C | 1229846 | + | | upstream Pnuc_1176, hypothetical protein |
| m4C | 1229864 | + | | upstream Pnuc_1176, hypothetical protein |
| m4C | 1230976 | - | | Pnuc_1171, Cellulose synthase (UDP-forming) |

| | | | | |
|---|---|---|---|---|
| m4C | 1233413 | + | | upstream Pnuc_1176, hypothetical protein |
| m4C | 1235766 | + | | upstream Pnuc_1184, Amidase |
| m4C | 1236389 | + | | upstream Pnuc_1184, Amidase |
| m6A | 1236712 | + | CTAYNNNNNNNNTRTC | upstream Pnuc_1184, Amidase |
| m6A | 1236722 | - | GAYANNNNNNNNRTAG | upstream Pnuc_1177, tRNA-U16,U17-dihydrouridine synthase |
| m6A | 1239968 | + | GAYANNNNNNNNRTAG | upstream Pnuc_1184, Amidase |
| m6A | 1239978 | - | CTAYNNNNNNNNTRTC | Pnuc_1180, putative ABC transporter, periplasmic protein |
| m6A | 1240127 | + | GAYANNNNNNNNRTAG | upstream Pnuc_1184, Amidase |
| m6A | 1240137 | - | CTAYNNNNNNNNTRTC | Pnuc_1180, putative ABC transporter, periplasmic protein |
| m4C | 1241022 | + | | upstream Pnuc_1184, Amidase |
| m4C | 1245515 | - | | upstream Pnuc_1183, conserved hypothetical protein |
| m6A | 1247968 | + | CTAYNNNNNNNNTRTC | upstream Pnuc_1210, Glutamate synthase (NADPH) |
| m6A | 1247978 | - | GAYANNNNNNNNRTAG | Pnuc_1190, urease accessory protein UreG |
| m4C | 1250367 | - | | Pnuc_1193, urease, Metallo peptidase, MEROPS family M38 |
| m4C | 1251140 | - | | Pnuc_1193, urease, Metallo peptidase, MEROPS family M39 |
| m6A | 1252457 | + | GAYANNNNNNNNRTAG | upstream Pnuc_1210, Glutamate synthase (NADPH) |
| m6A | 1252467 | - | CTAYNNNNNNNNTRTC | Pnuc_1196, urease, gamma subunit |
| m6A | 1253887 | + | | upstream Pnuc_1210, Glutamate synthase (NADPH) |
| m4C | 1260810 | - | | Pnuc_1203, secretory lipase |
| m6A | 1262425 | + | CTAYNNNNNNNNTRTC | upstream Pnuc_1210, Glutamate synthase (NADPH) |
| m6A | 1262435 | - | GAYANNNNNNNNRTAG | Pnuc_1205, protein of unknown function DUF1501 |
| m6A | 1263897 | - | | Pnuc_1206, conserved hypothetical protein |
| m4C | 1266245 | + | | Pnuc_1210, Glutamate synthase (NADPH) |
| m6A | 1268677 | + | | upstream Pnuc_1218, transglutaminase, N-terminal domain protein |
| m6A | 1269593 | + | | upstream Pnuc_1218, transglutaminase, N-terminal domain protein |
| m4C | 1270788 | - | | Pnuc_1214, intracellular polyhydroxyalkanoate depolymerase |
| m6A | 1273368 | - | | upstream Pnuc_1217, Glutathione S-transferase, N-terminal domain protein |
| m4C | 1278459 | - | | upstream Pnuc_1217, Glutathione S-transferase, N-terminal domain protein |
| m6A | 1278858 | - | | upstream Pnuc_1217, Glutathione S-transferase, N-terminal domain protein |
| m4C | 1280244 | - | | Pnuc_1221, conserved hypothetical protein |
| m4C | 1281881 | + | | upstream Pnuc_1223, major facilitator superfamily MFS_1 |
| m4C | 1287845 | + | | upstream Pnuc_1232, Patatin |
| m4C | 1289753 | + | | upstream Pnuc_1232, Patatin |
| m4C | 1291441 | - | | Pnuc_1229, protein of unknown function DUF150 |
| m6A | 1292600 | + | | upstream Pnuc_1232, Patatin |
| m4C | 1294041 | - | | Pnuc_1231, condensin subunit ScpB |
| m4C | 1294062 | - | | Pnuc_1231, condensin subunit ScpB |
| m4C | 1295793 | - | | Pnuc_1233, NLP/P60 protein |
| m6A | 1296429 | + | CTAYNNNNNNNNTRTC | upstream Pnuc_1238, Enoyl-[acyl-carrier-protein] reductase (NADH) |
| m6A | 1296439 | - | GAYANNNNNNNNRTAG | upstream Pnuc_1233, NLP/P60 protein |
| m6A | 1296443 | - | | upstream Pnuc_1233, NLP/P60 protein |
| m6A | 1296502 | - | | Pnuc_1234, ABC transporter related protein |
| m4C | 1296561 | + | | upstream Pnuc_1238, Enoyl-[acyl-carrier-protein] reductase (NADH) |
| m4C | 1296881 | + | | upstream Pnuc_1238, Enoyl-[acyl-carrier-protein] reductase (NADH) |

| | | | | |
|---|---|---|---|---|
| m4C | 1302181 | - | | upstream Pnuc_1237, extracellular solute-binding protein, family 5 |
| m4C | 1302290 | + | | Pnuc_1238, Enoyl-[acyl-carrier-protein] reductase (NADH) |
| m4C | 1302491 | + | | Pnuc_1238, Enoyl-[acyl-carrier-protein] reductase (NADH) |
| m4C | 1302831 | - | | upstream Pnuc_1237, extracellular solute-binding protein, family 5 |
| m6A | 1307149 | + | GAYANNNNNNNRTAG | Pnuc_1242, MmgE/PrpD family protein |
| m6A | 1307159 | - | CTAYNNNNNNNTRTC | upstream Pnuc_1237, extracellular solute-binding protein, family 5 |
| m4C | 1309811 | + | | upstream Pnuc_1249, integrase, catalytic region |
| m6A | 1311023 | - | | Pnuc_1246, BNR/Asp-box repeat protein |
| m6A | 1311414 | + | CTAYNNNNNNNTRTC | upstream Pnuc_1249, integrase, catalytic region |
| m6A | 1311424 | - | GAYANNNNNNNRTAG | Pnuc_1246, BNR/Asp-box repeat protein |
| m4C | 1315036 | - | | upstream Pnuc_1248, hypothetical protein |
| m6A | 1317420 | + | CTAYNNNNNNNTRTC | upstream Pnuc_1262, N-acetylglutamate synthase |
| m6A | 1317430 | - | GAYANNNNNNNRTAG | Pnuc_1252, Exodeoxyribonuclease III |
| m4C | 1318116 | + | | upstream Pnuc_1262, N-acetylglutamate synthase |
| m4C | 1320245 | - | | Pnuc_1254, signal transduction histidine kinase, nitrogen specific, NtrB |
| m6A | 1322292 | - | | Pnuc_1256, molybdopterin binding domain protein |
| m6A | 1330103 | + | | upstream Pnuc_1262, N-acetylglutamate synthase |
| m4C | 1330799 | + | | Pnuc_1262, N-acetylglutamate synthase |
| m4C | 1330809 | + | | Pnuc_1262, N-acetylglutamate synthase |
| m4C | 1330848 | - | | upstream Pnuc_1261, ATP-dependent helicase HrpA |
| m4C | 1332282 | - | | upstream Pnuc_1261, ATP-dependent helicase HrpA |
| m4C | 1336306 | + | | upstream Pnuc_R0038, tRNA-Leu3 |
| m4C | 1336698 | - | | Pnuc_1267, propionate CoA-transferase |
| m4C | 1337345 | + | | upstream Pnuc_R0038, tRNA-Leu3 |
| m4C | 1337444 | - | | Pnuc_1267, propionate CoA-transferase |
| m4C | 1344160 | + | | Pnuc_1272, amino acid permease-associated region |
| m4C | 1349236 | + | | upstream Pnuc_1294, protein of unknown function UPF0005 |
| m4C | 1353321 | + | | upstream Pnuc_1294, protein of unknown function UPF0005 |
| m4C | 1356193 | - | | Pnuc_1284, protease FtsH subunit HflK |
| m4C | 1357004 | + | | upstream Pnuc_1294, protein of unknown function UPF0005 |
| m4C | 1357054 | + | | upstream Pnuc_1294, protein of unknown function UPF0005 |
| m4C | 1357572 | - | | upstream Pnuc_1285, GTP-binding protein HflX |
| m4C | 1357629 | + | | upstream Pnuc_1294, protein of unknown function UPF0005 |
| m4C | 1358238 | - | | Pnuc_1287, small GTP-binding protein |
| m4C | 1358779 | - | | Pnuc_1287, small GTP-binding protein |
| m4C | 1358810 | - | | Pnuc_1287, small GTP-binding protein |
| m4C | 1367849 | + | | upstream Pnuc_1307, thioredoxin |
| m4C | 1372071 | + | | upstream Pnuc_1307, thioredoxin |
| m6A | 1373052 | + | | upstream Pnuc_1307, thioredoxin |
| m6A | 1373136 | + | | upstream Pnuc_1307, thioredoxin |
| m4C | 1375771 | - | | Pnuc_1302, outer membrane efflux protein |
| m4C | 1376040 | + | | upstream Pnuc_1307, thioredoxin |
| m4C | 1376075 | + | | upstream Pnuc_1307, thioredoxin |
| m4C | 1377069 | - | | Pnuc_1302, outer membrane efflux protein |

| | | | | |
|---|---|---|---|---|
| m6A | 1377692 | + | | upstream Pnuc_1307, thioredoxin |
| m4C | 1379860 | + | | upstream Pnuc_1307, thioredoxin |
| m4C | 1380179 | + | | upstream Pnuc_1307, thioredoxin |
| m4C | 1383112 | + | | upstream Pnuc_1307, thioredoxin |
| m4C | 1383881 | - | | Pnuc_1306, DNA helicase/exodeoxyribonuclease V, subunit B |
| m6A | 1384156 | + | | upstream Pnuc_1307, thioredoxin |
| m4C | 1387718 | - | | upstream Pnuc_1306, DNA helicase/exodeoxyribonuclease V, subunit B |
| m6A | 1387744 | - | | upstream Pnuc_1306, DNA helicase/exodeoxyribonuclease V, subunit B |
| m4C | 1387948 | + | | Pnuc_1309, LSU ribosomal protein L31P |
| m4C | 1391088 | - | | upstream Pnuc_1306, DNA helicase/exodeoxyribonuclease V, subunit B |
| m4C | 1391890 | + | | upstream Pnuc_1318, Enoyl-CoA hydratase/isomerase |
| m4C | 1392764 | + | | upstream Pnuc_1318, Enoyl-CoA hydratase/isomerase |
| m4C | 1393403 | - | | Pnuc_1314, conserved hypothetical protein |
| m6A | 1393669 | - | | Pnuc_1315, conserved hypothetical protein |
| m4C | 1396783 | + | | upstream Pnuc_1318, Enoyl-CoA hydratase/isomerase |
| m6A | 1399985 | - | | Pnuc_1321, fumarylacetoacetate (FAA) hydrolase |
| m4C | 1400775 | + | | Pnuc_1323, Uncharacterized protein UPF0065 |
| m4C | 1405924 | - | | Pnuc_1328, 5-oxopent-3-ene-1,2,5-tricarboxylate decarboxylase |
| m6A | 1406779 | - | | Pnuc_1329, ferrous iron transport protein B |
| m6A | 1407240 | + | | upstream Pnuc_1331, Carbohydrate-selective porin OprB |
| m6A | 1410626 | + | GAYANNNNNNNRTAG | Pnuc_1332, negative transcriptional regulator |
| m6A | 1410636 | - | CTAYNNNNNNNTRTC | upstream Pnuc_1330, FeoA family protein |
| m6A | 1412582 | + | | upstream Pnuc_1336, putative sulfate transport system substrate-binding protein |
| m6A | 1416244 | + | | upstream Pnuc_1339, GCN5-related N-acetyltransferase |
| m6A | 1416333 | + | | Pnuc_1339, GCN5-related N-acetyltransferase |
| m6A | 1416762 | + | CTAYNNNNNNNTRTC | Pnuc_1340, thioesterase superfamily protein |
| m6A | 1416772 | - | GAYANNNNNNNRTAG | upstream Pnuc_1338, hypothetical protein |
| m6A | 1417579 | + | GAYANNNNNNNRTAG | Pnuc_1341, conserved hypothetical protein |
| m6A | 1417589 | - | CTAYNNNNNNNTRTC | upstream Pnuc_1338, hypothetical protein |
| m4C | 1419056 | + | | Pnuc_1342, DEAD/DEAH box helicase domain protein |
| m6A | 1419263 | + | CTAYNNNNNNNTRTC | Pnuc_1343, transglutaminase domain protein |
| m6A | 1419273 | - | GAYANNNNNNNRTAG | upstream Pnuc_1338, hypothetical protein |
| m6A | 1419805 | + | | Pnuc_1343, transglutaminase domain protein |
| m4C | 1419935 | - | | upstream Pnuc_1338, hypothetical protein |
| m6A | 1420785 | - | | upstream Pnuc_1338, hypothetical protein |
| m6A | 1421798 | - | | upstream Pnuc_1338, hypothetical protein |
| m6A | 1422263 | + | CTAYNNNNNNNTRTC | upstream Pnuc_1347, protein of unknown function DUF1330 |
| m6A | 1422273 | - | GAYANNNNNNNRTAG | upstream Pnuc_1346, conserved hypothetical protein |
| m6A | 1422850 | + | | upstream Pnuc_1349, Uncharacterized protein UPF0065 |
| m4C | 1427357 | + | | Pnuc_1353, hypothetical protein |
| m4C | 1429892 | + | | upstream Pnuc_1360, hypothetical protein |
| m6A | 1430406 | - | | Pnuc_1357, HI0933 family protein |
| m4C | 1433867 | + | | upstream Pnuc_1363, carbohydrate kinase, YjeF related protein |
| m6A | 1434445 | + | CTAYNNNNNNNTRTC | upstream Pnuc_1363, carbohydrate kinase, YjeF related protein |

| | | | | |
|---|---|---|---|---|
| m6A | 1434455 | - | GAYANNNNNNNRTAG | Pnuc_1362, RNA-metabolising metallo-beta-lactamase |
| m6A | 1436527 | + | | Pnuc_1364, protein of unknown function DUF6, transmembrane |
| m4C | 1438928 | - | | Pnuc_1368, Uncharacterized protein UPF0065 |
| m4C | 1440052 | + | | upstream Pnuc_1371, L-lactate dehydrogenase (cytochrome) |
| m4C | 1444140 | - | | upstream Pnuc_1370, protein of unknown function DUF6, transmembrane |
| m6A | 1444335 | - | | upstream Pnuc_1370, protein of unknown function DUF6, transmembrane |
| m6A | 1460924 | + | | upstream Pnuc_1392, binding-protein-dependent transport systems inner membrane component |
| m4C | 1461233 | - | | Pnuc_1390, cardiolipin synthetase 2 |
| m4C | 1462691 | + | | upstream Pnuc_1392, binding-protein-dependent transport systems inner membrane component |
| m6A | 1463623 | - | | upstream Pnuc_1391, Endonuclease/exonuclease/phosphatase |
| m6A | 1464729 | + | GAYANNNNNNNRTAG | upstream Pnuc_1407, heavy metal translocating P-type ATPase |
| m6A | 1464739 | - | CTAYNNNNNNNTRTC | Pnuc_1395, formate dehydrogenase gamma subunit |
| m4C | 1468422 | - | | Pnuc_1398, molybdopterin oxidoreductase |
| m4C | 1470425 | + | | upstream Pnuc_1407, heavy metal translocating P-type ATPase |
| m4C | 1470899 | + | | upstream Pnuc_1407, heavy metal translocating P-type ATPase |
| m6A | 1471413 | - | | Pnuc_1400, 4Fe-4S ferredoxin, iron-sulfur binding domain protein |
| m6A | 1471884 | + | | upstream Pnuc_1407, heavy metal translocating P-type ATPase |
| m6A | 1473559 | + | | upstream Pnuc_1407, heavy metal translocating P-type ATPase |
| m6A | 1473645 | + | GAYANNNNNNNRTAG | upstream Pnuc_1407, heavy metal translocating P-type ATPase |
| m6A | 1473655 | - | CTAYNNNNNNNTRTC | Pnuc_1402, conserved hypothetical protein |
| m4C | 1473939 | - | | Pnuc_1403, protein of unknown function DUF59 |
| m4C | 1475423 | + | | upstream Pnuc_1407, heavy metal translocating P-type ATPase |
| m4C | 1475449 | - | | Pnuc_1404, phosphate transporter |
| m4C | 1477177 | + | | Pnuc_1407, heavy metal translocating P-type ATPase |
| m6A | 1478429 | - | | upstream Pnuc_1406, Heavy metal transport/detoxification protein |
| m4C | 1483414 | - | | upstream Pnuc_1408, conserved hypothetical protein |
| m6A | 1483806 | - | | upstream Pnuc_1408, conserved hypothetical protein |
| m4C | 1484502 | - | | upstream Pnuc_1408, conserved hypothetical protein |
| m6A | 1485343 | + | | Pnuc_1413, protein of unknown function DUF519 |
| m4C | 1488371 | - | | Pnuc_1418, Radical SAM domain protein |
| m6A | 1488814 | + | | upstream Pnuc_1426, SsrA-binding protein |
| m4C | 1493575 | - | | Pnuc_1422, GMP synthase (glutamine-hydrolyzing) |
| m4C | 1495693 | - | | Pnuc_1424, protein of unknown function UPF0125 |
| m6A | 1496168 | + | CTAYNNNNNNNTRTC | upstream Pnuc_1426, SsrA-binding protein |
| m6A | 1496178 | - | GAYANNNNNNNRTAG | Pnuc_1425, cyclase/dehydrase |
| m4C | 1496785 | - | | upstream Pnuc_1425, cyclase/dehydrase |
| m4C | 1498734 | + | | Pnuc_1428, protein of unknown function DUF403 |
| m4C | 1498747 | + | | Pnuc_1428, protein of unknown function DUF403 |
| m4C | 1500795 | - | | upstream Pnuc_1425, cyclase/dehydrase |
| m4C | 1502448 | + | | Pnuc_1432, Integrase, catalytic region |
| m4C | 1503975 | - | | Pnuc_1433, phosphoenolpyruvate synthase |
| m6A | 1505598 | + | CTAYNNNNNNNTRTC | Pnuc_1434, protein of unknown function DUF299 |
| m6A | 1505608 | - | GAYANNNNNNNRTAG | Pnuc_1433, phosphoenolpyruvate synthase |
| m4C | 1510301 | + | | upstream Pnuc_1452, methionine aminopeptidase, type I |

| | | | | |
|---|---|---|---|---|
| m6A | 1519356 | + | CTAYNNNNNNNTRTC | upstream Pnuc_1452, methionine aminopeptidase, type I |
| m6A | 1519366 | - | GAYANNNNNNNRTAG | Pnuc_1448, ribosome recycling factor |
| m4C | 1522785 | + | | Pnuc_1452, methionine aminopeptidase, type I |
| m4C | 1523178 | + | | Pnuc_1452, methionine aminopeptidase, type I |
| m6A | 1526556 | + | | upstream Pnuc_1458, 2,3,4,5-tetrahydropyridine-2,6-dicarboxylate N-succinyltransferase |
| m4C | 1529296 | - | | Pnuc_1456, ZipA, C-terminal FtsZ-binding region |
| m4C | 1530031 | + | | upstream Pnuc_1458, 2,3,4,5-tetrahydropyridine-2,6-dicarboxylate N-succinyltransferase |
| m4C | 1530327 | + | | upstream Pnuc_1458, 2,3,4,5-tetrahydropyridine-2,6-dicarboxylate N-succinyltransferase |
| m4C | 1530420 | + | | upstream Pnuc_1458, 2,3,4,5-tetrahydropyridine-2,6-dicarboxylate N-succinyltransferase |
| m6A | 1530582 | - | | Pnuc_1457, chromosome segregation protein SMC |
| m6A | 1532841 | - | | Pnuc_1457, chromosome segregation protein SMC |
| m4C | 1537844 | + | | upstream Pnuc_R0039, tRNA-Met1 |
| m4C | 1538994 | + | | upstream Pnuc_R0039, tRNA-Met1 |
| m6A | 1540249 | + | | upstream Pnuc_R0039, tRNA-Met1 |
| m4C | 1541125 | - | | Pnuc_1465, DNA internalization-related competence protein ComEC/Rec2 |
| m4C | 1544837 | - | | Pnuc_1467, ABC transporter related protein |
| m6A | 1550047 | + | | Pnuc_1471, bacterial peptide chain release factor 2(bRF-2) |
| m6A | 1550543 | + | | Pnuc_1472, lysyl-tRNA synthetase |
| m4C | 1551848 | + | | upstream Pnuc_1477, peptide deformylase |
| m6A | 1552414 | + | GAYANNNNNNNRTAG | upstream Pnuc_1477, peptide deformylase |
| m6A | 1552424 | - | CTAYNNNNNNNTRTC | Pnuc_1473, hypothetical protein |
| m6A | 1554164 | - | | Pnuc_1476, cysteine synthase A |
| m6A | 1557612 | + | CTAYNNNNNNNTRTC | Pnuc_1480, protein of unknown function DUF81 |
| m6A | 1557622 | - | GAYANNNNNNNRTAG | upstream Pnuc_1478, disulfide bond formation protein DsbB |
| m4C | 1565765 | - | | Pnuc_1489, Fe-S protein assembly chaperone HscA |
| m4C | 1566604 | - | | Pnuc_1489, Fe-S protein assembly chaperone HscA |
| m4C | 1566777 | + | | upstream Pnuc_1496, aminotransferase |
| m4C | 1568086 | + | | upstream Pnuc_1496, aminotransferase |
| m4C | 1568266 | - | | upstream Pnuc_1492, FeS cluster assembly scaffold IscU |
| m6A | 1568411 | + | | upstream Pnuc_1496, aminotransferase |
| m6A | 1569027 | + | | upstream Pnuc_1496, aminotransferase |
| m4C | 1569924 | - | | Pnuc_1494, transcriptional regulator, BadM/Rrf2 family |
| m4C | 1570822 | - | | Pnuc_1495, Excinuclease ABC subunit B |
| m6A | 1570842 | + | CTAYNNNNNNNTRTC | upstream Pnuc_1496, aminotransferase |
| m6A | 1570852 | - | GAYANNNNNNNRTAG | Pnuc_1495, Excinuclease ABC subunit B |
| m4C | 1571584 | - | | Pnuc_1495, Excinuclease ABC subunit B |
| m4C | 1572033 | - | | Pnuc_1495, Excinuclease ABC subunit B |
| m4C | 1574196 | - | | upstream Pnuc_1495, Excinuclease ABC subunit B |
| m4C | 1575410 | + | | Pnuc_1498, electron transport complex, RnfABCDGE type, B subunit |
| m4C | 1578103 | + | | upstream Pnuc_1528, conserved hypothetical protein |
| m4C | 1579573 | - | | Pnuc_1504, 3-oxoadipate CoA-transferase alpha subunit |
| m6A | 1580069 | + | | upstream Pnuc_1528, conserved hypothetical protein |
| m4C | 1581325 | + | | upstream Pnuc_1528, conserved hypothetical protein |
| m6A | 1582960 | + | GAYANNNNNNNRTAG | upstream Pnuc_1528, conserved hypothetical protein |

| | | | | |
|---|---|---|---|---|
| m6A | 1582970 | - | CTAYNNNNNNNTRTC | Pnuc_1507, major facilitator superfamily MFS_1 |
| m4C | 1583023 | - | | Pnuc_1507, major facilitator superfamily MFS_1 |
| m4C | 1586886 | + | | upstream Pnuc_1528, conserved hypothetical protein |
| m4C | 1587174 | - | | Pnuc_1511, cytochrome C biogenesis protein |
| m4C | 1588023 | - | | Pnuc_1513, cytochrome c-type biogenesis protein CcmF |
| m4C | 1588814 | - | | upstream Pnuc_1513, cytochrome c-type biogenesis protein CcmF |
| m4C | 1591196 | - | | Pnuc_1517, heme exporter protein CcmB |
| m4C | 1591513 | - | | Pnuc_1517, heme exporter protein CcmB |
| m6A | 1591897 | + | CTAYNNNNNNNTRTC | upstream Pnuc_1528, conserved hypothetical protein |
| m6A | 1591907 | - | GAYANNNNNNNRTAG | Pnuc_1518, heme exporter protein CcmA |
| m4C | 1592173 | + | | upstream Pnuc_1528, conserved hypothetical protein |
| m4C | 1595365 | + | | upstream Pnuc_1528, conserved hypothetical protein |
| m6A | 1595390 | - | | upstream Pnuc_1523, transcriptional regulator, MarR family |
| m4C | 1595553 | - | | Pnuc_1524, beta-lactamase domain protein |
| m4C | 1595665 | - | | Pnuc_1524, beta-lactamase domain protein |
| m4C | 1595861 | - | | Pnuc_1524, beta-lactamase domain protein |
| m6A | 1596988 | + | | upstream Pnuc_1528, conserved hypothetical protein |
| m4C | 1598464 | + | | upstream Pnuc_1528, conserved hypothetical protein |
| m6A | 1599496 | + | CTAYNNNNNNNTRTC | Pnuc_1529, histone deacetylase superfamily |
| m6A | 1599506 | - | GAYANNNNNNNRTAG | upstream Pnuc_1527, hypothetical protein |
| m6A | 1601203 | + | GAYANNNNNNNRTAG | upstream Pnuc_1534, alkyl hydroperoxide reductase/ Thiol specific antioxidant/ Mal allergen |
| m6A | 1601213 | - | CTAYNNNNNNNTRTC | Pnuc_1531, putative lipoprotein |
| m6A | 1602318 | - | | Pnuc_1533, hypothetical protein |
| m4C | 1602809 | + | | upstream Pnuc_1534, alkyl hydroperoxide reductase/ Thiol specific antioxidant/ Mal allergen |
| m4C | 1605257 | + | | Pnuc_1535, FAD-dependent pyridine nucleotide-disulfide oxidoreductase |
| m4C | 1606133 | + | | upstream Pnuc_1544, transcriptional regulator, GntR family |
| m6A | 1606433 | + | GAYANNNNNNNRTAG | upstream Pnuc_1544, transcriptional regulator, GntR family |
| m6A | 1606443 | - | CTAYNNNNNNNTRTC | Pnuc_1538, TRAP C4-dicarboxylate transport system permease DctM subunit |
| m4C | 1606519 | - | | Pnuc_1538, TRAP C4-dicarboxylate transport system permease DctM subunit |
| m6A | 1606979 | - | | Pnuc_1538, TRAP C4-dicarboxylate transport system permease DctM subunit |
| m4C | 1608288 | - | | Pnuc_1538, TRAP C4-dicarboxylate transport system permease DctM subunit |
| m4C | 1609032 | + | | upstream Pnuc_1544, transcriptional regulator, GntR family |
| m4C | 1611649 | - | | Pnuc_1542, ferredoxin |
| m4C | 1615808 | + | | Pnuc_1546, transcriptional regulator, LysR family |
| m4C | 1616536 | + | | Pnuc_1546, transcriptional regulator, LysR family |
| m6A | 1617388 | + | GAYANNNNNNNRTAG | Pnuc_1547, protein of unknown function DUF6, transmembrane |
| m6A | 1617398 | - | CTAYNNNNNNNTRTC | upstream Pnuc_1545, class II aldolase/adducin family protein |
| m4C | 1619064 | - | | Pnuc_1549, amino acid/amide ABC transporter substrate-binding protein, HAAT family |
| m4C | 1619386 | + | | upstream Pnuc_1553, transcriptional regulator, AsnC family |
| m6A | 1622660 | + | | upstream Pnuc_1553, transcriptional regulator, AsnC family |
| m4C | 1625078 | + | | upstream Pnuc_1553, transcriptional regulator, AsnC family |
| m4C | 1627175 | + | | upstream Pnuc_1560, transcriptional regulator, LysR family |
| m6A | 1628517 | + | GAYANNNNNNNRTAG | upstream Pnuc_1560, transcriptional regulator, LysR family |
| m6A | 1628527 | - | CTAYNNNNNNNTRTC | Pnuc_1556, Enoyl-CoA hydratase/isomerase |

| | | | | |
|---|---|---|---|---|
| m4C | 1630294 | + | | upstream Pnuc_1560, transcriptional regulator, LysR family |
| m4C | 1635495 | - | | upstream Pnuc_1561, major facilitator superfamily MFS_1 |
| m6A | 1638119 | + | CTAYNNNNNNNNTRTC | Pnuc_1564, FMN-binding domain protein |
| m6A | 1638129 | - | GAYANNNNNNNNRTAG | upstream Pnuc_1563, conserved hypothetical protein |
| m4C | 1645105 | + | | upstream Pnuc_1581, hypothetical protein |
| m4C | 1646478 | + | | upstream Pnuc_1581, hypothetical protein |
| m4C | 1647150 | - | | Pnuc_1574, Acetamidase/Formamidase |
| m4C | 1648753 | - | | Pnuc_1576, penicillin amidase, Cysteine peptidase, MEROPS family C59 |
| m4C | 1649413 | + | | upstream Pnuc_1581, hypothetical protein |
| m4C | 1653477 | - | | upstream Pnuc_1580, transcriptional regulator, ArsR family |
| m6A | 1654685 | + | | upstream Pnuc_1585, metallophosphoesterase |
| m6A | 1656328 | + | | Pnuc_1586, glycosyl transferase, group 1 |
| m6A | 1658012 | + | GAYANNNNNNNNRTAG | Pnuc_1588, ornithine-acyl[acyl carrier protein]N-acyltransferase |
| m6A | 1658022 | - | CTAYNNNNNNNNTRTC | upstream Pnuc_1584, lyso-ornithine lipid acyltransferase |
| m6A | 1660045 | + | | upstream Pnuc_1595, MgtC/SapB transporter |
| m4C | 1660384 | + | | upstream Pnuc_1595, MgtC/SapB transporter |
| m6A | 1660810 | + | CTAYNNNNNNNNTRTC | upstream Pnuc_1595, MgtC/SapB transporter |
| m6A | 1660820 | - | GAYANNNNNNNNRTAG | Pnuc_1591, MgtC/SapB transporter |
| m4C | 1663965 | - | | Pnuc_1593, conserved hypothetical protein |
| m4C | 1665691 | + | | Pnuc_1595, MgtC/SapB transporter |
| m4C | 1667725 | + | | upstream Pnuc_1606, short-chain dehydrogenase/reductase SDR |
| m4C | 1672413 | + | | upstream Pnuc_1606, short-chain dehydrogenase/reductase SDR |
| m4C | 1673396 | + | | upstream Pnuc_1606, short-chain dehydrogenase/reductase SDR |
| m4C | 1675386 | - | | Pnuc_1607, conserved hypothetical protein |
| m4C | 1683718 | + | | upstream Pnuc_1622, hypothetical protein |
| m6A | 1685160 | - | | Pnuc_1616, malate-quinone oxidoreductase |
| m4C | 1685584 | - | | Pnuc_1617, conserved hypothetical protein |
| m4C | 1687143 | - | | Pnuc_1618, fumarase |
| m6A | 1690825 | - | | upstream Pnuc_1621, Mg2+ transporter protein, CorA family protein |
| m4C | 1692780 | - | | Pnuc_1624, Nitrilase/cyanide hydratase and apolipoprotein N-acyltransferase |
| m4C | 1696150 | + | | upstream Pnuc_1636, transcriptional regulator, GntR family |
| m4C | 1698796 | + | | upstream Pnuc_1636, transcriptional regulator, GntR family |
| m6A | 1699838 | + | CTAYNNNNNNNNTRTC | upstream Pnuc_1636, transcriptional regulator, GntR family |
| m4C | 1700632 | + | | upstream Pnuc_1636, transcriptional regulator, GntR family |
| m4C | 1701764 | + | | upstream Pnuc_1636, transcriptional regulator, GntR family |
| m6A | 1705954 | + | CTAYNNNNNNNNTRTC | Pnuc_1638, protein of unknown function DUF6, transmembrane |
| m6A | 1705964 | - | GAYANNNNNNNNRTAG | upstream Pnuc_1637, NAD-dependent epimerase/dehydratase |
| m6A | 1708596 | - | | upstream Pnuc_1641, conserved hypothetical protein |
| m6A | 1712078 | + | CTAYNNNNNNNNTRTC | upstream Pnuc_1649, hypothetical protein |
| m6A | 1712088 | - | GAYANNNNNNNNRTAG | Pnuc_1647, DNA polymerase III, epsilon subunit |
| m6A | 1712327 | - | | Pnuc_1647, DNA polymerase III, epsilon subunit |
| m6A | 1712464 | - | | Pnuc_1647, DNA polymerase III, epsilon subunit |
| m6A | 1712822 | + | CTAYNNNNNNNNTRTC | upstream Pnuc_1649, hypothetical protein |
| m6A | 1712832 | - | GAYANNNNNNNNRTAG | Pnuc_1647, DNA polymerase III, epsilon subunit |

| | | | | |
|---|---|---|---|---|
| m4C | 1713392 | - | | Pnuc_1647, DNA polymerase III, epsilon subunit |
| m4C | 1715805 | - | | upstream Pnuc_1648, transcriptional regulator, BadM/Rrf2 family |
| m6A | 1716381 | - | | upstream Pnuc_1648, transcriptional regulator, BadM/Rrf2 family |
| m4C | 1717994 | - | | upstream Pnuc_1653, membrane-associated protein in eicosanoid and glutathione metabolism (MAPEG) |
| m6A | 1719842 | - | | upstream Pnuc_1653, membrane-associated protein in eicosanoid and glutathione metabolism (MAPEG) |
| m4C | 1731126 | + | | upstream Pnuc_1678, putative transcriptional regulator, MerR family |
| m4C | 1734371 | + | | upstream Pnuc_1678, putative transcriptional regulator, MerR family |
| m6A | 1734414 | + | | upstream Pnuc_1678, putative transcriptional regulator, MerR family |
| m4C | 1734534 | - | | Pnuc_1669, K+-transporting ATPase, B subunit |
| m6A | 1736959 | + | GAYANNNNNNNRTAG | upstream Pnuc_1678, putative transcriptional regulator, MerR family |
| m6A | 1736969 | - | CTAYNNNNNNNTRTC | Pnuc_1670, potassium-transporting ATPase, A subunit |
| m6A | 1738403 | + | CTAYNNNNNNNTRTC | upstream Pnuc_1678, putative transcriptional regulator, MerR family |
| m6A | 1738413 | - | GAYANNNNNNNRTAG | Pnuc_1672, Heavy metal transport/detoxification protein |
| m4C | 1739306 | - | | Pnuc_1673, conserved hypothetical protein |
| m4C | 1740979 | + | | upstream Pnuc_1678, putative transcriptional regulator, MerR family |
| m4C | 1742637 | + | | upstream Pnuc_1678, putative transcriptional regulator, MerR family |
| m4C | 1744638 | - | | Pnuc_1679, conserved hypothetical protein |
| m4C | 1748401 | + | | upstream Pnuc_1688, hypothetical protein |
| m6A | 1748827 | + | GAYANNNNNNNRTAG | upstream Pnuc_1688, hypothetical protein |
| m6A | 1748837 | - | CTAYNNNNNNNTRTC | Pnuc_1681, efflux transporter, RND family, MFP subunit |
| m4C | 1754380 | - | | Pnuc_1689, phage integrase family protein |
| m4C | 1756634 | + | | Pnuc_1692, hypothetical protein |
| m6A | 1758014 | + | GAYANNNNNNNRTAG | upstream Pnuc_1702, O-sialoglycoprotein endopeptidase |
| m4C | 1758023 | - | | Pnuc_1694, phage integrase family protein |
| m4C | 1760544 | - | | Pnuc_1696, hypothetical protein |
| m4C | 1760689 | + | | upstream Pnuc_1702, O-sialoglycoprotein endopeptidase |
| m6A | 1764285 | - | | Pnuc_1698, RNA polymerase, sigma 70 subunit, RpoD |
| m4C | 1765765 | - | | Pnuc_1699, DNA primase |
| m4C | 1769633 | + | | upstream Pnuc_1707, Rieske (2Fe-2S) domain protein |
| m6A | 1770001 | - | | Pnuc_1703, GTP cyclohydrolase I |
| m4C | 1772603 | + | | upstream Pnuc_1707, Rieske (2Fe-2S) domain protein |
| m4C | 1773113 | + | | upstream Pnuc_1707, Rieske (2Fe-2S) domain protein |
| m4C | 1773158 | + | | upstream Pnuc_1707, Rieske (2Fe-2S) domain protein |
| m4C | 1773375 | + | | upstream Pnuc_1707, Rieske (2Fe-2S) domain protein |
| m6A | 1774091 | + | | Pnuc_1707, Rieske (2Fe-2S) domain protein |
| m4C | 1775687 | - | | Pnuc_1709, zinc/iron permease |
| m4C | 1778142 | + | | upstream Pnuc_1712, putative transmembrane protein |
| m4C | 1779168 | - | | Pnuc_1711, DNA polymerase I |
| m4C | 1782990 | - | | Pnuc_1715, ATP-dependent DNA helicase UvrD |
| m6A | 1785209 | - | | upstream Pnuc_1715, ATP-dependent DNA helicase UvrD |
| m4C | 1785614 | + | | Pnuc_1716, valyl-tRNA synthetase |
| m4C | 1788121 | - | | upstream Pnuc_1715, ATP-dependent DNA helicase UvrD |
| m4C | 1790241 | + | | upstream Pnuc_1720, protein of unknown function DUF1468 |
| m4C | 1792420 | - | | upstream Pnuc_1719, alanyl-tRNA synthetase |

| | | | | |
|---|---|---|---|---|
| m6A | 1792900 | - | | upstream Pnuc_1719, alanyl-tRNA synthetase |
| m4C | 1793918 | - | | upstream Pnuc_1719, alanyl-tRNA synthetase |
| m6A | 1797655 | + | | Pnuc_1725, acetylornithine aminotransferase apoenzyme |
| m6A | 1797750 | + | CTAYNNNNNNNNTRTC | Pnuc_1725, acetylornithine aminotransferase apoenzyme |
| m6A | 1797760 | - | GAYANNNNNNNNRTAG | upstream Pnuc_1723, NAD-dependent epimerase/dehydratase |
| m4C | 1797852 | + | | Pnuc_1725, acetylornithine aminotransferase apoenzyme |
| m4C | 1800152 | - | | Pnuc_1727, amino acid/amide ABC transporter ATP-binding protein 1, HAAT family |
| m6A | 1804734 | + | | upstream Pnuc_1733, DNA replication and repair protein RadC |
| m4C | 1805763 | - | | upstream Pnuc_1732, peptidylprolyl isomerase, FKBP-type |
| m6A | 1806482 | - | | Pnuc_1736, fatty acid desaturase |
| m6A | 1808144 | + | | upstream Pnuc_1739, FMN adenylyltransferase / riboflavin kinase |
| m4C | 1809413 | - | | Pnuc_1738, formyltetrahydrofolate-dependent phosphoribosylglycinamide formyltransferase |
| m4C | 1816551 | + | | upstream Pnuc_1750, cold-shock DNA-binding protein family |
| m6A | 1821620 | - | | Pnuc_1748, ATP-dependent Clp protease ATP-binding subunit ClpA |
| m4C | 1823682 | - | | Pnuc_1753, argininosuccinate synthase |
| m4C | 1824982 | - | | Pnuc_1754, ornithine carbamoyltransferase |
| m4C | 1827467 | - | | upstream Pnuc_1756, SSU ribosomal protein S20P |
| m6A | 1829138 | - | | upstream Pnuc_1756, SSU ribosomal protein S20P |
| m4C | 1832132 | - | | upstream Pnuc_1761, phosphoribosylformylglycinamidine cyclo-ligase |
| m6A | 1832665 | + | CTAYNNNNNNNNTRTC | Pnuc_1762, protein of unknown function UPF0118 |
| m6A | 1832675 | - | GAYANNNNNNNNRTAG | upstream Pnuc_1761, phosphoribosylformylglycinamidine cyclo-ligase |
| m6A | 1832846 | - | | upstream Pnuc_1761, phosphoribosylformylglycinamidine cyclo-ligase |
| m6A | 1835641 | + | | Pnuc_1765, poly(A) polymerase |
| m4C | 1835657 | - | | upstream Pnuc_1761, phosphoribosylformylglycinamidine cyclo-ligase |
| m6A | 1836536 | + | | Pnuc_1766, 2-amino-4-hydroxy-6-hydroxymethyldihydropteridine pyrophosphokinase |
| m4C | 1836846 | - | | upstream Pnuc_1761, phosphoribosylformylglycinamidine cyclo-ligase |
| m4C | 1837887 | - | | Pnuc_1768, chaperone protein DnaJ |
| m4C | 1838959 | + | | upstream Pnuc_1773, NAD(+) kinase |
| m4C | 1839179 | - | | Pnuc_1769, chaperone protein DnaK |
| m4C | 1840063 | + | | upstream Pnuc_1773, NAD(+) kinase |
| m4C | 1840629 | + | | upstream Pnuc_1773, NAD(+) kinase |
| m4C | 1842862 | - | | Pnuc_1772, heat-inducible transcription repressor HrcA |
| m6A | 1843285 | + | GAYANNNNNNNNRTAG | upstream Pnuc_1773, NAD(+) kinase |
| m6A | 1843295 | - | CTAYNNNNNNNNTRTC | Pnuc_1772, heat-inducible transcription repressor HrcA |
| m4C | 1846249 | + | | upstream Pnuc_1776, membrane protein-like protein |
| m4C | 1849483 | - | | upstream Pnuc_1775, (Glutamate-ammonia-ligase) adenylyltransferase |
| m6A | 1852400 | + | | upstream Pnuc_1776, membrane protein-like protein |
| m6A | 1853226 | + | CTAYNNNNNNNNTRTC | Pnuc_1777, Nitrilase/cyanide hydratase and apolipoprotein N-acyltransferase |
| m6A | 1853236 | - | GAYANNNNNNNNRTAG | upstream Pnuc_1775, (Glutamate-ammonia-ligase) adenylyltransferase |
| m4C | 1855577 | - | | upstream Pnuc_1775, (Glutamate-ammonia-ligase) adenylyltransferase |
| m6A | 1861824 | + | GAYANNNNNNNNRTAG | Pnuc_1784, alanine racemase domain protein |
| m6A | 1861834 | - | CTAYNNNNNNNNTRTC | upstream Pnuc_1780, ATP:cob(I)alamin adenosyltransferase |
| m4C | 1862042 | - | | upstream Pnuc_1780, ATP:cob(I)alamin adenosyltransferase |
| m6A | 1862177 | - | | upstream Pnuc_1780, ATP:cob(I)alamin adenosyltransferase |

| | | | | |
|---|---|---|---|---|
| m4C | 1864988 | + | | upstream Pnuc_1788, S-adenosylmethionine--tRNA-ribosyltransferase-isomerase |
| m6A | 1865372 | + | GAYANNNNNNNRTAG | upstream Pnuc_1788, S-adenosylmethionine--tRNA-ribosyltransferase-isomerase |
| m6A | 1865382 | - | CTAYNNNNNNNTRTC | Pnuc_1787, ATP-dependent DNA helicase RecG |
| m6A | 1865725 | + | | upstream Pnuc_1788, S-adenosylmethionine--tRNA-ribosyltransferase-isomerase |
| m6A | 1867542 | + | CTAYNNNNNNNTRTC | Pnuc_1789, tRNA-guanine transglycosylase |
| m6A | 1867552 | - | GAYANNNNNNNRTAG | upstream Pnuc_1787, ATP-dependent DNA helicase RecG |
| m4C | 1867847 | - | | upstream Pnuc_1787, ATP-dependent DNA helicase RecG |
| m4C | 1868106 | - | | upstream Pnuc_1787, ATP-dependent DNA helicase RecG |
| m6A | 1868132 | + | | upstream Pnuc_1790, protein translocase subunit yajC |
| m4C | 1874377 | - | | upstream Pnuc_1794, Adenylosuccinate lyase |
| m4C | 1874784 | - | | upstream Pnuc_1794, Adenylosuccinate lyase |
| m6A | 1874872 | + | GAYANNNNNNNRTAG | upstream Pnuc_1797, alanine dehydrogenase/PNT domain protein |
| m6A | 1874882 | - | CTAYNNNNNNNTRTC | Pnuc_1796, tRNA (5-methylaminomethyl-2-thiouridylate)-methyltransferase |
| m6A | 1880808 | + | GAYANNNNNNNRTAG | upstream Pnuc_1802, hypothetical protein |
| m6A | 1880818 | - | CTAYNNNNNNNTRTC | Pnuc_1801, glycosyl transferase, family 11 |
| m4C | 1881748 | + | | upstream Pnuc_1802, hypothetical protein |
| m6A | 1881762 | + | CTAYNNNNNNNTRTC | upstream Pnuc_1802, hypothetical protein |
| m6A | 1881772 | - | GAYANNNNNNNRTAG | upstream Pnuc_1801, glycosyl transferase, family 11 |
| m6A | 1883472 | + | | upstream Pnuc_1807, putative diguanylate phosphodiesterase |
| m4C | 1884110 | - | | Pnuc_1805, chaperonin GroEL |
| m4C | 1886632 | + | | Pnuc_1808, two component transcriptional regulator, LuxR family |
| m4C | 1887063 | - | | upstream Pnuc_1805, chaperonin GroEL |
| m4C | 1891141 | - | | upstream Pnuc_1806, chaperonin Cpn10 |
| m4C | 1893231 | + | | upstream Pnuc_1818, branched chain amino acid aminotransferase apoenzyme |
| m4C | 1894041 | + | | upstream Pnuc_1818, branched chain amino acid aminotransferase apoenzyme |
| m6A | 1894804 | + | GAYANNNNNNNRTAG | upstream Pnuc_1818, branched chain amino acid aminotransferase apoenzyme |
| m6A | 1894814 | - | CTAYNNNNNNNTRTC | Pnuc_1814, phosphoribosylaminoimidazole-succinocarboxamide synthase |
| m4C | 1895880 | + | | upstream Pnuc_1818, branched chain amino acid aminotransferase apoenzyme |
| m4C | 1895901 | + | | upstream Pnuc_1818, branched chain amino acid aminotransferase apoenzyme |
| m6A | 1896432 | + | | upstream Pnuc_1818, branched chain amino acid aminotransferase apoenzyme |
| m4C | 1898142 | + | | upstream Pnuc_1818, branched chain amino acid aminotransferase apoenzyme |
| m4C | 1898342 | - | | Pnuc_1817, phosphoglycerate kinase |
| m6A | 1899165 | + | CTAYNNNNNNNTRTC | Pnuc_1818, branched chain amino acid aminotransferase apoenzyme |
| m6A | 1899175 | - | GAYANNNNNNNRTAG | upstream Pnuc_1817, phosphoglycerate kinase |
| m4C | 1899601 | - | | upstream Pnuc_1817, phosphoglycerate kinase |
| m4C | 1902681 | + | | upstream Pnuc_1824, GTP cyclohydrolase subunit MoaC |
| m4C | 1903168 | + | | upstream Pnuc_1824, GTP cyclohydrolase subunit MoaC |
| m4C | 1906490 | - | | upstream Pnuc_1825, TPR repeat-containing protein |
| m4C | 1907280 | + | | upstream Pnuc_1835, hypothetical protein |
| m6A | 1908212 | - | | Pnuc_1827, TPR repeat-containing protein |
| m6A | 1910712 | + | | upstream Pnuc_1835, hypothetical protein |
| m4C | 1910853 | - | | upstream Pnuc_1829, Integral membrane protein TerC |
| m4C | 1911260 | + | | upstream Pnuc_1835, hypothetical protein |
| m4C | 1911642 | + | | upstream Pnuc_1835, hypothetical protein |

| | | | | |
|---|---|---|---|---|
| m4C | 1913021 | + | | upstream Pnuc_1835, hypothetical protein |
| m6A | 1915736 | + | | Pnuc_1835, hypothetical protein |
| m6A | 1916434 | + | GAYANNNNNNNRTAG | Pnuc_1835, hypothetical protein |
| m6A | 1916444 | - | CTAYNNNNNNNTRTC | upstream Pnuc_1834, conserved hypothetical protein |
| m4C | 1918023 | - | | upstream Pnuc_1834, conserved hypothetical protein |
| m6A | 1918050 | + | GAYANNNNNNNRTAG | Pnuc_1838, hypothetical protein |
| m6A | 1918060 | - | CTAYNNNNNNNTRTC | upstream Pnuc_1834, conserved hypothetical protein |
| m4C | 1919243 | - | | upstream Pnuc_1839, phage integrase family protein |
| m6A | 1921973 | + | GAYANNNNNNNRTAG | upstream Pnuc_1847, secreted protein |
| m6A | 1921983 | - | CTAYNNNNNNNTRTC | upstream Pnuc_1843, hypothetical protein |
| m6A | 1922021 | - | | upstream Pnuc_1843, hypothetical protein |
| m4C | 1924413 | + | | upstream Pnuc_1847, secreted protein |
| m6A | 1926071 | - | | Pnuc_1849, major facilitator superfamily MFS_1 |
| m4C | 1928493 | - | | upstream Pnuc_1850, protein of unknown function UPF0126 |
| m6A | 1930518 | - | | Pnuc_1856, OsmC family protein |
| m4C | 1931091 | - | | Pnuc_1857, Rhodanese domain protein |
| m6A | 1931420 | - | | upstream Pnuc_1857, Rhodanese domain protein |
| m6A | 1931430 | - | | upstream Pnuc_1857, Rhodanese domain protein |
| m6A | 1932597 | + | | Pnuc_1859, tRNA-i(6)A37 thiotransferase enzyme MiaB |
| m6A | 1934625 | - | | upstream Pnuc_1857, Rhodanese domain protein |
| m6A | 1935097 | + | GAYANNNNNNNRTAG | Pnuc_1861, CBS domain containing protein |
| m6A | 1935107 | - | CTAYNNNNNNNTRTC | upstream Pnuc_1857, Rhodanese domain protein |
| m4C | 1936507 | + | | Pnuc_1862, apolipoprotein N-acyltransferase |
| m4C | 1938110 | + | | Pnuc_1864, glycyl-tRNA synthetase beta chain |
| m6A | 1938699 | - | | upstream Pnuc_1857, Rhodanese domain protein |
| m4C | 1938967 | - | | upstream Pnuc_1857, Rhodanese domain protein |
| m6A | 1940032 | + | CTAYNNNNNNNTRTC | Pnuc_1865,D-alpha,beta-D-heptose 1,7-bisphosphate phosphatase |
| m6A | 1940042 | - | GAYANNNNNNNRTAG | upstream Pnuc_1857, Rhodanese domain protein |
| m4C | 1943052 | - | | Pnuc_1869, PpiC-type peptidyl-prolyl cis-trans isomerase |
| m4C | 1943876 | + | | upstream Pnuc_1871, aminoglycoside phosphotransferase |
| m6A | 1944178 | + | | upstream Pnuc_1871, aminoglycoside phosphotransferase |
| m6A | 1944517 | + | | upstream Pnuc_1871, aminoglycoside phosphotransferase |
| m4C | 1944681 | + | | upstream Pnuc_1871, aminoglycoside phosphotransferase |
| m4C | 1948306 | - | | upstream Pnuc_1870, Organic solvent tolerance protein |
| m4C | 1949187 | - | | upstream Pnuc_1870, Organic solvent tolerance protein |
| m6A | 1949603 | - | | upstream Pnuc_1870, Organic solvent tolerance protein |
| m4C | 1950025 | + | | Pnuc_1873, aminopeptidase P, Metallo peptidase, MEROPS family M24B |
| m4C | 1955548 | - | | upstream Pnuc_1870, Organic solvent tolerance protein |
| m4C | 1956019 | - | | upstream Pnuc_1870, Organic solvent tolerance protein |
| m6A | 1956076 | + | | Pnuc_1880, Holliday junction DNA helicase RuvB |
| m6A | 1961700 | + | | upstream Pnuc_1887, OsmC family protein |
| m4C | 1962595 | - | | upstream Pnuc_R0042, RNase P |
| m6A | 1962717 | + | | upstream Pnuc_1900, L-threonine ammonia-lyase |
| m6A | 1964679 | + | | upstream Pnuc_1900, L-threonine ammonia-lyase |

| | | | | |
|---|---|---|---|---|
| m6A | 1968003 | + | CTAYNNNNNNNNTRTC | upstream Pnuc_1900, L-threonine ammonia-lyase |
| m6A | 1968013 | - | GAYANNNNNNNNRTAG | Pnuc_1892, aspartyl-tRNA synthetase |
| m6A | 1968185 | + | GAYANNNNNNNNRTAG | upstream Pnuc_1900, L-threonine ammonia-lyase |
| m6A | 1968195 | - | CTAYNNNNNNNNTRTC | Pnuc_1892, aspartyl-tRNA synthetase |
| m6A | 1968822 | + | | upstream Pnuc_1900, L-threonine ammonia-lyase |
| m4C | 1970066 | + | | upstream Pnuc_1900, L-threonine ammonia-lyase |
| m4C | 1972116 | + | | upstream Pnuc_1900, L-threonine ammonia-lyase |
| m4C | 1973150 | - | | upstream Pnuc_1896, import inner membrane translocase, subunit Tim44 |
| m4C | 1973296 | + | | upstream Pnuc_1900, L-threonine ammonia-lyase |
| m6A | 1974979 | + | GAYANNNNNNNNRTAG | upstream Pnuc_1900, L-threonine ammonia-lyase |
| m6A | 1974989 | - | CTAYNNNNNNNNTRTC | Pnuc_1899, FAD linked oxidase domain protein |
| m4C | 1976243 | + | | upstream Pnuc_1900, L-threonine ammonia-lyase |
| m6A | 1978391 | + | | Pnuc_1900, L-threonine ammonia-lyase |
| m4C | 1980452 | - | | upstream Pnuc_1899, FAD linked oxidase domain protein |
| m4C | 1980584 | + | | Pnuc_1901, 5-nucleotidase |
| m4C | 1981472 | + | | Pnuc_1902, GTP cyclohydrolase I |
| m4C | 1983001 | - | | Pnuc_1904, major facilitator superfamily MFS_1 |
| m4C | 1984330 | + | | Pnuc_1905, Excinuclease ABC subunit A |
| m4C | 1984737 | - | | upstream Pnuc_1904, major facilitator superfamily MFS_1 |
| m4C | 1986308 | - | | upstream Pnuc_1904, major facilitator superfamily MFS_2 |
| m4C | 1986900 | + | | upstream Pnuc_1907, KpsF/GutQ family protein |
| m4C | 1987093 | - | | Pnuc_1906, Kef-type potassium/proton antiporter, CPA2 family |
| m4C | 1987298 | + | | upstream Pnuc_1907, KpsF/GutQ family protein |
| m6A | 1992074 | + | | upstream Pnuc_1912, SSU ribosomal protein S30P / sigma 54 modulation protein |
| m6A | 1995448 | - | | Pnuc_1916, A/G-specific adenine glycosylase |
| m6A | 1995828 | - | | Pnuc_1916, A/G-specific adenine glycosylase |
| m4C | 1999392 | + | | Pnuc_1920, ribose-phosphate pyrophosphokinase |
| m6A | 1999753 | + | GAYANNNNNNNNRTAG | Pnuc_1920, ribose-phosphate pyrophosphokinase |
| m6A | 1999763 | - | CTAYNNNNNNNNTRTC | upstream Pnuc_1917, DNA-(apurinic or apyrimidinic site) lyase |
| m4C | 1999854 | - | | upstream Pnuc_1917, DNA-(apurinic or apyrimidinic site) lyase |
| m4C | 2000666 | + | | Pnuc_1921, LSU ribosomal protein L25P |
| m4C | 2002827 | - | | Pnuc_1925, putative methyltransferase |
| m4C | 2003070 | + | | upstream Pnuc_1928, signal recognition particle-docking protein FtsY |
| m6A | 2004271 | + | | upstream Pnuc_1928, signal recognition particle-docking protein FtsY |
| m6A | 2004383 | + | | upstream Pnuc_1928, signal recognition particle-docking protein FtsY |
| m4C | 2004652 | + | | upstream Pnuc_1928, signal recognition particle-docking protein FtsY |
| m6A | 2005336 | - | | Pnuc_1927, peptidase M16 domain protein |
| m4C | 2007198 | + | | Pnuc_1929, RNA polymerase, sigma 32 subunit, RpoH |
| m6A | 2008414 | + | GAYANNNNNNNNRTAG | upstream Pnuc_1935, conserved hypothetical protein |
| m6A | 2008424 | - | CTAYNNNNNNNNTRTC | Pnuc_1931, protoheme IX farnesyltransferase |
| m4C | 2011071 | - | | Pnuc_1934, SURF1 family protein |
| m4C | 2013906 | + | | upstream Pnuc_1942, phosphoribosyltransferase |
| m6A | 2014371 | + | CTAYNNNNNNNNTRTC | upstream Pnuc_1942, phosphoribosyltransferase |
| m6A | 2014381 | - | GAYANNNNNNNNRTAG | Pnuc_1939, Cytochrome-c oxidase |

| | | | | |
|---|---|---|---|---|
| m6A | 2014641 | + | GAYANNNNNNNRTAG | upstream Pnuc_1942, phosphoribosyltransferase |
| m6A | 2014651 | - | CTAYNNNNNNNNTRTC | Pnuc_1939, Cytochrome-c oxidase |
| m6A | 2016148 | - | | Pnuc_1940, cytochrome c oxidase, subunit II |
| m4C | 2016872 | - | | upstream Pnuc_1940, cytochrome c oxidase, subunit II |
| m4C | 2017520 | + | | upstream Pnuc_1942, phosphoribosyltransferase |
| m6A | 2020628 | + | | upstream Pnuc_1948, phosphoglycerate mutase |
| m4C | 2022461 | + | | Pnuc_1949, carboxyl-terminal protease |
| m4C | 2025658 | + | | upstream Pnuc_1959, protein of unknown function DUF526 |
| m4C | 2027802 | + | | upstream Pnuc_1959, protein of unknown function DUF526 |
| m4C | 2029132 | - | | Pnuc_1956, ammonium transporter |
| m6A | 2030432 | - | | Pnuc_1956, ammonium transporter |
| m6A | 2033708 | - | | Pnuc_1961, lipoic acid synthetase |
| m4C | 2034782 | - | | Pnuc_1963, protein of unknown function DUF493 |
| m6A | 2035852 | + | | upstream Pnuc_1967, biotin--acetyl-CoA-carboxylase ligase |
| m4C | 2037041 | - | | upstream Pnuc_1966, VanZ family protein |
| m6A | 2040053 | - | | upstream Pnuc_1969, rfaE bifunctional protein |
| m4C | 2042759 | + | | Pnuc_1973, Cyclopropane-fatty-acyl-phospholipid synthase |
| m4C | 2045187 | + | | upstream Pnuc_1978, conserved hypothetical protein |
| m4C | 2052349 | + | | upstream Pnuc_1983, protein of unknown function DUF1289 |
| m4C | 2053009 | - | | Pnuc_1984, Alcohol dehydrogenase, zinc-binding domain protein |
| m6A | 2054201 | + | CTAYNNNNNNNNTRTC | Pnuc_1985, hypothetical protein |
| m6A | 2054211 | - | GAYANNNNNNNRTAG | upstream Pnuc_1984, Alcohol dehydrogenase, zinc-binding domain protein |
| m4C | 2054550 | - | | upstream Pnuc_1984, Alcohol dehydrogenase, zinc-binding domain protein |
| m6A | 2058112 | + | | upstream Pnuc_1996, protein of unknown function DUF185 |
| m4C | 2060692 | - | | Pnuc_1990, RND efflux system, outer membrane lipoprotein, NodT family |
| m4C | 2061208 | + | | upstream Pnuc_1996, protein of unknown function DUF185 |
| m4C | 2061233 | + | | upstream Pnuc_1996, protein of unknown function DUF185 |
| m6A | 2062540 | - | | Pnuc_1991, glutamine-fructose-6-phosphate transaminase |
| m4C | 2063429 | - | | Pnuc_1991, glutamine-fructose-6-phosphate transaminase |
| m4C | 2063555 | - | | Pnuc_1992, UDP-N-acetylglucosamine pyrophosphorylase / glucosamine-1-phosphate N-acetyltransferase |
| m6A | 2063777 | + | CTAYNNNNNNNNTRTC | upstream Pnuc_1996, protein of unknown function DUF185 |
| m6A | 2063787 | - | GAYANNNNNNNRTAG | Pnuc_1992, UDP-N-acetylglucosamine pyrophosphorylase / glucosamine-1-phosphate N-acetyltransferase |
| m6A | 2066312 | + | GAYANNNNNNNRTAG | upstream Pnuc_1996, protein of unknown function DUF185 |
| m6A | 2066322 | - | CTAYNNNNNNNNTRTC | Pnuc_1994, dihydroneopterin aldolase |
| m4C | 2068726 | - | | Pnuc_1997, Polynucleotide adenylyltransferase region |
| m6A | 2071863 | + | | upstream Pnuc_2000, 5-formyltetrahydrofolate cyclo-ligase |
| m4C | 2072459 | - | | upstream Pnuc_1999, Lytic transglycosylase, catalytic |
| m4C | 2072686 | + | | Pnuc_2000, 5-formyltetrahydrofolate cyclo-ligase |
| m4C | 2072991 | - | | upstream Pnuc_1999, Lytic transglycosylase, catalytic |
| m4C | 2073910 | + | | upstream Pnuc_2004, lipid A biosynthesis acyltransferase |
| m6A | 2073919 | + | | upstream Pnuc_2004, lipid A biosynthesis acyltransferase |
| m4C | 2075118 | - | | Pnuc_2002, adenosylhomocysteinase |
| m6A | 2078652 | - | | upstream Pnuc_2003, methionine adenosyltransferase |
| m4C | 2081507 | + | | upstream Pnuc_2021, rod shape-determining protein MreB |

| | | | | |
|---|---|---|---|---|
| m4C | 2083039 | + | | upstream Pnuc_2021, rod shape-determining protein MreB |
| m6A | 2083866 | + | | upstream Pnuc_2021, rod shape-determining protein MreB |
| m4C | 2085644 | - | | Pnuc_2013, N-acetylglutamate kinase |
| m4C | 2085776 | - | | Pnuc_2013, N-acetylglutamate kinase |
| m6A | 2085972 | + | GAYANNNNNNNRTAG | upstream Pnuc_2021, rod shape-determining protein MreB |
| m6A | 2085982 | - | CTAYNNNNNNNTRTC | Pnuc_2014, transcriptional regulator, TraR/DksA family |
| m4C | 2086807 | + | | upstream Pnuc_2021, rod shape-determining protein MreB |
| m4C | 2087642 | - | | upstream Pnuc_2015, cobalamin synthesis protein, P47K |
| m4C | 2088399 | - | | Pnuc_2016, tyrosine recombinase XerC subunit |
| m4C | 2089108 | - | | Pnuc_2017, protein of unknown function DUF484 |
| m6A | 2091308 | + | | upstream Pnuc_2021, rod shape-determining protein MreB |
| m6A | 2091399 | - | | Pnuc_2019, aspartyl/glutamyl-tRNA(Asn/Gln) amidotransferase subunit A |
| m4C | 2092208 | - | | Pnuc_2019, aspartyl/glutamyl-tRNA(Asn/Gln) amidotransferase subunit A |
| m4C | 2092865 | - | | upstream Pnuc_2020, aspartyl/glutamyl-tRNA(Asn/Gln) amidotransferase subunit C |
| m4C | 2093583 | - | | upstream Pnuc_2020, aspartyl/glutamyl-tRNA(Asn/Gln) amidotransferase subunit C |
| m4C | 2094788 | + | | Pnuc_2023, rod shape-determining protein MreD |
| m6A | 2098763 | - | | Pnuc_2027, conserved hypothetical protein |
| m4C | 2098991 | - | | Pnuc_2027, conserved hypothetical protein |
| m6A | 2102026 | + | CTAYNNNNNNNTRTC | Pnuc_2031, lipid A biosynthesis acyltransferase |
| m6A | 2102036 | - | GAYANNNNNNNRTAG | upstream Pnuc_2029, protein of unknown function DUF167 |
| m4C | 2104632 | - | | Pnuc_2034, sodium:dicarboxylate symporter |
| m4C | 2104656 | + | | upstream Pnuc_2035, ribosomal large subunit pseudouridine synthase F |
| m6A | 2106577 | - | | upstream Pnuc_2034, sodium:dicarboxylate symporter |
| m6A | 2107824 | + | | Pnuc_2036, bacterial peptide chain release factor 3(bRF-3) |
| m6A | 2115796 | - | | upstream Pnuc_2043, alpha/beta hydrolase fold protein |
| m6A | 2118474 | + | GAYANNNNNNNRTAG | upstream Pnuc_2056, protein of unknown function DUF897 |
| m6A | 2118484 | - | CTAYNNNNNNNTRTC | Pnuc_2049, hypothetical protein |
| m6A | 2124960 | - | | upstream Pnuc_2055, transcriptional regulator, LysR family |
| m4C | 2125512 | + | | Pnuc_2057, conserved hypothetical protein |
| m4C | 2126829 | + | | upstream Pnuc_2062, hypothetical protein |
| m6A | 2127286 | - | | Pnuc_2060, Patatin |
| m4C | 2128347 | + | | upstream Pnuc_2062, hypothetical protein |
| m6A | 2129286 | - | | upstream Pnuc_2060, Patatin |
| m6A | 2129770 | - | | upstream Pnuc_2060, Patatin |
| m6A | 2132918 | + | | upstream Pnuc_2065, hypothetical protein |
| m4C | 2132997 | + | | upstream Pnuc_2065, hypothetical protein |
| m4C | 2133900 | + | | Pnuc_2065, hypothetical protein |
| m6A | 2133944 | + | GAYANNNNNNNRTAG | Pnuc_2065, hypothetical protein |
| m6A | 2133954 | - | CTAYNNNNNNNTRTC | upstream Pnuc_2064, hypothetical protein |
| m6A | 2136475 | + | | upstream Pnuc_2071, conserved hypothetical protein |
| m4C | 2137501 | + | | upstream Pnuc_2071, conserved hypothetical protein |
| m4C | 2138678 | - | | upstream Pnuc_2070, conserved hypothetical protein |
| m4C | 2139332 | - | | Pnuc_2072, hypothetical protein |
| m6A | 2142268 | - | | Pnuc_2073, protein of unknown function DUF181 |

| | | | | |
|---|---|---|---|---|
| m6A | 2143348 | - | | Pnuc_2075, multi-sensor signal transduction histidine kinase |
| m6A | 2145506 | - | | Pnuc_2076, putative periplasmic ligand-binding sensor protein |
| m4C | 2145808 | + | | upstream Pnuc_2080, DNA protecting protein DprA |
| m4C | 2146159 | + | | upstream Pnuc_2080, DNA protecting protein DprA |
| m6A | 2146256 | + | CTAYNNNNNNNNTRTC | upstream Pnuc_2080, DNA protecting protein DprA |
| m6A | 2146266 | - | GAYANNNNNNNNRTAG | Pnuc_2077, sun protein |
| m4C | 2147414 | + | | upstream Pnuc_2080, DNA protecting protein DprA |
| m4C | 2149995 | + | | Pnuc_2081, DNA topoisomerase III |
| m4C | 2150376 | + | | Pnuc_2081, DNA topoisomerase III |
| m4C | 2151878 | + | | Pnuc_2081, DNA topoisomerase III |
| m4C | 2152508 | + | | upstream Pnuc_0001, chromosomal replication initiator protein DnaA |
| m4C | 2154297 | + | | upstream Pnuc_0001, chromosomal replication initiator protein DnaA |
| m4C | 2154427 | + | | upstream Pnuc_0001, chromosomal replication initiator protein DnaA |
| m4C | 2154602 | + | | upstream Pnuc_0001, chromosomal replication initiator protein DnaA |

| 26°C + UV | | | | | |
|---|---|---|---|---|---|
| **Methylation** | **Location** | **Strand** | **Motif** | **Gene / Intergenic region** | **Count** |
| m4C | 2036 | + | | Pnuc_0002, DNA polymerase III, beta subunit | 457(+) |
| m6A | 10210 | + | | Pnuc_0010, amino acid/amide ABC transporter membrane protein 2, HAAT family | 416(-) |
| m4C | 10516 | + | | Pnuc_0010, amino acid/amide ABC transporter membrane protein 2, HAAT family | **Total** = 873 (m6A + m4C) |
| m4C | 19024 | + | | upstream Pnuc_0019, putative ATP synthase protein I AtpI | |
| m6A | 22187 | - | | upstream Pnuc_0013, acyl-CoA dehydrogenase domain protein | 253(+) |
| m4C | 23579 | + | | Pnuc_0025, ATP synthase F1 subcomplex gamma subunit | 251(-) |
| m6A | 24264 | + | | Pnuc_0025, ATP synthase F1 subcomplex gamma subunit | **Total** = 504 (m6A) |
| m4C | 29191 | - | | upstream Pnuc_0013, acyl-CoA dehydrogenase domain protein | |
| m4C | 30971 | + | | upstream Pnuc_0033, cytochrome c, class I | 204(+) |
| m4C | 31989 | + | | upstream Pnuc_0033, cytochrome c, class I | 165(-) |
| m4C | 32567 | + | | Pnuc_0033, cytochrome c, class I | Total = 369 (m4C) |
| m4C | 35794 | + | | upstream Pnuc_R0002, 16S ribosomal RNA | |
| m4C | 37423 | - | | upstream Pnuc_0038, GCN5-related N-acetyltransferase | Methylated genes = 336 (16.09%) |
| m6A | 37807 | + | CTAYNNNNNNNNTRTC | Pnuc_R0002, 16S ribosomal RNA | |
| m6A | 37817 | - | GAYANNNNNNNNRTAG | upstream Pnuc_0038, GCN5-related N-acetyltransferase | |
| m4C | 37977 | - | | upstream Pnuc_0038, GCN5-related N-acetyltransferase | |
| m4C | 43404 | - | | upstream Pnuc_0038, GCN5-related N-acetyltransferase | |
| m4C | 46285 | + | | Pnuc_0044, LSU ribosomal protein L10P | |
| m4C | 47940 | - | | upstream Pnuc_0038, GCN5-related N-acetyltransferase | |
| m4C | 48187 | + | | Pnuc_0046, DNA-directed RNA polymerase subunit beta | |
| m4C | 54576 | + | | Pnuc_0047, DNA-directed RNA polymerase subunit beta | |
| m4C | 57483 | - | | upstream Pnuc_0038, GCN5-related N-acetyltransferase | |
| m6A | 58559 | - | | upstream Pnuc_0038, GCN5-related N-acetyltransferase | |
| m4C | 61205 | + | | Pnuc_0053, LSU ribosomal protein L3P | |
| m4C | 61904 | + | | Pnuc_0054, LSU ribosomal protein L4P | |

| | | | | |
|---|---|---|---|---|
| m6A | 69850 | - | | upstream Pnuc_0038, GCN5-related N-acetyltransferase |
| m4C | 70523 | - | | upstream Pnuc_0038, GCN5-related N-acetyltransferase |
| m4C | 70690 | + | | Pnuc_0073, protein translocase subunit secY/sec61 alpha |
| m4C | 72955 | - | | upstream Pnuc_0038, GCN5-related N-acetyltransferase |
| m4C | 74844 | + | | upstream Pnuc_0080, CutA1 divalent ion tolerance protein |
| m4C | 75359 | - | | upstream Pnuc_0038, GCN5-related N-acetyltransferase |
| m6A | 77105 | - | | upstream Pnuc_0038, GCN5-related N-acetyltransferase |
| m4C | 78311 | + | | upstream Pnuc_0085, Cytochrome c553-like protein |
| m4C | 80546 | - | | Pnuc_0086, diaminopimelate decarboxylase |
| m6A | 83003 | + | GAYANNNNNNNRTAG | upstream Pnuc_0092, penicillin-binding protein, 1A family |
| m6A | 83013 | - | CTAYNNNNNNNTRTC | Pnuc_0088, 3-dehydroquinate synthase |
| m6A | 84386 | + | CTAYNNNNNNNTRTC | upstream Pnuc_0092, penicillin-binding protein, 1A family |
| m6A | 84396 | - | GAYANNNNNNNRTAG | Pnuc_0090, type II and III secretion system protein |
| m6A | 85182 | - | | Pnuc_0091, hypothetical protein |
| m6A | 85246 | - | | Pnuc_0091, hypothetical protein |
| m4C | 91931 | + | | Pnuc_0094, glutamate synthase (NADH) large subunit |
| m4C | 101398 | + | | Pnuc_0105, UDP-N-acetylglucosamine 1-carboxyvinyltransferase |
| m6A | 101539 | + | | Pnuc_0105, UDP-N-acetylglucosamine 1-carboxyvinyltransferase |
| m6A | 102779 | + | GAYANNNNNNNRTAG | Pnuc_0107, histidinol dehydrogenase |
| m6A | 102789 | - | CTAYNNNNNNNTRTC | upstream Pnuc_0091, hypothetical protein |
| m6A | 103179 | + | CTAYNNNNNNNTRTC | Pnuc_0107, histidinol dehydrogenase |
| m6A | 103189 | - | GAYANNNNNNNRTAG | upstream Pnuc_0091, hypothetical protein |
| m4C | 117662 | - | | Pnuc_0122, Tetratricopeptide TPR_2 repeat protein |
| m6A | 122705 | + | CTAYNNNNNNNTRTC | Pnuc_0127, cytochrome c1 |
| m6A | 122715 | - | GAYANNNNNNNRTAG | upstream Pnuc_0124, large conductance mechanosensitive channel protein |
| m6A | 124671 | + | | Pnuc_0130, TPR domain protein |
| m4C | 126750 | - | | upstream Pnuc_0131, General substrate transporter |
| m6A | 130121 | + | GAYANNNNNNNRTAG | upstream Pnuc_0142, GTP-binding protein YchF |
| m6A | 130131 | - | CTAYNNNNNNNTRTC | Pnuc_0136, bacterial peptide chain release factor 1(bRF-1) |
| m4C | 130984 | + | | upstream Pnuc_0142, GTP-binding protein YchF |
| m4C | 136421 | - | | upstream Pnuc_0141, Ubiquinone biosynthesis hydroxylase, UbiH/UbiF/VisC/COQ6 family |
| m6A | 137802 | - | | Pnuc_0141, Ubiquinone biosynthesis hydroxylase, UbiH/UbiF/VisC/COQ6 family |
| m6A | 139555 | - | | Pnuc_0145, indole-3-glycerol phosphate synthase |
| m4C | 139957 | + | | upstream Pnuc_0150, ApaG domain protein |
| m6A | 141313 | - | | Pnuc_0147, anthranilate synthase, component II |
| m4C | 142593 | + | | upstream Pnuc_0150, ApaG domain protein |
| m4C | 143233 | - | | Pnuc_0149, ribulose-5-phosphate 3-epimerase |
| m4C | 143384 | + | | upstream Pnuc_0150, ApaG domain protein |
| m4C | 144036 | + | | upstream Pnuc_0151, MltA domain protein |
| m4C | 144075 | - | | upstream Pnuc_0149, ribulose-5-phosphate 3-epimerase |
| m4C | 145779 | + | | upstream Pnuc_0156, biotin synthase |
| m6A | 149067 | + | CTAYNNNNNNNTRTC | upstream Pnuc_0156, biotin synthase |
| m6A | 149077 | - | GAYANNNNNNNRTAG | Pnuc_0155, aminotransferase |
| m4C | 151293 | + | | Pnuc_0157, putative ubiquinone biosynthesis protein |

| | | | | |
|---|---|---|---|---|
| m6A | 157993 | + | GAYANNNNNNNRTAG | Pnuc_0163, UDP-N-acetylmuramoylalanyl-D-glutamyl-2,6-diaminopimelate--D-alanyl-D-alanyl ligase |
| m6A | 158003 | - | CTAYNNNNNNNTRTC | upstream Pnuc_0155, aminotransferase |
| m4C | 165880 | - | | upstream Pnuc_0155, aminotransferase |
| m4C | 167854 | - | | upstream Pnuc_0155, aminotransferase |
| m4C | 171063 | - | | Pnuc_0175, hypothetical protein |
| m4C | 178192 | - | | upstream Pnuc_0175, hypothetical protein |
| m6A | 182434 | - | | Pnuc_0185, type II secretion system protein |
| m4C | 186994 | - | | upstream Pnuc_0189, farnesyl-diphosphate farnesyltransferase |
| m4C | 190804 | - | | Pnuc_0193, Farnesyltranstransferase |
| m4C | 191543 | + | | Pnuc_0195, LSU ribosomal protein L27P |
| m6A | 199904 | + | | upstream Pnuc_0204, ribonucleoside-diphosphate reductase, alpha subunit |
| m6A | 199926 | + | | upstream Pnuc_0204, ribonucleoside-diphosphate reductase, alpha subunit |
| m6A | 200762 | + | CTAYNNNNNNNTRTC | Pnuc_0204, ribonucleoside-diphosphate reductase, alpha subunit |
| m6A | 200772 | - | GAYANNNNNNNRTAG | upstream Pnuc_0201, signal recognition particle subunit FFH/SRP54 (srp54) |
| m4C | 200904 | + | | Pnuc_0204, ribonucleoside-diphosphate reductase, alpha subunit |
| m4C | 205747 | + | | upstream Pnuc_0213, UDP-N-acetylmuramate |
| m4C | 206369 | + | | upstream Pnuc_0213, UDP-N-acetylmuramate |
| m4C | 206994 | + | | upstream Pnuc_0213, UDP-N-acetylmuramate |
| m4C | 208221 | - | | Pnuc_0209, acetyl-CoA carboxylase, biotin carboxylase |
| m4C | 208743 | - | | Pnuc_0209, acetyl-CoA carboxylase, biotin carboxylase |
| m4C | 209491 | - | | Pnuc_0210, biotin carboxyl carrier protein |
| m6A | 209668 | + | GAYANNNNNNNRTAG | upstream Pnuc_0213, UDP-N-acetylmuramate |
| m6A | 209678 | - | CTAYNNNNNNNTRTC | Pnuc_0211, 3-dehydroquinate dehydratase |
| m4C | 211180 | + | | Pnuc_0213, UDP-N-acetylmuramate |
| m4C | 214042 | + | | Pnuc_0215, ribonuclease II |
| m4C | 218189 | - | | Pnuc_0219, orotidine 5'-phosphate decarboxylase |
| m4C | 219189 | + | | Pnuc_0220, magnesium and cobalt transport protein CorA |
| m4C | 221772 | - | | upstream Pnuc_0223, thiamine-phosphate kinase |
| m6A | 227673 | + | GAYANNNNNNNRTAG | Pnuc_0228, glyceraldehyde-3-phosphate dehydrogenase |
| m6A | 227683 | - | CTAYNNNNNNNTRTC | upstream Pnuc_0226, protein of unknown function DUF558 |
| m6A | 228733 | - | | Pnuc_0229, ferric uptake regulator, Fur family |
| m4C | 229043 | - | | Pnuc_0229, ferric uptake regulator, Fur family |
| m4C | 229114 | + | | upstream Pnuc_0230, SmpA/OmlA domain protein |
| m4C | 229124 | + | | upstream Pnuc_0230, SmpA/OmlA domain protein |
| m6A | 229942 | - | | upstream Pnuc_0229, ferric uptake regulator, Fur family |
| m4C | 237850 | + | | Pnuc_0237, putative RNA methylase |
| m4C | 245244 | + | | Pnuc_0246, aspartate carbamoyltransferase |
| m4C | 247712 | - | | upstream Pnuc_0241, deoxyribodipyrimidine photo-lyase type I |
| m6A | 248429 | + | | Pnuc_0249, glycosyl transferase, family 25 |
| m6A | 248439 | - | GAYANNNNNNNRTAG | upstream Pnuc_0241, deoxyribodipyrimidine photo-lyase type I |
| m4C | 249513 | - | | Pnuc_0250, hypothetical protein |
| m6A | 249670 | + | | upstream Pnuc_0252, hypothetical protein |
| m4C | 250708 | - | | Pnuc_0251, hypothetical protein |
| m6A | 250899 | + | CTAYNNNNNNNTRTC | Pnuc_0252, hypothetical protein |

| | | | | |
|---|---|---|---|---|
| m6A | 250909 | - | GAYANNNNNNNRTAG | upstream Pnuc_0251, hypothetical protein |
| m6A | 251666 | - | CTAYNNNNNNNTRTC | upstream Pnuc_0251, hypothetical protein |
| m4C | 257150 | + | | upstream Pnuc_0259, Undecaprenyl-diphosphatase |
| m6A | 269370 | + | GAYANNNNNNNRTAG | upstream Pnuc_0279, serine hydroxymethyltransferase |
| m6A | 269380 | - | CTAYNNNNNNNTRTC | Pnuc_0272 (start codon), preQ(0) biosynthesis protein QueC |
| m6A | 271013 | + | | upstream Pnuc_0279, serine hydroxymethyltransferase |
| m6A | 271023 | - | CTAYNNNNNNNTRTC | Pnuc_0275, TolB, N-terminal domain protein |
| m6A | 277009 | + | CTAYNNNNNNNTRTC | upstream Pnuc_0287, Exodeoxyribonuclease VII large subunit |
| m6A | 277019 | - | GAYANNNNNNNRTAG | Pnuc_0282, 3-deoxy-D-manno-octulosonate cytidylyltransferase |
| m6A | 279077 | - | | Pnuc_0285, Biopolymer transport protein ExbD/TolR |
| m6A | 280100 | + | | upstream Pnuc_0287, Exodeoxyribonuclease VII large subunit |
| m6A | 283189 | + | GAYANNNNNNNRTAG | upstream Pnuc_0290, protein of unknown function DUF520 |
| m6A | 283199 | - | CTAYNNNNNNNTRTC | Pnuc_0289, UDP-N-acetylmuramate dehydrogenase |
| m4C | 283631 | + | | upstream Pnuc_0290, protein of unknown function DUF520 |
| m4C | 304396 | - | | upstream Pnuc_0310, sugar transferase |
| m6A | 305983 | - | | upstream Pnuc_0310, sugar transferase |
| m4C | 309403 | + | | Pnuc_0317, glycosyl transferase, family 2 |
| m6A | 310366 | + | CTAYNNNNNNNTRTC | Pnuc_0318, ABC-2 type transporter |
| m6A | 310376 | - | GAYANNNNNNNRTAG | upstream Pnuc_0316, glycosyl transferase, family 4 |
| m6A | 312933 | + | GAYANNNNNNNRTAG | Pnuc_0320, Methyltransferase type 11 |
| m6A | 312943 | - | CTAYNNNNNNNTRTC | upstream Pnuc_0316, glycosyl transferase, family 4 |
| m6A | 313185 | - | CTAYNNNNNNNTRTC | upstream Pnuc_0316, glycosyl transferase, family 4 |
| m6A | 318999 | + | | upstream Pnuc_0324, NAD-dependent epimerase/dehydratase |
| m6A | 319009 | - | GAYANNNNNNNRTAG | upstream Pnuc_0316, glycosyl transferase, family 4 |
| m4C | 320163 | + | | Pnuc_0325, amine oxidase |
| m6A | 320925 | + | GAYANNNNNNNRTAG | Pnuc_0325, amine oxidase |
| m6A | 320935 | - | CTAYNNNNNNNTRTC | upstream Pnuc_0316, glycosyl transferase, family 4 |
| m6A | 321154 | + | CTAYNNNNNNNTRTC | Pnuc_0325, amine oxidase |
| m6A | 321164 | - | GAYANNNNNNNRTAG | upstream Pnuc_0316, glycosyl transferase, family 4 |
| m4C | 322370 | - | | Pnuc_0327, short-chain dehydrogenase/reductase SDR |
| m6A | 324292 | + | GAYANNNNNNNRTAG | Pnuc_0329, UbiA prenyltransferase |
| m6A | 324302 | - | CTAYNNNNNNNTRTC | upstream Pnuc_0328, FAD linked oxidase domain protein |
| m6A | 335858 | + | CTAYNNNNNNNTRTC | Pnuc_0338, hypothetical protein |
| m6A | 335868 | - | GAYANNNNNNNRTAG | upstream Pnuc_0336, hypothetical protein |
| m6A | 345459 | + | GAYANNNNNNNRTAG | upstream Pnuc_0351, acyltransferase 3 |
| m6A | 345469 | - | CTAYNNNNNNNTRTC | upstream Pnuc_0349, hypothetical protein |
| m6A | 345472 | - | | upstream Pnuc_0349, hypothetical protein |
| m6A | 347772 | + | GAYANNNNNNNRTAG | Pnuc_0353, Choline dehydrogenase |
| m6A | 347782 | - | CTAYNNNNNNNTRTC | upstream Pnuc_0352, protein of unknown function DUF615 |
| m4C | 348956 | + | | Pnuc_0354, protein of unknown function DUF6, transmembrane |
| m6A | 349791 | - | | upstream Pnuc_0352, protein of unknown function DUF615 |
| m4C | 353886 | + | | Pnuc_0357, amino acid/amide ABC transporter ATP-binding protein 1, HAAT family |
| m4C | 355021 | - | | Pnuc_0359, hypothetical protein |
| m4C | 355781 | + | | upstream Pnuc_0362, conserved hypothetical protein |

| | | | | |
|---|---|---|---|---|
| m4C | 367208 | + | | Pnuc_0371, transcriptional regulator, TetR family |
| m4C | 375033 | + | | upstream Pnuc_0384, D-isomer specific 2-hydroxyacid dehydrogenase, NAD-binding protein |
| m4C | 378715 | - | | Pnuc_0387, molybdenum cofactor guanylyltransferase |
| m4C | 378809 | - | | Pnuc_0387, molybdenum cofactor guanylyltransferase |
| m4C | 380268 | + | | upstream Pnuc_0390, ribosomal large subunit pseudouridine synthase C |
| m4C | 382926 | + | | upstream Pnuc_0390, ribosomal large subunit pseudouridine synthase C |
| m4C | 387629 | + | | Pnuc_0395, protein of unknown function DUF177 |
| m4C | 387941 | + | | Pnuc_0395, protein of unknown function DUF177 |
| m4C | 395248 | - | | upstream Pnuc_0394, maf protein |
| m6A | 397041 | + | CTAYNNNNNNNTRTC | Pnuc_0404, GTP-binding protein LepA |
| m6A | 401086 | + | CTAYNNNNNNNTRTC | Pnuc_0409, pyridoxine 5'-phosphate synthase |
| m6A | 401096 | - | GAYANNNNNNNRTAG | upstream Pnuc_0394, maf protein |
| m6A | 401232 | + | GAYANNNNNNNRTAG | Pnuc_0410, holo-acyl-carrier-protein synthase |
| m6A | 401242 | - | CTAYNNNNNNNTRTC | upstream Pnuc_0394, maf protein |
| m6A | 403790 | - | | Pnuc_0413, translation elongation factor P (EF-P) |
| m6A | 403974 | - | | Pnuc_0413, translation elongation factor P (EF-P) |
| m4C | 404185 | + | | upstream Pnuc_0415, Excinuclease ABC subunit C |
| m4C | 404201 | - | | upstream Pnuc_0413, translation elongation factor P (EF-P) |
| m4C | 405407 | + | | Pnuc_0415, Excinuclease ABC subunit C |
| m4C | 406883 | - | | upstream Pnuc_0414, conserved hypothetical protein |
| m4C | 412334 | - | | upstream Pnuc_0418, SOS-response transcriptional repressor, LexA |
| m4C | 413567 | - | | upstream Pnuc_0418, SOS-response transcriptional repressor, LexA |
| m6A | 421455 | + | GAYANNNNNNNRTAG | upstream Pnuc_0437, aminotransferase |
| m6A | 421465 | - | CTAYNNNNNNNTRTC | Pnuc_0432, formyl transferase domain protein |
| m4C | 421490 | - | | Pnuc_0432, formyl transferase domain protein |
| m6A | 422836 | + | CTAYNNNNNNNTRTC | upstream Pnuc_0437, aminotransferase |
| m6A | 422846 | - | GAYANNNNNNNRTAG | Pnuc_0434, DegT/DnrJ/EryC1/StrS aminotransferase |
| m4C | 424370 | + | | upstream Pnuc_0437, aminotransferase |
| m4C | 427493 | - | | upstream Pnuc_0436, protein of unknown function DUF498 |
| m6A | 439040 | + | GAYANNNNNNNRTAG | Pnuc_0447, ATPase AAA-2 domain protein |
| m6A | 439050 | - | CTAYNNNNNNNTRTC | upstream Pnuc_0446, Excinuclease ABC, C subunit domain protein |
| m6A | 439614 | - | | upstream Pnuc_0446, Excinuclease ABC, C subunit domain protein |
| m6A | 440012 | + | GAYANNNNNNNRTAG | Pnuc_0447, ATPase AAA-2 domain protein |
| m6A | 442569 | + | GAYANNNNNNNRTAG | Pnuc_0450, major facilitator superfamily MFS_1 |
| m6A | 442579 | - | CTAYNNNNNNNTRTC | upstream Pnuc_0449, Uncharacterized protein UPF0065 |
| m6A | 445942 | + | CTAYNNNNNNNTRTC | upstream Pnuc_0453, cytochrome c oxidase, cbb3-type, subunit I |
| m6A | 445952 | - | GAYANNNNNNNRTAG | upstream Pnuc_0449, Uncharacterized protein UPF0065 |
| m4C | 451434 | + | | upstream Pnuc_0461, putative transmembrane protein |
| m4C | 452040 | + | | Pnuc_0461, putative transmembrane protein |
| m6A | 453512 | + | CTAYNNNNNNNTRTC | Pnuc_0463, UspA domain protein |
| m6A | 453522 | - | GAYANNNNNNNRTAG | upstream Pnuc_0462, conserved hypothetical protein |
| m4C | 458223 | + | | upstream Pnuc_0471, 3-hydroxyacyl-CoA dehydrogenase |
| m6A | 461835 | + | GAYANNNNNNNRTAG | upstream Pnuc_0471, 3-hydroxyacyl-CoA dehydrogenase |
| m4C | 463015 | - | | upstream Pnuc_0470, D-isomer specific 2-hydroxyacid dehydrogenase, NAD-binding protein |

| | | | | |
|---|---|---|---|---|
| m4C | 471398 | + | | upstream Pnuc_0485, Uncharacterized protein UPF0065 |
| m4C | 471940 | + | | upstream Pnuc_0485, Uncharacterized protein UPF0065 |
| m4C | 472091 | - | | Pnuc_0482, MmgE/PrpD family protein |
| m6A | 474614 | + | CTAYNNNNNNNNTRTC | upstream Pnuc_0485, Uncharacterized protein UPF0065 |
| m6A | 474624 | - | GAYANNNNNNNNRTAG | upstream Pnuc_0484, Rhodanese domain protein |
| m4C | 477378 | + | | Pnuc_0488, methyltransferase small |
| m6A | 477475 | - | | upstream Pnuc_0484, Rhodanese domain protein |
| m4C | 479258 | - | | upstream Pnuc_0484, Rhodanese domain protein |
| m4C | 480232 | - | | upstream Pnuc_0484, Rhodanese domain protein |
| m4C | 480774 | - | | Pnuc_0490, phosphoglycolate phosphatase |
| m4C | 486646 | + | | Pnuc_0494, phosphoserine aminotransferase apoenzyme |
| m6A | 499723 | + | | Pnuc_0507, helix-hairpin-helix motif protein |
| m4C | 501868 | + | | upstream Pnuc_0510, histone deacetylase superfamily |
| m6A | 511656 | + | GAYANNNNNNNNRTAG | Pnuc_0520, LSU ribosomal protein L19P |
| m6A | 511666 | - | CTAYNNNNNNNNTRTC | upstream Pnuc_0516, protein of unknown function DUF306, Meta and HslJ |
| m4C | 511707 | - | | upstream Pnuc_0516, protein of unknown function DUF306, Meta and HslJ |
| m6A | 513038 | - | | upstream Pnuc_0516, protein of unknown function DUF306, Meta and HslJ |
| m6A | 513793 | + | CTAYNNNNNNNNTRTC | upstream Pnuc_0525, Exonuclease, RNase T and DNA polymerase III |
| m6A | 513803 | - | GAYANNNNNNNNRTAG | Pnuc_0523, ribosome small subunit-dependent GTPase A |
| m6A | 516835 | + | GAYANNNNNNNNRTAG | upstream Pnuc_0527, microcin-processing peptidase 1, Unknown type peptidase, MEROPS family U62 |
| m6A | 516845 | - | CTAYNNNNNNNNTRTC | Pnuc_0526, molybdopterin adenylyltransferase |
| m4C | 528854 | + | | upstream Pnuc_0541, Glutathione S-transferase, N-terminal domain protein |
| m6A | 529410 | + | GAYANNNNNNNNRTAG | upstream Pnuc_0541, Glutathione S-transferase, N-terminal domain protein |
| m6A | 529420 | - | CTAYNNNNNNNNTRTC | upstream Pnuc_0538, hypothetical protein |
| m4C | 532214 | + | | upstream Pnuc_0541, Glutathione S-transferase, N-terminal domain protein |
| m6A | 552556 | + | | upstream Pnuc_0565, hypothetical protein |
| m6A | 553023 | + | CTAYNNNNNNNNTRTC | upstream Pnuc_0565, hypothetical protein |
| m6A | 553033 | - | GAYANNNNNNNNRTAG | Pnuc_0563, integral membrane sensor hybrid histidine kinase |
| m6A | 556728 | - | | Pnuc_0568, conserved hypothetical protein |
| m4C | 556883 | - | | Pnuc_0568, conserved hypothetical protein |
| m6A | 557691 | + | | upstream Pnuc_0570, cation diffusion facilitator family transporter |
| m6A | 560162 | + | GAYANNNNNNNNRTAG | Pnuc_0570, cation diffusion facilitator family transporter |
| m6A | 560172 | - | CTAYNNNNNNNNTRTC | upstream Pnuc_0569, GCN5-related N-acetyltransferase |
| m4C | 562505 | + | | Pnuc_0573, multiple antibiotic resistance (MarC)-related protein |
| m6A | 563811 | - | | upstream Pnuc_0571, beta-lactamase domain protein |
| m6A | 564182 | - | | upstream Pnuc_0571, beta-lactamase domain protein |
| m4C | 567773 | + | | upstream Pnuc_0580, hypothetical protein |
| m4C | 574015 | - | | upstream Pnuc_0582, hypothetical protein |
| m6A | 575931 | + | | upstream Pnuc_0594, 2-hydroxy-3-oxopropionate reductase |
| m6A | 581452 | - | | Pnuc_0595, N-acetylmuramoyl-L-alanine amidase |
| m4C | 585405 | - | | Pnuc_0600, FAD linked oxidase domain protein |
| m6A | 587229 | + | | upstream Pnuc_0602, Serine--glyoxylate transaminase |
| m6A | 587239 | - | CTAYNNNNNNNNTRTC | Pnuc_0600, FAD linked oxidase domain protein |
| m4C | 588024 | + | | upstream Pnuc_0602, Serine--glyoxylate transaminase |

| | | | | |
|---|---|---|---|---|
| m6A | 590041 | - | | upstream Pnuc_0601, transcriptional regulator, GntR family |
| m6A | 598932 | + | GAYANNNNNNNRTAG | upstream Pnuc_0610, lipid A ABC exporter, fused ATPase and inner membrane subunits MsbA |
| m4C | 600957 | - | | upstream Pnuc_0609, glycosyl transferase, family 2 |
| m4C | 601332 | - | | Pnuc_0611, RNAse G |
| m6A | 610145 | + | | upstream Pnuc_0623, aminopeptidase A, Metallo peptidase, MEROPS family M17 |
| m6A | 618648 | + | GAYANNNNNNNRTAG | Pnuc_0628, TRAP dicarboxylate transporter- DctP subunit |
| m6A | 618658 | - | CTAYNNNNNNNTRTC | upstream Pnuc_0622, permease YjgP/YjgQ family protein |
| m6A | 621941 | + | GAYANNNNNNNRTAG | Pnuc_0631, AMP-dependent synthetase and ligase |
| m6A | 621951 | - | CTAYNNNNNNNTRTC | upstream Pnuc_0622, permease YjgP/YjgQ family protein |
| m4C | 624499 | + | | upstream Pnuc_0636, cyclic nucleotide-binding protein |
| m6A | 630181 | + | | Pnuc_0638, L-carnitine dehydratase/bile acid-inducible protein F |
| m4C | 630740 | + | | Pnuc_0639, putative PAS/PAC sensor protein |
| m6A | 632692 | + | | Pnuc_0641, Methionyl-tRNA formyltransferase |
| m4C | 637747 | - | | Pnuc_0647, methionyl-tRNA synthetase |
| m4C | 638606 | - | | Pnuc_0647, methionyl-tRNA synthetase |
| m6A | 639054 | + | GAYANNNNNNNRTAG | Pnuc_0648, formate dehydrogenase, subunit FdhD |
| m6A | 639064 | - | CTAYNNNNNNNTRTC | upstream Pnuc_0647, methionyl-tRNA synthetase |
| m6A | 639706 | + | | Pnuc_0648, formate dehydrogenase, subunit FdhD |
| m6A | 641919 | - | | upstream Pnuc_0647, methionyl-tRNA synthetase |
| m6A | 642290 | + | CTAYNNNNNNNTRTC | Pnuc_0651, Lysine decarboxylase |
| m6A | 642300 | - | GAYANNNNNNNRTAG | upstream Pnuc_0647, methionyl-tRNA synthetase |
| m4C | 646068 | + | | upstream Pnuc_0656, porphobilinogen deaminase |
| m4C | 648337 | + | | Pnuc_0656, porphobilinogen deaminase |
| m4C | 656646 | - | | upstream Pnuc_0663, Pyridoxamine 5'-phosphate oxidase |
| m6A | 664401 | + | CTAYNNNNNNNTRTC | upstream Pnuc_0672, ABC transporter related protein |
| m6A | 664411 | - | | Pnuc_0670, cyanophycin synthetase |
| m4C | 665984 | - | | Pnuc_0671, cyanophycin synthetase |
| m6A | 685642 | + | CTAYNNNNNNNTRTC | Pnuc_0686, V-type H(+)-translocating pyrophosphatase |
| m6A | 685652 | - | GAYANNNNNNNRTAG | upstream Pnuc_0685, Inorganic diphosphatase |
| m4C | 696773 | + | | Pnuc_0696, amidohydrolase |
| m4C | 702310 | - | | upstream Pnuc_R0019, SRP RNA; RNA component of signal recognition particle |
| m6A | 706399 | + | | Pnuc_0705, DNA-directed DNA polymerase |
| m6A | 709971 | + | | Pnuc_0711, AsmA family protein |
| m4C | 714147 | - | | upstream Pnuc_0716, YaeQ family protein |
| m6A | 716815 | + | CTAYNNNNNNNTRTC | Pnuc_0718, coproporphyrinogen III oxidase, anaerobic |
| m6A | 716825 | - | GAYANNNNNNNRTAG | upstream Pnuc_0716, YaeQ family protein |
| m6A | 717198 | + | CTAYNNNNNNNTRTC | upstream Pnuc_0719, OmpW family protein |
| m6A | 717208 | - | GAYANNNNNNNRTAG | upstream Pnuc_0716, YaeQ family protein |
| m4C | 718382 | + | | upstream Pnuc_0721, ABC transporter related protein |
| m4C | 723601 | - | | upstream Pnuc_0724, Altronate dehydratase |
| m6A | 724976 | - | | upstream Pnuc_0725, Pirin domain protein |
| m6A | 726362 | - | | upstream Pnuc_0727, phosphate-starvation-inducible E |
| m4C | 727752 | + | | upstream Pnuc_0734, 2-oxo-acid dehydrogenase E1 subunit, homodimeric type |
| m6A | 730423 | + | CTAYNNNNNNNTRTC | upstream Pnuc_0734, 2-oxo-acid dehydrogenase E1 subunit, homodimeric type |

| | | | | |
|---|---|---|---|---|
| m6A | 730433 | - | GAYANNNNNNNRTAG | Pnuc_0731, methenyltetrahydrofolate cyclohydrolase / 5,10-methylenetetrahydrofolate dehydrogenase (NADP+) |
| m4C | 730466 | - | | Pnuc_0731, methenyltetrahydrofolate cyclohydrolase / 5,10-methylenetetrahydrofolate dehydrogenase (NADP+) |
| m4C | 737025 | - | | upstream Pnuc_0733, PAS/PAC sensor signal transduction histidine kinase |
| m4C | 737919 | + | | Pnuc_0735, pyruvate dehydrogenase complex dihydrolipoamide acetyltransferase |
| m4C | 738620 | - | | upstream Pnuc_0733, PAS/PAC sensor signal transduction histidine kinase |
| m6A | 739406 | + | CTAYNNNNNNNTRTC | Pnuc_0736, dihydrolipoamide dehydrogenase |
| m6A | 739416 | - | GAYANNNNNNNRTAG | upstream Pnuc_0733, PAS/PAC sensor signal transduction histidine kinase |
| m6A | 741400 | + | | upstream Pnuc_0738, protein of unknown function DUF6, transmembrane |
| m6A | 741643 | + | CTAYNNNNNNNTRTC | Pnuc_0738, protein of unknown function DUF6, transmembrane |
| m6A | 741653 | - | GAYANNNNNNNRTAG | upstream Pnuc_0737, phasin family protein |
| m4C | 743185 | - | | upstream Pnuc_0737, phasin family protein |
| m6A | 750406 | + | GAYANNNNNNNRTAG | upstream Pnuc_0753, protein of unknown function DUF328 |
| m6A | 750416 | - | CTAYNNNNNNNTRTC | Pnuc_0747, DSBA oxidoreductase |
| m6A | 757609 | + | | upstream Pnuc_0757, transcriptional regulator, GntR family |
| m4C | 760193 | + | | Pnuc_0758, succinate dehydrogenase subunit C |
| m6A | 762421 | + | | Pnuc_0761, succinate dehydrogenase subunit B |
| m6A | 762452 | + | CTAYNNNNNNNTRTC | Pnuc_0761, succinate dehydrogenase subunit B |
| m6A | 762462 | - | GAYANNNNNNNRTAG | upstream Pnuc_0756, malate dehydrogenase (NAD) |
| m6A | 767599 | + | | Pnuc_0767, 3-isopropylmalate dehydrogenase |
| m4C | 767953 | + | | Pnuc_0767, 3-isopropylmalate dehydrogenase |
| m6A | 768513 | + | CTAYNNNNNNNTRTC | Pnuc_0768, aspartate semialdehyde dehydrogenase |
| m6A | 768523 | - | GAYANNNNNNNRTAG | upstream Pnuc_0756, malate dehydrogenase (NAD) |
| m4C | 769803 | + | | Pnuc_0769, Tfp pilus assembly protein FimV-like protein |
| m6A | 771608 | + | | Pnuc_0770, tRNA pseudouridine synthase A |
| m6A | 774540 | + | CTAYNNNNNNNTRTC | Pnuc_0774, acetyl-CoA carboxylase carboxyltransferase subunit alpha |
| m6A | 774550 | - | GAYANNNNNNNRTAG | upstream Pnuc_0756, malate dehydrogenase (NAD) |
| m6A | 783973 | + | GAYANNNNNNNRTAG | Pnuc_0784, OmpA/MotB domain protein |
| m6A | 783983 | - | CTAYNNNNNNNTRTC | upstream Pnuc_0782, molybdate ABC transporter, inner membrane subunit |
| m4C | 784659 | + | | Pnuc_0785, hypothetical protein |
| m6A | 784667 | - | | upstream Pnuc_0782, molybdate ABC transporter, inner membrane subunit |
| m6A | 786874 | + | GAYANNNNNNNRTAG | upstream Pnuc_0788, protein of unknown function DUF395, YeeE/YedE |
| m6A | 786884 | - | CTAYNNNNNNNTRTC | Pnuc_0787, cytochrome c family protein |
| m6A | 789389 | + | GAYANNNNNNNRTAG | Pnuc_0792, Rhodanese domain protein |
| m6A | 789399 | - | CTAYNNNNNNNTRTC | upstream Pnuc_0790, hypothetical protein |
| m6A | 793988 | + | CTAYNNNNNNNTRTC | upstream Pnuc_0800, beta-lactamase domain protein |
| m4C | 794908 | - | | upstream Pnuc_0799, FAD-dependent pyridine nucleotide-disulfide oxidoreductase |
| m4C | 797305 | + | | Pnuc_0802, sulfur dehydrogenase subunit SoxD |
| m6A | 802163 | + | GAYANNNNNNNRTAG | Pnuc_0807, sulfate thiol esterase SoxB |
| m6A | 802173 | - | CTAYNNNNNNNTRTC | upstream Pnuc_0799, FAD-dependent pyridine nucleotide-disulfide oxidoreductase |
| m4C | 812378 | - | | upstream Pnuc_0818, acyl-CoA dehydrogenase domain protein |
| m4C | 814873 | + | | upstream Pnuc_0825, 3-oxoacid CoA-transferase, A subunit |
| m6A | 815395 | + | GAYANNNNNNNRTAG | upstream Pnuc_0825, 3-oxoacid CoA-transferase, A subunit |
| m4C | 817957 | - | | upstream Pnuc_0824, Electron-transferring-flavoprotein dehydrogenase |
| m4C | 818014 | + | | Pnuc_0825, 3-oxoacid CoA-transferase, A subunit |

| | | | | |
|---|---|---|---|---|
| m4C | 819483 | - | | upstream Pnuc_0824, Electron-transferring-flavoprotein dehydrogenase |
| m6A | 820903 | + | CTAYNNNNNNNTRTC | Pnuc_0827, cation diffusion facilitator family transporter |
| m6A | 820913 | - | GAYANNNNNNNRTAG | upstream Pnuc_0824, Electron-transferring-flavoprotein dehydrogenase |
| m4C | 825453 | + | | Pnuc_0830, bacterial translation initiation factor 3(bIF-3) |
| m6A | 837877 | + | GAYANNNNNNNRTAG | Pnuc_0842, dihydrolipoamide dehydrogenase |
| m6A | 837887 | - | CTAYNNNNNNNTRTC | upstream Pnuc_0839, DNA binding domain, excisionase family |
| m4C | 843785 | + | | Pnuc_0848, NlpBDapX family lipoprotein |
| m4C | 843787 | + | | Pnuc_0848, NlpBDapX family lipoprotein |
| m4C | 846473 | - | | upstream Pnuc_0850, Cupin 4 family protein |
| m4C | 849590 | - | | Pnuc_0855, UDP-2,3-diacylglucosamine hydrolase |
| m4C | 852244 | - | | upstream Pnuc_0857, Tetratricopeptide TPR_2 repeat protein |
| m4C | 855676 | - | | upstream Pnuc_0857, Tetratricopeptide TPR_2 repeat protein |
| m6A | 856210 | + | GAYANNNNNNNRTAG | Pnuc_0862, aspartate kinase |
| m6A | 856220 | - | CTAYNNNNNNNTRTC | upstream Pnuc_0857, Tetratricopeptide TPR_2 repeat protein |
| m4C | 862259 | - | | upstream Pnuc_0867, pseudo |
| m6A | 864766 | + | CTAYNNNNNNNTRTC | Pnuc_0870, transporter, hydrophobe/amphiphile efflux-1(HAE1) family |
| m6A | 864776 | - | GAYANNNNNNNRTAG | upstream Pnuc_0865, transcriptional regulator, XRE family |
| m6A | 865018 | + | CTAYNNNNNNNTRTC | Pnuc_0870, transporter, hydrophobe/amphiphile efflux-1(HAE1) family |
| m6A | 865028 | - | GAYANNNNNNNRTAG | upstream Pnuc_0865, transcriptional regulator, XRE family |
| m6A | 868410 | + | GAYANNNNNNNRTAG | Pnuc_0872, Formyl-CoA transferase |
| m6A | 873168 | + | CTAYNNNNNNNTRTC | Pnuc_0879, putative transmembrane protein |
| m6A | 873178 | - | GAYANNNNNNNRTAG | upstream Pnuc_0875, hypothetical protein |
| m4C | 878961 | - | | Pnuc_0885, conserved hypothetical protein |
| m6A | 882214 | + | GAYANNNNNNNRTAG | Pnuc_0889, phosphomethylpyrimidine kinase |
| m6A | 882224 | - | CTAYNNNNNNNTRTC | upstream Pnuc_R0023, tRNA-Asn |
| m4C | 889815 | + | | upstream Pnuc_0900, DNA topoisomerase IV subunit B |
| m6A | 894883 | + | GAYANNNNNNNRTAG | Pnuc_0901, DNA topoisomerase IV subunit A |
| m6A | 894893 | - | CTAYNNNNNNNTRTC | upstream Pnuc_0899, Formyl-CoA transferase |
| m4C | 896466 | + | | Pnuc_0901, DNA topoisomerase IV subunit A |
| m6A | 898244 | - | GAYANNNNNNNRTAG | upstream Pnuc_0902, predicted sulfurylase subunit, molybdopterin cytosine dinucleotide biosynthesis |
| m6A | 903152 | + | GAYANNNNNNNRTAG | upstream Pnuc_0910, methylmalonyl-CoA mutase |
| m6A | 903162 | - | CTAYNNNNNNNTRTC | upstream Pnuc_0909, transcriptional regulator, GntR family |
| m6A | 905502 | + | | Pnuc_0911, LAO/AO transport system ATPase |
| m4C | 906279 | - | | upstream Pnuc_0909, transcriptional regulator, GntR family |
| m4C | 907862 | + | | Pnuc_0912, carboxyl transferase |
| m6A | 909439 | + | CTAYNNNNNNNTRTC | Pnuc_0913, acetyl-CoA carboxylase, biotin carboxylase |
| m6A | 909449 | - | GAYANNNNNNNRTAG | upstream Pnuc_0909, transcriptional regulator, GntR family |
| m4C | 918085 | + | | Pnuc_0923, poly(R)-hydroxyalkanoic acid synthase, class I |
| m4C | 925188 | + | | upstream Pnuc_0930, 2-C-methyl-D-erythritol 4-phosphate cytidylyltransferase |
| m4C | 929165 | - | | upstream Pnuc_0929, transcription-repair coupling factor |
| m6A | 933218 | - | | upstream Pnuc_0929, transcription-repair coupling factor |
| m4C | 933555 | + | | Pnuc_0935, ATP-dependent Clp protease ATP-binding subunit ClpX |
| m4C | 934509 | - | | upstream Pnuc_0929, transcription-repair coupling factor |
| m4C | 934696 | - | | upstream Pnuc_0929, transcription-repair coupling factor |

| | | | | |
|---|---|---|---|---|
| m4C | 934777 | - | | upstream Pnuc_0929, transcription-repair coupling factor |
| m4C | 936875 | - | | upstream Pnuc_0929, transcription-repair coupling factor |
| m6A | 937001 | - | GAYANNNNNNNNRTAG | upstream Pnuc_0929, transcription-repair coupling factor |
| m4C | 939907 | + | | Pnuc_0940, phosphoribosylformylglycinamidine synthase |
| m4C | 943646 | - | | upstream Pnuc_0938, Arylesterase |
| m6A | 946091 | + | GAYANNNNNNNNRTAG | upstream Pnuc_0945, CTP synthase |
| m4C | 952437 | - | | Pnuc_0949, Hsp33 protein |
| m4C | 952855 | + | | upstream Pnuc_0950, glutamyl-tRNA synthetase |
| m6A | 959025 | - | | Pnuc_0957, hypothetical protein |
| m6A | 959740 | + | GAYANNNNNNNNRTAG | Pnuc_0958, 2-hydroxy-3-oxopropionate reductase |
| m6A | 959750 | - | CTAYNNNNNNNNTRTC | upstream Pnuc_0957, hypothetical protein |
| m4C | 963789 | - | | Pnuc_0963, Ankyrin |
| m6A | 966942 | + | CTAYNNNNNNNNTRTC | upstream Pnuc_0968, glycine cleavage T protein (aminomethyl transferase) |
| m6A | 967979 | + | CTAYNNNNNNNNTRTC | Pnuc_0968, glycine cleavage T protein (aminomethyl transferase) |
| m6A | 967989 | - | GAYANNNNNNNNRTAG | upstream Pnuc_0967, aminodeoxychorismate lyase |
| m4C | 969007 | - | | upstream Pnuc_0967, aminodeoxychorismate lyase |
| m4C | 973870 | - | | Pnuc_0976, major facilitator superfamily MFS_1 |
| m6A | 974583 | + | | upstream Pnuc_0980, flavodoxin/nitric oxide synthase |
| m6A | 976832 | + | | upstream Pnuc_0980, flavodoxin/nitric oxide synthase |
| m6A | 979516 | + | | Pnuc_0982, FAD linked oxidase domain protein |
| m4C | 980292 | - | | Pnuc_0983, alpha/beta hydrolase fold protein |
| m6A | 980984 | + | GAYANNNNNNNNRTAG | upstream Pnuc_0999, ABC-type transporter, periplasmic component, NitT family |
| m6A | 980994 | - | CTAYNNNNNNNNTRTC | Pnuc_0983, alpha/beta hydrolase fold protein |
| m4C | 982136 | + | | upstream Pnuc_0999, ABC-type transporter, periplasmic component, NitT family |
| m6A | 982286 | + | CTAYNNNNNNNNTRTC | upstream Pnuc_0999, ABC-type transporter, periplasmic component, NitT family |
| m6A | 982296 | - | GAYANNNNNNNNRTAG | Pnuc_0986, hypothetical protein |
| m4C | 986038 | - | | upstream Pnuc_0990, amino acid ABC transporter substrate-binding protein, PAAT family |
| m6A | 986047 | - | | upstream Pnuc_0990, amino acid ABC transporter substrate-binding protein, PAAT family |
| m4C | 987808 | + | | upstream Pnuc_0999, ABC-type transporter, periplasmic component, NitT family |
| m6A | 988136 | + | GAYANNNNNNNNRTAG | upstream Pnuc_0999, ABC-type transporter, periplasmic component, NitT family |
| m6A | 988146 | - | CTAYNNNNNNNNTRTC | Pnuc_0992, Choloylglycine hydrolase |
| m4C | 991025 | - | | Pnuc_0995, nitrate ABC transporter, ATPase subunits C and D |
| m6A | 995404 | + | CTAYNNNNNNNNTRTC | Pnuc_0999, ABC-type transporter, periplasmic component, NitT family |
| m6A | 995414 | - | | upstream Pnuc_0998, uroporphyrin-III C-methyltransferase |
| m6A | 996739 | + | | Pnuc_1000, assimilatory nitrite reductase (NAD(P)H) large subunit precursor |
| m6A | 1008651 | + | CTAYNNNNNNNNTRTC | Pnuc_1010, TonB-dependent receptor |
| m6A | 1008661 | - | GAYANNNNNNNNRTAG | upstream Pnuc_R0031, tRNA-Pro2 |
| m4C | 1009686 | + | | upstream Pnuc_1015, protein of unknown function UPF0044 |
| m4C | 1010247 | + | | upstream Pnuc_1015, protein of unknown function UPF0044 |
| m4C | 1015036 | - | | Pnuc_1017, transcription elongation factor GreA |
| m4C | 1015612 | - | | Pnuc_1018, carbamoyl-phosphate synthase large subunit |
| m4C | 1016688 | + | | upstream Pnuc_1023, Methyltransferase type 11 |
| m6A | 1017202 | + | | upstream Pnuc_1023, Methyltransferase type 11 |
| m6A | 1019053 | + | CTAYNNNNNNNNTRTC | upstream Pnuc_1023, Methyltransferase type 11 |

| | | | | |
|---|---|---|---|---|
| m6A | 1019063 | - | GAYANNNNNNNNRTAG | Pnuc_1019, carbamoyl-phosphate synthase small subunit |
| m6A | 1020256 | + | CTAYNNNNNNNNTRTC | upstream Pnuc_1023, Methyltransferase type 11 |
| m6A | 1020266 | - | GAYANNNNNNNNRTAG | upstream Pnuc_1019, carbamoyl-phosphate synthase small subunit |
| m6A | 1021303 | + | CTAYNNNNNNNNTRTC | upstream Pnuc_1023, Methyltransferase type 11 |
| m6A | 1021312 | + | | upstream Pnuc_1023, Methyltransferase type 11 |
| m6A | 1021313 | - | GAYANNNNNNNNRTAG | Pnuc_1020, propionyl-CoA synthetase |
| m6A | 1022212 | - | | Pnuc_1021, Lytic transglycosylase, catalytic |
| m6A | 1025742 | + | | upstream Pnuc_1025, efflux transporter, RND family, MFP subunit |
| m6A | 1027462 | + | GAYANNNNNNNNRTAG | Pnuc_1026, acriflavin resistance protein |
| m6A | 1027472 | - | CTAYNNNNNNNNTRTC | upstream Pnuc_1022, Hydroxyacylglutathione hydrolase |
| m4C | 1032904 | - | | Pnuc_1029, hypothetical protein |
| m6A | 1033357 | + | | upstream Pnuc_1032, Amidase |
| m6A | 1034060 | + | CTAYNNNNNNNNTRTC | upstream Pnuc_1032, Amidase |
| m6A | 1034070 | - | GAYANNNNNNNNRTAG | Pnuc_1031, hypothetical protein |
| m4C | 1034143 | + | | upstream Pnuc_1032, Amidase |
| m4C | 1036580 | + | | upstream Pnuc_1035, methylmalonate-semialdehyde dehydrogenase (acylating) |
| m4C | 1037210 | + | | upstream Pnuc_1035, methylmalonate-semialdehyde dehydrogenase (acylating) |
| m4C | 1037402 | - | | Pnuc_1034, NADH:flavin oxidoreductase/NADH oxidase |
| m4C | 1038133 | + | | upstream Pnuc_1035, methylmalonate-semialdehyde dehydrogenase (acylating) |
| m4C | 1039836 | + | | Pnuc_1035, methylmalonate-semialdehyde dehydrogenase (acylating) |
| m6A | 1047413 | - | | Pnuc_1043, NADH dehydrogenase subunit I |
| m4C | 1053797 | + | | upstream Pnuc_1057, putative lipoprotein |
| m6A | 1055538 | + | GAYANNNNNNNNRTAG | upstream Pnuc_1057, putative lipoprotein |
| m4C | 1058121 | - | | Pnuc_1054, Alcohol dehydrogenase, zinc-binding domain protein |
| m4C | 1059733 | + | | upstream Pnuc_1057, putative lipoprotein |
| m4C | 1063511 | - | | Pnuc_1059, CDP-diacylglycerol-serine O-phosphatidyltransferase |
| m4C | 1068845 | + | | Pnuc_1064, RNA polymerase, sigma-24 subunit, RpoE |
| m6A | 1075780 | + | GAYANNNNNNNNRTAG | upstream Pnuc_1075, transcription elongation factor GreB |
| m6A | 1075790 | - | CTAYNNNNNNNNTRTC | Pnuc_1071, hypothetical protein |
| m6A | 1076951 | + | CTAYNNNNNNNNTRTC | upstream Pnuc_1075, transcription elongation factor GreB |
| m6A | 1076961 | - | GAYANNNNNNNNRTAG | Pnuc_1073, ABC transporter related protein |
| m4C | 1082483 | + | | upstream Pnuc_1080, RNAse PH |
| m6A | 1092756 | - | | upstream Pnuc_1085, integral membrane sensor signal transduction histidine kinase |
| m6A | 1093084 | + | GAYANNNNNNNNRTAG | Pnuc_1087, hypothetical protein |
| m6A | 1093094 | - | CTAYNNNNNNNNTRTC | upstream Pnuc_1085, integral membrane sensor signal transduction histidine kinase |
| m6A | 1095712 | + | CTAYNNNNNNNNTRTC | upstream Pnuc_1091, glucose-6-phosphate isomerase |
| m6A | 1095722 | - | GAYANNNNNNNNRTAG | upstream Pnuc_1089, hypothetical protein |
| m4C | 1098640 | - | | upstream Pnuc_1090, FAD dependent oxidoreductase |
| m4C | 1099741 | + | | Pnuc_1092, phosphomannomutase |
| m4C | 1101992 | - | | Pnuc_1094, Polypeptide-transport-associated domain protein, ShlB-type |
| m6A | 1108023 | + | GAYANNNNNNNNRTAG | upstream Pnuc_1097, hypothetical protein |
| m6A | 1108033 | - | CTAYNNNNNNNNTRTC | Pnuc_1095, outer membrane autotransporter barrel domain protein |
| m6A | 1109515 | + | CTAYNNNNNNNNTRTC | upstream Pnuc_1097, hypothetical protein |
| m6A | 1109525 | - | GAYANNNNNNNNRTAG | Pnuc_1095, outer membrane autotransporter barrel domain protein |

| | | | | |
|---|---|---|---|---|
| m6A | 1111707 | + | GAYANNNNNNNRTAG | upstream Pnuc_1097, hypothetical protein |
| m6A | 1111717 | - | CTAYNNNNNNNTRTC | Pnuc_1095, outer membrane autotransporter barrel domain protein |
| m6A | 1115916 | + | GAYANNNNNNNRTAG | upstream Pnuc_1097, hypothetical protein |
| m6A | 1115926 | - | CTAYNNNNNNNTRTC | Pnuc_1095, outer membrane autotransporter barrel domain protein |
| m4C | 1116271 | - | | Pnuc_1095, outer membrane autotransporter barrel domain protein |
| m4C | 1116515 | + | | upstream Pnuc_1097, hypothetical protein |
| m4C | 1118009 | + | | upstream Pnuc_1097, hypothetical protein |
| m6A | 1120460 | + | | upstream Pnuc_1097, hypothetical protein |
| m6A | 1128653 | - | | Pnuc_1095, outer membrane autotransporter barrel domain protein |
| m6A | 1134205 | - | | upstream Pnuc_1095, outer membrane autotransporter barrel domain protein |
| m4C | 1136504 | - | | Pnuc_1096, sulfotransferase |
| m4C | 1137761 | + | | upstream Pnuc_1097, hypothetical protein |
| m4C | 1138186 | + | | upstream Pnuc_1097, hypothetical protein |
| m6A | 1143445 | + | | upstream Pnuc_1101, protein of unknown function DUF583 |
| m6A | 1143575 | - | | upstream Pnuc_1100, peptidase M48, Ste24p |
| m6A | 1150445 | + | GAYANNNNNNNRTAG | upstream Pnuc_1112, protein of unknown function DUF1080 |
| m6A | 1150455 | - | CTAYNNNNNNNTRTC | Pnuc_1107, transcriptional modulator of MazE/toxin, MazF |
| m6A | 1151487 | + | GAYANNNNNNNRTAG | upstream Pnuc_1112, protein of unknown function DUF1080 |
| m6A | 1151497 | - | CTAYNNNNNNNTRTC | Pnuc_1109, NAD-dependent formate dehydrogenase iron-sulfur protein (catalytic activity) |
| m6A | 1156356 | - | GAYANNNNNNNRTAG | Pnuc_1111, protein of unknown function DUF748 |
| m6A | 1157066 | - | | Pnuc_1111, protein of unknown function DUF748 |
| m6A | 1159304 | + | GAYANNNNNNNRTAG | Pnuc_1112, protein of unknown function DUF1080 |
| m6A | 1159314 | - | CTAYNNNNNNNTRTC | upstream Pnuc_1111, protein of unknown function DUF748 |
| m6A | 1167044 | + | CTAYNNNNNNNTRTC | Pnuc_1122, Exonuclease, RNase T and DNA polymerase III |
| m6A | 1167054 | - | GAYANNNNNNNRTAG | upstream Pnuc_1121, hypothetical protein |
| m4C | 1169873 | + | | upstream Pnuc_1127, hypothetical protein |
| m6A | 1173977 | + | GAYANNNNNNNRTAG | Pnuc_1127, hypothetical protein |
| m6A | 1173987 | - | CTAYNNNNNNNTRTC | upstream Pnuc_1126, hypothetical protein |
| m6A | 1177319 | + | GAYANNNNNNNRTAG | upstream Pnuc_1131, phage transcriptional regulator, AlpA |
| m6A | 1177329 | - | CTAYNNNNNNNTRTC | Pnuc_1130, protein of unknown function DUF1376 |
| m4C | 1190609 | + | | Pnuc_1139, fumarase |
| m6A | 1191165 | + | CTAYNNNNNNNTRTC | Pnuc_1140, glutamate racemase |
| m6A | 1191175 | - | GAYANNNNNNNRTAG | upstream Pnuc_1138, acetyl-coenzyme A synthetase |
| m4C | 1192583 | + | | Pnuc_1142, outer membrane transport energization protein ExbB |
| m6A | 1199103 | + | | upstream Pnuc_1150, Penicillin amidase |
| m6A | 1199870 | + | GAYANNNNNNNRTAG | upstream Pnuc_1150, Penicillin amidase |
| m6A | 1199880 | - | CTAYNNNNNNNTRTC | Pnuc_1149, filamentous hemagglutinin family outer membrane protein |
| m6A | 1206854 | + | CTAYNNNNNNNTRTC | Pnuc_1150, Penicillin amidase |
| m6A | 1206864 | - | GAYANNNNNNNRTAG | upstream Pnuc_R0036, tRNA-Leu |
| m4C | 1210903 | - | | Pnuc_1158, aldehyde oxidase and xanthine dehydrogenase, molybdopterin binding protein |
| m4C | 1216436 | + | | Pnuc_1163, gamma-glutamyltransferase 2, Threonine peptidase, MEROPS family T03 |
| m6A | 1218238 | + | CTAYNNNNNNNTRTC | upstream Pnuc_1167, Cellulase |
| m6A | 1218248 | - | GAYANNNNNNNRTAG | Pnuc_1164, conserved hypothetical protein |
| m4C | 1225329 | - | | Pnuc_1170, cellulose synthase operon C domain protein |

| | | | | |
|---|---|---|---|---|
| m6A | 1228000 | + | GAYANNNNNNNRTAG | upstream Pnuc_1176, hypothetical protein |
| m6A | 1228010 | - | CTAYNNNNNNNTRTC | Pnuc_1170, cellulose synthase operon C domain protein |
| m6A | 1228683 | + | GAYANNNNNNNRTAG | upstream Pnuc_1176, hypothetical protein |
| m6A | 1228693 | - | CTAYNNNNNNNTRTC | Pnuc_1170, cellulose synthase operon C domain protein |
| m6A | 1228990 | + | CTAYNNNNNNNTRTC | upstream Pnuc_1176, hypothetical protein |
| m4C | 1230189 | - | | Pnuc_1171, Cellulose synthase (UDP-forming) |
| m4C | 1235017 | + | | Pnuc_1176, hypothetical protein |
| m4C | 1235061 | - | | upstream Pnuc_1175, metal dependent phosphohydrolase |
| m6A | 1236722 | - | GAYANNNNNNNRTAG | upstream Pnuc_1177, tRNA-U16,U17-dihydrouridine synthase |
| m6A | 1239968 | + | GAYANNNNNNNRTAG | upstream Pnuc_1184, Amidase |
| m6A | 1239978 | - | CTAYNNNNNNNTRTC | Pnuc_1180, putative ABC transporter, periplasmic protein |
| m6A | 1240127 | + | GAYANNNNNNNRTAG | upstream Pnuc_1184, Amidase |
| m4C | 1246490 | + | | Pnuc_1187, ABC transporter related protein |
| m6A | 1247136 | + | | upstream Pnuc_1210, Glutamate synthase (NADPH) |
| m6A | 1247968 | + | CTAYNNNNNNNTRTC | upstream Pnuc_1210, Glutamate synthase (NADPH) |
| m6A | 1247978 | - | GAYANNNNNNNRTAG | Pnuc_1190, urease accessory protein UreG |
| m4C | 1251636 | + | | upstream Pnuc_1210, Glutamate synthase (NADPH) |
| m6A | 1252457 | + | GAYANNNNNNNRTAG | upstream Pnuc_1210, Glutamate synthase (NADPH) |
| m6A | 1252467 | - | CTAYNNNNNNNTRTC | Pnuc_1196, urease, gamma subunit |
| m4C | 1257725 | + | | upstream Pnuc_1210, Glutamate synthase (NADPH) |
| m4C | 1260318 | + | | upstream Pnuc_1210, Glutamate synthase (NADPH) |
| m4C | 1260888 | + | | upstream Pnuc_1210, Glutamate synthase (NADPH) |
| m4C | 1261107 | + | | upstream Pnuc_1210, Glutamate synthase (NADPH) |
| m6A | 1262425 | + | CTAYNNNNNNNTRTC | upstream Pnuc_1210, Glutamate synthase (NADPH) |
| m6A | 1262435 | - | GAYANNNNNNNRTAG | Pnuc_1205, protein of unknown function DUF1501 |
| m6A | 1263897 | - | | Pnuc_1206, conserved hypothetical protein |
| m6A | 1269593 | + | | upstream Pnuc_1218, transglutaminase, N-terminal domain protein |
| m4C | 1271444 | + | | upstream Pnuc_1218, transglutaminase, N-terminal domain protein |
| m4C | 1283494 | + | | Pnuc_1223, major facilitator superfamily MFS_1 |
| m4C | 1284569 | + | | upstream Pnuc_1232, Patatin |
| m4C | 1288419 | + | | upstream Pnuc_1232, Patatin |
| m4C | 1296340 | - | | Pnuc_1233, NLP/P60 protein |
| m6A | 1296429 | + | CTAYNNNNNNNTRTC | upstream Pnuc_1238, Enoyl-[acyl-carrier-protein] reductase (NADH) |
| m6A | 1296439 | - | GAYANNNNNNNRTAG | upstream Pnuc_1233, NLP/P60 protein |
| m6A | 1301267 | - | | Pnuc_1237, extracellular solute-binding protein, family 5 |
| m4C | 1303180 | + | | Pnuc_1239, chromate transporter, chromate ion transporter(CHR) family |
| m4C | 1304745 | + | | Pnuc_1240, Integrase, catalytic region |
| m4C | 1305997 | - | | upstream Pnuc_1237, extracellular solute-binding protein, family 5 |
| m6A | 1307149 | + | GAYANNNNNNNRTAG | Pnuc_1242, MmgE/PrpD family protein |
| m6A | 1307159 | - | CTAYNNNNNNNTRTC | upstream Pnuc_1237, extracellular solute-binding protein, family 5 |
| m6A | 1311414 | + | CTAYNNNNNNNTRTC | upstream Pnuc_1249, integrase, catalytic region |
| m6A | 1311424 | - | GAYANNNNNNNRTAG | Pnuc_1246, BNR/Asp-box repeat protein |
| m6A | 1312246 | - | | Pnuc_1247, glycosyl transferase, family 39 |
| m4C | 1313486 | + | | upstream Pnuc_1249, integrase, catalytic region |

| | | | | |
|---|---|---|---|---|
| m4C | 1315427 | + | | upstream Pnuc_1251, pseudo |
| m6A | 1317420 | + | CTAYNNNNNNNNTRTC | upstream Pnuc_1262, N-acetylglutamate synthase |
| m6A | 1317430 | - | GAYANNNNNNNNRTAG | Pnuc_1252, Exodeoxyribonuclease III |
| m4C | 1318201 | + | | upstream Pnuc_1262, N-acetylglutamate synthase |
| m4C | 1318723 | + | | upstream Pnuc_1262, N-acetylglutamate synthase |
| m4C | 1318907 | - | | Pnuc_1253, nitrogen metabolism transcriptional regulator, NtrC, Fis family |
| m6A | 1325390 | + | | upstream Pnuc_1262, N-acetylglutamate synthase |
| m4C | 1336966 | - | | Pnuc_1267, propionate CoA-transferase |
| m6A | 1341847 | + | | upstream Pnuc_R0038, tRNA-Leu |
| m4C | 1345263 | + | | upstream Pnuc_1278, hypothetical protein |
| m4C | 1350920 | - | | Pnuc_1280, malate synthase G |
| m4C | 1357004 | + | | upstream Pnuc_1294, protein of unknown function UPF0005 |
| m4C | 1363387 | + | | upstream Pnuc_1294, protein of unknown function UPF0005 |
| m4C | 1366237 | - | | upstream Pnuc_1293, nucleoside diphosphate kinase |
| m4C | 1369370 | + | | upstream Pnuc_1307, thioredoxin |
| m4C | 1378180 | - | | Pnuc_1304, Tetratricopeptide TPR_2 repeat protein |
| m4C | 1384960 | + | | upstream Pnuc_1307, thioredoxin |
| m4C | 1386002 | - | | upstream Pnuc_1306, DNA helicase/exodeoxyribonuclease V, subunit B |
| m6A | 1388390 | - | | upstream Pnuc_1306, DNA helicase/exodeoxyribonuclease V, subunit B |
| m6A | 1395251 | + | | upstream Pnuc_1318, Enoyl-CoA hydratase/isomerase |
| m6A | 1397943 | - | | Pnuc_1319, Lytic transglycosylase, catalytic |
| m4C | 1400832 | - | | upstream Pnuc_1322, conserved hypothetical protein |
| m4C | 1405083 | + | | upstream Pnuc_1331, Carbohydrate-selective porin OprB |
| m6A | 1410626 | + | GAYANNNNNNNNRTAG | Pnuc_1332, negative transcriptional regulator |
| m6A | 1413234 | + | | upstream Pnuc_1336, putative sulfate transport system substrate-binding protein |
| m6A | 1416762 | + | CTAYNNNNNNNNTRTC | Pnuc_1340, thioesterase superfamily protein |
| m6A | 1416772 | - | GAYANNNNNNNNRTAG | upstream Pnuc_1338, hypothetical protein |
| m6A | 1417579 | + | GAYANNNNNNNNRTAG | Pnuc_1341, conserved hypothetical protein |
| m6A | 1417589 | - | CTAYNNNNNNNNTRTC | upstream Pnuc_1338, hypothetical protein |
| m6A | 1419263 | + | CTAYNNNNNNNNTRTC | Pnuc_1343, transglutaminase domain protein |
| m6A | 1419273 | - | GAYANNNNNNNNRTAG | upstream Pnuc_1338, hypothetical protein |
| m6A | 1422263 | + | CTAYNNNNNNNNTRTC | upstream Pnuc_1347, protein of unknown function DUF1330 |
| m6A | 1422273 | - | GAYANNNNNNNNRTAG | upstream Pnuc_1346, conserved hypothetical protein |
| m4C | 1423119 | + | | Pnuc_1349, Uncharacterized protein UPF0065 |
| m6A | 1432389 | - | | upstream Pnuc_1359, conserved hypothetical protein 730 |
| m4C | 1433506 | + | | upstream Pnuc_1363, carbohydrate kinase, YjeF related protein |
| m6A | 1434445 | + | CTAYNNNNNNNNTRTC | upstream Pnuc_1363, carbohydrate kinase, YjeF related protein |
| m6A | 1434455 | - | GAYANNNNNNNNRTAG | Pnuc_1362, RNA-metabolising metallo-beta-lactamase |
| m6A | 1436527 | + | | Pnuc_1364, protein of unknown function DUF6, transmembrane |
| m4C | 1442524 | - | | upstream Pnuc_1370, protein of unknown function DUF6, transmembrane |
| m6A | 1448329 | + | | Pnuc_1378, FAD-dependent pyridine nucleotide-disulfide oxidoreductase |
| m6A | 1449009 | + | | Pnuc_1378, FAD-dependent pyridine nucleotide-disulfide oxidoreductase |
| m4C | 1451114 | - | | Pnuc_1381, type II and III secretion system protein |
| m6A | 1451211 | + | | upstream Pnuc_1382, DSBA oxidoreductase |

| | | | | |
|---|---|---|---|---|
| m6A | 1451215 | + | | upstream Pnuc_1382, DSBA oxidoreductase |
| m4C | 1451602 | + | | upstream Pnuc_1382, DSBA oxidoreductase |
| m4C | 1453033 | - | | Pnuc_1381, type II and III secretion system protein |
| m4C | 1454705 | + | | upstream Pnuc_1388, acyl-CoA dehydrogenase domain protein |
| m4C | 1462285 | - | | Pnuc_1391, Endonuclease/exonuclease/phosphatase |
| m4C | 1463215 | - | | upstream Pnuc_1391, Endonuclease/exonuclease/phosphatase |
| m6A | 1464604 | - | | Pnuc_1394, hypothetical protein |
| m6A | 1464729 | + | GAYANNNNNNNNRTAG | upstream Pnuc_1407, heavy metal translocating P-type ATPase |
| m6A | 1464739 | - | CTAYNNNNNNNNTRTC | Pnuc_1395, formate dehydrogenase gamma subunit |
| m4C | 1466348 | + | | upstream Pnuc_1407, heavy metal translocating P-type ATPase |
| m4C | 1470475 | + | | upstream Pnuc_1407, heavy metal translocating P-type ATPase |
| m6A | 1473645 | + | GAYANNNNNNNNRTAG | upstream Pnuc_1407, heavy metal translocating P-type ATPase |
| m6A | 1473655 | - | CTAYNNNNNNNNTRTC | Pnuc_1402, conserved hypothetical protein |
| m4C | 1476780 | - | | upstream Pnuc_1405, protein of unknown function DUF47 |
| m4C | 1479800 | + | | upstream Pnuc_1409, 4-amino-4-deoxy-L-arabinose transferase and related glycosyltransferase of PMT family-like protein |
| m4C | 1479959 | + | | upstream Pnuc_1409, 4-amino-4-deoxy-L-arabinose transferase and related glycosyltransferase of PMT family-like protein |
| m6A | 1488814 | + | | upstream Pnuc_1426, SsrA-binding protein |
| m6A | 1488824 | - | | Pnuc_1418, Radical SAM domain protein |
| m4C | 1496040 | + | | upstream Pnuc_1426, SsrA-binding protein |
| m6A | 1496168 | + | CTAYNNNNNNNNTRTC | upstream Pnuc_1426, SsrA-binding protein |
| m6A | 1496178 | - | GAYANNNNNNNNRTAG | Pnuc_1425, cyclase/dehydrase |
| m4C | 1497236 | - | | upstream Pnuc_1425, cyclase/dehydrase |
| m4C | 1498737 | - | | upstream Pnuc_1425, cyclase/dehydrase |
| m6A | 1503143 | - | | Pnuc_1433, phosphoenolpyruvate synthase |
| m6A | 1505598 | + | CTAYNNNNNNNNTRTC | Pnuc_1434, protein of unknown function DUF299 |
| m6A | 1505608 | - | GAYANNNNNNNNRTAG | Pnuc_1433, phosphoenolpyruvate synthase |
| m4C | 1506886 | - | | Pnuc_1435, secretory lipase |
| m6A | 1509758 | + | | upstream Pnuc_1452, methionine aminopeptidase, type I |
| m4C | 1515771 | + | | upstream Pnuc_1452, methionine aminopeptidase, type I |
| m6A | 1519366 | - | GAYANNNNNNNNRTAG | Pnuc_1448, ribosome recycling factor |
| m6A | 1532526 | + | | upstream Pnuc_1458, 2,3,4,5-tetrahydropyridine-2,6-dicarboxylate N-succinyltransferase |
| m4C | 1536151 | - | | upstream Pnuc_1457, chromosome segregation protein SMC |
| m6A | 1545568 | - | | Pnuc_1468, lipoprotein releasing system, transmembrane protein, LolC/E family |
| m4C | 1546532 | - | | upstream Pnuc_1468, lipoprotein releasing system, transmembrane protein, LolC/E family |
| m4C | 1547566 | - | | upstream Pnuc_1468, lipoprotein releasing system, transmembrane protein, LolC/E family |
| m6A | 1548857 | + | | Pnuc_1470, exonuclease RecJ |
| m6A | 1552414 | + | GAYANNNNNNNNRTAG | upstream Pnuc_1477, peptide deformylase |
| m6A | 1552424 | - | CTAYNNNNNNNNTRTC | Pnuc_1473, hypothetical protein |
| m6A | 1553318 | + | | upstream Pnuc_1477, peptide deformylase |
| m4C | 1554796 | - | | Pnuc_1476, cysteine synthase A |
| m4C | 1556507 | + | | Pnuc_1479, Ferredoxin-NADP(+) reductase |
| m6A | 1556599 | + | | Pnuc_1479, Ferredoxin-NADP(+) reductase |
| m6A | 1556884 | - | | upstream Pnuc_1478, disulfide bond formation protein DsbB |
| m6A | 1557612 | + | CTAYNNNNNNNNTRTC | Pnuc_1480, protein of unknown function DUF81 |

| | | | | |
|---|---|---|---|---|
| m6A | 1557622 | - | GAYANNNNNNNNRTAG | upstream Pnuc_1478, disulfide bond formation protein DsbB |
| m4C | 1569924 | - | | Pnuc_1494, transcriptional regulator, BadM/Rrf2 family |
| m4C | 1570062 | - | | Pnuc_1494, transcriptional regulator, BadM/Rrf2 family |
| m4C | 1570576 | - | | Pnuc_1495, Excinuclease ABC subunit B |
| m4C | 1570822 | - | | Pnuc_1495, Excinuclease ABC subunit B |
| m6A | 1570852 | - | GAYANNNNNNNNRTAG | Pnuc_1495, Excinuclease ABC subunit B |
| m4C | 1572033 | - | | Pnuc_1495, Excinuclease ABC subunit B |
| m4C | 1575003 | + | | Pnuc_1497, polyhydroxyalkanoate depolymerase,intracellular |
| m6A | 1576519 | + | | Pnuc_1500, conserved hypothetical protein |
| m4C | 1579164 | + | | upstream Pnuc_1528, conserved hypothetical protein |
| m4C | 1581561 | + | | upstream Pnuc_1528, conserved hypothetical protein |
| m6A | 1582960 | + | GAYANNNNNNNNRTAG | upstream Pnuc_1528, conserved hypothetical protein |
| m6A | 1582970 | - | CTAYNNNNNNNNTRTC | Pnuc_1507, major facilitator superfamily MFS_1 |
| m6A | 1591897 | + | CTAYNNNNNNNNTRTC | upstream Pnuc_1528, conserved hypothetical protein |
| m6A | 1591907 | - | GAYANNNNNNNNRTAG | Pnuc_1518, heme exporter protein CcmA |
| m4C | 1595553 | - | | Pnuc_1524, beta-lactamase domain protein |
| m6A | 1599496 | + | CTAYNNNNNNNNTRTC | Pnuc_1529, histone deacetylase superfamily |
| m6A | 1599506 | - | GAYANNNNNNNNRTAG | upstream Pnuc_1527, hypothetical protein |
| m4C | 1600146 | - | | Pnuc_1530, putative lipoprotein |
| m6A | 1601203 | + | GAYANNNNNNNNRTAG | upstream Pnuc_1534, alkyl hydroperoxide reductase/ Thiol specific antioxidant/ Mal allergen |
| m6A | 1601213 | - | CTAYNNNNNNNNTRTC | Pnuc_1531, putative lipoprotein |
| m6A | 1606433 | + | GAYANNNNNNNNRTAG | upstream Pnuc_1544, transcriptional regulator, GntR family |
| m6A | 1606443 | - | CTAYNNNNNNNNTRTC | Pnuc_1538, TRAP C4-dicarboxylate transport system permease DctM subunit |
| m6A | 1612846 | + | | upstream Pnuc_1544, transcriptional regulator, GntR family |
| m6A | 1614081 | + | | Pnuc_1544, transcriptional regulator, GntR family |
| m6A | 1617388 | + | GAYANNNNNNNNRTAG | Pnuc_1547, protein of unknown function DUF6, transmembrane |
| m6A | 1617398 | - | CTAYNNNNNNNNTRTC | upstream Pnuc_1545, class II aldolase/adducin family protein |
| m4C | 1618715 | + | | upstream Pnuc_1553, transcriptional regulator, AsnC family |
| m4C | 1619966 | + | | upstream Pnuc_1553, transcriptional regulator, AsnC family |
| m6A | 1623159 | + | | upstream Pnuc_1553, transcriptional regulator, AsnC family |
| m4C | 1626430 | + | | Pnuc_1554, short-chain dehydrogenase/reductase SDR |
| m6A | 1628517 | + | GAYANNNNNNNNRTAG | upstream Pnuc_1560, transcriptional regulator, LysR family |
| m6A | 1628527 | - | CTAYNNNNNNNNTRTC | Pnuc_1556, Enoyl-CoA hydratase/isomerase |
| m6A | 1635651 | - | | upstream Pnuc_1561, major facilitator superfamily MFS_1 |
| m6A | 1638119 | + | CTAYNNNNNNNNTRTC | Pnuc_1564, FMN-binding domain protein |
| m4C | 1646478 | + | | upstream Pnuc_1581, hypothetical protein |
| m6A | 1647640 | - | | Pnuc_1575, anti-ECFsigma factor, ChrR |
| m6A | 1650038 | + | | upstream Pnuc_1581, hypothetical protein |
| m4C | 1654385 | + | | upstream Pnuc_1585, metallophosphoesterase |
| m4C | 1654654 | + | | upstream Pnuc_1585, metallophosphoesterase |
| m6A | 1655064 | - | | Pnuc_1584, lyso-ornithine lipid acyltransferase |
| m6A | 1658012 | + | GAYANNNNNNNNRTAG | Pnuc_1588, ornithine-acyl[acyl carrier protein]N-acyltransferase |
| m6A | 1658022 | - | CTAYNNNNNNNNTRTC | upstream Pnuc_1584, lyso-ornithine lipid acyltransferase |
| m6A | 1660810 | + | CTAYNNNNNNNNTRTC | upstream Pnuc_1595, MgtC/SapB transporter |

| | | | | |
|---|---|---|---|---|
| m6A | 1660820 | - | GAYANNNNNNNRTAG | Pnuc_1591, MgtC/SapB transporter |
| m4C | 1661237 | + | | upstream Pnuc_1595, MgtC/SapB transporter |
| m4C | 1668542 | + | | upstream Pnuc_1606, short-chain dehydrogenase/reductase SDR |
| m4C | 1674039 | - | | Pnuc_1604, conserved hypothetical protein |
| m6A | 1681242 | - | | upstream Pnuc_1613, conserved hypothetical membrane spanning protein |
| m4C | 1683718 | + | | upstream Pnuc_1622, hypothetical protein |
| m4C | 1689135 | + | | upstream Pnuc_1622, hypothetical protein |
| m4C | 1690534 | - | | upstream Pnuc_1621, Mg2+ transporter protein, CorA family protein |
| m4C | 1698112 | - | | Pnuc_1631, Hydroxypyruvate isomerase |
| m6A | 1699838 | + | CTAYNNNNNNNTRTC | upstream Pnuc_1636, transcriptional regulator, GntR family |
| m6A | 1699848 | - | | Pnuc_1633, type III effector Hrp-dependent outer domain protein |
| m4C | 1702669 | - | | Pnuc_1635, major facilitator superfamily MFS_1 |
| m6A | 1705954 | + | CTAYNNNNNNNTRTC | Pnuc_1638, protein of unknown function DUF6, transmembrane |
| m6A | 1705964 | - | GAYANNNNNNNRTAG | upstream Pnuc_1637, NAD-dependent epimerase/dehydratase |
| m4C | 1707923 | + | | upstream Pnuc_1642, DSBA oxidoreductase |
| m6A | 1712078 | + | CTAYNNNNNNNTRTC | upstream Pnuc_1649, hypothetical protein |
| m6A | 1712088 | - | GAYANNNNNNNRTAG | Pnuc_1647, DNA polymerase III, epsilon subunit |
| m6A | 1712822 | + | CTAYNNNNNNNTRTC | upstream Pnuc_1649, hypothetical protein |
| m6A | 1712832 | - | GAYANNNNNNNRTAG | Pnuc_1647, DNA polymerase III, epsilon subunit |
| m6A | 1714221 | - | | upstream Pnuc_1648, transcriptional regulator, BadM/Rrf2 family |
| m4C | 1718598 | + | | Pnuc_1655, FAD dependent oxidoreductase |
| m4C | 1722369 | - | | upstream Pnuc_1659, protein of unknown function DUF427 |
| m4C | 1726936 | + | | upstream Pnuc_1664, spermine/spermidine N-acetyltransferase |
| m6A | 1730416 | - | | Pnuc_1667, osmosensitive K+ channel signal transduction histidine kinase |
| m6A | 1736959 | + | GAYANNNNNNNRTAG | upstream Pnuc_1678, putative transcriptional regulator, MerR family |
| m6A | 1736969 | - | CTAYNNNNNNNTRTC | Pnuc_1670, potassium-transporting ATPase, A subunit |
| m6A | 1738403 | + | CTAYNNNNNNNTRTC | upstream Pnuc_1678, putative transcriptional regulator, MerR family |
| m6A | 1738413 | - | GAYANNNNNNNRTAG | Pnuc_1672, Heavy metal transport/detoxification protein |
| m4C | 1742637 | + | | upstream Pnuc_1678, putative transcriptional regulator, MerR family |
| m4C | 1743423 | + | | Pnuc_1678, putative transcriptional regulator, MerR family |
| m4C | 1748045 | - | | Pnuc_1681, efflux transporter, RND family, MFP subunit |
| m6A | 1748827 | + | GAYANNNNNNNRTAG | upstream Pnuc_1688, hypothetical protein |
| m6A | 1748837 | - | CTAYNNNNNNNTRTC | Pnuc_1681, efflux transporter, RND family, MFP subunit |
| m4C | 1749332 | - | | Pnuc_1682, hypothetical protein |
| m6A | 1758014 | + | GAYANNNNNNNRTAG | upstream Pnuc_1702, O-sialoglycoprotein endopeptidase |
| m6A | 1758024 | - | CTAYNNNNNNNTRTC | Pnuc_1694, phage integrase family protein |
| m4C | 1765686 | + | | upstream Pnuc_1702, O-sialoglycoprotein endopeptidase |
| m4C | 1769216 | - | | upstream Pnuc_1701, SSU ribosomal protein S21P |
| m6A | 1769409 | + | | Pnuc_1702, O-sialoglycoprotein endopeptidase |
| m6A | 1771008 | - | | Pnuc_1704, 1-deoxy-D-xylulose-5-phosphate synthase |
| m4C | 1773158 | + | | upstream Pnuc_1707, Rieske (2Fe-2S) domain protein |
| m4C | 1782299 | + | | Pnuc_1714, putative transmembrane protein |
| m4C | 1784344 | + | | upstream Pnuc_1716, valyl-tRNA synthetase |
| m4C | 1789753 | - | | Pnuc_1719, alanyl-tRNA synthetase |

| | | | | |
|---|---|---|---|---|
| m4C | 1792905 | + | | Pnuc_1721, protein of unknown function DUF112,transmembrane |
| m6A | 1797655 | + | | Pnuc_1725, acetylornithine aminotransferase apoenzyme |
| m6A | 1797750 | + | CTAYNNNNNNNNTRTC | Pnuc_1725, acetylornithine aminotransferase apoenzyme |
| m6A | 1797760 | - | GAYANNNNNNNNRTAG | upstream Pnuc_1723, NAD-dependent epimerase/dehydratase |
| m4C | 1799103 | + | | upstream Pnuc_1733, DNA replication and repair protein RadC |
| m6A | 1805177 | + | | Pnuc_1733, DNA replication and repair protein RadC |
| m6A | 1809128 | - | | Pnuc_1738, formyltetrahydrofolate-dependent phosphoribosylglycinamide formyltransferase |
| m6A | 1809406 | + | | upstream Pnuc_1739, FMN adenylyltransferase / riboflavin kinase |
| m4C | 1810877 | + | | Pnuc_1740, Isoleucyl-tRNA synthetase |
| m4C | 1811212 | - | | upstream Pnuc_1738, formyltetrahydrofolate-dependent phosphoribosylglycinamide formyltransferase |
| m6A | 1813329 | - | | upstream Pnuc_1738, formyltetrahydrofolate-dependent phosphoribosylglycinamide formyltransferase |
| m4C | 1815519 | - | | upstream Pnuc_1738, formyltetrahydrofolate-dependent phosphoribosylglycinamide formyltransferase |
| m6A | 1816094 | - | | upstream Pnuc_1738, formyltetrahydrofolate-dependent phosphoribosylglycinamide formyltransferase |
| m4C | 1816646 | - | | Pnuc_1745, sodium:dicarboxylate symporter |
| m4C | 1822325 | - | | upstream Pnuc_1749, ATP-dependent Clp protease adaptor protein ClpS |
| m6A | 1823635 | + | | upstream Pnuc_1755, conserved hypothetical protein |
| m6A | 1832050 | - | | Pnuc_1761, phosphoribosylformylglycinamidine cyclo-ligase |
| m6A | 1832665 | + | CTAYNNNNNNNNTRTC | Pnuc_1762, protein of unknown function UPF0118 |
| m6A | 1832675 | - | GAYANNNNNNNNRTAG | upstream Pnuc_1761, phosphoribosylformylglycinamidine cyclo-ligase |
| m4C | 1840381 | + | | upstream Pnuc_1773, NAD(+) kinase |
| m4C | 1840741 | + | | upstream Pnuc_1773, NAD(+) kinase |
| m6A | 1843285 | + | GAYANNNNNNNNRTAG | upstream Pnuc_1773, NAD(+) kinase |
| m6A | 1843295 | - | CTAYNNNNNNNNTRTC | Pnuc_1772, heat-inducible transcription repressor HrcA |
| m4C | 1845196 | - | | upstream Pnuc_1772, heat-inducible transcription repressor HrcA |
| m4C | 1846682 | - | | Pnuc_1775, (Glutamate-ammonia-ligase) adenylyltransferase |
| m6A | 1850932 | - | | upstream Pnuc_1775, (Glutamate-ammonia-ligase) adenylyltransferase |
| m6A | 1853226 | + | CTAYNNNNNNNNTRTC | Pnuc_1777, Nitrilase/cyanide hydratase and apolipoprotein N-acyltransferase |
| m6A | 1853236 | - | GAYANNNNNNNNRTAG | upstream Pnuc_1775, (Glutamate-ammonia-ligase) adenylyltransferase |
| m4C | 1861820 | + | | Pnuc_1784, alanine racemase domain protein |
| m6A | 1861824 | + | GAYANNNNNNNNRTAG | Pnuc_1784, alanine racemase domain protein |
| m6A | 1861834 | - | CTAYNNNNNNNNTRTC | upstream Pnuc_1780, ATP:cob(I)alamin adenosyltransferase |
| m4C | 1863314 | - | | Pnuc_1786, 4-hydroxybenzoate octaprenyltransferase |
| m6A | 1865372 | + | GAYANNNNNNNNRTAG | upstream Pnuc_1788, S-adenosylmethionine-tRNA-ribosyltransferase-isomerase |
| m6A | 1865382 | - | CTAYNNNNNNNNTRTC | Pnuc_1787, ATP-dependent DNA helicase RecG |
| m6A | 1866093 | + | | Pnuc_1788, S-adenosylmethionine-tRNA-ribosyltransferase-isomerase |
| m6A | 1867542 | + | CTAYNNNNNNNNTRTC | Pnuc_1789, tRNA-guanine transglycosylase |
| m6A | 1867552 | - | GAYANNNNNNNNRTAG | upstream Pnuc_1787, ATP-dependent DNA helicase RecG |
| m4C | 1872268 | + | | upstream Pnuc_1795, Glutathione S-transferase, N-terminal domain protein |
| m4C | 1873321 | + | | upstream Pnuc_1795, Glutathione S-transferase, N-terminal domain protein |
| m6A | 1874872 | + | GAYANNNNNNNNRTAG | upstream Pnuc_1797, alanine dehydrogenase/PNT domain protein |
| m6A | 1874879 | + | | upstream Pnuc_1797, alanine dehydrogenase/PNT domain protein |
| m6A | 1874882 | - | CTAYNNNNNNNNTRTC | Pnuc_1796, tRNA (5-methylaminomethyl-2-thiouridylate)-methyltransferase |
| m4C | 1879220 | + | | Pnuc_1800, hypothetical protein |
| m6A | 1880808 | + | GAYANNNNNNNNRTAG | upstream Pnuc_1802, hypothetical protein |

| | | | | |
|---|---|---|---|---|
| m6A | 1880818 | - | CTAYNNNNNNNNTRTC | Pnuc_1801, glycosyl transferase, family 11 |
| m6A | 1881762 | + | CTAYNNNNNNNNTRTC | Pnuc_1802, hypothetical protein |
| m6A | 1881772 | - | GAYANNNNNNNNRTAG | upstream Pnuc_1801, glycosyl transferase, family 11 |
| m6A | 1885043 | - | | Pnuc_1805, chaperonin GroEL |
| m4C | 1886632 | + | | Pnuc_1808, two component transcriptional regulator, LuxR family |
| m6A | 1887429 | - | | upstream Pnuc_1806, chaperonin Cpn10 |
| m6A | 1894804 | + | GAYANNNNNNNNRTAG | upstream Pnuc_1818, branched chain amino acid aminotransferase apoenzyme |
| m6A | 1894814 | - | CTAYNNNNNNNNTRTC | Pnuc_1814, phosphoribosylaminoimidazole-succinocarboxamide synthase |
| m4C | 1895256 | - | | Pnuc_1815, fructose-bisphosphate aldolase |
| m4C | 1895901 | + | | upstream Pnuc_1818, branched chain amino acid aminotransferase apoenzyme |
| m4C | 1897764 | + | | upstream Pnuc_1818, branched chain amino acid aminotransferase apoenzyme |
| m6A | 1899165 | + | CTAYNNNNNNNNTRTC | Pnuc_1818, branched chain amino acid aminotransferase apoenzyme |
| m6A | 1899175 | - | GAYANNNNNNNNRTAG | upstream Pnuc_1817, phosphoglycerate kinase |
| m4C | 1906490 | - | | upstream Pnuc_1825, TPR repeat-containing protein |
| m4C | 1910853 | - | | upstream Pnuc_1829, Integral membrane protein TerC |
| m6A | 1911071 | - | | Pnuc_1830, succinyl-CoA synthetase (ADP-forming) alpha subunit |
| m6A | 1916434 | + | GAYANNNNNNNNRTAG | Pnuc_1835, hypothetical protein |
| m6A | 1916444 | - | CTAYNNNNNNNNTRTC | upstream Pnuc_1834, conserved hypothetical protein |
| m6A | 1918050 | + | GAYANNNNNNNNRTAG | Pnuc_1838, hypothetical protein |
| m6A | 1921973 | + | GAYANNNNNNNNRTAG | upstream Pnuc_1847, secreted protein |
| m6A | 1921983 | - | CTAYNNNNNNNNTRTC | upstream Pnuc_1843, hypothetical protein |
| m4C | 1924413 | + | | upstream Pnuc_1847, secreted protein |
| m6A | 1931364 | + | | Pnuc_1858, protein of unknown function DUF1504 |
| m6A | 1934625 | - | | upstream Pnuc_1857, Rhodanese domain protein |
| m6A | 1935097 | + | GAYANNNNNNNNRTAG | Pnuc_1861, CBS domain containing protein |
| m6A | 1935107 | - | CTAYNNNNNNNNTRTC | upstream Pnuc_1857, Rhodanese domain protein |
| m6A | 1940032 | + | CTAYNNNNNNNNTRTC | Pnuc_1865, D-alpha,beta-D-heptose 1,7-bisphosphate phosphatase |
| m6A | 1940042 | - | GAYANNNNNNNNRTAG | upstream Pnuc_1857, Rhodanese domain protein |
| m4C | 1946837 | - | | Pnuc_1870, Organic solvent tolerance protein |
| m4C | 1948095 | - | | upstream Pnuc_1870, Organic solvent tolerance protein |
| m4C | 1954595 | + | | Pnuc_1878, crossover junction endodeoxyribonuclease RuvC |
| m4C | 1961924 | - | | upstream Pnuc_1886, LSU ribosomal protein L13P |
| m4C | 1964457 | + | | upstream Pnuc_1900, L-threonine ammonia-lyase |
| m4C | 1965110 | + | | upstream Pnuc_1900, L-threonine ammonia-lyase |
| m6A | 1968003 | + | CTAYNNNNNNNNTRTC | upstream Pnuc_1900, L-threonine ammonia-lyase |
| m6A | 1968013 | - | GAYANNNNNNNNRTAG | Pnuc_1892, aspartyl-tRNA synthetase |
| m6A | 1968185 | + | GAYANNNNNNNNRTAG | upstream Pnuc_1900, L-threonine ammonia-lyase |
| m6A | 1968195 | - | CTAYNNNNNNNNTRTC | Pnuc_1892, aspartyl-tRNA synthetase |
| m6A | 1970858 | + | | upstream Pnuc_1900, L-threonine ammonia-lyase |
| m6A | 1972754 | + | | upstream Pnuc_1900, L-threonine ammonia-lyase |
| m4C | 1974527 | - | | Pnuc_1899, FAD linked oxidase domain protein |
| m6A | 1974979 | + | GAYANNNNNNNNRTAG | upstream Pnuc_1900, L-threonine ammonia-lyase |
| m6A | 1974989 | - | CTAYNNNNNNNNTRTC | Pnuc_1899, FAD linked oxidase domain protein |
| m4C | 1978655 | - | | upstream Pnuc_1899, FAD linked oxidase domain protein |

| | | | | |
|---|---|---|---|---|
| m4C | 1978846 | - | | upstream Pnuc_1899, FAD linked oxidase domain protein |
| m4C | 1981099 | + | | Pnuc_1902, GTP cyclohydrolase I |
| m4C | 1981571 | + | | Pnuc_1902, GTP cyclohydrolase I |
| m4C | 1982756 | + | | upstream Pnuc_1905, Excinuclease ABC subunit A |
| m6A | 1984673 | - | | upstream Pnuc_1904, major facilitator superfamily MFS_1 |
| m4C | 1990207 | - | | upstream Pnuc_1906, Kef-type potassium/proton antiporter, CPA2 family |
| m4C | 1994211 | - | | upstream Pnuc_1906, Kef-type potassium/proton antiporter, CPA2 family |
| m4C | 1996322 | - | | Pnuc_1917, DNA-(apurinic or apyrimidinic site) lyase |
| m4C | 1999383 | + | | Pnuc_1920, ribose-phosphate pyrophosphokinase |
| m4C | 1999599 | + | | Pnuc_1920, ribose-phosphate pyrophosphokinase |
| m6A | 1999753 | + | GAYANNNNNNNRTAG | Pnuc_1920, ribose-phosphate pyrophosphokinase |
| m6A | 1999763 | - | CTAYNNNNNNNTRTC | upstream Pnuc_1917, DNA-(apurinic or apyrimidinic site) lyase |
| m6A | 2000577 | - | | upstream Pnuc_1917, DNA-(apurinic or apyrimidinic site) lyase |
| m4C | 2006549 | - | | upstream Pnuc_1927, peptidase M16 domain protein |
| m6A | 2008414 | + | GAYANNNNNNNRTAG | upstream Pnuc_1935, conserved hypothetical protein |
| m6A | 2008424 | - | CTAYNNNNNNNTRTC | Pnuc_1931, protoheme IX farnesyltransferase |
| m4C | 2009132 | + | | upstream Pnuc_1935, conserved hypothetical protein |
| m6A | 2014371 | + | CTAYNNNNNNNTRTC | upstream Pnuc_1942, phosphoribosyltransferase |
| m6A | 2014381 | - | GAYANNNNNNNRTAG | Pnuc_1939, Cytochrome-c oxidase |
| m6A | 2014641 | + | GAYANNNNNNNRTAG | upstream Pnuc_1942, phosphoribosyltransferase |
| m6A | 2014651 | - | CTAYNNNNNNNTRTC | Pnuc_1939, Cytochrome-c oxidase |
| m4C | 2016549 | - | | Pnuc_1940, cytochrome c oxidase, subunit II |
| m6A | 2024520 | + | | upstream Pnuc_1959, protein of unknown function DUF526 |
| m4C | 2026859 | + | | upstream Pnuc_1959, protein of unknown function DUF526 |
| m4C | 2033529 | + | | upstream Pnuc_1967, biotin--acetyl-CoA-carboxylase ligase |
| m4C | 2041650 | - | | upstream Pnuc_1969, rfaE bifunctional protein |
| m4C | 2041656 | + | | Pnuc_1972, protein of unknown function DUF1365 |
| m4C | 2047910 | + | | upstream Pnuc_1981, pseudo |
| m4C | 2050418 | + | | upstream Pnuc_1981, pseudo |
| m6A | 2054211 | - | GAYANNNNNNNRTAG | upstream Pnuc_1984, Alcohol dehydrogenase, zinc-binding domain protein |
| m4C | 2055912 | + | | upstream Pnuc_1996, protein of unknown function DUF185 |
| m6A | 2063777 | + | CTAYNNNNNNNTRTC | upstream Pnuc_1996, protein of unknown function DUF185 |
| m6A | 2063787 | - | GAYANNNNNNNRTAG | Pnuc_1992, UDP-N-acetylglucosamine pyrophosphorylase / glucosamine-1-phosphate N-acetyltransferase |
| m4C | 2066268 | + | | upstream Pnuc_1996, protein of unknown function DUF185 |
| m6A | 2066312 | + | GAYANNNNNNNRTAG | upstream Pnuc_1996, protein of unknown function DUF185 |
| m6A | 2066322 | - | CTAYNNNNNNNTRTC | Pnuc_1994, dihydroneopterin aldolase |
| m4C | 2075843 | + | | upstream Pnuc_2004, lipid A biosynthesis acyltransferase |
| m4C | 2075980 | - | | Pnuc_2003, methionine adenosyltransferase |
| m4C | 2083827 | + | | upstream Pnuc_2021, rod shape-determining protein MreB |
| m6A | 2085972 | + | GAYANNNNNNNRTAG | upstream Pnuc_2021, rod shape-determining protein MreB |
| m6A | 2085982 | - | CTAYNNNNNNNTRTC | Pnuc_2014, transcriptional regulator, TraR/DksA family |
| m6A | 2089363 | - | | Pnuc_2018, aspartyl/glutamyl-tRNA(Asn/Gln) amidotransferase subunit B |
| m4C | 2092386 | + | | upstream Pnuc_2021, rod shape-determining protein MreB |
| m4C | 2093112 | - | | upstream Pnuc_2020, aspartyl/glutamyl-tRNA(Asn/Gln) amidotransferase subunit C |

| | | | | |
|---|---|---|---|---|
| m6A | 2102026 | + | CTAYNNNNNNNNTRTC | Pnuc_2031, lipid A biosynthesis acyltransferase |
| m6A | 2102036 | - | GAYANNNNNNNNRTAG | upstream Pnuc_2029, protein of unknown function DUF167 |
| m6A | 2104275 | - | | Pnuc_2034, sodium:dicarboxylate symporter |
| m6A | 2118474 | + | GAYANNNNNNNNRTAG | upstream Pnuc_2056, protein of unknown function DUF897 |
| m6A | 2118484 | - | CTAYNNNNNNNNTRTC | Pnuc_2049, hypothetical protein |
| m6A | 2133954 | - | CTAYNNNNNNNNTRTC | upstream Pnuc_2064, hypothetical protein |
| m6A | 2146256 | + | CTAYNNNNNNNNTRTC | upstream Pnuc_2080, DNA protecting protein DprA |
| m6A | 2146266 | - | GAYANNNNNNNNRTAG | Pnuc_2077, sun protein |
| m4C | 2148177 | + | | upstream Pnuc_2080, DNA protecting protein DprA |
| m4C | 2150602 | - | | upstream Pnuc_2079, peptide deformylase |
| m6A | 2151830 | - | | upstream Pnuc_2079, peptide deformylase |
| m6A | 2158779 | + | | upstream Pnuc_0001, chromosomal replication initiator protein DnaA |

**Supplementary Table S6.** Enlistment of $C^5$-methyl-cytosine (m5C) and $N^6$-methyl-adenine (m6A) DNA modification sites in the genome of *P. asymbioticus* str. QLW-P1DMWA-$1^T$, as revealed by Single Molecule, Real-Time (SMRT) sequencing.

**Supplementary Table S7. (a)**

| | Open Reading Frames | | 500-bp Upstream | |
|---|---|---|---|---|
| | UV m6A (%) | UV m4C (%) | UV m6A (%) | UV m4C (%) |
| Upregulated | 12 (17) | 8 (15) | 0 (0) | 2 (25) |
| Downregulated | 59 (83) | 44 (85) | 16 (100) | 6 (75) |

**Supplementary Table S7. (b)**

| | Open Reading Frames | | 500-bp Upstream | |
|---|---|---|---|---|
| | 4ºC m6A (%) | 4ºC m4C (%) | 4ºC m6A (%) | 4ºC m4C (%) |
| Upregulated | 10 (11) | 39 (19) | 11 (29) | 2 (25) |
| Downregulated | 82 (89) | 164 (81) | 27 (71) | 6 (75) |

**Supplementary Table S7.** Table showing counts of adenine and cytosine methylation within differentially expressed genes (and percentage of expressed genes in parenthesis) of: **(a**). UV plus 26ºC (26ºC*) treated sample and **(b)**. 4ºC incubated sample, within gene body and 500-bp upstream sequence.

**Supplementary Table S8.**

**4ºC incubated sample versus 26ºC sample**

| Gene id | Microarray (fold expression) | Proteome (fold expression) | Methylome 4°C | Methylome 26°C | upstream methylation (4ºC) |
|---|---|---|---|---|---|
| Pnuc_0008 | 3.69 | 0.08 | m6A | | |
| Pnuc_1750 | 3.66 | 0.80 | m4C | | m4C (222bp) |
| Pnuc_2021 | 2.69 | 1.26 | m4C | | |
| Pnuc_1676 | 2.64 | 1.40 | m6A | | |
| Pnuc_1494 | 2.44 | 0.23 | | m4C | |
| Pnuc_1362 | 1.80 | 0.59 | m6A | m6A | |
| Pnuc_0134 | 1.40 | 0.64 | | m4C | |
| Pnuc_1804 | 0.84 | 0.22 | m6A | | |
| Pnuc_1403 | 0.77 | 1.06 | | m4C | m4C (22bp) |
| Pnuc_0604 | 0.76 | 0.78 | m6A | | |
| Pnuc_1483 | 0.60 | 0.15 | m6A | | |
| Pnuc_0714 | 0.54 | 0.78 | | m4C | |
| Pnuc_0171 | 0.49 | 0.46 | m6A | | |
| Pnuc_0100 | 0.42 | 0.95 | | m4C | |
| Pnuc_1624 | 0.39 | 0.52 | | m4C | |
| Pnuc_1479 | -0.28 | -0.51 | | | m4C (320bp) |
| Pnuc_1261 | -0.29 | -4.99 | m6A | | m6A (88bp) |
| Pnuc_1726 | -0.30 | -1.34 | m4C | | |
| Pnuc_0434 | -0.31 | -0.47 | m6A | m6A | |
| Pnuc_0470 | -0.34 | -1.03 | m6A | m6A | m6A (234bp) |
| Pnuc_0641 | -0.36 | -0.74 | m6A | m6A | |
| Pnuc_1774 | -0.36 | -0.73 | m6A | | |
| Pnuc_0689 | -0.38 | -0.62 | | | m6A (304bp) |

| | | | | | |
|---|---|---|---|---|---|
| Pnuc_0527 | -0.39 | -1.47 | | | m6A (406bp) |
| Pnuc_1777 | -0.39 | -2.60 | m6A | m6A | |
| Pnuc_0699 | -0.40 | -1.12 | | | m4C (277bp) |
| Pnuc_1531 | -0.41 | -2.34 | m6A | m6A | |
| Pnuc_1382 | -0.42 | -0.75 | | | m6A (201bp) |
| Pnuc_0353 | -0.42 | -1.75 | m6A | m6A | |
| Pnuc_0384 | -0.44 | -0.73 | | | m6A (16bp) |
| Pnuc_0899 | -0.44 | -2.41 | m4C, m6A | | |
| Pnuc_1306 | -0.47 | -2.04 | | m4C | |
| Pnuc_0991 | -0.49 | -0.77 | | m4C | |
| Pnuc_1256 | -0.50 | -0.40 | | m6A | |
| Pnuc_0447 | -0.51 | -1.28 | m6A, m6A | m4C, m6A, m6A, m4C, m6A | |
| Pnuc_1246 | -0.51 | -2.36 | m6A | m6A, m6A | |
| Pnuc_2060 | -0.52 | -0.59 | m6A, m4C | m6A | |
| Pnuc_1205 | -0.53 | -3.09 | m6A, m4C, m4C | m6A | |
| Pnuc_0796 | -0.54 | -0.52 | | m4C | |
| Pnuc_0227 | -0.55 | -0.29 | | m4C | |
| Pnuc_0141 | -0.55 | -0.58 | | m4C | |
| Pnuc_1180 | -0.56 | -0.46 | m6A, m6A | m6A, m6A | |
| Pnuc_0997 | -0.56 | -0.42 | | | m4C (6bp) |
| Pnuc_1357 | -0.56 | -1.53 | | m6A | |
| Pnuc_1105 | -0.57 | -2.52 | | m6A | |
| Pnuc_0509 | -0.58 | -0.54 | m4C | | |
| Pnuc_1711 | -0.58 | -0.12 | | m4C | |
| Pnuc_0763 | -0.59 | -0.30 | | m6A, m4A | |
| Pnuc_0220 | -0.59 | -0.76 | | m4C, m4C | m4C (395bp) |
| Pnuc_0489 | -0.59 | -0.48 | m4C | | |
| Pnuc_1206 | -0.59 | -2.17 | | m6A | |
| Pnuc_1799 | -0.59 | -1.41 | m4C | | |
| Pnuc_1103 | -0.60 | -1.46 | m6A | | |

| | | | | | |
|---|---|---|---|---|---|
| Pnuc_0258 | -0.60 | -1.38 | | m4C, m4C | |
| Pnuc_0535 | -0.61 | -1.99 | | m4C | |
| Pnuc_0193 | -0.64 | -1.31 | | m6A | |
| Pnuc_0570 | -0.64 | -1.23 | m6A | m6A | |
| Pnuc_0451 | -0.65 | -1.04 | m6A | | |
| Pnuc_1111 | -0.65 | -3.08 | m6A, m6A | m6A, m6A, m4C | |
| Pnuc_1787 | -0.69 | -1.59 | m6A, m4C | m6A | |
| Pnuc_1026 | -0.71 | -1.65 | m6A, m6A | m6A | |
| Pnuc_1012 | -0.75 | -1.46 | | m4C | |
| Pnuc_1084 | -0.75 | -0.48 | m6A | | |
| Pnuc_1784 | -0.81 | -1.47 | m6A | m6A | |
| Pnuc_1407 | -0.81 | -2.22 | m4C | m4C | |
| Pnuc_1148 | -0.83 | -0.27 | | | |
| Pnuc_0958 | -0.83 | -0.37 | m6A, m6A | m6A | |
| Pnuc_1284 | -0.84 | -1.51 | | m4C | |
| Pnuc_2022 | -0.85 | -3.52 | m4C, m4C | | |
| Pnuc_0335 | -0.87 | -2.99 | | m4C | |
| Pnuc_0889 | -0.87 | -0.19 | | m6A, m4C | |
| Pnuc_1163 | -0.88 | -0.73 | | m4C | |
| Pnuc_1300 | -0.91 | -2.40 | m4C, m4C | | |
| Pnuc_0436 | -0.92 | -1.73 | | m4C, m4C | |
| Pnuc_0251 | -0.95 | -1.68 | m4C | m4C | m6A (55bp) |
| Pnuc_0842 | -1.13 | -0.89 | m6A | m6A | |
| Pnuc_0431 | -1.27 | -0.88 | | m6A | |
| Pnuc_1873 | -1.28 | -0.45 | | m4C | |
| Pnuc_0672 | -1.30 | -0.55 | | m4C | |
| Pnuc_1150 | -1.40 | -0.53 | m6A | m6A | m4C (384bp) |
| Pnuc_0775 | -1.42 | -0.66 | | m4C | |
| Pnuc_0690 | -1.47 | -3.13 | m6A | m4C | m4C (70bp) |
| Pnuc_0426 | -1.53 | -0.63 | m6A | | |

| | | | | |
|---|---|---|---|---|
| Pnuc_0747 | -1.64 | -2.72 | m6A | m6A |
| Pnuc_1510 | -1.68 | -0.60 | | |
| Pnuc_0668 | -1.69 | -0.58 | | m4C |

**UV treated sample comparison against 26ºC sample**

| Gene id | Microarray (fold expression) | Proteome (fold expression) | Methylome UV | Methylome 26°C | upstream methylation (UV) |
|---|---|---|---|---|---|
| Pnuc_1805 | 2.39 | 2.80 | m6A | m4C | |
| Pnuc_1769 | 2.12 | 2.20 | | m4C | |
| Pnuc_2021 | 1.17 | 0.85 | | | m4C (285bp) |
| Pnuc_1564 | 0.49 | 3.36 | m6A | m6A | |
| Pnuc_1332 | 0.19 | 3.85 | m6A | m6A | |
| Pnuc_1504 | 0.16 | 0.73 | | m4C | |
| Pnuc_1234 | -0.20 | -0.53 | | m6A | |
| Pnuc_0735 | -0.23 | -0.89 | m4C | | |
| Pnuc_0990 | -0.33 | -2.50 | | | m4C (211bp), m6A (220bp) |
| Pnuc_0913 | -0.34 | -1.14 | m6A | m6A | |
| Pnuc_0570 | -0.36 | -1.32 | m6A | m6A | |
| Pnuc_0775 | -0.40 | -1.24 | | m4C | |
| Pnuc_0488 | -0.47 | -1.90 | m4C | | |
| Pnuc_0842 | -0.51 | -0.97 | m6A | m6A | |
| Pnuc_0737 | -0.52 | -0.43 | | | m6A (446bp) |
| Pnuc_0335 | -0.53 | -0.98 | | m4C | |
| Pnuc_1357 | -0.55 | -0.47 | | m6A | |
| Pnuc_1012 | -0.57 | -2.59 | | m4C | |
| Pnuc_0923 | -0.62 | -0.50 | m4C | m6A | |
| Pnuc_0092 | -0.70 | -0.05 | | m6A | |

| | | | | |
|---|---|---|---|---|
| Pnuc_0400 | -0.82 | -0.15 | | m4C |
| Pnuc_0958 | -0.98 | -0.30 | m6A | m6A |
| Pnuc_0668 | -1.06 | -0.23 | | m4C |
| Pnuc_0227 | -1.19 | -0.06 | | m4C |
| Pnuc_1150 | -1.22 | -0.34 | m6A | m6A |
| Pnuc_0526 | -1.61 | -0.39 | m6A | m6A |

**Supplementary Table S8.** Co-occurence of gene methylation, gene expression and corresponding protein detection.

### Supplementary Table S9.

| Bacteria | Genome (bp) | DNA methylation (all types detected) | Methylated genome (%) | References |
|---|---|---|---|---|
| *Helicobacter pylori* UM032 | 1593537 | 63299 | 3.97 | Lee, W. C., Anton, B. P., Wang, S., Baybayan, P., Singh, S., Ashby, M., et al. (2015). The complete methylome of *Helicobacter pylori* UM032. *BMC Genomics*. 16,424. |
| Helicobacter pylori J99-R3 | 1643831 | 57345 | 3.49 | Krebes, J., Morgan, R. D., Bunk, B., Spröer, C., Luong, K., Parusel, R., et al. (2013). The complex methylome of the human gastric pathogen *Helicobacter pylori*. *Nucleic Acids Res*. 42,1-18. |
| *Helicobacter pylori* 26695 | 1667867 | 50533 | 3.02 | Krebes, J., Morgan, R. D., Bunk, B., Spröer, C., Luong, K., Parusel, R., et al. (2013). The complex methylome of the human gastric *pathogen Helicobacter pylori*. *Nucleic Acids Res*. 42,1-18. |
| *Campylobacter jejuni* NCTC11168 | 1641481 | 32863 | 2.00 | Murray, I. A., Clark, T. A., Morgan, R. D., Boitano, M., Anton, B. P., Luong, K., et al. (2012). The methylomes of six bacteria. *Nucleic Acids Res*. 40,11450-11462. |
| *Campylobacter jejuni* 81-176 | 1699052 | 32023 | 1.88 | Murray, I. A., Clark, T. A., Morgan, R. D., Boitano, M., Anton, B. P., Luong, K., et al. (2012). The methylomes of six bacteria. *Nucleic Acids Res*. 40,11450-11462. |
| *Escherichia coli* O25b:K100:H4-ST131 EC958 | 5249449 | 52081 | 0.99 | Forde, B. M., Phan, M. D., Gawthorne, J. A., Ashcroft, M. M., Stanton-Cook, M., Sarkar, S., et al. (2015). Lineage-specific methyltransferases define the methylome of the globally disseminated *Escherichia coli* ST131 clone. *mBio*. 6:e01602-15. |

| | | | | |
|---|---|---|---|---|
| *Escherichia coli* O104:H4 C227-11 | 5542971 | 51972 | 0.94 | Fang, G., Munera, D., Friedman, D. I., Mandlik, A., Chao, M. C., Banerjee, O, et al. (2012). Genome-wide mapping of methylated adenine residues in pathogenic *Escherichia coli* using single-molecule real-time sequencing. *Nat. Biotechnol*. 30,1232-1239. |
| *Mycoplasma pneumoniae* M129 | 816394 | 6660 | 0.81 | Lluch-Senar, M., Luong, K., Lloréns-Rico, V., Delgado, J., Fang, G., Spittle, K., et al. (2013). Comprehensive methylome *characterization of Mycoplasma genitalium* and *Mycoplasma pneumoniae* at single-base resolution. *PLoS Genet*. 9,e1003191. |
| *Mycoplasma genitalium* G-37 | 580076 | 4568 | 0.79 | Lluch-Senar, M., Luong, K., Lloréns-Rico, V., Delgado, J., Fang, G., Spittle, K., et al. (2013). Comprehensive methylome *characterization of Mycoplasma genitalium* and *Mycoplasma pneumoniae* at single-base resolution. *PLoS Genet*. 9,e1003191. |
| *Vibrio breoganii* 1C-10 | 4116728 | 30020 | 0.72 | Murray, I. A., Clark, T. A., Morgan, R. D., Boitano, M., Anton, B. P., Luong, K., et al. (2012). The methylomes of six bacteria. *Nucleic Acids Res*. 40,11450-11462. |
| *Geobacter metallireducens* GS-15 | 4011182 | 29166 | 0.72 | Murray, I. A., Clark, T. A., Morgan, R. D., Boitano, M., Anton, B. P., Luong, K., et al. (2012). The methylomes of six bacteria. *Nucleic Acids Res*. 40,11450-11462. |
| *Caulobacter crescentus* | 4016947 | 13471 - 25927 | 0.33 - 0.64 | Kozdon, J. B., Melfi, M.D., Luong, K., Clark, T. A., Boitano, M., Wang, S, et al. (2013). Global methylation state at base-pair resolution *of the Caulobacter* genome throughout the cell cycle. *Proc. Natl. Acad. Sci. USA*. 110,E4658. |
| *Bacillus cereus* ATCC 10987 | 5432652 | 10583 | 0.19 | Murray, I. A., Clark, T. A., Morgan, R. D., Boitano, M., Anton, B. P., Luong, K., et al. (2012). The methylomes of six bacteria. *Nucleic* |

| | | | | |
|---|---|---|---|---|
| | | | | *Acids Res*. 40,11450-11462. |
| *Chromohalobacter salexigens* | 3696649 | 5541 | 0.15 | Murray, I. A., Clark, T. A., Morgan, R. D., Boitano, M., Anton, B. P., Luong, K., et al. (2012). The methylomes of six bacteria. *Nucleic Acids Res*. 40,11450-11462. |
| *Polynucleobacter asymbioticus* QLW-P1DMWA-1$^{T}$ | 2159490 | 873 - 1504 | 0.04 - 0.07 | This study. |

**Supplementary Table S9.** Methylation percentage in selected bacterial genome.

### Supplementary Table S10.

### Selected genes in Beta proteobacterium CB

| Sequence | Method | Ka | Ks | Ka/Ks | P-Value(Fisher) | Length | S-Sites | N-Sites | Substitutions | S-Substitutions | N-Substitutions |
|---|---|---|---|---|---|---|---|---|---|---|---|
| Alkyl | MS | 0.0490 | 2.3656 | 0.0207 | 4.64E-65 | 420 | 99.9824 | 320.018 | 89 | 83.4638 | 5.53621 |
| Alkyl | GMYN | 0.0526 | 2.1569 | 0.0244 | 3.92E-41 | 420 | 103.032 | 316.968 | 89 | 72.9183 | 16.0817 |
| Carbo | MS | 0.0513 | 4.1831 | 0.0123 | 6.84E-123 | 615 | 141.208 | 473.792 | 141 | 135.426 | 5.57402 |
| Carbo | GMYN | 0.0554 | 3.0948 | 0.0179 | 4.18E-65 | 615 | 158.128 | 456.872 | 141 | 116.682 | 24.3185 |
| Cold | MS | 0.0000 | 0.1975 | 0.0000 | 0 | 156 | 45.9512 | 110.049 | 7 | 6.99998 | 1.68E-05 |
| Cold | GMYN | 0.0000 | 0.2277 | 0.0000 | 0 | 156 | 38.2496 | 117.75 | 7 | 7 | 0 |
| DnaK | MS | 0.0142 | 0.8334 | 0.0170 | 3.87E-139 | 1878 | 517.35 | 1360.65 | 248 | 237.374 | 10.626 |
| DnaK | GMYN | 0.0133 | 1.5135 | 0.0088 | 5.94E-149 | 1878 | 423.783 | 1454.22 | 248 | 228.79 | 19.2105 |
| DSBA | MS | 0.2138 | 3.7619 | 0.0568 | NA | 594 | 131.622 | 462.378 | 197 | 164.216 | 32.7843 |
| DSBA | GMYN | 0.2140 | 3.6227 | 0.0591 | 3.91E-49 | 594 | 125.244 | 468.756 | 197 | 110.641 | 86.3591 |
| Fatty | MS | 0.0441 | 1.2511 | 0.0353 | 7.41E-116 | 1143 | 298.721 | 844.279 | 212 | 192.785 | 19.2146 |
| Fatty | GMYN | 0.0439 | 1.2500 | 0.0352 | 4.39E-91 | 1143 | 288.675 | 854.325 | 212 | 175.616 | 36.3844 |
| Fimb | MS | 0.0537 | 3.7249 | 0.0144 | 3.32E-86 | 465 | 119.856 | 345.144 | 112 | 107.533 | 4.46737 |
| Fimb | GMYN | 0.0545 | 3.5996 | 0.0151 | 4.43E-54 | 465 | 121.442 | 343.558 | 112 | 93.9862 | 18.0138 |
| FtsK | MS | 0.0298 | 1.2058 | 0.0247 | 5.23E-226 | 2256 | 594.393 | 1661.61 | 392 | 366.665 | 25.3354 |
| FtsK | GMYN | 0.0298 | 2.2884 | 0.0130 | 1.48E-191 | 2256 | 569.943 | 1686.06 | 392 | 342.76 | 49.2405 |
| GroL | MS | NA | NA | NA | NA | NA | NA | NA | NA | NA | NA |
| GroL | GMYN | 0.0239 | 1.3477 | 0.0177 | 5.70E-89 | 1596 | 357.271 | 1238.73 | 191 | 161.964 | 29.0361 |
| Hfq | MS | 0.0505 | 1.1646 | 0.0434 | 6.17E-15 | 195 | 67.7019 | 127.298 | 33 | 30.511 | 2.48904 |
| Hfq | GMYN | 0.0442 | 1.1378 | 0.0388 | 1.42E-13 | 195 | 51.3775 | 143.623 | 33 | 26.9232 | 6.07677 |
| Hyp | MS | 0.0429 | 1.5402 | 0.0279 | 2.74E-22 | 219 | 69.9217 | 149.078 | 42 | 39.6458 | 2.35418 |
| Hyp | GMYN | 0.0398 | 1.0712 | 0.0371 | 4.83E-18 | 219 | 63.1353 | 155.865 | 42 | 36 | 6 |
| MscS | MS | 0.1362 | 3.6868 | 0.0369 | NA | 768 | 186.848 | 581.152 | 228 | 204.509 | 23.4909 |
| MscS | GMYN | 0.1339 | 3.9083 | 0.0343 | 1.80E-79 | 768 | 183.276 | 584.724 | 228 | 157.031 | 70.9689 |
| Piri | MS | 0.1708 | 3.6202 | 0.0472 | NA | 684 | 164.432 | 519.568 | 208 | 181.022 | 26.9781 |
| Piri | GMYN | 0.1563 | 3.7993 | 0.0411 | 3.33E-63 | 684 | 158.501 | 525.499 | 208 | 134.636 | 73.3645 |
| RpoE | MS | 0.0421 | 0.8878 | 0.0475 | 9.16E-42 | 522 | 130.159 | 391.841 | 83 | 72.6232 | 10.3768 |
| RpoE | GMYN | 0.0403 | 1.5532 | 0.0260 | 3.76E-34 | 522 | 129.844 | 392.156 | 83 | 67.632 | 15.368 |
| RpoH | MS | 0.0576 | 1.8488 | 0.0311 | 5.55E-108 | 888 | 261.831 | 626.169 | 196 | 182.414 | 13.586 |

| | | | | | | | | | | | |
|---|---|---|---|---|---|---|---|---|---|---|---|
| RpoH | GMYN | 0.0530 | 3.3685 | 0.0157 | 1.12E-92 | 888 | 218.22 | 669.78 | 196 | 161.831 | 34.1686 |
| Rubre | MS | 0.0714 | 3.2578 | 0.0219 | 7.09E-66 | 438 | 127.645 | 310.355 | 105 | 99.6893 | 5.3107 |
| Rubre | GMYN | 0.0676 | 2.1995 | 0.0307 | 6.65E-43 | 438 | 118.953 | 319.047 | 105 | 84.4634 | 20.5366 |
| RuvB | MS | 0.0216 | 1.7691 | 0.0122 | 7.10E-134 | 1014 | 262.888 | 751.112 | 195 | 188.427 | 6.57318 |
| RuvB | GMYN | 0.0207 | 4.0690 | 0.0051 | 3.83E-132 | 1014 | 227.079 | 786.921 | 195 | 178.926 | 16.0736 |
| | | | **maximum** | 0.0591 | | | | | | | |
| | | | **minimum** | 0.0000 | | | | | | | |
| | | | **average** | 0.0263 | | | | | | | |
| | | | **median** | 0.0247 | | | | | | | |

### Housekeeping genes in Beta proteobacterium CB

| Sequence | Method | Ka | Ks | Ka/Ks | P-Value(Fisher) | Length | S-Sites | N-Sites | Substitutions | S-Substitutions | N-Substitutions |
|---|---|---|---|---|---|---|---|---|---|---|---|
| iso | MS | 0.0367 | 1.0320 | 0.0355 | 7.66E-192 | 2220 | 555.089 | 1664.91 | 358 | 323.532 | 34.4682 |
| iso | GMYN | 0.0364 | 2.4546 | 0.0148 | 4.93E-164 | 2220 | 511.193 | 1708.81 | 358 | 297.458 | 60.5418 |
| acsA | MS | 0.0191 | 0.7978 | 0.0240 | 7.18E-143 | 1959 | 537.515 | 1421.49 | 267 | 251.083 | 15.9166 |
| acsA | GMYN | 0.0182 | 1.1909 | 0.0153 | 7.15E-145 | 1959 | 455.441 | 1503.56 | 267 | 239.983 | 27.0171 |
| succ | MS | 0.0213 | 1.1644 | 0.0183 | 1.66E-166 | 1764 | 411.916 | 1352.08 | 259 | 244.355 | 14.6446 |
| succ | GMYN | 0.0204 | 1.6253 | 0.0126 | 2.26E-165 | 1764 | 347.102 | 1416.9 | 259 | 230.516 | 28.4839 |
| CTP | MS | 0.0431 | 2.5545 | 0.0169 | 4.83E-258 | 1647 | 388.169 | 1258.83 | 348 | 329.961 | 18.0385 |
| CTP | GMYN | 0.0446 | 4.4242 | 0.0101 | 1.94E-193 | 1647 | 364.641 | 1282.36 | 348 | 292.589 | 55.4106 |
| atpA | MS | 0.0058 | 0.5249 | 0.0110 | 3.44E-87 | 1527 | 408.415 | 1118.59 | 150 | 145.618 | 4.3816 |
| atpA | GMYN | 0.0054 | 0.5244 | 0.0103 | 9.82E-82 | 1527 | 415.773 | 1111.23 | 150 | 143.998 | 6.00155 |
| gyr | MS | 0.0421 | 2.8067 | 0.0150 | 1.76E-219 | 1422 | 370.516 | 1051.48 | 312 | 299.273 | 12.7274 |
| gyr | GMYN | 0.0433 | 2.0001 | 0.0216 | 9.49E-147 | 1422 | 384.668 | 1037.33 | 312 | 268.459 | 43.5413 |
| glnA | MS | 0.0262 | 0.6366 | 0.0412 | 7.71E-80 | 1401 | 327.913 | 1073.09 | 159 | 140.12 | 18.8796 |
| glnA | GMYN | 0.0257 | 1.3264 | 0.0193 | 8.04E-75 | 1401 | 284.568 | 1116.43 | 159 | 130.875 | 28.1248 |
| cit | MS | 0.0248 | 1.0818 | 0.0229 | 1.70E-120 | 1299 | 314.94 | 984.06 | 199 | 185.712 | 13.2882 |
| cit | GMYN | 0.0235 | 1.7439 | 0.0135 | 6.66E-123 | 1299 | 258.548 | 1040.45 | 199 | 174.954 | 24.0462 |
| adk | MS | 0.0394 | 0.8747 | 0.0450 | 1.55E-47 | 651 | 167.084 | 483.916 | 95 | 84.0386 | 10.9614 |
| adk | GMYN | 0.0356 | 1.0140 | 0.0351 | 3.30E-39 | 651 | 158.644 | 492.356 | 95 | 77.933 | 17.067 |
| ndk | MS | 0.0349 | 0.6510 | 0.0537 | 7.66E-23 | 411 | 78.9238 | 332.076 | 43 | 35.0774 | 7.92264 |
| ndk | GMYN | 0.0382 | 0.4817 | 0.0794 | 1.12E-13 | 411 | 98.3222 | 312.678 | 43 | 31.3974 | 11.6026 |
| | | | **maximum** | 0.0794 | | | | | | | |

| | |
|---|---|
| **minimum** | 0.0101 |
| **average** | 0.0258 |
| **median** | 0.0188 |

**Selected genes in FNE-F8 bin 6 1 PnecC**

| Sequence | Method | Ka | Ks | Ka/Ks | P-Value(Fisher) | Length | S-Sites | N-Sites | Substitutions | S-Substitutions | N-Substitutions |
|---|---|---|---|---|---|---|---|---|---|---|---|
| FtsK | MS | 0.0252 | 1.5749 | 0.0160 | 1.24E-259 | 2205 | 596.985 | 1608.01 | 416 | 398.838 | 17.1616 |
| FtsK | GMYN | 0.0248 | 1.3630 | 0.0182 | 7.24E-214 | 2205 | 599.942 | 1605.06 | 416 | 376.857 | 39.1426 |
| DnaK | MS | 0.0185 | 1.0103 | 0.0183 | 4.45E-141 | 1830 | 550.86 | 1279.14 | 267 | 256.134 | 10.8657 |
| DnaK | GMYN | 0.0168 | 1.9225 | 0.0087 | 2.32E-155 | 1830 | 423.986 | 1406.01 | 267 | 243.718 | 23.2819 |
| GroL | MS | 0.0241 | 0.8124 | 0.0296 | 2.24E-102 | 1551 | 417.608 | 1133.39 | 199 | 184.183 | 14.817 |
| GroL | GMYN | 0.0239 | 0.7832 | 0.0305 | 2.93E-85 | 1551 | 401.001 | 1150 | 199 | 171.99 | 27.0099 |
| Fatty | MS | 0.0511 | 1.6921 | 0.0302 | 2.84E-146 | 1101 | 260.493 | 840.507 | 227 | 206.861 | 20.1391 |
| Fatty | GMYN | 0.0510 | 2.4188 | 0.0211 | 1.44E-107 | 1101 | 257.707 | 843.293 | 227 | 185.597 | 41.4029 |
| RuvB | MS | 0.0248 | 2.0379 | 0.0122 | 7.82E-130 | 978 | 256.851 | 721.149 | 191 | 184.694 | 6.30582 |
| RuvB | GMYN | 0.0213 | 1.8020 | 0.0118 | 1.39E-109 | 978 | 257.85 | 720.15 | 191 | 175.891 | 15.1094 |
| RpoH | MS | 0.0443 | 2.2379 | 0.0198 | 1.43E-101 | 849 | 248.834 | 600.166 | 172 | 164.159 | 7.84083 |
| RpoH | GMYN | 0.0440 | 1.7936 | 0.0245 | 4.01E-81 | 849 | 212.649 | 636.351 | 172 | 144.894 | 27.1059 |
| Piri | MS | 0.1562 | 3.5478 | 0.0440 | NA | 615 | 153.004 | 461.996 | 188 | 165.936 | 22.0642 |
| Piri | GMYN | 0.1554 | 3.6912 | 0.0421 | 7.69E-62 | 615 | 137.219 | 477.781 | 188 | 121.503 | 66.4971 |
| Carbo | MS | 0.0490 | 2.3829 | 0.0206 | 7.60E-86 | 579 | 137.615 | 441.385 | 120 | 112.579 | 7.42133 |
| Carbo | GMYN | 0.0530 | 1.5878 | 0.0334 | 2.63E-52 | 579 | 148.626 | 430.374 | 120 | 98.0499 | 21.9501 |
| DSBA | MS | 0.2287 | 3.5739 | 0.0640 | NA | 543 | 125.194 | 417.806 | 197 | 162.327 | 34.6731 |
| DSBA | GMYN | 0.2308 | 3.5685 | 0.0647 | 4.05E-59 | 543 | 116.507 | 426.493 | 197 | 113.199 | 83.8006 |
| RpoE | MS | 0.0373 | 0.9192 | 0.0406 | 6.41E-39 | 489 | 120.406 | 368.594 | 74 | 65.8281 | 8.1719 |
| RpoE | GMYN | 0.0408 | 0.6439 | 0.0634 | 2.68E-22 | 489 | 157.596 | 331.404 | 74 | 60.869 | 13.131 |
| Rubre | MS | 0.0796 | 3.1231 | 0.0255 | 1.44E-52 | 405 | 122.483 | 282.517 | 92 | 86.8934 | 5.1066 |
| Rubre | GMYN | 0.0745 | 1.6575 | 0.0449 | 1.76E-33 | 405 | 106.147 | 298.853 | 92 | 70.8764 | 21.1236 |
| Alkyl | MS | 0.0477 | 2.8312 | 0.0169 | 1.18E-67 | 387 | 80.7862 | 306.214 | 79 | 74.2535 | 4.74646 |
| Alkyl | GMYN | 0.0556 | 1.5720 | 0.0353 | 1.44E-30 | 387 | 105.061 | 281.939 | 79 | 63.94 | 15.06 |
| Hyp | MS | 0.1081 | 1.7632 | 0.0613 | 3.83E-30 | 258 | 63.4016 | 194.598 | 56 | 47.1288 | 8.87121 |
| Hyp | GMYN | 0.1130 | 1.3727 | 0.0823 | 6.17E-13 | 258 | 67.9994 | 190.001 | 56 | 36.1982 | 19.8018 |

| | | | | | | | | | | | |
|---|---|---|---|---|---|---|---|---|---|---|---|
| Hfq | MS | 0.0162 | 4.1595 | 0.0039 | 0 | 165 | 39.1784 | 125.822 | 35 | 34.568 | 0.431958 |
| Hfq | GMYN | 0.0166 | 2.8099 | 0.0059 | 7.18E-24 | 165 | 42.3758 | 122.624 | 35 | 33 | 2 |
| Cold | MS | 0.0587 | 0.3859 | 0.1520 | 0.000282898 | 126 | 33.8772 | 92.1228 | 14 | 9.90516 | 4.09484 |
| Cold | GMYN | 0.0555 | 0.3986 | 0.1392 | 0.000979928 | 126 | 31.0745 | 94.9255 | 14 | 9 | 5 |
| | | | **maximum** | 0.1520 | | | | | | | |
| | | | **minimum** | 0.0039 | | | | | | | |
| | | | **average** | 0.0394 | | | | | | | |
| | | | **median** | 0.0299 | | | | | | | |

### Housekeeping genes in FNE-F8 bin 6 1 PneccC

| Sequence | Method | Ka | Ks | Ka/Ks | P-Value(Fisher) | Length | S-Sites | N-Sites | Substitutions | S-Substitutions | N-Substitutions |
|---|---|---|---|---|---|---|---|---|---|---|---|
| iso | MS | 0.0377 | 1.1919 | 0.0317 | 4.93E-205 | 2229 | 613.557 | 1615.44 | 390 | 359.982 | 30.0182 |
| iso | GMYN | 0.0367 | 1.1967 | 0.0306 | 2.46E-178 | 2229 | 552.028 | 1676.97 | 390 | 330.066 | 59.9338 |
| acsA | MS | 0.0451 | 0.9224 | 0.0489 | 1.22E-147 | 1968 | 505.133 | 1462.87 | 312 | 273.294 | 38.7059 |
| acsA | GMYN | 0.0450 | 1.1273 | 0.0399 | 2.22E-115 | 1968 | 483.851 | 1484.15 | 312 | 247.275 | 64.7248 |
| succ | MS | 0.0244 | 1.4442 | 0.0169 | 1.68E-197 | 1773 | 415.09 | 1357.91 | 294 | 278.629 | 15.3715 |
| succ | GMYN | 0.0231 | 4.3822 | 0.0053 | 2.19E-198 | 1773 | 344.8 | 1428.2 | 294 | 261.573 | 32.4274 |
| CTP | MS | 0.0445 | 2.0753 | 0.0215 | 2.13E-220 | 1656 | 426.505 | 1229.5 | 340 | 320.192 | 19.8082 |
| CTP | GMYN | 0.0430 | 4.4527 | 0.0097 | 6.11E-180 | 1656 | 378.766 | 1277.23 | 340 | 286.693 | 53.3066 |
| atpA | MS | 0.0109 | 0.5337 | 0.0205 | 3.59E-84 | 1536 | 431.827 | 1104.17 | 163 | 154.883 | 8.11725 |
| atpA | GMYN | 0.0103 | 0.8107 | 0.0128 | 2.82E-90 | 1536 | 366.785 | 1169.21 | 163 | 150.998 | 12.0025 |
| gyr | MS | 0.0394 | 2.9300 | 0.0135 | 8.63E-225 | 1431 | 361.43 | 1069.57 | 308 | 296.203 | 11.797 |
| gyr | GMYN | 0.0392 | 4.3704 | 0.0090 | 5.64E-173 | 1431 | 339.397 | 1091.6 | 308 | 266.397 | 41.6028 |
| glnA | MS | 0.0212 | 0.5916 | 0.0358 | 1.08E-79 | 1413 | 366.88 | 1046.12 | 161 | 146.086 | 14.9141 |
| glnA | GMYN | 0.0212 | 0.6623 | 0.0321 | 1.63E-69 | 1413 | 355.65 | 1057.35 | 161 | 138.89 | 22.1102 |
| cit | MS | 0.0226 | 1.3813 | 0.0163 | 2.13E-148 | 1308 | 317.063 | 990.937 | 222 | 211.209 | 10.7907 |
| cit | GMYN | 0.0227 | 1.4395 | 0.0158 | 6.41E-126 | 1308 | 312.118 | 995.882 | 222 | 199.746 | 22.2536 |
| adk | MS | 0.0582 | 1.3849 | 0.0421 | 6.22E-68 | 660 | 172.253 | 487.747 | 127 | 113.485 | 13.5149 |
| adk | GMYN | 0.0551 | 1.8739 | 0.0294 | 5.14E-57 | 660 | 144.849 | 515.151 | 127 | 99.7069 | 27.2931 |
| ndk | MS | 0.0321 | 0.7148 | 0.0448 | 1.13E-25 | 420 | 101.298 | 318.702 | 51 | 44.6942 | 6.30577 |
| ndk | GMYN | 0.0330 | 0.6499 | 0.0508 | 1.03E-19 | 420 | 102.274 | 317.726 | 51 | 40.795 | 10.205 |
| | | | **maximum** | 0.0508 | | | | | | | |
| | | | **minimum** | 0.0053 | | | | | | | |

| | |
|---|---|
| **average** | 0.0264 |
| **median** | 0.0254 |

### Selected genes in MWH-Adler-W8

| Sequence | Method | Ka | Ks | Ka/Ks | P-Value(Fisher) | Length | S-Sites | N-Sites | Substitutions | S-Substitutions | N-Substitutions |
|---|---|---|---|---|---|---|---|---|---|---|---|
| FtsK | MS | 0.0134 | 0.9306 | 0.0144 | 1.17E-188 | 2232 | 655.371 | 1576.63 | 339 | 327.62 | 11.3798 |
| FtsK | GMYN | 0.0133 | 1.0223 | 0.0130 | 1.03E-174 | 2232 | 635.419 | 1596.58 | 339 | 317.958 | 21.042 |
| DnaK | MS | 0.0084 | 0.5584 | 0.0150 | 5.98E-105 | 1857 | 534.311 | 1322.69 | 195 | 188.012 | 6.98776 |
| DnaK | GMYN | 0.0079 | 0.7183 | 0.0110 | 7.00E-114 | 1857 | 443.221 | 1413.78 | 195 | 183.936 | 11.0639 |
| GroL | MS | 0.0038 | 0.4759 | 0.0080 | 2.62E-82 | 1575 | 433.088 | 1141.91 | 140 | 137.114 | 2.88573 |
| GroL | GMYN | 0.0033 | 0.7345 | 0.0045 | 3.09E-89 | 1575 | 369.8 | 1205.2 | 140 | 136 | 4.00009 |
| Fatty | MS | 0.0600 | 1.7005 | 0.0353 | 1.43E-146 | 1125 | 255.687 | 869.313 | 230 | 205.347 | 24.6531 |
| Fatty | GMYN | 0.0594 | 4.1670 | 0.0143 | 1.01E-106 | 1125 | 240.517 | 884.483 | 230 | 179.691 | 50.3093 |
| RuvB | MS | 0.0154 | 1.5575 | 0.0099 | 1.30E-124 | 996 | 268.221 | 727.779 | 185 | 180.16 | 4.84025 |
| RuvB | GMYN | 0.0154 | 1.4935 | 0.0103 | 1.32E-108 | 996 | 268.642 | 727.358 | 185 | 173.976 | 11.0241 |
| RpoH | MS | 0.0165 | 1.4136 | 0.0117 | 2.64E-88 | 855 | 237.894 | 617.106 | 143 | 138.795 | 4.20463 |
| RpoH | GMYN | 0.0158 | 1.3085 | 0.0121 | 1.04E-84 | 855 | 214.857 | 640.143 | 143 | 132.991 | 10.0092 |
| Piri | MS | 0.1356 | 3.3205 | 0.0408 | NA | 633 | 135.082 | 497.918 | 171 | 148.624 | 22.3756 |
| Piri | GMYN | 0.1362 | 3.6944 | 0.0369 | 1.26E-51 | 633 | 137.803 | 495.197 | 171 | 109.728 | 61.2724 |
| Carbo | MS | 0.0362 | 2.6543 | 0.0137 | 3.55E-88 | 603 | 141.123 | 461.877 | 116 | 111.038 | 4.9616 |
| Carbo | GMYN | 0.0387 | 1.9678 | 0.0197 | 2.17E-60 | 603 | 141.779 | 461.221 | 116 | 98.6223 | 17.3777 |
| DSBA | MS | 0.1338 | 3.8000 | 0.0352 | NA | 564 | 133.261 | 430.739 | 155 | 139.167 | 15.8333 |
| DSBA | GMYN | 0.1354 | 3.6545 | 0.0370 | 7.52E-46 | 564 | 130.663 | 433.337 | 155 | 101.764 | 53.2363 |
| RpoE | MS | 0.0153 | 0.8914 | 0.0172 | 4.23E-44 | 507 | 115.697 | 391.303 | 66 | 62.3697 | 3.63032 |
| RpoE | GMYN | 0.0148 | 2.0241 | 0.0073 | 7.32E-45 | 507 | 97.0133 | 409.987 | 66 | 59.9993 | 6.00071 |
| Superox | MS | 0.0731 | 1.6862 | 0.0433 | 9.78E-61 | 471 | 104.752 | 366.248 | 99 | 85.9714 | 13.0286 |
| Superox | GMYN | 0.0771 | 1.6481 | 0.0468 | 8.99E-32 | 471 | 123.061 | 347.939 | 99 | 73.6828 | 25.3172 |
| Fimb | MS | 0.0323 | 3.8914 | 0.0083 | 5.03E-88 | 465 | 116.601 | 348.399 | 107 | 104.409 | 2.5913 |
| Fimb | GMYN | 0.0335 | 3.5954 | 0.0093 | 1.69E-62 | 465 | 120.768 | 344.232 | 107 | 95.9016 | 11.0984 |
| Rubre | MS | 0.0245 | 2.5196 | 0.0097 | 5.37E-53 | 483 | 146.595 | 336.405 | 86 | 84.1211 | 1.87894 |
| Rubre | GMYN | 0.0244 | 1.1113 | 0.0219 | 1.15E-38 | 483 | 148.491 | 334.509 | 86 | 77.9986 | 8.00139 |
| Alkyl | MS | 0.0366 | 1.6574 | 0.0221 | 7.85E-52 | 405 | 96.7634 | 308.237 | 75 | 70.0759 | 4.92413 |

| | | | | | | | | | | | |
|---|---|---|---|---|---|---|---|---|---|---|---|
| Alkyl | GMYN | 0.0349 | 1.2734 | 0.0274 | 4.78E-36 | 405 | 105.767 | 299.233 | 75 | 64.8017 | 10.1983 |
| Hyp | MS | 0.0425 | 1.8575 | 0.0229 | 1.78E-19 | 204 | 53.2076 | 150.792 | 30 | 28.1741 | 1.82585 |
| Hyp | GMYN | 0.0384 | 1.4289 | 0.0269 | 1.25E-14 | 204 | 42.1986 | 161.801 | 30 | 24 | 6 |
| Hfq | MS | 0.0000 | 0.9622 | 0.0000 | 0 | 189 | 54.6278 | 134.372 | 24 | 23.9999 | 5.90E-05 |
| Hfq | GMYN | 0.0000 | 1.0078 | 0.0000 | 0 | 189 | 43.2947 | 145.705 | 24 | 24 | 0 |
| Cold | MS | 0.0000 | 0.0619 | 0.0000 | 0 | 141 | 32.2973 | 108.703 | 2 | 1.99999 | 6.73E-06 |
| Cold | GMYN | 0.0000 | 0.0629 | 0.0000 | 0 | 141 | 35.7646 | 105.235 | 2 | 2 | 0 |
| | | | **maximum** | 0.0468 | | | | | | | |
| | | | **minimum** | 0.0000 | | | | | | | |
| | | | **average** | 0.0178 | | | | | | | |
| | | | **median** | 0.0140 | | | | | | | |

### Housekeeping genes in MWH-Adler-W8

| Sequence | Method | Ka | Ks | Ka/Ks | P-Value(Fisher) | Length | S-Sites | N-Sites | Substitutions | S-Substitutions | N-Substitutions |
|---|---|---|---|---|---|---|---|---|---|---|---|
| iso | MS | 0.0241 | 0.9105 | 0.0265 | 6.78E-177 | 2232 | 583.67 | 1648.33 | 324 | 301.438 | 22.5624 |
| iso | GMYN | 0.0245 | 1.4186 | 0.0172 | 2.73E-144 | 2232 | 595.329 | 1636.67 | 324 | 284.646 | 39.3536 |
| acsA | MS | 0.0271 | 0.7413 | 0.0365 | 1.82E-132 | 1971 | 537.997 | 1433 | 274 | 249.709 | 24.2911 |
| acsA | GMYN | 0.0253 | 1.5192 | 0.0167 | 9.39E-140 | 1971 | 433.419 | 1537.58 | 274 | 235.81 | 38.1899 |
| succ | MS | 0.0121 | 0.7966 | 0.0152 | 2.38E-123 | 1776 | 527.008 | 1248.99 | 232 | 223.912 | 8.08822 |
| succ | GMYN | 0.0113 | 1.4411 | 0.0078 | 5.91E-136 | 1776 | 427.859 | 1348.14 | 232 | 216.947 | 15.0531 |
| CTP | MS | 0.0441 | 1.6630 | 0.0265 | 5.29E-200 | 1659 | 406.643 | 1252.36 | 318 | 294.014 | 23.9861 |
| CTP | GMYN | 0.0409 | 4.3859 | 0.0093 | 3.89E-177 | 1659 | 346.475 | 1312.52 | 318 | 265.895 | 52.1048 |
| atpA | MS | 0.0027 | 0.2263 | 0.0120 | 4.25E-44 | 1539 | 438.488 | 1100.51 | 83 | 80.5698 | 2.43025 |
| atpA | GMYN | 0.0026 | 0.2655 | 0.0098 | 1.55E-49 | 1539 | 381.204 | 1157.8 | 83 | 80 | 3 |
| pyr | MS | 0.0363 | 3.6677 | 0.0099 | 5.11E-210 | 1434 | 380.539 | 1053.46 | 297 | 289.072 | 7.92841 |
| pyr | GMYN | 0.0365 | 2.0799 | 0.0175 | 1.84E-152 | 1434 | 368.689 | 1065.31 | 297 | 259.245 | 37.7552 |
| glnA | MS | 0.0079 | 0.3491 | 0.0225 | 2.74E-54 | 1413 | 404.929 | 1008.07 | 109 | 103.207 | 5.79339 |
| glnA | GMYN | 0.0077 | 0.4163 | 0.0184 | 6.23E-55 | 1413 | 360.822 | 1052.18 | 109 | 100.997 | 8.00264 |
| cit | MS | 0.0071 | 0.6569 | 0.0108 | 2.40E-89 | 1311 | 330.451 | 980.549 | 144 | 139.534 | 4.46591 |
| cit | GMYN | 0.0072 | 0.6880 | 0.0105 | 2.31E-82 | 1311 | 333.787 | 977.213 | 144 | 136.999 | 7.00117 |
| adk | MS | 0.0250 | 0.7580 | 0.0330 | 1.77E-41 | 663 | 208.577 | 454.423 | 91 | 84.9041 | 6.09593 |
| adk | GMYN | 0.0230 | 0.9005 | 0.0256 | 2.82E-41 | 663 | 175.583 | 487.417 | 91 | 79.9576 | 11.0424 |
| ndk | MS | 0.0030 | 0.4310 | 0.0070 | 0 | 423 | 98.4718 | 324.528 | 29 | 28.3486 | 0.651395 |

| | | | | | | | | | | | |
|---|---|---|---|---|---|---|---|---|---|---|---|
| ndk | GMYN | 0.0031 | 0.3955 | 0.0078 | 7.47E-19 | 423 | 96.2147 | 326.785 | 29 | 28 | 1 |
| | | | **maximum** | 0.0365 | | | | | | | |
| | | | **minimum** | 0.0070 | | | | | | | |
| | | | **average** | 0.0170 | | | | | | | |
| | | | **median** | 0.0159 | | | | | | | |

### Selected genes in MWH-HuW1

| Sequence | Method | Ka | Ks | Ka/Ks | P-Value(Fisher) | Length | S-Sites | N-Sites | Substitutions | S-Substitutions | N-Substitutions |
|---|---|---|---|---|---|---|---|---|---|---|---|
| Alkyl | MS | 0.0578 | 2.0329 | 0.0285 | 1.81E-60 | 432 | 103.051 | 328.949 | 90 | 82.5059 | 7.49409 |
| Alkyl | GMYN | 0.0616 | 1.7443 | 0.0353 | 1.49E-37 | 432 | 104.463 | 327.537 | 90 | 70.6916 | 19.3084 |
| Carbo | MS | 0.0658 | 4.1225 | 0.0160 | NA | 627 | 144.387 | 482.613 | 154 | 146.196 | 7.80388 |
| Carbo | GMYN | 0.0653 | 3.6205 | 0.0180 | 5.11E-98 | 627 | 124.875 | 502.125 | 154 | 122.929 | 31.0711 |
| Cold | MS | 0.0000 | 0.1302 | 0.0000 | 0 | 171 | 38.4118 | 132.588 | 5 | 4.99998 | 1.73E-05 |
| Cold | GMYN | 0.0000 | 0.1473 | 0.0000 | 0 | 171 | 38.8647 | 132.135 | 5 | 5 | 0 |
| DnaK | MS | 0.0176 | 1.0387 | 0.0169 | 4.08E-156 | 1890 | 548.263 | 1341.74 | 284 | 272.71 | 11.2896 |
| DnaK | GMYN | 0.0161 | 1.2408 | 0.0130 | 1.65E-172 | 1890 | 429.855 | 1460.14 | 284 | 260.749 | 23.2506 |
| DSBA | MS | 0.2284 | 3.7082 | 0.0616 | NA | 600 | 133.047 | 466.953 | 217 | 178.435 | 38.5649 |
| DSBA | GMYN | 0.2311 | 3.6198 | 0.0638 | 1.92E-67 | 600 | 124.76 | 475.24 | 217 | 123.477 | 93.5227 |
| Fatty | MS | 0.0488 | 1.7769 | 0.0275 | 1.86E-158 | 1155 | 283.951 | 871.049 | 243 | 224.103 | 18.8968 |
| Fatty | GMYN | 0.0500 | 1.7359 | 0.0288 | 1.15E-106 | 1155 | 298.506 | 856.494 | 243 | 201.757 | 41.2426 |
| Fimb | MS | 0.0922 | 3.6166 | 0.0255 | NA | 489 | 125.954 | 363.046 | 143 | 133.21 | 9.78959 |
| Fimb | GMYN | 0.0955 | 3.6445 | 0.0262 | 2.71E-60 | 489 | 128.943 | 360.057 | 143 | 110.897 | 32.1029 |
| FtsK | MS | 0.0308 | 1.6376 | 0.0188 | 7.34E-283 | 2265 | 600.328 | 1664.67 | 448 | 425.805 | 22.1948 |
| FtsK | GMYN | 0.0317 | 1.7850 | 0.0178 | 4.45E-232 | 2265 | 581.485 | 1683.51 | 448 | 395.749 | 52.2514 |
| GroL | MS | 0.0255 | 0.8289 | 0.0308 | 9.77E-113 | 1608 | 433.234 | 1174.77 | 219 | 202.11 | 16.8905 |
| GroL | GMYN | 0.0237 | 2.8105 | 0.0084 | 2.00E-110 | 1608 | 363.96 | 1244.04 | 219 | 189.962 | 29.0379 |
| Hfq | MS | 0.0500 | 1.4481 | 0.0345 | 2.03E-22 | 210 | 60.6699 | 149.33 | 41 | 37.7899 | 3.21014 |
| Hfq | GMYN | 0.0478 | 1.2818 | 0.0373 | 5.32E-18 | 210 | 55.2251 | 154.775 | 41 | 33.9205 | 7.0795 |
| Hyp | MS | 0.0839 | 3.2142 | 0.0261 | 2.06E-35 | 237 | 69.3616 | 167.638 | 57 | 53.6181 | 3.38187 |
| Hyp | GMYN | 0.0874 | 1.9998 | 0.0437 | 1.49E-16 | 237 | 74.8291 | 162.171 | 57 | 43.7518 | 13.2482 |
| MscS | MS | 0.1593 | 3.6796 | 0.0433 | NA | 783 | 186.984 | 596.016 | 252 | 221.435 | 30.5654 |
| MscS | GMYN | 0.1594 | 3.9277 | 0.0406 | 3.07E-82 | 783 | 188.102 | 594.898 | 252 | 167.729 | 84.271 |

| | | | | | | | | | | | |
|---|---|---|---|---|---|---|---|---|---|---|---|
| Piri | MS | 0.2023 | 3.8314 | 0.0528 | NA | 663 | 145.735 | 517.265 | 212 | 178.54 | 33.4597 |
| Piri | GMYN | 0.2025 | 3.7300 | 0.0543 | 1.38E-49 | 663 | 144.516 | 518.484 | 212 | 120.756 | 91.2439 |
| RpoE | MS | 0.0271 | 1.1903 | 0.0227 | 2.71E-52 | 534 | 117.046 | 416.954 | 78 | 72.1561 | 5.84387 |
| RpoE | GMYN | 0.0268 | 1.8250 | 0.0147 | 8.89E-43 | 534 | 112.498 | 421.502 | 78 | 66.9366 | 11.0634 |
| RpoH | MS | 0.0579 | 2.0835 | 0.0278 | 2.69E-122 | 897 | 255.006 | 641.994 | 206 | 192.529 | 13.4715 |
| RpoH | GMYN | 0.0556 | 2.5579 | 0.0217 | 4.37E-92 | 897 | 235.377 | 661.623 | 206 | 170.703 | 35.2973 |
| | | | **maximum** | 0.0638 | | | | | | | |
| | | | **minimum** | 0.0000 | | | | | | | |
| | | | **average** | 0.0285 | | | | | | | |
| | | | **median** | 0.0268 | | | | | | | |

### Housekeeping genes in MWH-HuW1

| Sequence | Method | Ka | Ks | Ka/Ks | P-Value(Fisher) | Length | S-Sites | N-Sites | Substitutions | S-Substitutions | N-Substitutions |
|---|---|---|---|---|---|---|---|---|---|---|---|
| iso | MS | 0.0307 | 1.1857 | 0.0259 | 7.20E-202 | 2232 | 621.318 | 1610.68 | 373 | 349.512 | 23.4885 |
| iso | GMYN | 0.0300 | 1.1779 | 0.0255 | 2.24E-183 | 2232 | 544.351 | 1687.65 | 373 | 323.368 | 49.6319 |
| acsA | MS | 0.0190 | 0.9509 | 0.0199 | 6.22E-165 | 1971 | 534.467 | 1436.53 | 291 | 276.202 | 14.7979 |
| acsA | GMYN | 0.0179 | 2.7781 | 0.0065 | 3.70E-171 | 1971 | 443.175 | 1527.82 | 291 | 263.979 | 27.0206 |
| succ | MS | 0.0244 | 1.1871 | 0.0206 | 1.84E-169 | 1776 | 457.36 | 1318.64 | 283 | 267.154 | 15.8464 |
| succ | GMYN | 0.0232 | 2.2408 | 0.0104 | 6.18E-182 | 1776 | 351.706 | 1424.29 | 283 | 250.484 | 32.5163 |
| CTP | MS | 0.0469 | 2.3755 | 0.0197 | 1.85E-244 | 1659 | 395.744 | 1263.26 | 345 | 324.552 | 20.4483 |
| CTP | GMYN | 0.0468 | 4.4215 | 0.0106 | 5.03E-186 | 1659 | 363.337 | 1295.66 | 345 | 286.354 | 58.6459 |
| atpA | MS | 0.0049 | 0.5582 | 0.0088 | 3.11E-94 | 1539 | 415.525 | 1123.47 | 158 | 154.339 | 3.66115 |
| atpA | GMYN | 0.0044 | 0.7333 | 0.0060 | 2.83E-94 | 1539 | 394.052 | 1144.95 | 158 | 152.999 | 5.00095 |
| gyr | MS | 0.0425 | 3.6752 | 0.0116 | 7.32E-232 | 1434 | 365.786 | 1068.21 | 314 | 303.75 | 10.2497 |
| gyr | GMYN | 0.0438 | 2.7416 | 0.0160 | 4.14E-155 | 1434 | 367.735 | 1066.27 | 314 | 268.672 | 45.3283 |
| glnA | MS | 0.0260 | 0.7578 | 0.0343 | 2.59E-97 | 1413 | 365.432 | 1047.57 | 189 | 172.082 | 16.9183 |
| glnA | GMYN | 0.0252 | 1.1563 | 0.0218 | 5.25E-93 | 1413 | 316.056 | 1096.94 | 189 | 161.881 | 27.1187 |
| cit | MS | 0.0187 | 0.8896 | 0.0210 | 6.26E-103 | 1311 | 344.629 | 966.371 | 182 | 171.865 | 10.1346 |
| cit | GMYN | 0.0172 | 1.5183 | 0.0113 | 1.66E-120 | 1311 | 251.927 | 1059.07 | 182 | 163.991 | 18.0088 |
| adk | MS | NA | NA | NA | NA | NA | NA | NA | NA | NA | NA |
| adk | GMYN | 0.0336 | 1.3159 | 0.0255 | 6.14E-54 | 663 | 141.762 | 521.238 | 105 | 87.9287 | 17.0713 |
| ndk | MS | 0.0121 | 0.5806 | 0.0209 | 6.22E-24 | 423 | 99.3507 | 323.649 | 39 | 36.5139 | 2.48615 |
| ndk | GMYN | 0.0125 | 0.5149 | 0.0243 | 1.76E-20 | 423 | 99.3462 | 323.654 | 39 | 35 | 4 |

| | |
|---|---|
| **maximum** | 0.0343 |
| **minimum** | 0.0060 |
| **average** | 0.0179 |
| **median** | 0.0199 |

### Selected genes in MWH-JaK3

| Sequence | Method | Ka | Ks | Ka/Ks | P-Value(Fisher) | Length | S-Sites | N-Sites | Substitutions | S-Substitutions | N-Substitutions |
|---|---|---|---|---|---|---|---|---|---|---|---|
| Alkyl | MS | 0.0498 | 2.0586 | 0.0242 | 1.22E-50 | 444 | 125.475 | 318.525 | 85 | 80.0813 | 4.91868 |
| Alkyl | GMYN | 0.0518 | 1.9869 | 0.0261 | 8.43E-33 | 444 | 119.039 | 324.961 | 85 | 68.7605 | 16.2395 |
| Carbo | MS | 0.0711 | 2.6137 | 0.0272 | 2.92E-105 | 639 | 145.466 | 493.534 | 144 | 131.831 | 12.1692 |
| Carbo | GMYN | 0.0644 | 3.7198 | 0.0173 | 2.17E-69 | 639 | 142.548 | 496.452 | 144 | 113.49 | 30.5099 |
| Cold | MS | 0.0000 | 0.0485 | 0.0000 | 0 | 180 | 41.2123 | 138.788 | 2 | 1.99999 | 6.74E-06 |
| Cold | GMYN | NA | 0.0506 | NA | 0 | 180 | 43.1023 | 136.898 | 2 | 2 | 0 |
| DnaK | MS | 0.0096 | 0.5981 | 0.0160 | 2.37E-111 | 1896 | 497.188 | 1398.81 | 197 | 188.51 | 8.48977 |
| DnaK | GMYN | 0.0091 | 0.7509 | 0.0121 | 2.72E-114 | 1896 | 439.163 | 1456.84 | 197 | 183.896 | 13.1045 |
| DSBA | MS | 0.1303 | 3.6993 | 0.0352 | NA | 600 | 146.215 | 453.785 | 163 | 146.94 | 16.0595 |
| DSBA | GMYN | 0.1316 | 3.7388 | 0.0352 | 2.41E-46 | 600 | 145.769 | 454.231 | 163 | 108.579 | 54.4209 |
| Fatty | MS | 0.0671 | 1.9638 | 0.0342 | 9.31E-170 | 1161 | 271.284 | 889.716 | 257 | 231.106 | 25.8945 |
| Fatty | GMYN | 0.0672 | 4.1854 | 0.0161 | 7.91E-112 | 1161 | 265.218 | 895.782 | 257 | 199.624 | 57.3765 |
| Fimb | MS | 0.0581 | 3.1576 | 0.0184 | 5.75E-82 | 504 | 129.606 | 374.394 | 114 | 108.243 | 5.75668 |
| Fimb | GMYN | 0.0580 | 3.6129 | 0.0160 | 3.61E-53 | 504 | 123.612 | 380.388 | 114 | 92.8365 | 21.1635 |
| FtsK | MS | NA | NA | NA | NA | NA | NA | NA | NA | NA | NA |
| FtsK | GMYN | 0.0184 | 0.8855 | 0.0208 | 1.53E-158 | 2271 | 671.97 | 1599.03 | 347 | 317.888 | 29.1116 |
| GroL | MS | 0.0043 | 0.4033 | 0.0107 | 1.37E-72 | 1614 | 461.508 | 1152.49 | 133 | 129.544 | 3.45647 |
| GroL | GMYN | 0.0041 | 0.5418 | 0.0075 | 1.55E-81 | 1614 | 378.782 | 1235.22 | 133 | 128 | 5.00025 |
| Hfq | MS | 0.0067 | 0.9608 | 0.0070 | 0 | 219 | 65.6215 | 153.379 | 32 | 31.4868 | 0.51322 |
| Hfq | GMYN | 0.0068 | 0.8991 | 0.0075 | 4.58E-17 | 219 | 70.2969 | 148.703 | 32 | 31 | 1 |
| Hyp | MS | 0.0620 | 1.2128 | 0.0511 | 5.61E-22 | 243 | 68.3067 | 174.693 | 43 | 38.0284 | 4.97162 |
| Hyp | GMYN | 0.0627 | 0.9219 | 0.0680 | 5.24E-13 | 243 | 70.2652 | 172.735 | 43 | 32.7181 | 10.2819 |
| MscS | MS | 0.1391 | 2.8398 | 0.0490 | 5.40E-147 | 789 | 183.979 | 605.021 | 212 | 182.586 | 29.4139 |
| MscS | GMYN | 0.1338 | 3.8727 | 0.0346 | 7.12E-64 | 789 | 174.787 | 614.213 | 212 | 137.427 | 74.5728 |
| Piri | MS | 0.1090 | 2.1400 | 0.0509 | 4.45E-100 | 669 | 150.938 | 518.062 | 160 | 136.19 | 23.8101 |

| | | | | | | | | | | | |
|---|---|---|---|---|---|---|---|---|---|---|---|
| Piri | GMYN | 0.1105 | 3.7016 | 0.0298 | 1.67E-52 | 669 | 139.138 | 529.862 | 160 | 105.835 | 54.1649 |
| RpoE | MS | 0.0243 | 0.9293 | 0.0262 | 2.06E-46 | 543 | 126.226 | 416.774 | 77 | 70.8697 | 6.13029 |
| RpoE | GMYN | 0.0224 | 1.1076 | 0.0202 | 2.31E-39 | 543 | 131.462 | 411.538 | 77 | 67.9305 | 9.06949 |
| RpoH | MS | 0.0176 | 1.1343 | 0.0155 | 6.01E-84 | 891 | 253.7 | 637.3 | 144 | 138.604 | 5.39645 |
| RpoH | GMYN | 0.0168 | 1.8747 | 0.0090 | 1.14E-81 | 891 | 226.384 | 664.616 | 144 | 132.978 | 11.0217 |
| RuvB | MS | 0.0195 | 1.5113 | 0.0129 | 1.17E-122 | 1032 | 274.545 | 757.455 | 188 | 181.547 | 6.45305 |
| RuvB | GMYN | 0.0176 | 1.3142 | 0.0134 | 2.34E-104 | 1032 | 282.23 | 749.77 | 188 | 174.974 | 13.0262 |
| | | | **maximum** | 0.0680 | | | | | | | |
| | | | **minimum** | 0.0000 | | | | | | | |
| | | | **average** | 0.0237 | | | | | | | |
| | | | **median** | 0.0193 | | | | | | | |

### Housekeeping genes in MWH-JaK3

| Sequence | Method | Ka | Ks | Ka/Ks | P-Value(Fisher) | Length | S-Sites | N-Sites | Substitutions | S-Substitutions | N-Substitutions |
|---|---|---|---|---|---|---|---|---|---|---|---|
| iso | MS | 0.0161 | 0.8209 | 0.0196 | 1.85E-170 | 2232 | 595.338 | 1636.66 | 303 | 287.544 | 15.4558 |
| iso | GMYN | 0.0154 | 1.8312 | 0.0084 | 3.51E-174 | 2232 | 513.728 | 1718.27 | 303 | 276.868 | 26.1316 |
| acsA | MS | 0.0259 | 0.7986 | 0.0324 | 1.53E-138 | 1971 | 559.195 | 1411.81 | 283 | 261.597 | 21.4034 |
| acsA | GMYN | 0.0240 | 1.8174 | 0.0132 | 1.06E-149 | 1971 | 437.394 | 1533.61 | 283 | 246.884 | 36.1162 |
| succ | MS | 0.0176 | 0.8302 | 0.0212 | 4.27E-135 | 1776 | 438.421 | 1337.58 | 230 | 216.015 | 13.9846 |
| succ | GMYN | 0.0161 | 1.7823 | 0.0090 | 7.29E-132 | 1776 | 393.031 | 1382.97 | 230 | 207.96 | 22.0404 |
| CTP | MS | 0.0383 | 1.4202 | 0.0270 | 1.53E-182 | 1659 | 419.317 | 1239.68 | 305 | 282.456 | 22.5437 |
| CTP | GMYN | 0.0379 | 1.4861 | 0.0255 | 3.88E-147 | 1659 | 399.829 | 1259.17 | 305 | 258.527 | 46.4732 |
| atpA | MS | 0.0019 | 0.1442 | 0.0129 | 2.04E-30 | 1539 | 464.41 | 1074.59 | 59 | 57.2907 | 1.70933 |
| atpA | GMYN | 0.0018 | 0.1646 | 0.0107 | 2.22E-32 | 1539 | 400.875 | 1138.13 | 59 | 57 | 2 |
| gyr | MS | 0.0394 | 3.4594 | 0.0114 | 1.43E-214 | 1434 | 402.787 | 1031.21 | 319 | 309.964 | 9.03628 |
| gyr | GMYN | 0.0402 | 2.2461 | 0.0179 | 1.53E-156 | 1434 | 390.631 | 1043.37 | 319 | 278.312 | 40.6879 |
| glnA | MS | 0.0104 | 0.4212 | 0.0248 | 1.15E-62 | 1413 | 361.456 | 1051.54 | 118 | 110.068 | 7.93202 |
| glnA | GMYN | 0.0108 | 0.3915 | 0.0275 | 5.94E-53 | 1413 | 382.426 | 1030.57 | 118 | 106.994 | 11.0057 |
| cit | MS | 0.0118 | 0.6805 | 0.0173 | 3.34E-86 | 1311 | 376.81 | 934.19 | 160 | 153.414 | 6.58645 |
| cit | GMYN | 0.0117 | 0.7571 | 0.0155 | 1.91E-79 | 1311 | 365.424 | 945.576 | 160 | 149 | 11 |
| adk | MS | 0.0231 | 1.0322 | 0.0224 | 1.05E-56 | 663 | 176.891 | 486.109 | 98 | 92.3184 | 5.68161 |
| adk | GMYN | 0.0217 | 2.3886 | 0.0091 | 2.01E-57 | 663 | 142.78 | 520.22 | 98 | 86.9152 | 11.0848 |
| ndk | MS | 0.0068 | 0.3892 | 0.0176 | 1.40E-18 | 423 | 108.122 | 314.878 | 31 | 29.4898 | 1.51023 |

| | | | | | | | | | | | |
|---|---|---|---|---|---|---|---|---|---|---|---|
| ndk | GMYN | 0.0063 | 0.3899 | 0.0160 | 9.94E-18 | 423 | 101.342 | 321.658 | 31 | 29 | 2 |
| | | | **maximum** | 0.0324 | | | | | | | |
| | | | **minimum** | 0.0084 | | | | | | | |
| | | | **average** | 0.0180 | | | | | | | |
| | | | **median** | 0.0175 | | | | | | | |

### Selected genes in MWH-MoK4

| Sequence | Method | Ka | Ks | Ka/Ks | P-Value(Fisher) | Length | S-Sites | N-Sites | Substitutions | S-Substitutions | N-Substitutions |
|---|---|---|---|---|---|---|---|---|---|---|---|
| Alkyl | MS | 0.0501 | 1.4584 | 0.0343 | 1.09E-49 | 435 | 105.063 | 329.937 | 81 | 73.1168 | 7.88318 |
| Alkyl | GMYN | 0.0501 | 1.6571 | 0.0302 | 1.16E-37 | 435 | 96.8239 | 338.176 | 81 | 64.647 | 16.353 |
| Carbo | MS | 0.0640 | 4.0729 | 0.0157 | 4.79E-129 | 630 | 147.092 | 482.908 | 151 | 143.591 | 7.40934 |
| Carbo | GMYN | 0.0695 | 2.4956 | 0.0279 | 1.39E-61 | 630 | 166.479 | 463.521 | 151 | 120.379 | 30.621 |
| Cold | MS | 0.0000 | 0.3117 | 0.0000 | 0 | 171 | 45.7305 | 125.269 | 10 | 9.99997 | 2.74E-05 |
| Cold | GMYN | NA | 0.3436 | NA | 0 | 171 | 39.0396 | 131.96 | 10 | 10 | 0 |
| DnaK | MS | 0.0191 | 0.8982 | 0.0213 | 1.60E-137 | 1893 | 554.357 | 1338.64 | 266 | 252.992 | 13.0085 |
| DnaK | GMYN | 0.0182 | 1.2457 | 0.0146 | 3.65E-135 | 1893 | 483.934 | 1409.07 | 266 | 240.74 | 25.2598 |
| DSBA | MS | 0.2109 | 3.8950 | 0.0541 | NA | 609 | 130.442 | 478.558 | 197 | 164.351 | 32.6492 |
| DSBA | GMYN | 0.2078 | 3.5746 | 0.0581 | 2.10E-54 | 609 | 117.458 | 491.542 | 197 | 108.721 | 88.2793 |
| Fatty | MS | 0.0546 | 1.6296 | 0.0335 | 4.31E-147 | 1161 | 280.18 | 880.82 | 238 | 215.329 | 22.6708 |
| Fatty | GMYN | 0.0510 | 1.8746 | 0.0272 | 9.80E-107 | 1161 | 282.956 | 878.044 | 238 | 194.787 | 43.2133 |
| Fimb | MS | 0.0550 | 3.6437 | 0.0151 | 3.75E-96 | 480 | 126.395 | 353.605 | 122 | 117.054 | 4.94559 |
| Fimb | GMYN | 0.0563 | 3.6370 | 0.0155 | 2.90E-60 | 480 | 127.652 | 352.348 | 122 | 102.96 | 19.0403 |
| FtsK | MS | NA | NA | NA | NA | NA | NA | NA | NA | NA | NA |
| FtsK | GMYN | 0.0330 | 1.2264 | 0.0269 | 1.94E-196 | 2271 | 620.571 | 1650.43 | 428 | 374.709 | 53.2915 |
| GroL | MS | 0.0315 | 0.6930 | 0.0454 | 1.64E-95 | 1614 | 436.828 | 1177.17 | 210 | 187.113 | 22.8874 |
| GroL | GMYN | 0.0295 | 0.8056 | 0.0366 | 4.83E-85 | 1614 | 398.948 | 1215.05 | 210 | 174.942 | 35.0579 |
| Hfq | MS | 0.0346 | 1.3306 | 0.0260 | 1.16E-21 | 210 | 58.9698 | 151.03 | 37 | 34.6928 | 2.30717 |
| Hfq | GMYN | 0.0331 | 1.2354 | 0.0268 | 9.32E-19 | 210 | 52.7496 | 157.25 | 37 | 31.943 | 5.057 |
| Piri | MS | 0.1525 | 3.7022 | 0.0412 | NA | 666 | 159.009 | 506.991 | 190 | 167.943 | 22.057 |
| Piri | GMYN | 0.1515 | 2.7019 | 0.0561 | 4.72E-47 | 666 | 167.49 | 498.51 | 190 | 122.194 | 67.8064 |
| RpoE | MS | 0.0346 | 0.8836 | 0.0391 | 3.37E-41 | 537 | 122.034 | 414.966 | 74 | 65.3127 | 8.68725 |
| RpoE | GMYN | 0.0343 | 0.7338 | 0.0467 | 6.69E-29 | 537 | 138.458 | 398.542 | 74 | 60.6677 | 13.3323 |

| | | | | | | | | | | | |
|---|---|---|---|---|---|---|---|---|---|---|---|
| RpoH | MS | 0.0654 | 1.7953 | 0.0364 | 2.41E-113 | 906 | 254.257 | 651.743 | 204 | 186.581 | 17.4195 |
| RpoH | GMYN | 0.0616 | 3.2919 | 0.0187 | 1.53E-90 | 906 | 220.911 | 685.089 | 204 | 163.627 | 40.3726 |
| RuvB | MS | 0.0226 | 2.6409 | 0.0085 | 3.67E-159 | 1032 | 265.99 | 766.01 | 216 | 210.81 | 5.18987 |
| RuvB | GMYN | 0.0218 | 4.0990 | 0.0053 | 8.15E-151 | 1032 | 236.349 | 795.651 | 216 | 198.91 | 17.0905 |
| Superox | MS | 0.1478 | 3.9317 | 0.0376 | NA | 501 | 112.829 | 388.171 | 147 | 130.161 | 16.8386 |
| Superox | GMYN | 0.1545 | 3.5797 | 0.0432 | 2.51E-40 | 501 | 118.272 | 382.728 | 147 | 93.8519 | 53.1481 |
| | | | **maximum** | 0.0581 | | | | | | | |
| | | | **minimum** | 0.0053 | | | | | | | |
| | | | **average** | 0.0332 | | | | | | | |
| | | | **median** | 0.0364 | | | | | | | |

### Housekeeping genes in MWH-MoK4

| Sequence | Method | Ka | Ks | Ka/Ks | P-Value(Fisher) | Length | S-Sites | N-Sites | Substitutions | S-Substitutions | N-Substitutions |
|---|---|---|---|---|---|---|---|---|---|---|---|
| iso | MS | 0.0449 | 1.3134 | 0.0342 | 7.77E-219 | 2232 | 594.613 | 1637.39 | 406 | 371.058 | 34.942 |
| iso | GMYN | 0.0436 | 1.3547 | 0.0322 | 1.17E-180 | 2232 | 533.201 | 1698.8 | 406 | 334.209 | 71.7912 |
| acsA | MS | 0.0332 | 0.8538 | 0.0389 | 3.33E-145 | 1971 | 529.54 | 1441.46 | 297 | 268.591 | 28.4093 |
| acsA | GMYN | 0.0316 | 1.4453 | 0.0218 | 9.23E-142 | 1971 | 442.811 | 1528.19 | 297 | 249.879 | 47.1209 |
| succ | MS | 0.0264 | 1.2285 | 0.0215 | 2.73E-172 | 1776 | 450.743 | 1325.26 | 288 | 270.908 | 17.0919 |
| succ | GMYN | 0.0252 | 1.7650 | 0.0143 | 1.62E-172 | 1776 | 373.114 | 1402.89 | 288 | 253.238 | 34.7617 |
| CTP | MS | 0.0452 | 2.1968 | 0.0206 | 1.82E-239 | 1659 | 389.105 | 1269.89 | 339 | 317.671 | 21.3295 |
| CTP | GMYN | 0.0455 | 4.3991 | 0.0103 | 4.97E-188 | 1659 | 352.653 | 1306.35 | 339 | 281.475 | 57.5253 |
| atpA | MS | 0.0073 | 0.5753 | 0.0127 | 1.77E-92 | 1539 | 437.524 | 1101.48 | 168 | 162.798 | 5.20194 |
| atpA | GMYN | 0.0074 | 0.5742 | 0.0129 | 2.82E-84 | 1539 | 449.593 | 1089.41 | 168 | 159.998 | 8.00241 |
| gyr | MS | 0.0440 | 2.8297 | 0.0156 | 1.63E-214 | 1434 | 397.871 | 1036.13 | 323 | 310.426 | 12.5738 |
| gyr | GMYN | 0.0432 | 3.3716 | 0.0128 | 2.55E-161 | 1434 | 375.774 | 1058.23 | 323 | 278.685 | 44.3148 |
| glnA | MS | 0.0234 | 0.5632 | 0.0415 | 9.55E-74 | 1413 | 379.737 | 1033.26 | 161 | 144.653 | 16.3467 |
| glnA | GMYN | 0.0223 | 0.9748 | 0.0228 | 1.26E-75 | 1413 | 314.461 | 1098.54 | 161 | 136.929 | 24.0706 |
| cit | MS | 0.0248 | 1.0765 | 0.0230 | 1.78E-104 | 1311 | 403.661 | 907.339 | 207 | 196.811 | 10.1895 |
| cit | GMYN | 0.0236 | 1.3596 | 0.0173 | 3.04E-96 | 1311 | 359.966 | 951.034 | 207 | 184.961 | 22.0394 |
| adk | MS | 0.0423 | 0.8510 | 0.0497 | 2.46E-46 | 663 | 175.563 | 487.437 | 100 | 87.8641 | 12.1359 |
| adk | GMYN | 0.0392 | 2.1557 | 0.0182 | 1.77E-47 | 663 | 133.984 | 529.016 | 100 | 79.8747 | 20.1253 |
| ndk | MS | 0.0271 | 0.6862 | 0.0394 | 9.61E-26 | 423 | 92.2277 | 330.772 | 46 | 40.3017 | 5.69831 |
| ndk | GMYN | 0.0263 | 0.9812 | 0.0268 | 2.48E-24 | 423 | 73.4068 | 349.593 | 46 | 36.9863 | 9.01365 |

| | |
|---|---|
| **maximum** | 0.0497 |
| **minimum** | 0.0103 |
| **average** | 0.0243 |
| **median** | 0.0217 |

### Selected genes in MWH-Recht1

| Sequence | Method | Ka | Ks | Ka/Ks | P-Value(Fisher) | Length | S-Sites | N-Sites | Substitutions | S-Substitutions | N-Substitutions |
|---|---|---|---|---|---|---|---|---|---|---|---|
| Methyl | MS | 0.0754 | 1.1880 | 0.0635 | 1.18E-128 | 1470 | 350.71 | 1119.29 | 276 | 229.493 | 46.5069 |
| Methyl | GMYN | 0.0697 | 2.3825 | 0.0292 | 7.38E-89 | 1470 | 348.153 | 1121.85 | 276 | 201.683 | 74.3171 |
| RpoE | MS | 0.0045 | 0.0336 | 0.1354 | 0.00299834 | 567 | 132.04 | 434.96 | 6 | 4.14927 | 1.85073 |
| RpoE | GMYN | 0.0047 | 0.0289 | 0.1636 | 0.0042709 | 567 | 142.236 | 424.764 | 6 | 4 | 2 |
| | | | **maximum** | 0.1636 | | | | | | | |
| | | | **minimum** | 0.0292 | | | | | | | |
| | | | **average** | 0.0979 | | | | | | | |
| | | | **median** | 0.0995 | | | | | | | |

### Housekeeping genes in MWH-Recht1

| Sequence | Method | Ka | Ks | Ka/Ks | P-Value(Fisher) | Length | S-Sites | N-Sites | Substitutions | S-Substitutions | N-Substitutions |
|---|---|---|---|---|---|---|---|---|---|---|---|
| iso | MS | 0.0000 | 0.0000 | 0.8609 | 0 | 2232 | 517.539 | 1714.46 | 0 | 0 | 0 |
| iso | GMYN | NA | 0.0000 | NA | 0 | 2232 | 555.654 | 1676.35 | 0 | 0 | 0 |
| acsA | MS | 0.0013 | 0.0022 | 0.5994 | 0.137823 | 1971 | 487 | 1484 | 3 | 1.06134 | 1.93866 |
| acsA | GMYN | 0.0014 | 0.0020 | 0.6865 | 0.571049 | 1971 | 503.725 | 1467.28 | 3 | 1 | 2 |
| succ | MS | 0.0000 | 0.0000 | 0.8609 | 0 | 1776 | 410.966 | 1365.03 | 0 | 0 | 0 |
| succ | GMYN | 0.0000 | NA | NA | 0 | 1776 | 437.879 | 1338.12 | 0 | 0 | 0 |
| CTP | MS | 0.0000 | 0.0000 | 0.8609 | 0 | 1659 | 396.751 | 1262.25 | 0 | 0 | 0 |
| CTP | GMYN | 0.0000 | 0.0000 | NA | 0 | 1659 | 418.508 | 1240.49 | 0 | 0 | 0 |
| atpA | MS | 0.0000 | 0.0000 | 0.8609 | 0 | 1539 | 380.541 | 1158.46 | 0 | 0 | 0 |
| atpA | GMYN | NA | 0.0000 | NA | 0 | 1539 | 407.726 | 1131.27 | 0 | 0 | 0 |
| gyr | MS | 0.0000 | 0.0000 | 0.8610 | 0 | 1434 | 378.656 | 1055.34 | 0 | 0 | 0 |
| gyr | GMYN | 0.0000 | NA | NA | 0 | 1434 | 387.291 | 1046.71 | 0 | 0 | 0 |
| glnA | MS | 0.0000 | 0.0030 | 0.0000 | 0 | 1413 | 332.108 | 1080.89 | 1 | 0.999997 | 3.25E-06 |

| | | | | | | | | | | | |
|---|---|---|---|---|---|---|---|---|---|---|---|
| glnA | GMYN | 0.0000 | 0.0029 | 0.0000 | 0 | 1413 | 349.801 | 1063.2 | 1 | 1 | 0 |
| cit | MS | 0.0000 | 0.0034 | 0.0000 | 0 | 1311 | 291.309 | 1019.69 | 1 | 0.999996 | 3.50E-06 |
| cit | GMYN | 0.0000 | 0.0032 | 0.0000 | 0 | 1311 | 312.796 | 998.204 | 1 | 1 | 0 |
| adk | MS | 0.0079 | 0.1992 | 0.0397 | 6.71E-15 | 663 | 159.472 | 503.528 | 30 | 26.6596 | 3.34038 |
| adk | GMYN | 0.0078 | 0.2078 | 0.0375 | 1.59E-14 | 663 | 146.132 | 516.868 | 30 | 26 | 4 |
| ndk | MS | 0.0000 | 0.0000 | 0.8603 | 0 | 423 | 97.5858 | 325.414 | 0 | 0 | 0 |
| ndk | GMYN | NA | NA | NA | 0 | 423 | 97.8687 | 325.131 | 0 | 0 | 0 |
| | | | **maximum** | 0.8610 | | | | | | | |
| | | | **minimum** | 0.0000 | | | | | | | |
| | | | **average** | 0.4663 | | | | | | | |
| | | | **median** | 0.6430 | | | | | | | |

### Selected genes in MWH-RechtKol4

| Sequence | Method | Ka | Ks | Ka/Ks | P-Value(Fisher) | Length | S-Sites | N-Sites | Substitutions | S-Substitutions | N-Substitutions |
|---|---|---|---|---|---|---|---|---|---|---|---|
| FtsK | MS | 0.0024 | 0.0345 | 0.0709 | 4.98E-10 | 2310 | 682.337 | 1627.66 | 26 | 22.2388 | 3.76118 |
| FtsK | GMYN | 0.0024 | 0.0362 | 0.0657 | 1.18E-09 | 2310 | 625.81 | 1684.19 | 26 | 22 | 4 |
| Alkyl | MS | 0.0031 | 0.0443 | 0.0708 | 0 | 465 | 144.282 | 320.718 | 7 | 6.04791 | 0.952093 |
| Alkyl | GMYN | 0.0029 | 0.0532 | 0.0549 | 0.00146005 | 465 | 120.669 | 344.331 | 7 | 6 | 1 |
| DSBA | MS | 0.0069 | 0.0285 | 0.2402 | 0.0100676 | 624 | 184.867 | 439.133 | 8 | 5.09395 | 2.90605 |
| DSBA | GMYN | 0.0065 | 0.0321 | 0.2029 | 0.0295253 | 624 | 160.064 | 463.936 | 8 | 5 | 3 |
| Fimb | MS | 0.0086 | 0.1329 | 0.0650 | 9.77E-09 | 510 | 147.434 | 362.566 | 20 | 17.2435 | 2.75652 |
| Fimb | GMYN | 0.0107 | 0.1371 | 0.0777 | 2.66E-07 | 510 | 129.6 | 380.4 | 20 | 16 | 4 |
| GroL | MS | 0.0009 | 0.0020 | 0.4437 | 0 | 1650 | 523.849 | 1126.15 | 2 | 1.02367 | 0.976329 |
| GroL | GMYN | 0.0008 | 0.0023 | 0.3496 | 0.384551 | 1650 | 428.172 | 1221.83 | 2 | 1 | 1 |
| Hyp | MS | 0.0049 | 0.0001 | 50.0000 | 0 | 264 | 62.6998 | 201.3 | 1 | 0.00619091 | 0.993809 |
| Hyp | GMYN | 0.0050 | 0.0000 | NA | 0 | 264 | 64.3212 | 199.679 | 1 | 0 | 1 |
| MscS | MS | 0.0000 | 0.0437 | 0.0000 | 0 | 819 | 216.856 | 602.144 | 9 | 8.99998 | 2.50E-05 |
| MscS | GMYN | NA | 0.0455 | NA | 0 | 819 | 205.048 | 613.952 | 9 | 9 | 0 |
| Piri | MS | 0.0055 | 0.0070 | 0.7869 | 0.214849 | 699 | 156.124 | 542.876 | 4 | 1.07058 | 2.92942 |
| Piri | GMYN | 0.0056 | 0.0061 | 0.9257 | 0.658823 | 699 | 165.625 | 533.375 | 4 | 1 | 3 |
| RpoE | MS | 0.0000 | 0.0151 | 0.0000 | 0 | 567 | 132.016 | 434.984 | 2 | 1.99999 | 6.59E-06 |
| RpoE | GMYN | NA | 0.0142 | NA | 0 | 567 | 142.534 | 424.466 | 2 | 2 | 0 |

| | | | | | | | | | | | |
|---|---|---|---|---|---|---|---|---|---|---|---|
| RuvB | MS | 0.0013 | 0.0715 | 0.0187 | 0 | 1068 | 318.884 | 749.116 | 22 | 21.0732 | 0.926773 |
| RuvB | GMYN | 0.0013 | 0.0836 | 0.0150 | 2.76E-12 | 1068 | 270.522 | 797.478 | 22 | 21 | 1 |
| Superox | MS | 0.0000 | 0.0082 | 0.0000 | 0 | 531 | 122.67 | 408.33 | 1 | 0.999997 | 3.33E-06 |
| Superox | GMYN | 0.0000 | 0.0078 | 0.0000 | 0 | 531 | 130.956 | 400.044 | 1 | 1 | 0 |
| | | | **maximum** | 50.0000 | | | | | | | |
| | | | **minimum** | 0.0000 | | | | | | | |
| | | | **average** | 2.8099 | | | | | | | |
| | | | **median** | 0.0708 | | | | | | | |

### Housekeeping genes in MWH-RechtKol4

| Sequence | Method | Ka | Ks | Ka/Ks | P-Value(Fisher) | Length | S-Sites | N-Sites | Substitutions | S-Substitutions | N-Substitutions |
|---|---|---|---|---|---|---|---|---|---|---|---|
| iso | MS | 0.0019 | 0.0942 | 0.0201 | 1.10E-28 | 2232 | 646.227 | 1585.77 | 58 | 55.2741 | 2.72588 |
| iso | GMYN | 0.0019 | 0.0891 | 0.0216 | 4.46E-26 | 2232 | 670.713 | 1561.29 | 58 | 54.9983 | 3.0017 |
| acsA | MS | 0.0028 | 0.0711 | 0.0392 | 1.51E-17 | 1971 | 547.583 | 1423.42 | 39 | 35.392 | 3.60804 |
| acsA | GMYN | 0.0028 | 0.0699 | 0.0399 | 1.30E-16 | 1971 | 530.919 | 1440.08 | 39 | 35 | 4 |
| succ | MS | 0.0000 | 0.0366 | 0.0000 | 0 | 1776 | 430.6 | 1345.4 | 15 | 15 | 4.69E-05 |
| succ | GMYN | 0.0000 | 0.0369 | 0.0000 | 0 | 1776 | 421.093 | 1354.91 | 15 | 15 | 0 |
| CTP | MS | 0.0018 | 0.0190 | 0.0931 | 7.22E-05 | 1659 | 525.286 | 1133.71 | 11 | 9.15969 | 1.84031 |
| CTP | GMYN | 0.0016 | 0.0221 | 0.0730 | 0.000125077 | 1659 | 417.189 | 1241.81 | 11 | 9 | 2 |
| atpA | MS | 0.0000 | 0.0096 | 0.0000 | 0 | 1539 | 533.542 | 1005.46 | 5 | 4.99999 | 9.42E-06 |
| atpA | GMYN | 0.0000 | 0.0124 | 0.0000 | 0 | 1539 | 407.975 | 1131.02 | 5 | 5 | 0 |
| gyr | MS | 0.0000 | 0.0079 | 0.0000 | 0 | 1434 | 378.849 | 1055.15 | 3 | 2.99999 | 8.36E-06 |
| gyr | GMYN | NA | 0.0076 | NA | 0 | 1434 | 395.131 | 1038.87 | 3 | 3 | 0 |
| glnA | MS | 0.0000 | 0.0247 | 0.0000 | 0 | 1413 | 393.521 | 1019.48 | 9 | 8.99998 | 2.33E-05 |
| glnA | GMYN | 0.0000 | 0.0264 | 0.0000 | 0 | 1413 | 348.816 | 1064.18 | 9 | 9 | 0 |
| cit | MS | 0.0016 | 0.0234 | 0.0684 | 0.000382964 | 1311 | 267.118 | 1043.88 | 7 | 5.52364 | 1.47636 |
| cit | GMYN | 0.0021 | 0.0160 | 0.1290 | 0.0101498 | 1311 | 312.415 | 998.585 | 7 | 4.9408 | 2.0592 |
| adk | MS | 0.0083 | 0.1850 | 0.0449 | 2.00E-14 | 663 | 181.819 | 481.181 | 32 | 28.5993 | 3.40068 |
| adk | GMYN | 0.0090 | 0.1475 | 0.0609 | 5.40E-11 | 663 | 215.007 | 447.993 | 32 | 28 | 4 |
| ndk | MS | 0.0000 | 0.0000 | 0.8603 | 0 | 423 | 97.5858 | 325.414 | 0 | 0 | 0 |
| ndk | GMYN | NA | NA | NA | 0 | 423 | 97.8687 | 325.131 | 0 | 0 | 0 |
| | | | **maximum** | 0.8603 | | | | | | | |
| | | | **minimum** | 0.0000 | | | | | | | |

| | |
|---|---|
| **average** | 0.0806 |
| **median** | 0.0304 |

### Selected genes in MWH-RechtKolB

| Sequence | Method | Ka | Ks | Ka/Ks | P-Value(Fisher) | Length | S-Sites | N-Sites | Substitutions | S-Substitutions | N-Substitutions |
|---|---|---|---|---|---|---|---|---|---|---|---|
| FtsK | MS | 0.0024 | 0.0345 | 0.0709 | 4.98E-10 | 2310 | 682.337 | 1627.66 | 26 | 22.2388 | 3.76118 |
| FtsK | GMYN | 0.0024 | 0.0362 | 0.0657 | 1.18E-09 | 2310 | 625.81 | 1684.19 | 26 | 22 | 4 |
| Alkyl | MS | 0.0031 | 0.0443 | 0.0708 | 0 | 465 | 144.282 | 320.718 | 7 | 6.04791 | 0.952093 |
| Alkyl | GMYN | 0.0029 | 0.0532 | 0.0549 | 0.00146005 | 465 | 120.669 | 344.331 | 7 | 6 | 1 |
| DSBA | MS | 0.0069 | 0.0285 | 0.2402 | 0.0100676 | 624 | 184.867 | 439.133 | 8 | 5.09395 | 2.90605 |
| DSBA | GMYN | 0.0065 | 0.0321 | 0.2029 | 0.0295253 | 624 | 160.064 | 463.936 | 8 | 5 | 3 |
| Fimb | MS | 0.0086 | 0.1329 | 0.0650 | 9.77E-09 | 510 | 147.434 | 362.566 | 20 | 17.2435 | 2.75652 |
| Fimb | GMYN | 0.0107 | 0.1371 | 0.0777 | 2.66E-07 | 510 | 129.6 | 380.4 | 20 | 16 | 4 |
| GroL | MS | 0.0009 | 0.0020 | 0.4437 | 0 | 1650 | 523.849 | 1126.15 | 2 | 1.02367 | 0.976329 |
| GroL | GMYN | 0.0008 | 0.0023 | 0.3496 | 0.384551 | 1650 | 428.172 | 1221.83 | 2 | 1 | 1 |
| Hyp | MS | 0.0049 | 0.0001 | 50.0000 | 0 | 264 | 62.6998 | 201.3 | 1 | 0.00619091 | 0.993809 |
| Hyp | GMYN | 0.0050 | 0.0000 | NA | 0 | 264 | 64.3212 | 199.679 | 1 | 0 | 1 |
| MscS | MS | 0.0000 | 0.0437 | 0.0000 | 0 | 819 | 216.856 | 602.144 | 9 | 8.99998 | 2.50E-05 |
| MscS | GMYN | NA | 0.0455 | NA | 0 | 819 | 205.048 | 613.952 | 9 | 9 | 0 |
| Piri | MS | 0.0055 | 0.0070 | 0.7869 | 0.214849 | 699 | 156.124 | 542.876 | 4 | 1.07058 | 2.92942 |
| Piri | GMYN | 0.0056 | 0.0061 | 0.9257 | 0.658823 | 699 | 165.625 | 533.375 | 4 | 1 | 3 |
| RpoE | MS | 0.0000 | 0.0151 | 0.0000 | 0 | 567 | 132.016 | 434.984 | 2 | 1.99999 | 6.59E-06 |
| RpoE | GMYN | NA | 0.0142 | NA | 0 | 567 | 142.534 | 424.466 | 2 | 2 | 0 |
| RuvB | MS | 0.0013 | 0.0715 | 0.0187 | 0 | 1068 | 318.884 | 749.116 | 22 | 21.0732 | 0.926773 |
| RuvB | GMYN | 0.0013 | 0.0836 | 0.0150 | 2.76E-12 | 1068 | 270.522 | 797.478 | 22 | 21 | 1 |
| Superox | MS | 0.0000 | 0.0082 | 0.0000 | 0 | 531 | 122.67 | 408.33 | 1 | 0.999997 | 3.33E-06 |
| Superox | GMYN | 0.0000 | 0.0078 | 0.0000 | 0 | 531 | 130.956 | 400.044 | 1 | 1 | 0 |
| | | | **maximum** | 50.0000 | | | | | | | |
| | | | **minimum** | 0.0000 | | | | | | | |
| | | | **average** | 2.8099 | | | | | | | |
| | | | **median** | 0.0708 | | | | | | | |

### Housekeeping genes in MWH-RechtKolB

| Sequence | Method | Ka | Ks | Ka/Ks | P-Value(Fisher) | Length | S-Sites | N-Sites | Substitutions | S-Substitutions | N-Substitutions |
|---|---|---|---|---|---|---|---|---|---|---|---|
| iso | MS | 0.0019 | 0.0942 | 0.0201 | 1.10E-28 | 2232 | 646.227 | 1585.77 | 58 | 55.2741 | 2.72588 |
| iso | GMYN | 0.0019 | 0.0891 | 0.0216 | 4.46E-26 | 2232 | 670.713 | 1561.29 | 58 | 54.9983 | 3.0017 |
| acsA | MS | 0.0028 | 0.0711 | 0.0392 | 1.51E-17 | 1971 | 547.583 | 1423.42 | 39 | 35.392 | 3.60804 |
| acsA | GMYN | 0.0028 | 0.0699 | 0.0399 | 1.30E-16 | 1971 | 530.919 | 1440.08 | 39 | 35 | 4 |
| succ | MS | 0.0000 | 0.0366 | 0.0000 | 0 | 1776 | 430.6 | 1345.4 | 15 | 15 | 4.69E-05 |
| succ | GMYN | 0.0000 | 0.0369 | 0.0000 | 0 | 1776 | 421.093 | 1354.91 | 15 | 15 | 0 |
| CTP | MS | 0.0018 | 0.0190 | 0.0931 | 7.22E-05 | 1659 | 525.286 | 1133.71 | 11 | 9.15969 | 1.84031 |
| CTP | GMYN | 0.0016 | 0.0221 | 0.0730 | 0.000125077 | 1659 | 417.189 | 1241.81 | 11 | 9 | 2 |
| atpA | MS | 0.0000 | 0.0096 | 0.0000 | 0 | 1539 | 533.542 | 1005.46 | 5 | 4.99999 | 9.42E-06 |
| atpA | GMYN | 0.0000 | 0.0124 | 0.0000 | 0 | 1539 | 407.975 | 1131.02 | 5 | 5 | 0 |
| gyr | MS | 0.0000 | 0.0079 | 0.0000 | 0 | 1434 | 378.849 | 1055.15 | 3 | 2.99999 | 8.36E-06 |
| gyr | GMYN | NA | 0.0076 | NA | 0 | 1434 | 395.131 | 1038.87 | 3 | 3 | 0 |
| glnA | MS | 0.0000 | 0.0247 | 0.0000 | 0 | 1413 | 393.521 | 1019.48 | 9 | 8.99998 | 2.33E-05 |
| glnA | GMYN | 0.0000 | 0.0264 | 0.0000 | 0 | 1413 | 348.816 | 1064.18 | 9 | 9 | 0 |
| cit | MS | 0.0016 | 0.0234 | 0.0684 | 0.000382964 | 1311 | 267.118 | 1043.88 | 7 | 5.52364 | 1.47636 |
| cit | GMYN | 0.0021 | 0.0160 | 0.1290 | 0.0101498 | 1311 | 312.415 | 998.585 | 7 | 4.9408 | 2.0592 |
| adk | MS | 0.0083 | 0.1850 | 0.0449 | 2.00E-14 | 663 | 181.819 | 481.181 | 32 | 28.5993 | 3.40068 |
| adk | GMYN | 0.0090 | 0.1475 | 0.0609 | 5.40E-11 | 663 | 215.007 | 447.993 | 32 | 28 | 4 |
| ndk | MS | 0.0000 | 0.0000 | 0.8603 | 0 | 423 | 97.5858 | 325.414 | 0 | 0 | 0 |
| ndk | GMYN | NA | NA | NA | 0 | 423 | 97.8687 | 325.131 | 0 | 0 | 0 |
| | | | **maximum** | 0.8603 | | | | | | | |
| | | | **minimum** | 0.0000 | | | | | | | |
| | | | **average** | 0.0806 | | | | | | | |
| | | | **median** | 0.0304 | | | | | | | |

### Selected genes in MWH-Tro7-1-4

| Sequence | Method | Ka | Ks | Ka/Ks | P-Value(Fisher) | Length | S-Sites | N-Sites | Substitutions | S-Substitutions | N-Substitutions |
|---|---|---|---|---|---|---|---|---|---|---|---|
| RuvB | MS | 0.0000 | 0.0037 | 0.0000 | 0 | 1068 | 272.786 | 795.214 | 1 | 0.999997 | 2.92E-06 |
| RuvB | GMYN | 0.0000 | 0.0036 | 0.0000 | 0 | 1068 | 282.739 | 785.261 | 1 | 1 | 0 |

| | | | | | | | | | | | |
|---|---|---|---|---|---|---|---|---|---|---|---|
| Fimb | MS | 0.0055 | 0.0540 | 0.1012 | 0.00017699 | 510 | 139.271 | 370.729 | 9 | 7.09043 | 1.90957 |
| Fimb | GMYN | 0.0054 | 0.0552 | 0.0972 | 0.00180129 | 510 | 135.493 | 374.507 | 9 | 7 | 2 |
| RpoE | MS | 0.0045 | 0.0336 | 0.1354 | 0.00299834 | 567 | 132.04 | 434.96 | 6 | 4.14927 | 1.85073 |
| RpoE | GMYN | 0.0047 | 0.0289 | 0.1636 | 0.0042709 | 567 | 142.236 | 424.764 | 6 | 4 | 2 |
| Superox | MS | 0.0083 | 0.0606 | 0.1365 | 0.000221587 | 531 | 163.432 | 367.568 | 12 | 9.18071 | 2.81929 |
| Superox | GMYN | 0.0075 | 0.0772 | 0.0974 | 0.00026245 | 531 | 129.114 | 401.886 | 12 | 9 | 3 |
| | | | **maximum** | 0.1636 | | | | | | | |
| | | | **minimum** | 0.0000 | | | | | | | |
| | | | **average** | 0.0914 | | | | | | | |
| | | | **median** | 0.0993 | | | | | | | |

### Housekeeping genes in MWH-Tro7-1-4

| Sequence | Method | Ka | Ks | Ka/Ks | P-Value(Fisher) | Length | S-Sites | N-Sites | Substitutions | S-Substitutions | N-Substitutions |
|---|---|---|---|---|---|---|---|---|---|---|---|
| iso | MS | 0.0006 | 0.0236 | 0.0252 | 0 | 2232 | 561.142 | 1670.86 | 14 | 13.0219 | 0.978057 |
| iso | GMYN | 0.0006 | 0.0239 | 0.0250 | 1.35E-07 | 2232 | 555.944 | 1676.06 | 14 | 13 | 1 |
| acsA | MS | 0.0015 | 0.0000 | 50.0000 | 0.434142 | 1971 | 627.238 | 1343.76 | 2 | 0.0184984 | 1.9815 |
| acsA | GMYN | 0.0014 | 0.0000 | 1.2303E+12 | 0 | 1971 | 503.109 | 1467.89 | 2 | 0 | 2 |
| succ | MS | 0.0000 | 0.0000 | 0.8609 | 0 | 1776 | 410.966 | 1365.03 | 0 | 0 | 0 |
| succ | GMYN | 0.0000 | NA | NA | 0 | 1776 | 437.879 | 1338.12 | 0 | 0 | 0 |
| CTP | MS | 0.0000 | 0.0000 | 0.8609 | 0 | 1659 | 396.751 | 1262.25 | 0 | 0 | 0 |
| CTP | GMYN | 0.0000 | 0.0000 | NA | 0 | 1659 | 418.508 | 1240.49 | 0 | 0 | 0 |
| atpA | MS | 0.0000 | 0.0000 | 0.8609 | 0 | 1539 | 380.541 | 1158.46 | 0 | 0 | 0 |
| atpA | GMYN | NA | 0.0000 | NA | 0 | 1539 | 407.726 | 1131.27 | 0 | 0 | 0 |
| gyr | MS | 0.0000 | 0.0144 | 0.0000 | 0 | 1434 | 430.428 | 1003.57 | 6 | 5.99999 | 1.40E-05 |
| gyr | GMYN | NA | 0.0149 | NA | 0 | 1434 | 408.105 | 1025.9 | 6 | 6 | 0 |
| glnA | MS | 0.0000 | 0.0060 | 0.0000 | 0 | 1413 | 332.456 | 1080.54 | 2 | 1.99999 | 6.50E-06 |
| glnA | GMYN | NA | 0.0057 | NA | 0 | 1413 | 350.238 | 1062.76 | 2 | 2 | 0 |
| cit | MS | 0.0000 | 0.0000 | 0.8607 | 0 | 1311 | 291.476 | 1019.52 | 0 | 0 | 0 |
| cit | GMYN | NA | NA | NA | 0 | 1311 | 313.061 | 997.939 | 0 | 0 | 0 |
| adk | MS | 0.0000 | 0.0000 | 0.8610 | 0 | 663 | 160.97 | 502.03 | 0 | 0 | 0 |
| adk | GMYN | 0.0000 | NA | NA | 0 | 663 | 170.565 | 492.435 | 0 | 0 | 0 |
| ndk | MS | 0.0031 | 0.0001 | 50.0000 | 0 | 423 | 97.5862 | 325.414 | 1 | 0.00596191 | 0.994038 |
| ndk | GMYN | 0.0031 | NA | NA | 0 | 423 | 97.7649 | 325.235 | 1 | 0 | 1 |

| | |
|---|---|
| **maximum** | 1.2303E+12 |
| **minimum** | 0.0000 |
| **average** | 1.0252E+11 |
| **median** | 0.8609 |

### Selected genes in MWH-Tro8-2-5gr

| Sequence | Method | Ka | Ks | Ka/Ks | P-Value(Fisher) | Length | S-Sites | N-Sites | Substitutions | S-Substitutions | N-Substitutions |
|---|---|---|---|---|---|---|---|---|---|---|---|
| Alkyl | MS | 0.0309 | 2.5430 | 0.0121 | 3.46E-65 | 444 | 108.251 | 335.749 | 88 | 84.8045 | 3.19553 |
| Alkyl | GMYN | 0.0333 | 4.3104 | 0.0077 | 4.72E-51 | 444 | 102.89 | 341.11 | 88 | 76.9209 | 11.0791 |
| Cold | MS | 0.0000 | 0.0495 | 0.0000 | 0 | 180 | 40.3807 | 139.619 | 2 | 1.99999 | 6.92E-06 |
| Cold | GMYN | 0.0000 | 0.0516 | 0.0000 | 0 | 180 | 42.6501 | 137.35 | 2 | 2 | 0 |
| DnaK | MS | 0.0120 | 0.7478 | 0.0161 | 6.32E-128 | 1899 | 549.353 | 1349.65 | 239 | 229.928 | 9.07215 |
| DnaK | GMYN | 0.0115 | 0.8884 | 0.0130 | 2.93E-129 | 1899 | 489.13 | 1409.87 | 239 | 222.916 | 16.0843 |
| DSBA | MS | 0.1276 | 2.9315 | 0.0435 | NA | 603 | 138.561 | 464.439 | 163 | 142.251 | 20.7487 |
| DSBA | GMYN | 0.1248 | 3.6412 | 0.0343 | 1.15E-57 | 603 | 128.368 | 474.632 | 163 | 108.823 | 54.1775 |
| Fatty | MS | 0.0642 | 2.4045 | 0.0267 | 2.80E-189 | 1167 | 268.666 | 898.334 | 262 | 240.539 | 21.4612 |
| Fatty | GMYN | 0.0623 | 4.1310 | 0.0151 | 1.00E-135 | 1167 | 246.651 | 920.349 | 262 | 207.187 | 54.8133 |
| Fimb | MS | 0.0559 | 3.8195 | 0.0146 | 1.50E-86 | 498 | 124.921 | 373.079 | 113 | 108.266 | 4.73378 |
| Fimb | GMYN | 0.0562 | 3.6049 | 0.0156 | 4.11E-54 | 498 | 122.299 | 375.701 | 113 | 92.7167 | 20.2833 |
| FtsK | MS | 0.0149 | 1.0308 | 0.0145 | 1.69E-209 | 2277 | 623.013 | 1653.99 | 354 | 340.878 | 13.1224 |
| FtsK | GMYN | 0.0139 | 1.3083 | 0.0106 | 7.42E-195 | 2277 | 606.13 | 1670.87 | 354 | 330.962 | 23.038 |
| GroL | MS | 0.0060 | 0.5393 | 0.0111 | 7.01E-90 | 1620 | 460.493 | 1159.51 | 162 | 157.582 | 4.41769 |
| GroL | GMYN | 0.0058 | 0.6509 | 0.0088 | 1.40E-95 | 1620 | 399.484 | 1220.52 | 162 | 155 | 7.00048 |
| Hfq | MS | 0.0330 | 0.5506 | 0.0598 | 2.25E-12 | 228 | 67.352 | 160.648 | 28 | 24.5025 | 3.49753 |
| Hfq | GMYN | 0.0307 | 1.0094 | 0.0304 | 7.28E-11 | 228 | 61.2365 | 166.763 | 28 | 23 | 5 |
| Piri | MS | 0.1350 | 3.1715 | 0.0426 | NA | 675 | 144.253 | 530.747 | 176 | 152.168 | 23.8322 |
| Piri | GMYN | 0.1387 | 6.1000 | 0.0227 | 6.44E-48 | 675 | 146.142 | 528.858 | 176 | 109.574 | 66.426 |
| RpoE | MS | NA | NA | NA | NA | NA | NA | NA | NA | NA | NA |
| RpoE | GMYN | 0.0305 | 0.7767 | 0.0393 | 3.39E-31 | 546 | 136.588 | 409.412 | 74 | 61.7805 | 12.2195 |
| RpoH | MS | 0.0322 | 1.1725 | 0.0274 | 1.47E-82 | 903 | 262.057 | 640.943 | 155 | 145.254 | 9.74615 |
| RpoH | GMYN | 0.0298 | 1.3640 | 0.0218 | 1.54E-80 | 903 | 214.772 | 688.228 | 155 | 134.944 | 20.0557 |
| Rubre | MS | 0.0367 | 2.7185 | 0.0135 | 6.40E-50 | 462 | 147.886 | 314.114 | 88 | 85.5493 | 2.45071 |

| | | | | | | | | | | | |
|---|---|---|---|---|---|---|---|---|---|---|---|
| Rubre | GMYN | 0.0345 | 1.7929 | 0.0192 | 2.66E-38 | 462 | 134.721 | 327.279 | 88 | 77 | 11 |
| | | | **maximum** | 0.0598 | | | | | | | |
| | | | **minimum** | 0.0000 | | | | | | | |
| | | | **average** | 0.0208 | | | | | | | |
| | | | **median** | 0.0156 | | | | | | | |

### Housekeeping genes in MWH-Tro8-2-5gr

| Sequence | Method | Ka | Ks | Ka/Ks | P-Value(Fisher) | Length | S-Sites | N-Sites | Substitutions | S-Substitutions | N-Substitutions |
|---|---|---|---|---|---|---|---|---|---|---|---|
| iso | MS | 0.0223 | 0.9998 | 0.0223 | 1.13E-190 | 2232 | 596.507 | 1635.49 | 339 | 319.447 | 19.5526 |
| iso | GMYN | 0.0222 | 1.3559 | 0.0164 | 2.72E-166 | 2232 | 575.916 | 1656.08 | 339 | 302.855 | 36.1454 |
| acsA | MS | 0.0303 | 0.8617 | 0.0351 | 6.64E-147 | 1971 | 493.945 | 1477.05 | 281 | 254.288 | 26.7124 |
| acsA | GMYN | 0.0272 | 1.5565 | 0.0175 | 3.77E-141 | 1971 | 432.585 | 1538.41 | 281 | 239.927 | 41.0729 |
| succ | MS | 0.0126 | 0.8678 | 0.0145 | 1.08E-139 | 1776 | 459.187 | 1316.81 | 233 | 223.668 | 9.33209 |
| succ | GMYN | 0.0111 | 1.6003 | 0.0069 | 3.14E-141 | 1776 | 413.624 | 1362.38 | 233 | 218 | 15.0005 |
| CTP | MS | 0.0291 | 1.8451 | 0.0158 | 1.39E-216 | 1659 | 396.265 | 1262.73 | 309 | 294.203 | 14.7971 |
| CTP | GMYN | 0.0292 | 2.0363 | 0.0143 | 9.17E-174 | 1659 | 389.391 | 1269.61 | 309 | 272.709 | 36.2908 |
| atpA | MS | 0.0029 | 0.2471 | 0.0119 | 5.50E-48 | 1539 | 416.33 | 1122.67 | 86 | 83.3313 | 2.66866 |
| atpA | GMYN | 0.0026 | 0.2821 | 0.0092 | 8.33E-50 | 1539 | 378.591 | 1160.41 | 86 | 82.9997 | 3.00028 |
| gyr | MS | 0.0306 | 1.9569 | 0.0156 | 6.70E-173 | 1434 | 394.482 | 1039.52 | 275 | 264.113 | 10.8867 |
| gyr | GMYN | 0.0314 | 1.7556 | 0.0179 | 3.20E-125 | 1434 | 410.89 | 1023.11 | 275 | 243.578 | 31.4223 |
| glnA | MS | 0.0163 | 0.4674 | 0.0349 | 3.72E-65 | 1413 | 369.219 | 1043.78 | 136 | 123.8 | 12.2004 |
| glnA | GMYN | 0.0140 | 0.8743 | 0.0161 | 5.05E-67 | 1413 | 332.257 | 1080.74 | 136 | 120.992 | 15.0082 |
| cit | MS | 0.0140 | 0.8290 | 0.0169 | 3.17E-103 | 1311 | 318.87 | 992.13 | 168 | 159.61 | 8.38997 |
| cit | GMYN | 0.0128 | 2.1119 | 0.0061 | 3.95E-105 | 1311 | 281.381 | 1029.62 | 168 | 154.948 | 13.0524 |
| adk | MS | 0.0281 | 0.8887 | 0.0316 | 2.92E-49 | 663 | 190.603 | 472.397 | 96 | 89.0224 | 6.97763 |
| adk | GMYN | 0.0270 | 1.0241 | 0.0263 | 2.16E-43 | 663 | 169.874 | 493.126 | 96 | 82.9971 | 13.0029 |
| ndk | MS | 0.0029 | 0.4709 | 0.0063 | 0 | 423 | 87.6668 | 335.333 | 29 | 28.3217 | 0.678306 |
| ndk | GMYN | 0.0030 | 0.4702 | 0.0065 | 3.64E-19 | 423 | 94.1205 | 328.879 | 29 | 28 | 1 |
| | | | **maximum** | 0.0351 | | | | | | | |
| | | | **minimum** | 0.0061 | | | | | | | |
| | | | **average** | 0.0171 | | | | | | | |
| | | | **median** | 0.0159 | | | | | | | |

**Selected genes in MWH-Tro8-2-9**

| Sequence | Method | Ka | Ks | Ka/Ks | P-Value(Fisher) | Length | S-Sites | N-Sites | Substitutions | S-Substitutions | N-Substitutions |
|---|---|---|---|---|---|---|---|---|---|---|---|
| RpoE | MS | 0.0000 | 0.0076 | 0.0000 | 0 | 567 | 131.79 | 435.21 | 1 | 0.999997 | 3.30E-06 |
| RpoE | GMYN | 0.0000 | 0.0071 | 0.0000 | 0 | 567 | 142.392 | 424.608 | 1 | 1 | 0 |
| | | | **maximum** | 0.0000 | | | | | | | |
| | | | **minimum** | 0.0000 | | | | | | | |
| | | | **average** | 0.0000 | | | | | | | |
| | | | **median** | 0.0000 | | | | | | | |

**Housekeeping genes in MWH-Tro8-2-9**

| Sequence | Method | Ka | Ks | Ka/Ks | P-Value(Fisher) | Length | S-Sites | N-Sites | Substitutions | S-Substitutions | N-Substitutions |
|---|---|---|---|---|---|---|---|---|---|---|---|
| iso | MS | 0.0000 | 0.0000 | 0.8609 | 0 | 2232 | 517.539 | 1714.46 | 0 | 0 | 0 |
| iso | GMYN | NA | 0.0000 | NA | 0 | 2232 | 555.654 | 1676.35 | 0 | 0 | 0 |
| acsA | MS | 0.0007 | 0.0247 | 0.0269 | 0 | 1971 | 486.858 | 1484.14 | 12 | 11.0915 | 0.908467 |
| acsA | GMYN | 0.0007 | 0.0222 | 0.0307 | 2.52E-06 | 1971 | 503.811 | 1467.19 | 12 | 11 | 1 |
| succ | MS | 0.0000 | 0.0000 | 0.8609 | 0 | 1776 | 410.966 | 1365.03 | 0 | 0 | 0 |
| succ | GMYN | 0.0000 | NA | NA | 0 | 1776 | 437.879 | 1338.12 | 0 | 0 | 0 |
| CTP | MS | 0.0000 | 0.0000 | 0.8609 | 0 | 1659 | 396.751 | 1262.25 | 0 | 0 | 0 |
| CTP | GMYN | 0.0000 | 0.0000 | NA | 0 | 1659 | 418.508 | 1240.49 | 0 | 0 | 0 |
| atpA | MS | 0.0000 | 0.0000 | 0.8609 | 0 | 1539 | 380.541 | 1158.46 | 0 | 0 | 0 |
| atpA | GMYN | NA | 0.0000 | NA | 0 | 1539 | 407.726 | 1131.27 | 0 | 0 | 0 |
| gyr | MS | 0.0000 | 0.0000 | 0.8610 | 0 | 1434 | 378.656 | 1055.34 | 0 | 0 | 0 |
| gyr | GMYN | 0.0000 | NA | NA | 0 | 1434 | 387.291 | 1046.71 | 0 | 0 | 0 |
| glnA | MS | 0.0000 | 0.0000 | 0.8607 | 0 | 1413 | 332.453 | 1080.55 | 0 | 0 | 0 |
| glnA | GMYN | 0.0000 | 0.0000 | NA | 0 | 1413 | 350.104 | 1062.9 | 0 | 0 | 0 |
| cit | MS | 0.0000 | 0.0034 | 0.0000 | 0 | 1311 | 291.309 | 1019.69 | 1 | 0.999996 | 3.50E-06 |
| cit | GMYN | 0.0000 | 0.0032 | 0.0000 | 0 | 1311 | 312.796 | 998.204 | 1 | 1 | 0 |
| adk | MS | 0.0000 | 0.0000 | 0.8610 | 0 | 663 | 160.97 | 502.03 | 0 | 0 | 0 |
| adk | GMYN | 0.0000 | NA | NA | 0 | 663 | 170.565 | 492.435 | 0 | 0 | 0 |
| ndk | MS | 0.0000 | 0.0102 | 0.0000 | 0 | 423 | 98.1022 | 324.898 | 1 | 0.999997 | 3.31E-06 |
| ndk | GMYN | 0.0000 | 0.0103 | 0.0000 | 0 | 423 | 98.4044 | 324.596 | 1 | 1 | 0 |

| | |
|---|---|
| **maximum** | 0.8610 |
| **minimum** | 0.0000 |
| **average** | 0.4680 |
| **median** | 0.8607 |

### Selected genes in MWH-UH21B

| Sequence | Method | Ka | Ks | Ka/Ks | P-Value(Fisher) | Length | S-Sites | N-Sites | Substitutions | S-Substitutions | N-Substitutions |
|---|---|---|---|---|---|---|---|---|---|---|---|
| FtsK | MS | 0.0222 | 1.2567 | 0.0177 | 1.08E-233 | 2292 | 651.722 | 1640.28 | 405 | 387.739 | 17.2606 |
| FtsK | GMYN | 0.0229 | 1.2204 | 0.0188 | 1.41E-185 | 2292 | 685.669 | 1606.33 | 405 | 368.807 | 36.1931 |
| Alkyl | MS | 0.0605 | 2.5784 | 0.0235 | 5.54E-73 | 459 | 112.785 | 346.215 | 102 | 95.1455 | 6.85449 |
| Alkyl | GMYN | 0.0621 | 3.4839 | 0.0178 | 1.74E-48 | 459 | 104.089 | 354.911 | 102 | 80.9172 | 21.0828 |
| Carbo | MS | 0.0326 | 2.0699 | 0.0158 | 4.94E-94 | 654 | 149.461 | 504.539 | 126 | 119.631 | 6.36862 |
| Carbo | GMYN | 0.0327 | 2.6131 | 0.0125 | 2.31E-71 | 654 | 151.246 | 502.754 | 126 | 109.954 | 16.0458 |
| Cold | MS | 0.0000 | 0.1104 | 0.0000 | 0 | 195 | 45.2978 | 149.702 | 5 | 4.99998 | 1.65E-05 |
| Cold | GMYN | NA | 0.1144 | NA | 0 | 195 | 48.3411 | 146.659 | 5 | 5 | 0 |
| DnaK | MS | 0.0163 | 0.9043 | 0.0180 | 3.02E-146 | 1917 | 539.196 | 1377.8 | 265 | 253.358 | 11.6424 |
| DnaK | GMYN | 0.0156 | 1.1806 | 0.0132 | 2.47E-141 | 1917 | 483.017 | 1433.98 | 265 | 242.828 | 22.1722 |
| DSBA | MS | 0.1640 | 3.8153 | 0.0430 | NA | 618 | 141.504 | 476.496 | 187 | 163.361 | 23.6387 |
| DSBA | GMYN | 0.1667 | 3.6609 | 0.0455 | 1.10E-55 | 618 | 131.796 | 486.204 | 187 | 114.925 | 72.0753 |
| Fatty | MS | 0.0708 | 1.9511 | 0.0363 | 3.10E-164 | 1182 | 312.083 | 869.917 | 276 | 250.66 | 25.3403 |
| Fatty | GMYN | 0.0672 | 4.1833 | 0.0161 | 2.82E-133 | 1182 | 264.481 | 917.519 | 276 | 217.206 | 58.7944 |
| Fimb | MS | 0.0778 | 3.6881 | 0.0211 | 1.30E-91 | 510 | 130.276 | 379.724 | 125 | 117.764 | 7.23638 |
| Fimb | GMYN | 0.0788 | 3.6154 | 0.0218 | 5.38E-53 | 510 | 124.028 | 385.972 | 125 | 96.2343 | 28.7657 |
| GroL | MS | 0.0098 | 0.7316 | 0.0134 | 6.69E-107 | 1635 | 445.156 | 1189.84 | 187 | 180.532 | 6.46815 |
| GroL | GMYN | 0.0096 | 0.8940 | 0.0108 | 1.68E-111 | 1635 | 377.041 | 1257.96 | 187 | 174.998 | 12.0019 |
| Hfq | MS | 0.0422 | 0.9785 | 0.0431 | 6.20E-20 | 234 | 60.883 | 173.117 | 38 | 33.8495 | 4.15045 |
| Hfq | GMYN | 0.0414 | 1.2440 | 0.0333 | 8.08E-16 | 234 | 59.4434 | 174.557 | 38 | 30.9876 | 7.01241 |
| Piri | MS | 0.1323 | 4.4088 | 0.0300 | NA | 690 | 139.996 | 550.004 | 191 | 170.853 | 20.1475 |
| Piri | GMYN | 0.1361 | 3.7244 | 0.0365 | 2.62E-65 | 690 | 143.423 | 546.577 | 191 | 123.533 | 67.4672 |
| RpoH | MS | 0.0392 | 1.2508 | 0.0313 | 1.71E-91 | 912 | 255.122 | 656.878 | 169 | 156.386 | 12.6136 |
| RpoH | GMYN | 0.0378 | 2.9140 | 0.0130 | 1.69E-78 | 912 | 230.945 | 681.055 | 169 | 143.987 | 25.0134 |
| RuvB | MS | 0.0229 | 3.3218 | 0.0069 | 1.49E-179 | 1053 | 255.307 | 797.693 | 222 | 217.314 | 4.68602 |

| | | | | | | | | | | | |
|---|---|---|---|---|---|---|---|---|---|---|---|
| RuvB | GMYN | 0.0227 | 4.1202 | 0.0055 | 7.49E-153 | 1053 | 243.125 | 809.875 | 222 | 203.947 | 18.0532 |
| | | | **maximum** | 0.0455 | | | | | | | |
| | | | **minimum** | 0.0000 | | | | | | | |
| | | | **average** | 0.0218 | | | | | | | |
| | | | **median** | 0.0180 | | | | | | | |

### Housekeeping genes in MWH-UH21B

| Sequence | Method | Ka | Ks | Ka/Ks | P-Value(Fisher) | Length | S-Sites | N-Sites | Substitutions | S-Substitutions | N-Substitutions |
|---|---|---|---|---|---|---|---|---|---|---|---|
| iso | MS | 0.0246 | 1.0173 | 0.0242 | 7.82E-196 | 2232 | 578.399 | 1653.6 | 345 | 322.713 | 22.287 |
| iso | GMYN | 0.0239 | 2.2572 | 0.0106 | 1.91E-182 | 2232 | 522.921 | 1709.08 | 345 | 304.774 | 40.2261 |
| acsA | MS | 0.0335 | 0.9679 | 0.0346 | 1.03E-150 | 1971 | 592.697 | 1378.3 | 318 | 294.294 | 23.7064 |
| acsA | GMYN | 0.0307 | 1.1675 | 0.0263 | 7.83E-157 | 1971 | 460.746 | 1510.25 | 318 | 272.702 | 45.2985 |
| succ | MS | 0.0249 | 0.8416 | 0.0296 | 3.14E-131 | 1776 | 464.767 | 1311.23 | 249 | 229.811 | 19.1886 |
| succ | GMYN | 0.0235 | 2.8881 | 0.0081 | 2.81E-134 | 1776 | 383.146 | 1392.85 | 249 | 216.877 | 32.1227 |
| CTP | MS | 0.0311 | 2.2809 | 0.0136 | 7.07E-232 | 1659 | 402.797 | 1256.2 | 324 | 310.796 | 13.2037 |
| CTP | GMYN | 0.0283 | 2.8327 | 0.0100 | 1.49E-188 | 1659 | 394.251 | 1264.75 | 324 | 288.919 | 35.0806 |
| atpA | MS | NA | NA | NA | NA | NA | NA | NA | NA | NA | NA |
| atpA | GMYN | 0.0099 | 0.3724 | 0.0265 | 1.84E-55 | 1539 | 417.596 | 1121.4 | 123 | 112 | 11 |
| gyr | MS | 0.0400 | 2.7876 | 0.0144 | 2.61E-202 | 1434 | 380.038 | 1053.96 | 299 | 287.548 | 11.4519 |
| gyr | GMYN | 0.0413 | 1.5850 | 0.0260 | 5.04E-137 | 1434 | 390.054 | 1043.95 | 299 | 257.19 | 41.8102 |
| glnA | MS | 0.0245 | 0.4959 | 0.0494 | 3.22E-65 | 1413 | 347.415 | 1065.59 | 144 | 125.049 | 18.9507 |
| glnA | GMYN | 0.0231 | 0.7397 | 0.0312 | 5.55E-64 | 1413 | 301.413 | 1111.59 | 144 | 118.749 | 25.2505 |
| cit | MS | 0.0220 | 0.9288 | 0.0237 | 2.36E-108 | 1311 | 325.072 | 985.928 | 186 | 173.521 | 12.4791 |
| cit | GMYN | 0.0194 | 1.5580 | 0.0124 | 2.85E-98 | 1311 | 316.295 | 994.705 | 186 | 166.995 | 19.0047 |
| adk | MS | 0.0273 | 1.6085 | 0.0170 | 2.16E-71 | 663 | 178.29 | 484.71 | 116 | 110.878 | 5.12185 |
| adk | GMYN | 0.0278 | 1.8006 | 0.0154 | 1.89E-66 | 663 | 149.552 | 513.448 | 116 | 101.997 | 14.0029 |
| ndk | MS | 0.0300 | 0.5242 | 0.0573 | 2.44E-19 | 423 | 112.061 | 310.939 | 45 | 38.829 | 6.17102 |
| ndk | GMYN | 0.0281 | 0.6091 | 0.0461 | 1.96E-18 | 423 | 94.9409 | 328.059 | 45 | 35.9672 | 9.03279 |
| | | | **maximum** | 0.0573 | | | | | | | |
| | | | **minimum** | 0.0081 | | | | | | | |
| | | | **average** | 0.0251 | | | | | | | |
| | | | **median** | 0.0242 | | | | | | | |

### Selected genes in MWH-Weng1-1

| Sequence | Method | Ka | Ks | Ka/Ks | P-Value(Fisher) | Length | S-Sites | N-Sites | Substitutions | S-Substitutions | N-Substitutions |
|---|---|---|---|---|---|---|---|---|---|---|---|
| FtsK | MS | 0.0368 | 1.3299 | 0.0276 | 1.73E-229 | 2244 | 653.403 | 1590.6 | 430 | 402.885 | 27.1152 |
| FtsK | GMYN | 0.0365 | 1.2947 | 0.0282 | 9.82E-195 | 2244 | 602.918 | 1641.08 | 430 | 371.722 | 58.2781 |
| DnaK | MS | 0.0160 | 1.0334 | 0.0155 | 1.01E-143 | 1866 | 583.693 | 1282.31 | 277 | 267.866 | 9.13392 |
| DnaK | GMYN | 0.0160 | 0.7616 | 0.0210 | 6.21E-123 | 1866 | 585.487 | 1280.51 | 277 | 256.799 | 20.2014 |
| GroL | MS | 0.0173 | 0.6752 | 0.0256 | 5.00E-95 | 1590 | 479.487 | 1110.51 | 198 | 186.937 | 11.0633 |
| GroL | GMYN | 0.0163 | 0.7967 | 0.0204 | 2.08E-97 | 1590 | 408.032 | 1181.97 | 198 | 178.996 | 19.0042 |
| Fatty | MS | 0.0614 | 2.1489 | 0.0286 | 6.27E-186 | 1137 | 254.923 | 882.077 | 256 | 232.957 | 23.0428 |
| Fatty | GMYN | 0.0598 | 4.0524 | 0.0148 | 7.34E-151 | 1137 | 222.11 | 914.89 | 256 | 203.582 | 52.4181 |
| RuvB | MS | 0.0289 | 3.2835 | 0.0088 | 5.08E-168 | 1008 | 224.501 | 783.499 | 202 | 195.973 | 6.02699 |
| RuvB | GMYN | 0.0289 | 4.0509 | 0.0071 | 1.21E-130 | 1008 | 221.661 | 786.339 | 202 | 179.927 | 22.0727 |
| RpoH | MS | 0.0967 | 2.8548 | 0.0339 | 1.83E-142 | 885 | 232.111 | 652.889 | 222 | 202.684 | 19.3164 |
| RpoH | GMYN | 0.0880 | 4.0555 | 0.0217 | 8.64E-83 | 885 | 223.028 | 661.972 | 222 | 167.291 | 54.7091 |
| MscS | MS | 0.1795 | 3.6195 | 0.0496 | NA | 774 | 184.613 | 589.387 | 249 | 214.968 | 34.0321 |
| MscS | GMYN | 0.1792 | 3.9383 | 0.0455 | 4.08E-65 | 774 | 190.775 | 583.225 | 249 | 157.243 | 91.7568 |
| Piri | MS | 0.1342 | 4.1084 | 0.0327 | NA | 654 | 142.481 | 511.519 | 185 | 165.585 | 19.4154 |
| Piri | GMYN | 0.1383 | 3.7079 | 0.0373 | 1.97E-62 | 654 | 140.313 | 513.687 | 185 | 120.648 | 64.3521 |
| Carbo | MS | 0.0692 | 3.1345 | 0.0221 | 1.38E-109 | 624 | 144.509 | 479.491 | 142 | 132.305 | 9.6952 |
| Carbo | GMYN | 0.0699 | 3.6402 | 0.0192 | 1.67E-72 | 624 | 128.208 | 495.792 | 142 | 109.07 | 32.9296 |
| DSBA | MS | 0.2520 | 3.4307 | 0.0735 | NA | 591 | 139.069 | 451.931 | 224 | 180.83 | 43.1698 |
| DSBA | GMYN | 0.2600 | 3.6288 | 0.0717 | 2.50E-59 | 591 | 126.273 | 464.727 | 224 | 122.845 | 101.155 |
| RpoE | MS | 0.0425 | 1.6226 | 0.0262 | 1.53E-66 | 534 | 123.714 | 410.286 | 101 | 92.9237 | 8.07626 |
| RpoE | GMYN | 0.0453 | 1.4523 | 0.0312 | 2.25E-42 | 534 | 140.738 | 393.262 | 101 | 83.734 | 17.266 |
| Fimb | MS | 0.0546 | 3.8451 | 0.0142 | 4.05E-84 | 489 | 121.966 | 367.034 | 110 | 105.496 | 4.50405 |
| Fimb | GMYN | 0.0541 | 3.5853 | 0.0151 | 7.74E-54 | 489 | 119.156 | 369.844 | 110 | 90.7937 | 19.2063 |
| Rubre | MS | 0.0488 | 3.3624 | 0.0145 | 5.52E-58 | 450 | 129.181 | 320.819 | 92 | 88.8017 | 3.19828 |
| Rubre | GMYN | 0.0465 | 1.5976 | 0.0291 | 1.76E-41 | 450 | 116.512 | 333.488 | 92 | 77 | 15 |
| Alkyl | MS | 0.0440 | 1.6646 | 0.0264 | 1.07E-55 | 432 | 103.259 | 328.741 | 84 | 77.4847 | 6.51525 |
| Alkyl | GMYN | 0.0451 | 1.5451 | 0.0292 | 5.34E-39 | 432 | 106.656 | 325.344 | 84 | 69.7976 | 14.2024 |
| Hfq | MS | 0.0128 | 1.4557 | 0.0088 | 0 | 207 | 55.9527 | 151.047 | 32 | 31.2594 | 0.740612 |
| Hfq | GMYN | 0.0131 | 2.0777 | 0.0063 | 8.37E-20 | 207 | 52.0075 | 154.992 | 32 | 29.9994 | 2.00059 |

| | | | | | | | | | | | |
|---|---|---|---|---|---|---|---|---|---|---|---|
| Cold | MS | 0.0000 | 0.1806 | 0.0000 | 0 | 168 | 48.6253 | 119.375 | 7 | 6.99998 | 1.72E-05 |
| Cold | GMYN | 0.0000 | 0.2167 | 0.0000 | 0 | 168 | 40.1796 | 127.82 | 7 | 7 | 0 |
| | | | **maximum** | 0.0735 | | | | | | | |
| | | | **minimum** | 0.0000 | | | | | | | |
| | | | **average** | 0.0252 | | | | | | | |
| | | | **median** | 0.0238 | | | | | | | |

### Housekeeping genes in MWH-Weng1-1

| Sequence | Method | Ka | Ks | Ka/Ks | P-Value(Fisher) | Length | S-Sites | N-Sites | Substitutions | S-Substitutions | N-Substitutions |
|---|---|---|---|---|---|---|---|---|---|---|---|
| iso | MS | 0.0126 | 1.4558 | 0.0087 | 1.58E-164 | 1527 | 417.299 | 1109.7 | 255 | 249.242 | 5.75769 |
| iso | GMYN | 0.0121 | 1.7223 | 0.0070 | 8.02E-170 | 1527 | 357.192 | 1169.81 | 255 | 240.939 | 14.0612 |
| acsA | MS | 0.0123 | 1.2940 | 0.0095 | 1.95E-137 | 1323 | 346.255 | 976.745 | 210 | 204.51 | 5.48991 |
| acsA | GMYN | 0.0120 | 1.3882 | 0.0086 | 3.91E-135 | 1323 | 313.179 | 1009.82 | 210 | 197.983 | 12.0165 |
| succ | MS | 0.0052 | 1.2822 | 0.0041 | 6.65E-120 | 1137 | 286.368 | 850.632 | 171 | 168.951 | 2.04913 |
| succ | GMYN | 0.0047 | 1.2350 | 0.0038 | 6.38E-119 | 1137 | 275.818 | 861.182 | 171 | 167 | 4.00027 |
| CTP | MS | 0.0157 | 1.5813 | 0.0099 | 1.40E-127 | 1044 | 271.359 | 772.641 | 188 | 182.824 | 5.1758 |
| CTP | GMYN | 0.0163 | 2.9439 | 0.0055 | 1.49E-126 | 1044 | 238.02 | 805.98 | 188 | 174.992 | 13.0082 |
| atpA | MS | 0.0000 | 0.7320 | 0.0000 | 0 | 963 | 252.127 | 710.873 | 110 | 110 | 0.000310145 |
| atpA | GMYN | 0.0000 | 0.9391 | 0.0000 | 0 | 963 | 238.516 | 724.484 | 110 | 110 | 9.42E-10 |
| gyr | MS | 0.0198 | 2.5768 | 0.0077 | 6.94E-108 | 894 | 236.303 | 657.697 | 158 | 154.7 | 3.30038 |
| gyr | GMYN | 0.0201 | 1.4627 | 0.0137 | 4.18E-91 | 894 | 225.047 | 668.953 | 158 | 144.779 | 13.2208 |
| glnA | MS | 0.0174 | 1.2335 | 0.0141 | 6.92E-87 | 873 | 232.469 | 640.531 | 140 | 134.776 | 5.22418 |
| glnA | GMYN | 0.0166 | 1.4398 | 0.0116 | 8.00E-87 | 873 | 201.482 | 671.518 | 140 | 128.951 | 11.0489 |
| cit | MS | 0.0055 | 1.8270 | 0.0030 | 0 | 816 | 267.637 | 548.363 | 146 | 145.107 | 0.892631 |
| cit | GMYN | 0.0053 | 1.1520 | 0.0046 | 4.48E-88 | 816 | 242.962 | 573.038 | 146 | 142.999 | 3.00097 |
| adk | MS | 0.0077 | 1.7370 | 0.0044 | 0 | 375 | 112.06 | 262.94 | 68 | 67.3013 | 0.698709 |
| adk | GMYN | 0.0073 | 1.6208 | 0.0045 | 1.83E-45 | 375 | 99.4572 | 275.543 | 68 | 65.9998 | 2.00022 |
| ndk | MS | 0.0000 | 0.8468 | 0.0000 | 0 | 225 | 71.2132 | 153.787 | 25 | 24.9999 | 5.40E-05 |
| ndk | GMYN | NA | 0.8193 | NA | 0 | 225 | 58.0371 | 166.963 | 25 | 25 | 0 |
| | | | **maximum** | 0.0141 | | | | | | | |
| | | | **minimum** | 0.0000 | | | | | | | |
| | | | **average** | 0.0064 | | | | | | | |
| | | | **median** | 0.0055 | | | | | | | |

### Selected genes in P1-Kol8

| Sequence | Method | Ka | Ks | Ka/Ks | P-Value(Fisher) | Length | S-Sites | N-Sites | Substitutions | S-Substitutions | N-Substitutions |
|---|---|---|---|---|---|---|---|---|---|---|---|
| Fimb | MS | 0.0084 | 0.1283 | 0.0655 | 1.37E-08 | 510 | 140.079 | 369.921 | 19 | 16.199 | 2.80101 |
| Fimb | GMYN | 0.0107 | 0.1227 | 0.0876 | 1.54E-06 | 510 | 134.308 | 375.692 | 19 | 15 | 4 |
| | | | **maximum** | 0.0876 | | | | | | | |
| | | | **minimum** | 0.0655 | | | | | | | |
| | | | **average** | 0.0765 | | | | | | | |
| | | | **median** | 0.0765 | | | | | | | |

### Housekeeping genes in P1-Kol8

| Sequence | Method | Ka | Ks | Ka/Ks | P-Value(Fisher) | Length | S-Sites | N-Sites | Substitutions | S-Substitutions | N-Substitutions |
|---|---|---|---|---|---|---|---|---|---|---|---|
| iso | MS | NA | NA | NA | NA | NA | NA | NA | NA | NA | NA |
| iso | GMYN | 0.0006 | 0.0392 | 0.0152 | 0 | 2232 | 553.663 | 1678.34 | 22 | 21 | 1 |
| acsA | MS | 0.0000 | 0.0000 | 0.8609 | 0 | 1971 | 487.169 | 1483.83 | 0 | 0 | 0 |
| acsA | GMYN | 0.0000 | 0.0000 | 0.0000 | 0 | 1971 | 503.592 | 1467.41 | 0 | 0 | 0 |
| succ | MS | 0.0000 | 0.0000 | 0.8609 | 0 | 1776 | 410.966 | 1365.03 | 0 | 0 | 0 |
| succ | GMYN | 0.0000 | NA | NA | 0 | 1776 | 437.879 | 1338.12 | 0 | 0 | 0 |
| CTP | MS | 0.0000 | 0.0000 | 0.8609 | 0 | 1659 | 396.751 | 1262.25 | 0 | 0 | 0 |
| CTP | GMYN | 0.0000 | 0.0000 | NA | 0 | 1659 | 418.508 | 1240.49 | 0 | 0 | 0 |
| atpA | MS | 0.0000 | 0.0000 | 0.8609 | 0 | 1539 | 380.541 | 1158.46 | 0 | 0 | 0 |
| atpA | GMYN | NA | 0.0000 | NA | 0 | 1539 | 407.726 | 1131.27 | 0 | 0 | 0 |
| gyr | MS | 0.0000 | 0.0000 | 0.8610 | 0 | 1434 | 378.656 | 1055.34 | 0 | 0 | 0 |
| gyr | GMYN | 0.0000 | NA | NA | 0 | 1434 | 387.291 | 1046.71 | 0 | 0 | 0 |
| glnA | MS | 0.0000 | 0.0000 | 0.8607 | 0 | 1413 | 332.453 | 1080.55 | 0 | 0 | 0 |
| glnA | GMYN | 0.0000 | 0.0000 | NA | 0 | 1413 | 350.104 | 1062.9 | 0 | 0 | 0 |
| cit | MS | 0.0000 | 0.0034 | 0.0000 | 0 | 1311 | 291.309 | 1019.69 | 1 | 0.999996 | 3.50E-06 |
| cit | GMYN | 0.0000 | 0.0032 | 0.0000 | 0 | 1311 | 312.796 | 998.204 | 1 | 1 | 0 |
| adk | MS | 0.0059 | 0.1633 | 0.0364 | 3.45E-13 | 663 | 160.211 | 502.789 | 25 | 22.4396 | 2.56039 |
| adk | GMYN | 0.0061 | 0.1459 | 0.0419 | 3.77E-11 | 663 | 170.116 | 492.884 | 25 | 22 | 3 |
| ndk | MS | 0.0000 | 0.0000 | 0.8603 | 0 | 423 | 97.5858 | 325.414 | 0 | 0 | 0 |

| | | | | | | | | | | | |
|---|---|---|---|---|---|---|---|---|---|---|---|
| ndk | GMYN | NA | NA | NA | 0 | 423 | 97.8687 | 325.131 | 0 | 0 | 0 |
| | | | **maximum** | 0.8610 | | | | | | | |
| | | | **minimum** | 0.0000 | | | | | | | |
| | | | **average** | 0.4707 | | | | | | | |
| | | | **median** | 0.8603 | | | | | | | |

**Selected genes in QLW-P1DATA-2**

| Sequence | Method | Ka | Ks | Ka/Ks | P-Value(Fisher) | Length | S-Sites | N-Sites | Substitutions | S-Substitutions | N-Substitutions |
|---|---|---|---|---|---|---|---|---|---|---|---|
| Alkyl | MS | 0.0309 | 2.5430 | 0.0121 | 3.46E-65 | 444 | 108.251 | 335.749 | 88 | 84.8045 | 3.19553 |
| Alkyl | GMYN | 0.0333 | 4.3104 | 0.0077 | 4.72E-51 | 444 | 102.89 | 341.11 | 88 | 76.9209 | 11.0791 |
| Carbo | MS | 0.0217 | 1.6203 | 0.0134 | 1.31E-73 | 639 | 169.63 | 469.37 | 114 | 109.927 | 4.07271 |
| Carbo | GMYN | 0.0215 | 1.4251 | 0.0151 | 3.55E-64 | 639 | 162.959 | 476.041 | 114 | 103.941 | 10.0589 |
| Cold | MS | 0.0000 | 0.0495 | 0.0000 | 0 | 180 | 40.3807 | 139.619 | 2 | 1.99999 | 6.92E-06 |
| Cold | GMYN | 0.0000 | 0.0516 | 0.0000 | 0 | 180 | 42.6501 | 137.35 | 2 | 2 | 0 |
| DnaK | MS | 0.0121 | 0.7511 | 0.0161 | 3.00E-127 | 1890 | 547.109 | 1342.89 | 238 | 228.969 | 9.03078 |
| DnaK | GMYN | 0.0116 | 0.9016 | 0.0128 | 2.25E-129 | 1890 | 483.897 | 1406.1 | 238 | 221.917 | 16.0828 |
| DSBA | MS | 0.1276 | 2.9315 | 0.0435 | NA | 603 | 138.561 | 464.439 | 163 | 142.251 | 20.7487 |
| DSBA | GMYN | 0.1248 | 3.6412 | 0.0343 | 1.15E-57 | 603 | 128.368 | 474.632 | 163 | 108.823 | 54.1775 |
| Fatty | MS | 0.0647 | 2.4607 | 0.0263 | 1.27E-190 | 1158 | 264.889 | 893.111 | 261 | 239.749 | 21.2512 |
| Fatty | GMYN | 0.0629 | 4.1281 | 0.0152 | 1.96E-134 | 1158 | 245.709 | 912.291 | 261 | 206.176 | 54.8241 |
| Fimb | MS | 0.0559 | 3.8195 | 0.0146 | 1.50E-86 | 498 | 124.921 | 373.079 | 113 | 108.266 | 4.73378 |
| Fimb | GMYN | 0.0562 | 3.6049 | 0.0156 | 4.11E-54 | 498 | 122.299 | 375.701 | 113 | 92.7167 | 20.2833 |
| FtsK | MS | 0.0150 | 1.0321 | 0.0146 | 5.19E-209 | 2268 | 620.198 | 1647.8 | 353 | 339.852 | 13.1481 |
| FtsK | GMYN | 0.0140 | 1.3271 | 0.0105 | 1.31E-194 | 2268 | 602.647 | 1665.35 | 353 | 329.962 | 23.0377 |
| GroL | MS | 0.0060 | 0.5348 | 0.0113 | 1.07E-88 | 1611 | 457.404 | 1153.6 | 160 | 155.571 | 4.4293 |
| GroL | GMYN | 0.0058 | 0.6384 | 0.0091 | 7.74E-94 | 1611 | 398.76 | 1212.24 | 160 | 152.999 | 7.00051 |
| Piri | MS | 0.1350 | 3.1715 | 0.0426 | NA | 675 | 144.253 | 530.747 | 176 | 152.168 | 23.8322 |
| Piri | GMYN | 0.1387 | 6.1000 | 0.0227 | 6.44E-48 | 675 | 146.142 | 528.858 | 176 | 109.574 | 66.426 |
| RpoE | MS | NA | NA | NA | NA | NA | NA | NA | NA | NA | NA |
| RpoE | GMYN | 0.0305 | 0.7767 | 0.0393 | 3.39E-31 | 546 | 136.588 | 409.412 | 74 | 61.7805 | 12.2195 |
| RpoH | MS | 0.0438 | 1.2207 | 0.0359 | 1.02E-82 | 894 | 254.367 | 639.633 | 162 | 148.596 | 13.4035 |
| RpoH | GMYN | 0.0413 | 1.2961 | 0.0318 | 6.98E-73 | 894 | 218.78 | 675.22 | 162 | 134.941 | 27.0586 |

| | | | | | | | | | | | |
|---|---|---|---|---|---|---|---|---|---|---|---|
| Rubre | MS | 0.0367 | 2.7185 | 0.0135 | 6.40E-50 | 462 | 147.886 | 314.114 | 88 | 85.5493 | 2.45071 |
| Rubre | GMYN | 0.0345 | 1.7929 | 0.0192 | 2.66E-38 | 462 | 134.721 | 327.279 | 88 | 77 | 11 |
| RuvB | MS | 0.0095 | 1.2635 | 0.0075 | 9.59E-112 | 1029 | 286.963 | 742.037 | 174 | 170.69 | 3.31038 |
| RuvB | GMYN | 0.0094 | 1.8698 | 0.0050 | 1.27E-105 | 1029 | 278.984 | 750.016 | 174 | 166.992 | 7.00753 |
| | | | **maximum** | 0.0435 | | | | | | | |
| | | | **minimum** | 0.0000 | | | | | | | |
| | | | **average** | 0.0181 | | | | | | | |
| | | | **median** | 0.0146 | | | | | | | |

### Housekeeping genes in QLW-P1DATA-2

| Sequence | Method | Ka | Ks | Ka/Ks | P-Value(Fisher) | Length | S-Sites | N-Sites | Substitutions | S-Substitutions | N-Substitutions |
|---|---|---|---|---|---|---|---|---|---|---|---|
| iso | MS | 0.0223 | 0.9998 | 0.0223 | 1.13E-190 | 2232 | 596.507 | 1635.49 | 339 | 319.447 | 19.5526 |
| iso | GMYN | 0.0222 | 1.3559 | 0.0164 | 2.72E-166 | 2232 | 575.916 | 1656.08 | 339 | 302.855 | 36.1454 |
| acsA | MS | 0.0303 | 0.8617 | 0.0351 | 6.64E-147 | 1971 | 493.945 | 1477.05 | 281 | 254.288 | 26.7124 |
| acsA | GMYN | 0.0272 | 1.5565 | 0.0175 | 3.77E-141 | 1971 | 432.585 | 1538.41 | 281 | 239.927 | 41.0729 |
| succ | MS | 0.0126 | 0.8678 | 0.0145 | 1.08E-139 | 1776 | 459.187 | 1316.81 | 233 | 223.668 | 9.33209 |
| succ | GMYN | 0.0111 | 1.6003 | 0.0069 | 3.14E-141 | 1776 | 413.624 | 1362.38 | 233 | 218 | 15.0005 |
| CTP | MS | 0.0291 | 1.8451 | 0.0158 | 1.39E-216 | 1659 | 396.265 | 1262.73 | 309 | 294.203 | 14.7971 |
| CTP | GMYN | 0.0292 | 2.0363 | 0.0143 | 9.17E-174 | 1659 | 389.391 | 1269.61 | 309 | 272.709 | 36.2908 |
| atpA | MS | 0.0029 | 0.2471 | 0.0119 | 5.50E-48 | 1539 | 416.33 | 1122.67 | 86 | 83.3313 | 2.66866 |
| atpA | GMYN | 0.0026 | 0.2821 | 0.0092 | 8.33E-50 | 1539 | 378.591 | 1160.41 | 86 | 82.9997 | 3.00028 |
| gyr | MS | 0.0292 | 1.8620 | 0.0157 | 1.38E-164 | 1434 | 412.584 | 1021.42 | 275 | 264.715 | 10.2845 |
| gyr | GMYN | 0.0295 | 1.9973 | 0.0148 | 5.21E-126 | 1434 | 417.706 | 1016.29 | 275 | 245.663 | 29.3372 |
| glnA | MS | 0.0163 | 0.4674 | 0.0349 | 3.72E-65 | 1413 | 369.219 | 1043.78 | 136 | 123.8 | 12.2004 |
| glnA | GMYN | 0.0140 | 0.8743 | 0.0161 | 5.05E-67 | 1413 | 332.257 | 1080.74 | 136 | 120.992 | 15.0082 |
| cit | MS | 0.0140 | 0.8290 | 0.0169 | 3.17E-103 | 1311 | 318.87 | 992.13 | 168 | 159.61 | 8.38997 |
| cit | GMYN | 0.0128 | 2.1119 | 0.0061 | 3.95E-105 | 1311 | 281.381 | 1029.62 | 168 | 154.948 | 13.0524 |
| adk | MS | 0.0281 | 0.8887 | 0.0316 | 2.92E-49 | 663 | 190.603 | 472.397 | 96 | 89.0224 | 6.97763 |
| adk | GMYN | 0.0270 | 1.0241 | 0.0263 | 2.16E-43 | 663 | 169.874 | 493.126 | 96 | 82.9971 | 13.0029 |
| ndk | MS | 0.0029 | 0.4709 | 0.0063 | 0 | 423 | 87.6668 | 335.333 | 29 | 28.3217 | 0.678306 |
| ndk | GMYN | 0.0030 | 0.4702 | 0.0065 | 3.64E-19 | 423 | 94.1205 | 328.879 | 29 | 28 | 1 |
| | | | **maximum** | 0.0351 | | | | | | | |
| | | | **minimum** | 0.0061 | | | | | | | |

| | |
|---|---|
| **average** | 0.0170 |
| **median** | 0.0157 |

### Selected genes in QLW-P1FAT50C-4

| Sequence | Method | Ka | Ks | Ka/Ks | P-Value(Fisher) | Length | S-Sites | N-Sites | Substitutions | S-Substitutions | N-Substitutions |
|---|---|---|---|---|---|---|---|---|---|---|---|
| FtsK | MS | 0.0125 | 1.0848 | 0.0115 | 8.69E-202 | 2202 | 661.527 | 1540.47 | 356 | 346.733 | 9.26654 |
| FtsK | GMYN | 0.0123 | 1.4236 | 0.0086 | 3.02E-189 | 2202 | 635.725 | 1566.27 | 356 | 336.975 | 19.0248 |
| Alkyl | MS | 0.0224 | 1.7097 | 0.0131 | 9.32E-46 | 363 | 80.6686 | 282.331 | 60 | 57.3708 | 2.62924 |
| Alkyl | GMYN | 0.0258 | 2.3727 | 0.0109 | 8.14E-33 | 363 | 86.3952 | 276.605 | 60 | 52.9968 | 7.00321 |
| Carbo | MS | 0.0266 | 2.0222 | 0.0131 | 4.16E-79 | 558 | 127.269 | 430.731 | 104 | 99.5746 | 4.42543 |
| Carbo | GMYN | 0.0295 | 1.6056 | 0.0183 | 9.12E-56 | 558 | 138.878 | 419.122 | 104 | 91.9133 | 12.0867 |
| Cold | MS | 0.0000 | 0.0816 | 0.0000 | 0 | 99 | 24.5139 | 74.4861 | 2 | 1.99999 | 6.08E-06 |
| Cold | GMYN | 0.0000 | 0.0818 | 0.0000 | 0 | 99 | 28.0974 | 70.9026 | 2 | 2 | 0 |
| DnaK | MS | 0.0109 | 0.7159 | 0.0152 | 2.12E-122 | 1824 | 524.785 | 1299.22 | 226 | 217.802 | 8.19835 |
| DnaK | GMYN | 0.0102 | 1.6663 | 0.0061 | 1.63E-134 | 1824 | 431.263 | 1392.74 | 226 | 211.953 | 14.0469 |
| DSBA | MS | 0.1384 | 3.8109 | 0.0363 | NA | 522 | 122.466 | 399.534 | 153 | 136.792 | 16.208 |
| DSBA | GMYN | 0.1401 | 3.6040 | 0.0389 | 3.66E-50 | 522 | 122.167 | 399.833 | 153 | 102.162 | 50.8375 |
| Fatty | MS | 0.0597 | 2.1779 | 0.0274 | 6.46E-171 | 1092 | 249.207 | 842.793 | 240 | 219.643 | 20.3566 |
| Fatty | GMYN | 0.0590 | 4.0728 | 0.0145 | 2.00E-127 | 1092 | 228.241 | 863.759 | 240 | 191.199 | 48.8012 |
| Fimb | MS | 0.0307 | 3.5496 | 0.0087 | 3.34E-64 | 417 | 114.861 | 302.139 | 91 | 88.974 | 2.02604 |
| Fimb | GMYN | 0.0311 | 3.0580 | 0.0102 | 4.80E-51 | 417 | 110.971 | 306.029 | 91 | 81.8172 | 9.18276 |
| GroL | MS | 0.0099 | 0.6139 | 0.0162 | 9.83E-92 | 1545 | 435.919 | 1109.08 | 169 | 162.326 | 6.67375 |
| GroL | GMYN | 0.0095 | 0.7438 | 0.0128 | 2.28E-94 | 1545 | 378.873 | 1166.13 | 169 | 157.998 | 11.0017 |
| Hfq | MS | 0.0309 | 0.5873 | 0.0527 | 2.46E-08 | 144 | 42.0409 | 101.959 | 18 | 15.9613 | 2.0387 |
| Hfq | GMYN | 0.0292 | 0.7239 | 0.0403 | 7.47E-08 | 144 | 37.722 | 106.278 | 18 | 15 | 3 |
| Hyp | MS | 0.0329 | 4.2779 | 0.0077 | 0 | 162 | 36.9056 | 125.094 | 33 | 32.1607 | 0.839256 |
| Hyp | GMYN | 0.0335 | 3.0362 | 0.0110 | 3.22E-19 | 162 | 39.3534 | 122.647 | 33 | 29 | 4 |
| Piri | MS | 0.1626 | 3.5973 | 0.0452 | NA | 594 | 144.815 | 449.185 | 176 | 154.353 | 21.647 |
| Piri | GMYN | 0.1614 | 3.6715 | 0.0440 | 4.43E-49 | 594 | 133.658 | 460.342 | 176 | 109.705 | 66.2949 |
| RpoE | MS | 0.0224 | 0.8582 | 0.0261 | 3.76E-38 | 465 | 103.751 | 361.249 | 62 | 56.8306 | 5.16941 |
| RpoE | GMYN | 0.0228 | 1.1131 | 0.0205 | 3.71E-32 | 465 | 106.014 | 358.986 | 62 | 53.9583 | 8.04168 |
| RpoH | MS | 0.0333 | 1.2678 | 0.0263 | 3.24E-79 | 822 | 231.935 | 590.065 | 145 | 135.908 | 9.09161 |

| | | | | | | | | | | | |
|---|---|---|---|---|---|---|---|---|---|---|---|
| RpoH | GMYN | 0.0309 | 1.6432 | 0.0188 | 1.47E-78 | 822 | 188.964 | 633.036 | 145 | 125.878 | 19.1225 |
| Rubre | MS | 0.0362 | 3.0609 | 0.0118 | 1.38E-42 | 381 | 121.404 | 259.596 | 72 | 70.2252 | 1.7748 |
| Rubre | GMYN | 0.0357 | 1.0941 | 0.0326 | 2.11E-29 | 381 | 122.203 | 258.797 | 72 | 63 | 9 |
| RuvB | MS | 0.0161 | 1.5556 | 0.0104 | 3.38E-115 | 963 | 272.316 | 690.684 | 180 | 175.384 | 4.61607 |
| RuvB | GMYN | 0.0161 | 1.3747 | 0.0117 | 1.25E-101 | 963 | 268.211 | 694.789 | 180 | 168.983 | 11.0174 |
| Superox | MS | 0.0476 | 2.2384 | 0.0213 | 3.12E-56 | 429 | 118.796 | 310.204 | 89 | 84.3182 | 4.68179 |
| Superox | GMYN | 0.0486 | 2.4670 | 0.0197 | 4.26E-34 | 429 | 128.64 | 300.36 | 89 | 74.9082 | 14.0918 |
| | | | **maximum** | 0.0527 | | | | | | | |
| | | | **minimum** | 0.0000 | | | | | | | |
| | | | **average** | 0.0195 | | | | | | | |
| | | | **median** | 0.0148 | | | | | | | |

### Housekeeping genes in QLW-P1FAT50C-4

| Sequence | Method | Ka | Ks | Ka/Ks | P-Value(Fisher) | Length | S-Sites | N-Sites | Substitutions | S-Substitutions | N-Substitutions |
|---|---|---|---|---|---|---|---|---|---|---|---|
| iso | MS | 0.0194 | 0.9274 | 0.0209 | 3.43E-180 | 2232 | 612.155 | 1619.84 | 327 | 309.838 | 17.1624 |
| iso | GMYN | 0.0190 | 1.2797 | 0.0149 | 1.91E-167 | 2232 | 569.193 | 1662.81 | 327 | 295.802 | 31.1981 |
| acsA | MS | 0.0254 | 0.9576 | 0.0265 | 4.92E-162 | 1971 | 532.638 | 1438.36 | 299 | 279 | 19.9996 |
| acsA | GMYN | 0.0241 | 2.8319 | 0.0085 | 2.40E-161 | 1971 | 444.845 | 1526.16 | 299 | 262.838 | 36.1616 |
| succ | MS | 0.0228 | 0.7962 | 0.0287 | 6.53E-126 | 1776 | 506.432 | 1269.57 | 250 | 233.227 | 16.7728 |
| succ | GMYN | 0.0216 | 1.7305 | 0.0125 | 1.09E-131 | 1776 | 408.645 | 1367.35 | 250 | 220.971 | 29.029 |
| CTP | MS | 0.0287 | 1.4080 | 0.0204 | 5.77E-183 | 1659 | 410.376 | 1248.62 | 288 | 271.188 | 16.812 |
| CTP | GMYN | 0.0287 | 1.3198 | 0.0218 | 1.33E-147 | 1659 | 407.071 | 1251.93 | 288 | 252.764 | 35.2364 |
| atpA | MS | 0.0069 | 0.3281 | 0.0210 | 2.25E-58 | 1539 | 414.089 | 1124.91 | 111 | 105.009 | 5.99135 |
| atpA | GMYN | 0.0063 | 0.3240 | 0.0194 | 4.47E-54 | 1539 | 420.3 | 1118.7 | 111 | 104 | 7 |
| gyr | MS | 0.0294 | 2.9137 | 0.0101 | 9.18E-210 | 1434 | 389.71 | 1044.29 | 302 | 294.044 | 7.95621 |
| gyr | GMYN | 0.0295 | 2.2954 | 0.0128 | 1.71E-164 | 1434 | 379.947 | 1054.05 | 302 | 271.606 | 30.3935 |
| glnA | MS | 0.0009 | 0.3288 | 0.0029 | 0 | 1413 | 376.781 | 1036.22 | 93 | 92.2678 | 0.732182 |
| glnA | GMYN | 0.0010 | 0.3391 | 0.0028 | 1.14E-57 | 1413 | 360.354 | 1052.65 | 93 | 91.9999 | 1.00008 |
| cit | MS | 0.0228 | 0.6920 | 0.0329 | 1.20E-84 | 1311 | 342.568 | 968.432 | 167 | 152.781 | 14.2187 |
| cit | GMYN | 0.0225 | 0.9200 | 0.0244 | 3.22E-78 | 1311 | 316.177 | 994.823 | 167 | 144.987 | 22.0131 |
| adk | MS | 0.0266 | 1.0241 | 0.0260 | 1.51E-54 | 663 | 196.497 | 466.503 | 103 | 97.0207 | 5.97934 |
| adk | GMYN | 0.0245 | 1.9126 | 0.0128 | 6.89E-54 | 663 | 159.511 | 503.489 | 103 | 90.8916 | 12.1084 |
| ndk | MS | 0.0097 | 0.4362 | 0.0222 | 9.09E-21 | 423 | 116.198 | 306.802 | 36 | 34.0073 | 1.99266 |

| | | | | | | | | | | | |
|---|---|---|---|---|---|---|---|---|---|---|---|
| ndk | GMYN | 0.0093 | 0.5018 | 0.0186 | 4.30E-20 | 423 | 98.377 | 324.623 | 36 | 33 | 3 |
| | | | **maximum** | 0.0329 | | | | | | | |
| | | | **minimum** | 0.0028 | | | | | | | |
| | | | **average** | 0.0180 | | | | | | | |
| | | | **median** | 0.0199 | | | | | | | |

### Selected genes in STIR1

| Sequence | Method | Ka | Ks | Ka/Ks | P-Value(Fisher) | Length | S-Sites | N-Sites | Substitutions | S-Substitutions | N-Substitutions |
|---|---|---|---|---|---|---|---|---|---|---|---|
| Alkyl | MS | 0.0374733 | 4.36149 | 0.00859187 | 4.13E-71 | 387 | 86.1463 | 300.854 | 81 | 78.6403 | 2.35967 |
| Alkyl | GMYN | 0.0421388 | 3.35097 | 0.0125751 | 2.70E-46 | 387 | 87.1785 | 299.822 | 81 | 68.7847 | 12.2153 |
| Carbo | MS | NA | NA | NA | NA | NA | NA | NA | NA | NA | NA |
| Carbo | GMYN | 0.0806866 | 3.5869 | 0.0224948 | 5.76E-63 | 522 | 119.406 | 402.594 | 133 | 102.329 | 30.6713 |
| Cold | MS | 5.04E-08 | 0.0504104 | 1.00E-06 | 0 | 195 | 65.3257 | 129.674 | 3 | 2.99999 | 5.96E-06 |
| Cold | GMYN | 1.11E-15 | 0.0675801 | 1.64E-14 | 0 | 195 | 47.762 | 147.238 | 3 | 3 | 0 |
| DnaK | MS | 0.0176324 | 0.752496 | 0.0234318 | 2.69E-118 | 1764 | 495.011 | 1268.99 | 226 | 213.194 | 12.8063 |
| DnaK | GMYN | 0.0162898 | 1.39952 | 0.0116396 | 1.41E-128 | 1764 | 390.355 | 1373.64 | 226 | 203.891 | 22.1088 |
| Fatty | MS | 0.0686111 | 1.57088 | 0.0436769 | 1.40E-131 | 1050 | 263.549 | 786.451 | 231 | 204.364 | 26.6359 |
| Fatty | GMYN | 0.0674697 | 2.72085 | 0.0247973 | 1.48E-97 | 1050 | 245.528 | 804.472 | 231 | 179.253 | 51.7474 |
| FtsK | MS | 0.0186167 | 1.07178 | 0.0173699 | 9.31E-196 | 2142 | 623.939 | 1518.06 | 352 | 337.727 | 14.2728 |
| FtsK | GMYN | 0.0191907 | 1.01163 | 0.01897 | 1.43E-159 | 2142 | 655.893 | 1486.11 | 352 | 323.879 | 28.1214 |
| GroL | MS | 0.0132982 | 0.594931 | 0.0223525 | 7.80E-85 | 1494 | 418.369 | 1075.63 | 163 | 154.142 | 8.85828 |
| GroL | GMYN | 0.0126462 | 0.531829 | 0.0237787 | 2.22E-69 | 1494 | 456.241 | 1037.76 | 163 | 149.998 | 13.0021 |
| Hfq | MS | 0.0118487 | 0.954838 | 0.0124092 | 2.40E-20 | 234 | 61.682 | 172.318 | 32 | 30.9278 | 1.07217 |
| Hfq | GMYN | 0.0112376 | 2.02923 | 0.00553787 | 4.09E-21 | 234 | 53.0886 | 180.911 | 32 | 29.9916 | 2.00839 |
| RpoE | MS | 0.142481 | 3.63299 | 0.0392187 | NA | 426 | 95.0634 | 330.937 | 111 | 97.6658 | 13.3342 |
| RpoE | GMYN | 0.150827 | 1.68973 | 0.0892605 | 1.74E-24 | 426 | 99.2193 | 326.781 | 111 | 66.5945 | 44.4055 |
| RpoH | MS | 0.0368315 | 1.0868 | 0.03389 | 3.08E-69 | 780 | 218.821 | 561.179 | 133 | 122.365 | 10.6351 |
| RpoH | GMYN | 0.0360859 | 1.24944 | 0.0288816 | 5.77E-56 | 780 | 209.234 | 570.766 | 133 | 112.938 | 20.062 |
| RuvB | MS | 0.0222919 | 2.61062 | 0.00853893 | 3.84E-144 | 921 | 191.99 | 729.01 | 169 | 163.693 | 5.30748 |
| RuvB | GMYN | 0.0233258 | 1.98957 | 0.0117241 | 1.43E-100 | 921 | 219.415 | 701.585 | 169 | 152.967 | 16.0331 |
| | | | **maximum** | 0.0893 | | | | | | | |
| | | | **minimum** | 0.0000 | | | | | | | |

| | |
|---|---|
| **average** | 0.0219 |
| **median** | 0.0190 |

### Housekeeping genes in STIR1

| Sequence | Method | Ka | Ks | Ka/Ks | P-Value(Fisher) | Length | S-Sites | N-Sites | Substitutions | S-Substitutions | N-Substitutions |
|---|---|---|---|---|---|---|---|---|---|---|---|
| iso | MS | 0.0246 | 0.8577 | 0.0286 | 5.64E-158 | 2148 | 607.868 | 1540.13 | 313 | 291.834 | 21.166 |
| iso | GMYN | 0.0244 | 0.8963 | 0.0272 | 3.15E-134 | 2148 | 596.948 | 1551.05 | 313 | 275.79 | 37.2096 |
| acsA | MS | 0.0365 | 0.9353 | 0.0390 | 4.32E-149 | 1887 | 507.18 | 1379.82 | 300 | 271.218 | 28.782 |
| acsA | GMYN | 0.0356 | 1.4655 | 0.0243 | 3.08E-127 | 1887 | 469.867 | 1417.13 | 300 | 250.775 | 49.2249 |
| atpA | MS | NA | NA | NA | NA | NA | NA | NA | NA | NA | NA |
| atpA | GMYN | 0.0118 | 0.3687 | 0.0320 | 5.77E-48 | 1470 | 357.029 | 1112.97 | 107 | 94 | 13 |
| gyr | MS | 0.0484 | 3.6212 | 0.0134 | 1.86E-215 | 1365 | 363.563 | 1001.44 | 305 | 294.174 | 10.8262 |
| gyr | GMYN | 0.0496 | 2.0975 | 0.0237 | 2.78E-138 | 1365 | 365.201 | 999.799 | 305 | 257.189 | 47.8108 |
| glnA | MS | 0.0110 | 0.4696 | 0.0234 | 9.82E-66 | 1344 | 344.318 | 999.682 | 122 | 114.235 | 7.76513 |
| glnA | GMYN | 0.0107 | 0.5838 | 0.0184 | 1.10E-64 | 1344 | 311.139 | 1032.86 | 122 | 110.994 | 11.0057 |
| cit | MS | 0.0190 | 0.6864 | 0.0277 | 1.37E-78 | 1242 | 345.667 | 896.333 | 155 | 144.602 | 10.3981 |
| cit | GMYN | 0.0175 | 1.6527 | 0.0106 | 2.62E-89 | 1242 | 259.875 | 982.125 | 155 | 137.996 | 17.0036 |
| adk | MS | 0.0336 | 1.2359 | 0.0272 | 4.20E-55 | 594 | 152.53 | 441.47 | 94 | 87.1361 | 6.86386 |
| adk | GMYN | 0.0320 | 0.8913 | 0.0359 | 2.95E-37 | 594 | 172.334 | 421.666 | 94 | 80.8497 | 13.1503 |
| ndk | MS | 0.0108 | 0.3335 | 0.0325 | 8.68E-12 | 354 | 84.6678 | 269.332 | 22 | 19.9386 | 2.06145 |
| ndk | GMYN | 0.0112 | 0.2822 | 0.0398 | 1.89E-10 | 354 | 83.7145 | 270.285 | 22 | 19 | 3 |
| | | | **maximum** | 0.0398 | | | | | | | |
| | | | **minimum** | 0.0106 | | | | | | | |
| | | | **average** | 0.0269 | | | | | | | |
| | | | **median** | 0.0272 | | | | | | | |

### Selected genes in Tro8F10W22

| Sequence | Method | Ka | Ks | Ka/Ks | P-Value(Fisher) | Length | S-Sites | N-Sites | Substitutions | S-Substitutions | N-Substitutions |
|---|---|---|---|---|---|---|---|---|---|---|---|
| GroL | MS | 0.0009 | 0.0000 | 50.0000 | 0 | 1650 | 524.594 | 1125.41 | 1 | 0.00923665 | 0.990763 |
| GroL | GMYN | 0.0008 | 0.0000 | 2.5E+11 | 0 | 1650 | 428.744 | 1221.26 | 1 | 0 | 1 |
| Alkyl | MS | 0.0032 | 0.0730 | 0.0439 | 0 | 465 | 149.252 | 315.748 | 11 | 10.0655 | 0.93448 |

| | | | | | | | | | | | |
|---|---|---|---|---|---|---|---|---|---|---|---|
| Alkyl | GMYN | 0.0030 | 0.0841 | 0.0356 | 1.76E-05 | 465 | 129.538 | 335.462 | 11 | 10 | 1 |
| Fimb | MS | 0.0359 | 0.6772 | 0.0530 | 3.91E-30 | 510 | 134.073 | 375.927 | 66 | 57.4616 | 8.53837 |
| Fimb | GMYN | 0.0355 | 0.7096 | 0.0500 | 8.03E-24 | 510 | 131.263 | 378.737 | 66 | 52.9318 | 13.0682 |
| RpoE | MS | 0.0000 | 0.0076 | 0.0000 | 0 | 567 | 131.79 | 435.21 | 1 | 0.999997 | 3.30E-06 |
| RpoE | GMYN | 0.0000 | 0.0071 | 0.0000 | 0 | 567 | 142.392 | 424.608 | 1 | 1 | 0 |
| Superox | MS | 0.0034 | 0.0001 | 50.0000 | 0 | 531 | 235.19 | 295.81 | 1 | 0.0156525 | 0.984347 |
| Superox | GMYN | 0.0025 | NA | NA | 0 | 531 | 130.855 | 400.145 | 1 | 0 | 1 |
| | | | **maximum** | 2.5E+11 | | | | | | | |
| | | | **minimum** | 0.0000 | | | | | | | |
| | | | **average** | 2.7E+10 | | | | | | | |
| | | | **median** | 0.0500 | | | | | | | |

### Housekeeping genes in Tro8F10W22

| Sequence | Method | Ka | Ks | Ka/Ks | P-Value(Fisher) | Length | S-Sites | N-Sites | Substitutions | S-Substitutions | N-Substitutions |
|---|---|---|---|---|---|---|---|---|---|---|---|
| iso | MS | 0.0000 | 0.0000 | 0.8609 | 0 | 2232 | 517.539 | 1714.46 | 0 | 0 | 0 |
| iso | GMYN | NA | 0.0000 | NA | 0 | 2232 | 555.654 | 1676.35 | 0 | 0 | 0 |
| acsA | MS | 0.0007 | 0.0017 | 0.4511 | 0 | 1971 | 626.959 | 1344.04 | 2 | 1.0167 | 0.983301 |
| acsA | GMYN | 0.0007 | 0.0020 | 0.3417 | 0.38025 | 1971 | 502.821 | 1468.18 | 2 | 1 | 1 |
| succ | MS | 0.0000 | 0.0000 | 0.8609 | 0 | 1776 | 410.966 | 1365.03 | 0 | 0 | 0 |
| succ | GMYN | 0.0000 | NA | NA | 0 | 1776 | 437.879 | 1338.12 | 0 | 0 | 0 |
| CTP | MS | 0.0000 | 0.0000 | 0.8609 | 0 | 1659 | 396.751 | 1262.25 | 0 | 0 | 0 |
| CTP | GMYN | 0.0000 | 0.0000 | NA | 0 | 1659 | 418.508 | 1240.49 | 0 | 0 | 0 |
| atpA | MS | 0.0010 | 0.0058 | 0.1735 | 0 | 1539 | 533.218 | 1005.78 | 4 | 3.01356 | 0.986439 |
| atpA | GMYN | 0.0009 | 0.0074 | 0.1194 | 0.0544848 | 1539 | 407.728 | 1131.27 | 4 | 3 | 1 |
| gyr | MS | 0.0000 | 0.0000 | 0.8610 | 0 | 1434 | 378.656 | 1055.34 | 0 | 0 | 0 |
| gyr | GMYN | 0.0000 | NA | NA | 0 | 1434 | 387.291 | 1046.71 | 0 | 0 | 0 |
| glnA | MS | 0.0000 | 0.0000 | 0.8607 | 0 | 1413 | 332.453 | 1080.55 | 0 | 0 | 0 |
| glnA | GMYN | 0.0000 | 0.0000 | NA | 0 | 1413 | 350.104 | 1062.9 | 0 | 0 | 0 |
| cit | MS | 0.0000 | 0.0034 | 0.0000 | 0 | 1311 | 291.309 | 1019.69 | 1 | 0.999996 | 3.50E-06 |
| cit | GMYN | 0.0000 | 0.0032 | 0.0000 | 0 | 1311 | 312.796 | 998.204 | 1 | 1 | 0 |
| adk | MS | 0.0079 | 0.1992 | 0.0397 | 6.71E-15 | 663 | 159.472 | 503.528 | 30 | 26.6596 | 3.34038 |
| adk | GMYN | 0.0078 | 0.2078 | 0.0375 | 1.59E-14 | 663 | 146.132 | 516.868 | 30 | 26 | 4 |
| ndk | MS | 0.0000 | 0.0102 | 0.0000 | 0 | 423 | 98.1022 | 324.898 | 1 | 0.999997 | 3.31E-06 |

| | | | | | | | | | | | |
|---|---|---|---|---|---|---|---|---|---|---|---|
| ndk | GMYN | 0.0000 | 0.0103 | 0.0000 | 0 | 423 | 98.4044 | 324.596 | 1 | 1 | 0 |
| | | | **maximum** | 0.8610 | | | | | | | |
| | | | **minimum** | 0.0000 | | | | | | | |
| | | | **average** | 0.3645 | | | | | | | |
| | | | **median** | 0.1735 | | | | | | | |

GMYN = Approximate method (γ-MYN)
MS = Maximum-likelihood method

**Supplementary Table S10.** Ka/Ks analysis of highly expressed genes and housekeeping genes in selected strains of genus *Polynucleobacter*. Ka/Ks values were calculated twice; once by approximate method (γ-MYN) and then using a maximum-likelihood method (MS, i.e. the substitution model with the best score was selected).

### Supplementary Table S11.

| Genomes | Ka | Ks | Ka/Ks | Length | S-Sites | N-Sites | Substitutions | S-Substitutions | N-Substitutions |
|---|---|---|---|---|---|---|---|---|---|
| QLW-P1DMWA-1 vs. MWH-Tro7-1-4 | 0.00195056 | 0.023324 | 0.083629 | 1706745 | 454441 | 1.25E+06 | 12836 | 10396.8 | 2439.17 |
| P1-4-10KL vs. MWH-Tro7-1-4 | 0.00196293 | 0.0232851 | 0.0842999 | 1706853 | 455223 | 1.25E+06 | 12851 | 10397.7 | 2453.29 |
| P1-Kol8 vs. MWH-Tro7-1-4 | 0.00198399 | 0.0234529 | 0.0845945 | 1706637 | 457937 | 1.25E+06 | 13007 | 10533.2 | 2473.75 |
| MWH-Recht1 vs. MWH-Tro7-1-4 | 0.00221659 | 0.0258409 | 0.0857784 | 1706475 | 453292 | 1.25E+06 | 14240 | 11466.7 | 2773.26 |
| MWH-Tro7-1-4 vs. Tro8F10W22 | 0.00244588 | 0.0284838 | 0.0858691 | 1705725 | 455481 | 1.25E+06 | 15724 | 12671.6 | 3052.41 |
| MWH-Tro7-1-4 vs. MWH-Tro8-2-9 | 0.00202447 | 0.023507 | 0.0861219 | 1703907 | 455736 | 1.25E+06 | 13029 | 10505.9 | 2523.1 |
| P1-Kol8 vs. MWH-Recht1 | 0.00104664 | 0.0121318 | 0.086273 | 1707705 | 453078 | 1.25E+06 | 6754 | 5441.87 | 1312.13 |
| QLW-P1DMWA-1 vs. P1-Kol8 | 0.000725487 | 0.00834921 | 0.0868929 | 1707858 | 457364 | 1.25E+06 | 4699 | 3792.27 | 906.726 |
| QLW-P1DMWA-1 vs. MWH-Recht1 | 0.0012077 | 0.0138429 | 0.0872431 | 1707726 | 449003 | 1.26E+06 | 7664 | 6145.19 | 1518.81 |
| MWH-Recht1 vs. Tro8F10W22 | 0.00249435 | 0.0284729 | 0.0876046 | 1705869 | 456741 | 1.25E+06 | 15813 | 12702.9 | 3110.06 |
| P1-4-10KL vs. MWH-Recht1 | 0.0012184 | 0.01381 | 0.0882259 | 1707789 | 450122 | 1.26E+06 | 7677 | 6146.03 | 1530.97 |
| P1-Kol8 vs. P1-4-10KL | 0.000736141 | 0.00831726 | 0.0885076 | 1707975 | 459092 | 1.25E+06 | 4711 | 3792.15 | 918.849 |
| MWH-Recht1 vs. MWH-Tro8-2-9 | 0.00144906 | 0.0162686 | 0.0890708 | 1704606 | 459228 | 1.25E+06 | 9174 | 7371.3 | 1802.7 |
| P1-Kol8 vs. Tro8F10W22 | 0.00238612 | 0.0264698 | 0.0901451 | 1705965 | 457107 | 1.25E+06 | 14813 | 11838.3 | 2974.69 |
| QLW-P1DMWA-1 vs. Tro8F10W22 | 0.00257755 | 0.0283585 | 0.0908917 | 1706103 | 455977 | 1.25E+06 | 15849 | 12632.8 | 3216.15 |
| P1-4-10KL vs. Tro8F10W22 | 0.00258822 | 0.0283283 | 0.0913654 | 1706166 | 456483 | 1.25E+06 | 15862 | 12633.7 | 3228.3 |
| MWH-Tro8-2-9 vs. Tro8F10W22 | 0.0022336 | 0.0243685 | 0.0916592 | 1704483 | 454515 | 1.25E+06 | 13644 | 10856.6 | 2787.36 |
| MWH-RechtKol4 vs. MWH-RechtKolB | 8.93E-05 | 0.000974467 | 0.0916854 | 1704036 | 487083 | 1.22E+06 | 583 | 474.28 | 108.72 |
| MWH-RechtKol4 vs. Tro8F10W22 | 0.00658013 | 0.0696538 | 0.094469 | 1701204 | 449438 | 1.25E+06 | 37776 | 29578.9 | 8197.09 |
| MWH-RechtKolB vs. Tro8F10W22 | 0.00654865 | 0.0691745 | 0.0946685 | 1701531 | 449685 | 1.25E+06 | 37560 | 29401.4 | 8158.59 |
| QLW-P1DMWA-1 vs. MWH-RechtKol4 | 0.00588615 | 0.0617279 | 0.0953563 | 1702494 | 450910 | 1.25E+06 | 33797 | 26461.8 | 7335.17 |
| P1-4-10KL vs. MWH-RechtKol4 | 0.00589486 | 0.0617132 | 0.0955202 | 1702494 | 451027 | 1.25E+06 | 33808 | 26462.7 | 7345.28 |
| QLW-P1DMWA-1 vs. MWH-RechtKolB | 0.00585996 | 0.0612638 | 0.0956513 | 1702890 | 451304 | 1.25E+06 | 33597 | 26294.3 | 7302.7 |
| P1-4-10KL vs. MWH-RechtKolB | 0.00586868 | 0.0612441 | 0.0958245 | 1702890 | 451423 | 1.25E+06 | 33606 | 26293.2 | 7312.81 |
| MWH-Recht1 vs. MWH-RechtKol4 | 0.00601424 | 0.0626273 | 0.0960321 | 1702197 | 450690 | 1.25E+06 | 34310 | 26816.3 | 7493.71 |
| MWH-Recht1 vs. MWH-RechtKolB | 0.00598426 | 0.0621824 | 0.0962372 | 1702593 | 450975 | 1.25E+06 | 34108 | 26650.8 | 7457.18 |
| P1-Kol8 vs. MWH-RechtKol4 | 0.00587588 | 0.0608286 | 0.0965973 | 1702464 | 450401 | 1.25E+06 | 33393 | 26067.7 | 7325.28 |
| P1-Kol8 vs. MWH-RechtKolB | 0.00584633 | 0.0603935 | 0.0968039 | 1702755 | 450666 | 1.25E+06 | 33193 | 25904.3 | 7288.75 |
| MWH-RechtKol4 vs. MWH-Tro8-2-9 | 0.00604144 | 0.061833 | 0.0977057 | 1699644 | 451725 | 1.25E+06 | 34061 | 26555.2 | 7505.84 |
| MWH-RechtKolB vs. MWH-Tro8-2-9 | 0.00601881 | 0.0612945 | 0.098195 | 1699956 | 452263 | 1.25E+06 | 33842 | 26365.5 | 7476.5 |
| MWH-RechtKol4 vs. MWH-Tro7-1-4 | 0.00594435 | 0.0598772 | 0.0992758 | 1701645 | 448774 | 1.25E+06 | 33005 | 25589.9 | 7415.11 |

| | | | | | | | | | |
|---|---|---|---|---|---|---|---|---|---|
| MWH-RechtKolB vs. MWH-Tro7-1-4 | 0.00591963 | 0.0594336 | 0.0996007 | 1702062 | 449149 | 1.25E+06 | 32814 | 25429.4 | 7384.65 |
| QLW-P1DMWA-1 vs. MWH-Tro8-2-9 | 0.00125452 | 0.0125479 | 0.0999781 | 1705017 | 462707 | 1.24E+06 | 7303 | 5745.94 | 1557.06 |
| P1-Kol8 vs. MWH-Tro8-2-9 | 0.0011471 | 0.0114233 | 0.100418 | 1705023 | 466364 | 1.24E+06 | 6697 | 5277.34 | 1419.66 |
| P1-4-10KL vs. MWH-Tro8-2-9 | 0.00126523 | 0.0125232 | 0.101031 | 1705080 | 463678 | 1.24E+06 | 7316 | 5746.8 | 1569.2 |
| | | **Median** | **0.0916854** | | | | | | |

Ka = Numbers of substitutions per nonsynonymous site

Ks = Numbers of substitutions per synonymous site

**Supplementary Table S11.** Ka/Ks analysis among the nine strains of *Polynucleobacter asymbioticus* using their whole core genomes.

### Supplementary Table S12.

**Selected genes in *Polynucleobacter asymbioticus* strain QLW-P1DMWA-1[T]**

| Sequence | Method | Ka | Ks | Ka/Ks | P-Value(Fisher) | Length | S-Sites | N-Sites | Substitutions | S-Substitutions | N-Substitutions |
|---|---|---|---|---|---|---|---|---|---|---|---|
| DSBA | MS | 0.0069 | 0.0285 | 0.2402 | 0.0100676 | 624 | 184.867 | 439.133 | 8 | 5.09395 | 2.90605 |
| DSBA | MS | 0.0069 | 0.0285 | 0.2402 | 0.0100676 | 624 | 184.867 | 439.133 | 8 | 5.09395 | 2.90605 |
| DSBA | GMYN | 0.0065 | 0.0321 | 0.2029 | 0.0295253 | 624 | 160.064 | 463.936 | 8 | 5 | 3 |
| DSBA | GMYN | 0.0065 | 0.0321 | 0.2029 | 0.0295253 | 624 | 160.064 | 463.936 | 8 | 5 | 3 |
| RpoE | GMYN | 0.0047 | 0.0289 | 0.1636 | 0.0042709 | 567 | 142.236 | 424.764 | 6 | 4 | 2 |
| RpoE | GMYN | 0.0047 | 0.0289 | 0.1636 | 0.0042709 | 567 | 142.236 | 424.764 | 6 | 4 | 2 |
| Superox | MS | 0.0083 | 0.0606 | 0.1365 | 0.000221587 | 531 | 163.432 | 367.568 | 12 | 9.18071 | 2.81929 |
| RpoE | MS | 0.0045 | 0.0336 | 0.1354 | 0.00299834 | 567 | 132.04 | 434.96 | 6 | 4.14927 | 1.85073 |
| RpoE | MS | 0.0045 | 0.0336 | 0.1354 | 0.00299834 | 567 | 132.04 | 434.96 | 6 | 4.14927 | 1.85073 |
| Fimb | MS | 0.0055 | 0.0540 | 0.1012 | 0.00017699 | 510 | 139.271 | 370.729 | 9 | 7.09043 | 1.90957 |
| Superox | GMYN | 0.0075 | 0.0772 | 0.0974 | 0.00026245 | 531 | 129.114 | 401.886 | 12 | 9 | 3 |
| Fimb | GMYN | 0.0054 | 0.0552 | 0.0972 | 0.00180129 | 510 | 135.493 | 374.507 | 9 | 7 | 2 |
| Fimb | GMYN | 0.0107 | 0.1227 | 0.0876 | 1.54E-06 | 510 | 134.308 | 375.692 | 19 | 15 | 4 |
| Fimb | GMYN | 0.0107 | 0.1371 | 0.0777 | 2.66E-07 | 510 | 129.6 | 380.4 | 20 | 16 | 4 |
| Fimb | GMYN | 0.0107 | 0.1371 | 0.0777 | 2.66E-07 | 510 | 129.6 | 380.4 | 20 | 16 | 4 |
| FtsK | MS | 0.0024 | 0.0345 | 0.0709 | 4.98E-10 | 2310 | 682.337 | 1627.66 | 26 | 22.2388 | 3.76118 |
| FtsK | MS | 0.0024 | 0.0345 | 0.0709 | 4.98E-10 | 2310 | 682.337 | 1627.66 | 26 | 22.2388 | 3.76118 |
| Alkyl | MS | 0.0031 | 0.0443 | 0.0708 | 0 | 465 | 144.282 | 320.718 | 7 | 6.04791 | 0.952093 |
| Alkyl | MS | 0.0031 | 0.0443 | 0.0708 | 0 | 465 | 144.282 | 320.718 | 7 | 6.04791 | 0.952093 |
| FtsK | GMYN | 0.0024 | 0.0362 | 0.0657 | 1.18E-09 | 2310 | 625.81 | 1684.19 | 26 | 22 | 4 |
| FtsK | GMYN | 0.0024 | 0.0362 | 0.0657 | 1.18E-09 | 2310 | 625.81 | 1684.19 | 26 | 22 | 4 |
| Fimb | MS | 0.0084 | 0.1283 | 0.0655 | 1.37E-08 | 510 | 140.079 | 369.921 | 19 | 16.199 | 2.80101 |
| Fimb | MS | 0.0086 | 0.1329 | 0.0650 | 9.77E-09 | 510 | 147.434 | 362.566 | 20 | 17.2435 | 2.75652 |
| Fimb | MS | 0.0086 | 0.1329 | 0.0650 | 9.77E-09 | 510 | 147.434 | 362.566 | 20 | 17.2435 | 2.75652 |
| Methyl | MS | 0.0754 | 1.1880 | 0.0635 | 1.18E-128 | 1470 | 350.71 | 1119.29 | 276 | 229.493 | 46.5069 |
| Alkyl | GMYN | 0.0029 | 0.0532 | 0.0549 | 0.00146005 | 465 | 120.669 | 344.331 | 7 | 6 | 1 |
| Alkyl | GMYN | 0.0029 | 0.0532 | 0.0549 | 0.00146005 | 465 | 120.669 | 344.331 | 7 | 6 | 1 |
| Fimb | MS | 0.0359 | 0.6772 | 0.0530 | 3.91E-30 | 510 | 134.073 | 375.927 | 66 | 57.4616 | 8.53837 |

| | | | | | | | | | | | |
|---|---|---|---|---|---|---|---|---|---|---|---|
| Fimb | GMYN | 0.0355 | 0.7096 | 0.0500 | 8.03E-24 | 510 | 131.263 | 378.737 | 66 | 52.9318 | 13.0682 |
| Alkyl | MS | 0.0032 | 0.0730 | 0.0439 | 0 | 465 | 149.252 | 315.748 | 11 | 10.0655 | 0.93448 |
| Alkyl | GMYN | 0.0030 | 0.0841 | 0.0356 | 1.76E-05 | 465 | 129.538 | 335.462 | 11 | 10 | 1 |
| Methyl | GMYN | 0.0697 | 2.3825 | 0.0292 | 7.38E-89 | 1470 | 348.153 | 1121.85 | 276 | 201.683 | 74.3171 |
| RuvB | MS | 0.0013 | 0.0715 | 0.0187 | 0 | 1068 | 318.884 | 749.116 | 22 | 21.0732 | 0.926773 |
| RuvB | MS | 0.0013 | 0.0715 | 0.0187 | 0 | 1068 | 318.884 | 749.116 | 22 | 21.0732 | 0.926773 |
| RuvB | GMYN | 0.0013 | 0.0836 | 0.0150 | 2.76E-12 | 1068 | 270.522 | 797.478 | 22 | 21 | 1 |
| RuvB | GMYN | 0.0013 | 0.0836 | 0.0150 | 2.76E-12 | 1068 | 270.522 | 797.478 | 22 | 21 | 1 |
| MscS | MS | 0.0000 | 0.0437 | 0.0000 | 0 | 819 | 216.856 | 602.144 | 9 | 8.99998 | 2.50E-05 |
| MscS | MS | 0.0000 | 0.0437 | 0.0000 | 0 | 819 | 216.856 | 602.144 | 9 | 8.99998 | 2.50E-05 |

### Housekeeping genes in *Polynucleobacter asymbioticus* strain QLW-P1DMWA-1T

| Sequence | Method | Ka | Ks | Ka/Ks | P-Value(Fisher) | Length | S-Sites | N-Sites | Substitutions | S-Substitutions | N-Substitutions |
|---|---|---|---|---|---|---|---|---|---|---|---|
| atpA | MS | 0.0010 | 0.0058 | 0.1735 | 0 | 1539 | 533.218 | 1005.78 | 4 | 3.01356 | 0.986439 |
| cit | GMYN | 0.0021 | 0.0160 | 0.1290 | 0.0101498 | 1311 | 312.415 | 998.585 | 7 | 4.9408 | 2.0592 |
| cit | GMYN | 0.0021 | 0.0160 | 0.1290 | 0.0101498 | 1311 | 312.415 | 998.585 | 7 | 4.9408 | 2.0592 |
| CTP | MS | 0.0018 | 0.0190 | 0.0931 | 7.22E-05 | 1659 | 525.286 | 1133.71 | 11 | 9.15969 | 1.84031 |
| CTP | MS | 0.0018 | 0.0190 | 0.0931 | 7.22E-05 | 1659 | 525.286 | 1133.71 | 11 | 9.15969 | 1.84031 |
| CTP | GMYN | 0.0016 | 0.0221 | 0.0730 | 0.000125077 | 1659 | 417.189 | 1241.81 | 11 | 9 | 2 |
| CTP | GMYN | 0.0016 | 0.0221 | 0.0730 | 0.000125077 | 1659 | 417.189 | 1241.81 | 11 | 9 | 2 |
| cit | MS | 0.0016 | 0.0234 | 0.0684 | 0.000382964 | 1311 | 267.118 | 1043.88 | 7 | 5.52364 | 1.47636 |
| cit | MS | 0.0016 | 0.0234 | 0.0684 | 0.000382964 | 1311 | 267.118 | 1043.88 | 7 | 5.52364 | 1.47636 |
| adk | GMYN | 0.0090 | 0.1475 | 0.0609 | 5.40E-11 | 663 | 215.007 | 447.993 | 32 | 28 | 4 |
| adk | GMYN | 0.0090 | 0.1475 | 0.0609 | 5.40E-11 | 663 | 215.007 | 447.993 | 32 | 28 | 4 |
| adk | MS | 0.0083 | 0.1850 | 0.0449 | 2.00E-14 | 663 | 181.819 | 481.181 | 32 | 28.5993 | 3.40068 |
| adk | MS | 0.0083 | 0.1850 | 0.0449 | 2.00E-14 | 663 | 181.819 | 481.181 | 32 | 28.5993 | 3.40068 |
| adk | GMYN | 0.0061 | 0.1459 | 0.0419 | 3.77E-11 | 663 | 170.116 | 492.884 | 25 | 22 | 3 |
| acsA | GMYN | 0.0028 | 0.0699 | 0.0399 | 1.30E-16 | 1971 | 530.919 | 1440.08 | 39 | 35 | 4 |
| acsA | GMYN | 0.0028 | 0.0699 | 0.0399 | 1.30E-16 | 1971 | 530.919 | 1440.08 | 39 | 35 | 4 |
| adk | MS | 0.0079 | 0.1992 | 0.0397 | 6.71E-15 | 663 | 159.472 | 503.528 | 30 | 26.6596 | 3.34038 |
| adk | MS | 0.0079 | 0.1992 | 0.0397 | 6.71E-15 | 663 | 159.472 | 503.528 | 30 | 26.6596 | 3.34038 |
| acsA | MS | 0.0028 | 0.0711 | 0.0392 | 1.51E-17 | 1971 | 547.583 | 1423.42 | 39 | 35.392 | 3.60804 |

| | | | | | | | | | | | |
|---|---|---|---|---|---|---|---|---|---|---|---|
| acsA | MS | 0.0028 | 0.0711 | 0.0392 | 1.51E-17 | 1971 | 547.583 | 1423.42 | 39 | 35.392 | 3.60804 |
| adk | GMYN | 0.0078 | 0.2078 | 0.0375 | 1.59E-14 | 663 | 146.132 | 516.868 | 30 | 26 | 4 |
| adk | GMYN | 0.0078 | 0.2078 | 0.0375 | 1.59E-14 | 663 | 146.132 | 516.868 | 30 | 26 | 4 |
| adk | MS | 0.0059 | 0.1633 | 0.0364 | 3.45E-13 | 663 | 160.211 | 502.789 | 25 | 22.4396 | 2.56039 |
| acsA | GMYN | 0.0007 | 0.0222 | 0.0307 | 2.52E-06 | 1971 | 503.811 | 1467.19 | 12 | 11 | 1 |
| acsA | MS | 0.0007 | 0.0247 | 0.0269 | 0 | 1971 | 486.858 | 1484.14 | 12 | 11.0915 | 0.908467 |
| iso | MS | 0.0006 | 0.0236 | 0.0252 | 0 | 2232 | 561.142 | 1670.86 | 14 | 13.0219 | 0.978057 |
| iso | GMYN | 0.0006 | 0.0239 | 0.0250 | 1.35E-07 | 2232 | 555.944 | 1676.06 | 14 | 13 | 1 |
| iso | GMYN | 0.0019 | 0.0891 | 0.0216 | 4.46E-26 | 2232 | 670.713 | 1561.29 | 58 | 54.9983 | 3.0017 |
| iso | GMYN | 0.0019 | 0.0891 | 0.0216 | 4.46E-26 | 2232 | 670.713 | 1561.29 | 58 | 54.9983 | 3.0017 |
| iso | MS | 0.0019 | 0.0942 | 0.0201 | 1.10E-28 | 2232 | 646.227 | 1585.77 | 58 | 55.2741 | 2.72588 |
| iso | MS | 0.0019 | 0.0942 | 0.0201 | 1.10E-28 | 2232 | 646.227 | 1585.77 | 58 | 55.2741 | 2.72588 |
| iso | GMYN | 0.0006 | 0.0392 | 0.0152 | 0 | 2232 | 553.663 | 1678.34 | 22 | 21 | 1 |
| gyr | MS | 0.0000 | 0.0079 | 0.0000 | 0 | 1434 | 378.849 | 1055.15 | 3 | 2.99999 | 8.36E-06 |
| gyr | MS | 0.0000 | 0.0079 | 0.0000 | 0 | 1434 | 378.849 | 1055.15 | 3 | 2.99999 | 8.36E-06 |
| atpA | MS | 0.0000 | 0.0096 | 0.0000 | 0 | 1539 | 533.542 | 1005.46 | 5 | 4.99999 | 9.42E-06 |
| atpA | MS | 0.0000 | 0.0096 | 0.0000 | 0 | 1539 | 533.542 | 1005.46 | 5 | 4.99999 | 9.42E-06 |
| gyr | MS | 0.0000 | 0.0144 | 0.0000 | 0 | 1434 | 430.428 | 1003.57 | 6 | 5.99999 | 1.40E-05 |
| glnA | MS | 0.0000 | 0.0247 | 0.0000 | 0 | 1413 | 393.521 | 1019.48 | 9 | 8.99998 | 2.33E-05 |
| glnA | MS | 0.0000 | 0.0247 | 0.0000 | 0 | 1413 | 393.521 | 1019.48 | 9 | 8.99998 | 2.33E-05 |
| succ | MS | 0.0000 | 0.0366 | 0.0000 | 0 | 1776 | 430.6 | 1345.4 | 15 | 15 | 4.69E-05 |
| succ | MS | 0.0000 | 0.0366 | 0.0000 | 0 | 1776 | 430.6 | 1345.4 | 15 | 15 | 4.69E-05 |
| atpA | GMYN | 0.0000 | 0.0124 | 0.0000 | 0 | 1539 | 407.975 | 1131.02 | 5 | 5 | 0 |
| atpA | GMYN | 0.0000 | 0.0124 | 0.0000 | 0 | 1539 | 407.975 | 1131.02 | 5 | 5 | 0 |
| glnA | GMYN | 0.0000 | 0.0264 | 0.0000 | 0 | 1413 | 348.816 | 1064.18 | 9 | 9 | 0 |
| glnA | GMYN | 0.0000 | 0.0264 | 0.0000 | 0 | 1413 | 348.816 | 1064.18 | 9 | 9 | 0 |
| succ | GMYN | 0.0000 | 0.0369 | 0.0000 | 0 | 1776 | 421.093 | 1354.91 | 15 | 15 | 0 |
| succ | GMYN | 0.0000 | 0.0369 | 0.0000 | 0 | 1776 | 421.093 | 1354.91 | 15 | 15 | 0 |

### Selected genes in PnecC subcluster

| Sequence | Method | Ka | Ks | Ka/Ks | P-Value(Fisher) | Length | S-Sites | N-Sites | Substitutions | S-Substitutions | N-Substitutions |
|---|---|---|---|---|---|---|---|---|---|---|---|
| Cold | MS | 0.0587 | 0.3859 | 0.1520 | 0.000282898 | 126 | 33.8772 | 92.1228 | 14 | 9.90516 | 4.09484 |

| | | | | | | | | | | | |
|---|---|---|---|---|---|---|---|---|---|---|---|
| Cold | GMYN | 0.0555 | 0.3986 | 0.1392 | 0.000979928 | 126 | 31.0745 | 94.9255 | 14 | 9 | 5 |
| RpoE | GMYN | 0.150827 | 1.68973 | 0.0892605 | 1.74E-24 | 426 | 99.2193 | 326.781 | 111 | 66.5945 | 44.4055 |
| Hyp | GMYN | 0.1130 | 1.3727 | 0.0823 | 6.17E-13 | 258 | 67.9994 | 190.001 | 56 | 36.1982 | 19.8018 |
| DSBA | MS | 0.2520 | 3.4307 | 0.0735 | NA | 591 | 139.069 | 451.931 | 224 | 180.83 | 43.1698 |
| DSBA | GMYN | 0.2600 | 3.6288 | 0.0717 | 2.50E-59 | 591 | 126.273 | 464.727 | 224 | 122.845 | 101.155 |
| Hyp | GMYN | 0.0627 | 0.9219 | 0.0680 | 5.24E-13 | 243 | 70.2652 | 172.735 | 43 | 32.7181 | 10.2819 |
| DSBA | GMYN | 0.2308 | 3.5685 | 0.0647 | 4.05E-59 | 543 | 116.507 | 426.493 | 197 | 113.199 | 83.8006 |
| DSBA | MS | 0.2287 | 3.5739 | 0.0640 | NA | 543 | 125.194 | 417.806 | 197 | 162.327 | 34.6731 |
| DSBA | GMYN | 0.2311 | 3.6198 | 0.0638 | 1.92E-67 | 600 | 124.76 | 475.24 | 217 | 123.477 | 93.5227 |
| RpoE | GMYN | 0.0408 | 0.6439 | 0.0634 | 2.68E-22 | 489 | 157.596 | 331.404 | 74 | 60.869 | 13.131 |
| DSBA | MS | 0.2284 | 3.7082 | 0.0616 | NA | 600 | 133.047 | 466.953 | 217 | 178.435 | 38.5649 |
| Hyp | MS | 0.1081 | 1.7632 | 0.0613 | 3.83E-30 | 258 | 63.4016 | 194.598 | 56 | 47.1288 | 8.87121 |
| Hfq | MS | 0.0330 | 0.5506 | 0.0598 | 2.25E-12 | 228 | 67.352 | 160.648 | 28 | 24.5025 | 3.49753 |
| DSBA | GMYN | 0.2140 | 3.6227 | 0.0591 | 3.91E-49 | 594 | 125.244 | 468.756 | 197 | 110.641 | 86.3591 |
| DSBA | GMYN | 0.2078 | 3.5746 | 0.0581 | 2.10E-54 | 609 | 117.458 | 491.542 | 197 | 108.721 | 88.2793 |
| DSBA | MS | 0.2138 | 3.7619 | 0.0568 | NA | 594 | 131.622 | 462.378 | 197 | 164.216 | 32.7843 |
| Piri | GMYN | 0.1515 | 2.7019 | 0.0561 | 4.72E-47 | 666 | 167.49 | 498.51 | 190 | 122.194 | 67.8064 |
| Piri | GMYN | 0.2025 | 3.7300 | 0.0543 | 1.38E-49 | 663 | 144.516 | 518.484 | 212 | 120.756 | 91.2439 |
| DSBA | MS | 0.2109 | 3.8950 | 0.0541 | NA | 609 | 130.442 | 478.558 | 197 | 164.351 | 32.6492 |
| Piri | MS | 0.2023 | 3.8314 | 0.0528 | NA | 663 | 145.735 | 517.265 | 212 | 178.54 | 33.4597 |
| Hfq | MS | 0.0309 | 0.5873 | 0.0527 | 2.46E-08 | 144 | 42.0409 | 101.959 | 18 | 15.9613 | 2.0387 |
| Hyp | MS | 0.0620 | 1.2128 | 0.0511 | 5.61E-22 | 243 | 68.3067 | 174.693 | 43 | 38.0284 | 4.97162 |
| Piri | MS | 0.1090 | 2.1400 | 0.0509 | 4.45E-100 | 669 | 150.938 | 518.062 | 160 | 136.19 | 23.8101 |
| MscS | MS | 0.1795 | 3.6195 | 0.0496 | NA | 774 | 184.613 | 589.387 | 249 | 214.968 | 34.0321 |
| MscS | MS | 0.1391 | 2.8398 | 0.0490 | 5.40E-147 | 789 | 183.979 | 605.021 | 212 | 182.586 | 29.4139 |
| RpoE | MS | 0.0421 | 0.8878 | 0.0475 | 9.16E-42 | 522 | 130.159 | 391.841 | 83 | 72.6232 | 10.3768 |
| Piri | MS | 0.1708 | 3.6202 | 0.0472 | NA | 684 | 164.432 | 519.568 | 208 | 181.022 | 26.9781 |
| Superox | GMYN | 0.0771 | 1.6481 | 0.0468 | 8.99E-32 | 471 | 123.061 | 347.939 | 99 | 73.6828 | 25.3172 |
| RpoE | GMYN | 0.0343 | 0.7338 | 0.0467 | 6.69E-29 | 537 | 138.458 | 398.542 | 74 | 60.6677 | 13.3323 |
| DSBA | GMYN | 0.1667 | 3.6609 | 0.0455 | 1.10E-55 | 618 | 131.796 | 486.204 | 187 | 114.925 | 72.0753 |
| MscS | GMYN | 0.1792 | 3.9383 | 0.0455 | 4.08E-65 | 774 | 190.775 | 583.225 | 249 | 157.243 | 91.7568 |
| GroL | MS | 0.0315 | 0.6930 | 0.0454 | 1.64E-95 | 1614 | 436.828 | 1177.17 | 210 | 187.113 | 22.8874 |
| Piri | MS | 0.1626 | 3.5973 | 0.0452 | NA | 594 | 144.815 | 449.185 | 176 | 154.353 | 21.647 |
| Rubre | GMYN | 0.0745 | 1.6575 | 0.0449 | 1.76E-33 | 405 | 106.147 | 298.853 | 92 | 70.8764 | 21.1236 |

| | | | | | | | | | | | |
|---|---|---|---|---|---|---|---|---|---|---|---|
| Piri | MS | 0.1562 | 3.5478 | 0.0440 | NA | 615 | 153.004 | 461.996 | 188 | 165.936 | 22.0642 |
| Piri | GMYN | 0.1614 | 3.6715 | 0.0440 | 4.43E-49 | 594 | 133.658 | 460.342 | 176 | 109.705 | 66.2949 |
| Hyp | GMYN | 0.0874 | 1.9998 | 0.0437 | 1.49E-16 | 237 | 74.8291 | 162.171 | 57 | 43.7518 | 13.2482 |
| Fatty | MS | 0.0686111 | 1.57088 | 0.0436769 | 1.40E-131 | 1050 | 263.549 | 786.451 | 231 | 204.364 | 26.6359 |
| DSBA | MS | 0.1276 | 2.9315 | 0.0435 | NA | 603 | 138.561 | 464.439 | 163 | 142.251 | 20.7487 |
| DSBA | MS | 0.1276 | 2.9315 | 0.0435 | NA | 603 | 138.561 | 464.439 | 163 | 142.251 | 20.7487 |
| Hfq | MS | 0.0505 | 1.1646 | 0.0434 | 6.17E-15 | 195 | 67.7019 | 127.298 | 33 | 30.511 | 2.48904 |
| Superox | MS | 0.0731 | 1.6862 | 0.0433 | 9.78E-61 | 471 | 104.752 | 366.248 | 99 | 85.9714 | 13.0286 |
| MscS | MS | 0.1593 | 3.6796 | 0.0433 | NA | 783 | 186.984 | 596.016 | 252 | 221.435 | 30.5654 |
| Superox | GMYN | 0.1545 | 3.5797 | 0.0432 | 2.51E-40 | 501 | 118.272 | 382.728 | 147 | 93.8519 | 53.1481 |
| Hfq | MS | 0.0422 | 0.9785 | 0.0431 | 6.20E-20 | 234 | 60.883 | 173.117 | 38 | 33.8495 | 4.15045 |
| DSBA | MS | 0.1640 | 3.8153 | 0.0430 | NA | 618 | 141.504 | 476.496 | 187 | 163.361 | 23.6387 |
| Piri | MS | 0.1350 | 3.1715 | 0.0426 | NA | 675 | 144.253 | 530.747 | 176 | 152.168 | 23.8322 |
| Piri | MS | 0.1350 | 3.1715 | 0.0426 | NA | 675 | 144.253 | 530.747 | 176 | 152.168 | 23.8322 |
| Piri | GMYN | 0.1554 | 3.6912 | 0.0421 | 7.69E-62 | 615 | 137.219 | 477.781 | 188 | 121.503 | 66.4971 |
| Piri | MS | 0.1525 | 3.7022 | 0.0412 | NA | 666 | 159.009 | 506.991 | 190 | 167.943 | 22.057 |
| Piri | GMYN | 0.1563 | 3.7993 | 0.0411 | 3.33E-63 | 684 | 158.501 | 525.499 | 208 | 134.636 | 73.3645 |
| Piri | MS | 0.1356 | 3.3205 | 0.0408 | NA | 633 | 135.082 | 497.918 | 171 | 148.624 | 22.3756 |
| MscS | GMYN | 0.1594 | 3.9277 | 0.0406 | 3.07E-82 | 783 | 188.102 | 594.898 | 252 | 167.729 | 84.271 |
| RpoE | MS | 0.0373 | 0.9192 | 0.0406 | 6.41E-39 | 489 | 120.406 | 368.594 | 74 | 65.8281 | 8.1719 |
| Hfq | GMYN | 0.0292 | 0.7239 | 0.0403 | 7.47E-08 | 144 | 37.722 | 106.278 | 18 | 15 | 3 |
| RpoE | GMYN | 0.0305 | 0.7767 | 0.0393 | 3.39E-31 | 546 | 136.588 | 409.412 | 74 | 61.7805 | 12.2195 |
| RpoE | GMYN | 0.0305 | 0.7767 | 0.0393 | 3.39E-31 | 546 | 136.588 | 409.412 | 74 | 61.7805 | 12.2195 |
| RpoE | MS | 0.142481 | 3.63299 | 0.0392187 | NA | 426 | 95.0634 | 330.937 | 111 | 97.6658 | 13.3342 |
| RpoE | MS | 0.0346 | 0.8836 | 0.0391 | 3.37E-41 | 537 | 122.034 | 414.966 | 74 | 65.3127 | 8.68725 |
| DSBA | GMYN | 0.1401 | 3.6040 | 0.0389 | 3.66E-50 | 522 | 122.167 | 399.833 | 153 | 102.162 | 50.8375 |
| Hfq | GMYN | 0.0442 | 1.1378 | 0.0388 | 1.42E-13 | 195 | 51.3775 | 143.623 | 33 | 26.9232 | 6.07677 |
| Superox | MS | 0.1478 | 3.9317 | 0.0376 | NA | 501 | 112.829 | 388.171 | 147 | 130.161 | 16.8386 |
| Piri | GMYN | 0.1383 | 3.7079 | 0.0373 | 1.97E-62 | 654 | 140.313 | 513.687 | 185 | 120.648 | 64.3521 |
| Hfq | GMYN | 0.0478 | 1.2818 | 0.0373 | 5.32E-18 | 210 | 55.2251 | 154.775 | 41 | 33.9205 | 7.0795 |
| Hyp | GMYN | 0.0398 | 1.0712 | 0.0371 | 4.83E-18 | 219 | 63.1353 | 155.865 | 42 | 36 | 6 |
| DSBA | GMYN | 0.1354 | 3.6545 | 0.0370 | 7.52E-46 | 564 | 130.663 | 433.337 | 155 | 101.764 | 53.2363 |
| MscS | MS | 0.1362 | 3.6868 | 0.0369 | NA | 768 | 186.848 | 581.152 | 228 | 204.509 | 23.4909 |
| Piri | GMYN | 0.1362 | 3.6944 | 0.0369 | 1.26E-51 | 633 | 137.803 | 495.197 | 171 | 109.728 | 61.2724 |

| | | | | | | | | | | | |
|---|---|---|---|---|---|---|---|---|---|---|---|
| GroL | GMYN | 0.0295 | 0.8056 | 0.0366 | 4.83E-85 | 1614 | 398.948 | 1215.05 | 210 | 174.942 | 35.0579 |
| Piri | GMYN | 0.1361 | 3.7244 | 0.0365 | 2.62E-65 | 690 | 143.423 | 546.577 | 191 | 123.533 | 67.4672 |
| RpoH | MS | 0.0654 | 1.7953 | 0.0364 | 2.41E-113 | 906 | 254.257 | 651.743 | 204 | 186.581 | 17.4195 |
| DSBA | MS | 0.1384 | 3.8109 | 0.0363 | NA | 522 | 122.466 | 399.534 | 153 | 136.792 | 16.208 |
| Fatty | MS | 0.0708 | 1.9511 | 0.0363 | 3.10E-164 | 1182 | 312.083 | 869.917 | 276 | 250.66 | 25.3403 |
| RpoH | MS | 0.0438 | 1.2207 | 0.0359 | 1.02E-82 | 894 | 254.367 | 639.633 | 162 | 148.596 | 13.4035 |
| Alkyl | GMYN | 0.0556 | 1.5720 | 0.0353 | 1.44E-30 | 387 | 105.061 | 281.939 | 79 | 63.94 | 15.06 |
| Fatty | MS | 0.0600 | 1.7005 | 0.0353 | 1.43E-146 | 1125 | 255.687 | 869.313 | 230 | 205.347 | 24.6531 |
| Alkyl | GMYN | 0.0616 | 1.7443 | 0.0353 | 1.49E-37 | 432 | 104.463 | 327.537 | 90 | 70.6916 | 19.3084 |
| Fatty | MS | 0.0441 | 1.2511 | 0.0353 | 7.41E-116 | 1143 | 298.721 | 844.279 | 212 | 192.785 | 19.2146 |
| DSBA | MS | 0.1303 | 3.6993 | 0.0352 | NA | 600 | 146.215 | 453.785 | 163 | 146.94 | 16.0595 |
| DSBA | MS | 0.1338 | 3.8000 | 0.0352 | NA | 564 | 133.261 | 430.739 | 155 | 139.167 | 15.8333 |
| DSBA | GMYN | 0.1316 | 3.7388 | 0.0352 | 2.41E-46 | 600 | 145.769 | 454.231 | 163 | 108.579 | 54.4209 |
| Fatty | GMYN | 0.0439 | 1.2500 | 0.0352 | 4.39E-91 | 1143 | 288.675 | 854.325 | 212 | 175.616 | 36.3844 |
| MscS | GMYN | 0.1338 | 3.8727 | 0.0346 | 7.12E-64 | 789 | 174.787 | 614.213 | 212 | 137.427 | 74.5728 |
| Hfq | MS | 0.0500 | 1.4481 | 0.0345 | 2.03E-22 | 210 | 60.6699 | 149.33 | 41 | 37.7899 | 3.21014 |
| Alkyl | MS | 0.0501 | 1.4584 | 0.0343 | 1.09E-49 | 435 | 105.063 | 329.937 | 81 | 73.1168 | 7.88318 |
| DSBA | GMYN | 0.1248 | 3.6412 | 0.0343 | 1.15E-57 | 603 | 128.368 | 474.632 | 163 | 108.823 | 54.1775 |
| DSBA | GMYN | 0.1248 | 3.6412 | 0.0343 | 1.15E-57 | 603 | 128.368 | 474.632 | 163 | 108.823 | 54.1775 |
| MscS | GMYN | 0.1339 | 3.9083 | 0.0343 | 1.80E-79 | 768 | 183.276 | 584.724 | 228 | 157.031 | 70.9689 |
| Fatty | MS | 0.0671 | 1.9638 | 0.0342 | 9.31E-170 | 1161 | 271.284 | 889.716 | 257 | 231.106 | 25.8945 |
| RpoH | MS | 0.0368315 | 1.0868 | 0.03389 | 3.08E-69 | 780 | 218.821 | 561.179 | 133 | 122.365 | 10.6351 |
| RpoH | MS | 0.0967 | 2.8548 | 0.0339 | 1.83E-142 | 885 | 232.111 | 652.889 | 222 | 202.684 | 19.3164 |
| Fatty | MS | 0.0546 | 1.6296 | 0.0335 | 4.31E-147 | 1161 | 280.18 | 880.82 | 238 | 215.329 | 22.6708 |
| Carbo | GMYN | 0.0530 | 1.5878 | 0.0334 | 2.63E-52 | 579 | 148.626 | 430.374 | 120 | 98.0499 | 21.9501 |
| Hfq | GMYN | 0.0414 | 1.2440 | 0.0333 | 8.08E-16 | 234 | 59.4434 | 174.557 | 38 | 30.9876 | 7.01241 |
| Piri | MS | 0.1342 | 4.1084 | 0.0327 | NA | 654 | 142.481 | 511.519 | 185 | 165.585 | 19.4154 |
| Rubre | GMYN | 0.0357 | 1.0941 | 0.0326 | 2.11E-29 | 381 | 122.203 | 258.797 | 72 | 63 | 9 |
| RpoH | GMYN | 0.0413 | 1.2961 | 0.0318 | 6.98E-73 | 894 | 218.78 | 675.22 | 162 | 134.941 | 27.0586 |
| RpoH | MS | 0.0392 | 1.2508 | 0.0313 | 1.71E-91 | 912 | 255.122 | 656.878 | 169 | 156.386 | 12.6136 |
| RpoE | GMYN | 0.0453 | 1.4523 | 0.0312 | 2.25E-42 | 534 | 140.738 | 393.262 | 101 | 83.734 | 17.266 |
| RpoH | MS | 0.0576 | 1.8488 | 0.0311 | 5.55E-108 | 888 | 261.831 | 626.169 | 196 | 182.414 | 13.586 |
| GroL | MS | 0.0255 | 0.8289 | 0.0308 | 9.77E-113 | 1608 | 433.234 | 1174.77 | 219 | 202.11 | 16.8905 |
| Rubre | GMYN | 0.0676 | 2.1995 | 0.0307 | 6.65E-43 | 438 | 118.953 | 319.047 | 105 | 84.4634 | 20.5366 |

| | | | | | | | | | | | |
|---|---|---|---|---|---|---|---|---|---|---|---|
| GroL | GMYN | 0.0239 | 0.7832 | 0.0305 | 2.93E-85 | 1551 | 401.001 | 1150 | 199 | 171.99 | 27.0099 |
| Hfq | GMYN | 0.0307 | 1.0094 | 0.0304 | 7.28E-11 | 228 | 61.2365 | 166.763 | 28 | 23 | 5 |
| Alkyl | GMYN | 0.0501 | 1.6571 | 0.0302 | 1.16E-37 | 435 | 96.8239 | 338.176 | 81 | 64.647 | 16.353 |
| Fatty | MS | 0.0511 | 1.6921 | 0.0302 | 2.84E-146 | 1101 | 260.493 | 840.507 | 227 | 206.861 | 20.1391 |
| Piri | MS | 0.1323 | 4.4088 | 0.0300 | NA | 690 | 139.996 | 550.004 | 191 | 170.853 | 20.1475 |
| Piri | GMYN | 0.1105 | 3.7016 | 0.0298 | 1.67E-52 | 669 | 139.138 | 529.862 | 160 | 105.835 | 54.1649 |
| GroL | MS | 0.0241 | 0.8124 | 0.0296 | 2.24E-102 | 1551 | 417.608 | 1133.39 | 199 | 184.183 | 14.817 |
| Alkyl | GMYN | 0.0451 | 1.5451 | 0.0292 | 5.34E-39 | 432 | 106.656 | 325.344 | 84 | 69.7976 | 14.2024 |
| Rubre | GMYN | 0.0465 | 1.5976 | 0.0291 | 1.76E-41 | 450 | 116.512 | 333.488 | 92 | 77 | 15 |
| RpoH | GMYN | 0.0360859 | 1.24944 | 0.0288816 | 5.77E-56 | 780 | 209.234 | 570.766 | 133 | 112.938 | 20.062 |
| Fatty | GMYN | 0.0500 | 1.7359 | 0.0288 | 1.15E-106 | 1155 | 298.506 | 856.494 | 243 | 201.757 | 41.2426 |
| Fatty | MS | 0.0614 | 2.1489 | 0.0286 | 6.27E-186 | 1137 | 254.923 | 882.077 | 256 | 232.957 | 23.0428 |
| Alkyl | MS | 0.0578 | 2.0329 | 0.0285 | 1.81E-60 | 432 | 103.051 | 328.949 | 90 | 82.5059 | 7.49409 |
| FtsK | GMYN | 0.0365 | 1.2947 | 0.0282 | 9.82E-195 | 2244 | 602.918 | 1641.08 | 430 | 371.722 | 58.2781 |
| Carbo | GMYN | 0.0695 | 2.4956 | 0.0279 | 1.39E-61 | 630 | 166.479 | 463.521 | 151 | 120.379 | 30.621 |
| Hyp | MS | 0.0429 | 1.5402 | 0.0279 | 2.74E-22 | 219 | 69.9217 | 149.078 | 42 | 39.6458 | 2.35418 |
| RpoH | MS | 0.0579 | 2.0835 | 0.0278 | 2.69E-122 | 897 | 255.006 | 641.994 | 206 | 192.529 | 13.4715 |
| FtsK | MS | 0.0368 | 1.3299 | 0.0276 | 1.73E-229 | 2244 | 653.403 | 1590.6 | 430 | 402.885 | 27.1152 |
| Fatty | MS | 0.0488 | 1.7769 | 0.0275 | 1.86E-158 | 1155 | 283.951 | 871.049 | 243 | 224.103 | 18.8968 |
| Alkyl | GMYN | 0.0349 | 1.2734 | 0.0274 | 4.78E-36 | 405 | 105.767 | 299.233 | 75 | 64.8017 | 10.1983 |
| RpoH | MS | 0.0322 | 1.1725 | 0.0274 | 1.47E-82 | 903 | 262.057 | 640.943 | 155 | 145.254 | 9.74615 |
| Fatty | MS | 0.0597 | 2.1779 | 0.0274 | 6.46E-171 | 1092 | 249.207 | 842.793 | 240 | 219.643 | 20.3566 |
| Fatty | GMYN | 0.0510 | 1.8746 | 0.0272 | 9.80E-107 | 1161 | 282.956 | 878.044 | 238 | 194.787 | 43.2133 |
| Carbo | MS | 0.0711 | 2.6137 | 0.0272 | 2.92E-105 | 639 | 145.466 | 493.534 | 144 | 131.831 | 12.1692 |
| FtsK | GMYN | 0.0330 | 1.2264 | 0.0269 | 1.94E-196 | 2271 | 620.571 | 1650.43 | 428 | 374.709 | 53.2915 |
| Hyp | GMYN | 0.0384 | 1.4289 | 0.0269 | 1.25E-14 | 204 | 42.1986 | 161.801 | 30 | 24 | 6 |
| Hfq | GMYN | 0.0331 | 1.2354 | 0.0268 | 9.32E-19 | 210 | 52.7496 | 157.25 | 37 | 31.943 | 5.057 |
| Fatty | MS | 0.0642 | 2.4045 | 0.0267 | 2.80E-189 | 1167 | 268.666 | 898.334 | 262 | 240.539 | 21.4612 |
| Alkyl | MS | 0.0440 | 1.6646 | 0.0264 | 1.07E-55 | 432 | 103.259 | 328.741 | 84 | 77.4847 | 6.51525 |
| RpoH | MS | 0.0333 | 1.2678 | 0.0263 | 3.24E-79 | 822 | 231.935 | 590.065 | 145 | 135.908 | 9.09161 |
| Fatty | MS | 0.0647 | 2.4607 | 0.0263 | 1.27E-190 | 1158 | 264.889 | 893.111 | 261 | 239.749 | 21.2512 |
| RpoE | MS | 0.0425 | 1.6226 | 0.0262 | 1.53E-66 | 534 | 123.714 | 410.286 | 101 | 92.9237 | 8.07626 |
| Fimb | GMYN | 0.0955 | 3.6445 | 0.0262 | 2.71E-60 | 489 | 128.943 | 360.057 | 143 | 110.897 | 32.1029 |
| RpoE | MS | 0.0243 | 0.9293 | 0.0262 | 2.06E-46 | 543 | 126.226 | 416.774 | 77 | 70.8697 | 6.13029 |

| | | | | | | | | | | | |
|---|---|---|---|---|---|---|---|---|---|---|---|
| RpoE | MS | 0.0224 | 0.8582 | 0.0261 | 3.76E-38 | 465 | 103.751 | 361.249 | 62 | 56.8306 | 5.16941 |
| Hyp | MS | 0.0839 | 3.2142 | 0.0261 | 2.06E-35 | 237 | 69.3616 | 167.638 | 57 | 53.6181 | 3.38187 |
| Alkyl | GMYN | 0.0518 | 1.9869 | 0.0261 | 8.43E-33 | 444 | 119.039 | 324.961 | 85 | 68.7605 | 16.2395 |
| Hfq | MS | 0.0346 | 1.3306 | 0.0260 | 1.16E-21 | 210 | 58.9698 | 151.03 | 37 | 34.6928 | 2.30717 |
| RpoE | GMYN | 0.0403 | 1.5532 | 0.0260 | 3.76E-34 | 522 | 129.844 | 392.156 | 83 | 67.632 | 15.368 |
| GroL | MS | 0.0173 | 0.6752 | 0.0256 | 5.00E-95 | 1590 | 479.487 | 1110.51 | 198 | 186.937 | 11.0633 |
| Fimb | MS | 0.0922 | 3.6166 | 0.0255 | NA | 489 | 125.954 | 363.046 | 143 | 133.21 | 9.78959 |
| Rubre | MS | 0.0796 | 3.1231 | 0.0255 | 1.44E-52 | 405 | 122.483 | 282.517 | 92 | 86.8934 | 5.1066 |
| Fatty | GMYN | 0.0674697 | 2.72085 | 0.0247973 | 1.48E-97 | 1050 | 245.528 | 804.472 | 231 | 179.253 | 51.7474 |
| FtsK | MS | 0.0298 | 1.2058 | 0.0247 | 5.23E-226 | 2256 | 594.393 | 1661.61 | 392 | 366.665 | 25.3354 |
| RpoH | GMYN | 0.0440 | 1.7936 | 0.0245 | 4.01E-81 | 849 | 212.649 | 636.351 | 172 | 144.894 | 27.1059 |
| Alkyl | GMYN | 0.0526 | 2.1569 | 0.0244 | 3.92E-41 | 420 | 103.032 | 316.968 | 89 | 72.9183 | 16.0817 |
| Alkyl | MS | 0.0498 | 2.0586 | 0.0242 | 1.22E-50 | 444 | 125.475 | 318.525 | 85 | 80.0813 | 4.91868 |
| GroL | GMYN | 0.0126462 | 0.531829 | 0.0237787 | 2.22E-69 | 1494 | 456.241 | 1037.76 | 163 | 149.998 | 13.0021 |
| Alkyl | MS | 0.0605 | 2.5784 | 0.0235 | 5.54E-73 | 459 | 112.785 | 346.215 | 102 | 95.1455 | 6.85449 |
| DnaK | MS | 0.0176324 | 0.752496 | 0.0234318 | 2.69E-118 | 1764 | 495.011 | 1268.99 | 226 | 213.194 | 12.8063 |
| Hyp | MS | 0.0425 | 1.8575 | 0.0229 | 1.78E-19 | 204 | 53.2076 | 150.792 | 30 | 28.1741 | 1.82585 |
| Piri | GMYN | 0.1387 | 6.1000 | 0.0227 | 6.44E-48 | 675 | 146.142 | 528.858 | 176 | 109.574 | 66.426 |
| Piri | GMYN | 0.1387 | 6.1000 | 0.0227 | 6.44E-48 | 675 | 146.142 | 528.858 | 176 | 109.574 | 66.426 |
| RpoE | MS | 0.0271 | 1.1903 | 0.0227 | 2.71E-52 | 534 | 117.046 | 416.954 | 78 | 72.1561 | 5.84387 |
| Carbo | GMYN | 0.0806866 | 3.5869 | 0.0224948 | 5.76E-63 | 522 | 119.406 | 402.594 | 133 | 102.329 | 30.6713 |
| GroL | MS | 0.0132982 | 0.594931 | 0.0223525 | 7.80E-85 | 1494 | 418.369 | 1075.63 | 163 | 154.142 | 8.85828 |
| Carbo | MS | 0.0692 | 3.1345 | 0.0221 | 1.38E-109 | 624 | 144.509 | 479.491 | 142 | 132.305 | 9.6952 |
| Alkyl | MS | 0.0366 | 1.6574 | 0.0221 | 7.85E-52 | 405 | 96.7634 | 308.237 | 75 | 70.0759 | 4.92413 |
| Rubre | GMYN | 0.0244 | 1.1113 | 0.0219 | 1.15E-38 | 483 | 148.491 | 334.509 | 86 | 77.9986 | 8.00139 |
| Rubre | MS | 0.0714 | 3.2578 | 0.0219 | 7.09E-66 | 438 | 127.645 | 310.355 | 105 | 99.6893 | 5.3107 |
| RpoH | GMYN | 0.0298 | 1.3640 | 0.0218 | 1.54E-80 | 903 | 214.772 | 688.228 | 155 | 134.944 | 20.0557 |
| Fimb | GMYN | 0.0788 | 3.6154 | 0.0218 | 5.38E-53 | 510 | 124.028 | 385.972 | 125 | 96.2343 | 28.7657 |
| RpoH | GMYN | 0.0556 | 2.5579 | 0.0217 | 4.37E-92 | 897 | 235.377 | 661.623 | 206 | 170.703 | 35.2973 |
| RpoH | GMYN | 0.0880 | 4.0555 | 0.0217 | 8.64E-83 | 885 | 223.028 | 661.972 | 222 | 167.291 | 54.7091 |
| DnaK | MS | 0.0191 | 0.8982 | 0.0213 | 1.60E-137 | 1893 | 554.357 | 1338.64 | 266 | 252.992 | 13.0085 |
| Superox | MS | 0.0476 | 2.2384 | 0.0213 | 3.12E-56 | 429 | 118.796 | 310.204 | 89 | 84.3182 | 4.68179 |
| Fimb | MS | 0.0778 | 3.6881 | 0.0211 | 1.30E-91 | 510 | 130.276 | 379.724 | 125 | 117.764 | 7.23638 |
| Fatty | GMYN | 0.0510 | 2.4188 | 0.0211 | 1.44E-107 | 1101 | 257.707 | 843.293 | 227 | 185.597 | 41.4029 |

| | | | | | | | | | | | |
|---|---|---|---|---|---|---|---|---|---|---|---|
| DnaK | GMYN | 0.0160 | 0.7616 | 0.0210 | 6.21E-123 | 1866 | 585.487 | 1280.51 | 277 | 256.799 | 20.2014 |
| FtsK | GMYN | 0.0184 | 0.8855 | 0.0208 | 1.53E-158 | 2271 | 671.97 | 1599.03 | 347 | 317.888 | 29.1116 |
| Alkyl | MS | 0.0490 | 2.3656 | 0.0207 | 4.64E-65 | 420 | 99.9824 | 320.018 | 89 | 83.4638 | 5.53621 |
| Carbo | MS | 0.0490 | 2.3829 | 0.0206 | 7.60E-86 | 579 | 137.615 | 441.385 | 120 | 112.579 | 7.42133 |
| RpoE | GMYN | 0.0228 | 1.1131 | 0.0205 | 3.71E-32 | 465 | 106.014 | 358.986 | 62 | 53.9583 | 8.04168 |
| GroL | GMYN | 0.0163 | 0.7967 | 0.0204 | 2.08E-97 | 1590 | 408.032 | 1181.97 | 198 | 178.996 | 19.0042 |
| RpoE | GMYN | 0.0224 | 1.1076 | 0.0202 | 2.31E-39 | 543 | 131.462 | 411.538 | 77 | 67.9305 | 9.06949 |
| RpoH | MS | 0.0443 | 2.2379 | 0.0198 | 1.43E-101 | 849 | 248.834 | 600.166 | 172 | 164.159 | 7.84083 |
| Superox | GMYN | 0.0486 | 2.4670 | 0.0197 | 4.26E-34 | 429 | 128.64 | 300.36 | 89 | 74.9082 | 14.0918 |
| Carbo | GMYN | 0.0387 | 1.9678 | 0.0197 | 2.17E-60 | 603 | 141.779 | 461.221 | 116 | 98.6223 | 17.3777 |
| Rubre | GMYN | 0.0345 | 1.7929 | 0.0192 | 2.66E-38 | 462 | 134.721 | 327.279 | 88 | 77 | 11 |
| Rubre | GMYN | 0.0345 | 1.7929 | 0.0192 | 2.66E-38 | 462 | 134.721 | 327.279 | 88 | 77 | 11 |
| Carbo | GMYN | 0.0699 | 3.6402 | 0.0192 | 1.67E-72 | 624 | 128.208 | 495.792 | 142 | 109.07 | 32.9296 |
| FtsK | GMYN | 0.0191907 | 1.01163 | 0.01897 | 1.43E-159 | 2142 | 655.893 | 1486.11 | 352 | 323.879 | 28.1214 |
| RpoH | GMYN | 0.0309 | 1.6432 | 0.0188 | 1.47E-78 | 822 | 188.964 | 633.036 | 145 | 125.878 | 19.1225 |
| FtsK | MS | 0.0308 | 1.6376 | 0.0188 | 7.34E-283 | 2265 | 600.328 | 1664.67 | 448 | 425.805 | 22.1948 |
| FtsK | GMYN | 0.0229 | 1.2204 | 0.0188 | 1.41E-185 | 2292 | 685.669 | 1606.33 | 405 | 368.807 | 36.1931 |
| RpoH | GMYN | 0.0616 | 3.2919 | 0.0187 | 1.53E-90 | 906 | 220.911 | 685.089 | 204 | 163.627 | 40.3726 |
| Fimb | MS | 0.0581 | 3.1576 | 0.0184 | 5.75E-82 | 504 | 129.606 | 374.394 | 114 | 108.243 | 5.75668 |
| Carbo | GMYN | 0.0295 | 1.6056 | 0.0183 | 9.12E-56 | 558 | 138.878 | 419.122 | 104 | 91.9133 | 12.0867 |
| DnaK | MS | 0.0185 | 1.0103 | 0.0183 | 4.45E-141 | 1830 | 550.86 | 1279.14 | 267 | 256.134 | 10.8657 |
| FtsK | GMYN | 0.0248 | 1.3630 | 0.0182 | 7.24E-214 | 2205 | 599.942 | 1605.06 | 416 | 376.857 | 39.1426 |
| Carbo | GMYN | 0.0653 | 3.6205 | 0.0180 | 5.11E-98 | 627 | 124.875 | 502.125 | 154 | 122.929 | 31.0711 |
| DnaK | MS | 0.0163 | 0.9043 | 0.0180 | 3.02E-146 | 1917 | 539.196 | 1377.8 | 265 | 253.358 | 11.6424 |
| Carbo | GMYN | 0.0554 | 3.0948 | 0.0179 | 4.18E-65 | 615 | 158.128 | 456.872 | 141 | 116.682 | 24.3185 |
| Alkyl | GMYN | 0.0621 | 3.4839 | 0.0178 | 1.74E-48 | 459 | 104.089 | 354.911 | 102 | 80.9172 | 21.0828 |
| FtsK | GMYN | 0.0317 | 1.7850 | 0.0178 | 4.45E-232 | 2265 | 581.485 | 1683.51 | 448 | 395.749 | 52.2514 |
| GroL | GMYN | 0.0239 | 1.3477 | 0.0177 | 5.70E-89 | 1596 | 357.271 | 1238.73 | 191 | 161.964 | 29.0361 |
| FtsK | MS | 0.0222 | 1.2567 | 0.0177 | 1.08E-233 | 2292 | 651.722 | 1640.28 | 405 | 387.739 | 17.2606 |
| FtsK | MS | 0.0186167 | 1.07178 | 0.0173699 | 9.31E-196 | 2142 | 623.939 | 1518.06 | 352 | 337.727 | 14.2728 |
| Carbo | GMYN | 0.0644 | 3.7198 | 0.0173 | 2.17E-69 | 639 | 142.548 | 496.452 | 144 | 113.49 | 30.5099 |
| RpoE | MS | 0.0153 | 0.8914 | 0.0172 | 4.23E-44 | 507 | 115.697 | 391.303 | 66 | 62.3697 | 3.63032 |
| DnaK | MS | 0.0142 | 0.8334 | 0.0170 | 3.87E-139 | 1878 | 517.35 | 1360.65 | 248 | 237.374 | 10.626 |
| DnaK | MS | 0.0176 | 1.0387 | 0.0169 | 4.08E-156 | 1890 | 548.263 | 1341.74 | 284 | 272.71 | 11.2896 |

| | | | | | | | | | | | |
|---|---|---|---|---|---|---|---|---|---|---|---|
| Alkyl | MS | 0.0477 | 2.8312 | 0.0169 | 1.18E-67 | 387 | 80.7862 | 306.214 | 79 | 74.2535 | 4.74646 |
| GroL | MS | 0.0099 | 0.6139 | 0.0162 | 9.83E-92 | 1545 | 435.919 | 1109.08 | 169 | 162.326 | 6.67375 |
| Fatty | GMYN | 0.0672 | 4.1833 | 0.0161 | 2.82E-133 | 1182 | 264.481 | 917.519 | 276 | 217.206 | 58.7944 |
| DnaK | MS | 0.0121 | 0.7511 | 0.0161 | 3.00E-127 | 1890 | 547.109 | 1342.89 | 238 | 228.969 | 9.03078 |
| Fatty | GMYN | 0.0672 | 4.1854 | 0.0161 | 7.91E-112 | 1161 | 265.218 | 895.782 | 257 | 199.624 | 57.3765 |
| DnaK | MS | 0.0120 | 0.7478 | 0.0161 | 6.32E-128 | 1899 | 549.353 | 1349.65 | 239 | 229.928 | 9.07215 |
| Fimb | GMYN | 0.0580 | 3.6129 | 0.0160 | 3.61E-53 | 504 | 123.612 | 380.388 | 114 | 92.8365 | 21.1635 |
| DnaK | MS | 0.0096 | 0.5981 | 0.0160 | 2.37E-111 | 1896 | 497.188 | 1398.81 | 197 | 188.51 | 8.48977 |
| FtsK | MS | 0.0252 | 1.5749 | 0.0160 | 1.24E-259 | 2205 | 596.985 | 1608.01 | 416 | 398.838 | 17.1616 |
| Carbo | MS | 0.0658 | 4.1225 | 0.0160 | NA | 627 | 144.387 | 482.613 | 154 | 146.196 | 7.80388 |
| Carbo | MS | 0.0326 | 2.0699 | 0.0158 | 4.94E-94 | 654 | 149.461 | 504.539 | 126 | 119.631 | 6.36862 |
| RpoH | GMYN | 0.0530 | 3.3685 | 0.0157 | 1.12E-92 | 888 | 218.22 | 669.78 | 196 | 161.831 | 34.1686 |
| Carbo | MS | 0.0640 | 4.0729 | 0.0157 | 4.79E-129 | 630 | 147.092 | 482.908 | 151 | 143.591 | 7.40934 |
| Fimb | GMYN | 0.0562 | 3.6049 | 0.0156 | 4.11E-54 | 498 | 122.299 | 375.701 | 113 | 92.7167 | 20.2833 |
| Fimb | GMYN | 0.0562 | 3.6049 | 0.0156 | 4.11E-54 | 498 | 122.299 | 375.701 | 113 | 92.7167 | 20.2833 |
| DnaK | MS | 0.0160 | 1.0334 | 0.0155 | 1.01E-143 | 1866 | 583.693 | 1282.31 | 277 | 267.866 | 9.13392 |
| RpoH | MS | 0.0176 | 1.1343 | 0.0155 | 6.01E-84 | 891 | 253.7 | 637.3 | 144 | 138.604 | 5.39645 |
| Fimb | GMYN | 0.0563 | 3.6370 | 0.0155 | 2.90E-60 | 480 | 127.652 | 352.348 | 122 | 102.96 | 19.0403 |
| Fatty | GMYN | 0.0629 | 4.1281 | 0.0152 | 1.96E-134 | 1158 | 245.709 | 912.291 | 261 | 206.176 | 54.8241 |
| DnaK | MS | 0.0109 | 0.7159 | 0.0152 | 2.12E-122 | 1824 | 524.785 | 1299.22 | 226 | 217.802 | 8.19835 |
| Fimb | GMYN | 0.0545 | 3.5996 | 0.0151 | 4.43E-54 | 465 | 121.442 | 343.558 | 112 | 93.9862 | 18.0138 |
| Carbo | GMYN | 0.0215 | 1.4251 | 0.0151 | 3.55E-64 | 639 | 162.959 | 476.041 | 114 | 103.941 | 10.0589 |
| Fimb | MS | 0.0550 | 3.6437 | 0.0151 | 3.75E-96 | 480 | 126.395 | 353.605 | 122 | 117.054 | 4.94559 |
| Fimb | GMYN | 0.0541 | 3.5853 | 0.0151 | 7.74E-54 | 489 | 119.156 | 369.844 | 110 | 90.7937 | 19.2063 |
| Fatty | GMYN | 0.0623 | 4.1310 | 0.0151 | 1.00E-135 | 1167 | 246.651 | 920.349 | 262 | 207.187 | 54.8133 |
| DnaK | MS | 0.0084 | 0.5584 | 0.0150 | 5.98E-105 | 1857 | 534.311 | 1322.69 | 195 | 188.012 | 6.98776 |
| Fatty | GMYN | 0.0598 | 4.0524 | 0.0148 | 7.34E-151 | 1137 | 222.11 | 914.89 | 256 | 203.582 | 52.4181 |
| RpoE | GMYN | 0.0268 | 1.8250 | 0.0147 | 8.89E-43 | 534 | 112.498 | 421.502 | 78 | 66.9366 | 11.0634 |
| Fimb | MS | 0.0559 | 3.8195 | 0.0146 | 1.50E-86 | 498 | 124.921 | 373.079 | 113 | 108.266 | 4.73378 |
| Fimb | MS | 0.0559 | 3.8195 | 0.0146 | 1.50E-86 | 498 | 124.921 | 373.079 | 113 | 108.266 | 4.73378 |
| DnaK | GMYN | 0.0182 | 1.2457 | 0.0146 | 3.65E-135 | 1893 | 483.934 | 1409.07 | 266 | 240.74 | 25.2598 |
| FtsK | MS | 0.0150 | 1.0321 | 0.0146 | 5.19E-209 | 2268 | 620.198 | 1647.8 | 353 | 339.852 | 13.1481 |
| Rubre | MS | 0.0488 | 3.3624 | 0.0145 | 5.52E-58 | 450 | 129.181 | 320.819 | 92 | 88.8017 | 3.19828 |
| FtsK | MS | 0.0149 | 1.0308 | 0.0145 | 1.69E-209 | 2277 | 623.013 | 1653.99 | 354 | 340.878 | 13.1224 |

| | | | | | | | | | | | |
|---|---|---|---|---|---|---|---|---|---|---|---|
| Fatty | GMYN | 0.0590 | 4.0728 | 0.0145 | 2.00E-127 | 1092 | 228.241 | 863.759 | 240 | 191.199 | 48.8012 |
| FtsK | MS | 0.0134 | 0.9306 | 0.0144 | 1.17E-188 | 2232 | 655.371 | 1576.63 | 339 | 327.62 | 11.3798 |
| Fimb | MS | 0.0537 | 3.7249 | 0.0144 | 3.32E-86 | 465 | 119.856 | 345.144 | 112 | 107.533 | 4.46737 |
| Fatty | GMYN | 0.0594 | 4.1670 | 0.0143 | 1.01E-106 | 1125 | 240.517 | 884.483 | 230 | 179.691 | 50.3093 |
| Fimb | MS | 0.0546 | 3.8451 | 0.0142 | 4.05E-84 | 489 | 121.966 | 367.034 | 110 | 105.496 | 4.50405 |
| Carbo | MS | 0.0362 | 2.6543 | 0.0137 | 3.55E-88 | 603 | 141.123 | 461.877 | 116 | 111.038 | 4.9616 |
| Rubre | MS | 0.0367 | 2.7185 | 0.0135 | 6.40E-50 | 462 | 147.886 | 314.114 | 88 | 85.5493 | 2.45071 |
| Rubre | MS | 0.0367 | 2.7185 | 0.0135 | 6.40E-50 | 462 | 147.886 | 314.114 | 88 | 85.5493 | 2.45071 |
| GroL | MS | 0.0098 | 0.7316 | 0.0134 | 6.69E-107 | 1635 | 445.156 | 1189.84 | 187 | 180.532 | 6.46815 |
| RuvB | GMYN | 0.0176 | 1.3142 | 0.0134 | 2.34E-104 | 1032 | 282.23 | 749.77 | 188 | 174.974 | 13.0262 |
| Carbo | MS | 0.0217 | 1.6203 | 0.0134 | 1.31E-73 | 639 | 169.63 | 469.37 | 114 | 109.927 | 4.07271 |
| DnaK | GMYN | 0.0156 | 1.1806 | 0.0132 | 2.47E-141 | 1917 | 483.017 | 1433.98 | 265 | 242.828 | 22.1722 |
| Carbo | MS | 0.0266 | 2.0222 | 0.0131 | 4.16E-79 | 558 | 127.269 | 430.731 | 104 | 99.5746 | 4.42543 |
| Alkyl | MS | 0.0224 | 1.7097 | 0.0131 | 9.32E-46 | 363 | 80.6686 | 282.331 | 60 | 57.3708 | 2.62924 |
| FtsK | GMYN | 0.0298 | 2.2884 | 0.0130 | 1.48E-191 | 2256 | 569.943 | 1686.06 | 392 | 342.76 | 49.2405 |
| FtsK | GMYN | 0.0133 | 1.0223 | 0.0130 | 1.03E-174 | 2232 | 635.419 | 1596.58 | 339 | 317.958 | 21.042 |
| DnaK | GMYN | 0.0115 | 0.8884 | 0.0130 | 2.93E-129 | 1899 | 489.13 | 1409.87 | 239 | 222.916 | 16.0843 |
| DnaK | GMYN | 0.0161 | 1.2408 | 0.0130 | 1.65E-172 | 1890 | 429.855 | 1460.14 | 284 | 260.749 | 23.2506 |
| RpoH | GMYN | 0.0378 | 2.9140 | 0.0130 | 1.69E-78 | 912 | 230.945 | 681.055 | 169 | 143.987 | 25.0134 |
| RuvB | MS | 0.0195 | 1.5113 | 0.0129 | 1.17E-122 | 1032 | 274.545 | 757.455 | 188 | 181.547 | 6.45305 |
| DnaK | GMYN | 0.0116 | 0.9016 | 0.0128 | 2.25E-129 | 1890 | 483.897 | 1406.1 | 238 | 221.917 | 16.0828 |
| GroL | GMYN | 0.0095 | 0.7438 | 0.0128 | 2.28E-94 | 1545 | 378.873 | 1166.13 | 169 | 157.998 | 11.0017 |
| Alkyl | GMYN | 0.0421388 | 3.35097 | 0.0125751 | 2.70E-46 | 387 | 87.1785 | 299.822 | 81 | 68.7847 | 12.2153 |
| Carbo | GMYN | 0.0327 | 2.6131 | 0.0125 | 2.31E-71 | 654 | 151.246 | 502.754 | 126 | 109.954 | 16.0458 |
| Hfq | MS | 0.0118487 | 0.954838 | 0.0124092 | 2.40E-20 | 234 | 61.682 | 172.318 | 32 | 30.9278 | 1.07217 |
| Carbo | MS | 0.0513 | 4.1831 | 0.0123 | 6.84E-123 | 615 | 141.208 | 473.792 | 141 | 135.426 | 5.57402 |
| RuvB | MS | 0.0216 | 1.7691 | 0.0122 | 7.10E-134 | 1014 | 262.888 | 751.112 | 195 | 188.427 | 6.57318 |
| RuvB | MS | 0.0248 | 2.0379 | 0.0122 | 7.82E-130 | 978 | 256.851 | 721.149 | 191 | 184.694 | 6.30582 |
| Alkyl | MS | 0.0309 | 2.5430 | 0.0121 | 3.46E-65 | 444 | 108.251 | 335.749 | 88 | 84.8045 | 3.19553 |
| Alkyl | MS | 0.0309 | 2.5430 | 0.0121 | 3.46E-65 | 444 | 108.251 | 335.749 | 88 | 84.8045 | 3.19553 |
| RpoH | GMYN | 0.0158 | 1.3085 | 0.0121 | 1.04E-84 | 855 | 214.857 | 640.143 | 143 | 132.991 | 10.0092 |
| DnaK | GMYN | 0.0091 | 0.7509 | 0.0121 | 2.72E-114 | 1896 | 439.163 | 1456.84 | 197 | 183.896 | 13.1045 |
| RuvB | GMYN | 0.0213 | 1.8020 | 0.0118 | 1.39E-109 | 978 | 257.85 | 720.15 | 191 | 175.891 | 15.1094 |
| Rubre | MS | 0.0362 | 3.0609 | 0.0118 | 1.38E-42 | 381 | 121.404 | 259.596 | 72 | 70.2252 | 1.7748 |

| | | | | | | | | | | | |
|---|---|---|---|---|---|---|---|---|---|---|---|
| RuvB | GMYN | 0.0233258 | 1.98957 | 0.0117241 | 1.43E-100 | 921 | 219.415 | 701.585 | 169 | 152.967 | 16.0331 |
| RuvB | GMYN | 0.0161 | 1.3747 | 0.0117 | 1.25E-101 | 963 | 268.211 | 694.789 | 180 | 168.983 | 11.0174 |
| RpoH | MS | 0.0165 | 1.4136 | 0.0117 | 2.64E-88 | 855 | 237.894 | 617.106 | 143 | 138.795 | 4.20463 |
| DnaK | GMYN | 0.0162898 | 1.39952 | 0.0116396 | 1.41E-128 | 1764 | 390.355 | 1373.64 | 226 | 203.891 | 22.1088 |
| FtsK | MS | 0.0125 | 1.0848 | 0.0115 | 8.69E-202 | 2202 | 661.527 | 1540.47 | 356 | 346.733 | 9.26654 |
| GroL | MS | 0.0060 | 0.5348 | 0.0113 | 1.07E-88 | 1611 | 457.404 | 1153.6 | 160 | 155.571 | 4.4293 |
| GroL | MS | 0.0060 | 0.5393 | 0.0111 | 7.01E-90 | 1620 | 460.493 | 1159.51 | 162 | 157.582 | 4.41769 |
| Hyp | GMYN | 0.0335 | 3.0362 | 0.0110 | 3.22E-19 | 162 | 39.3534 | 122.647 | 33 | 29 | 4 |
| DnaK | GMYN | 0.0079 | 0.7183 | 0.0110 | 7.00E-114 | 1857 | 443.221 | 1413.78 | 195 | 183.936 | 11.0639 |
| Alkyl | GMYN | 0.0258 | 2.3727 | 0.0109 | 8.14E-33 | 363 | 86.3952 | 276.605 | 60 | 52.9968 | 7.00321 |
| GroL | GMYN | 0.0096 | 0.8940 | 0.0108 | 1.68E-111 | 1635 | 377.041 | 1257.96 | 187 | 174.998 | 12.0019 |
| GroL | MS | 0.0043 | 0.4033 | 0.0107 | 1.37E-72 | 1614 | 461.508 | 1152.49 | 133 | 129.544 | 3.45647 |
| FtsK | GMYN | 0.0139 | 1.3083 | 0.0106 | 7.42E-195 | 2277 | 606.13 | 1670.87 | 354 | 330.962 | 23.038 |
| FtsK | GMYN | 0.0140 | 1.3271 | 0.0105 | 1.31E-194 | 2268 | 602.647 | 1665.35 | 353 | 329.962 | 23.0377 |
| RuvB | MS | 0.0161 | 1.5556 | 0.0104 | 3.38E-115 | 963 | 272.316 | 690.684 | 180 | 175.384 | 4.61607 |
| RuvB | GMYN | 0.0154 | 1.4935 | 0.0103 | 1.32E-108 | 996 | 268.642 | 727.358 | 185 | 173.976 | 11.0241 |
| Fimb | GMYN | 0.0311 | 3.0580 | 0.0102 | 4.80E-51 | 417 | 110.971 | 306.029 | 91 | 81.8172 | 9.18276 |
| RuvB | MS | 0.0154 | 1.5575 | 0.0099 | 1.30E-124 | 996 | 268.221 | 727.779 | 185 | 180.16 | 4.84025 |
| Rubre | MS | 0.0245 | 2.5196 | 0.0097 | 5.37E-53 | 483 | 146.595 | 336.405 | 86 | 84.1211 | 1.87894 |
| Fimb | GMYN | 0.0335 | 3.5954 | 0.0093 | 1.69E-62 | 465 | 120.768 | 344.232 | 107 | 95.9016 | 11.0984 |
| GroL | GMYN | 0.0058 | 0.6384 | 0.0091 | 7.74E-94 | 1611 | 398.76 | 1212.24 | 160 | 152.999 | 7.00051 |
| RpoH | GMYN | 0.0168 | 1.8747 | 0.0090 | 1.14E-81 | 891 | 226.384 | 664.616 | 144 | 132.978 | 11.0217 |
| GroL | GMYN | 0.0058 | 0.6509 | 0.0088 | 1.40E-95 | 1620 | 399.484 | 1220.52 | 162 | 155 | 7.00048 |
| DnaK | GMYN | 0.0133 | 1.5135 | 0.0088 | 5.94E-149 | 1878 | 423.783 | 1454.22 | 248 | 228.79 | 19.2105 |
| RuvB | MS | 0.0289 | 3.2835 | 0.0088 | 5.08E-168 | 1008 | 224.501 | 783.499 | 202 | 195.973 | 6.02699 |
| Hfq | MS | 0.0128 | 1.4557 | 0.0088 | 0 | 207 | 55.9527 | 151.047 | 32 | 31.2594 | 0.740612 |
| DnaK | GMYN | 0.0168 | 1.9225 | 0.0087 | 2.32E-155 | 1830 | 423.986 | 1406.01 | 267 | 243.718 | 23.2819 |
| Fimb | MS | 0.0307 | 3.5496 | 0.0087 | 3.34E-64 | 417 | 114.861 | 302.139 | 91 | 88.974 | 2.02604 |
| FtsK | GMYN | 0.0123 | 1.4236 | 0.0086 | 3.02E-189 | 2202 | 635.725 | 1566.27 | 356 | 336.975 | 19.0248 |
| Alkyl | MS | 0.0374733 | 4.36149 | 0.00859187 | 4.13E-71 | 387 | 86.1463 | 300.854 | 81 | 78.6403 | 2.35967 |
| RuvB | MS | 0.0226 | 2.6409 | 0.0085 | 3.67E-159 | 1032 | 265.99 | 766.01 | 216 | 210.81 | 5.18987 |
| RuvB | MS | 0.0222919 | 2.61062 | 0.00853893 | 3.84E-144 | 921 | 191.99 | 729.01 | 169 | 163.693 | 5.30748 |
| GroL | GMYN | 0.0237 | 2.8105 | 0.0084 | 2.00E-110 | 1608 | 363.96 | 1244.04 | 219 | 189.962 | 29.0379 |
| Fimb | MS | 0.0323 | 3.8914 | 0.0083 | 5.03E-88 | 465 | 116.601 | 348.399 | 107 | 104.409 | 2.5913 |

| | | | | | | | | | | | |
|---|---|---|---|---|---|---|---|---|---|---|---|
| GroL | MS | 0.0038 | 0.4759 | 0.0080 | 2.62E-82 | 1575 | 433.088 | 1141.91 | 140 | 137.114 | 2.88573 |
| Alkyl | GMYN | 0.0333 | 4.3104 | 0.0077 | 4.72E-51 | 444 | 102.89 | 341.11 | 88 | 76.9209 | 11.0791 |
| Alkyl | GMYN | 0.0333 | 4.3104 | 0.0077 | 4.72E-51 | 444 | 102.89 | 341.11 | 88 | 76.9209 | 11.0791 |
| Hyp | MS | 0.0329 | 4.2779 | 0.0077 | 0 | 162 | 36.9056 | 125.094 | 33 | 32.1607 | 0.839256 |
| Hfq | GMYN | 0.0068 | 0.8991 | 0.0075 | 4.58E-17 | 219 | 70.2969 | 148.703 | 32 | 31 | 1 |
| RuvB | MS | 0.0095 | 1.2635 | 0.0075 | 9.59E-112 | 1029 | 286.963 | 742.037 | 174 | 170.69 | 3.31038 |
| GroL | GMYN | 0.0041 | 0.5418 | 0.0075 | 1.55E-81 | 1614 | 378.782 | 1235.22 | 133 | 128 | 5.00025 |
| RpoE | GMYN | 0.0148 | 2.0241 | 0.0073 | 7.32E-45 | 507 | 97.0133 | 409.987 | 66 | 59.9993 | 6.00071 |
| RuvB | GMYN | 0.0289 | 4.0509 | 0.0071 | 1.21E-130 | 1008 | 221.661 | 786.339 | 202 | 179.927 | 22.0727 |
| Hfq | MS | 0.0067 | 0.9608 | 0.0070 | 0 | 219 | 65.6215 | 153.379 | 32 | 31.4868 | 0.51322 |
| RuvB | MS | 0.0229 | 3.3218 | 0.0069 | 1.49E-179 | 1053 | 255.307 | 797.693 | 222 | 217.314 | 4.68602 |
| Hfq | GMYN | 0.0131 | 2.0777 | 0.0063 | 8.37E-20 | 207 | 52.0075 | 154.992 | 32 | 29.9994 | 2.00059 |
| DnaK | GMYN | 0.0102 | 1.6663 | 0.0061 | 1.63E-134 | 1824 | 431.263 | 1392.74 | 226 | 211.953 | 14.0469 |
| Hfq | GMYN | 0.0166 | 2.8099 | 0.0059 | 7.18E-24 | 165 | 42.3758 | 122.624 | 35 | 33 | 2 |
| Hfq | GMYN | 0.0112376 | 2.02923 | 0.00553787 | 4.09E-21 | 234 | 53.0886 | 180.911 | 32 | 29.9916 | 2.00839 |
| RuvB | GMYN | 0.0227 | 4.1202 | 0.0055 | 7.49E-153 | 1053 | 243.125 | 809.875 | 222 | 203.947 | 18.0532 |
| RuvB | GMYN | 0.0218 | 4.0990 | 0.0053 | 8.15E-151 | 1032 | 236.349 | 795.651 | 216 | 198.91 | 17.0905 |
| RuvB | GMYN | 0.0207 | 4.0690 | 0.0051 | 3.83E-132 | 1014 | 227.079 | 786.921 | 195 | 178.926 | 16.0736 |
| RuvB | GMYN | 0.0094 | 1.8698 | 0.0050 | 1.27E-105 | 1029 | 278.984 | 750.016 | 174 | 166.992 | 7.00753 |
| GroL | GMYN | 0.0033 | 0.7345 | 0.0045 | 3.09E-89 | 1575 | 369.8 | 1205.2 | 140 | 136 | 4.00009 |
| Hfq | MS | 0.0162 | 4.1595 | 0.0039 | 0 | 165 | 39.1784 | 125.822 | 35 | 34.568 | 0.431958 |
| Hfq | MS | 0.0000 | 0.9622 | 0.0000 | 0 | 189 | 54.6278 | 134.372 | 24 | 23.9999 | 5.90E-05 |
| Cold | MS | 0.0000 | 0.3117 | 0.0000 | 0 | 171 | 45.7305 | 125.269 | 10 | 9.99997 | 2.74E-05 |
| Cold | MS | 0.0000 | 0.1806 | 0.0000 | 0 | 168 | 48.6253 | 119.375 | 7 | 6.99998 | 1.72E-05 |
| Cold | MS | 0.0000 | 0.1975 | 0.0000 | 0 | 156 | 45.9512 | 110.049 | 7 | 6.99998 | 1.68E-05 |
| Cold | MS | 0.0000 | 0.1104 | 0.0000 | 0 | 195 | 45.2978 | 149.702 | 5 | 4.99998 | 1.65E-05 |
| Cold | MS | 0.0000 | 0.1302 | 0.0000 | 0 | 171 | 38.4118 | 132.588 | 5 | 4.99998 | 1.73E-05 |
| Cold | MS | 5.04E-08 | 0.0504104 | 1.00E-06 | 0 | 195 | 65.3257 | 129.674 | 3 | 2.99999 | 5.96E-06 |
| Cold | GMYN | 1.11E-15 | 0.0675801 | 1.64E-14 | 0 | 195 | 47.762 | 147.238 | 3 | 3 | 0 |
| Cold | GMYN | 0.0000 | 0.2167 | 0.0000 | 0 | 168 | 40.1796 | 127.82 | 7 | 7 | 0 |
| Hfq | GMYN | 0.0000 | 1.0078 | 0.0000 | 0 | 189 | 43.2947 | 145.705 | 24 | 24 | 0 |
| Cold | GMYN | 0.0000 | 0.2277 | 0.0000 | 0 | 156 | 38.2496 | 117.75 | 7 | 7 | 0 |
| Cold | GMYN | 0.0000 | 0.1473 | 0.0000 | 0 | 171 | 38.8647 | 132.135 | 5 | 5 | 0 |

**Housekeeping genes  in PnecC subcluster**

| Sequence | Method | Ka | Ks | Ka/Ks | P-Value(Fisher) | Length | S-Sites | N-Sites | Substitutions | S-Substitutions | N-Substitutions |
|---|---|---|---|---|---|---|---|---|---|---|---|
| ndk | GMYN | 0.0382 | 0.4817 | 0.0794 | 1.12E-13 | 411 | 98.3222 | 312.678 | 43 | 31.3974 | 11.6026 |
| ndk | MS | 0.0300 | 0.5242 | 0.0573 | 2.44E-19 | 423 | 112.061 | 310.939 | 45 | 38.829 | 6.17102 |
| ndk | MS | 0.0349 | 0.6510 | 0.0537 | 7.66E-23 | 411 | 78.9238 | 332.076 | 43 | 35.0774 | 7.92264 |
| ndk | GMYN | 0.0330 | 0.6499 | 0.0508 | 1.03E-19 | 420 | 102.274 | 317.726 | 51 | 40.795 | 10.205 |
| adk | MS | 0.0423 | 0.8510 | 0.0497 | 2.46E-46 | 663 | 175.563 | 487.437 | 100 | 87.8641 | 12.1359 |
| glnA | MS | 0.0245 | 0.4959 | 0.0494 | 3.22E-65 | 1413 | 347.415 | 1065.59 | 144 | 125.049 | 18.9507 |
| acsA | MS | 0.0451 | 0.9224 | 0.0489 | 1.22E-147 | 1968 | 505.133 | 1462.87 | 312 | 273.294 | 38.7059 |
| ndk | GMYN | 0.0281 | 0.6091 | 0.0461 | 1.96E-18 | 423 | 94.9409 | 328.059 | 45 | 35.9672 | 9.03279 |
| adk | MS | 0.0394 | 0.8747 | 0.0450 | 1.55E-47 | 651 | 167.084 | 483.916 | 95 | 84.0386 | 10.9614 |
| ndk | MS | 0.0321 | 0.7148 | 0.0448 | 1.13E-25 | 420 | 101.298 | 318.702 | 51 | 44.6942 | 6.30577 |
| adk | MS | 0.0582 | 1.3849 | 0.0421 | 6.22E-68 | 660 | 172.253 | 487.747 | 127 | 113.485 | 13.5149 |
| glnA | MS | 0.0234 | 0.5632 | 0.0415 | 9.55E-74 | 1413 | 379.737 | 1033.26 | 161 | 144.653 | 16.3467 |
| glnA | MS | 0.0262 | 0.6366 | 0.0412 | 7.71E-80 | 1401 | 327.913 | 1073.09 | 159 | 140.12 | 18.8796 |
| acsA | GMYN | 0.0450 | 1.1273 | 0.0399 | 2.22E-115 | 1968 | 483.851 | 1484.15 | 312 | 247.275 | 64.7248 |
| ndk | GMYN | 0.0112 | 0.2822 | 0.0398 | 1.89E-10 | 354 | 83.7145 | 270.285 | 22 | 19 | 3 |
| ndk | MS | 0.0271 | 0.6862 | 0.0394 | 9.61E-26 | 423 | 92.2277 | 330.772 | 46 | 40.3017 | 5.69831 |
| acsA | MS | 0.0365 | 0.9353 | 0.0390 | 4.32E-149 | 1887 | 507.18 | 1379.82 | 300 | 271.218 | 28.782 |
| acsA | MS | 0.0332 | 0.8538 | 0.0389 | 3.33E-145 | 1971 | 529.54 | 1441.46 | 297 | 268.591 | 28.4093 |
| acsA | MS | 0.0271 | 0.7413 | 0.0365 | 1.82E-132 | 1971 | 537.997 | 1433 | 274 | 249.709 | 24.2911 |
| adk | GMYN | 0.0320 | 0.8913 | 0.0359 | 2.95E-37 | 594 | 172.334 | 421.666 | 94 | 80.8497 | 13.1503 |
| glnA | MS | 0.0212 | 0.5916 | 0.0358 | 1.08E-79 | 1413 | 366.88 | 1046.12 | 161 | 146.086 | 14.9141 |
| iso | MS | 0.0367 | 1.0320 | 0.0355 | 7.66E-192 | 2220 | 555.089 | 1664.91 | 358 | 323.532 | 34.4682 |
| acsA | MS | 0.0303 | 0.8617 | 0.0351 | 6.64E-147 | 1971 | 493.945 | 1477.05 | 281 | 254.288 | 26.7124 |
| acsA | MS | 0.0303 | 0.8617 | 0.0351 | 6.64E-147 | 1971 | 493.945 | 1477.05 | 281 | 254.288 | 26.7124 |
| adk | GMYN | 0.0356 | 1.0140 | 0.0351 | 3.30E-39 | 651 | 158.644 | 492.356 | 95 | 77.933 | 17.067 |
| glnA | MS | 0.0163 | 0.4674 | 0.0349 | 3.72E-65 | 1413 | 369.219 | 1043.78 | 136 | 123.8 | 12.2004 |
| glnA | MS | 0.0163 | 0.4674 | 0.0349 | 3.72E-65 | 1413 | 369.219 | 1043.78 | 136 | 123.8 | 12.2004 |
| acsA | MS | 0.0335 | 0.9679 | 0.0346 | 1.03E-150 | 1971 | 592.697 | 1378.3 | 318 | 294.294 | 23.7064 |
| glnA | MS | 0.0260 | 0.7578 | 0.0343 | 2.59E-97 | 1413 | 365.432 | 1047.57 | 189 | 172.082 | 16.9183 |
| iso | MS | 0.0449 | 1.3134 | 0.0342 | 7.77E-219 | 2232 | 594.613 | 1637.39 | 406 | 371.058 | 34.942 |
| adk | MS | 0.0250 | 0.7580 | 0.0330 | 1.77E-41 | 663 | 208.577 | 454.423 | 91 | 84.9041 | 6.09593 |

| | | | | | | | | | | | |
|---|---|---|---|---|---|---|---|---|---|---|---|
| cit | MS | 0.0228 | 0.6920 | 0.0329 | 1.20E-84 | 1311 | 342.568 | 968.432 | 167 | 152.781 | 14.2187 |
| ndk | MS | 0.0108 | 0.3335 | 0.0325 | 8.68E-12 | 354 | 84.6678 | 269.332 | 22 | 19.9386 | 2.06145 |
| acsA | MS | 0.0259 | 0.7986 | 0.0324 | 1.53E-138 | 1971 | 559.195 | 1411.81 | 283 | 261.597 | 21.4034 |
| iso | GMYN | 0.0436 | 1.3547 | 0.0322 | 1.17E-180 | 2232 | 533.201 | 1698.8 | 406 | 334.209 | 71.7912 |
| glnA | GMYN | 0.0212 | 0.6623 | 0.0321 | 1.63E-69 | 1413 | 355.65 | 1057.35 | 161 | 138.89 | 22.1102 |
| atpA | GMYN | 0.0118 | 0.3687 | 0.0320 | 5.77E-48 | 1470 | 357.029 | 1112.97 | 107 | 94 | 13 |
| iso | MS | 0.0377 | 1.1919 | 0.0317 | 4.93E-205 | 2229 | 613.557 | 1615.44 | 390 | 359.982 | 30.0182 |
| adk | MS | 0.0281 | 0.8887 | 0.0316 | 2.92E-49 | 663 | 190.603 | 472.397 | 96 | 89.0224 | 6.97763 |
| adk | MS | 0.0281 | 0.8887 | 0.0316 | 2.92E-49 | 663 | 190.603 | 472.397 | 96 | 89.0224 | 6.97763 |
| glnA | GMYN | 0.0231 | 0.7397 | 0.0312 | 5.55E-64 | 1413 | 301.413 | 1111.59 | 144 | 118.749 | 25.2505 |
| iso | GMYN | 0.0367 | 1.1967 | 0.0306 | 2.46E-178 | 2229 | 552.028 | 1676.97 | 390 | 330.066 | 59.9338 |
| succ | MS | 0.0249 | 0.8416 | 0.0296 | 3.14E-131 | 1776 | 464.767 | 1311.23 | 249 | 229.811 | 19.1886 |
| adk | GMYN | 0.0551 | 1.8739 | 0.0294 | 5.14E-57 | 660 | 144.849 | 515.151 | 127 | 99.7069 | 27.2931 |
| succ | MS | 0.0228 | 0.7962 | 0.0287 | 6.53E-126 | 1776 | 506.432 | 1269.57 | 250 | 233.227 | 16.7728 |
| iso | MS | 0.0246 | 0.8577 | 0.0286 | 5.64E-158 | 2148 | 607.868 | 1540.13 | 313 | 291.834 | 21.166 |
| cit | MS | 0.0190 | 0.6864 | 0.0277 | 1.37E-78 | 1242 | 345.667 | 896.333 | 155 | 144.602 | 10.3981 |
| glnA | GMYN | 0.0108 | 0.3915 | 0.0275 | 5.94E-53 | 1413 | 382.426 | 1030.57 | 118 | 106.994 | 11.0057 |
| iso | GMYN | 0.0244 | 0.8963 | 0.0272 | 3.15E-134 | 2148 | 596.948 | 1551.05 | 313 | 275.79 | 37.2096 |
| adk | MS | 0.0336 | 1.2359 | 0.0272 | 4.20E-55 | 594 | 152.53 | 441.47 | 94 | 87.1361 | 6.86386 |
| CTP | MS | 0.0383 | 1.4202 | 0.0270 | 1.53E-182 | 1659 | 419.317 | 1239.68 | 305 | 282.456 | 22.5437 |
| ndk | GMYN | 0.0263 | 0.9812 | 0.0268 | 2.48E-24 | 423 | 73.4068 | 349.593 | 46 | 36.9863 | 9.01365 |
| acsA | MS | 0.0254 | 0.9576 | 0.0265 | 4.92E-162 | 1971 | 532.638 | 1438.36 | 299 | 279 | 19.9996 |
| atpA | GMYN | 0.0099 | 0.3724 | 0.0265 | 1.84E-55 | 1539 | 417.596 | 1121.4 | 123 | 112 | 11 |
| iso | MS | 0.0241 | 0.9105 | 0.0265 | 6.78E-177 | 2232 | 583.67 | 1648.33 | 324 | 301.438 | 22.5624 |
| CTP | MS | 0.0441 | 1.6630 | 0.0265 | 5.29E-200 | 1659 | 406.643 | 1252.36 | 318 | 294.014 | 23.9861 |
| adk | GMYN | 0.0270 | 1.0241 | 0.0263 | 2.16E-43 | 663 | 169.874 | 493.126 | 96 | 82.9971 | 13.0029 |
| adk | GMYN | 0.0270 | 1.0241 | 0.0263 | 2.16E-43 | 663 | 169.874 | 493.126 | 96 | 82.9971 | 13.0029 |
| acsA | GMYN | 0.0307 | 1.1675 | 0.0263 | 7.83E-157 | 1971 | 460.746 | 1510.25 | 318 | 272.702 | 45.2985 |
| gyr | GMYN | 0.0413 | 1.5850 | 0.0260 | 5.04E-137 | 1434 | 390.054 | 1043.95 | 299 | 257.19 | 41.8102 |
| adk | MS | 0.0266 | 1.0241 | 0.0260 | 1.51E-54 | 663 | 196.497 | 466.503 | 103 | 97.0207 | 5.97934 |
| iso | MS | 0.0307 | 1.1857 | 0.0259 | 7.20E-202 | 2232 | 621.318 | 1610.68 | 373 | 349.512 | 23.4885 |
| adk | GMYN | 0.0230 | 0.9005 | 0.0256 | 2.82E-41 | 663 | 175.583 | 487.417 | 91 | 79.9576 | 11.0424 |
| adk | GMYN | 0.0336 | 1.3159 | 0.0255 | 6.14E-54 | 663 | 141.762 | 521.238 | 105 | 87.9287 | 17.0713 |
| CTP | GMYN | 0.0379 | 1.4861 | 0.0255 | 3.88E-147 | 1659 | 399.829 | 1259.17 | 305 | 258.527 | 46.4732 |

| | | | | | | | | | | | |
|---|---|---|---|---|---|---|---|---|---|---|---|
| iso | GMYN | 0.0300 | 1.1779 | 0.0255 | 2.24E-183 | 2232 | 544.351 | 1687.65 | 373 | 323.368 | 49.6319 |
| glnA | MS | 0.0104 | 0.4212 | 0.0248 | 1.15E-62 | 1413 | 361.456 | 1051.54 | 118 | 110.068 | 7.93202 |
| cit | GMYN | 0.0225 | 0.9200 | 0.0244 | 3.22E-78 | 1311 | 316.177 | 994.823 | 167 | 144.987 | 22.0131 |
| acsA | GMYN | 0.0356 | 1.4655 | 0.0243 | 3.08E-127 | 1887 | 469.867 | 1417.13 | 300 | 250.775 | 49.2249 |
| ndk | GMYN | 0.0125 | 0.5149 | 0.0243 | 1.76E-20 | 423 | 99.3462 | 323.654 | 39 | 35 | 4 |
| iso | MS | 0.0246 | 1.0173 | 0.0242 | 7.82E-196 | 2232 | 578.399 | 1653.6 | 345 | 322.713 | 22.287 |
| acsA | MS | 0.0191 | 0.7978 | 0.0240 | 7.18E-143 | 1959 | 537.515 | 1421.49 | 267 | 251.083 | 15.9166 |
| cit | MS | 0.0220 | 0.9288 | 0.0237 | 2.36E-108 | 1311 | 325.072 | 985.928 | 186 | 173.521 | 12.4791 |
| gyr | GMYN | 0.0496 | 2.0975 | 0.0237 | 2.78E-138 | 1365 | 365.201 | 999.799 | 305 | 257.189 | 47.8108 |
| glnA | MS | 0.0110 | 0.4696 | 0.0234 | 9.82E-66 | 1344 | 344.318 | 999.682 | 122 | 114.235 | 7.76513 |
| cit | MS | 0.0248 | 1.0765 | 0.0230 | 1.78E-104 | 1311 | 403.661 | 907.339 | 207 | 196.811 | 10.1895 |
| cit | MS | 0.0248 | 1.0818 | 0.0229 | 1.70E-120 | 1299 | 314.94 | 984.06 | 199 | 185.712 | 13.2882 |
| glnA | GMYN | 0.0223 | 0.9748 | 0.0228 | 1.26E-75 | 1413 | 314.461 | 1098.54 | 161 | 136.929 | 24.0706 |
| glnA | MS | 0.0079 | 0.3491 | 0.0225 | 2.74E-54 | 1413 | 404.929 | 1008.07 | 109 | 103.207 | 5.79339 |
| adk | MS | 0.0231 | 1.0322 | 0.0224 | 1.05E-56 | 663 | 176.891 | 486.109 | 98 | 92.3184 | 5.68161 |
| iso | MS | 0.0223 | 0.9998 | 0.0223 | 1.13E-190 | 2232 | 596.507 | 1635.49 | 339 | 319.447 | 19.5526 |
| iso | MS | 0.0223 | 0.9998 | 0.0223 | 1.13E-190 | 2232 | 596.507 | 1635.49 | 339 | 319.447 | 19.5526 |
| ndk | MS | 0.0097 | 0.4362 | 0.0222 | 9.09E-21 | 423 | 116.198 | 306.802 | 36 | 34.0073 | 1.99266 |
| acsA | GMYN | 0.0316 | 1.4453 | 0.0218 | 9.23E-142 | 1971 | 442.811 | 1528.19 | 297 | 249.879 | 47.1209 |
| glnA | GMYN | 0.0252 | 1.1563 | 0.0218 | 5.25E-93 | 1413 | 316.056 | 1096.94 | 189 | 161.881 | 27.1187 |
| CTP | GMYN | 0.0287 | 1.3198 | 0.0218 | 1.33E-147 | 1659 | 407.071 | 1251.93 | 288 | 252.764 | 35.2364 |
| gyr | GMYN | 0.0433 | 2.0001 | 0.0216 | 9.49E-147 | 1422 | 384.668 | 1037.33 | 312 | 268.459 | 43.5413 |
| CTP | MS | 0.0445 | 2.0753 | 0.0215 | 2.13E-220 | 1656 | 426.505 | 1229.5 | 340 | 320.192 | 19.8082 |
| succ | MS | 0.0264 | 1.2285 | 0.0215 | 2.73E-172 | 1776 | 450.743 | 1325.26 | 288 | 270.908 | 17.0919 |
| succ | MS | 0.0176 | 0.8302 | 0.0212 | 4.27E-135 | 1776 | 438.421 | 1337.58 | 230 | 216.015 | 13.9846 |
| cit | MS | 0.0187 | 0.8896 | 0.0210 | 6.26E-103 | 1311 | 344.629 | 966.371 | 182 | 171.865 | 10.1346 |
| atpA | MS | 0.0069 | 0.3281 | 0.0210 | 2.25E-58 | 1539 | 414.089 | 1124.91 | 111 | 105.009 | 5.99135 |
| iso | MS | 0.0194 | 0.9274 | 0.0209 | 3.43E-180 | 2232 | 612.155 | 1619.84 | 327 | 309.838 | 17.1624 |
| ndk | MS | 0.0121 | 0.5806 | 0.0209 | 6.22E-24 | 423 | 99.3507 | 323.649 | 39 | 36.5139 | 2.48615 |
| CTP | MS | 0.0452 | 2.1968 | 0.0206 | 1.82E-239 | 1659 | 389.105 | 1269.89 | 339 | 317.671 | 21.3295 |
| succ | MS | 0.0244 | 1.1871 | 0.0206 | 1.84E-169 | 1776 | 457.36 | 1318.64 | 283 | 267.154 | 15.8464 |
| atpA | MS | 0.0109 | 0.5337 | 0.0205 | 3.59E-84 | 1536 | 431.827 | 1104.17 | 163 | 154.883 | 8.11725 |
| CTP | MS | 0.0287 | 1.4080 | 0.0204 | 5.77E-183 | 1659 | 410.376 | 1248.62 | 288 | 271.188 | 16.812 |
| acsA | MS | 0.0190 | 0.9509 | 0.0199 | 6.22E-165 | 1971 | 534.467 | 1436.53 | 291 | 276.202 | 14.7979 |

| | | | | | | | | | | | |
|---|---|---|---|---|---|---|---|---|---|---|---|
| CTP | MS | 0.0469 | 2.3755 | 0.0197 | 1.85E-244 | 1659 | 395.744 | 1263.26 | 345 | 324.552 | 20.4483 |
| iso | MS | 0.0161 | 0.8209 | 0.0196 | 1.85E-170 | 2232 | 595.338 | 1636.66 | 303 | 287.544 | 15.4558 |
| atpA | GMYN | 0.0063 | 0.3240 | 0.0194 | 4.47E-54 | 1539 | 420.3 | 1118.7 | 111 | 104 | 7 |
| glnA | GMYN | 0.0257 | 1.3264 | 0.0193 | 8.04E-75 | 1401 | 284.568 | 1116.43 | 159 | 130.875 | 28.1248 |
| ndk | GMYN | 0.0093 | 0.5018 | 0.0186 | 4.30E-20 | 423 | 98.377 | 324.623 | 36 | 33 | 3 |
| glnA | GMYN | 0.0107 | 0.5838 | 0.0184 | 1.10E-64 | 1344 | 311.139 | 1032.86 | 122 | 110.994 | 11.0057 |
| glnA | GMYN | 0.0077 | 0.4163 | 0.0184 | 6.23E-55 | 1413 | 360.822 | 1052.18 | 109 | 100.997 | 8.00264 |
| succ | MS | 0.0213 | 1.1644 | 0.0183 | 1.66E-166 | 1764 | 411.916 | 1352.08 | 259 | 244.355 | 14.6446 |
| adk | GMYN | 0.0392 | 2.1557 | 0.0182 | 1.77E-47 | 663 | 133.984 | 529.016 | 100 | 79.8747 | 20.1253 |
| gyr | GMYN | 0.0314 | 1.7556 | 0.0179 | 3.20E-125 | 1434 | 410.89 | 1023.11 | 275 | 243.578 | 31.4223 |
| gyr | GMYN | 0.0402 | 2.2461 | 0.0179 | 1.53E-156 | 1434 | 390.631 | 1043.37 | 319 | 278.312 | 40.6879 |
| ndk | MS | 0.0068 | 0.3892 | 0.0176 | 1.40E-18 | 423 | 108.122 | 314.878 | 31 | 29.4898 | 1.51023 |
| pyr | GMYN | 0.0365 | 2.0799 | 0.0175 | 1.84E-152 | 1434 | 368.689 | 1065.31 | 297 | 259.245 | 37.7552 |
| acsA | GMYN | 0.0272 | 1.5565 | 0.0175 | 3.77E-141 | 1971 | 432.585 | 1538.41 | 281 | 239.927 | 41.0729 |
| acsA | GMYN | 0.0272 | 1.5565 | 0.0175 | 3.77E-141 | 1971 | 432.585 | 1538.41 | 281 | 239.927 | 41.0729 |
| cit | GMYN | 0.0236 | 1.3596 | 0.0173 | 3.04E-96 | 1311 | 359.966 | 951.034 | 207 | 184.961 | 22.0394 |
| cit | MS | 0.0118 | 0.6805 | 0.0173 | 3.34E-86 | 1311 | 376.81 | 934.19 | 160 | 153.414 | 6.58645 |
| iso | GMYN | 0.0245 | 1.4186 | 0.0172 | 2.73E-144 | 2232 | 595.329 | 1636.67 | 324 | 284.646 | 39.3536 |
| adk | MS | 0.0273 | 1.6085 | 0.0170 | 2.16E-71 | 663 | 178.29 | 484.71 | 116 | 110.878 | 5.12185 |
| cit | MS | 0.0140 | 0.8290 | 0.0169 | 3.17E-103 | 1311 | 318.87 | 992.13 | 168 | 159.61 | 8.38997 |
| cit | MS | 0.0140 | 0.8290 | 0.0169 | 3.17E-103 | 1311 | 318.87 | 992.13 | 168 | 159.61 | 8.38997 |
| succ | MS | 0.0244 | 1.4442 | 0.0169 | 1.68E-197 | 1773 | 415.09 | 1357.91 | 294 | 278.629 | 15.3715 |
| CTP | MS | 0.0431 | 2.5545 | 0.0169 | 4.83E-258 | 1647 | 388.169 | 1258.83 | 348 | 329.961 | 18.0385 |
| acsA | GMYN | 0.0253 | 1.5192 | 0.0167 | 9.39E-140 | 1971 | 433.419 | 1537.58 | 274 | 235.81 | 38.1899 |
| iso | GMYN | 0.0222 | 1.3559 | 0.0164 | 2.72E-166 | 2232 | 575.916 | 1656.08 | 339 | 302.855 | 36.1454 |
| iso | GMYN | 0.0222 | 1.3559 | 0.0164 | 2.72E-166 | 2232 | 575.916 | 1656.08 | 339 | 302.855 | 36.1454 |
| cit | MS | 0.0226 | 1.3813 | 0.0163 | 2.13E-148 | 1308 | 317.063 | 990.937 | 222 | 211.209 | 10.7907 |
| glnA | GMYN | 0.0140 | 0.8743 | 0.0161 | 5.05E-67 | 1413 | 332.257 | 1080.74 | 136 | 120.992 | 15.0082 |
| glnA | GMYN | 0.0140 | 0.8743 | 0.0161 | 5.05E-67 | 1413 | 332.257 | 1080.74 | 136 | 120.992 | 15.0082 |
| ndk | GMYN | 0.0063 | 0.3899 | 0.0160 | 9.94E-18 | 423 | 101.342 | 321.658 | 31 | 29 | 2 |
| gyr | GMYN | 0.0438 | 2.7416 | 0.0160 | 4.14E-155 | 1434 | 367.735 | 1066.27 | 314 | 268.672 | 45.3283 |
| cit | GMYN | 0.0227 | 1.4395 | 0.0158 | 6.41E-126 | 1308 | 312.118 | 995.882 | 222 | 199.746 | 22.2536 |
| CTP | MS | 0.0291 | 1.8451 | 0.0158 | 1.39E-216 | 1659 | 396.265 | 1262.73 | 309 | 294.203 | 14.7971 |
| CTP | MS | 0.0291 | 1.8451 | 0.0158 | 1.39E-216 | 1659 | 396.265 | 1262.73 | 309 | 294.203 | 14.7971 |

| | | | | | | | | | | | |
|---|---|---|---|---|---|---|---|---|---|---|---|
| gyr | MS | 0.0292 | 1.8620 | 0.0157 | 1.38E-164 | 1434 | 412.584 | 1021.42 | 275 | 264.715 | 10.2845 |
| gyr | MS | 0.0306 | 1.9569 | 0.0156 | 6.70E-173 | 1434 | 394.482 | 1039.52 | 275 | 264.113 | 10.8867 |
| gyr | MS | 0.0440 | 2.8297 | 0.0156 | 1.63E-214 | 1434 | 397.871 | 1036.13 | 323 | 310.426 | 12.5738 |
| cit | GMYN | 0.0117 | 0.7571 | 0.0155 | 1.91E-79 | 1311 | 365.424 | 945.576 | 160 | 149 | 11 |
| adk | GMYN | 0.0278 | 1.8006 | 0.0154 | 1.89E-66 | 663 | 149.552 | 513.448 | 116 | 101.997 | 14.0029 |
| acsA | GMYN | 0.0182 | 1.1909 | 0.0153 | 7.15E-145 | 1959 | 455.441 | 1503.56 | 267 | 239.983 | 27.0171 |
| succ | MS | 0.0121 | 0.7966 | 0.0152 | 2.38E-123 | 1776 | 527.008 | 1248.99 | 232 | 223.912 | 8.08822 |
| gyr | MS | 0.0421 | 2.8067 | 0.0150 | 1.76E-219 | 1422 | 370.516 | 1051.48 | 312 | 299.273 | 12.7274 |
| iso | GMYN | 0.0190 | 1.2797 | 0.0149 | 1.91E-167 | 2232 | 569.193 | 1662.81 | 327 | 295.802 | 31.1981 |
| iso | GMYN | 0.0364 | 2.4546 | 0.0148 | 4.93E-164 | 2220 | 511.193 | 1708.81 | 358 | 297.458 | 60.5418 |
| gyr | GMYN | 0.0295 | 1.9973 | 0.0148 | 5.21E-126 | 1434 | 417.706 | 1016.29 | 275 | 245.663 | 29.3372 |
| succ | MS | 0.0126 | 0.8678 | 0.0145 | 1.08E-139 | 1776 | 459.187 | 1316.81 | 233 | 223.668 | 9.33209 |
| succ | MS | 0.0126 | 0.8678 | 0.0145 | 1.08E-139 | 1776 | 459.187 | 1316.81 | 233 | 223.668 | 9.33209 |
| gyr | MS | 0.0400 | 2.7876 | 0.0144 | 2.61E-202 | 1434 | 380.038 | 1053.96 | 299 | 287.548 | 11.4519 |
| CTP | GMYN | 0.0292 | 2.0363 | 0.0143 | 9.17E-174 | 1659 | 389.391 | 1269.61 | 309 | 272.709 | 36.2908 |
| CTP | GMYN | 0.0292 | 2.0363 | 0.0143 | 9.17E-174 | 1659 | 389.391 | 1269.61 | 309 | 272.709 | 36.2908 |
| succ | GMYN | 0.0252 | 1.7650 | 0.0143 | 1.62E-172 | 1776 | 373.114 | 1402.89 | 288 | 253.238 | 34.7617 |
| glnA | MS | 0.0174 | 1.2335 | 0.0141 | 6.92E-87 | 873 | 232.469 | 640.531 | 140 | 134.776 | 5.22418 |
| gyr | GMYN | 0.0201 | 1.4627 | 0.0137 | 4.18E-91 | 894 | 225.047 | 668.953 | 158 | 144.779 | 13.2208 |
| CTP | MS | 0.0311 | 2.2809 | 0.0136 | 7.07E-232 | 1659 | 402.797 | 1256.2 | 324 | 310.796 | 13.2037 |
| cit | GMYN | 0.0235 | 1.7439 | 0.0135 | 6.66E-123 | 1299 | 258.548 | 1040.45 | 199 | 174.954 | 24.0462 |
| gyr | MS | 0.0394 | 2.9300 | 0.0135 | 8.63E-225 | 1431 | 361.43 | 1069.57 | 308 | 296.203 | 11.797 |
| gyr | MS | 0.0484 | 3.6212 | 0.0134 | 1.86E-215 | 1365 | 363.563 | 1001.44 | 305 | 294.174 | 10.8262 |
| acsA | GMYN | 0.0240 | 1.8174 | 0.0132 | 1.06E-149 | 1971 | 437.394 | 1533.61 | 283 | 246.884 | 36.1162 |
| atpA | MS | 0.0019 | 0.1442 | 0.0129 | 2.04E-30 | 1539 | 464.41 | 1074.59 | 59 | 57.2907 | 1.70933 |
| atpA | GMYN | 0.0074 | 0.5742 | 0.0129 | 2.82E-84 | 1539 | 449.593 | 1089.41 | 168 | 159.998 | 8.00241 |
| gyr | GMYN | 0.0295 | 2.2954 | 0.0128 | 1.71E-164 | 1434 | 379.947 | 1054.05 | 302 | 271.606 | 30.3935 |
| gyr | GMYN | 0.0432 | 3.3716 | 0.0128 | 2.55E-161 | 1434 | 375.774 | 1058.23 | 323 | 278.685 | 44.3148 |
| adk | GMYN | 0.0245 | 1.9126 | 0.0128 | 6.89E-54 | 663 | 159.511 | 503.489 | 103 | 90.8916 | 12.1084 |
| atpA | GMYN | 0.0103 | 0.8107 | 0.0128 | 2.82E-90 | 1536 | 366.785 | 1169.21 | 163 | 150.998 | 12.0025 |
| atpA | MS | 0.0073 | 0.5753 | 0.0127 | 1.77E-92 | 1539 | 437.524 | 1101.48 | 168 | 162.798 | 5.20194 |
| succ | GMYN | 0.0204 | 1.6253 | 0.0126 | 2.26E-165 | 1764 | 347.102 | 1416.9 | 259 | 230.516 | 28.4839 |
| succ | GMYN | 0.0216 | 1.7305 | 0.0125 | 1.09E-131 | 1776 | 408.645 | 1367.35 | 250 | 220.971 | 29.029 |
| cit | GMYN | 0.0194 | 1.5580 | 0.0124 | 2.85E-98 | 1311 | 316.295 | 994.705 | 186 | 166.995 | 19.0047 |

| | | | | | | | | | | | |
|---|---|---|---|---|---|---|---|---|---|---|---|
| atpA | MS | 0.0027 | 0.2263 | 0.0120 | 4.25E-44 | 1539 | 438.488 | 1100.51 | 83 | 80.5698 | 2.43025 |
| atpA | MS | 0.0029 | 0.2471 | 0.0119 | 5.50E-48 | 1539 | 416.33 | 1122.67 | 86 | 83.3313 | 2.66866 |
| atpA | MS | 0.0029 | 0.2471 | 0.0119 | 5.50E-48 | 1539 | 416.33 | 1122.67 | 86 | 83.3313 | 2.66866 |
| glnA | GMYN | 0.0166 | 1.4398 | 0.0116 | 8.00E-87 | 873 | 201.482 | 671.518 | 140 | 128.951 | 11.0489 |
| gyr | MS | 0.0425 | 3.6752 | 0.0116 | 7.32E-232 | 1434 | 365.786 | 1068.21 | 314 | 303.75 | 10.2497 |
| gyr | MS | 0.0394 | 3.4594 | 0.0114 | 1.43E-214 | 1434 | 402.787 | 1031.21 | 319 | 309.964 | 9.03628 |
| cit | GMYN | 0.0172 | 1.5183 | 0.0113 | 1.66E-120 | 1311 | 251.927 | 1059.07 | 182 | 163.991 | 18.0088 |
| atpA | MS | 0.0058 | 0.5249 | 0.0110 | 3.44E-87 | 1527 | 408.415 | 1118.59 | 150 | 145.618 | 4.3816 |
| cit | MS | 0.0071 | 0.6569 | 0.0108 | 2.40E-89 | 1311 | 330.451 | 980.549 | 144 | 139.534 | 4.46591 |
| atpA | GMYN | 0.0018 | 0.1646 | 0.0107 | 2.22E-32 | 1539 | 400.875 | 1138.13 | 59 | 57 | 2 |
| cit | GMYN | 0.0175 | 1.6527 | 0.0106 | 2.62E-89 | 1242 | 259.875 | 982.125 | 155 | 137.996 | 17.0036 |
| iso | GMYN | 0.0239 | 2.2572 | 0.0106 | 1.91E-182 | 2232 | 522.921 | 1709.08 | 345 | 304.774 | 40.2261 |
| CTP | GMYN | 0.0468 | 4.4215 | 0.0106 | 5.03E-186 | 1659 | 363.337 | 1295.66 | 345 | 286.354 | 58.6459 |
| cit | GMYN | 0.0072 | 0.6880 | 0.0105 | 2.31E-82 | 1311 | 333.787 | 977.213 | 144 | 136.999 | 7.00117 |
| succ | GMYN | 0.0232 | 2.2408 | 0.0104 | 6.18E-182 | 1776 | 351.706 | 1424.29 | 283 | 250.484 | 32.5163 |
| atpA | GMYN | 0.0054 | 0.5244 | 0.0103 | 9.82E-82 | 1527 | 415.773 | 1111.23 | 150 | 143.998 | 6.00155 |
| CTP | GMYN | 0.0455 | 4.3991 | 0.0103 | 4.97E-188 | 1659 | 352.653 | 1306.35 | 339 | 281.475 | 57.5253 |
| gyr | MS | 0.0294 | 2.9137 | 0.0101 | 9.18E-210 | 1434 | 389.71 | 1044.29 | 302 | 294.044 | 7.95621 |
| CTP | GMYN | 0.0446 | 4.4242 | 0.0101 | 1.94E-193 | 1647 | 364.641 | 1282.36 | 348 | 292.589 | 55.4106 |
| CTP | GMYN | 0.0283 | 2.8327 | 0.0100 | 1.49E-188 | 1659 | 394.251 | 1264.75 | 324 | 288.919 | 35.0806 |
| CTP | MS | 0.0157 | 1.5813 | 0.0099 | 1.40E-127 | 1044 | 271.359 | 772.641 | 188 | 182.824 | 5.1758 |
| pyr | MS | 0.0363 | 3.6677 | 0.0099 | 5.11E-210 | 1434 | 380.539 | 1053.46 | 297 | 289.072 | 7.92841 |
| atpA | GMYN | 0.0026 | 0.2655 | 0.0098 | 1.55E-49 | 1539 | 381.204 | 1157.8 | 83 | 80 | 3 |
| CTP | GMYN | 0.0430 | 4.4527 | 0.0097 | 6.11E-180 | 1656 | 378.766 | 1277.23 | 340 | 286.693 | 53.3066 |
| acsA | MS | 0.0123 | 1.2940 | 0.0095 | 1.95E-137 | 1323 | 346.255 | 976.745 | 210 | 204.51 | 5.48991 |
| CTP | GMYN | 0.0409 | 4.3859 | 0.0093 | 3.89E-177 | 1659 | 346.475 | 1312.52 | 318 | 265.895 | 52.1048 |
| atpA | GMYN | 0.0026 | 0.2821 | 0.0092 | 8.33E-50 | 1539 | 378.591 | 1160.41 | 86 | 82.9997 | 3.00028 |
| atpA | GMYN | 0.0026 | 0.2821 | 0.0092 | 8.33E-50 | 1539 | 378.591 | 1160.41 | 86 | 82.9997 | 3.00028 |
| adk | GMYN | 0.0217 | 2.3886 | 0.0091 | 2.01E-57 | 663 | 142.78 | 520.22 | 98 | 86.9152 | 11.0848 |
| succ | GMYN | 0.0161 | 1.7823 | 0.0090 | 7.29E-132 | 1776 | 393.031 | 1382.97 | 230 | 207.96 | 22.0404 |
| gyr | GMYN | 0.0392 | 4.3704 | 0.0090 | 5.64E-173 | 1431 | 339.397 | 1091.6 | 308 | 266.397 | 41.6028 |
| atpA | MS | 0.0049 | 0.5582 | 0.0088 | 3.11E-94 | 1539 | 415.525 | 1123.47 | 158 | 154.339 | 3.66115 |
| iso | MS | 0.0126 | 1.4558 | 0.0087 | 1.58E-164 | 1527 | 417.299 | 1109.7 | 255 | 249.242 | 5.75769 |
| acsA | GMYN | 0.0120 | 1.3882 | 0.0086 | 3.91E-135 | 1323 | 313.179 | 1009.82 | 210 | 197.983 | 12.0165 |

| | | | | | | | | | | | |
|---|---|---|---|---|---|---|---|---|---|---|---|
| acsA | GMYN | 0.0241 | 2.8319 | 0.0085 | 2.40E-161 | 1971 | 444.845 | 1526.16 | 299 | 262.838 | 36.1616 |
| iso | GMYN | 0.0154 | 1.8312 | 0.0084 | 3.51E-174 | 2232 | 513.728 | 1718.27 | 303 | 276.868 | 26.1316 |
| succ | GMYN | 0.0235 | 2.8881 | 0.0081 | 2.81E-134 | 1776 | 383.146 | 1392.85 | 249 | 216.877 | 32.1227 |
| succ | GMYN | 0.0113 | 1.4411 | 0.0078 | 5.91E-136 | 1776 | 427.859 | 1348.14 | 232 | 216.947 | 15.0531 |
| ndk | GMYN | 0.0031 | 0.3955 | 0.0078 | 7.47E-19 | 423 | 96.2147 | 326.785 | 29 | 28 | 1 |
| gyr | MS | 0.0198 | 2.5768 | 0.0077 | 6.94E-108 | 894 | 236.303 | 657.697 | 158 | 154.7 | 3.30038 |
| iso | GMYN | 0.0121 | 1.7223 | 0.0070 | 8.02E-170 | 1527 | 357.192 | 1169.81 | 255 | 240.939 | 14.0612 |
| ndk | MS | 0.0030 | 0.4310 | 0.0070 | 0 | 423 | 98.4718 | 324.528 | 29 | 28.3486 | 0.651395 |
| succ | GMYN | 0.0111 | 1.6003 | 0.0069 | 3.14E-141 | 1776 | 413.624 | 1362.38 | 233 | 218 | 15.0005 |
| succ | GMYN | 0.0111 | 1.6003 | 0.0069 | 3.14E-141 | 1776 | 413.624 | 1362.38 | 233 | 218 | 15.0005 |
| ndk | GMYN | 0.0030 | 0.4702 | 0.0065 | 3.64E-19 | 423 | 94.1205 | 328.879 | 29 | 28 | 1 |
| ndk | GMYN | 0.0030 | 0.4702 | 0.0065 | 3.64E-19 | 423 | 94.1205 | 328.879 | 29 | 28 | 1 |
| acsA | GMYN | 0.0179 | 2.7781 | 0.0065 | 3.70E-171 | 1971 | 443.175 | 1527.82 | 291 | 263.979 | 27.0206 |
| ndk | MS | 0.0029 | 0.4709 | 0.0063 | 0 | 423 | 87.6668 | 335.333 | 29 | 28.3217 | 0.678306 |
| ndk | MS | 0.0029 | 0.4709 | 0.0063 | 0 | 423 | 87.6668 | 335.333 | 29 | 28.3217 | 0.678306 |
| cit | GMYN | 0.0128 | 2.1119 | 0.0061 | 3.95E-105 | 1311 | 281.381 | 1029.62 | 168 | 154.948 | 13.0524 |
| cit | GMYN | 0.0128 | 2.1119 | 0.0061 | 3.95E-105 | 1311 | 281.381 | 1029.62 | 168 | 154.948 | 13.0524 |
| atpA | GMYN | 0.0044 | 0.7333 | 0.0060 | 2.83E-94 | 1539 | 394.052 | 1144.95 | 158 | 152.999 | 5.00095 |
| CTP | GMYN | 0.0163 | 2.9439 | 0.0055 | 1.49E-126 | 1044 | 238.02 | 805.98 | 188 | 174.992 | 13.0082 |
| succ | GMYN | 0.0231 | 4.3822 | 0.0053 | 2.19E-198 | 1773 | 344.8 | 1428.2 | 294 | 261.573 | 32.4274 |
| cit | GMYN | 0.0053 | 1.1520 | 0.0046 | 4.48E-88 | 816 | 242.962 | 573.038 | 146 | 142.999 | 3.00097 |
| adk | GMYN | 0.0073 | 1.6208 | 0.0045 | 1.83E-45 | 375 | 99.4572 | 275.543 | 68 | 65.9998 | 2.00022 |
| adk | MS | 0.0077 | 1.7370 | 0.0044 | 0 | 375 | 112.06 | 262.94 | 68 | 67.3013 | 0.698709 |
| succ | MS | 0.0052 | 1.2822 | 0.0041 | 6.65E-120 | 1137 | 286.368 | 850.632 | 171 | 168.951 | 2.04913 |
| succ | GMYN | 0.0047 | 1.2350 | 0.0038 | 6.38E-119 | 1137 | 275.818 | 861.182 | 171 | 167 | 4.00027 |
| cit | MS | 0.0055 | 1.8270 | 0.0030 | 0 | 816 | 267.637 | 548.363 | 146 | 145.107 | 0.892631 |
| glnA | MS | 0.0009 | 0.3288 | 0.0029 | 0 | 1413 | 376.781 | 1036.22 | 93 | 92.2678 | 0.732182 |
| glnA | GMYN | 0.0010 | 0.3391 | 0.0028 | 1.14E-57 | 1413 | 360.354 | 1052.65 | 93 | 91.9999 | 1.00008 |
| ndk | MS | 0.0000 | 0.8468 | 0.0000 | 0 | 225 | 71.2132 | 153.787 | 25 | 24.9999 | 5.40E-05 |
| atpA | MS | 0.0000 | 0.7320 | 0.0000 | 0 | 963 | 252.127 | 710.873 | 110 | 110 | 0.000310145 |
| atpA | GMYN | 0.0000 | 0.9391 | 0.0000 | 0 | 963 | 238.516 | 724.484 | 110 | 110 | 9.42E-10 |

GMYN = Approximate method (γ-MYN). Wang, D. P., Wan, H. L., Zhang, S., Yu, J. (2009). γ-MYN: a new algorithm for estimating Ka and Ks with consideration of variable substitution rates. *Biol. Direct*. 4, 20.

MS = Maximum-likelihood method. Posada, D. (2003). "Using Modeltest and PAUP* to select a model of nucleotide substitution", in Current Protocols in Bioinformatics, ed. A. D. Baxevanis (New York, USA: JohnWiley & Sons), 6.5.1-6.5.14.

**Supplementary Table S12.** Ka/Ks calculation using highly expressed selected genes and housekeeping genes from *Polynucleobacter asymbioticus* st QLW-P1DMWA-1$^{T}$ and strains belonging to the PnecC subcluster.

**Details of the analysed genes related to *Polynucleobacter asymbioticus* strain QLW-P1DMWA-1$^{T}$.**

| Locus Tag | Gene Product Name | Abbreviation | DNA Sequence Length (bp) |
|---|---|---|---|
| **Gene set** | | | |
| Pnuc_0690 | DNA translocase FtsK | FtsK | 2313 |
| Pnuc_1769 | Chaperone protein DnaK | Dank | 1935 |
| Pnuc_1805 | Chaperonin GroEL | GroL | 1653 |
| Pnuc_1133 | N-6 DNA methylase | methyl | 1473 |
| Pnuc_1736 | Fatty acid desaturase | fatty | 1200 |
| Pnuc_1880 | Holliday junction DNA helicase subunit RuvB | RuvB | 1071 |
| Pnuc_1929 | RNA polymerase, sigma 32 subunit, RpoH | RpoH | 930 |
| Pnuc_0426 | MscS Mechanosensitive ion channel | MscS | 822 |
| Pnuc_0725 | Pirin domain protein | Piri | 702 |
| Pnuc_2030 | Carbonate dehydratase | Carbo | 666 |
| Pnuc_0747 | DSBA oxidoreductase | DSBA | 627 |
| Pnuc_0334 | Adenylyl-sulfate kinase | CysC | 588 |
| Pnuc_1064 | RNA polymerase, sigma-24 subunit, RpoE | RpoE | 570 |
| Pnuc_1626 | Superoxide dismutase, copper/zinc binding | Superox | 534 |
| Pnuc_1828 | Fimbrial protein pilin | Fimb | 513 |
| Pnuc_1376 | Rubrerythrin | Rubre | 486 |
| Pnuc_0429 | Alkyl hydroperoxide reductase | Alkyl | 468 |
| Pnuc_0425 | Hypothetical protein | Hyp | 267 |
| Pnuc_1286 | RNA-binding protein Hfq | Hfq | 243 |
| Pnuc_1750 | Cold-shock DNA-binding protein family | Cold | 204 |
| **Housekeeping genes** | | | |
| Pnuc_0366 | Isocitrate dehydrogenase, NADP-dependent | iso | 2235 |
| Pnuc_1138 | Acetyl-coenzyme A synthetase (EC 6.2.1.1) | acsA | 1974 |
| Pnuc_0760 | Succinate dehydrogenase subunit A (EC 1.3.5.1) | succ | 1779 |
| Pnuc_0945 | CTP synthase (EC 6.3.4.2) | CTP | 1662 |
| Pnuc_0024 | ATP synthase F1 subcomplex alpha subunit | atpA | 1542 |
| Pnuc_1816 | Pyruvate kinase (EC 2.7.1.40) | gyr | 1437 |
| Pnuc_1255 | L-glutamine synthetase (EC 6.3.1.2) | glnA | 1416 |
| Pnuc_0763 | Citrate synthase (EC 2.3.3.1) | cit | 1314 |
| Pnuc_0281 | Adenylate kinase (EC 2.7.4.3) | adk | 666 |
| Pnuc_1293 | Nucleoside diphosphate kinase (EC 2.7.4.6) | ndk | 426 |
